# Combining single-cell RNA-sequencing with a molecular atlas unveils new markers for *C. elegans* neuron classes

**DOI:** 10.1101/826560

**Authors:** Ramiro Lorenzo, Michiho Onizuka, Matthieu Defrance, Patrick Laurent

**Affiliations:** Laboratory of Neurophysiology, ULB Neuroscience Institute (UNI), Université Libre de Bruxelles (ULB), Brussels, Belgium; Centro de Investigación Veterinaria de Tandil (CIVETAN), CONICET-CICPBA-UNCPBA, Facultad de Ciencias Veterinarias, Universidad Nacional del Centro (FCV-UNCPBA), Tandil, Argentina; Interuniversity Institute of Bioinformatics in Brussels, Université Libre de Bruxelles, Brussels, Belgium

**Author notes:** These authors contributed equally to this work (Ramiro Lorenzo, Michiho Onizuka). Corresponding author: (PL).

## Abstract

Single-cell RNA-sequencing (scRNA-seq) of the *Caenorhabditis elegans* (*C. elegans*) nervous system offers the unique opportunity to obtain a partial expression profile for each neuron within a known connectome. Building on recent scRNA-seq data [1] and on a molecular atlas describing the expression pattern of ~800 genes at the single cell resolution [2], we designed an iterative clustering analysis aiming to match each cell-cluster to the ~100 anatomically defined neuron classes of *C. elegans*. This heuristic approach successfully assigned 58 clusters to their corresponding neuron class. Another 11 clusters grouped neuron classes sharing close molecular signatures and 7 clusters were not assigned. Based on these 76 molecular profiles, we designed 15 new neuron class-specific promoters validated *in vivo.* Amongst them, 10 represent the only specific promoter reported to this day, expanding the list of neurons amenable to genetic manipulations. Finally, we observed a differential expression of functionally relevant genes between sensory-, inter-, and motor neurons in *C. elegans*, suggesting the mode of functional diversification may vary accordingly to the neuronal modalities.

## Introduction

The human brain comprises around 100 billion neurons. Early classification based on morphology already distinguished pyramidal, stellate, granule, bipolar, Purkinje or basket cells [3]. Additional knowledge about their neurotransmitters, functional and electrophysiological properties further improved this classification. Recent progresses in molecular profiling using single-cell RNA-sequencing (scRNA-seq) allowed exploring neuronal diversity at the molecular level in human and mouse brain [4–6]. In comparison, the nervous system of *Caenorhabditis elegans* (*C. elegans*) adult hermaphrodite has a simple and well-described structure composed of only 302 neurons. Despite its anatomical simplicity, the neuronal diversity of *C. elegans* encompasses 118 neuron classes identified by the examination of their complete diagram of connectivity as revealed by serial sections and electron microscopy [7–9]. Based on expression patterns at the single-cell resolution, the 118 anatomically defined neuron classes in the adult hermaphrodite would correspond to 118 or more unique molecular identities [2, 10]. However, the transcriptional profiles are only known for a few neuron classes purified by FACS-sorting [11–14].

The exquisitely described small circuits composed by few neurons in *C. elegans* allow exploring how cell-to-cell communication can shape behaviour. The function of several *C. elegans* neuron classes has been defined using laser ablation, genetic and optogenetic methods [15, 16]. These later methods, however, rely on reliable promoters driving transgene expression in a single neuron class to be characterized. Currently, cell-specific promoters are described for only 29 neuron classes. This current lack of cell-specific promoters has limited the optogenetic and chemogenetic analysis of the neuronal functions to a fraction of the *C. elegans* nervous system.

Recently, scRNA-seq of the full *C. elegans* organism at L2 larval stage was delivered [1]. Out of the 42.000 sequenced cells, the author yielded ~7000 single neuron transcriptomes. Based on gene co-expression patterns, a semi-supervised clustering analysis segregated these 7000 neurons into ~40 clusters but assigned only 10 of them to their corresponding neuron class. However, 112 out of the 118 adult neuron classes are theoretically present in the L2 larval stage [17]. Hence, these 10 clusters assigned by the authors represent only one tenth of the expected neuronal diversity.

Therefore, our aims were 1) to define molecular profiles for the remaining neuron classes and 2) to determine new neuron-specific promoters. To these ends, a heuristic approach was used where we matched a molecular atlas made of ~800 neuron class-specific marker genes [2] to the scRNA-seq dataset generated by *Cao et al* [1]. We performed a clustering analysis aiming to segregate ~100 cell-clusters assigned to the expected ~100 neuronal identities. Through two clustering iterations, we segregated 76 neuronal clusters and assigned 58 clusters to a single neuron class, from which we extracted the enriched genes. Based on these expression profiles, we designed 15 new neuron-specific promoters validated *in vivo*. Finally, we explored whether and how the most differentially expressed genes drive neuronal diversity in *C. elegans*. We observed that sensory machineries genes are driving the sensory neurons diversity, while neuromodulatory genes were highly diverse in interneurons and genes involved in membrane potential tuning were over-represented in motor neurons.

## Results

### Matching the clustering analysis to a molecular atlas

Each neuron class is theoretically detectable by a specific combinatorial gene expression signature. To assign each cell cluster to an anatomically defined neuron class in *C. elegans*, we designed an automated assignment strategy where the genes expressed in each cell cluster are compared to a curated molecular atlas extracted from <u>Wormbase</u> [2]. For each of the 112 neuron classes present in L2 stage, a combinatorial gene expression signature made of 19 to 116 marker genes was generated from ~800 expression patterns at the single cell resolution (**S1 Table**). While these expression patterns do not necessarily capture the accurate expression of each of the respective gene loci, this database provided us a unique proxy to assign objectively a cell-cluster to a neuron class.

From the original dataset generated by *Cao et al.* [1], we extracted 9078 cells from all clusters expressing the neuron markers *egl-21, egl-3, ida-1, sbt-1, unc-104, unc-31, ric-4, snt-1* and *unc-10* [18]. Our clustering was realized using the Louvain algorithm implemented in Seurat R package [19]. Two parameters mostly affected the clustering output: the number of Principal Components (PCs) and the Resolution. Each PC essentially represents a ‘metagene’, combining information across a set of correlated genes. The Resolution parameter sets the ‘granularity’ of the clustering: increased Resolution facilitates clusters segregation. These two parameters were optimized in order to identify a maximum of clusters assigned to a single neuron class.

Using Louvain algorithm over 92 PCs and a Resolution of 4, we identified 64 cell-clusters visualized with *t-*distributed Stochastic Neighbour Embedding (t-SNE) (Fig 1a). Each cluster was assigned to a cell identity –neuron class or tissue– based on the cell markers presents in **S1 table**. We used a combined *z-score* (Stouffer’s method) for all cells in a given cluster. Neuron classes or tissues are ranked by the median value of their combined *z-scores*. A sequential *t*-test from the highest to lowest ranked neuron class indicates the best assignment. For example, the cluster 20 is assigned to PLN and ALN, but not SDQ (Figs 1b and 1c). These 64 identified cell clusters included 53 clusters corresponding to neurons and 11 clusters corresponding to glial, hypodermal, pharyngeal and muscle cells (Fig 1a). The comparison of the 54 segregated neuron clusters with the 112 signatures of each neuron class allowed us to directly assign 22 clusters to a single neuron class (Figs 1d and 3). We observed that ASI or ASJ passed our threshold for assignment significance in clusters 38 and 61. However, ASJ matched noticeably better with cluster 38 and ASI with cluster 61, assigning ASI and ASJ to clusters 61 and 38, respectively (Fig 1e). We also observed that 7 clusters matched well with a subset of molecularly similar neuron classes (Figs 1e and 3). These clusters with mixed identities could be explained by the molecular proximity of the included neuron classes, making them hard to be segregated. For example, cluster 28 is assigned to three nociceptive neuron classes (ASH, PHA and PHB) co-expressing many chemosensory GPCRs and responding to similar cues [20]. Similarly, cluster 36 and cluster 4 contain a subset of sensory neurons sharing sensory machineries for oxygen or touch, respectively [21, 22]. Finally, 24 clusters were poorly or not assigned (Fig 1d).

**Figure 1:**
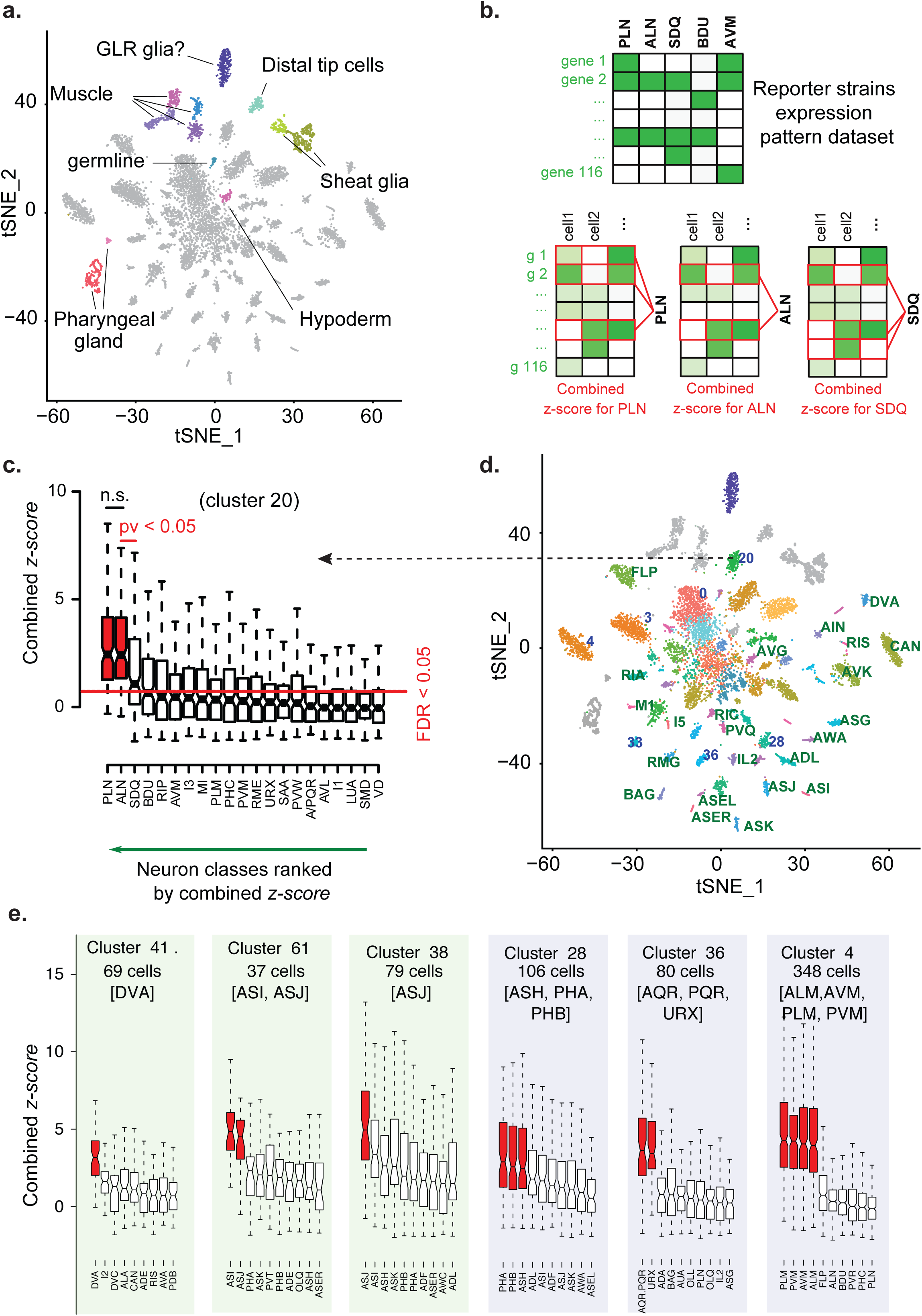
t-SNE projection and neuron-class assignment principle. **a.** Single cell profiles are projected in two dimensions using t-SNE. Cell-clusters corresponding to non-neuronal cells or to neurons are distinguished by colour and label. **b.** Neuron classes assignment to clusters using the *z-score* method. Gene’s *z-scores* from the centered scaled dataset are used to assign neuronal classes to the clusters. A combined *z-score* (Stouffer’s method) for all neuronal classes is obtained for each cell in the cluster, in this case PLN, ALN, SDQ and cluster 20. **c.** Neuron classes are treated as independent groups and ranked by the median value of their combined *z-scores*. In the notched box plots, the notches display the 95th confidence interval around the median (black horizontal line); the box contains the interquartile and the whiskers extend to the most extreme data points which are no more than 1.5 times the interquartile range from the box. A sequential *t*-test from the highest to lowest ranked neuron class indicates where the assignment of neuron classes should stop (p<0.05). An additional false discovery rate (FDR) filter of <5% is applied. **d.** Clusters assigned to a single neuron class are labelled in green (e.g., DVA); clusters assigned to a subset of molecularly similar neuron classes are labelled in blue (e.g. 36, corresponding to URX, AQR and PQR). **e.** Based on their combined *z-score*, clusters 41 is assigned to the DVA neuron class; clusters 61 and 38 are assigned to ASI and ASJ, respectively; clusters 28, 36 and 4 are assigned to a set of molecularly similar classes of sensory neurons, which we could not further segregate with our approach.

### A second clustering iteration identifies another 29 neuron classes

To validate the assignment results of the 22 clusters assigned with combined *z-scores* and to assess if a better cluster segregation could be obtained for the 24 poorly assigned clusters, each of the 64 parent clusters was isolated and treated independently for a second clustering iteration. Theoretically, this second iteration starting from a molecularly less complex collection of cells is expected to further divide each parent cluster. As we could not predict the best clustering parameters, we examined the results of 330 combinations of parameters from 3 to 92 PCs and Resolution from 0.1 to 3 (**S2 Table**). These 330 combinations often reached a consensus on the neuron classes’ content of the parent clusters: e.g. for the cluster 16, all 330 combinations identified one to many clusters assigned to the FLP neuron class (**S2 Table**). The quality of this re-clustering consensus as well as the *z-score*s for the neuron classes assigned to the parent cluster offers two ways to measure the assignment confidence (Figs 2 and 3). Interestingly, 93% ±19% (SD) of the re-clustering trials confirmed the neuron class of the 23 clusters directly assigned after the first iteration.

**Figure 2:**
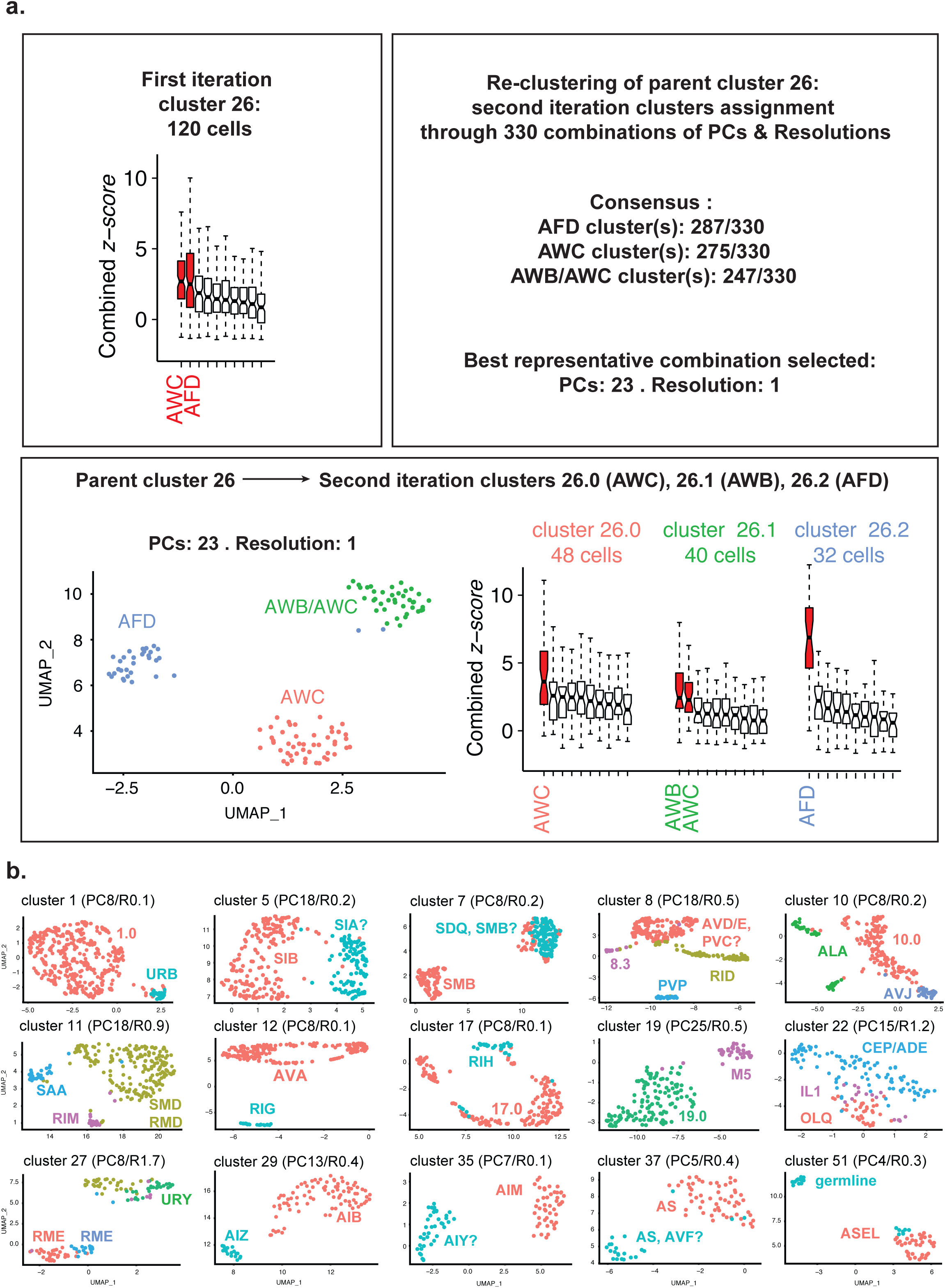
Second iteration clustering approach and results. **a.** Each parent cluster from the first iteration clustering was treated independently for re-clustering for all combinations of parameters ranging from 3 to 92 PCs and Resolution from 0.1 to 3. The neuron classes assigned to every sub-clusters generated by this large number of re-clustering trials are analysed. A consensus is reached for parent cluster 26: it suggests the re-clustering of cluster 26 should generate 3 sub-clusters. These 3 sub-clusters should be assigned to AFD, AWC and AWB/AWC. This result is observed for PCs: 23/Resolution: 1, used for representation and for gene expression lists. **b.** Uniform Manifold Approximation and Projection (UMAP) projections are used to represent the re-clustering of each parent cluster. Several of the second iteration clusters can be assigned to single neuron classes using the combined *z-scores*.

**Figure 3:**
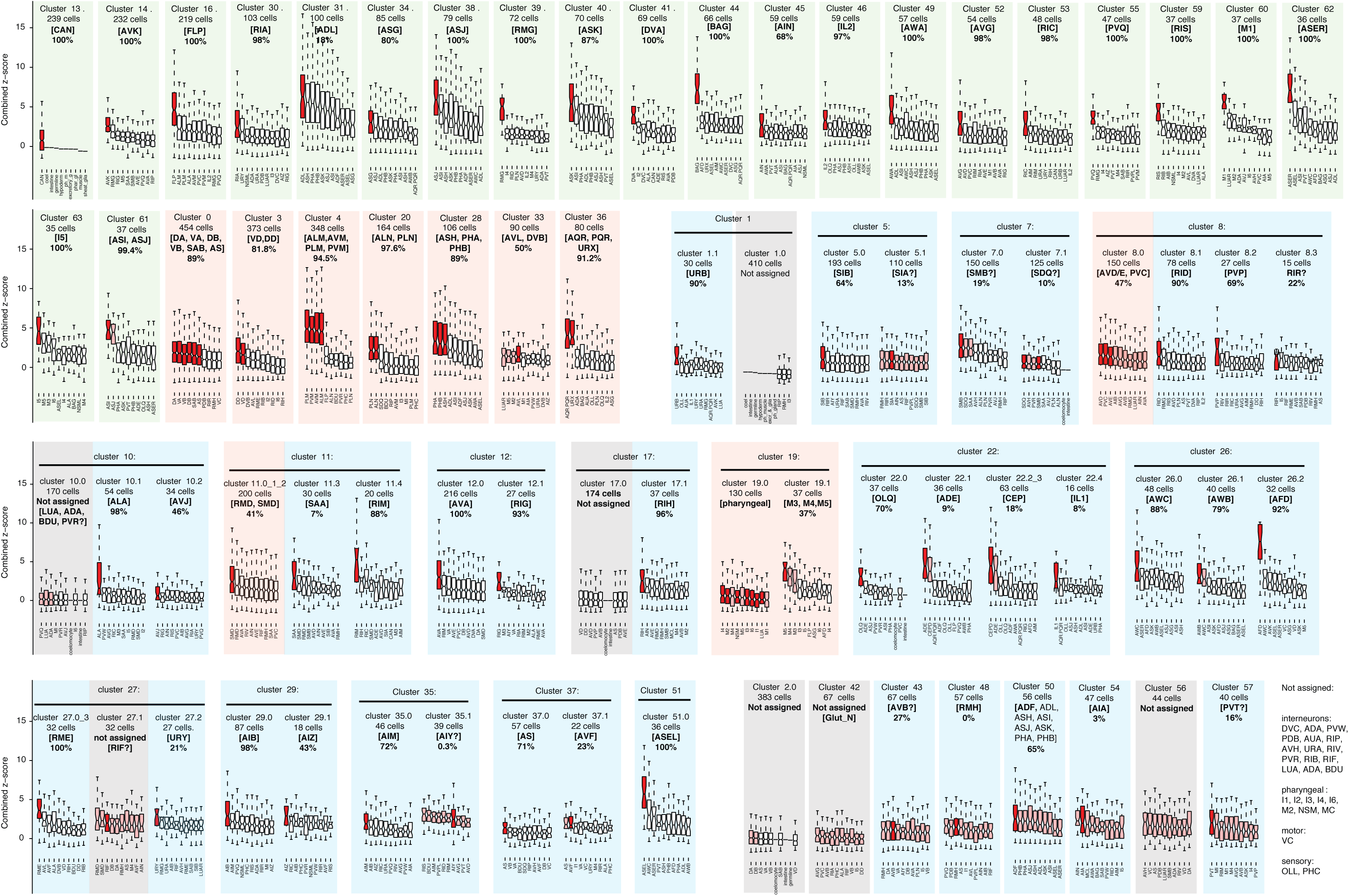
Clusters assignment by combined z-score and multiple iterations. For each cluster, the 10 best-ranked combined *z-scores* are displayed as well as the quality of the assignment consensus (%) reached by the re-clustering iterations of the parent cluster. These two assessments helped us to assign each cluster to a neuron class. The significant assignment by *z-scores* is displayed by the red colour within the notched box (pink when discarded for final assignment). The first iteration clusters directly assigned to a single neuron class identity are shaded in light green. The parent clusters assigned to a set of molecularly similar neuronal classes we could not segregate are shaded in orange. The parent clusters segregated by re-clustering are shaded in light blue. The parent and daughter clusters poorly assigned to a neuronal identity after the first iteration and not improved by re-clustering are shaded in grey.

In addition to this cluster assignment validation, the re-clustering also succeeded to segregate 16 of the poorly-assigned assigned parent clusters into better-assigned second iteration clusters. For example, the parent cluster 26 displayed a high *z-score* for AWC and AFD sensory neurons, suggesting it might contain several neuron classes. We observed that 285, 275 and 247 combinations of parameters segregated cluster 26 into 3 sub-clusters assigned to AFD, AWC and AWB/AWC, respectively (Fig 2a). This consensus is well represented by 23 PCs and a resolution of 1, splitting cluster 26 into AFD, AWC and a third sub-cluster assigned to both AWB and AWC (Fig 2a). Differential expression analysis shows AWB/AWC sub-cluster expresses AWB specific markers (e.g. *srab-24* and *srab-4*), suggesting this sub-cluster should have been assigned to AWB. A lack of differentially expressed genes between AWB and AWC in **S1 Table** potentially explains this inadequate assignment. Moreover, the re-clustering improved the *z-score* of AFD between the first and second iteration, suggesting the AFD sub-cluster is more homogeneously composed of AFD cells than the original parent cluster 26 (Fig 2a). This example above illustrates how re-clustering allowed us to split 16 of the poorly assigned clusters and/or to remove contaminating cells from the first iteration clusters. Altogether, the two clustering iterations coupled with the *z-score* assignment strategy automatically assigned 43 clusters to a single neuron class and 2 clusters to ASER and ASEL (Figs 2b and 3, **S1 folder)**. Removing the previously assigned neuron classes and examining the genes lists allowed us to assign another 11 clusters to a single identity. 11 clusters remained assigned to a set of 2 to 6 neuron classes molecularly too similar to be segregated (Fig 3). Three clusters of the first iteration and four clusters of the second iteration remained non-assigned (2, 42, 56, 1.0, 10.0, 17.0, and 27.1). Conversely, 26 neuron classes present in L2 stage are not confidently assigned to any cluster (e.g. AUA, URA, VC…). We assume the neurons of these 26 neuron classes might populate the seven non-assigned clusters or potentially contaminate the assigned clusters. Among the non-assigned neurons classes, several would require a better molecular signature in **S1 Table** to be assigned correctly to their corresponding cell-cluster.

### Expression profile validation for cluster 14 assigned to the AVK neuron class

We next extracted the genes expressed in each assigned cluster and computed their most enriched genes (**S3 Tables**). To validate one of these lists, we compared each of our clusters to the bulk sequencing of the AVK neurons purified by FACS-sorting [14]. Although both scRNA-seq and bulk sequencing have their own biases, we expect a strong match between cluster 14 we assigned to AVK and the AVK bulk sequencing. Indeed, the best match was observed between cluster 14 and the AVK bulk sequencing with 38 of the 100 most enriched genes present in both lists (p=9.83e-40). This overlap included the known AVK marker *flp-1* as well as new potential AVK markers such as *twk-47* (Fig 4).

**Figure 4:**
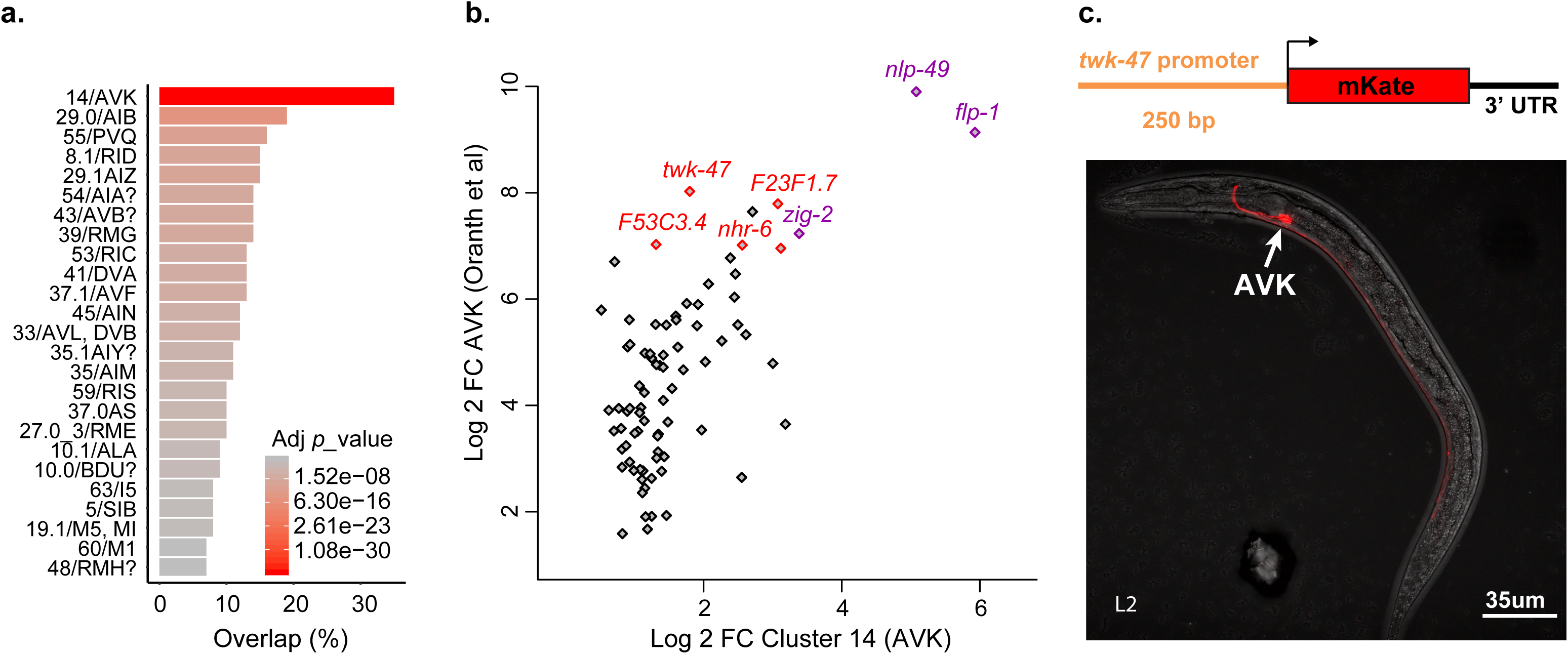
AVK profiling overlap and reporter strain design strategy. **a.** We observed an overlap of 35 genes between the 100 most enriched genes in cluster 14 assigned to AVK neuron class and the 100 most enriched genes from the AVK neuron bulk sequencing^16^ (adj. pv=9.82e-40; *p*-value adjusted by Bonferroni correction). **b.** The fold change comparison highlights the genes co-enriched in cluster 14 and in AVK bulk sequencing profille. The expression of *nlp-49* and *flp-1* mRNA in AVK were previously described (purple) [14, 33]. Several new potential AVK markers are highlighted in red, including *twk-47*. **c.** Generation of a reporter strain expressing a 250bp *twk-47* promoter fused to the mKate sequence. The confocal image shows the specific expression of mKate in the AVK neurons.

In *C. elegans*, the intergenic regions containing the promoters are short and in most case, a segment extending 1kb upstream the ATG was shown to drive the expression of a fluorescent protein reporter [23, 24]. We designed a reporter strain expressing mKate (a monomeric red fluorescent protein) under the control of a 250bp *twk-47* promoter. As expected, we observed the expression of mKate only in two neurons that we could identify as the two AVK neurons, supporting the accuracy of our gene enrichment list for cluster 14 (Fig 4).

### Gene enrichment lists provide 15 new neuron class-specific promoters

Identifying suitable promoters to specifically drive the expression of genetic tools in a given neuron class is an important step toward its functional characterization in vivo. Currently, only 29 neuron classes can be targeted using specific promoters (Table 1). Theoretically, each of the clusters assigned to a single neuron class could deliver potential specific markers within enriched genes. To validate the approach, we designed short (<1kb) and medium (<2kb) size promoters upstream the ATG of 17 genes highly enriched in clusters of interest (Fig 5 and **S1 Fig**). These promoters were fused to a monomeric EGFP (mEGFP) or mKate sequence in order to visualize their expression patterns *in vivo* (Fig 5 and **S1 Fig**). We confirmed the expression patterns of these promoters based on their cellular position, morphology, and/or with NeuroPAL, a polychromatic strain expressing four fluorescent proteins under overlapping drivers, designed to unambiguously identify individual neurons [25] (**S1 Fig**). First, we confirmed the validity of the clustering prediction by designing two promoters for the cluster 31, poorly assigned to ADL. We designed promoter-mKate fusion for two enriched genes: *T09B9.3* and *C18H7.6*. Both constructions showed mKate expression in the two ADL neurons, confirming the identity of cluster 31 (Fig 5 and **S1 Fig**). We next designed promoters for a set of parent clusters in order to identify new promoters for neuron classes lacking specific promoters: AIM, AVK, CAN, FLP, PVQ, IL2 and RMG neurons. Indeed, all the seven designed reporter strains showed a specific expression of the fluorescent protein in their corresponding neuron class (Fig 5 and **S1 Fig**). Finally, we designed promoters for a set of clusters assigned after the second clustering iteration. Among those, we identified new specific promoters for DD, OLQ, PVP and RIS neurons (Fig 5 and **S1 Fig**). Interestingly, no specific promoters were described until now for FLP without PVD, or IL2 D/V without IL2 L/R (Table 1).

**Table 1:** Neuron class-specific promoters. As a resource to drive expression selectively in neurons of interest, we gathered all known neuron class-specific promoters published by the *C. elegans* community as well as the ones we identified in this work.

| Neuron | Genes | Ref | Neuron | Genes | Ref |
| --- | --- | --- | --- | --- | --- |
| ADE_CEP | *flp-33* | *us* | FLP | *Y48G10A.6* | *us* |
| ADF | *srh-142* | [35] | FLP_PVD | *deg-3* | [57] |
| ADL | *svh-1* | [36] | FLP_PVD | *des-2* | [58] |
| ADL | *T09B9.3* | *us* | IL1 | *aqp-6* | [59] |
| ADL | *C18H7.6* | *us* | IL1 | *Y111B2A.8* | [24] |
| AFD | *gcy-8* | [37] | IL1 | *flp-3* | [60] |
| AIA | *gcy-28* | [38] | IL2 | *klp-6* | [61] |
| AIM | *nlp-70* | *us* | IL2LR | *C18F10.2* | *us* |
| AIY | *ttx-3* | [39] | OLQ | *ocr-4* | [62] |
| ALN_PLN | *C50F7.5* | *us* | OLQ | *ttll-9* | *us* |
| ASEL | *gcy-6* | [40] | PHA | *srg-13* | [46]- |
| ASEL | *gcy-7* | [40] | PHA | *gcy-17* | [41] |
| ASER | *gcy-4* | [41] | PVD_FLP | *F49H12.4* | [58] |
| ASER | *gcy-5* | [40] | PVD_FLP | *dma-1* | [63] |
| ASG | *gcy-15* | [41] | PVP | *ocr-3* | *us* |
| ASI | *str-3* | [42] | PVQ | *nlp-17* | *us* |
| ASJ | *sptf-1* | [43] | PVT | *zig-2* | WB |
| ASJ | *ssu-1* | [44] | RIA | *glr-3* | [64] |
| ASJ | *trx-1* | [45] | RIA | *glr-6* | [64] |
| ASK | *sra-7* | [46] | RIC | *tbh-1* | [65] |
| ASK | *sra-9* | [46] | RIM | *gcy-13* | [41] |
| ASK | *srg-2* | [46] | RIM | *cex-1* | [66] |
| ASK | *srg-8* | [46] | RIS | *srsx-18* | *us* |
| ASK | *srbc-64* | [47] | RIS | *flp-11* | [67] |
| ASK | *srbc-66* | [47] | RMD | *C42D4.1* | *us* |
| AVJ | *hlh-34* | [48] | RMG | *nlp-56* | *us* |
| AVK | *flp-1* | [49] | RMH | *sem-2* | [60] and *us* |
| AVK | *twk-47* | *us* | URX_AQR_PQR | *gcy-32* | [40] |
| AWA | *odr-10* | [50] | SAA | *rig-5?* | [60] |
| AWB | *srab-16* | [51] |  |  |  |
| AWB | *sru-38* | [51] |  |  |  |
| AWB | *str-44* | [51] |  |  |  |
| AWB | *str-163* | [51] |  |  |  |
| AWC OFF | *srsx-3* | [52] |  |  |  |
| AWC OFF | *srt-26* | [52] |  |  |  |
| AWC OFF | *srt-28* | [52] |  |  |  |
| AWC ON | *str-2* | [52] |  |  |  |
| AWC ON | *srt-45* | [52] |  |  |  |
| AWC ON | *srt-47* | [52] |  |  |  |
| BAG | *gcy-9* | [53] |  |  |  |
| BAG | *flp-17* | [54] |  |  |  |
| CAN | *pks-1* | *us* |  |  |  |
| DD | *ttr-39* | *us* |  |  |  |
| DVA | *twk-16* | [55] |  |  |  |
| DVC | *ceh-63* | [56] |  |  |  |

**Figure 5:**
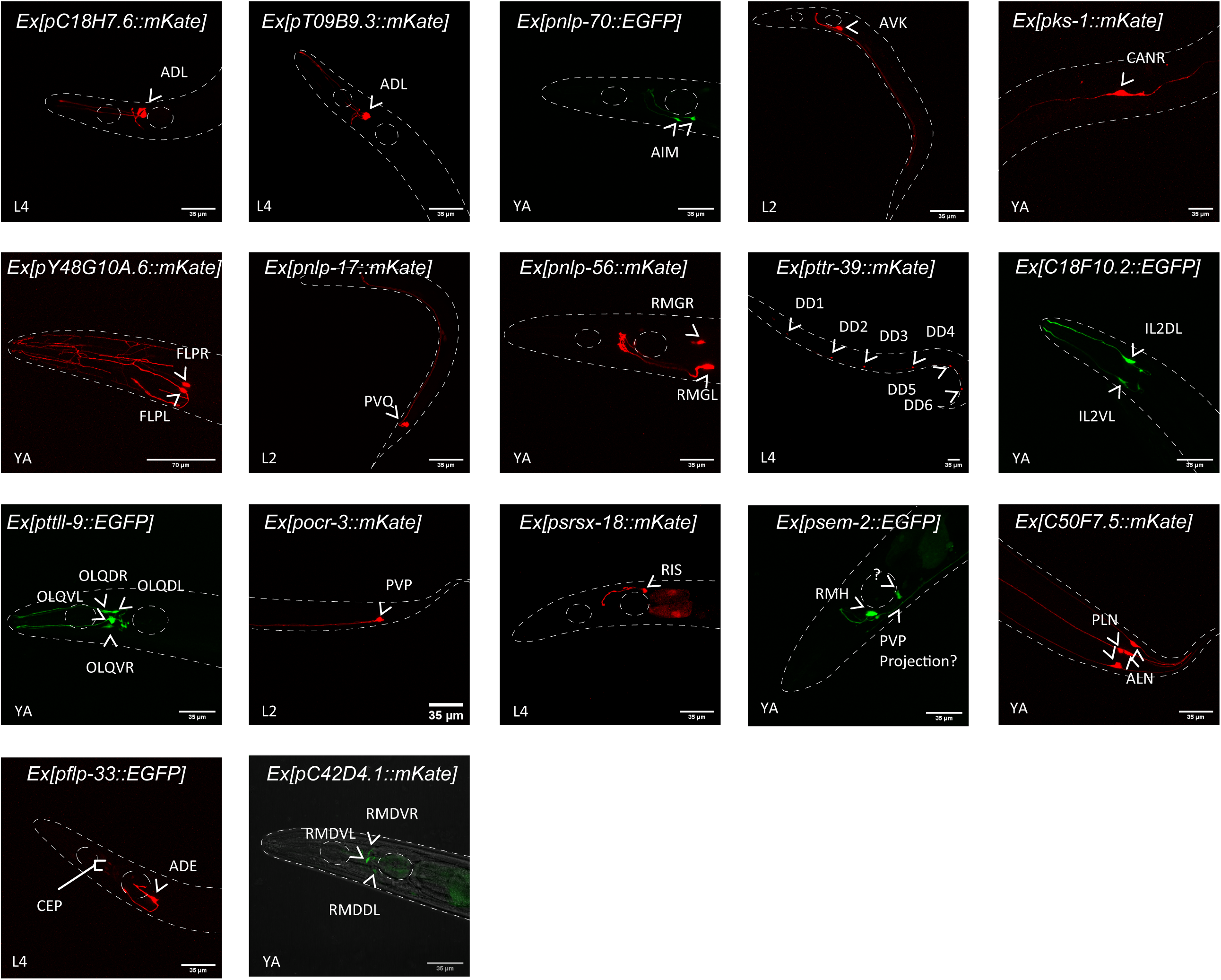
Confocal images of 17 designed reporter strains. The expression patterns of mKate (red) or mEGFP (green) driven by the corresponding gene promoter were analysed by confocal microscopy. White arrowheads indicate the neuron-classes expressing the fluorescent proteins. The animals were imaged between L2, L4 or young adult (YA) developmental stages. Details concerning neuron class identification, promoters’ size and primers’ sequences are described in **S1 Fig**.

This promoter screen also helped us to assign an identity to the poorly assigned cluster 48. A reporter strain for *sem-2*, enriched in cluster 48, showed a strong expression in RMH, suggesting cluster 48 can be assigned to RMH or does include RMH (Fig 5 and **S1 Fig**). Some of the selected promoters highlighted more than a single neuron class but confirmed our predictions. For example, *C50F7.5* was enriched in the cluster 20, assigned to the oxygen sensory neurons ALN and PLN. Accordingly to our prediction, the *C50F7.5-*mKate reporter strain showed a specific expression of mKate in both ALN and PLN neurons, allowing us to define a new marker for ALN and PLN (Fig 5 and **S1 Fig**). Similarly, *flp-33* was expressed in the dopaminergic neurons ADE and CEP, but more importantly enriched in the ADE neurons cluster. In fact, the *flp-33*-mKate reporter strain strongly highlighted the ADE neurons and showed a weak expression of mKate in the CEP neurons (Fig 5 and **S1 Fig**). Nevertheless, some of the selected promoters failed to highlight a single neuron class: *C42D4.1* was enriched in a cluster assigned to SMD and RMD. The reporter strain showed indeed an expression in RMD but also showed a weak expression in another non-identified neuron (Fig 5 and **S1 Fig**).

### Gene Ontology (GO) analysis reveals functional diversification within neuronal modalities

The most differentially expressed genes between neuron classes likely contribute to their functional diversity. We performed a GO analysis on the most differentially expressed genes between clusters. Similarly to the mouse brain [6], we observed that genes involved in sensory machineries, membrane potential, synaptic function and neuromodulation were over-represented in these most differentially expressed genes. To assess whether there was a difference in enriched genes ontology between neuronal modality (sensory-, inter- and motor neurons), we classified these gene lists accordingly to neuronal modality. To get an equivalent number of genes per modality (~200 genes per modality), we extracted the top 9 most enriched genes from each sensory neuron class, the top 8 most enriched genes from each interneuron class and the top 20 ones from each motor neuron class [9]. Interestingly, these genes were not equally distributed between neuron classes (Table 2). The genes required for the sensory machineries (guanylate-cyclases, chemosensory receptors, Transient Receptor Potential channels …) mostly contributed to the diversity of the sensory neuron classes. The genes involved in neuromodulation (bioamine synthetic pathways, neuropeptides and their receptors) mostly contributed to the diversity of the interneuron classes. Finally, the chloride and potassium channels contributed mostly to the diversity of the motor neuron classes (Table 2). These observations suggest the mode of functional diversification may vary accordingly to each neuronal modality.

**Table 2:** Differential expression of functional genes between neuronal modalities. The molecular and biological functions of the 200 most neuron class-specific genes were analysed independently for the 24 sensory neurons clusters, 27 interneurons clusters and 11 motor neurons clusters. Genes are segregated accordingly to their putative or demonstrated functions in *C. elegans*.

| Sensory neurons |  | Interneurons |  | Motorneurons |  |
| --- | --- | --- | --- | --- | --- |
| 55 sensory-like genes | 12 chemoreceptor GPCRs (srx, srd, srh, srj, srbc, srab, srv, odr-genes) | 15 sensory-like genes | 6 chemoreceptor GPCRs (srg, srx, srsx, sre, sru-genes) | 1 sensory-like genes | 0 chemoreceptor GPCR (srg, srx, srsx, sre, sru-genes) |
|  | 3 neuroglobins (glb-genes) |  | 4 neuroglobins (glb-genes) |  | 1 neuroglobin (glb-genes) |
|  | 10 mechanosensors (del, mec-genes) |  | 1 guanylate cyclase (gcy-genes) |  |  |
|  | 26 guanylate cyclases (odr, daf, gcy-genes) |  | 2 TRP channel (ocr & trp-genes) |  |  |
|  | 3 TRP channels (ocr, osm genes) |  | 2 other (tmc and lite-genes) |  |  |
|  | 1 other (cng-gene) |  |  |  |  |
| 18 neuro-modulatory genes | 16 neuropeptides (flp, nlp, ins, ntc, capa-genes) | 50 neuro-modulatory genes | 43 neuropeptides (pdf, flp, nlp, ins-genes) | 12 neuro-modulatory genes | 10 neuropeptides (flp, nlp, ins-genes) |
|  | 2 bioamine synthesis (dat, cat-genes) |  | 2 bioamine synthesis (tdc, tbh-genes) |  | 2 GPCR (gar-genes) |
|  |  |  | 5 GPCRs (ador,frpr, mgl, ser, daf-genes), probably more. |  |  |
| 3 membrane potential | 3 potassium channels (kcnl, twk-genes) | 2 membrane potential | 2 potassium channel (srk, kvs-genes) | 9 membrane potential | 8 potassium channels (twk, irk, kcnl-genes) |
|  |  |  |  |  | 1 chloride channel (clh-gene) |
| 4 synaptic genes | 4 ligand-gated channels (acr, glr, deg, des-genes) | 9 synaptic genes | 9 ligand-gated channels (glr, nmr, des, acr-genes) | 15 synaptic genes | 7 ligand gated channels (acr, glr, lgc, acc, sol-genes) |
|  |  |  |  |  | 8 presynaptic function (cho, unc, glt, sup, snt, ace-genes) |
| 14 developmental genes | 3 axon guidance (zig, eva-genes) | 14 developmental genes | 6 axon guidance (ncam, slt, hen, zig, eva, unc-genes) | 14 developmental genes | 3 axon guidance (unc, lad, madd-genes) |
|  | 11 transcription factors (sptf, xbx, nhr, pax, lim, ceh, egl, mec, dac, ahr, vab-genes) |  | 8 transcription factors (nhr, lim, ceh, fozi, mls, tab-genes) |  | 11 transcription factors (unc, ceh, fozi, bnc, jun, sox, tag, lin-genes) |

## Discussion

Although neuronal classification is a useful abstraction to neurophysiologists, the border between classes is difficult to resolve. Contrasting with complex brains, the full diversity of the *C. elegans* nervous system is already anatomically established at the single cell resolution. In order to generate the molecular profiles for each neuron class, we attempted to match the molecular information from scRNA-seq data to these anatomically defined neuron classes in *C. elegans*. Building on *Cao et al*. [1], our heuristic approach succeeded to assign 48 additional clusters to a single-neuron class. However, the simple one-to-one correspondence between neuron classes and cell-clusters did not apply to 11 of our clusters, assigned to a subset of neuron classes. This could be explained by limitations in expression profiling, preventing us to identify genes segregating these classes. Alternatively, some neuronal classes might be molecularly too similar to each other to be further separated despite their known differences in connectivity and functions (e.g. the mechanosensory neuron classes ALM, PLM, AVM, and PVM) [22]. In contrast, we easily segregated the chemosensory neuron class ASE in two sub-classes ASER and ASEL, known to display symmetric morphology but asymmetric gene expressions and functions [26]. These two examples above illustrate the abstract border between neuron classes and sub-classes.

We anticipate our work will serve as a resource to facilitate future works on *C. elegans* nervous system. Indeed, only a few markers were available for each neuron class, and only a few neuron classes were profiled in depth [11–14]. Our work provides a partial molecular profile for the 58 clusters we assigned to a single neuron class and 11 combined profiles for clusters assigned to a subset of neuron classes. Despite the shallow sequencing, zero-inflation effects and cluster size affecting the analysis [27], we confirmed the quality of the expression profiles by the comparison with bulk sequencing and by the generation of neuron-specific promoters. Although the regulatory *cis-*elements of every gene are not known, the restricted expression patterns of the 17 promoters we tested here support the strength of this approach to deliver additional specific promoters. Partly because of the lack of specific promoters, little is known about the functions of some neuron classes (e.g. ALN, PLN, AIM and CAN). The 15 new markers we validated here already provide to the community a set of new specific promoters, of which 10 are for neuron classes without any known specific marker to this day. Some of these new promoters will be valuable tools to manipulate neurons hard to access currently. For example, optogenetic manipulation of the DD neurons with the available promoters required a specialized illumination system [28]; manipulating specifically RMG or FLP required to use expression overlap between two promoters [29, 30].

Recently, two other datasets corresponding to ~85.000 embryonic cells and to ~52.000 L4 larval stage neurons were sequenced by [31] and [32], respectively. The authors used a small number of specific markers to assign each cell clusters to their corresponding neuron class. We observed an important overlap (~85%) between our assigned clusters and the corresponding ones in the L4 stage animal [32], supporting the results of both assignment efforts and suggesting additional markers for specific promoters (**S4 Table and S2 Folder**). Similarly, we observed an important overlap (~90%) between our assigned clusters and the neuron classes emerging at the end of the embryonic development [32] (**S4 Table**). While some neuron classes seem to converge in their expression profiles only after circuit assembly [32], these observations suggest most of them retain their transcriptomic identities throughout post-developmental stages.

Finally, we analysed the most differentially enriched gene ontologies between neuronal modalities (sensory-, inter- and motor neurons). While it is intuitively expected that genes involved in the distinctive sensory machineries diversify the sensory neurons, we observed that other specific gene families contributed to the diversity in interneurons and motor neurons. On one hand, genes involved in neuromodulation were highly diverse between interneurons, suggesting a complex neuromodulation soup is generated by each interneuron and differentially controls these neurons. On the other hand, genes involved in the tuning of membrane potential were highly diverse between motor neurons, suggesting their diversity involves a tight control of their membrane potential. Altogether, our work will contribute to the molecular characterisation of the neuronal diversity of *C. elegans*, a crucial step toward the understanding of its neuronal network.

## Methods

### Dataset pre-processing

As starting dataset, we used the neuron clusters defined in [1] and containing 7603 cells. This dataset is a sub-selection of clusters coming from a whole worm dataset using the neuroendocrine markers *egl-21, egl-3, ida-1* and *sbt-1*. We added 1475 cells included in clusters expressing the neuron markers *unc-104, unc-31, ric-4, snt-1* or *unc-10* [18], further expanding the dataset to 9078 cells. Dataset clustering was realised using Seurat v3, and visualized using t-SNE or UMAP. Louvain algorithm was used for all clustering settings using Seurat v3 default parameters, unless otherwise indicated.

### Clustering parameters

Two parameters mostly affect Louvain algorithm clustering output in Seurat: the number of PC used and the Resolution used. We ranked the PCs based on the percentage of variance they explain but did not observe any obvious elbow (**S1 Folder**). To determine dimensions and Resolution for the first iteration, we set the dimensions by repeating the clustering analysis with 40 to 100 PCs and Resolution ranging from 1 to 8. To evaluate the quality of each clustering result, we designed an automatic assignment of neuronal identities inside clusters using genes expressed in neurons (see cluster assignment method). We chose PC and Resolution parameters by selecting the clustering condition giving the maximum amount of clusters with one identity not being repeated in the clustering set. The best clustering parameters were set to 92 PCs dimensions and a Resolution value of 4; as a result, the dataset was divided in 64 clusters including 23 directly assigned to only one neuron class.

### *z-scores* assignment method

Each cell-cluster is compared to the signature of each 102 neuron class defined from **S1 Table**. These signatures are made by a combination of 19 to 116 genes. First, for each cell in the dataset, the *z-score* of the signature genes representing a specific neuron class is calculated and combined using Stouffer’s method [34], according to the formula:

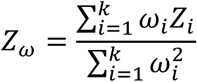

*Z*_*ω*_ is the combined *z-score* for a particular cell in the dataset for a specific neuron class signature. *k* represents the signature genes defining the neuron class. *Z*_*i*_ is the *z-score* for a cell for a particular gene (*i*) present in the neuron class signature, previously calculated by centring and scaling the dataset. *ω*_*i*_ is the weight for gene (*i*) calculated as (1-the fraction of cells expressing gene (*i*)). Hence, each neuron class is defined by a specific group of combined *z-scores* calculated from the cells present in a given cluster. For each cluster, the combined *z-scores* were ranked by median values. A sequential *t*-test from the highest to lowest ranked neuron class was applied; all the neuron classes before a significant difference (p<0.05) were assigned to the cluster. A second independent filter was applied by calculating the FDR for all neuron classes identities over the cluster; the threshold for FDR was set to <5%. Finally, only the neuron classes selected by both methods were assigned to a cluster.

### Second clustering iteration

We performed a second round of clustering on the 64 parent clusters obtained above. This second iteration potentially segregates non-homogenous clusters, allowing the assignment of more clusters to their neuron class. Single-cell profiles coming from parent clusters were independently re-clustered with Seurat v3. Multiple re-clustering trials were generated for Resolution values from 0.1 to 3 and for dimensions values from 3 to 92. These conditions generated a total number of clustering trials over each parent cluster ranging from 151 to 330, depending on the cell number within the parent clusters. In each trial, the identities of the daughter clusters are assigned with *z-score* assignment method (see above) and counted in **S2 Table.** The neuron classes most often detected in the parent cluster represent a consensus on the identity(ies) present in the parent clusters. The consensus on the identities present in each parent cluster was used to detect the neuron class most likely present in each parent cluster. Based on these predictions, we chose the lowest PC and Resolution among the most representative clustering parameters.

### Comparison between gene lists

The read count for each gene was extracted for each cluster. We also computed lists of enriched genes compared to all other neuronal clusters of our dataset. The AVK bulk sequencing profile was generated by FACS-sequencing and compared to all neurons to compute the enriched genes [14]. The comparison between the top 100 *p*-value ranked genes for each neuronal cluster versus AVK bulk sequencing was calculated by running a hypergeometric test for the overlap between lists considering a total of 5589 genes found in both datasets. The multiple comparisons between our enrichment lists and the enrichment lists from 31 and 32 were done similarly for the top 100 *p*-value ranked genes and considering a total of 5968 genes found in both datasets.

### Reporter strain generation

The primers used to generate the promoters collection and the amplified regions size are described in **S1 Fig**. The promoter sequences were amplified from N2 genomic DNA, purified from gel extraction then fused with mKate or mEGFP sequence followed by the 3’UTR sequence of *unc-54* or *let-858*, respectively. 30 ng/µL of the fusion PCR, 30 ng/µL of 1kb Plus DNA mass ladder (Invitrogen) and 30 ng/µL of the injection co-marker *punc-122::*mKate were co-injected in D1 animals to generate transgenic animals. Injections were carried out in N2, OH12312, OH13083, CZ13799 and/or OH15262. After injection, 3 to 5 independent transgenic strains were maintained for at least three generations before imaging.

### Microscopy

In order to identify the neurons expressing the reporter transgene, we performed confocal microscopy imaging (Zeiss LSM 510 and Zeiss LSM 780) with 20x or 40x objective. Animals were anesthetized with 25mM sodium azide in M9 solution, transferred to 2% agarose mounting slides and imaged within 30min of mounting. The images were analysed using Fiji imaging software. The expression patterns were analysed at L2, L4 and YA developmental stages for 3-5 independent transgenic strains. In order to identify the expression pattern, we used cell position, morphology and/or the NeuroPAL strain (OH15262), used as described [25]. The images obtained are available in **S1 Fig** together with the detailed evidences used to identify the neurons.

### GO analysis

We generated gene ontology lists for the genes enriched in:

- **32 sensory neurons/ 24 clusters:** 44.0: BAG, 62.0: ASER, 51.0:ASEL, 26.0: AWC, 26.1: AWB, 26.2: AFD, 36.0: URX, AQR, PQR, 46.0: IL2, 40.0: ASK, 50.0: ADF, 38.0: ASJ, 31.0: ADL, 61.0: ASI, 49.0: AWA, 34.0: ASG, 22.1: OLQ, 22.2: ADE, 22.3: CEP, 22.4: IL1, 27.2: URY, 28.0: ASH, PHB, PHA, 20.0: ALN,PLN, 16.0: FLP, 4.0: ALM, PLM, AVM, PVM.
- **30 interneurons/ 27 clusters:** 42.0: AVH?, 33.0: AVL, DVB, 37.1: AVF, 1.1: URB, 55.0: PVQ, 53.0: RIC, 29.0: AIB, 29.1: AIZ, 12.0: RIG, 12.1: AVA, 57.0: PVT, 35.0: AIY, 35.1: AIM, 14.0: AVK, 59.0: RIS, 45.0: AIN, 41.0: DVA, 52.0: AVG?, 43: AVB, 8.0: AVD, AVE, PVC, 8.1: RID, 8.2: PVP, 8.3: RIR?, 17.1: RIH, 30.0: RIA, 10.1: ALA, 10.2: AVJ. We did not include 39.0: RMG, 8.1: RID, 11.4: RIM, 37.1: AVF, 48: RMH that are interneuron but also motorneurons.
- **17 motorneurons/ 11 clusters:** 5.0: SIB, 5.1: SIA, 7.0: SMB, 7.1: SDQ, SMB, 27.0: RME, 37.0: AS, 11.3: SAA, 11.0: SMD, RMD, 0.0: VA, VB, DA, DB, SAB, 2.0: VA, DA, 3.0: VD, DD.

### Nematode strains

The primers used to generate the reporter strains are described in **S1 Fig**.

OQ180: CZ13799; ulb180 *Ex[psrsx-18::mKate; unc-122p::mKate]*

OQ181: CZ13799; ulb181 *Ex[pttr-39::mKate; unc-122p::mKate]*

OQ182: N2; ulb182 *Ex[pC18F10.2::mEGFP]*

OQ183: N2; ulb183 *Ex[pC41A3.1::mKate]*

OQ184: N2; ulb184 *Ex[pC50F7.5::mKate]*

OQ185: N2; ulb185 *Ex[pflp-33::mKate]*

OQ186: N2; ulb186 *Ex[pnlp-56::mKate; unc-122p::mKate]*

OQ187: N2; ulb187 *Ex[pocr-3::mKate; unc-122p::mKate]*

OQ188: N2; ulb188 *Ex[pY48G10A.6::mKate; unc-122p::mKate]*

OQ189: OH12312; ulb189 *Ex[pC18H7.6::mKate; unc-122p::mKate]*

OQ190: OH12312; ulb190 *Ex[pnlp-17::mKate; unc-122p::mKate]*

OQ191: OH12312; ulb191 *Ex[pT09B9.3::mKate; unc-122p::mKate]*

OQ192: OH12312; ulb192 *Ex[pY48G10A.6::mKate; unc-122p::mKate]*

OQ193: OH13083; ulb193 *Ex[ptwk-47::mKate; unc-122p::mKate]*

OQ194: OH15262; ulb194 *Ex[pC42D4.1::mEGFP]*

OQ195: OH15262; ulb195 *Ex[psem-2::mEGFP]*

OQ196: OH15262; ulb196 *Ex[pttll-9::mEGFP]*

OQ197: OH15262; ulb197 *Ex[pY58G8A.5::mEGFP]*

CZ13799: juIs76 *[unc-25p::GFP + lin-15(+)] II*

N2 CGC

OH15262: otIs669 V [NeuroPAL]

OH13083: otIs576 *[unc-17(fosmid)::GFP + lin-44::YFP]; him-5(e1490) V*

OH12312: *pha-1(e2123);* otIs388 *[eat-4(fosmid)::SL2::YFP::H2B+pha-1(+)]III; him-5(e1490) V*

## Supporting information

Supplementary Folder 1

Supplementary Folder 2

## Acknowledgment

Some strains were provided by the CGC, which is funded by NIH Office of Research Infrastructure Programs (P40 OD010440). We acknowledge the Shendure, Trapnell, and Waterston laboratories for the scRNA-seq dataset; WormBase for the molecular atlas; the Hobert lab for the Neuropal strain, the Satija Lab for the Seurat v3 R package. RL and MO performed experiments, RL analysed the database, MD supervised the analysis and PL supervised the experiments and analysis. PL, RL, MO & MD wrote the article.

