## Supplementary Figure 1 for "Combining single-cell RNA-sequencing with a molecular atlas unveils new markers for *C. elegans* neuron classes"

| N° | Neuron ID | Cluster | Gene | avg_logFC | pct.1 | pct.2 | Size (bp) | FW Primer | RV Primer |
|---|---|---|---|---|---|---|---|---|---|
| 1 | ADE_CEP | 22.1 | *flp-33* | 4.367916081 | 0.389 | 0.003 | 345 | ATACTTGCTTCCACCCGAAA | GGTAGGGGGACCCTGGAA |

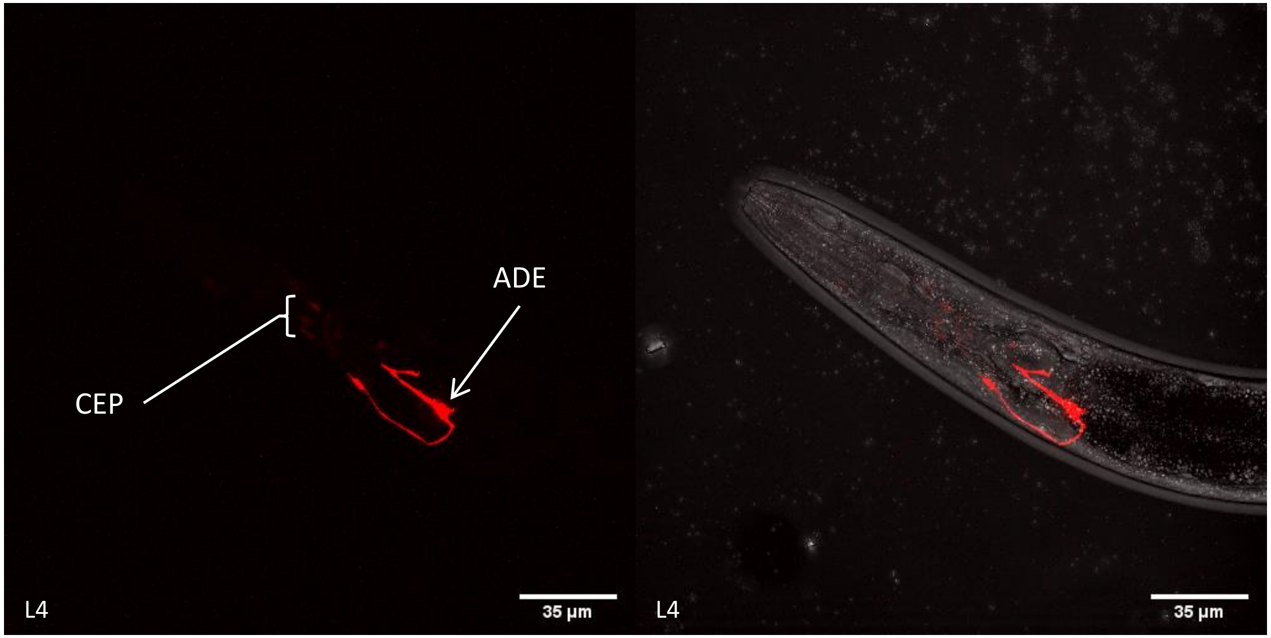


ADE_CEP (*flp-33*) – N2

Left lateral view of the ADE and CEP neurons: the two ADE cell bodies are located posteriorly and ventrally from the posterior pharynx. Their dorsal processes send a small branch laterally, while their ventral processes run into the ventral ganglia. The barely visible 3 CEP (VL, VR and one D) cell bodies are located between the anterior and the posterior pharynx, with dendritic processes reaching the tip of the animal.

| N° | Neuron ID | Cluster | Gene | avg_logFC | pct.1 | pct.2 | Size (bp) | FW Primer | RV Primer |
|---|---|---|---|---|---|---|---|---|---|
| 2 | ADL | 31 | *C18H7.6* | 3.275017302 | 0.49 | 0 | 196 | GTGATCAGTTCACGCCACAG | TTTTTCTTCAGGTTGACTCTCTGAC |

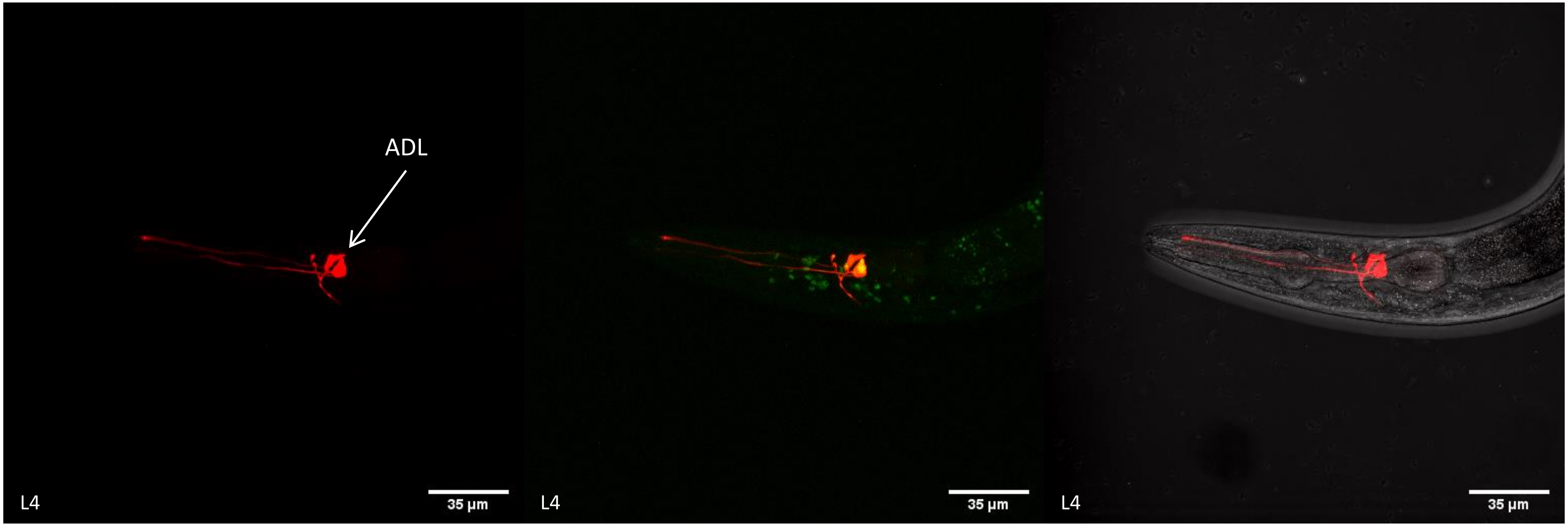


ADL (*C18H7.6*) – OH12312

Lateral view of the ADL neurons: the two cell bodies are located in the lateral ganglion, dorsally from the nerve ring (NR) and anteriorly from the posterior pharynx, collocalizing with peat-4::YFP signal which is in accord with their glutamatergic signature. Their longest dendritic processes run ventrally, while their shortest processes meet ventrally. Finally, their dual ciliated sensory endings run into the amphid channel.

| N° | Neuron ID | Cluster | Gene | avg_logFC | pct.1 | pct.2 | Size (bp) | FW Primer | RV Primer |
|---|---|---|---|---|---|---|---|---|---|
| 3 | ADL | 31 | *T09B9.3* | 3.693844231 | 0.64 | 0 | 145 | CAAAATCAGCTGCCAAAAA | TTTGAGTTCATTTCGGCTTTTAGT |

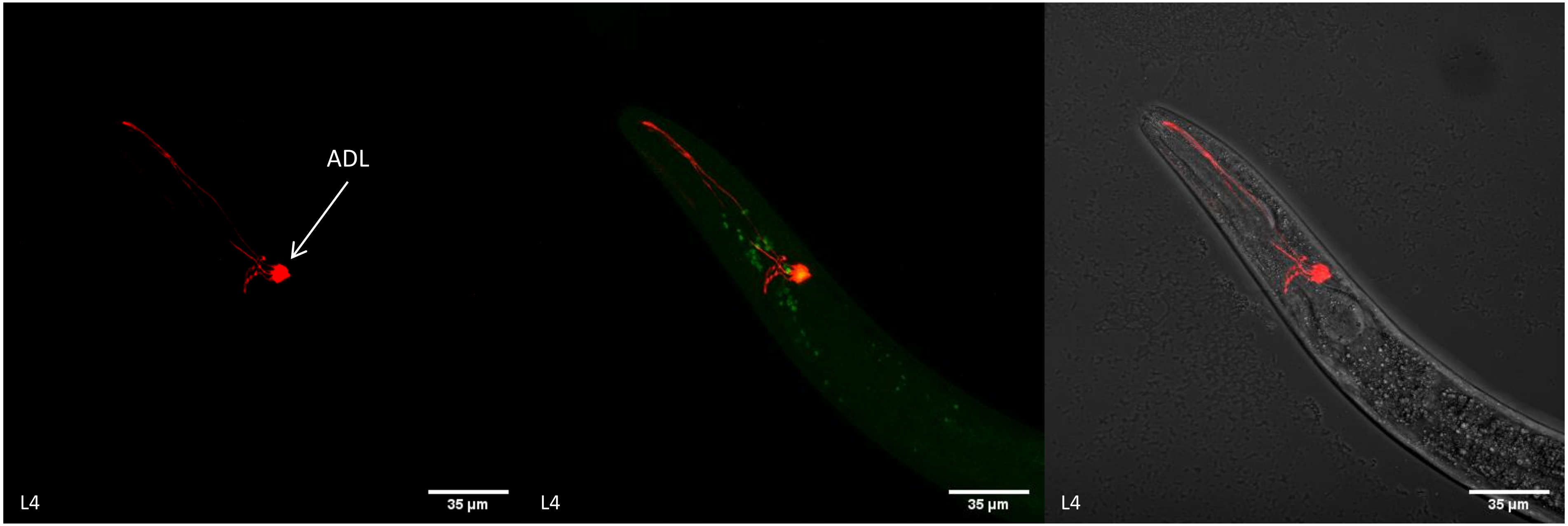


ADL (*T09B9.3*) – OH12312

Lateral view of the ADL neurons: the two cell bodies are located in the lateral ganglion, dorsally from the nerve ring (NR) and anteriorly from the posterior pharynx, collocalizing with peat-4::YFP signal which is in accord with their glutamatergic signature. Their longest dendritic processes run ventrally, while their shortest processes meet ventrally. Finally, their dual ciliated sensory endings run into the amphid channel.

| N° | Neuron ID | Cluster | Gene | avg_logFC | pct.1 | pct.2 | Size (bp) | FW Primer | RV Primer |
|---|---|---|---|---|---|---|---|---|---|
| 4 | AIM | 35 | *Y58G8A.5* | 3.956936273 | 0.543 | 0.003 | 1275 | TTCTGATGCACCCATTACATT | TCTGAAATGAGGAAAATGTTAGACG |

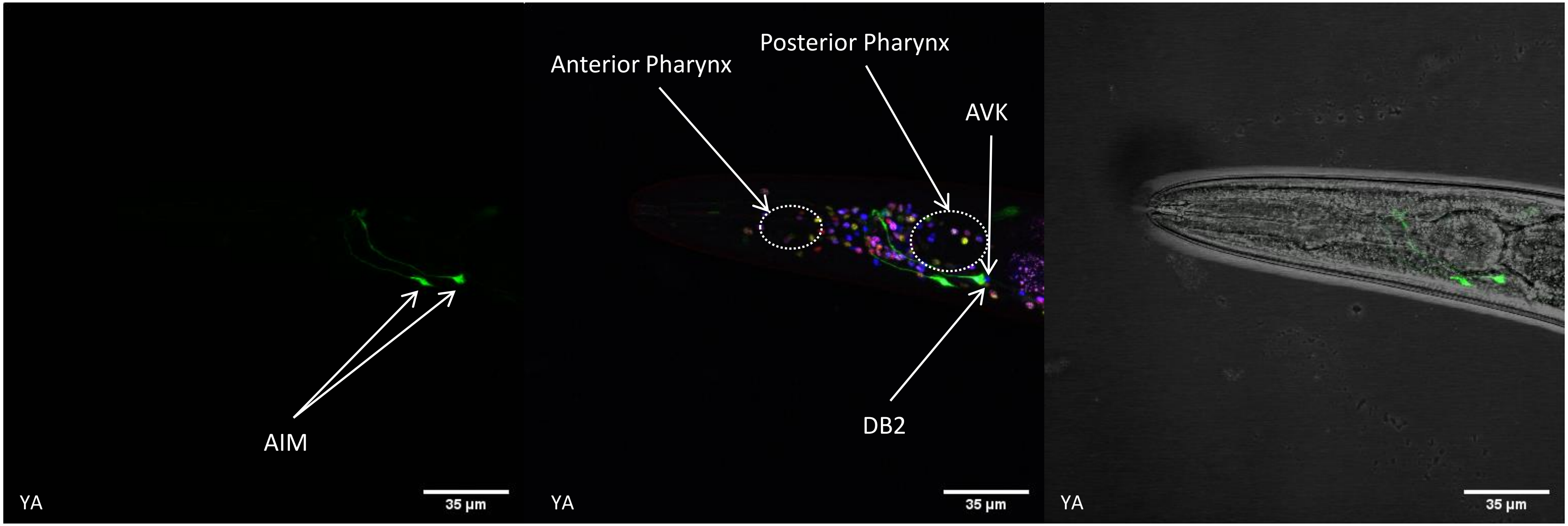


AIM (*Y58G8A.5*) – OH15262

Lateral view of the AIM neurons: the two cell bodies are located in the ventral ganglion, dorsally from the posterior pharynx and anteriorly from the AVK and DB2 neurons identified in the NeuroPAL strain. Both dendritic processes enter the NR from the ventral side and meet each other in the dorsal part.

| N° | Neuron ID | Cluster | Gene | avg_logFC | pct.1 | pct.2 | Size (bp) | FW Primer | RV Primer |
|---|---|---|---|---|---|---|---|---|---|
| 5 | ALN_PLN | 20.0 | *C50F7.5* | 2.754673512 | 0.22 | 0 | 614 | GCAAAGTGGGGAATCTAGCC | TCTGAAAATAGCTCACTGCAATT |

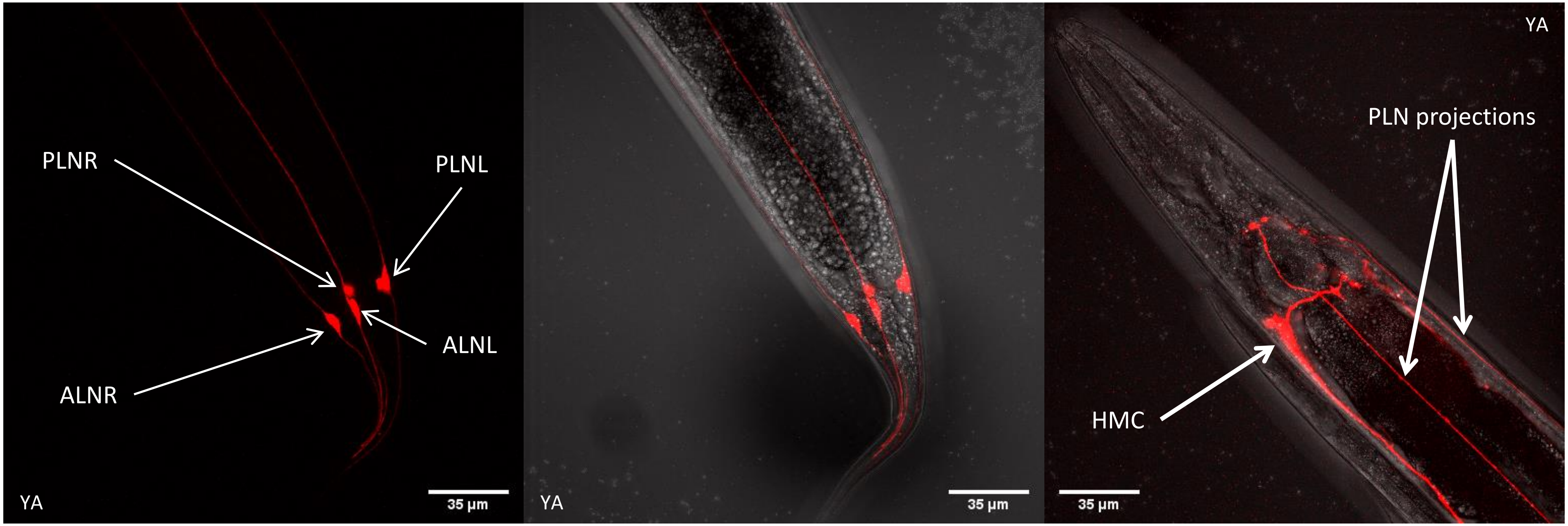


ALN – PLN (*C50F7.5*) – N2

Ventral view of the ALN and PLN neurons: the four cell bodies are located in the lumbar ganglion region, all sending a projection into the tailspike. The two PLN neurons cell bodies are located slightly anteriorly from the two ALN neurons cell bodies. The PLN projections perform a “hook” shape before entering the ventral cord. The ALNR projection is masked with the HMC cell projection.

| N° | Neuron ID | Cluster | Gene | avg_logFC | pct.1 | pct.2 | Size (bp) | FW Primer | RV Primer |
|---|---|---|---|---|---|---|---|---|---|
| 6 | AVK | 14 | *twk-47* | 1.800434757 | 0.134 | 0 | 250 | TTTATTGTTAGACCCTACTGAGGAA | TTTAGCGTATAGTATTCGGATAGAAATG |

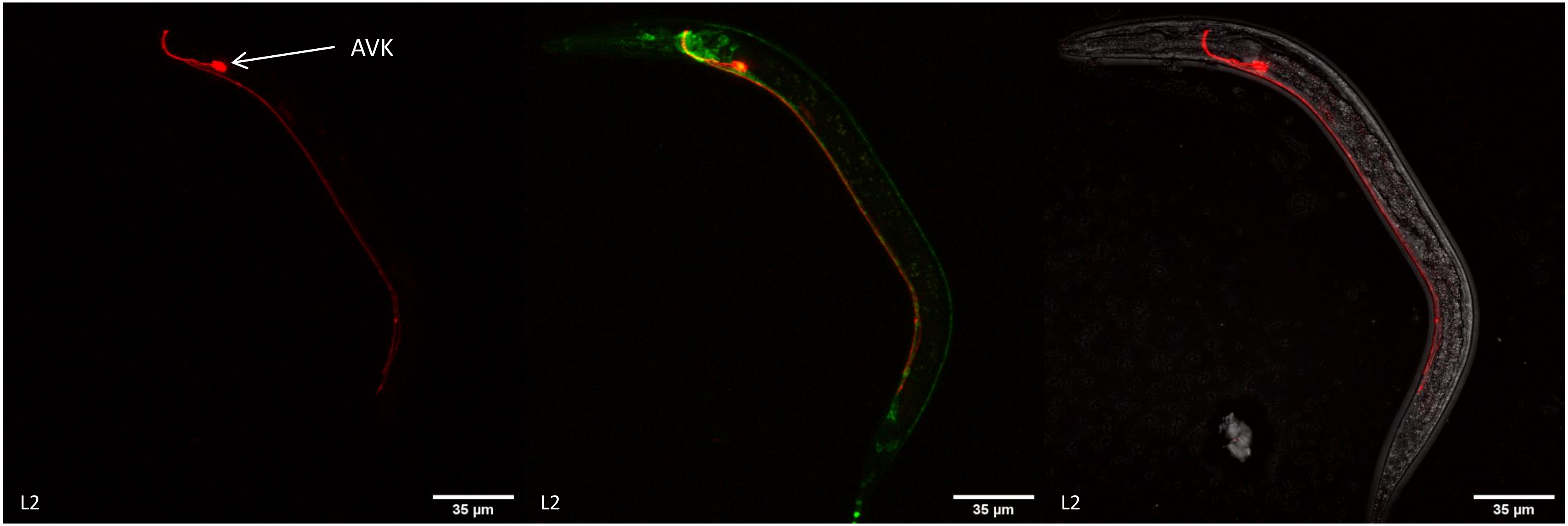


AVK (*twk-47*) – OH13087

Lateral view of the AVK neurons: the two cell bodies are located in the ventral ganglion region, ventrally and posteriorly from the posterior pharynx. Their projections enter the NR ventrally, perform a complete loop around the NR before going along the ventral cord and reach the tail where they stop before the rectum. The two cell bodies collocalize also with punc-17::GFP signal, in accord with the cholinergic signature of the corresponding cluster in our dataset.

| N° | Neuron ID | Cluster | Gene | avg_logFC | pct.1 | pct.2 | Size (bp) | FW Primer | RV Primer |
|---|---|---|---|---|---|---|---|---|---|
| 7 | CAN | 13 | *C41A3.1* | 4.949827134 | 0.925 | 0.003 | 541 | GAGGGTGCGGAGAAAGATTA | TTTCTCCAAATCTTAATACAAATTATA |

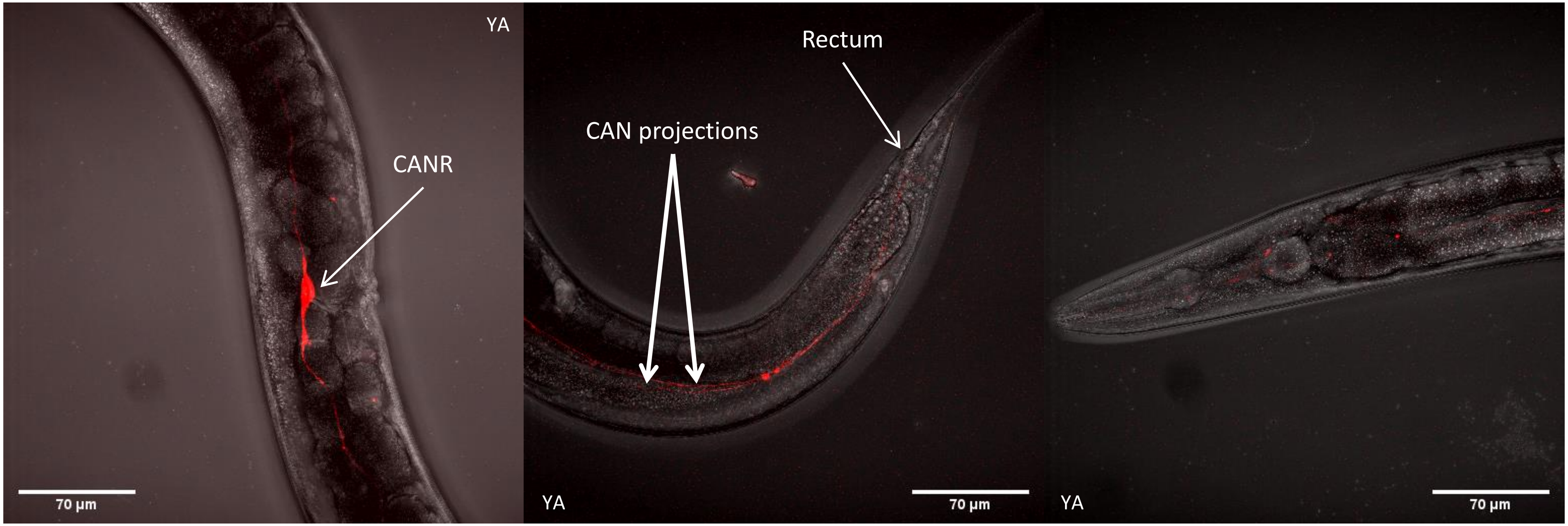


CAN (*C41A3.1*) – N2

Right lateral view of the CAN neuron: the right CAN neuron cell body is located in the mid body in front of the vulva of the animal (the left cell body is masked within the stack). The anterior projections stop before the anterior pharynx, while the posterior projections stop at the rectum.

| N° | Neuron ID | Cluster | Gene | avg_logFC | pct.1 | pct.2 | Size (bp) | FW Primer | RV Primer |
|---|---|---|---|---|---|---|---|---|---|
| 8 | DD | 3 | *ttr-39* | 2.619919683 | 0.226 | 0.013 | 252 | TGATGTTTGTAAAAGTCCTGCAA | GATTTTTTGTTTTAACAAAATTGTGA |

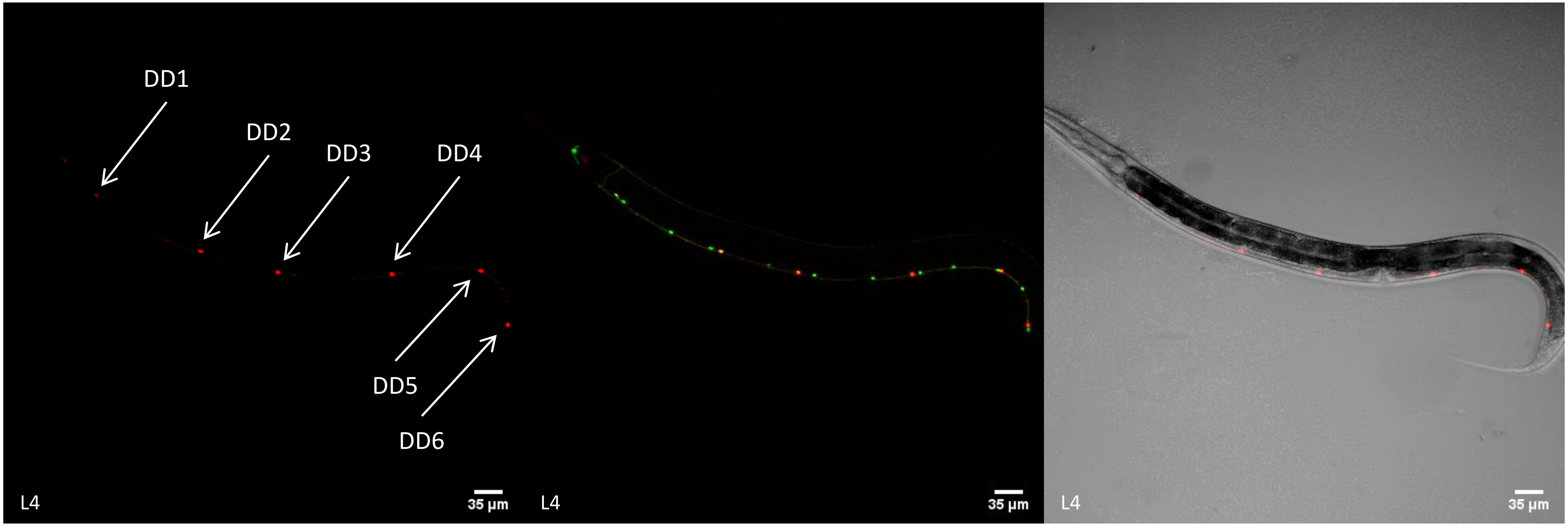


DD (*ttr-39*) – CZ13799

Left lateral view of the DD neurons: the six cell bodies are located within the ventral nerve cord, collocalizing with punc-25::GFP signal. Note that their commissures are hidden in the stacks.

| N° | Neuron ID | Cluster | Gene | avg_logFC | pct.1 | pct.2 | Size (bp) | FW Primer | RV Primer |
|---|---|---|---|---|---|---|---|---|---|
| 9 | FLP | 16 | *Y48G10A.6* | 2.988471297 | 0.425 | 0.007 | 315 | TTGGATTTTTGGCACTTTTT | TTTTTTTTGGGAAAAAACTTTTTCT |

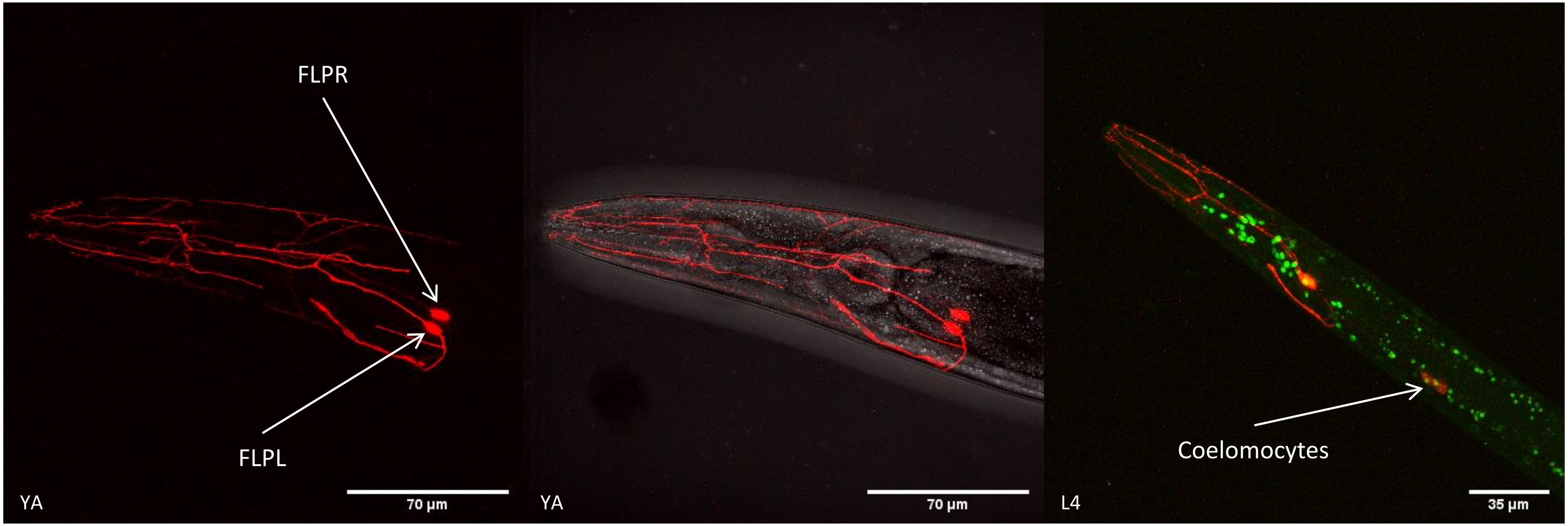


FLP (*Y48G10A.6*) – N2

Lateral view of the FLP neurons: the two cell bodies are located posteriorly from the posterior pharynx. Their multidendritic arborization structure is typical and covers the head and the neck of the animal. In the third image, the cell body is collocalizing with peat-4::YFP signal which is in accord with their glutamatergic signature.

| N° | Neuron ID | Cluster | Gene | avg_logFC | pct.1 | pct.2 | Size (bp) | FW Primer | RV Primer |
|---|---|---|---|---|---|---|---|---|---|
| 10 | IL2V/D | 46 | *C18F10.2* | 2.858799578 | 0.22 | 0 | 1035 | TCACAATCGGAAACACCAGA | TCAGAGCAAAAGAAATACAAATGAATT |

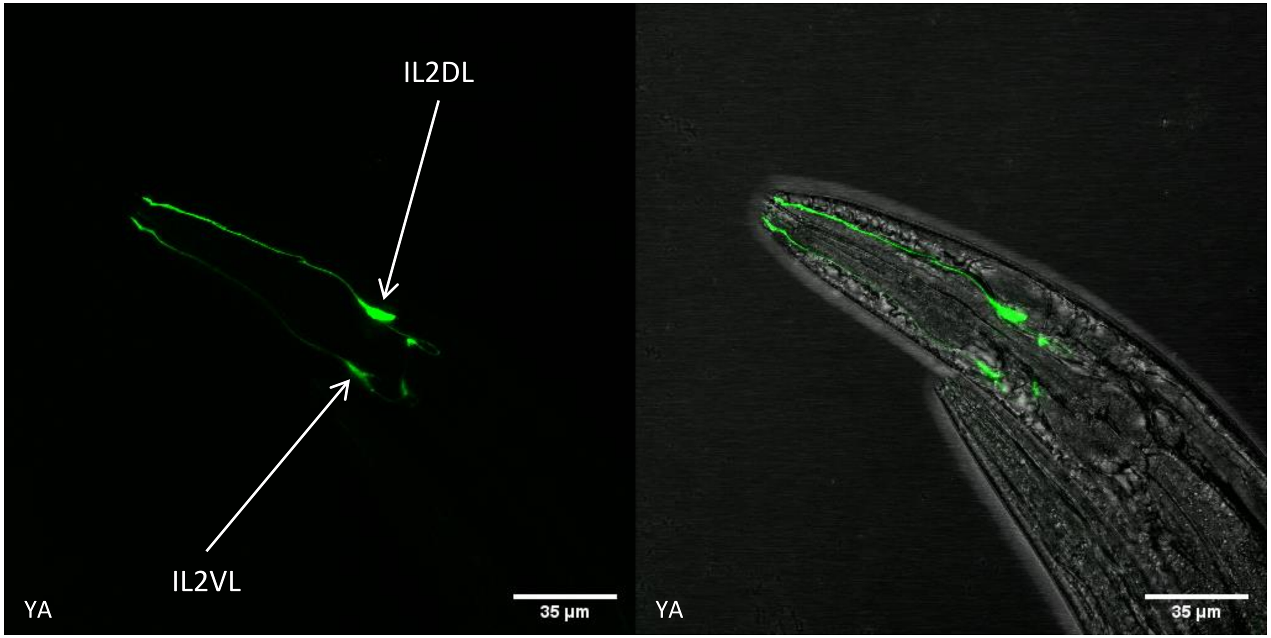


IL2V/D (*C18F10.2*) – N2

Left lateral view of the IL2V/D neurons: the four cell bodies are located in the anterior ganglion region right posteriorly from the anterior pharynx (the IL2VR and DR neurons are masked within the stacks). Their posterior projections are small and perform the typical "hook" shape of the IL2 neurons. Their anterior projections run into the tip of the animal.

| N° | Neuron ID | Cluster | Gene | avg_logFC | pct.1 | pct.2 | Size (bp) | FW Primer | RV Primer |
|---|---|---|---|---|---|---|---|---|---|
| 11 | OLQ | 22.0 | *ttll-9* | 3.129875591 | 0.27 | 0.001 | 501 | CACCAAGTTCGCTTATCAGTTG | GATTTTCACAACAAAAAAAATCCA |

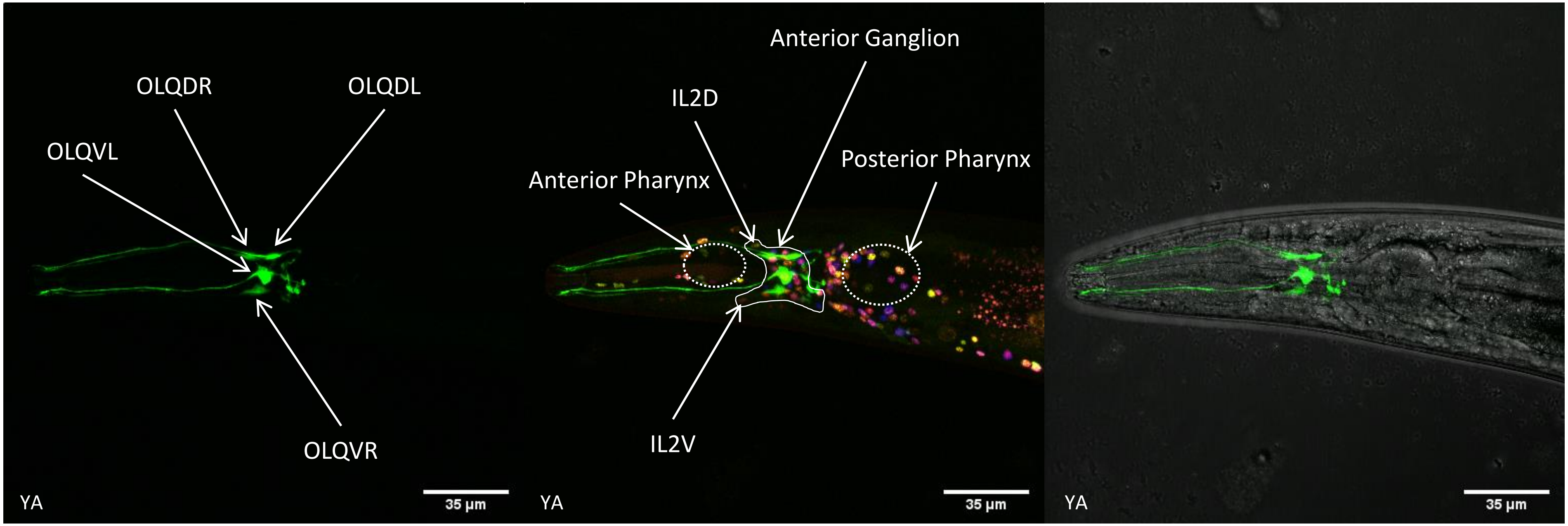


OLQ (*ttll-9*) – OH15262

Left lateral view of the OLQ neurons: the four cell bodies are located in the anterior ganglion, with typical posterior striated rootlets and anterior projections ending in the nose of the animal. Anteriorly from the OLQ neurons, the IL2 neurons were identified in the NeuroPAL strains.

| N° | Neuron ID | Cluster | Gene | avg_logFC | pct.1 | pct.2 | Size (bp) | FW Primer | RV Primer |
|---|---|---|---|---|---|---|---|---|---|
| 12 | PVP | 8.3 | *ocr-3* | 2.624196155 | 0.3 | 0.001 | 467 | TTTGGGAATGAAGTTGTTTAGAA | GTTGTTAGTAAGAGTAAAATGAGGAAG |

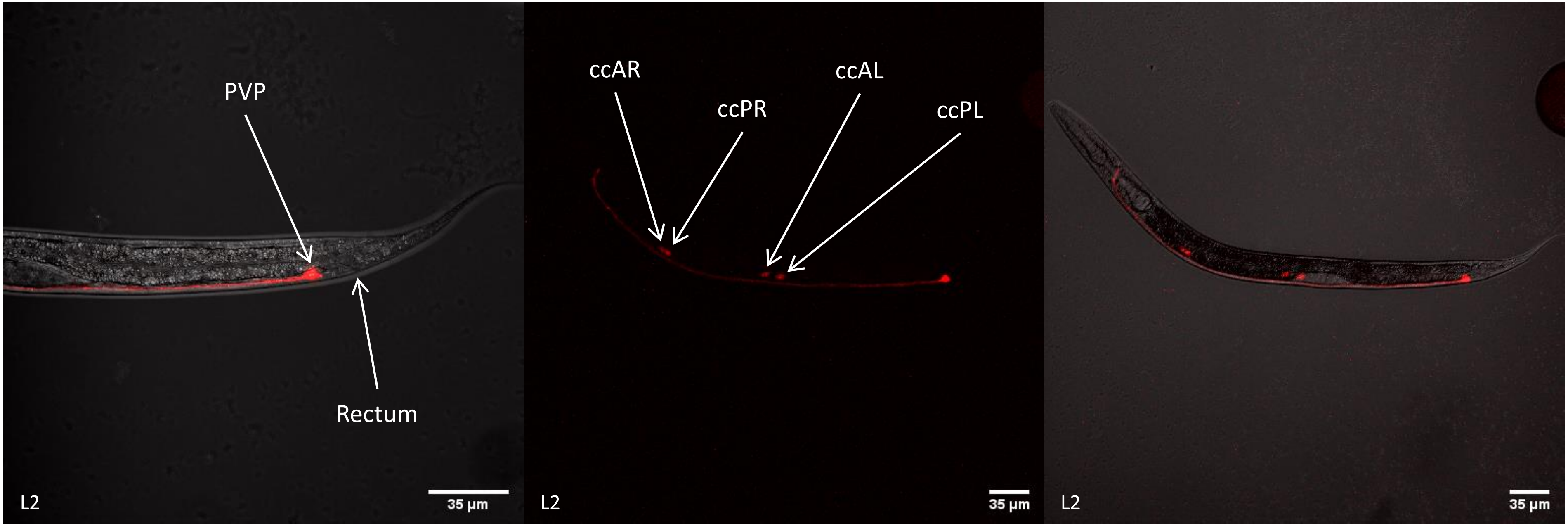


PVP (*ocr-3*) – N2

Lateral view of the PVP neurons: the two cell bodies (superposed each other) are located in the pre-anal ganglion region, just before the rectum. Their dendritic projections run ventrally before entering the NR where they reach the dorsal region.

| N° | Neuron ID | Cluster | Gene | avg_logFC | pct.1 | pct.2 | Size (bp) | FW Primer | RV Primer |
|---|---|---|---|---|---|---|---|---|---|
| 13 | PVQ | 55 | *nlp-17* | 5.501333306 | 0.936 | 0.001 | 339 | TGCAAAATTCCAAAAGGTGA | ATTTTCTGTGAAAAAGCCTGACT |

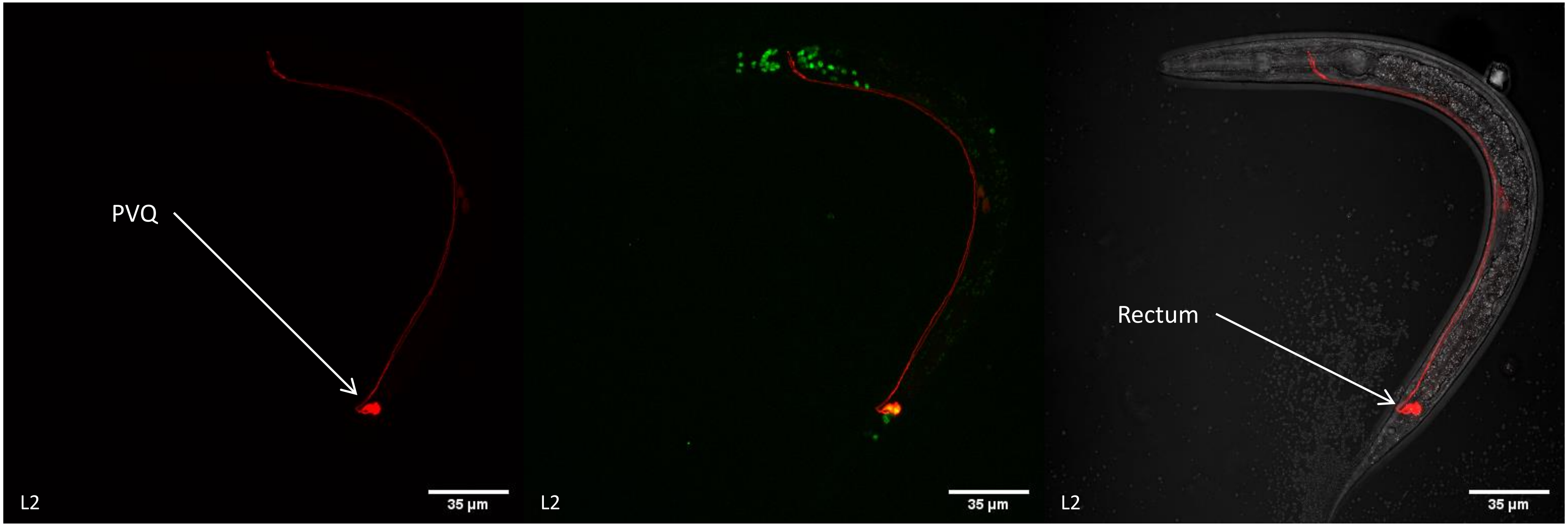


PVQ (*nlp-17*) – OH12312

Left lateral view of the PVQ neurons: the two cell bodies are located in the lumbar ganglion region, right after the rectum. They collocalize with peat-4::YFP signal, supporting their glutamatergic signature and their dendritic projections run ventrally before entering the NR where they reach the dorsal region.

| N° | Neuron ID | Cluster | Gene | avg_logFC | pct.1 | pct.2 | Size (bp) | FW Primer | RV Primer |
|---|---|---|---|---|---|---|---|---|---|
| 14 | RIS | 59 | *srsx-18* | 2.282598468 | 0.189 | 0 | 300 | GGGATATTCTGAAAGTCATCGAA | TGTCTGAAAATATAGATGGTAATGAAG |

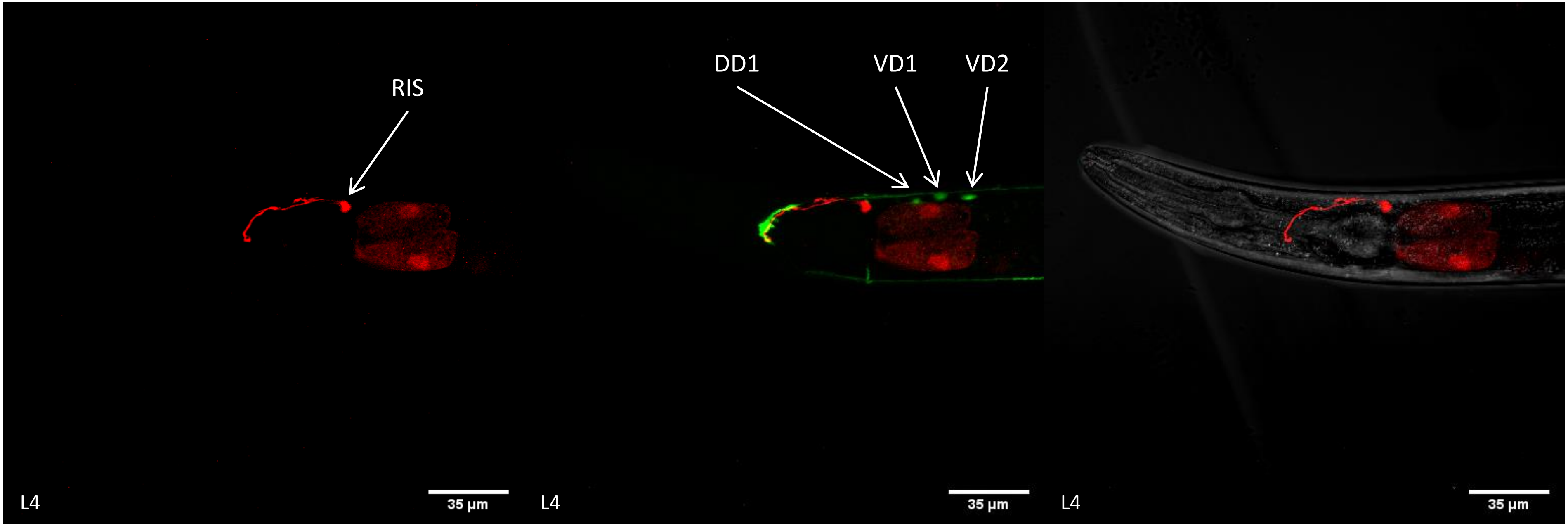


RIS (*srsx-18*) – CZ13799

Right lateral view of the RIS neuron: the cell body is located in the ventral ganglion, posteriorly from the posterior pharynx. Its dendritic process shows a loop around the NR from ventral to dorsal part. Note that the cell body doesn't collocalize with punc-25::GFP signal, this latest lacking the RIS, AVL and RME neurons. The three green neurons cell bodies correspond from left to right to DD1, VD1 and VD2.

| N° | Neuron ID | Cluster | Gene | avg_logFC | pct.1 | pct.2 | Size (bp) | FW Primer | RV Primer |
|---|---|---|---|---|---|---|---|---|---|
| 15 | RMD | 11.0 | *C42D4.1* | 2.018061865 | 0.102 | 0.009 | 565 | TCGCGGAACTCTAGTAAATTGAA | CCTGAAATCAACCATTTCCATC |

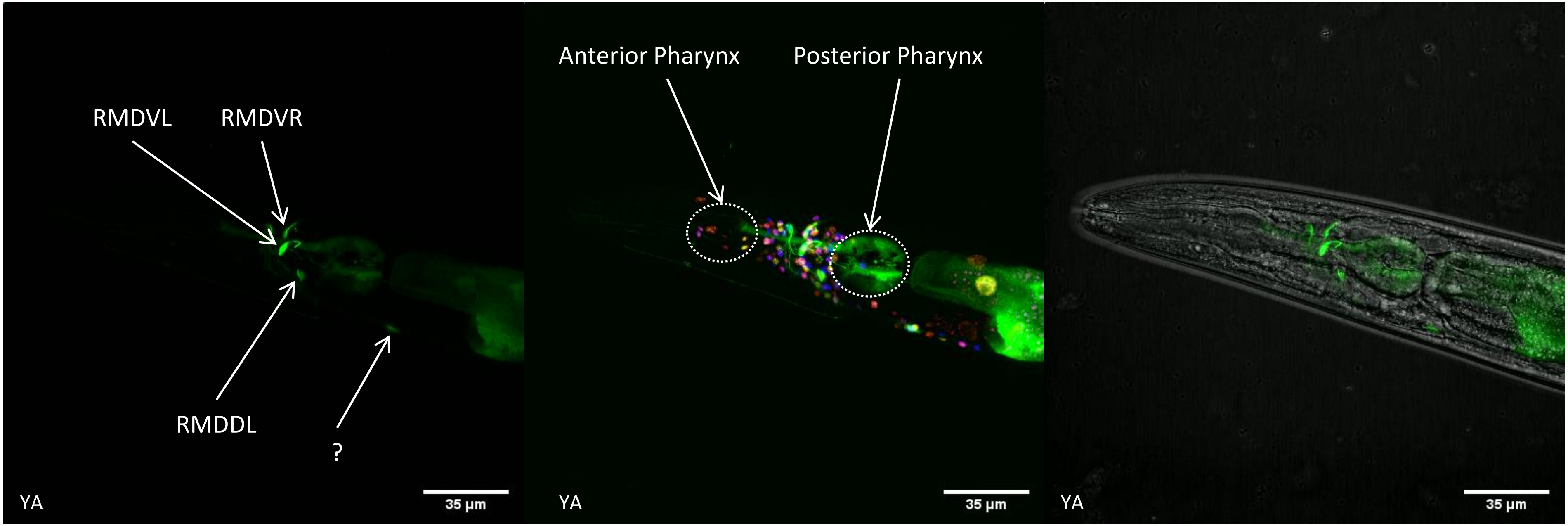


RMD (*C42D4.1)* – OH15262

Left lateral view of the RMD neurons: the two RMDV cell bodies are located in the lateral ganglion, in the most anterior part. The two RMDD cell bodies are located in the ventral ganglion (the RMDDR is too weak and masked in the stacks). We were able to distinguish them from SMDV and D with the NeuroPAL strain colour assignation: they perfectly collocalize with the anterior pink/red cells while SMDV and D are yellow/green cells. Note that an unidentified neuron is located in the retro-vesicular ganglion.

| N° | Neuron ID | Cluster | Gene | avg_logFC | pct.1 | pct.2 | Size (bp) | FW Primer | RV Primer |
|---|---|---|---|---|---|---|---|---|---|
| 16 | RMG | 39 | *nlp-56* | 4.610979105 | 0.694 | 0 | 721 | TTCCAAATCCGAACTTCCAG | CTGGAAGAGTTGAATCATATGG |

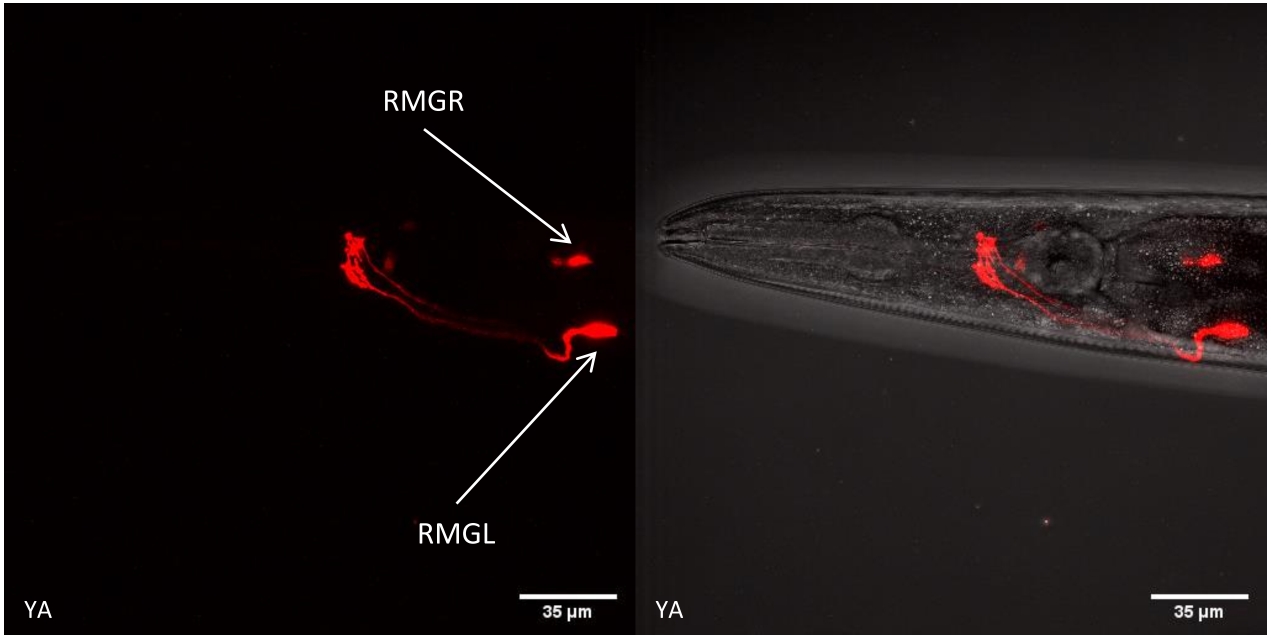


RMG (*nlp-56*) – N2

Left-dorsal view of the RMG neurons: the two cell bodies are located posteriorly from the posterior pharynx. Their dendritic projections perform a small "hook" ventrally before entering the NR in an ipsilateral manner.

| N° | Neuron ID | Cluster | Gene | avg_logFC | pct.1 | pct.2 | Size (bp) | FW Primer | RV Primer |
|---|---|---|---|---|---|---|---|---|---|
| 17 | RMH | 48 | *sem-2* | 4.033578579 | 0.719 | 0.004 | 1113 | TTTTCTTGGCACCCTTTTTG | TCCACAATCCAGCATGAAGTT |

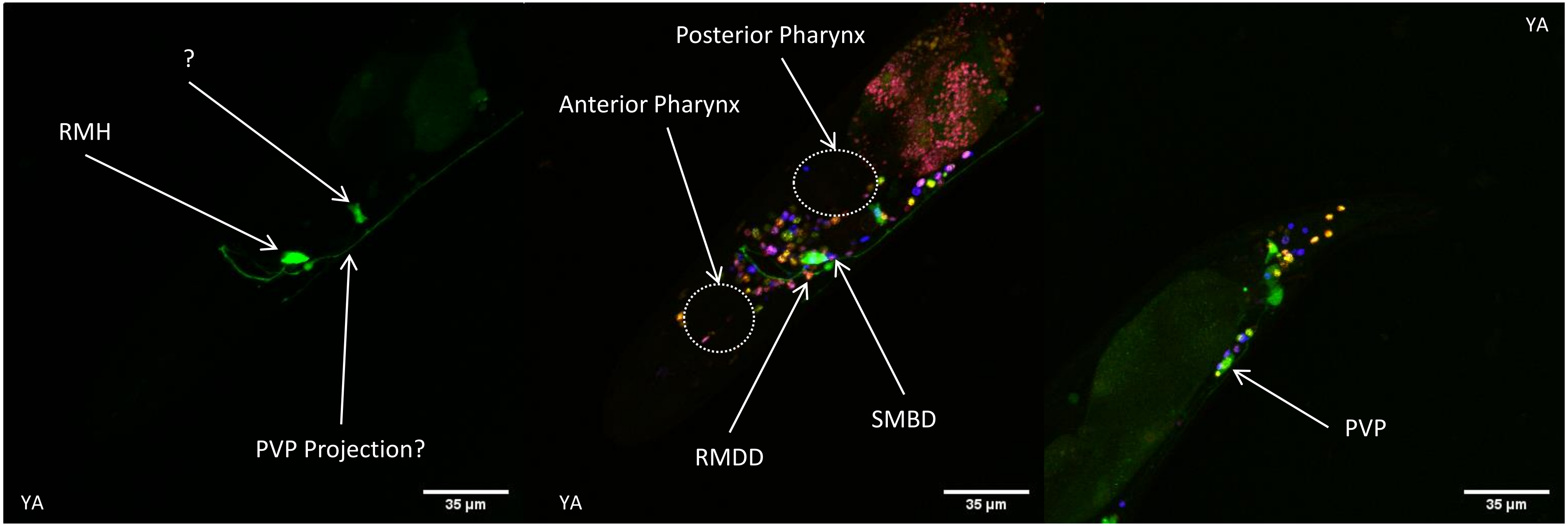


RMH (*sem-2*) – OH15262

Left lateral view of the RMH neurons: the two cell bodies are located in the ventral ganglion and close each other in the middle of the animal (the RMHR neuron is masked within the RMHL neuron signal). Ventrally from the two RMH neurons are located the RMDD (orange) and SMBD (blue) neurons. RMHL an R both send their projections anteriorly, and perform a loop around the NR. Note that an unidentified neuron is located in the anterior part of the retro-vesicular ganglion. eGFP signal is detected in 3 non-neuronal cells in the tail as well as in PVP neuron.
