## Supplementary Folder 1 for "Combining single-cell RNA-sequencing with a molecular atlas unveils new markers for *C. elegans* neuron classes"

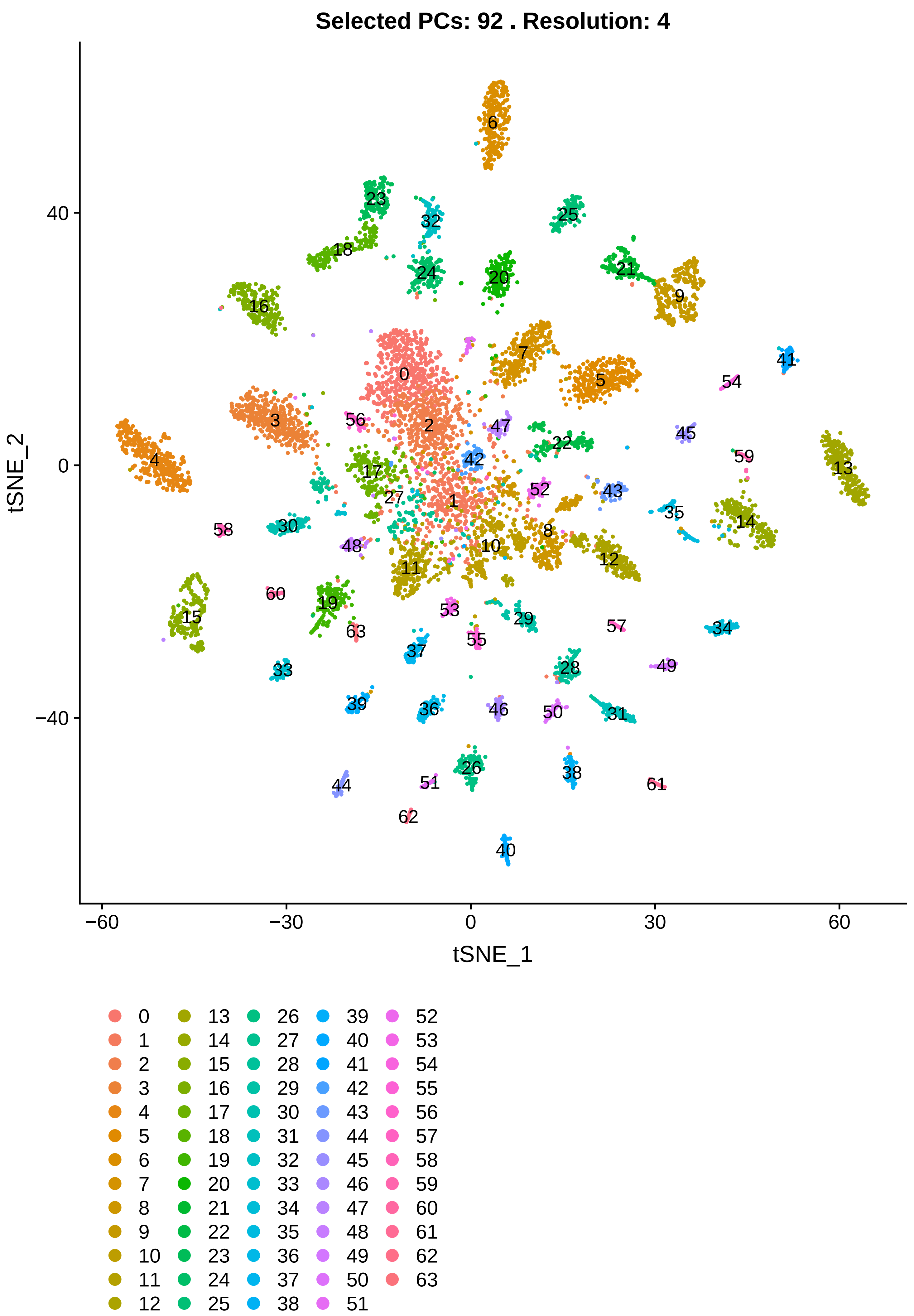
Selected PCs: 92 . Resolution: 4
tSNE_2
tSNE_1
40
0
−40
−60
−30
0
30
60
0 1 2 3 4 5 6 7 8 9 10 11 12 13 14 15 16 17 18 19 20 21 22 23 24 25 26 27 28 29 30 31 32 33 34 35 36 37 38 39 40 41 42 43 44 45 46 47 48 49 50 51 52 53 54 55 56 57 58 59 60 61 62 63

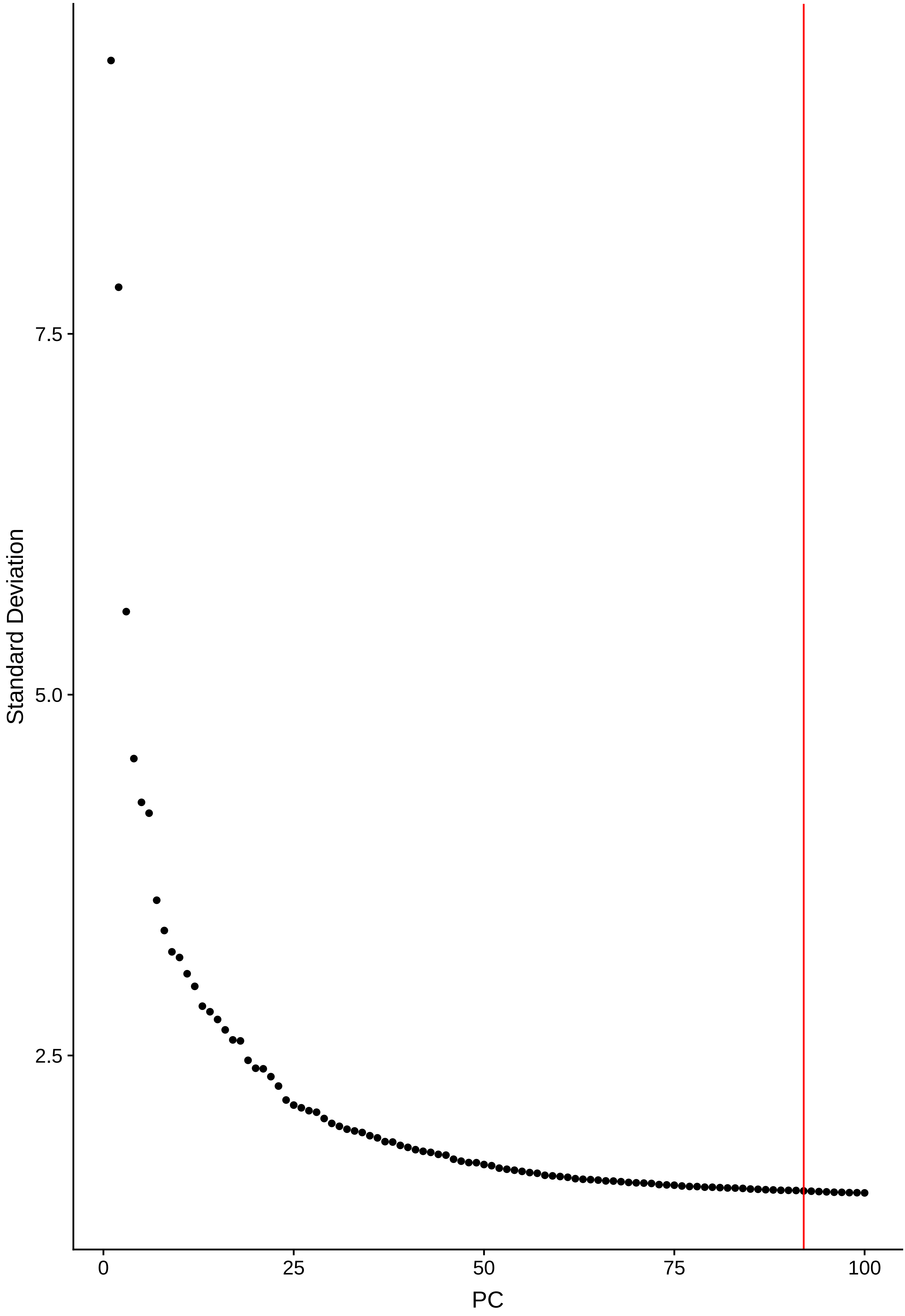

7.5
5.0
2.5
Standard Deviation
0
25
50
75
100
PC

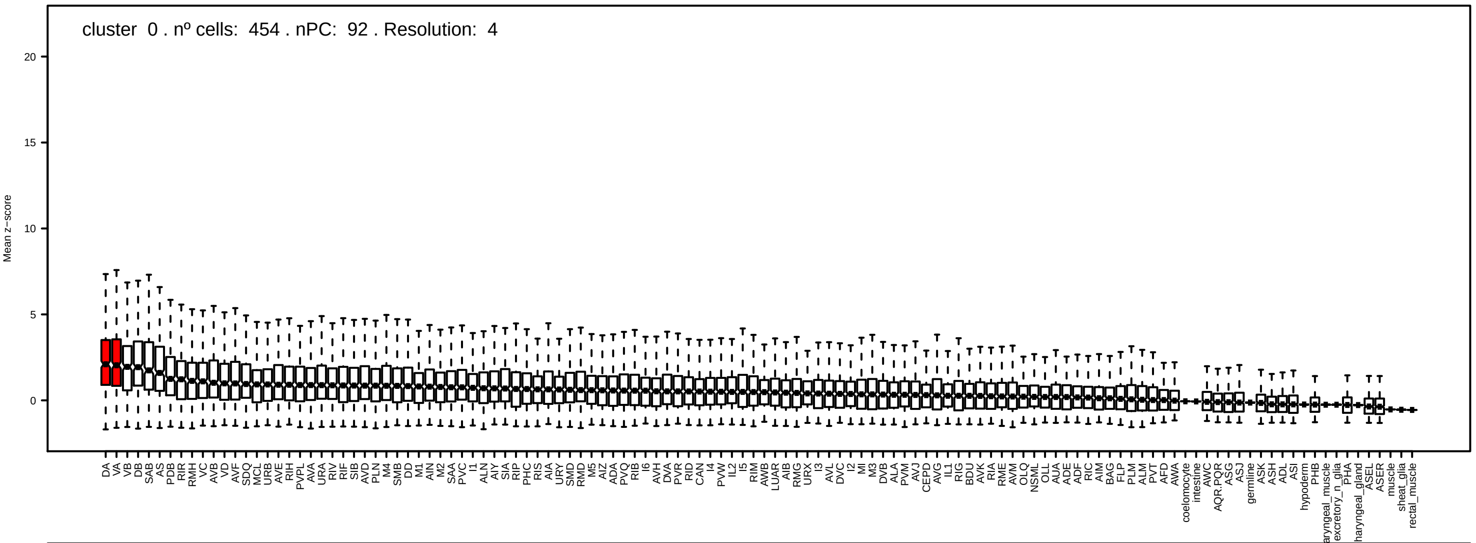
cluster 0 . nº cells: 454 . nPC: 92 . Resolution: 4
Mean z−score
20
15
10
5
0

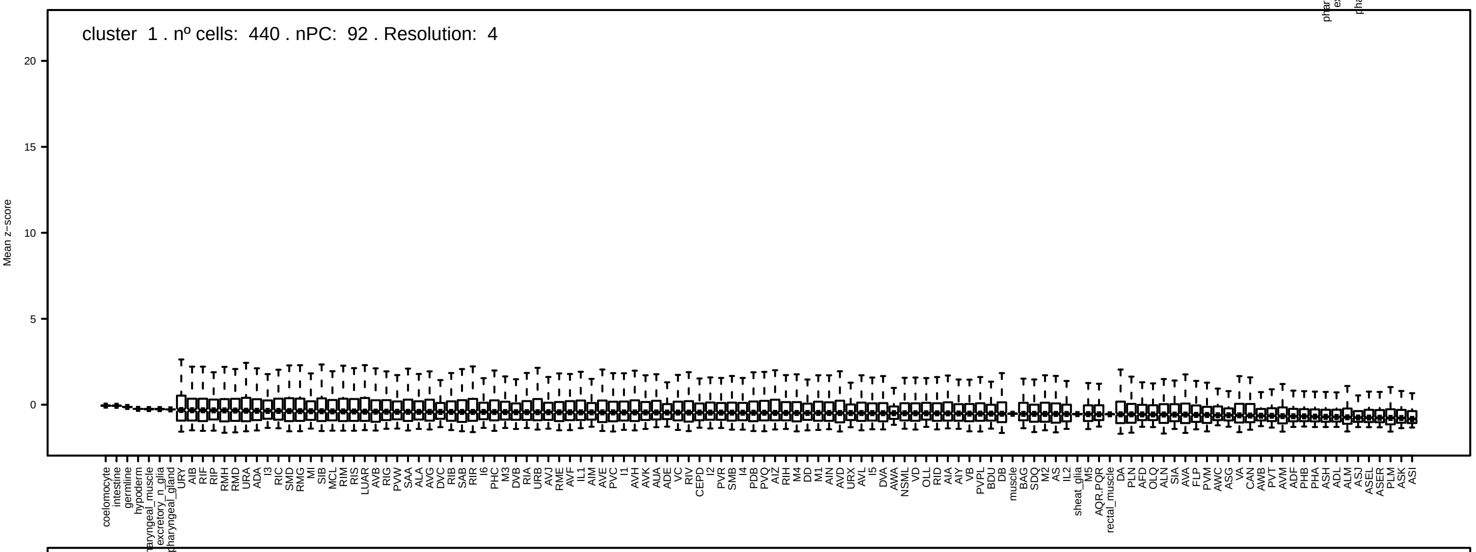
cluster 1 . nº cells: 440 . nPC: 92 . Resolution: 4
Mean z−score
20
15
10
5
0

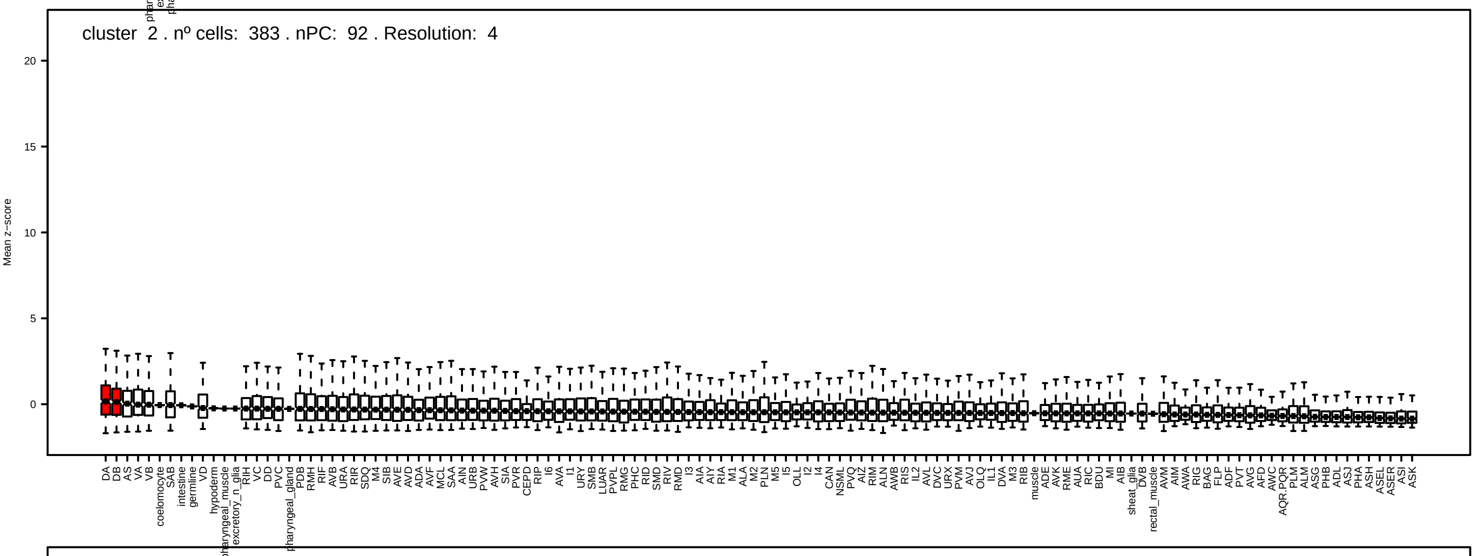
cluster 2 . nº cells: 383 . nPC: 92 . Resolution: 4
Mean z−score
20
15
10
5
0

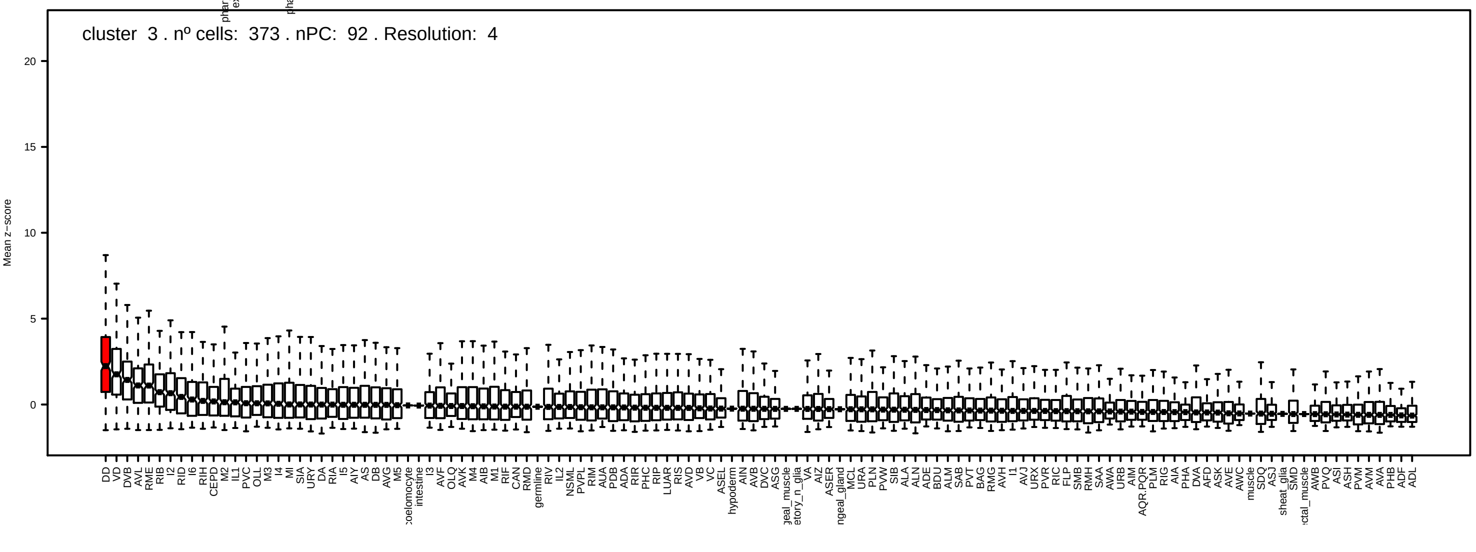
cluster 3 . nº cells: 373 . nPC: 92 . Resolution: 4
Mean z−score
20
15
10
5
0

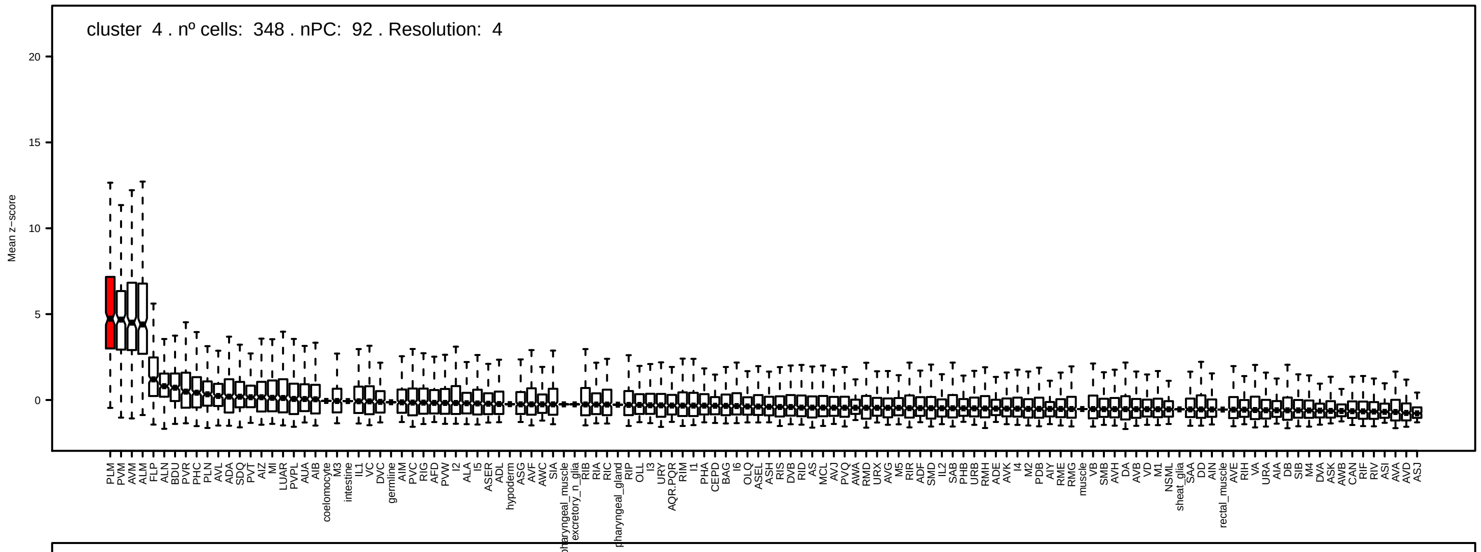
cluster 4 . nº cells: 348 . nPC: 92 . Resolution: 4
Mean z−score

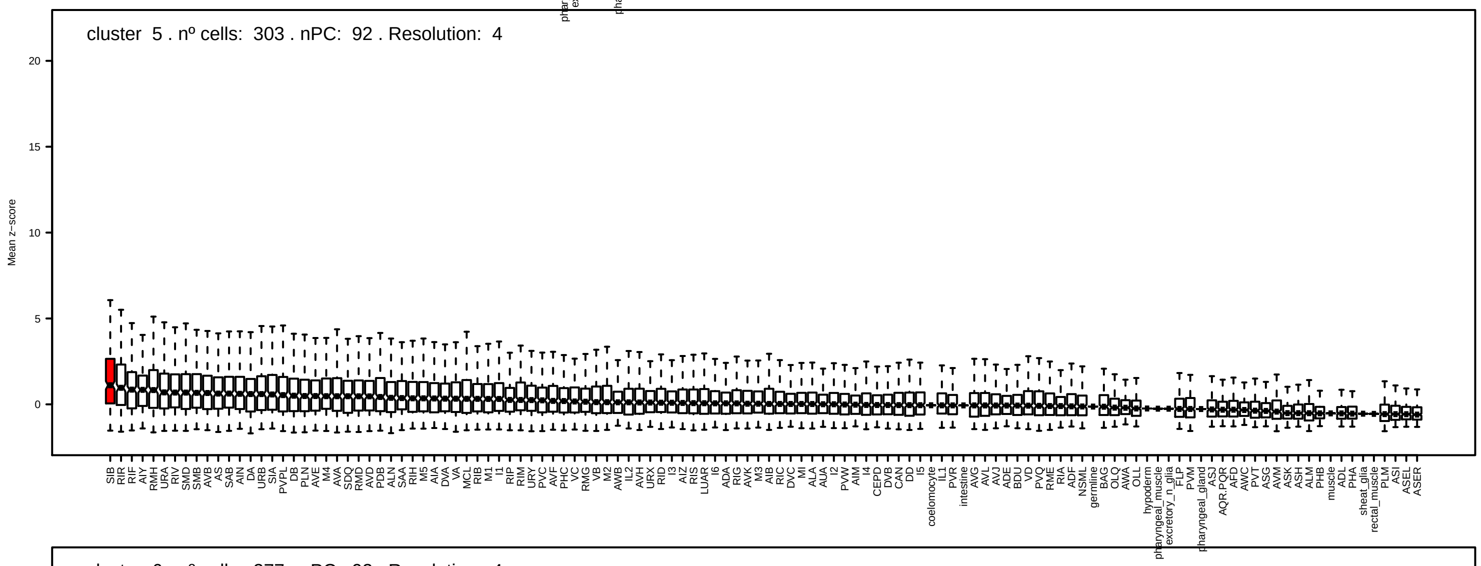
cluster 5 . nº cells: 303 . nPC: 92 . Resolution: 4
Mean z−score

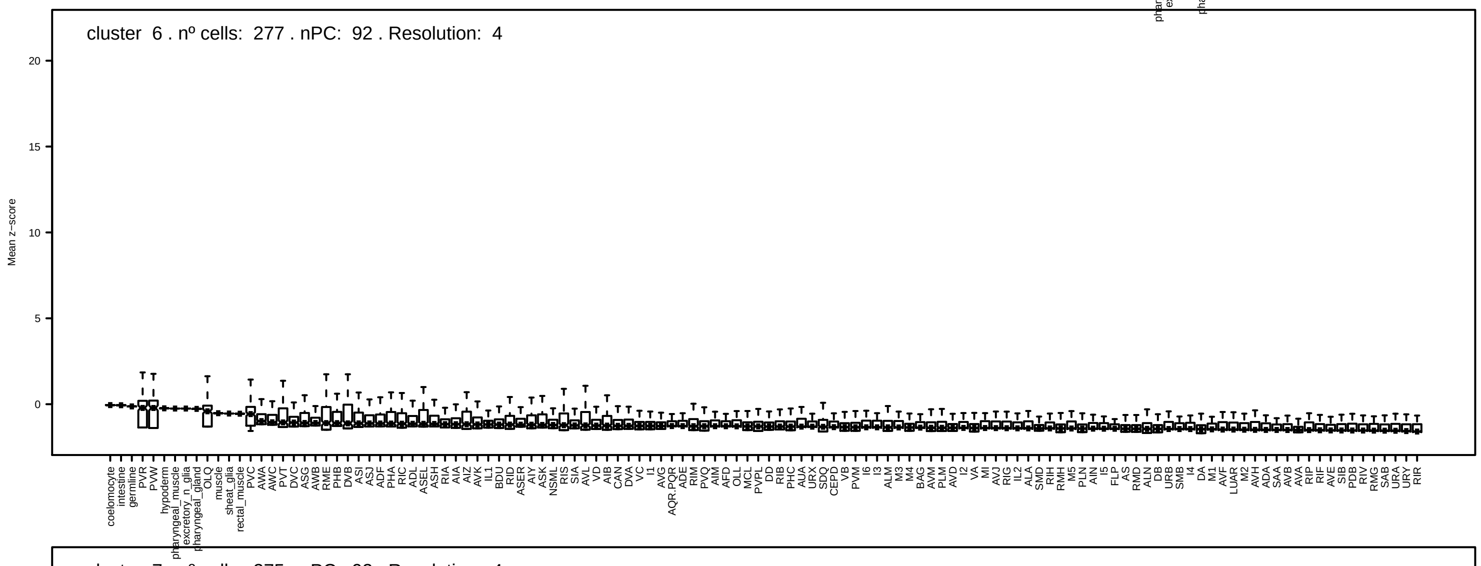
cluster 6 . nº cells: 277 . nPC: 92 . Resolution: 4
Mean z−score

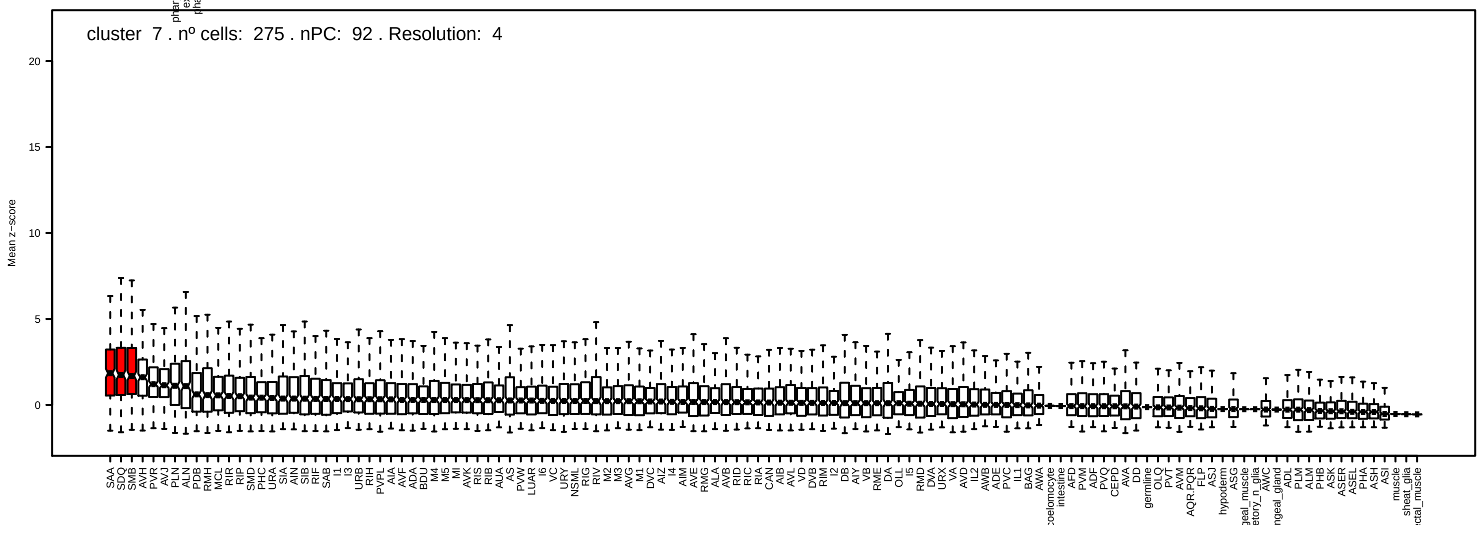
cluster 7 . nº cells: 275 . nPC: 92 . Resolution: 4
Mean z−score

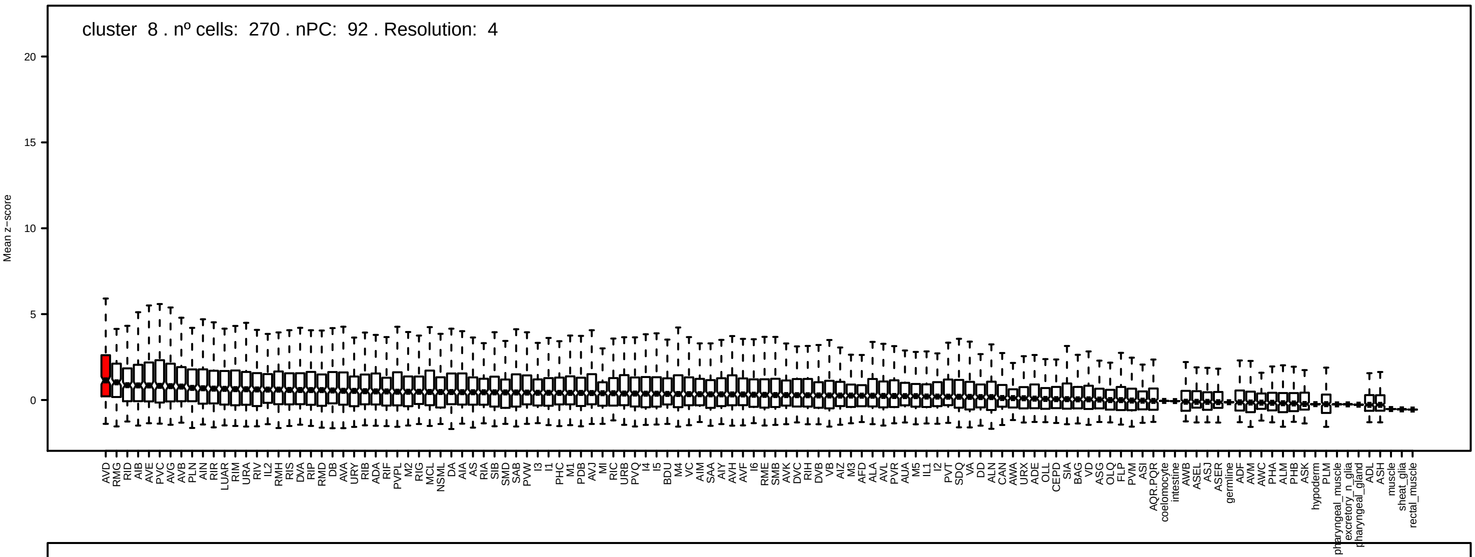
cluster 8 . nº cells: 270 . nPC: 92 . Resolution: 4
Mean z−score

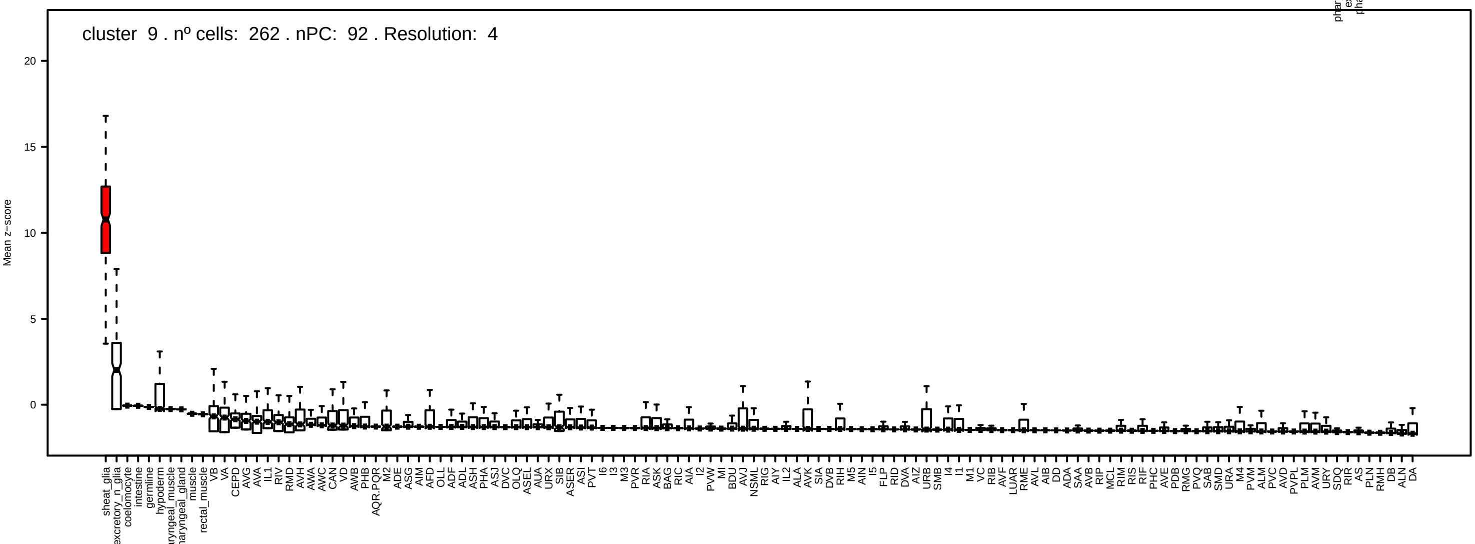
cluster 9 . nº cells: 262 . nPC: 92 . Resolution: 4
Mean z−score

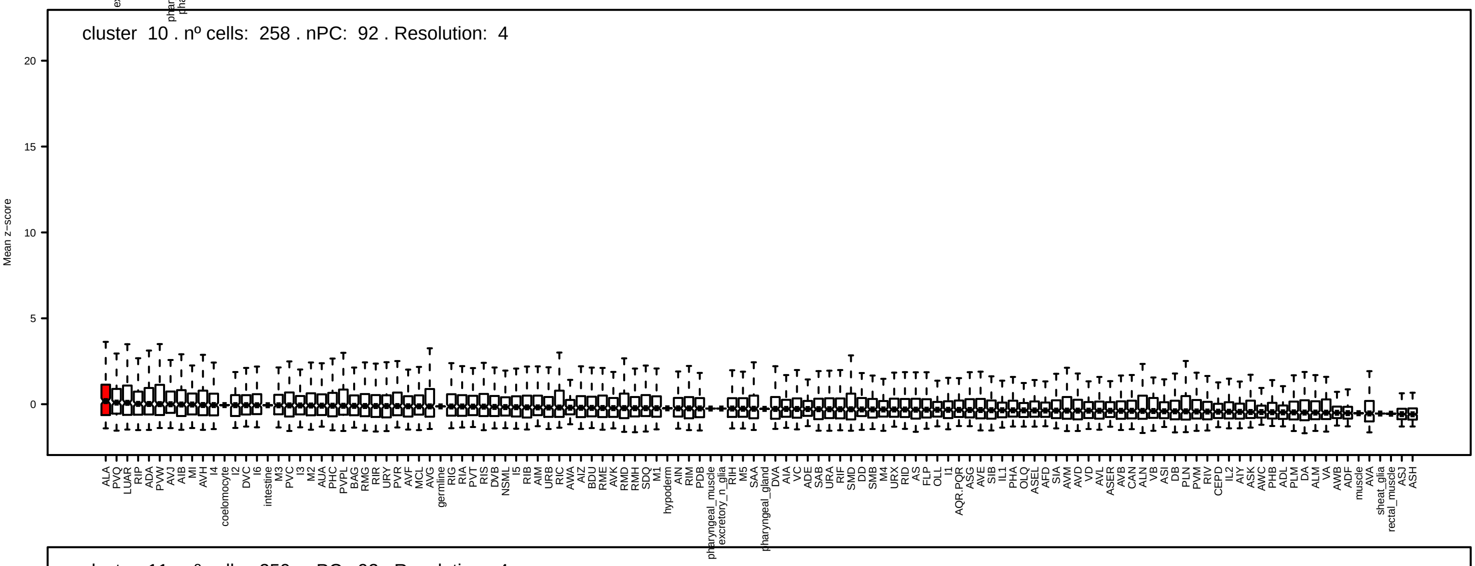
cluster 10 . nº cells: 258 . nPC: 92 . Resolution: 4
Mean z−score

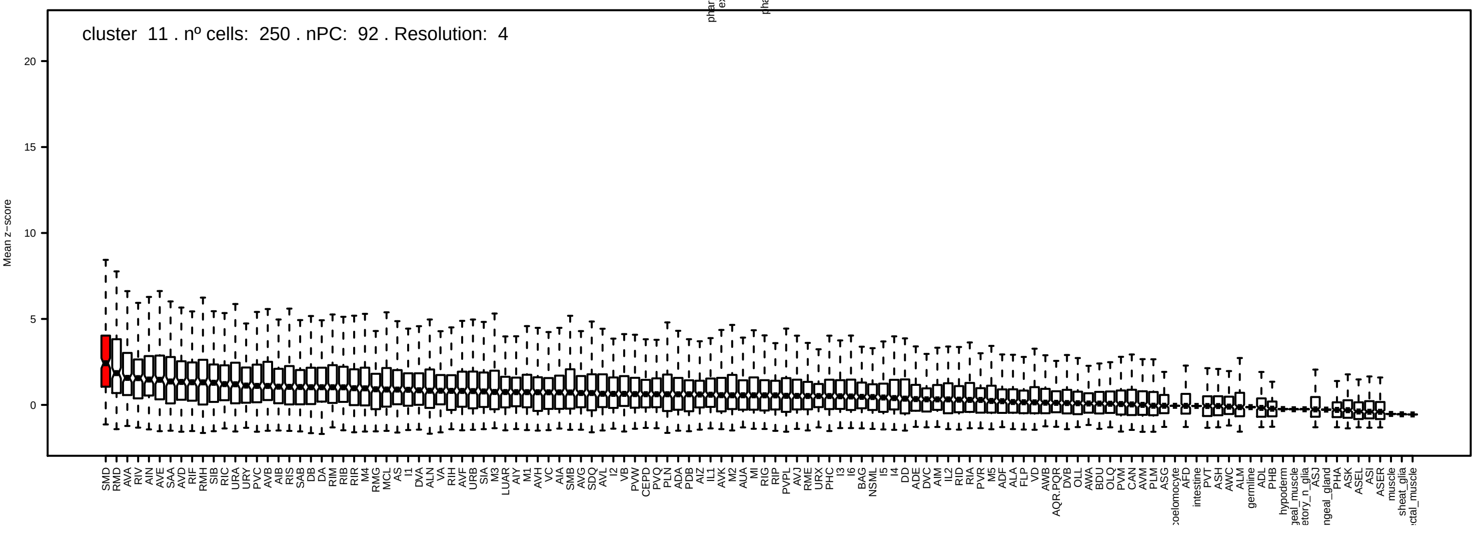
cluster 11 . nº cells: 250 . nPC: 92 . Resolution: 4
Mean z−score

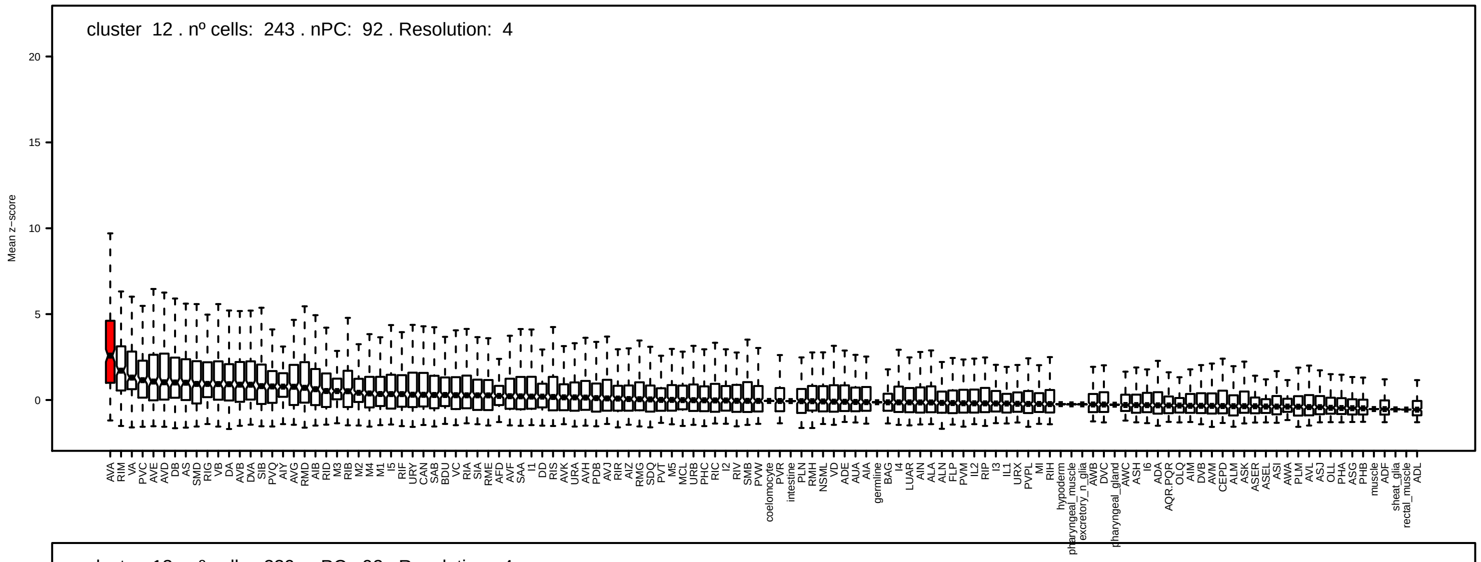
cluster 12 . nº cells: 243 . nPC: 92 . Resolution: 4
Mean z−score

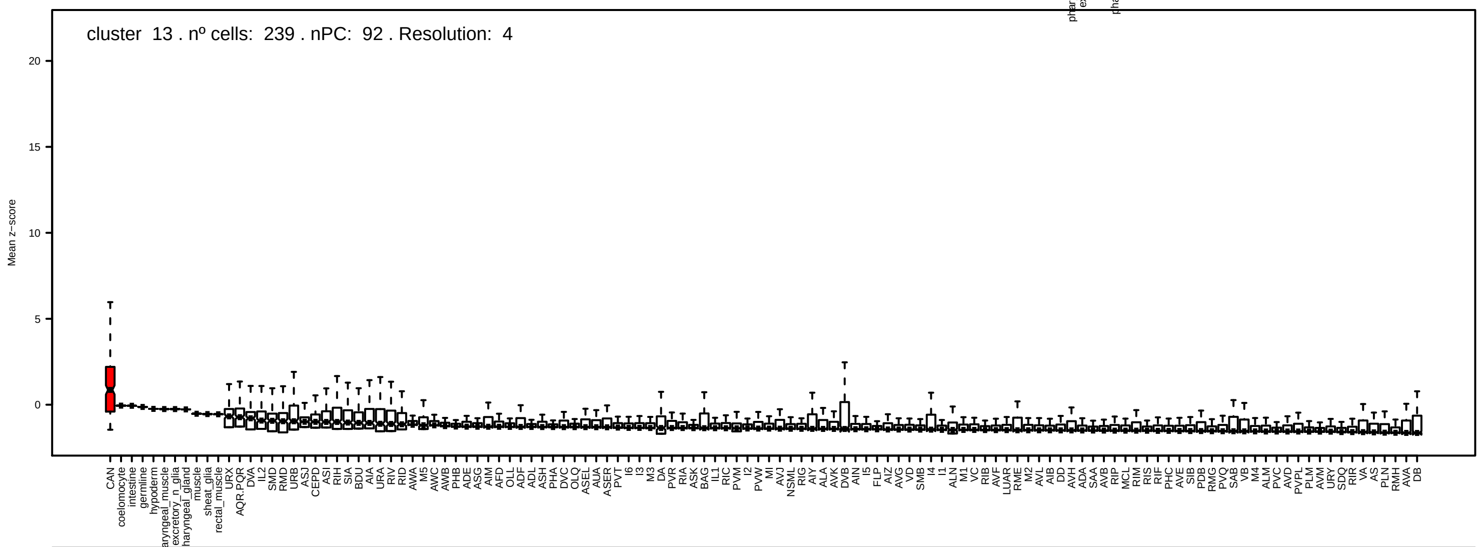
cluster 13 . nº cells: 239 . nPC: 92 . Resolution: 4
Mean z−score

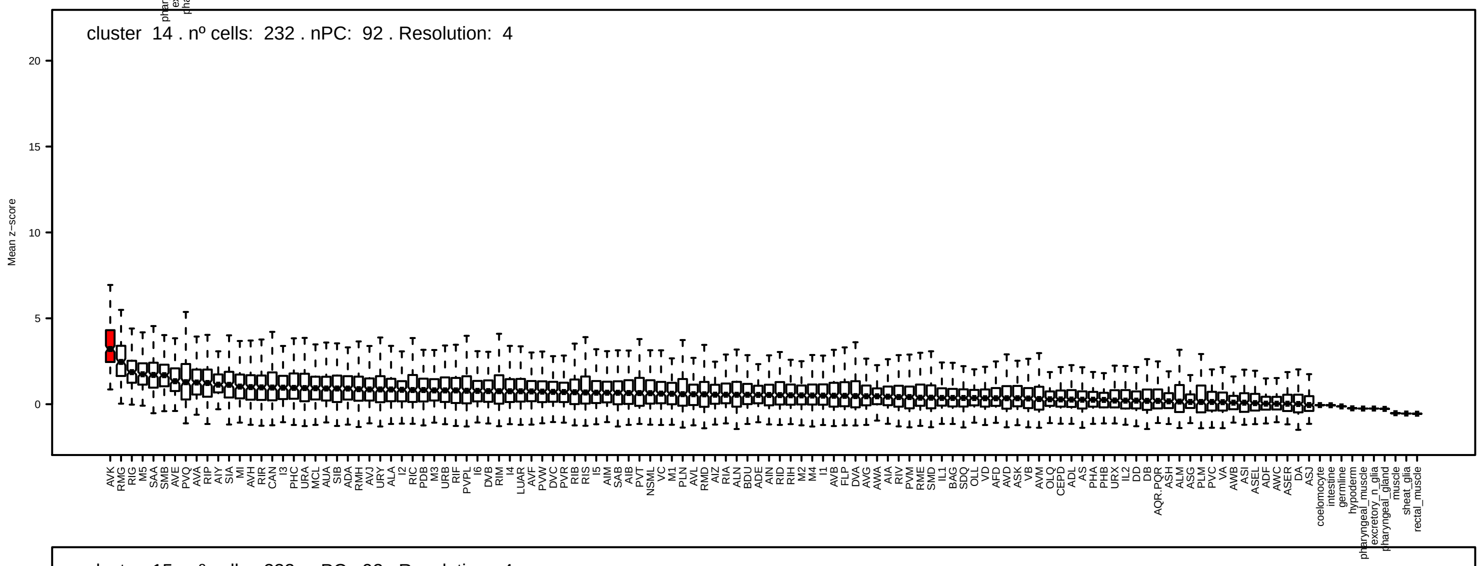
cluster 14 . nº cells: 232 . nPC: 92 . Resolution: 4
Mean z−score

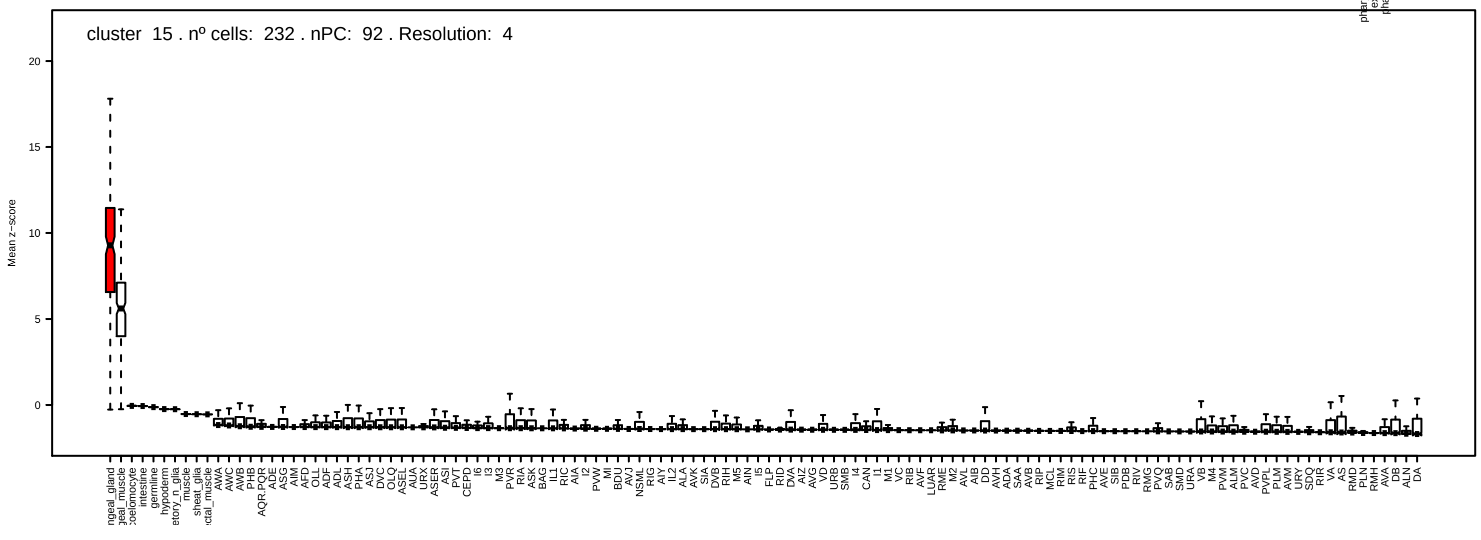
cluster 15 . nº cells: 232 . nPC: 92 . Resolution: 4
Mean z−score

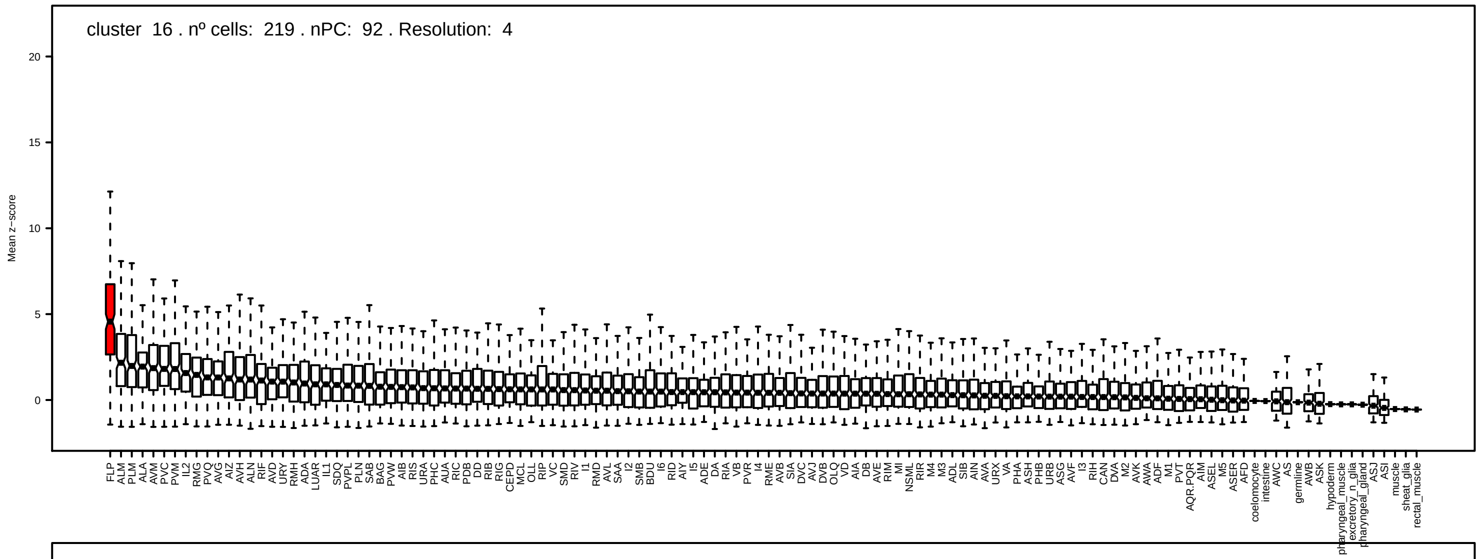
cluster 16 . nº cells: 219 . nPC: 92 . Resolution: 4
Mean z−score

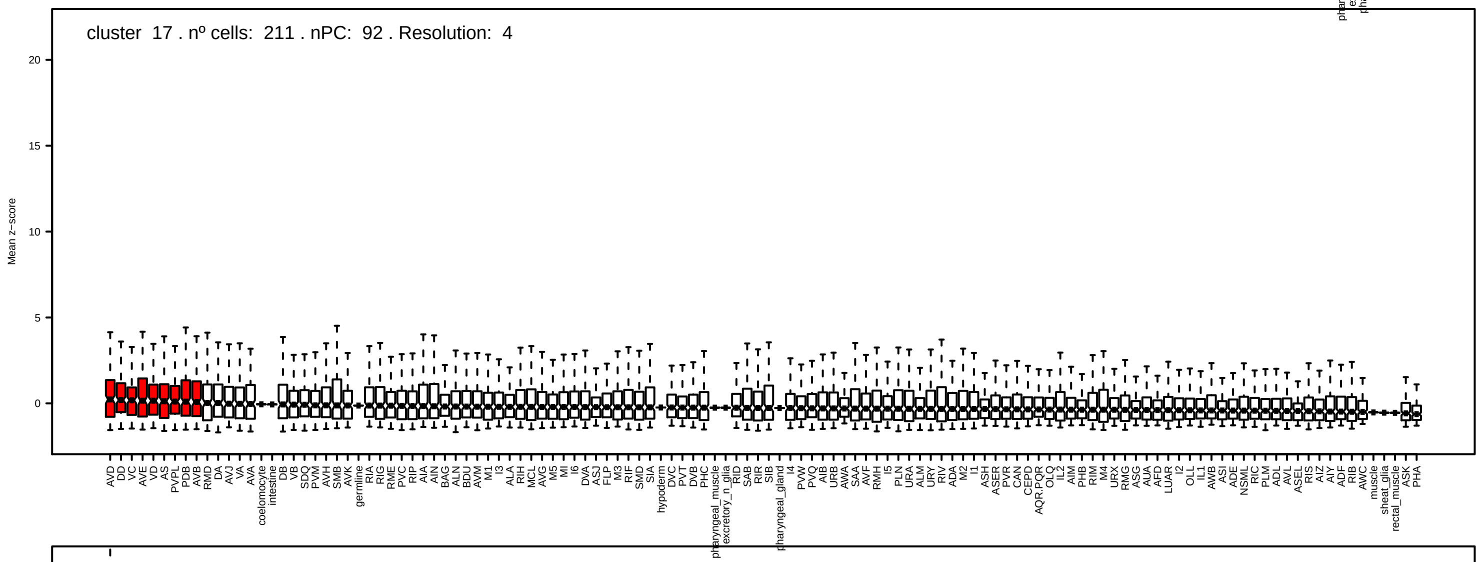
cluster 17 . nº cells: 211 . nPC: 92 . Resolution: 4
Mean z−score

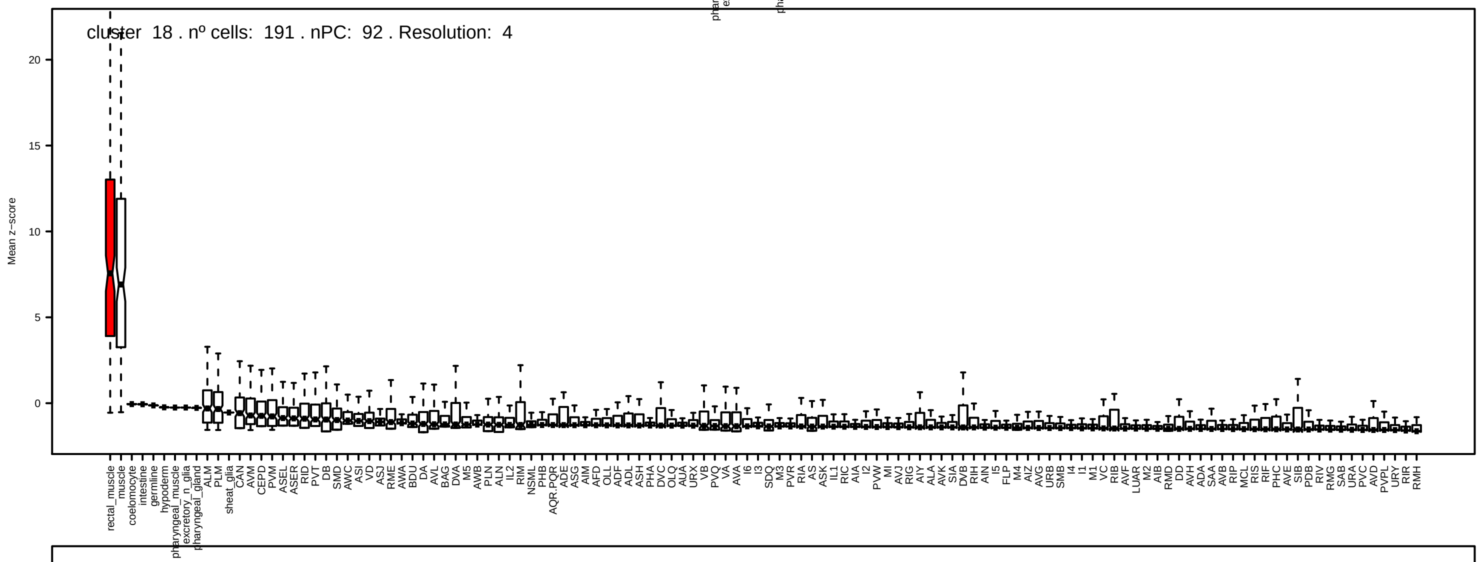
cluster 18 . nº cells: 191 . nPC: 92 . Resolution: 4
Mean z−score

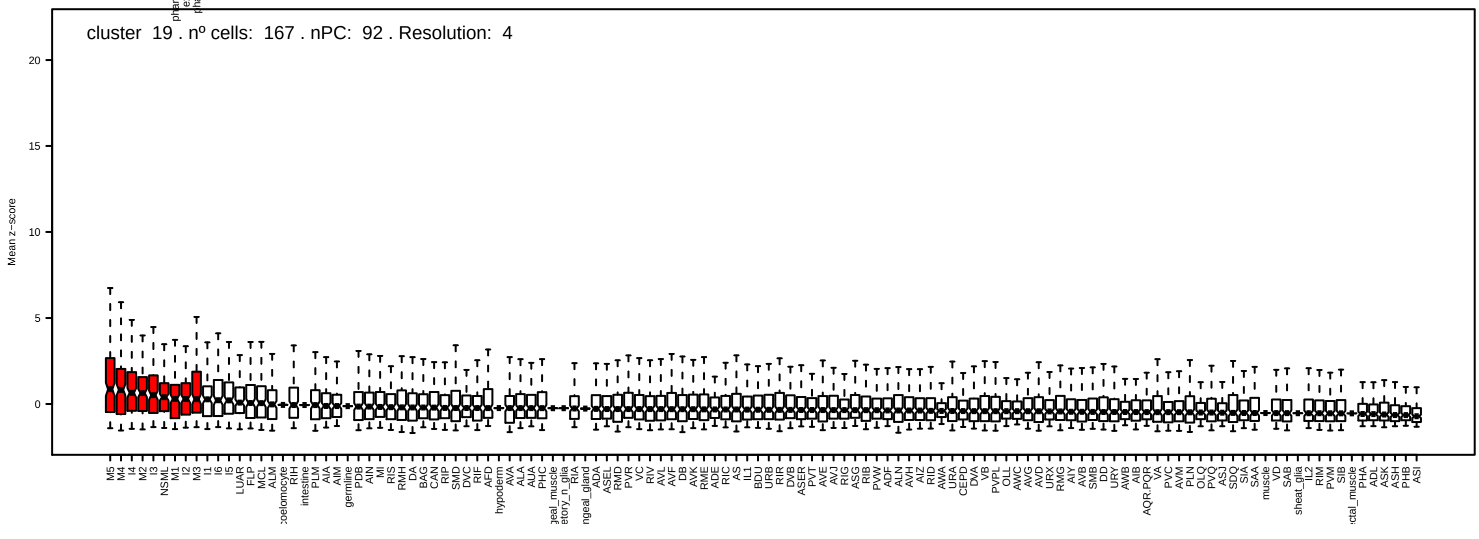
cluster 19 . nº cells: 167 . nPC: 92 . Resolution: 4
Mean z−score

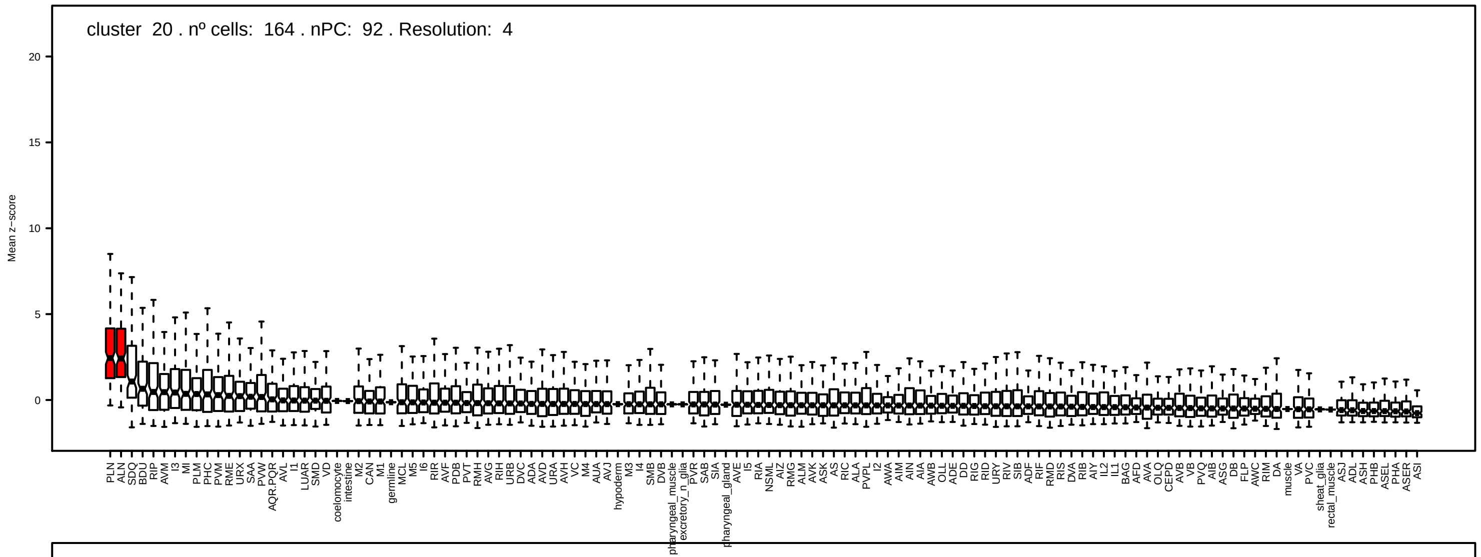
cluster 20 . nº cells: 164 . nPC: 92 . Resolution: 4
Mean z−score
20
15
10
5
0

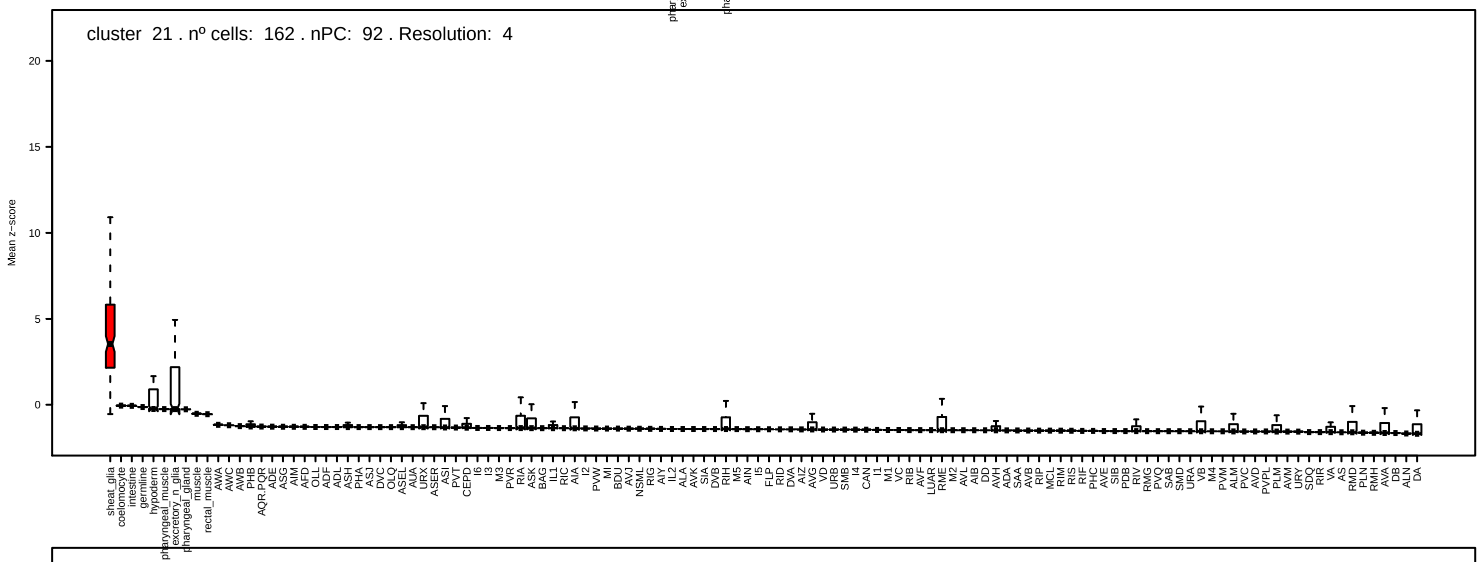
cluster 21 . nº cells: 162 . nPC: 92 . Resolution: 4
Mean z−score
20
15
10
5
0

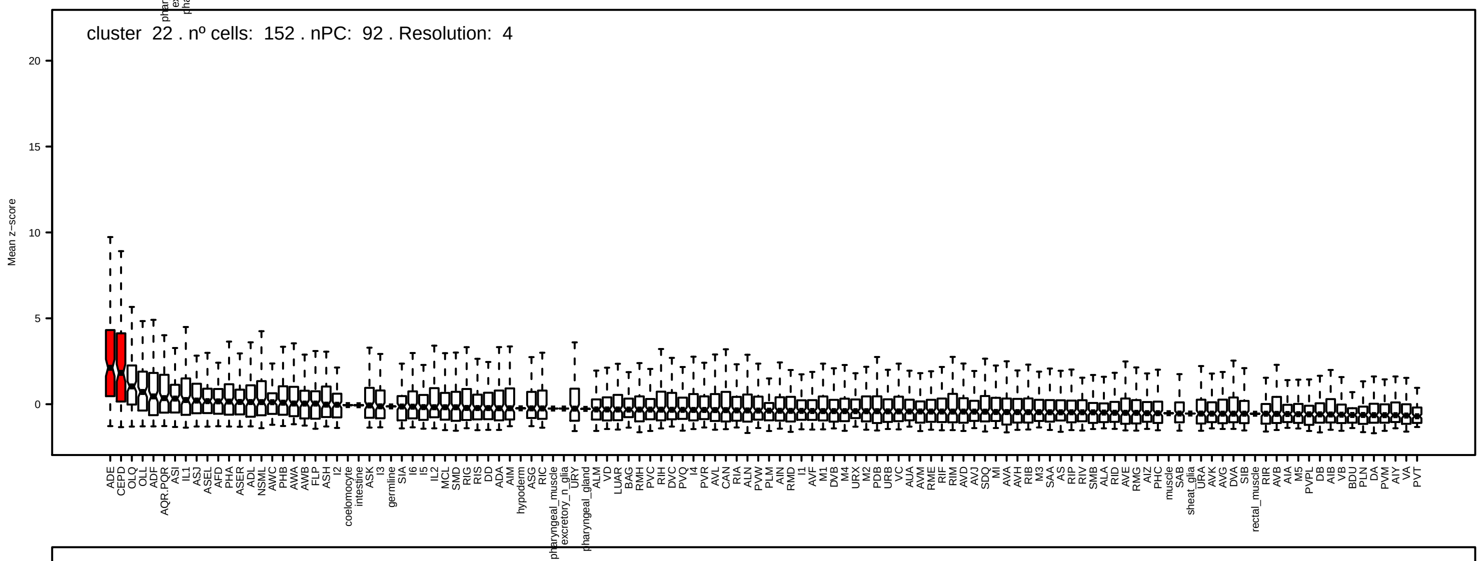
cluster 22 . nº cells: 152 . nPC: 92 . Resolution: 4
Mean z−score
20
15
10
5
0

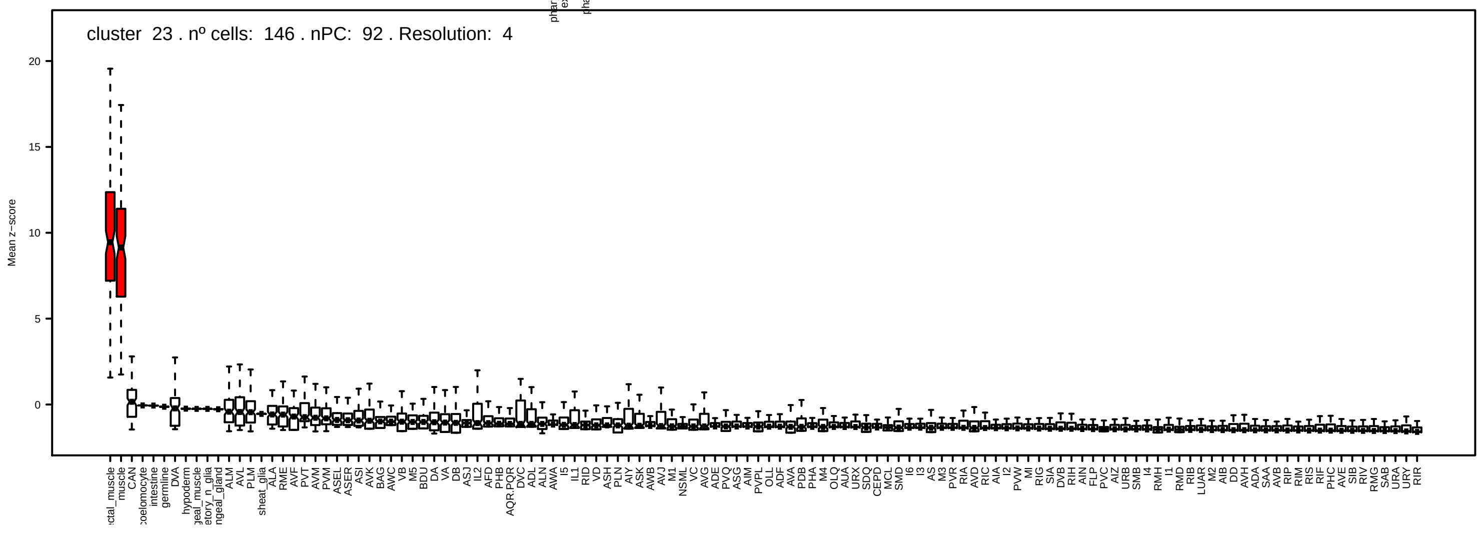
cluster 23 . nº cells: 146 . nPC: 92 . Resolution: 4
Mean z−score
20
15
10
5
0

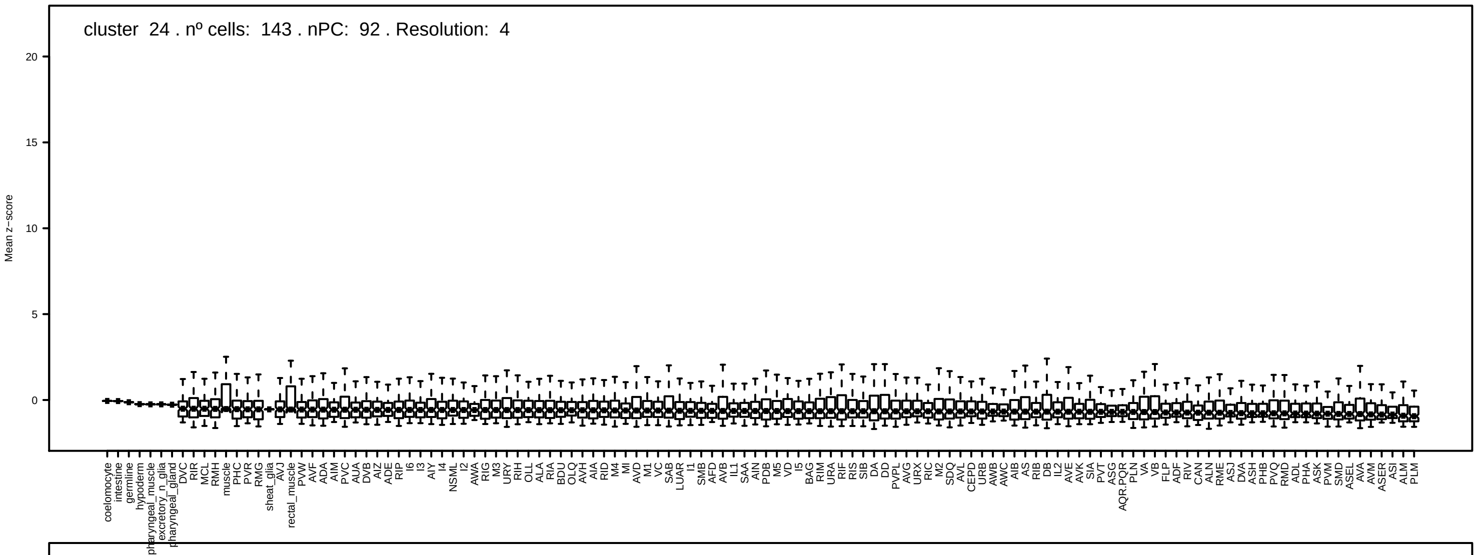
cluster 24 . nº cells: 143 . nPC: 92 . Resolution: 4
Mean z−score

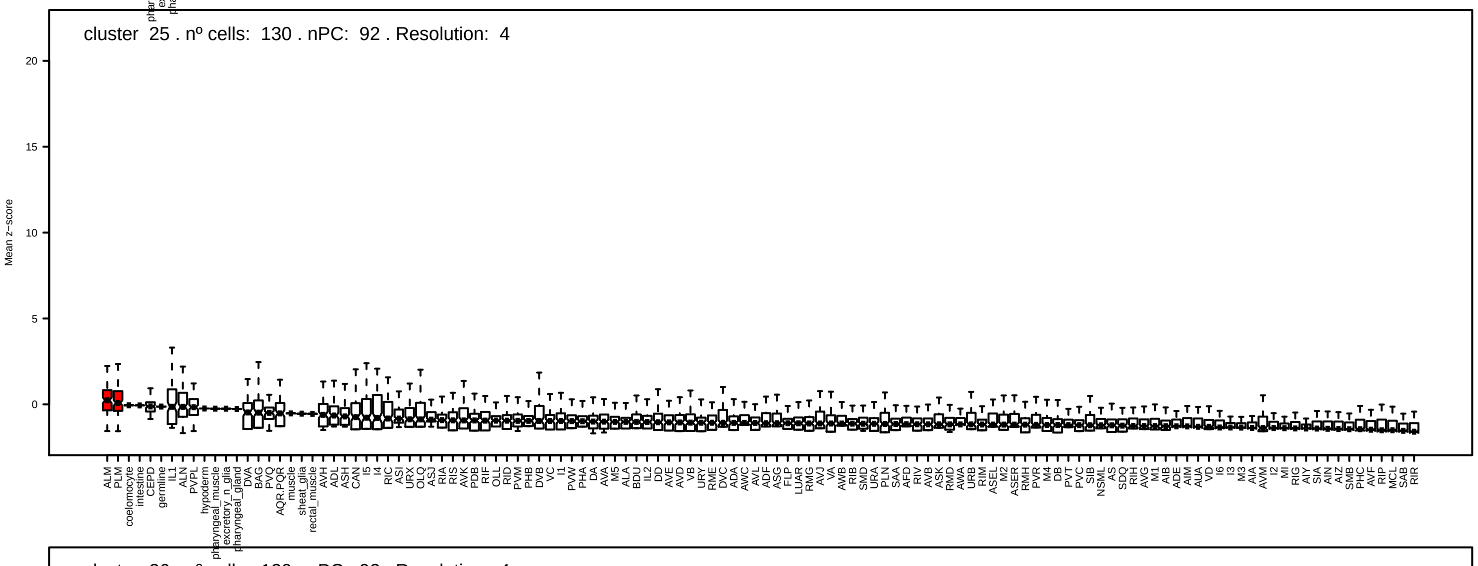
cluster 25 . nº cells: 130 . nPC: 92 . Resolution: 4
Mean z−score

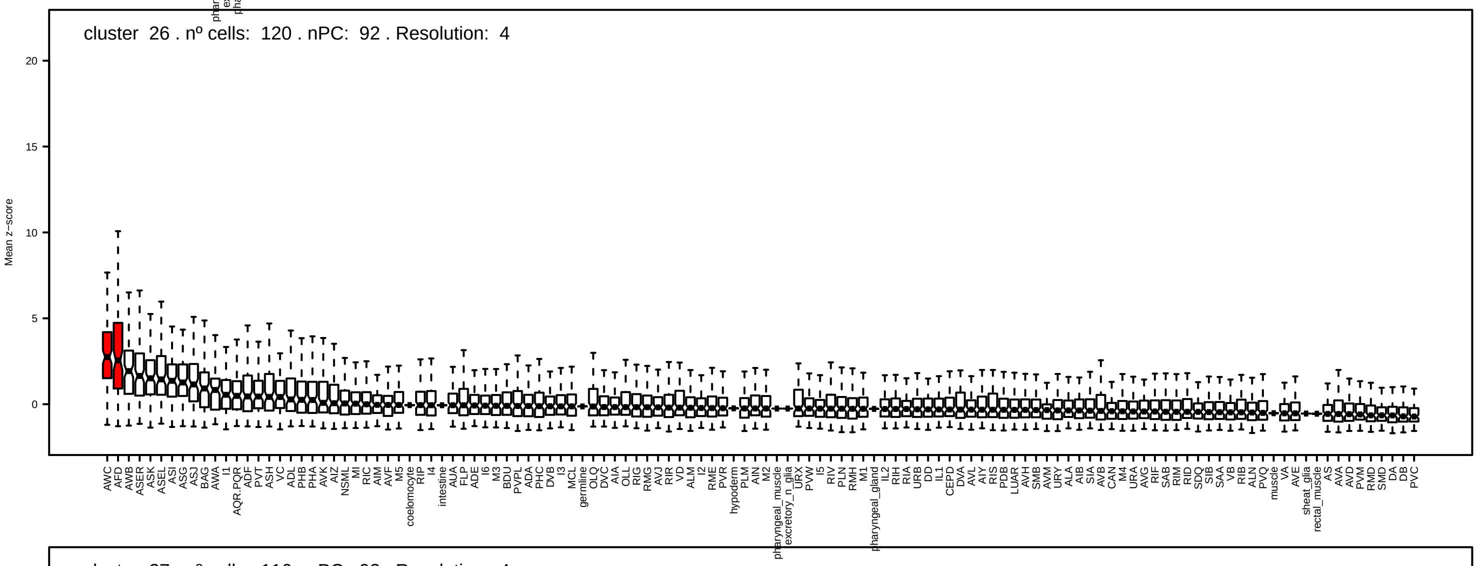
cluster 26 . nº cells: 120 . nPC: 92 . Resolution: 4
Mean z−score

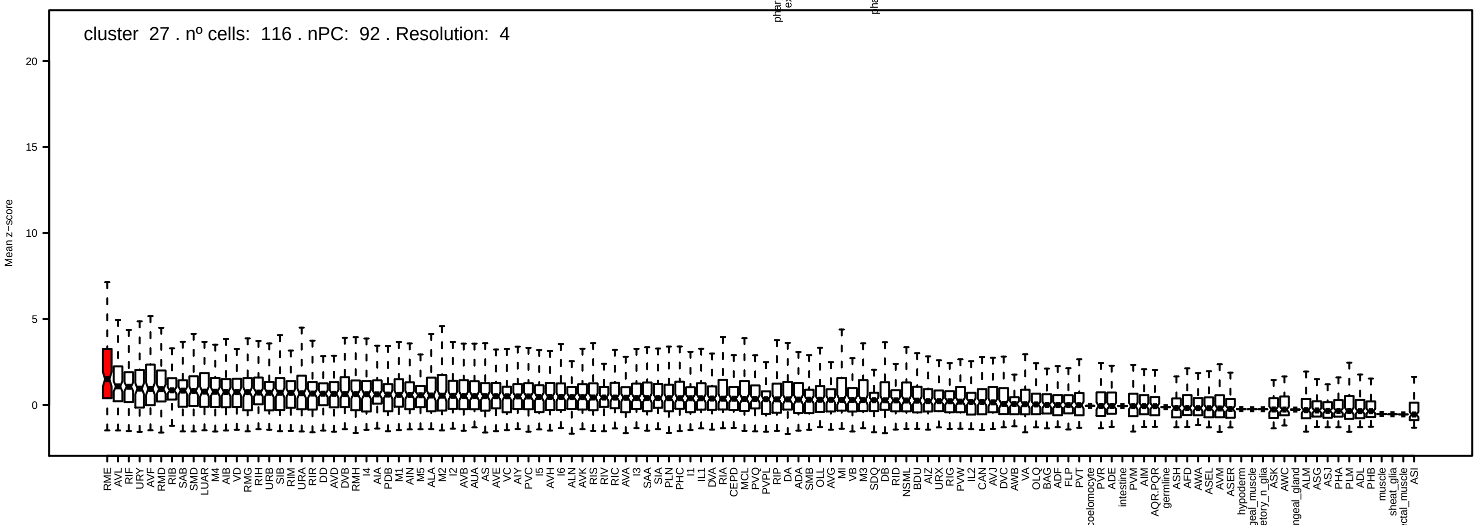
cluster 27 . nº cells: 116 . nPC: 92 . Resolution: 4
Mean z−score

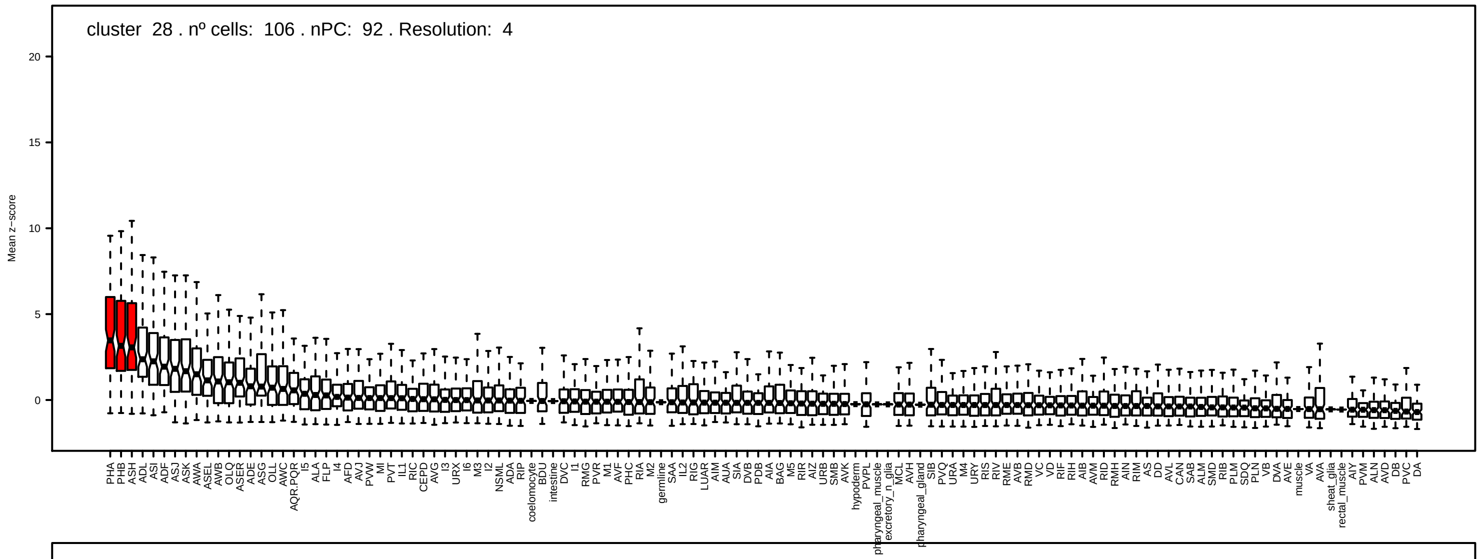
cluster 28 . nº cells: 106 . nPC: 92 . Resolution: 4
Mean z−score

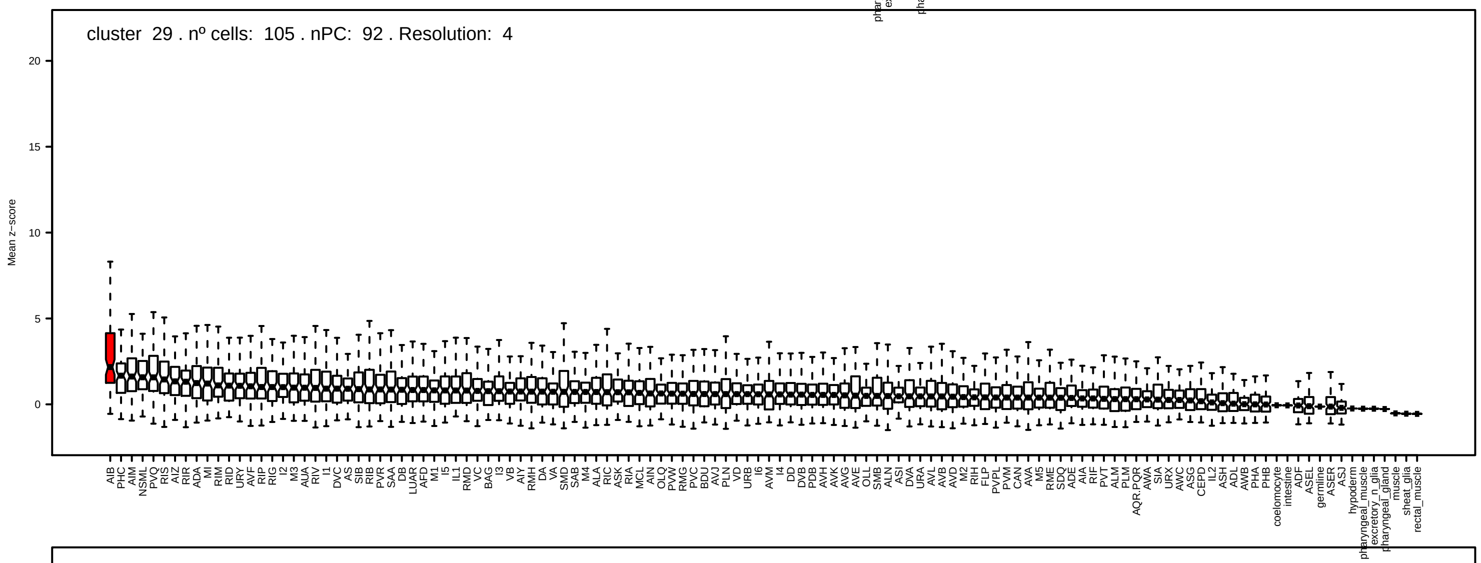
cluster 29 . nº cells: 105 . nPC: 92 . Resolution: 4
Mean z−score

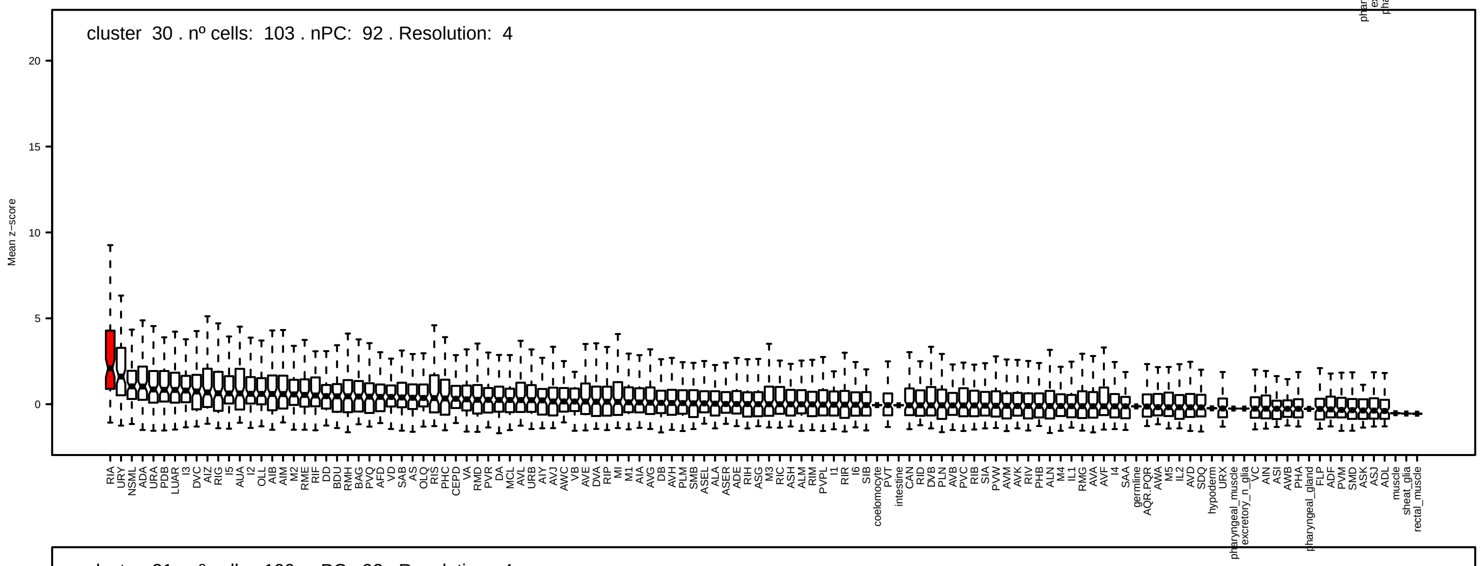
cluster 30 . nº cells: 103 . nPC: 92 . Resolution: 4
Mean z−score

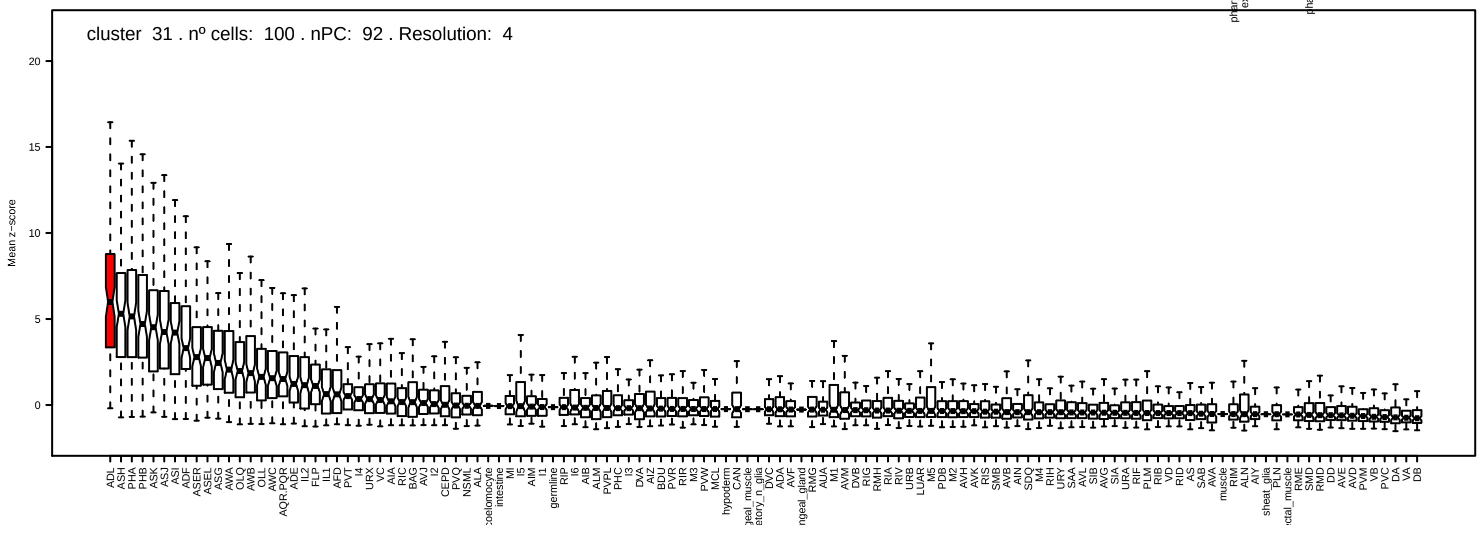
cluster 31 . nº cells: 100 . nPC: 92 . Resolution: 4
Mean z−score

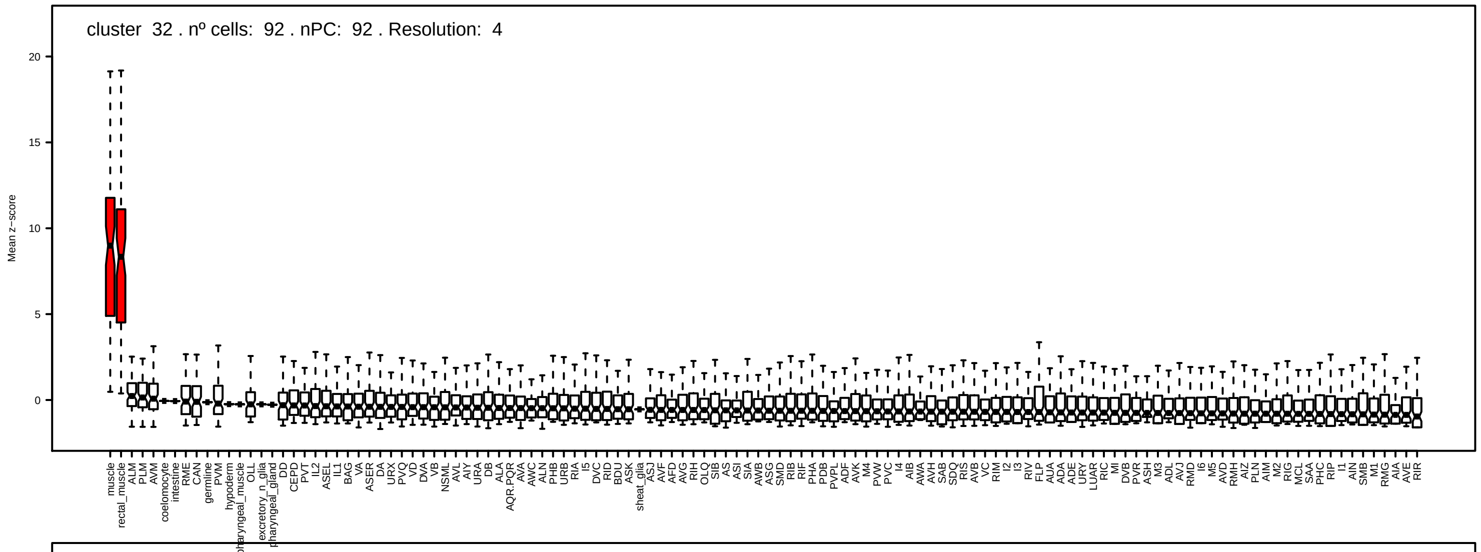
cluster 32 . nº cells: 92 . nPC: 92 . Resolution: 4
Mean z−score

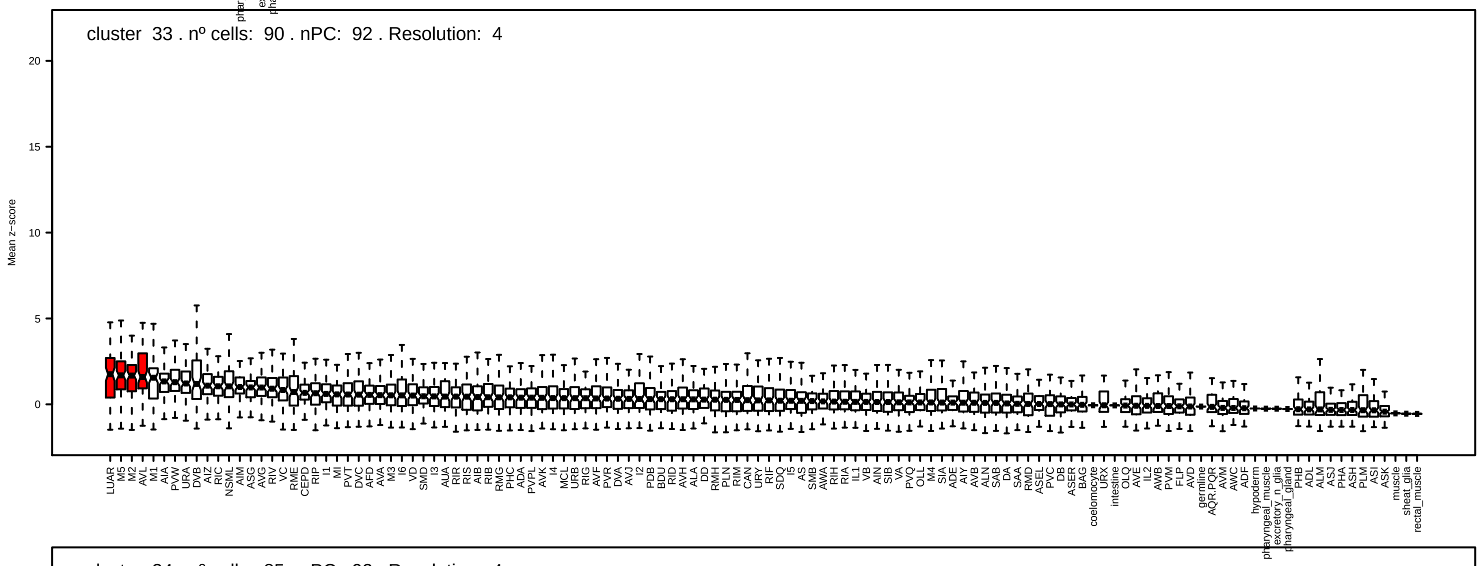
cluster 33 . nº cells: 90 . nPC: 92 . Resolution: 4
Mean z−score

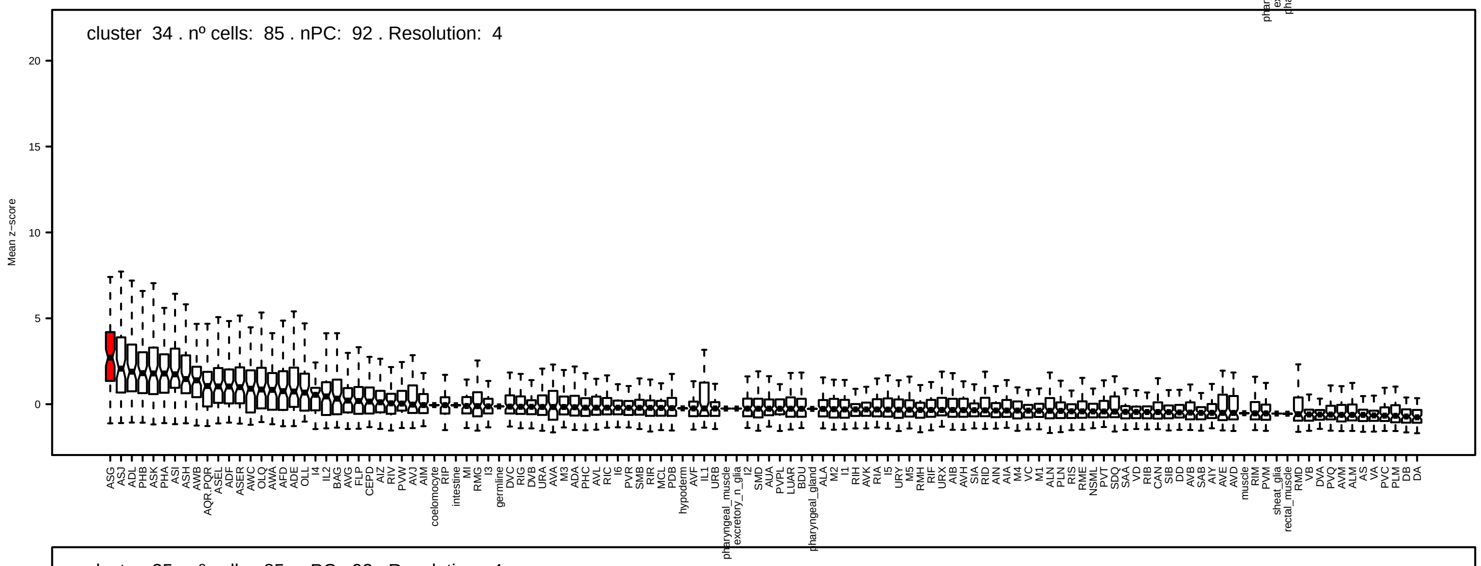
cluster 34 . nº cells: 85 . nPC: 92 . Resolution: 4
Mean z−score

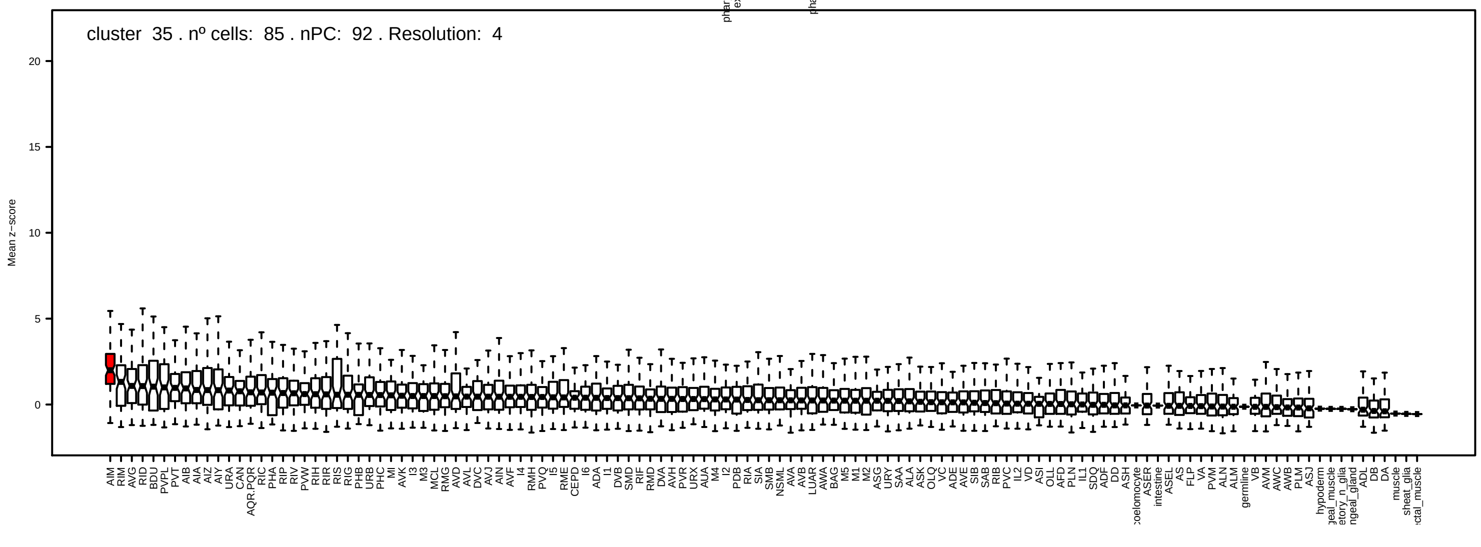
cluster 35 . nº cells: 85 . nPC: 92 . Resolution: 4
Mean z−score

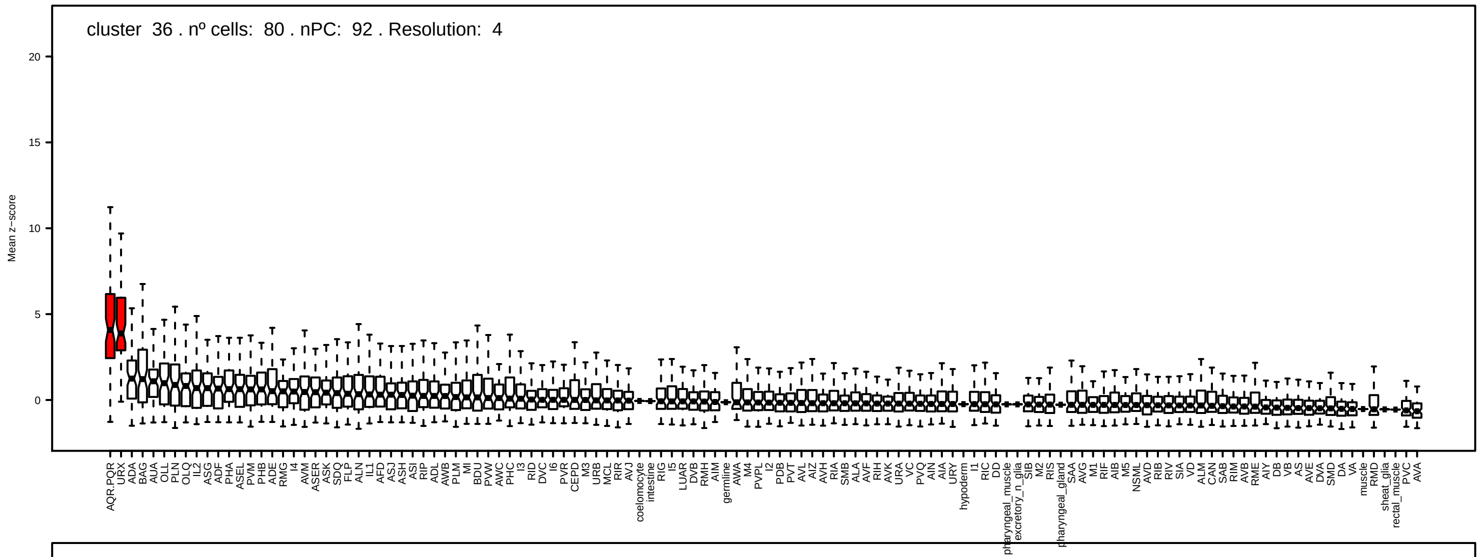
cluster 36 . nº cells: 80 . nPC: 92 . Resolution: 4
Mean z−score
20
15
10
5
0

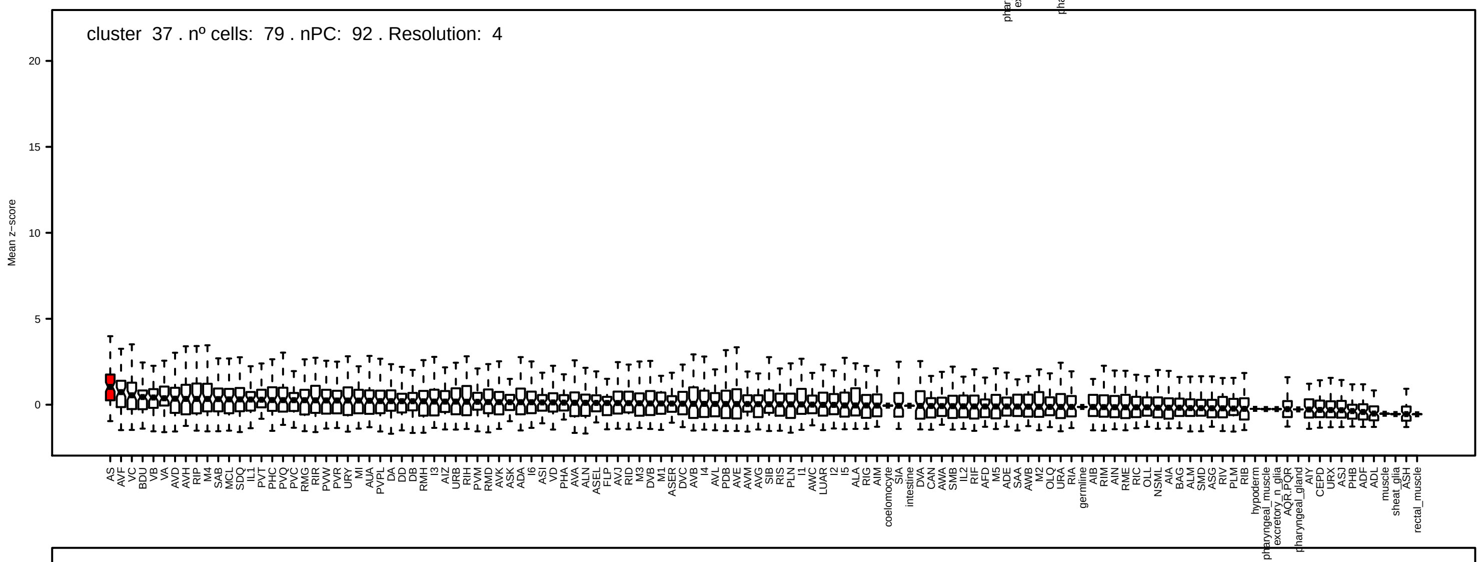
cluster 37 . nº cells: 79 . nPC: 92 . Resolution: 4
Mean z−score
20
15
10
5
0

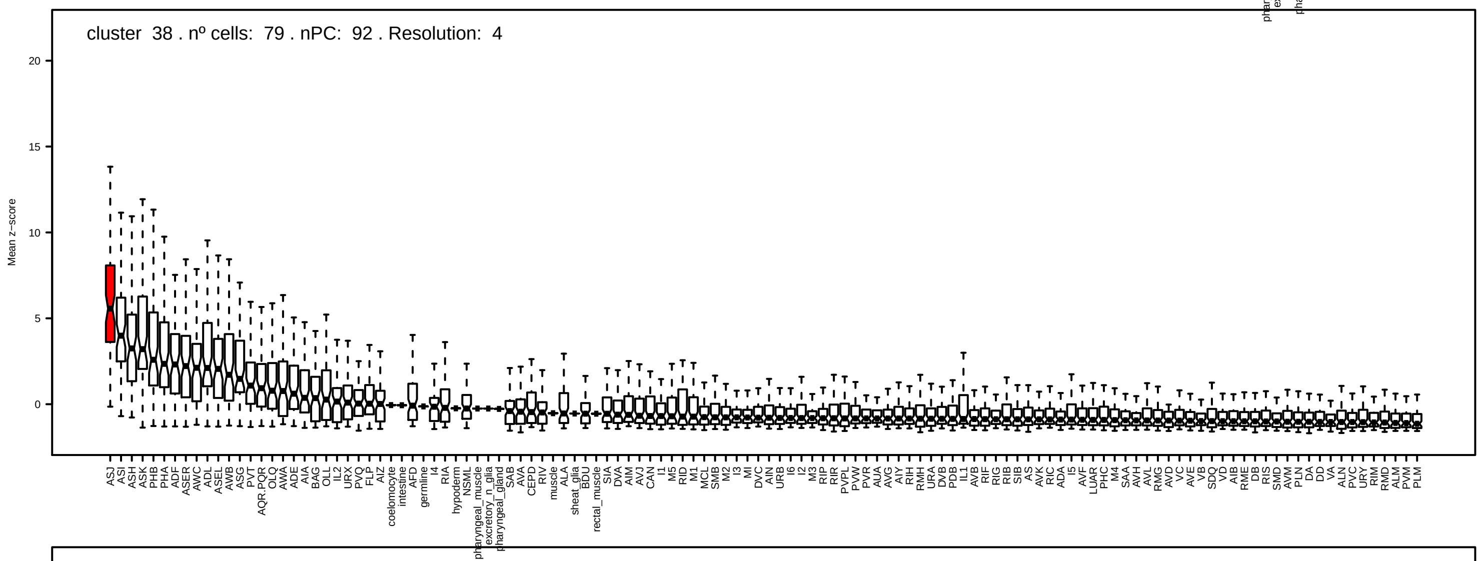
cluster 38 . nº cells: 79 . nPC: 92 . Resolution: 4
Mean z−score
20
15
10
5
0

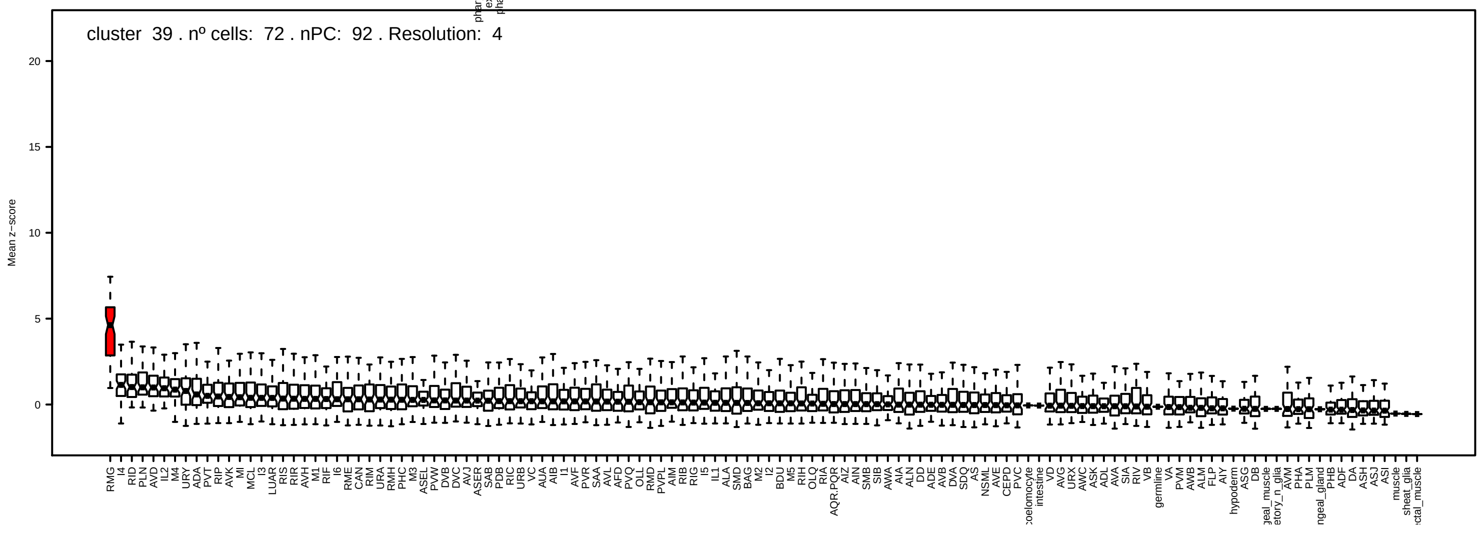
cluster 39 . nº cells: 72 . nPC: 92 . Resolution: 4
Mean z−score
20
15
10
5
0

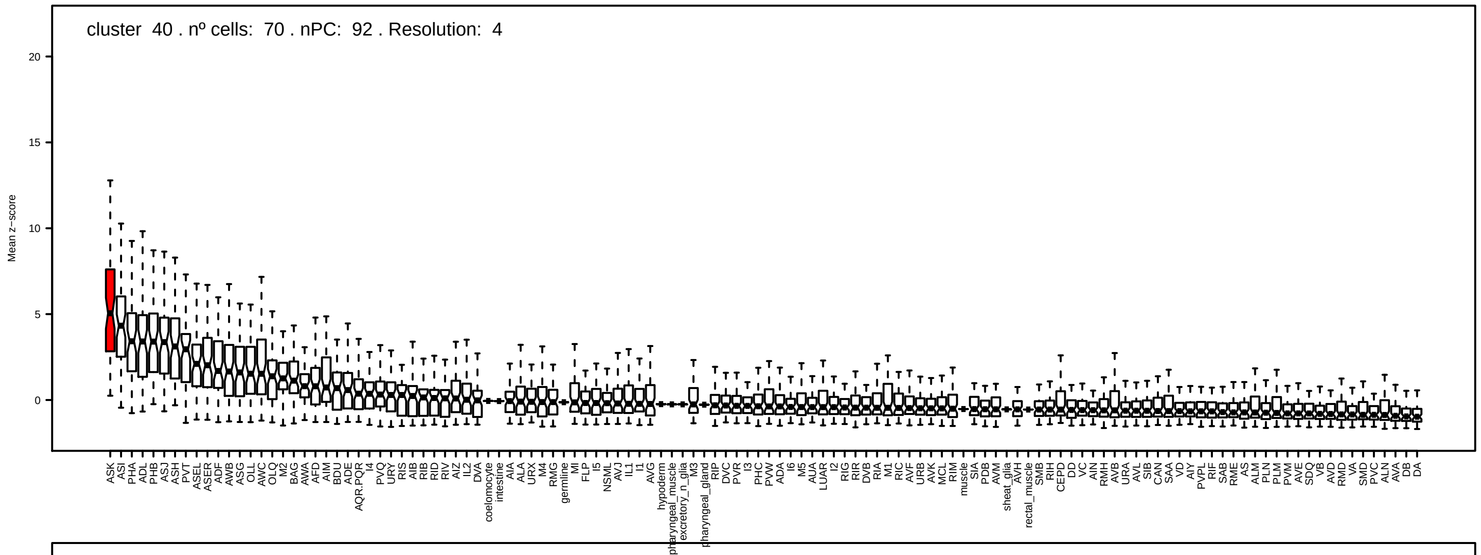
cluster 40 . nº cells: 70 . nPC: 92 . Resolution: 4
Mean z−score
20
15
10
5
0

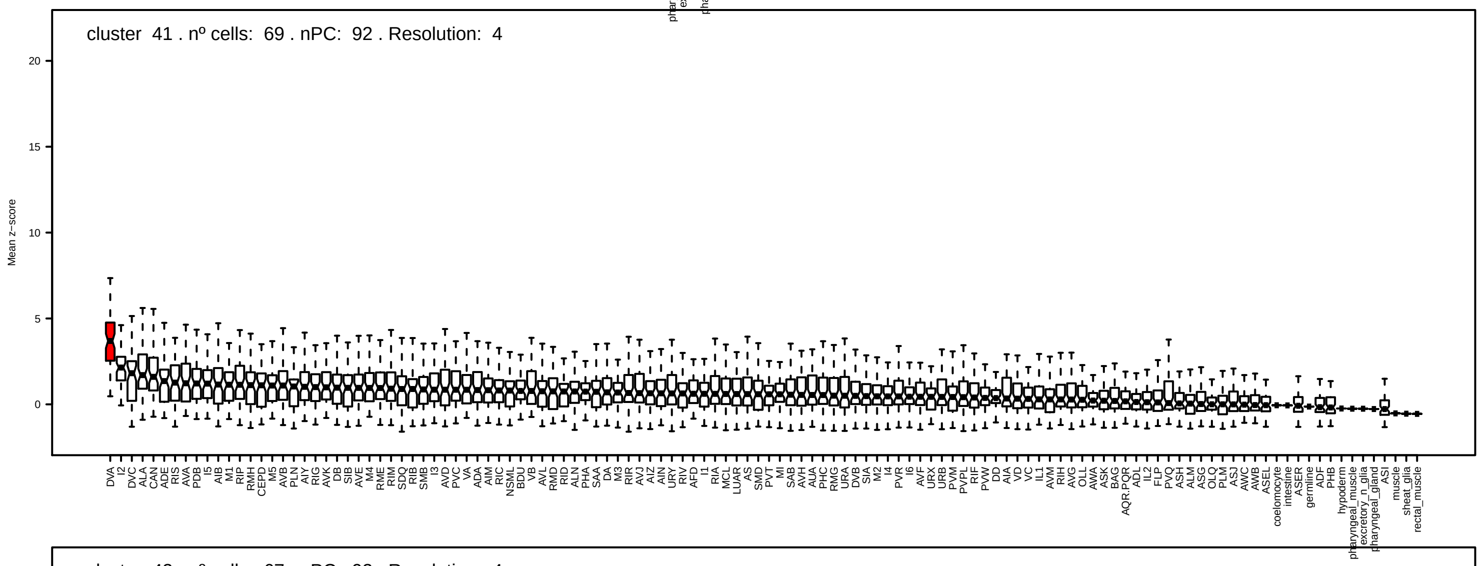
cluster 41 . nº cells: 69 . nPC: 92 . Resolution: 4
Mean z−score
20
15
10
5
0

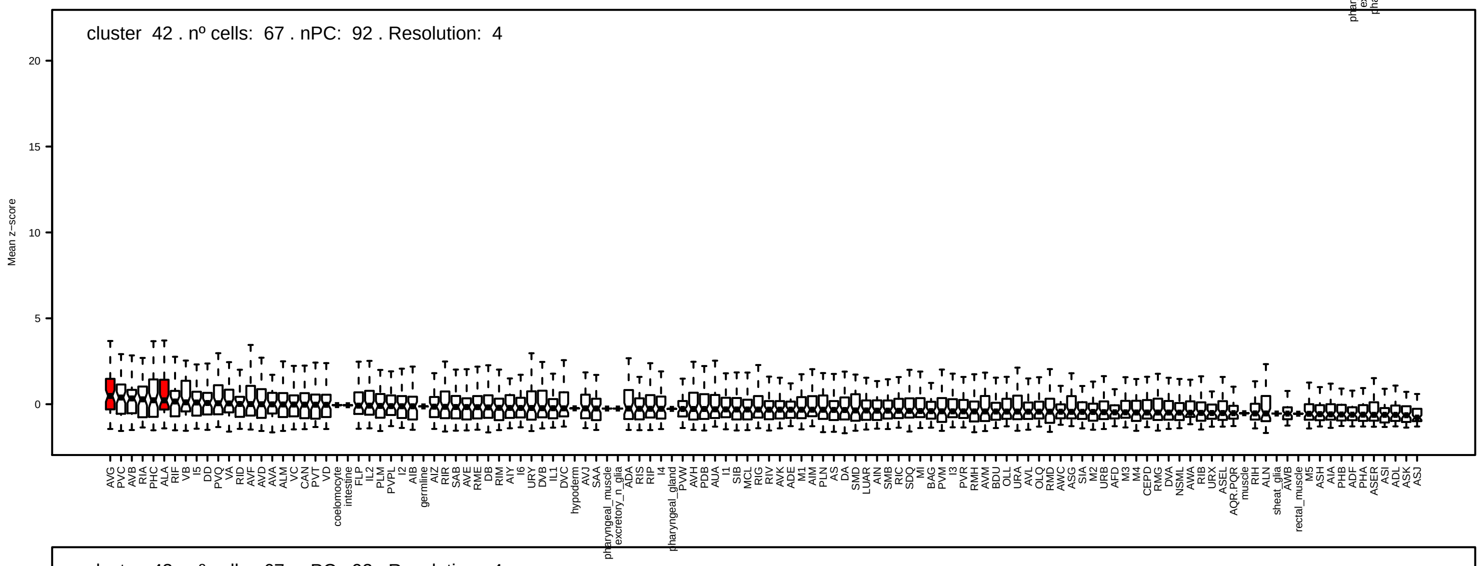
cluster 42 . nº cells: 67 . nPC: 92 . Resolution: 4
Mean z−score
20
15
10
5
0

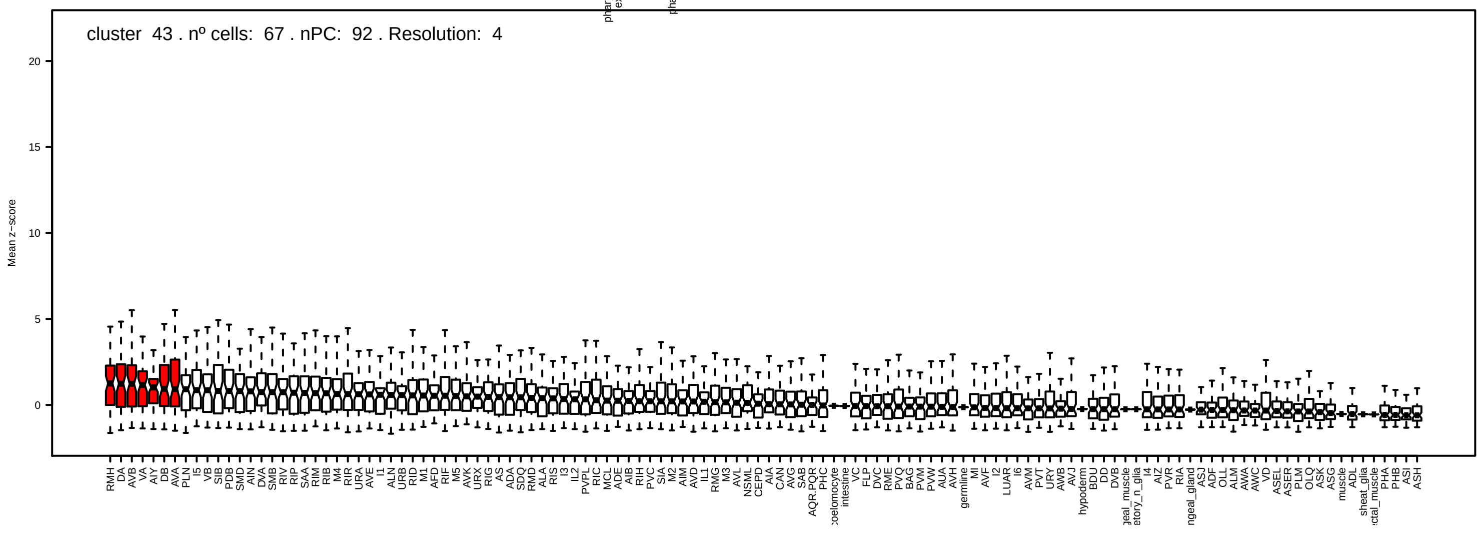
cluster 43 . nº cells: 67 . nPC: 92 . Resolution: 4
Mean z−score
20
15
10
5
0

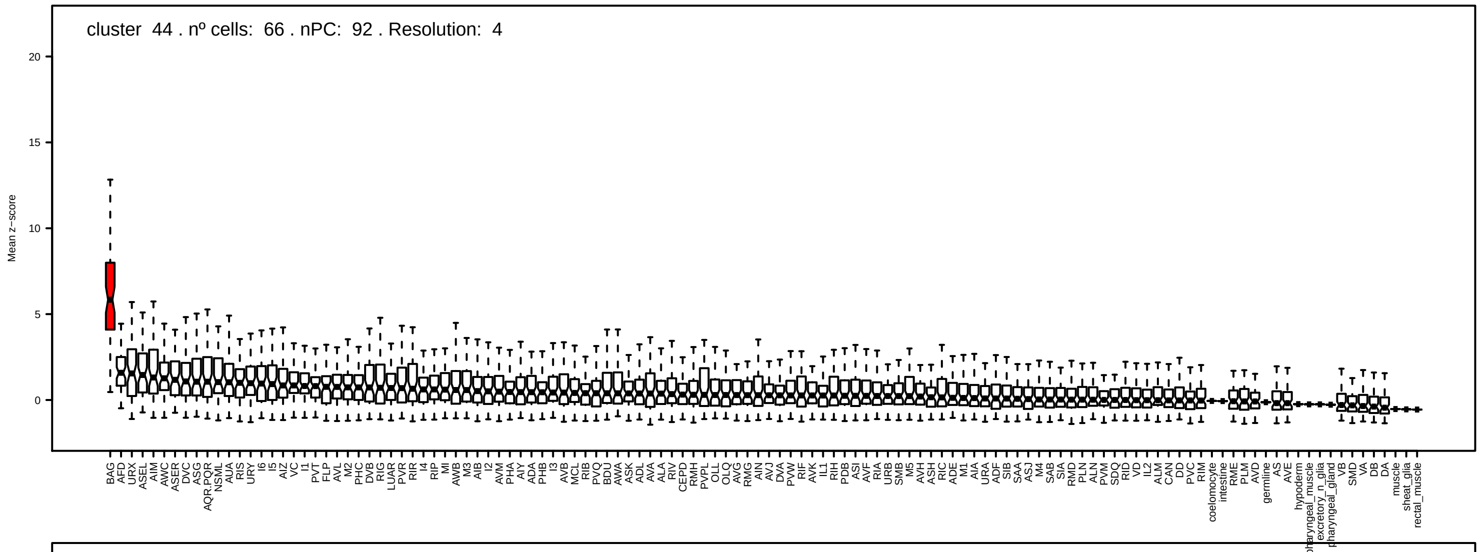
cluster 44 . nº cells: 66 . nPC: 92 . Resolution: 4
Mean z−score

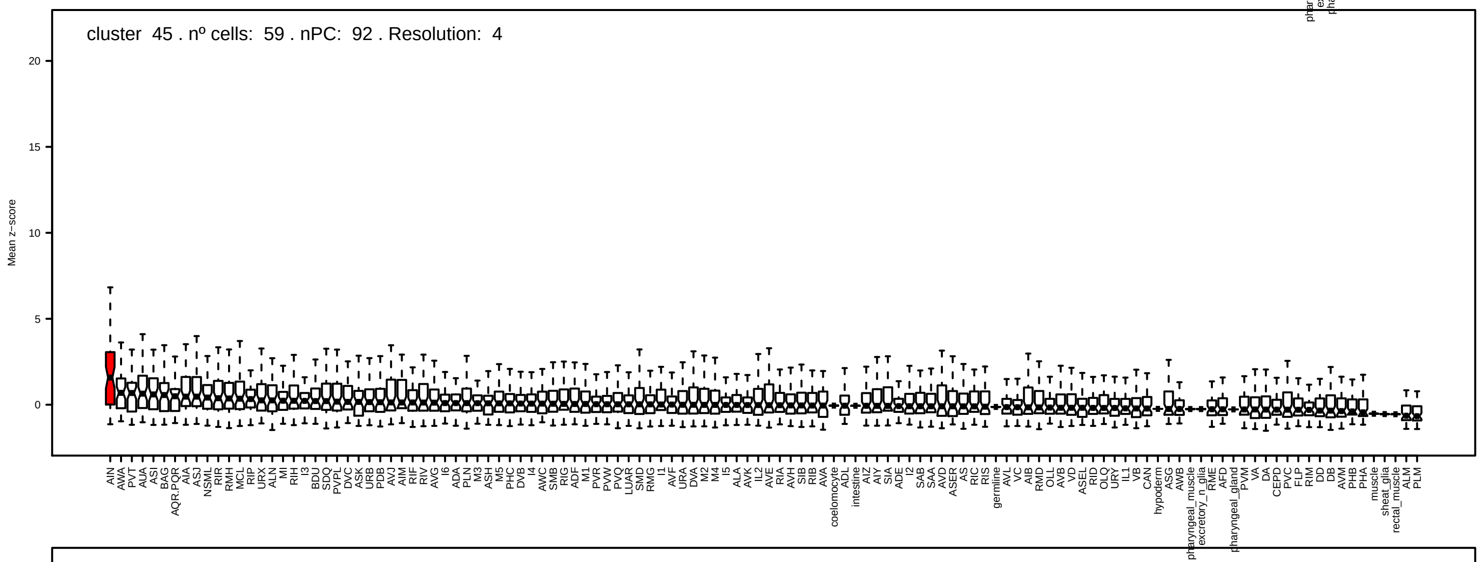
cluster 45 . nº cells: 59 . nPC: 92 . Resolution: 4
Mean z−score

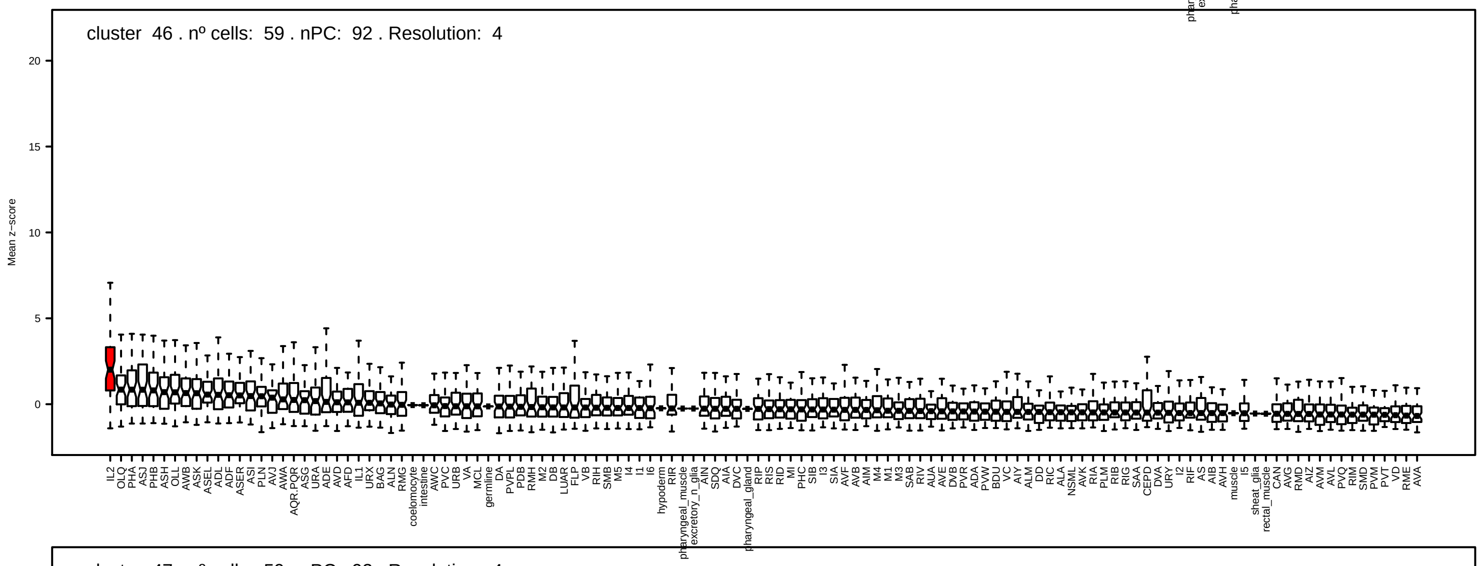
cluster 46 . nº cells: 59 . nPC: 92 . Resolution: 4
Mean z−score

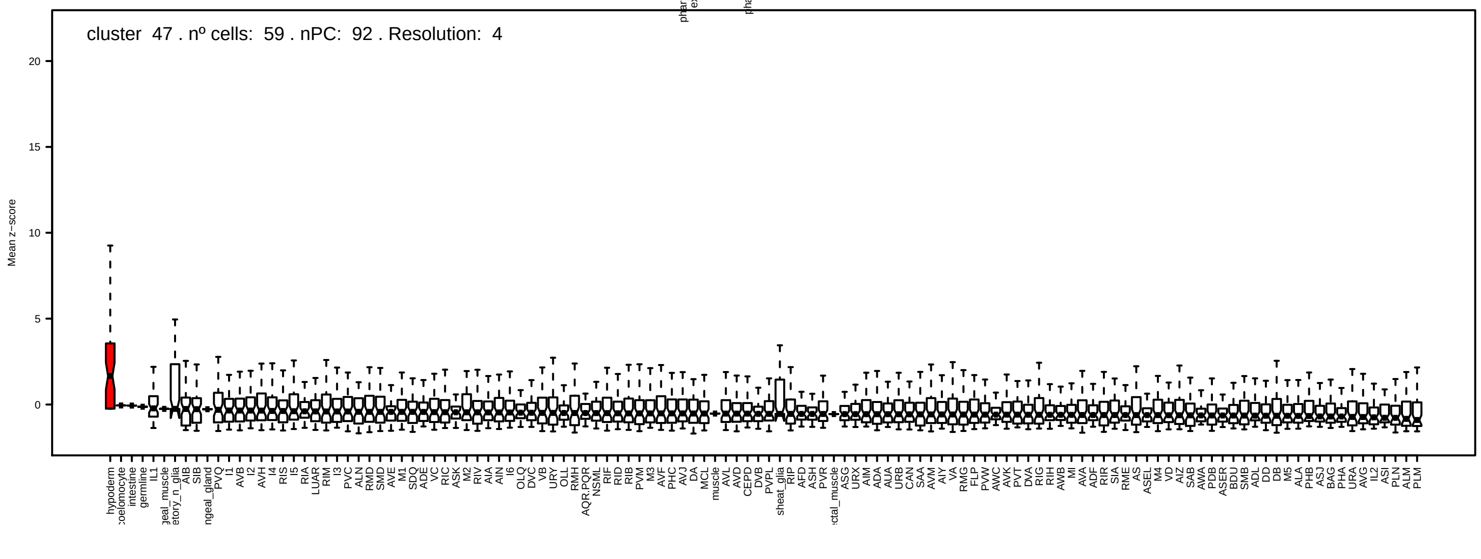
cluster 47 . nº cells: 59 . nPC: 92 . Resolution: 4
Mean z−score

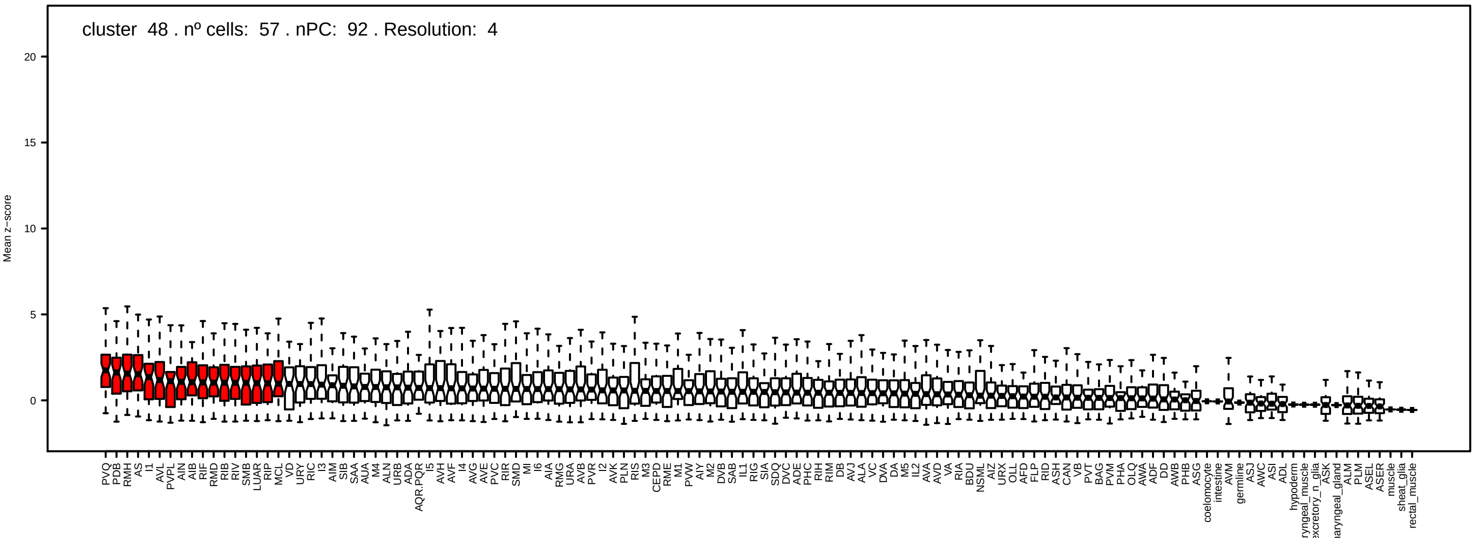
cluster 48 . nº cells: 57 . nPC: 92 . Resolution: 4
Mean z−score

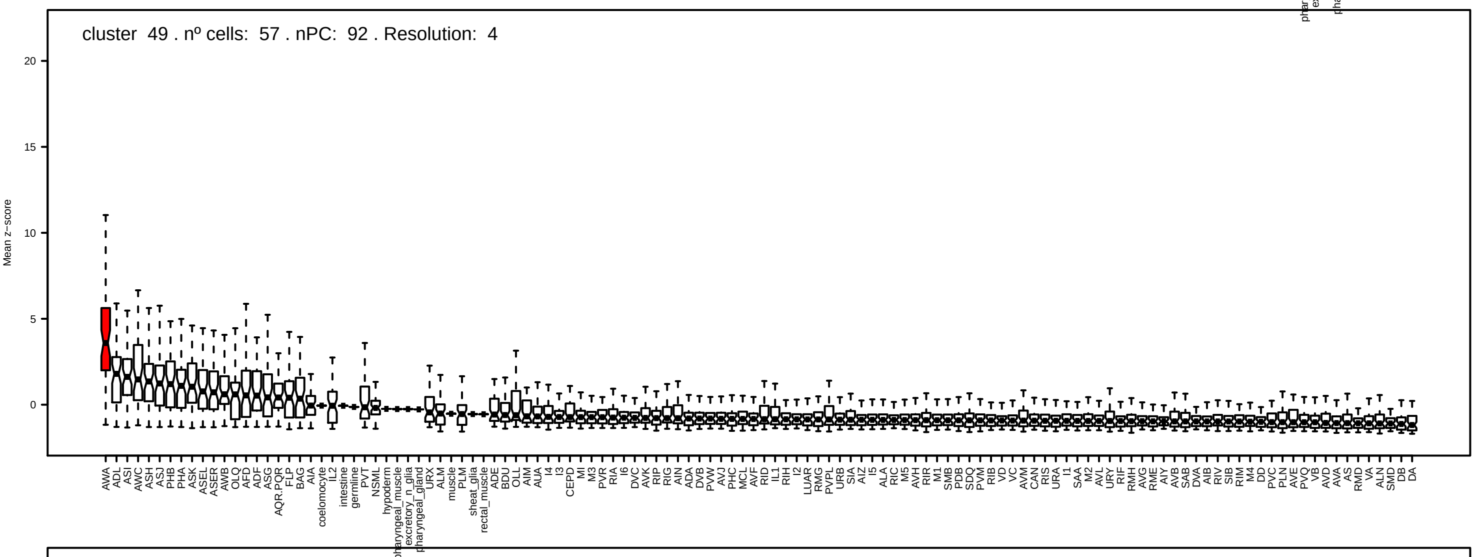
cluster 49 . nº cells: 57 . nPC: 92 . Resolution: 4
Mean z−score

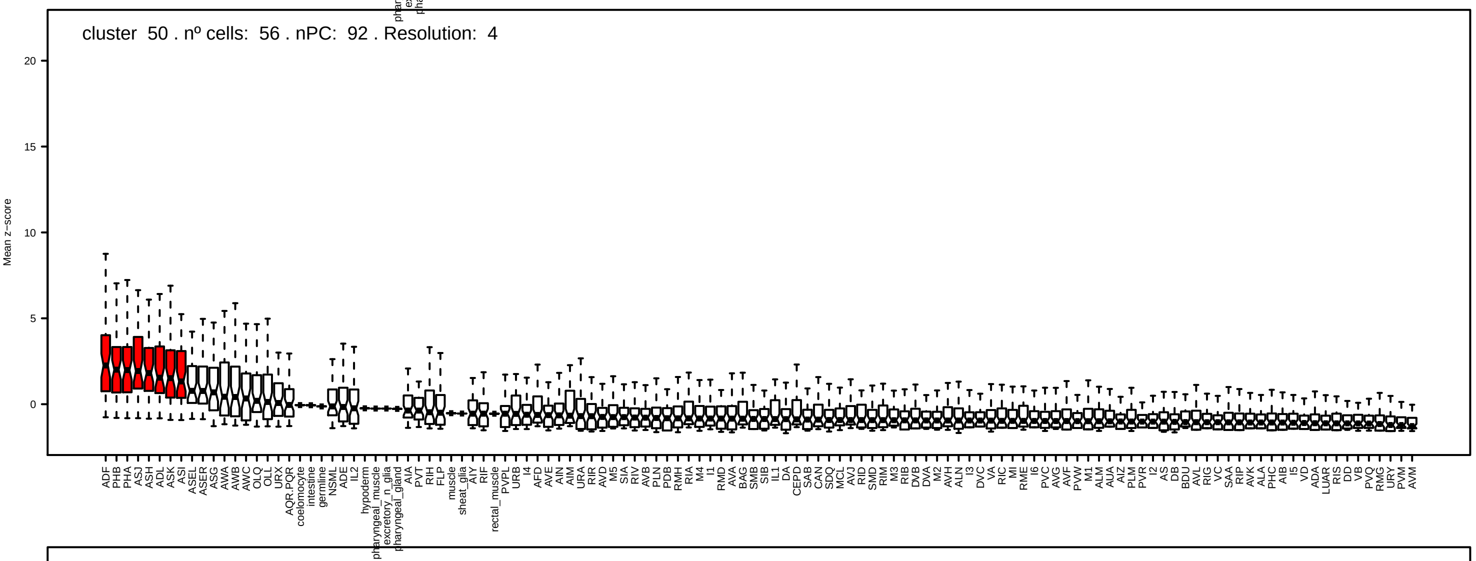
cluster 50 . nº cells: 56 . nPC: 92 . Resolution: 4
Mean z−score

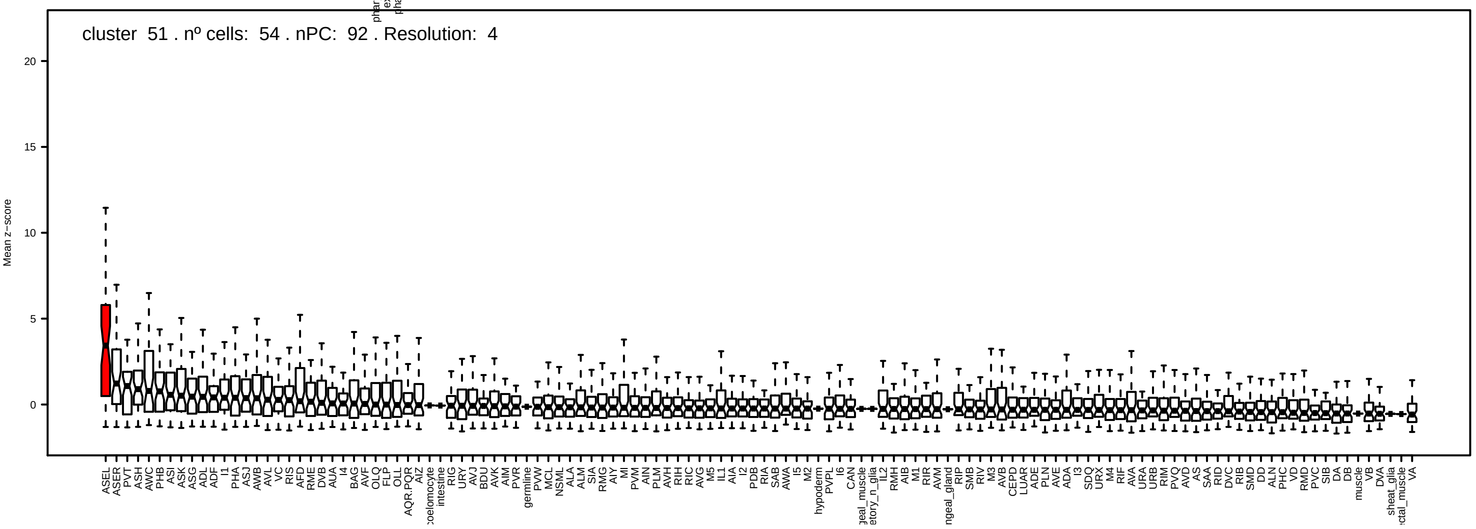
cluster 51 . nº cells: 54 . nPC: 92 . Resolution: 4
Mean z−score

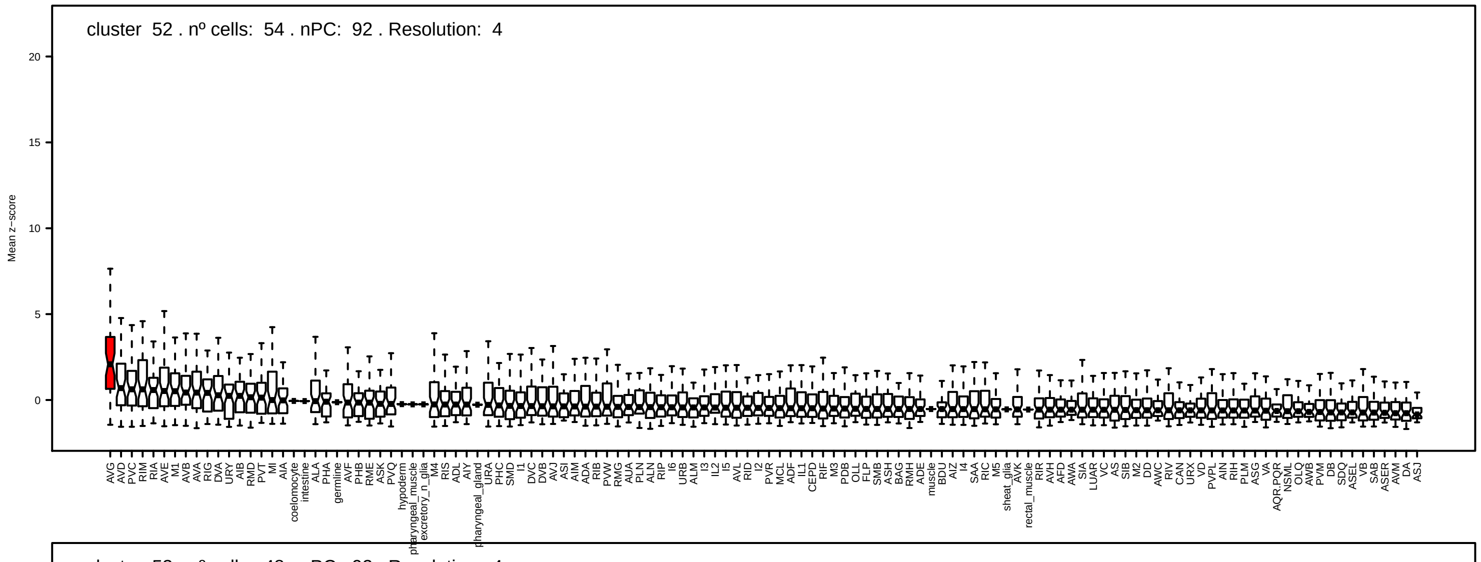
cluster 52 . nº cells: 54 . nPC: 92 . Resolution: 4
Mean z−score
20
15
10
5
0

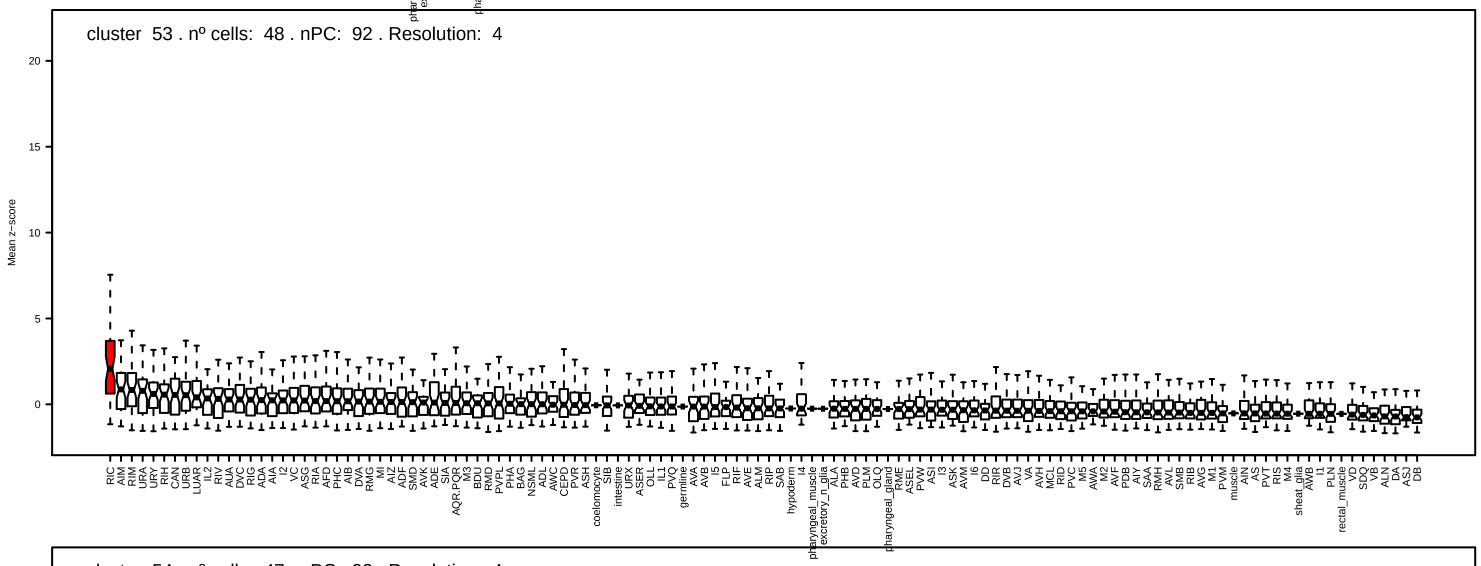
cluster 53 . nº cells: 48 . nPC: 92 . Resolution: 4
Mean z−score
20
15
10
5
0

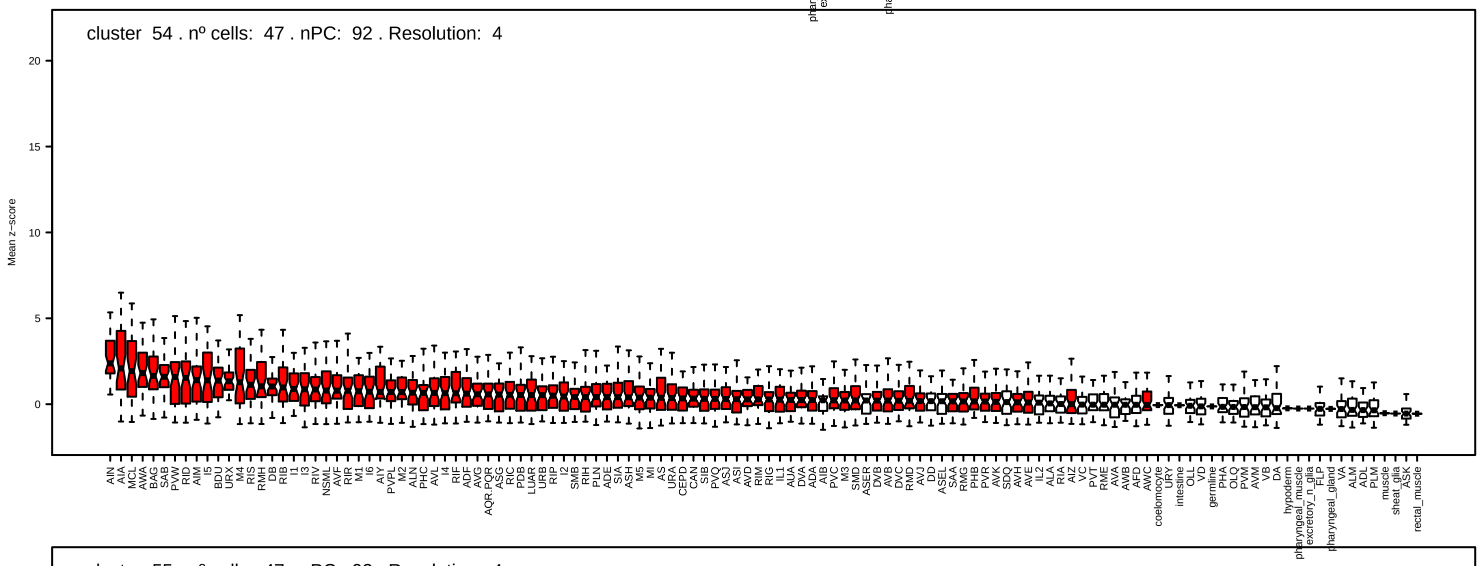
cluster 54 . nº cells: 47 . nPC: 92 . Resolution: 4
Mean z−score
20
15
10
5
0

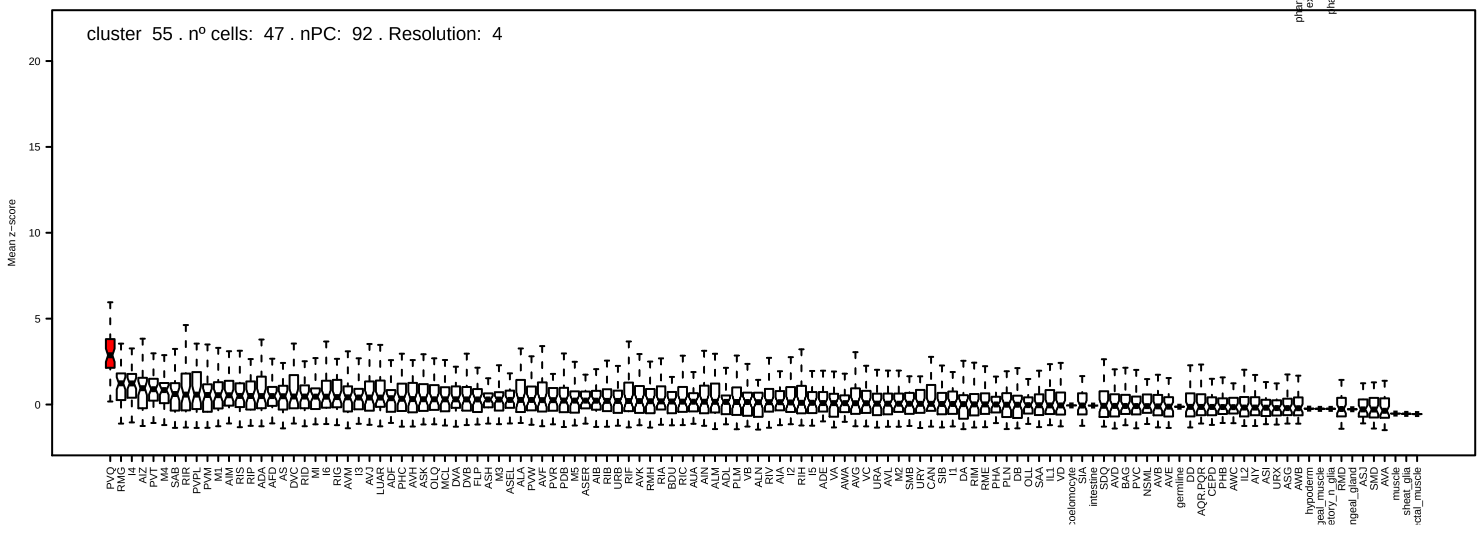
cluster 55 . nº cells: 47 . nPC: 92 . Resolution: 4
Mean z−score
20
15
10
5
0

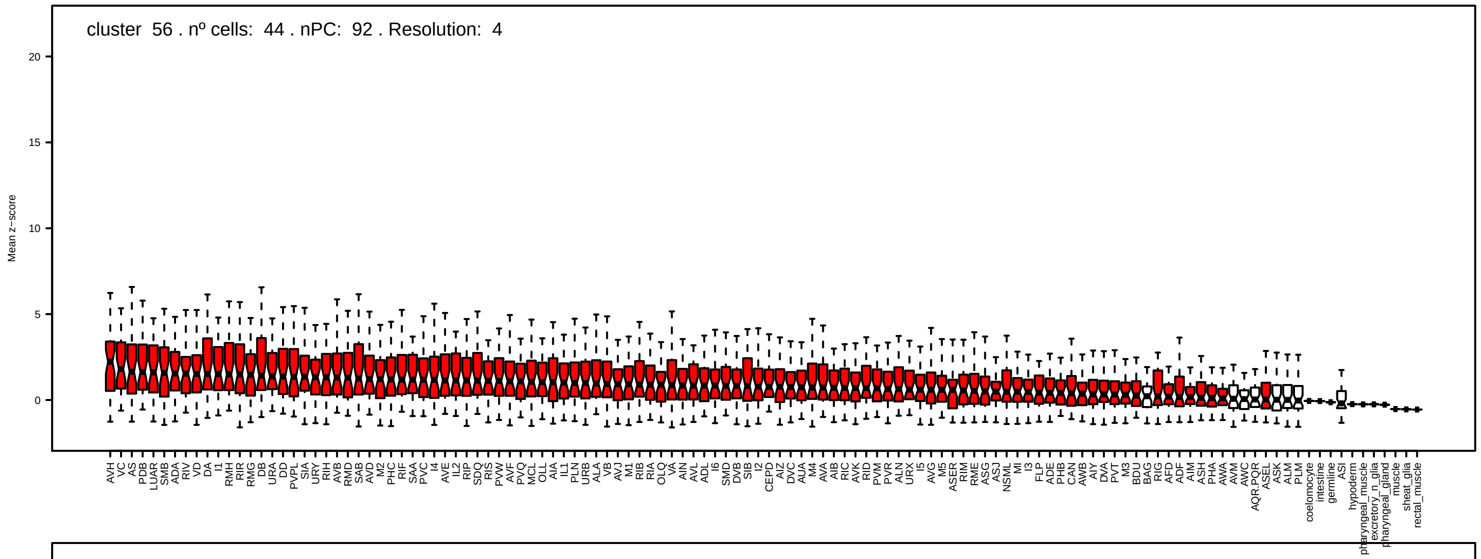
cluster 56 . nº cells: 44 . nPC: 92 . Resolution: 4
Mean z−score

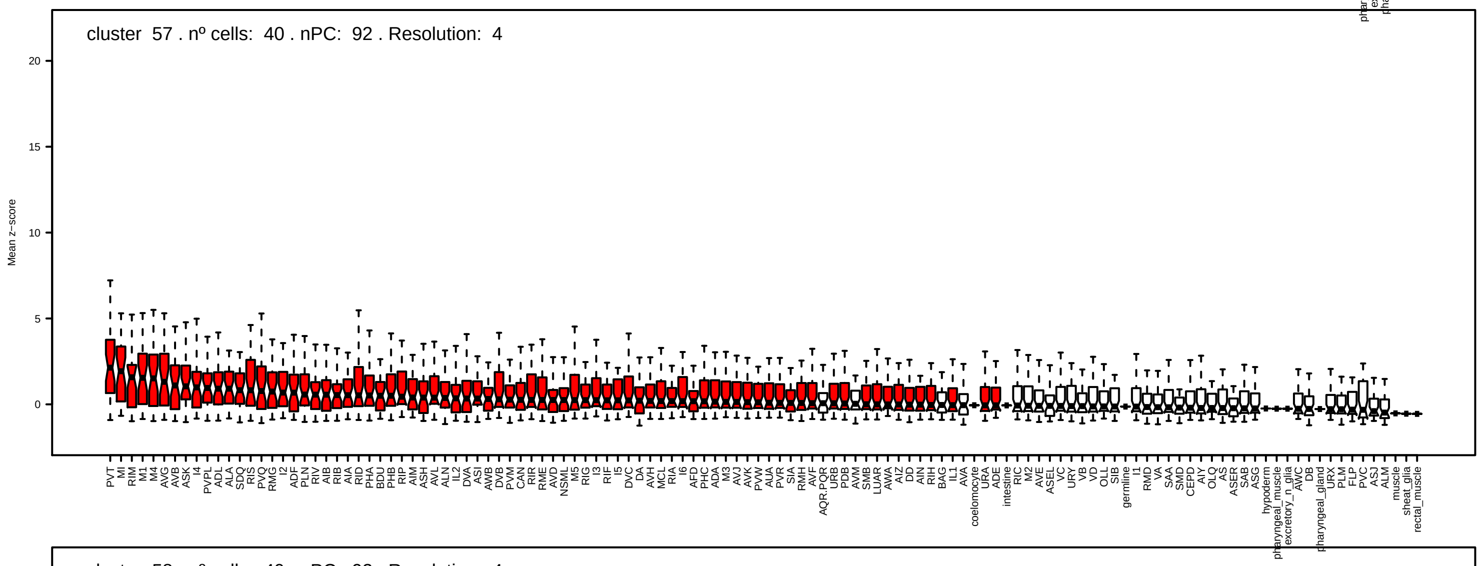
cluster 57 . nº cells: 40 . nPC: 92 . Resolution: 4
Mean z−score

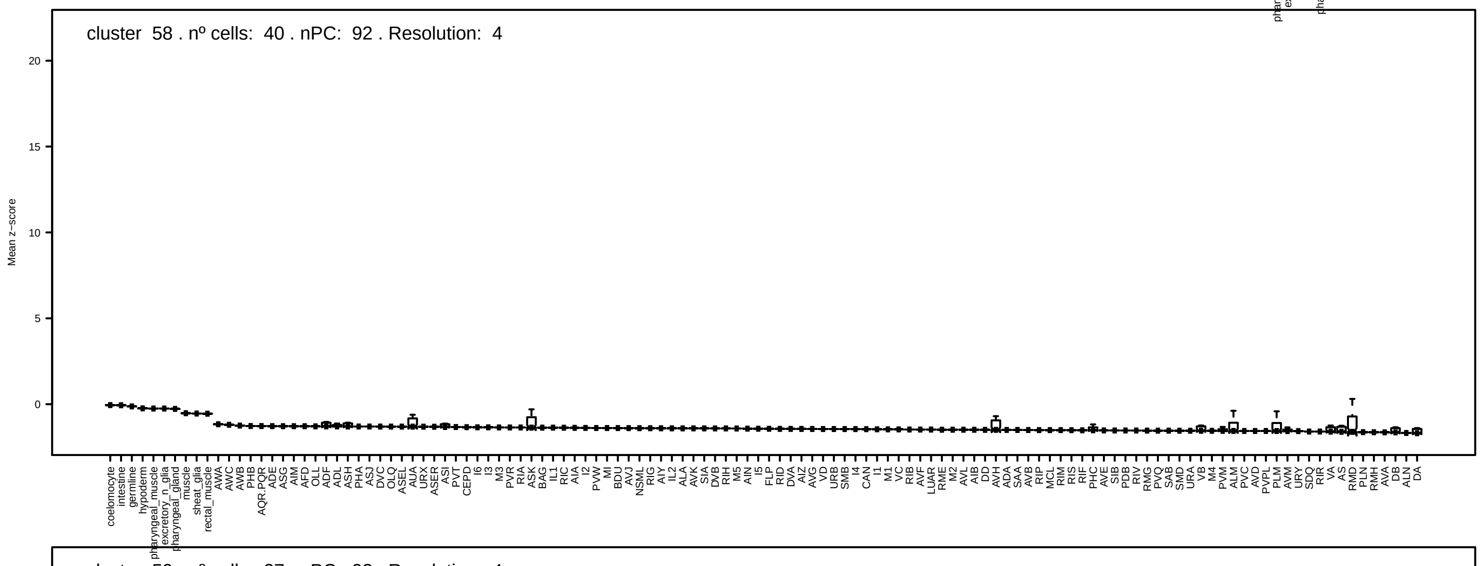
cluster 58 . nº cells: 40 . nPC: 92 . Resolution: 4
Mean z−score

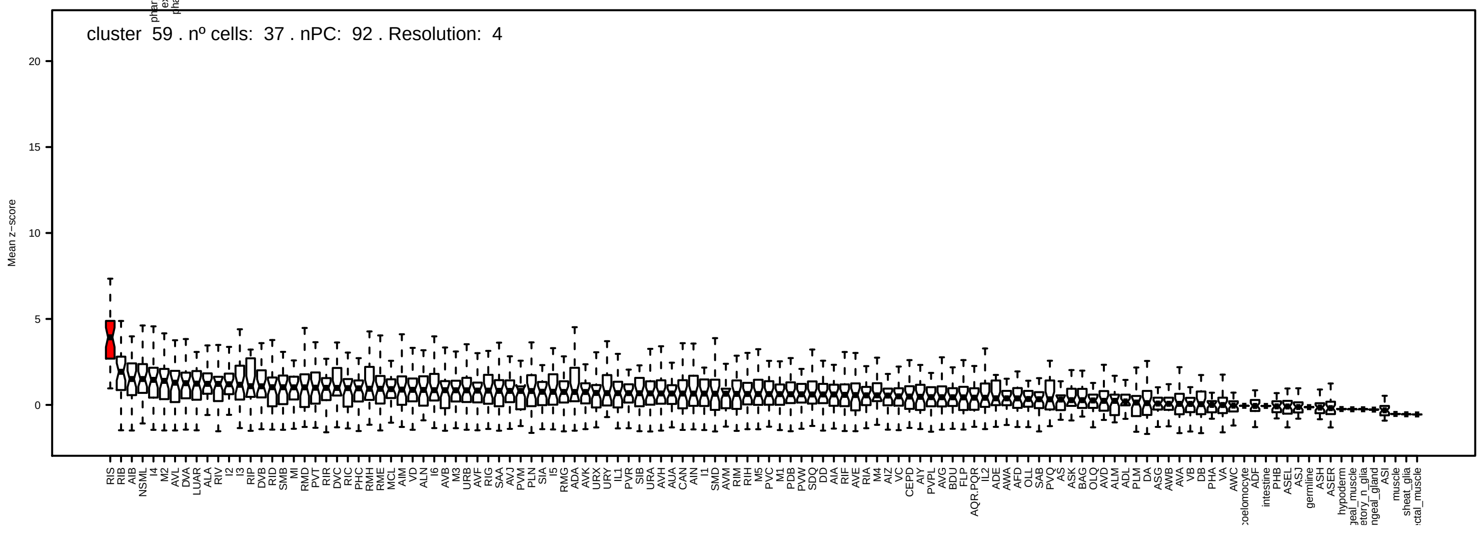
cluster 59 . nº cells: 37 . nPC: 92 . Resolution: 4
Mean z−score

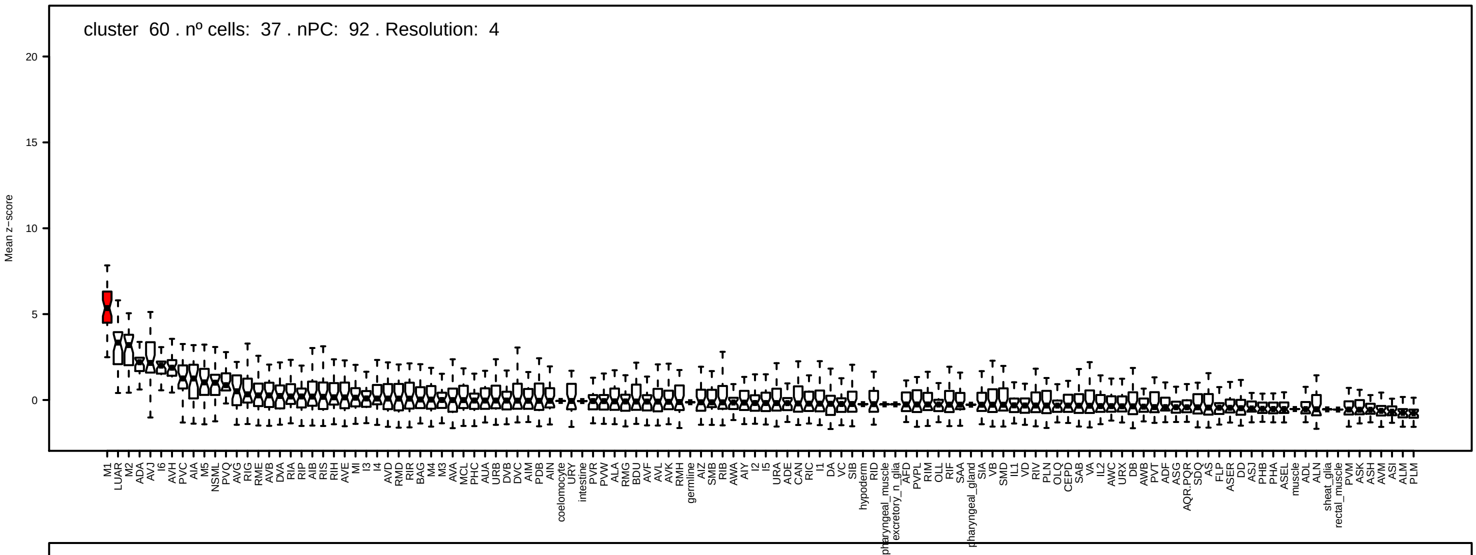
cluster 60 . nº cells: 37 . nPC: 92 . Resolution: 4
Mean z−score

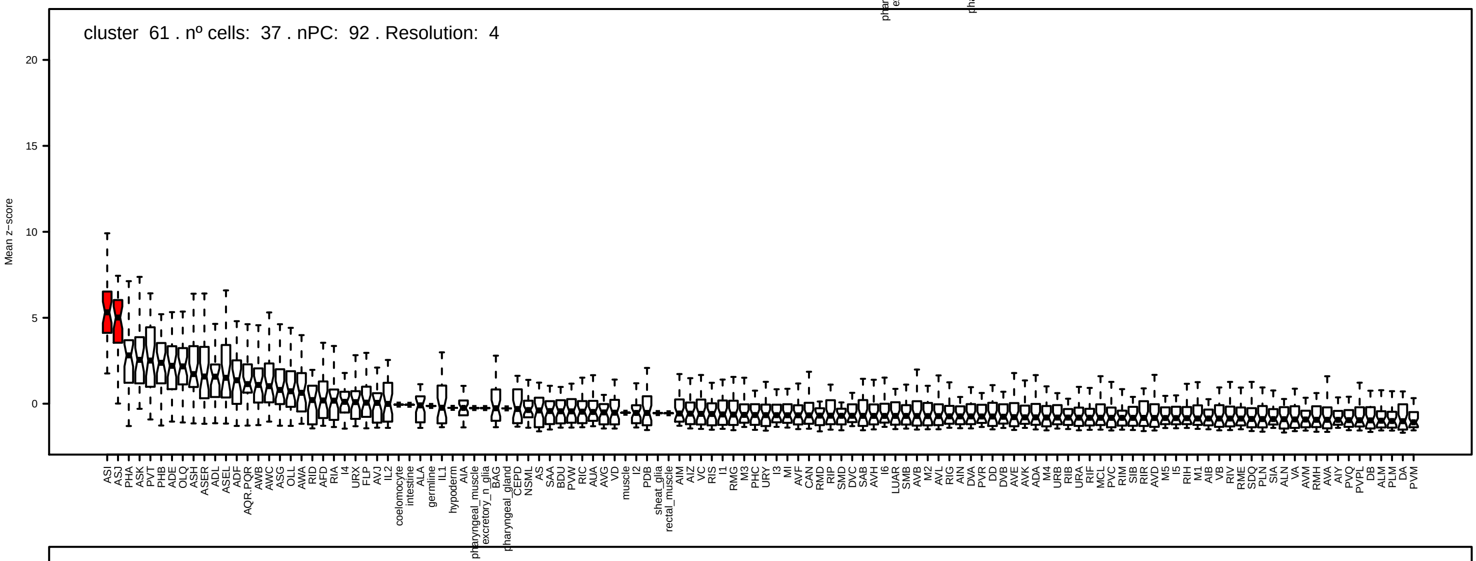
cluster 61 . nº cells: 37 . nPC: 92 . Resolution: 4
Mean z−score

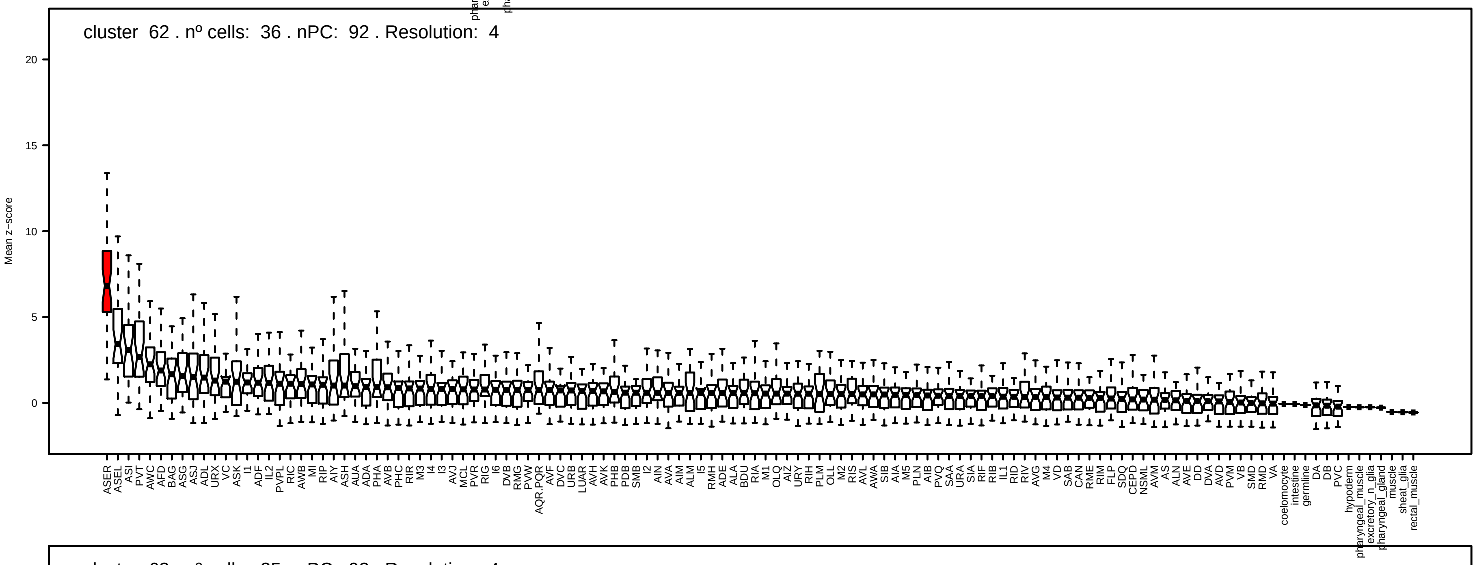
cluster 62 . nº cells: 36 . nPC: 92 . Resolution: 4
Mean z−score

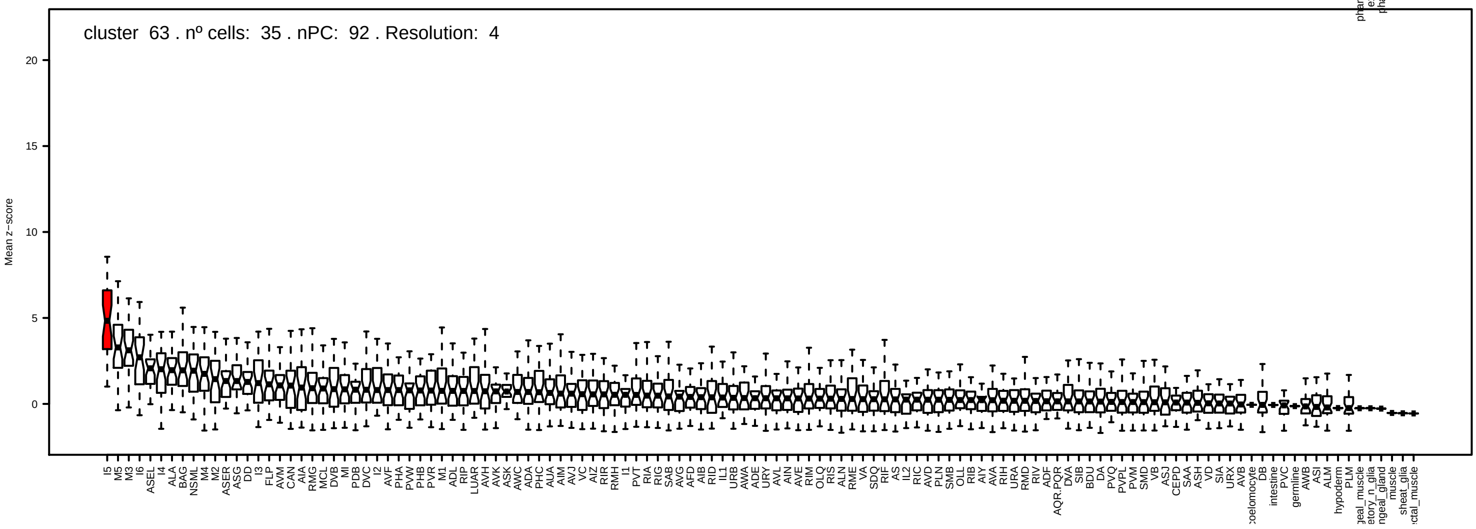
cluster 63 . nº cells: 35 . nPC: 92 . Resolution: 4
Mean z−score

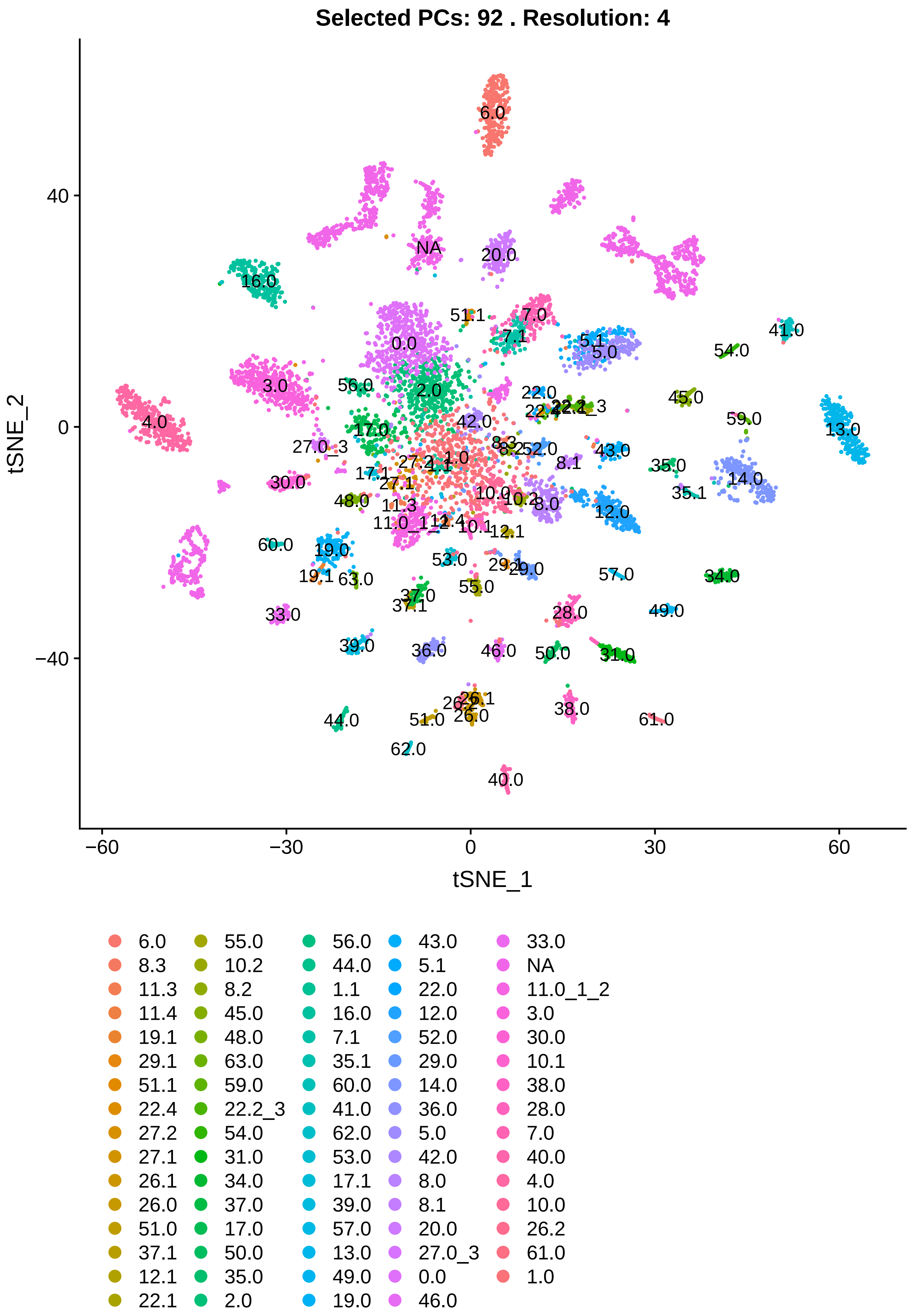
Selected PCs: 92 . Resolution: 4
tSNE_2
tSNE_1
40
0
−40
−60
−30
0
30
60
6.0
8.3
11.3
11.4
19.1
29.1
51.1
22.4
27.2
27.1
26.1
26.0
51.0
37.1
12.1
22.1
55.0
10.2
8.2
45.0
48.0
63.0
59.0
22.2_3
54.0
31.0
34.0
37.0
17.0
50.0
35.0
2.0
56.0
44.0
1.1
16.0
7.1
35.1
60.0
41.0
62.0
53.0
17.1
39.0
57.0
13.0
49.0
19.0
43.0
5.1
22.0
12.0
52.0
29.0
14.0
36.0
5.0
42.0
8.0
8.1
20.0
27.0_3
0.0
46.0
33.0
NA
11.0_1_2
3.0
30.0
10.1
38.0
28.0
7.0
40.0
4.0
10.0
26.2
61.0
1.0

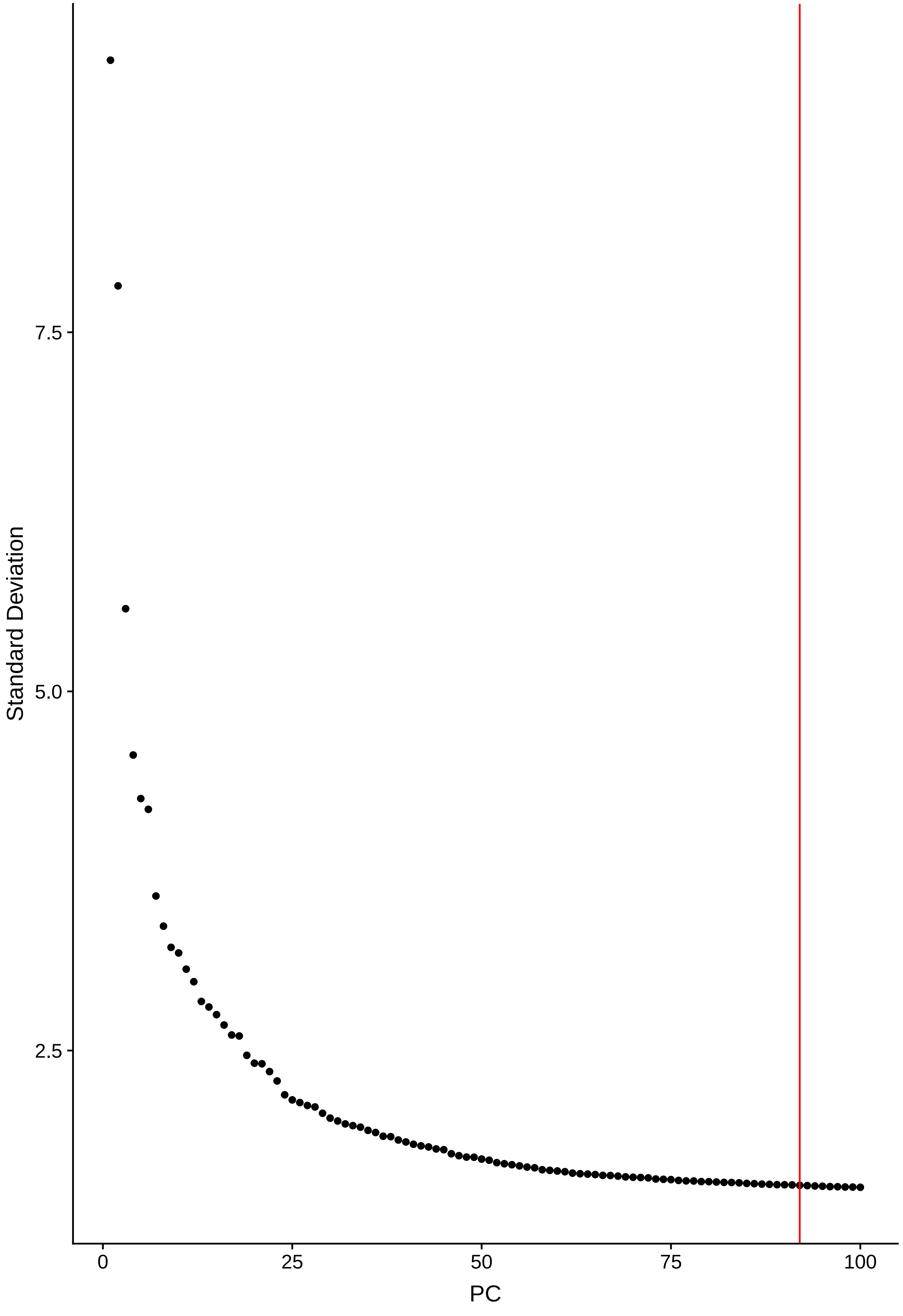
Standard Deviation
7.5
5.0
2.5
0
25
50
75
100
PC

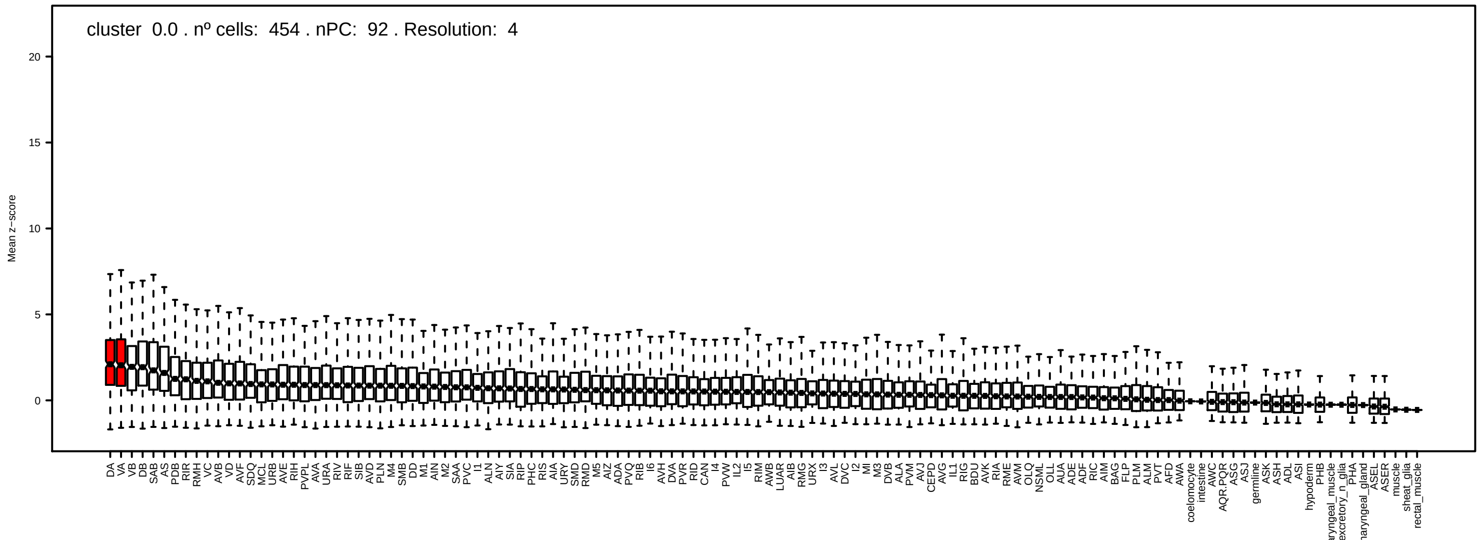
cluster 0.0 . nº cells: 454 . nPC: 92 . Resolution: 4
Mean z−score

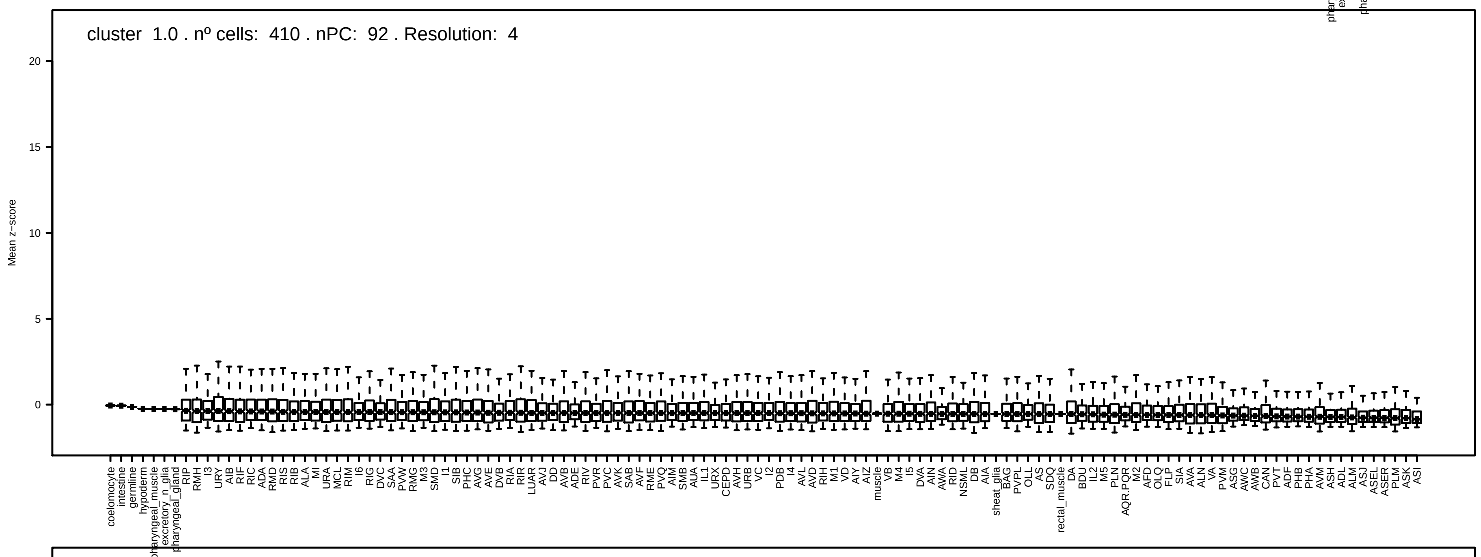
cluster 1.0 . nº cells: 410 . nPC: 92 . Resolution: 4
Mean z−score

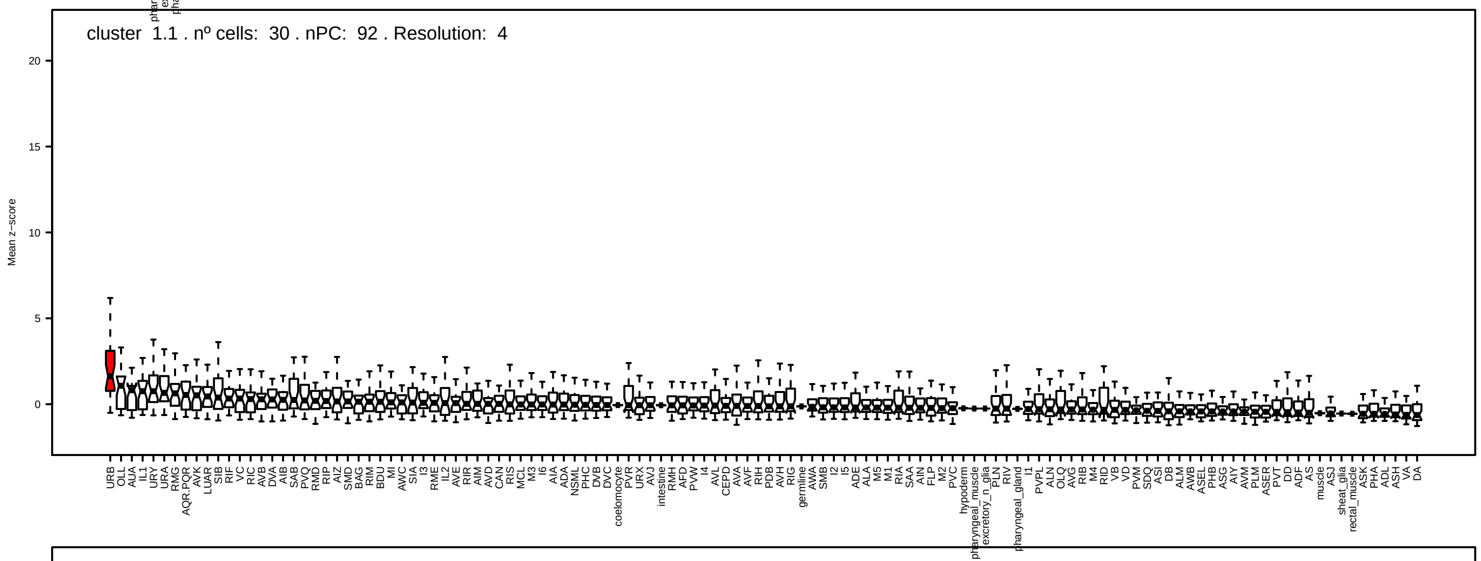
cluster 1.1 . nº cells: 30 . nPC: 92 . Resolution: 4
Mean z−score

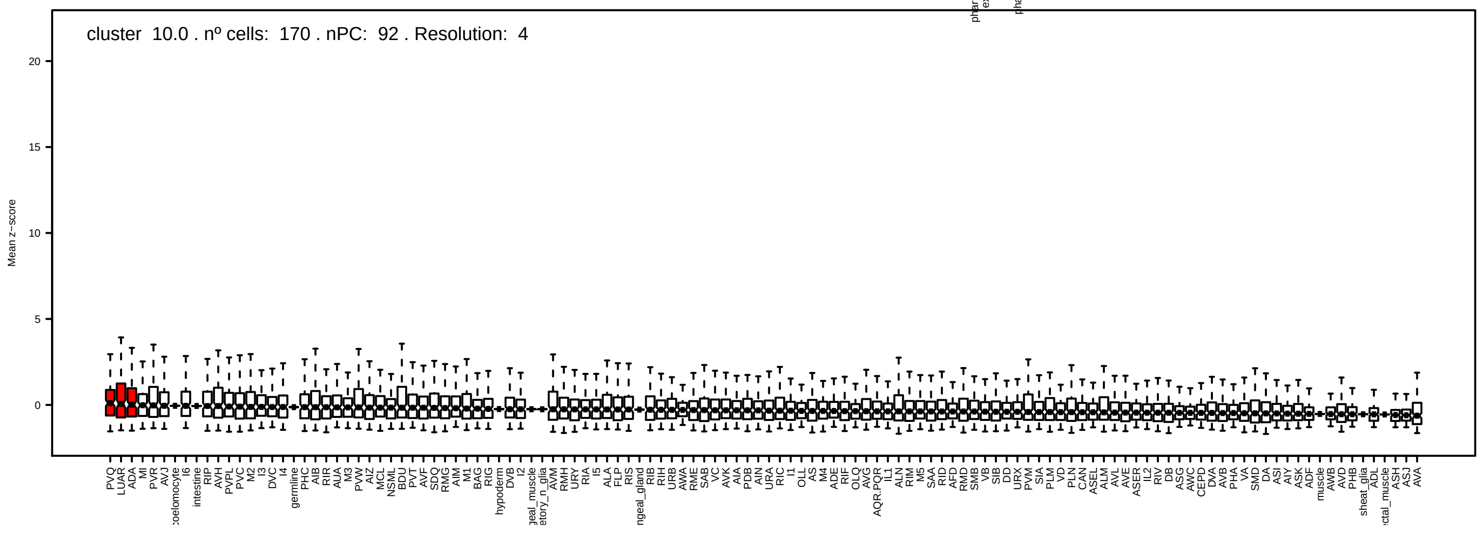
cluster 10.0 . nº cells: 170 . nPC: 92 . Resolution: 4
Mean z−score

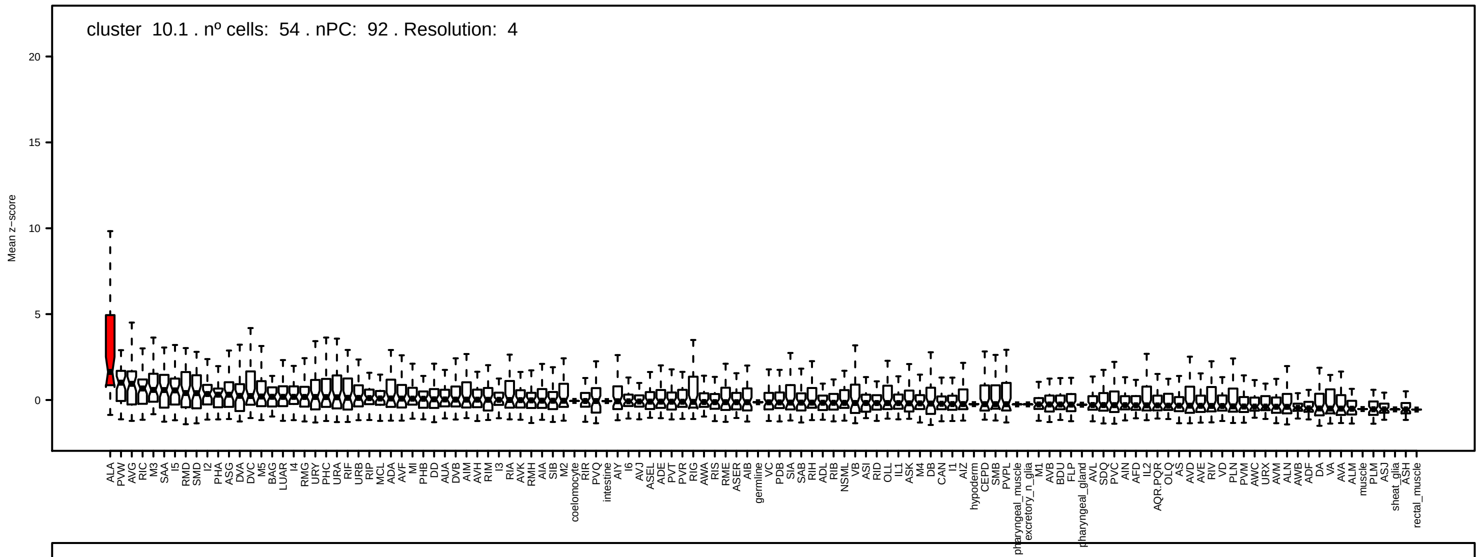
cluster 10.1 . nº cells: 54 . nPC: 92 . Resolution: 4
Mean z−score
20
15
10
5
0

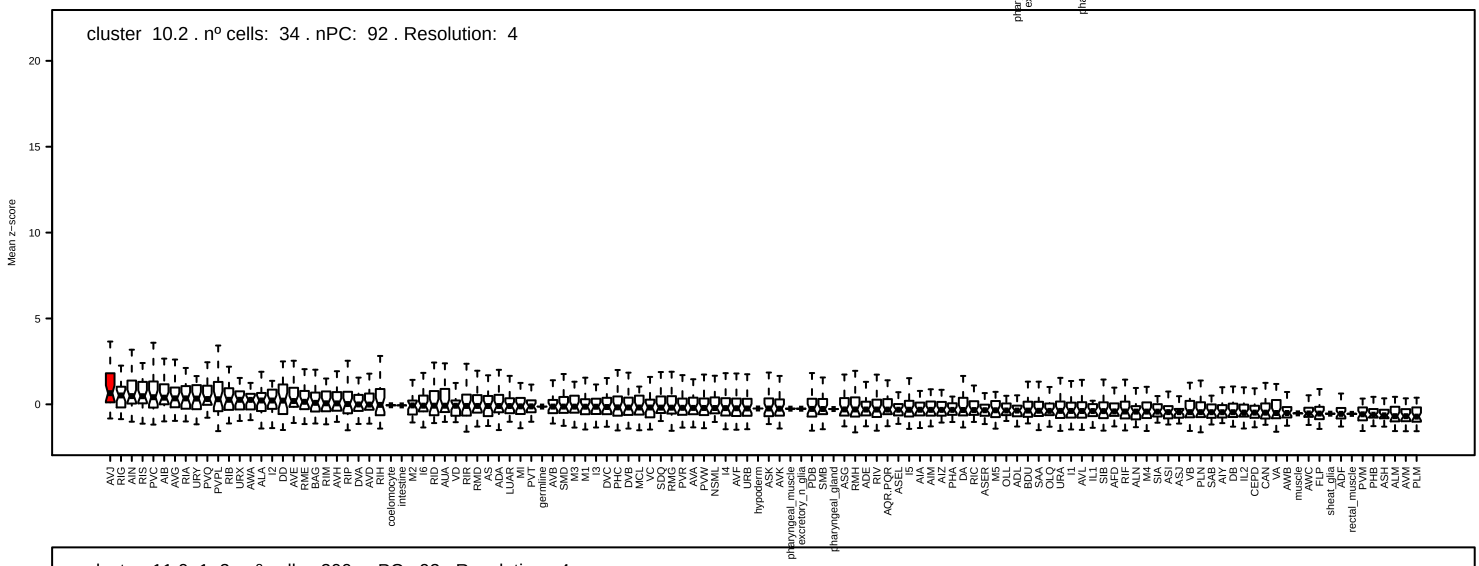
cluster 10.2 . nº cells: 34 . nPC: 92 . Resolution: 4
Mean z−score
20
15
10
5
0

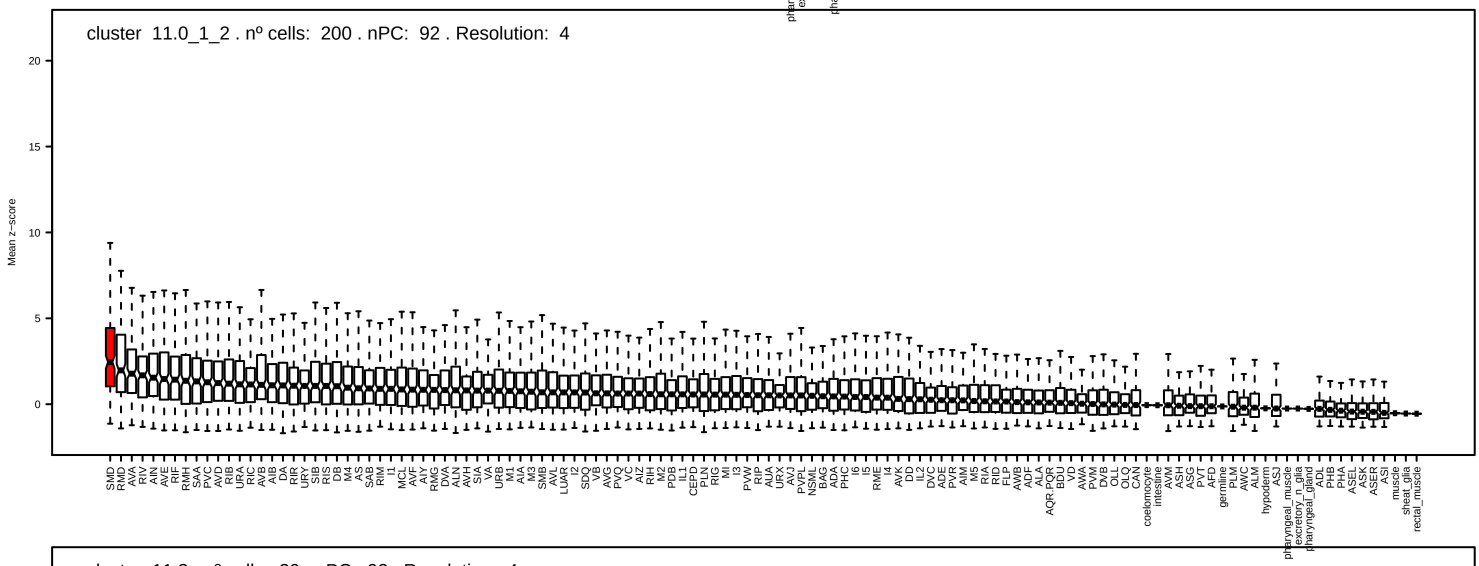
cluster 11.0_1_2 . nº cells: 200 . nPC: 92 . Resolution: 4
Mean z−score
20
15
10
5
0

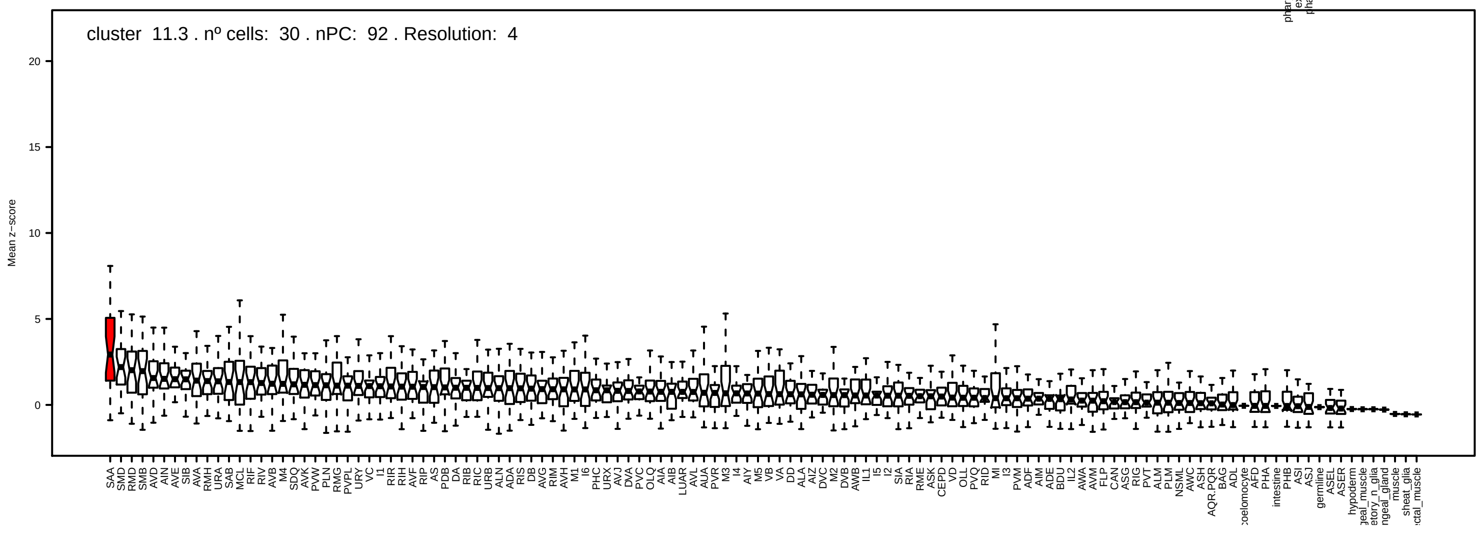
cluster 11.3 . nº cells: 30 . nPC: 92 . Resolution: 4
Mean z−score
20
15
10
5
0

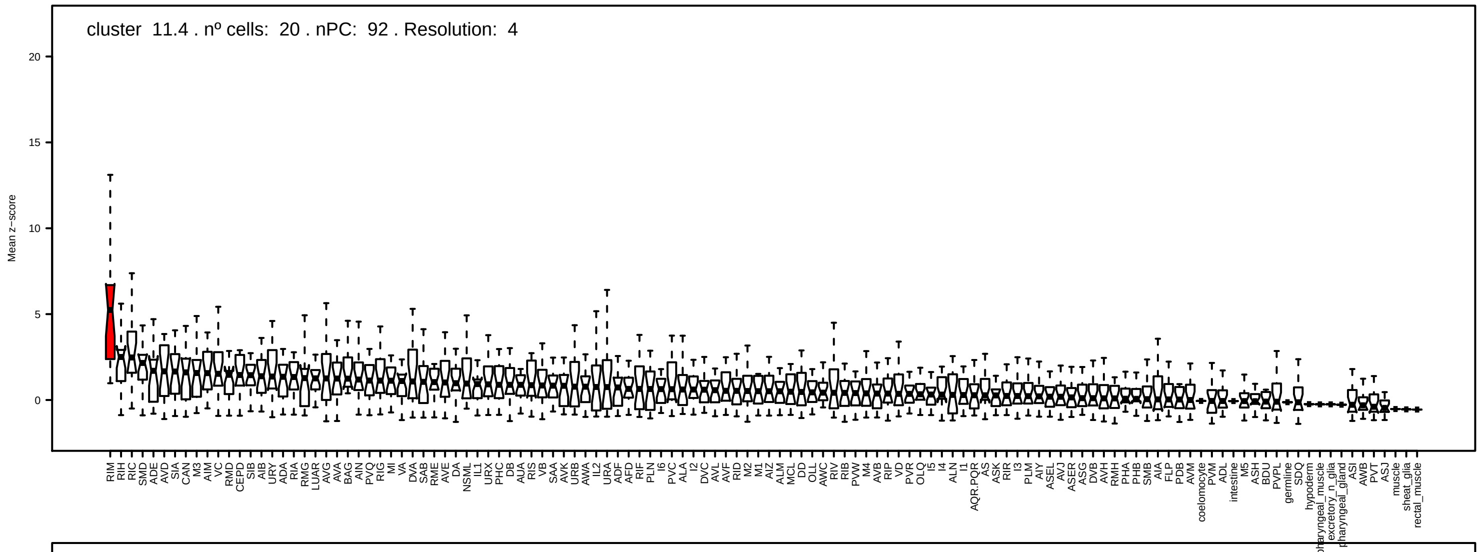
cluster 11.4 . nº cells: 20 . nPC: 92 . Resolution: 4
Mean z−score
0 5 10 15 20

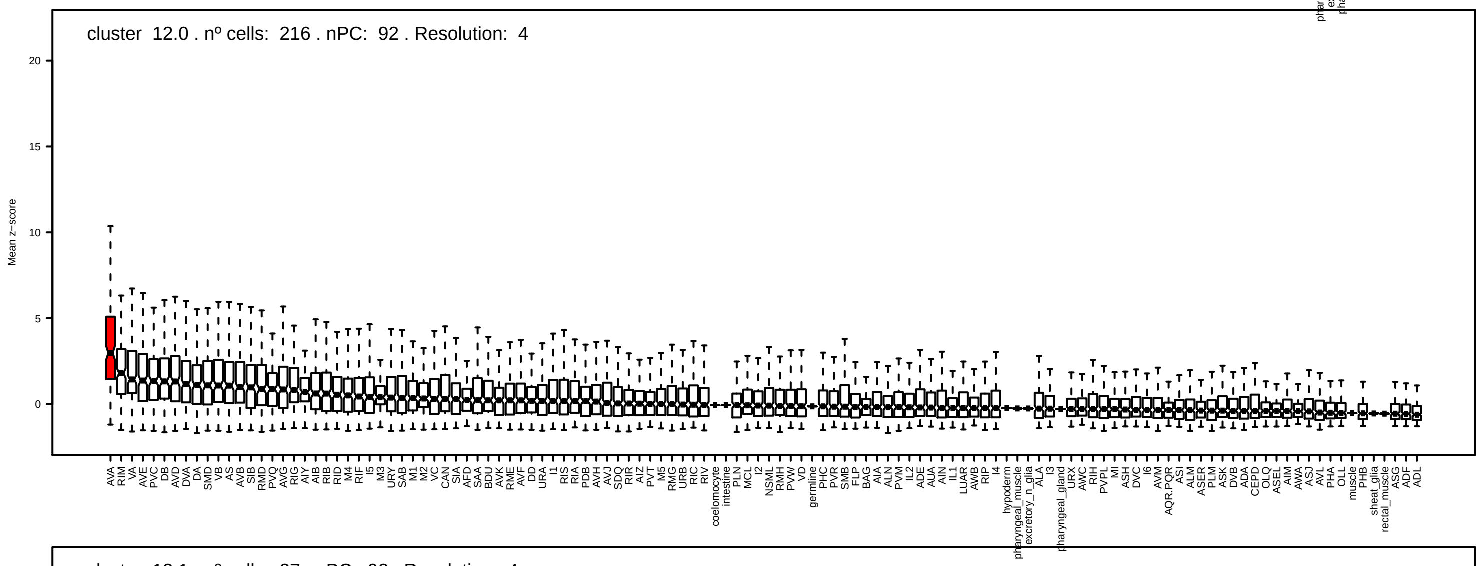
cluster 12.0 . nº cells: 216 . nPC: 92 . Resolution: 4
Mean z−score
0 5 10 15 20

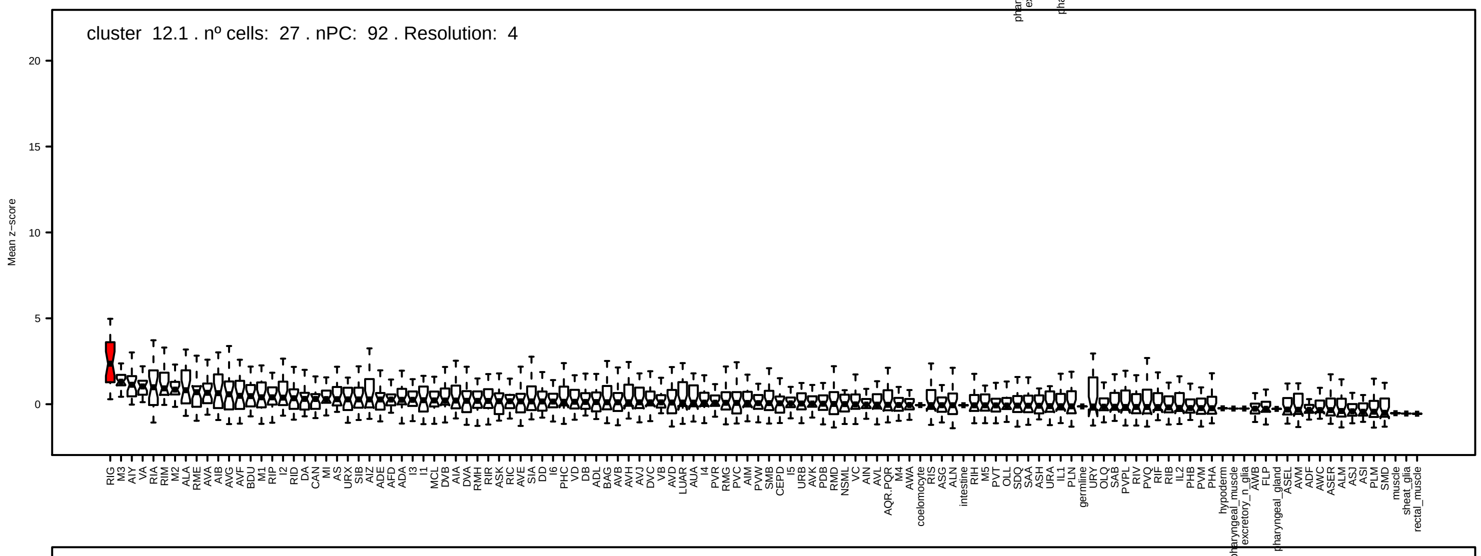
cluster 12.1 . nº cells: 27 . nPC: 92 . Resolution: 4
Mean z−score
0 5 10 15 20

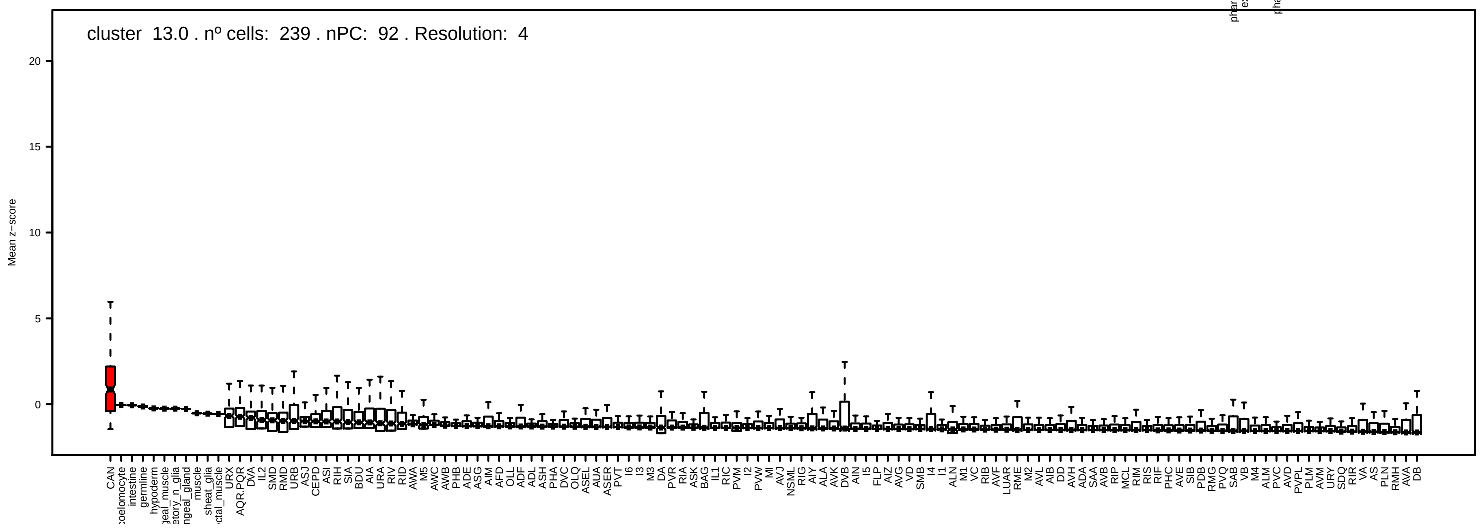
cluster 13.0 . nº cells: 239 . nPC: 92 . Resolution: 4
Mean z−score
0 5 10 15 20

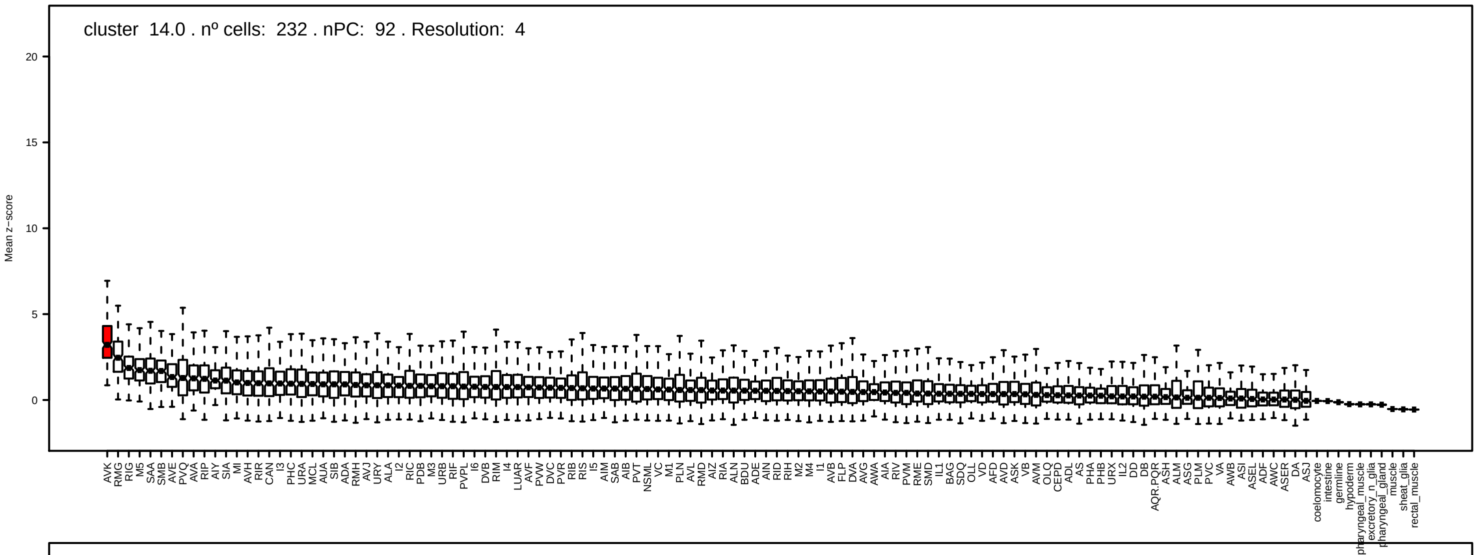
cluster 14.0 . nº cells: 232 . nPC: 92 . Resolution: 4
Mean z−score

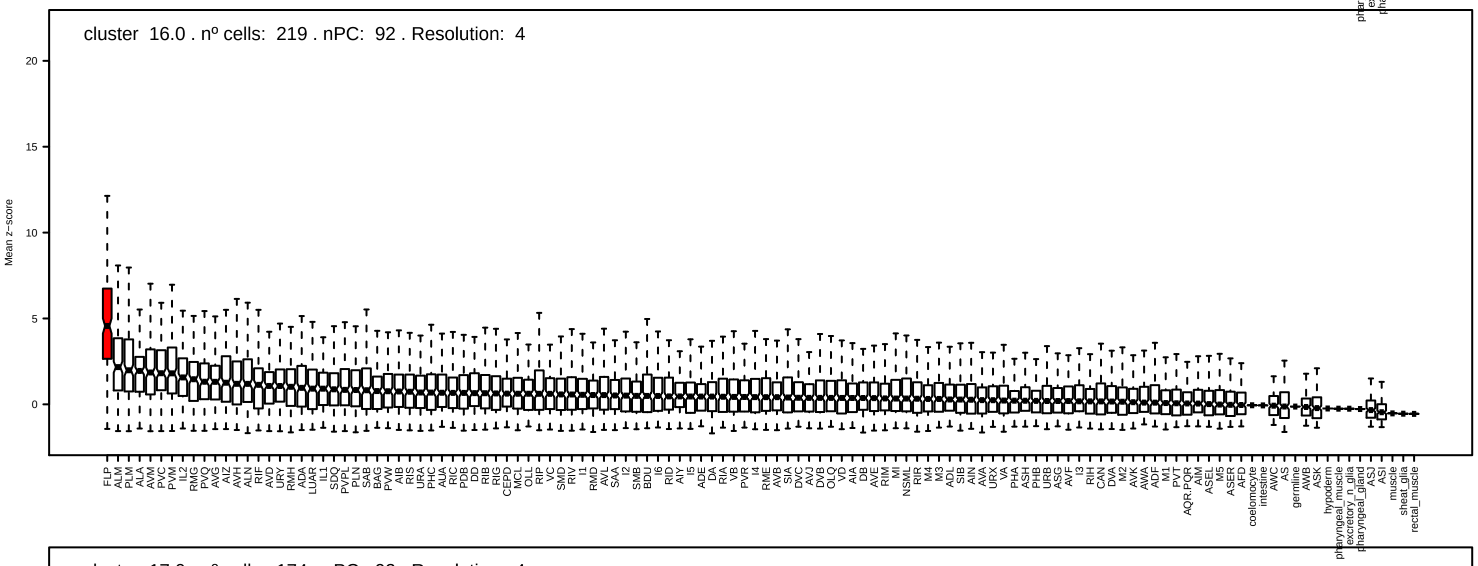
cluster 16.0 . nº cells: 219 . nPC: 92 . Resolution: 4
Mean z−score

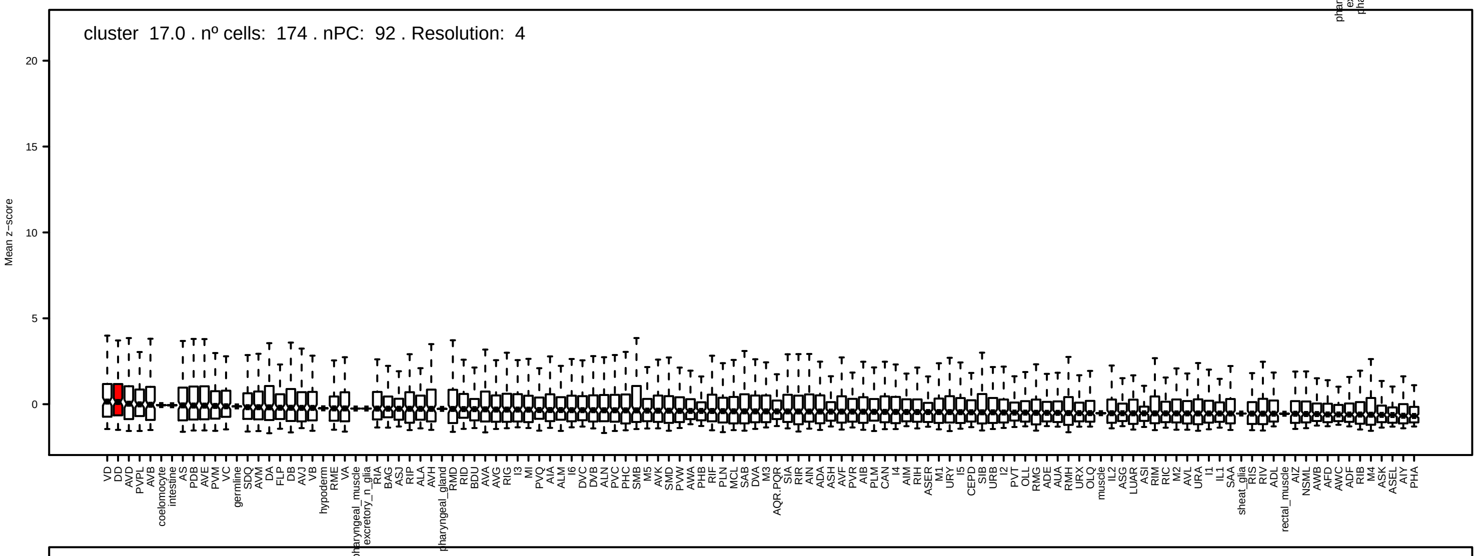
cluster 17.0 . nº cells: 174 . nPC: 92 . Resolution: 4
Mean z−score

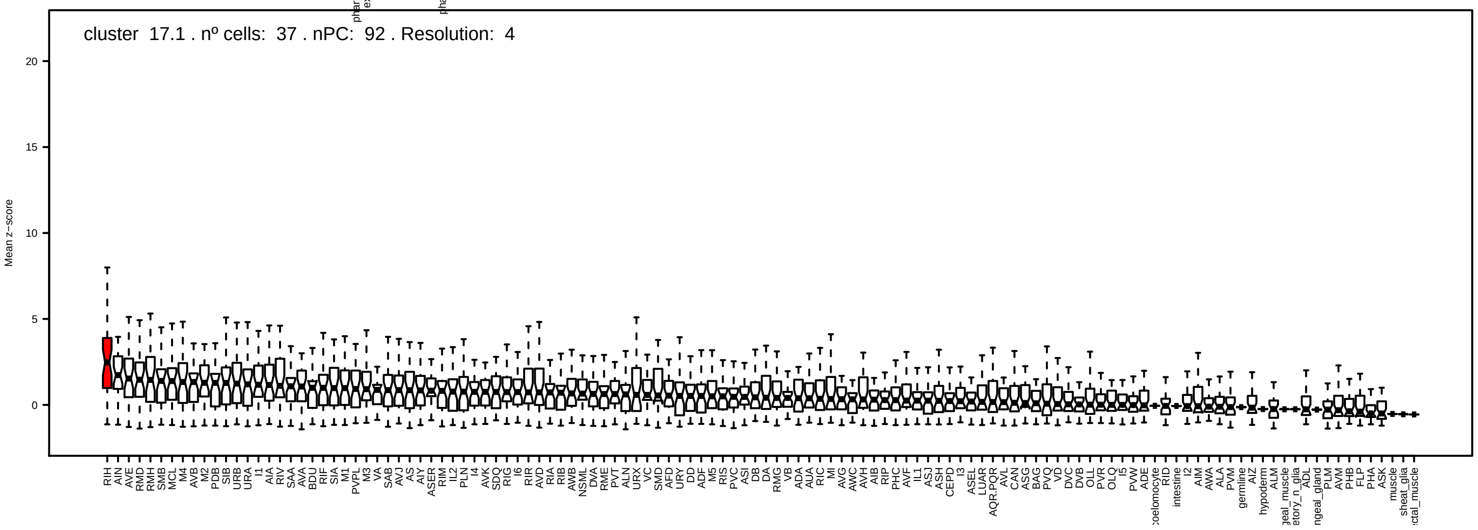
cluster 17.1 . nº cells: 37 . nPC: 92 . Resolution: 4
Mean z−score

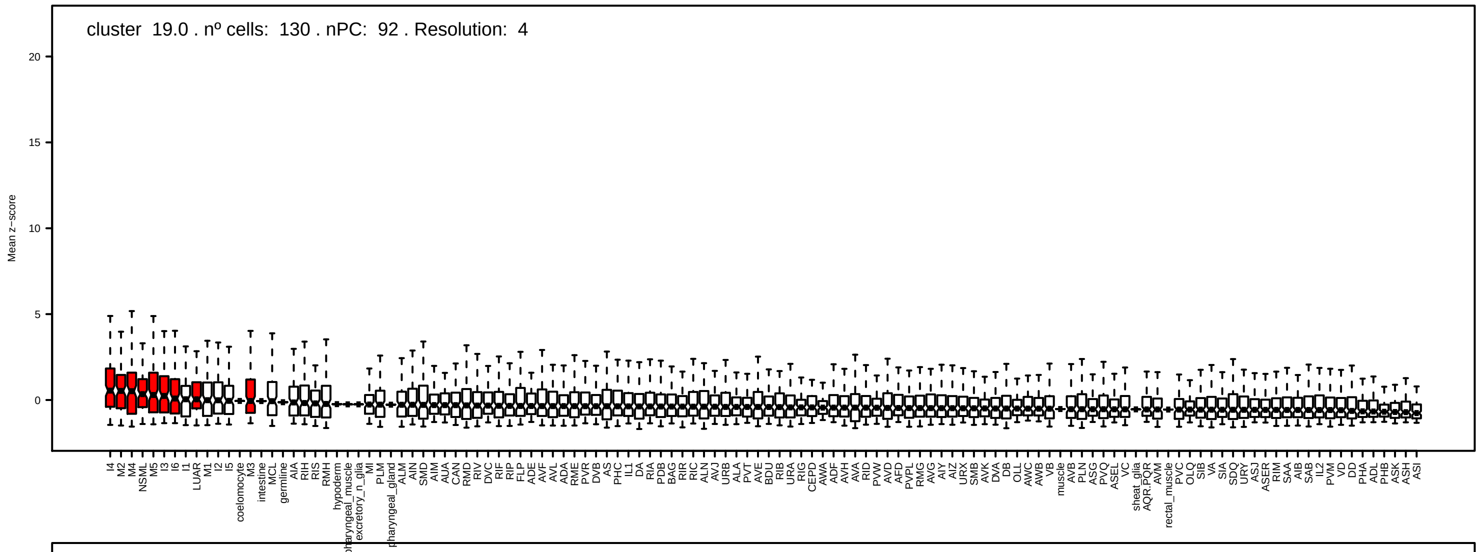
cluster 19.0 . nº cells: 130 . nPC: 92 . Resolution: 4
Mean z−score

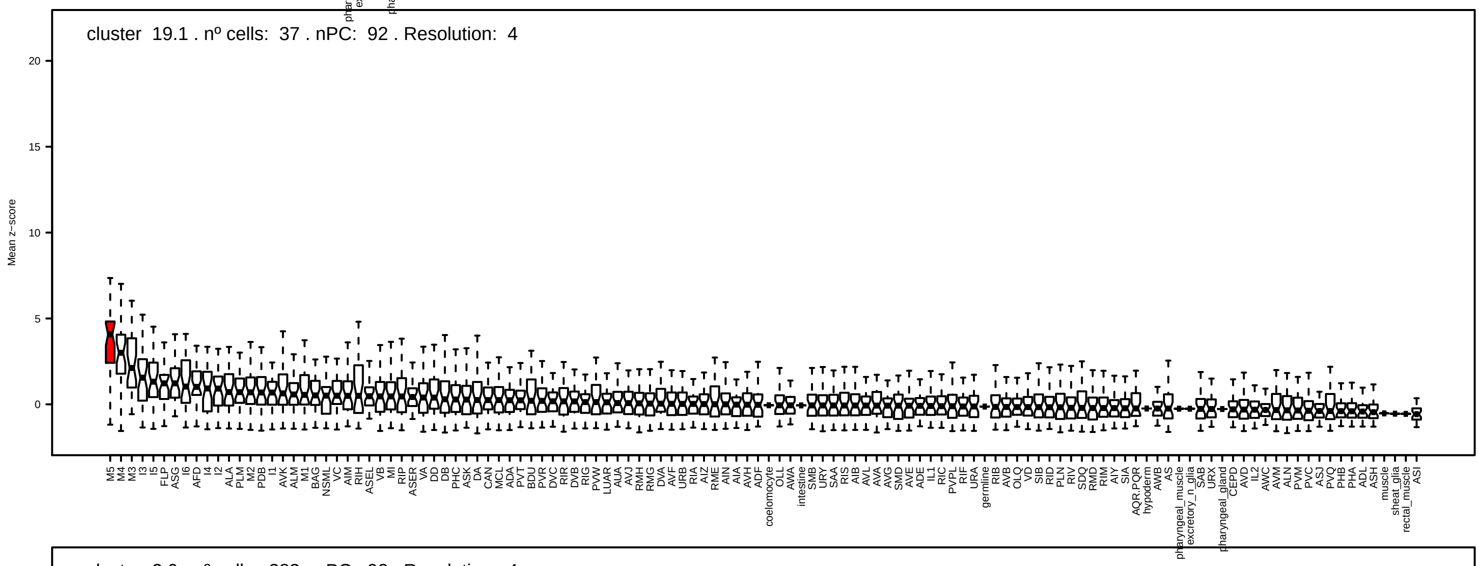
cluster 19.1 . nº cells: 37 . nPC: 92 . Resolution: 4
Mean z−score

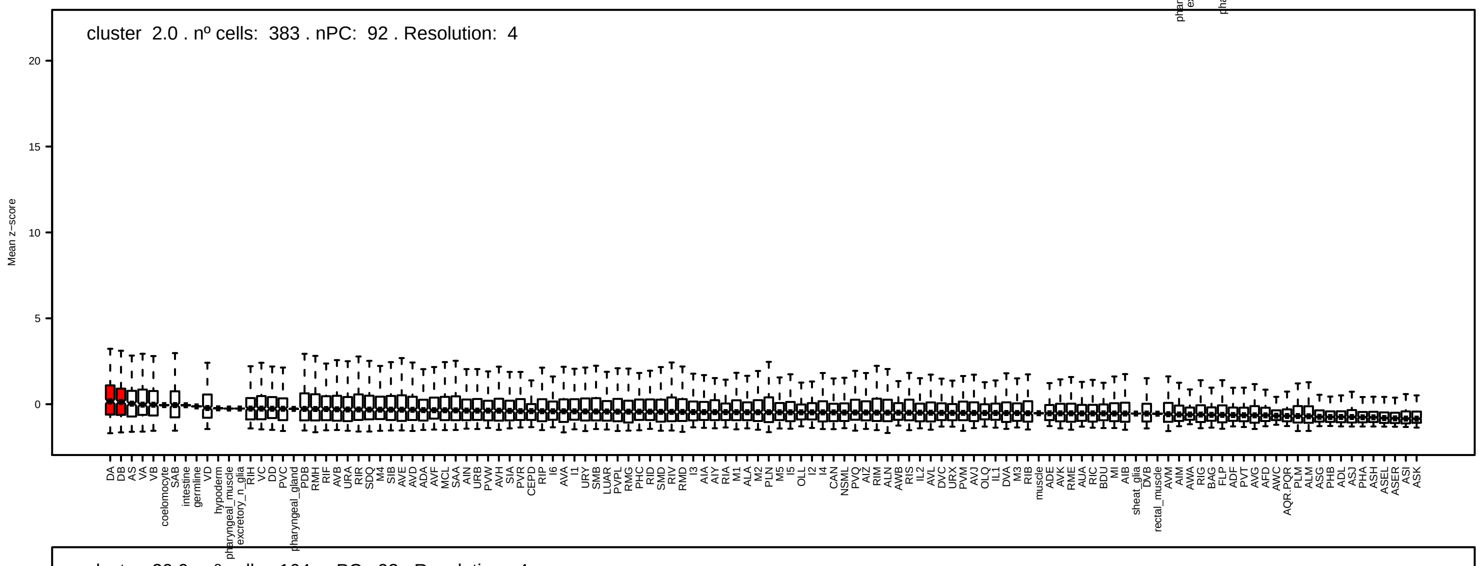
cluster 2.0 . nº cells: 383 . nPC: 92 . Resolution: 4
Mean z−score

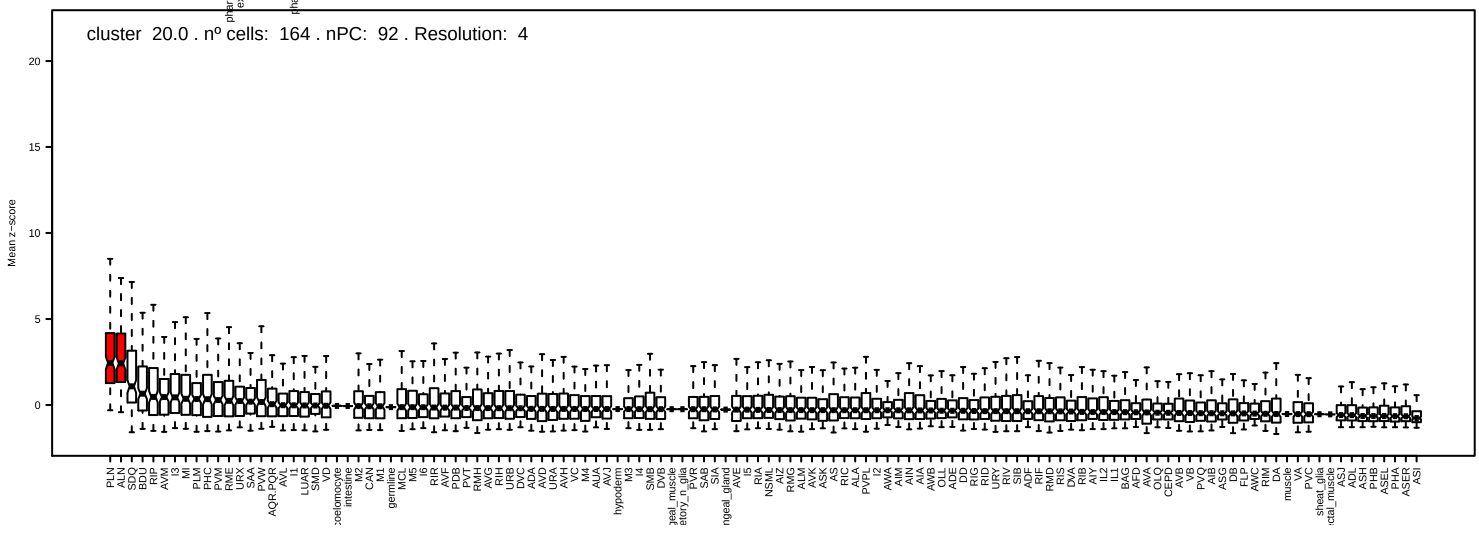
cluster 20.0 . nº cells: 164 . nPC: 92 . Resolution: 4
Mean z−score

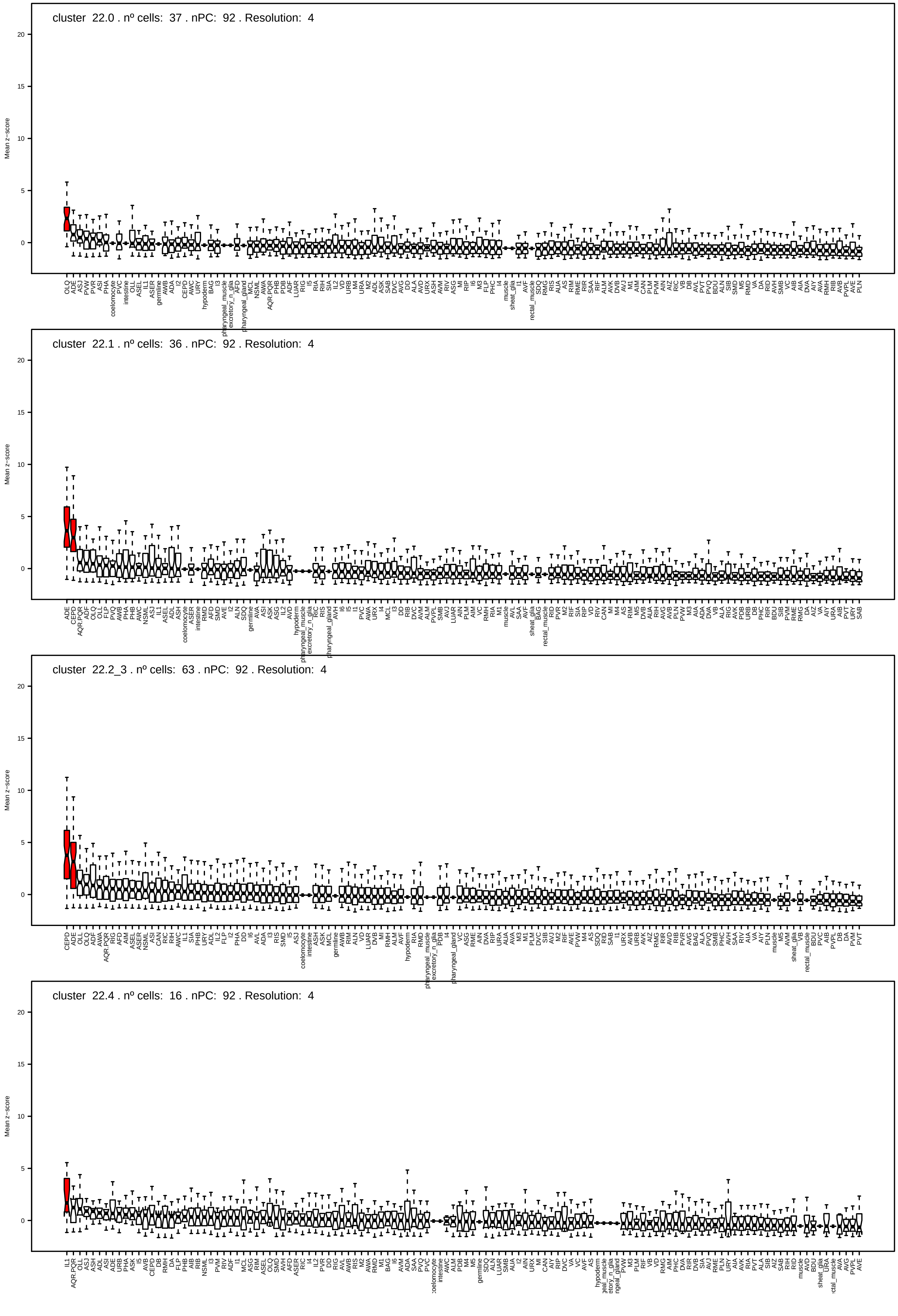
cluster 22.0 . nº cells: 37 . nPC: 92 . Resolution: 4
cluster 22.1 . nº cells: 36 . nPC: 92 . Resolution: 4
cluster 22.2_3 . nº cells: 63 . nPC: 92 . Resolution: 4
cluster 22.4 . nº cells: 16 . nPC: 92 . Resolution: 4
Mean z−score

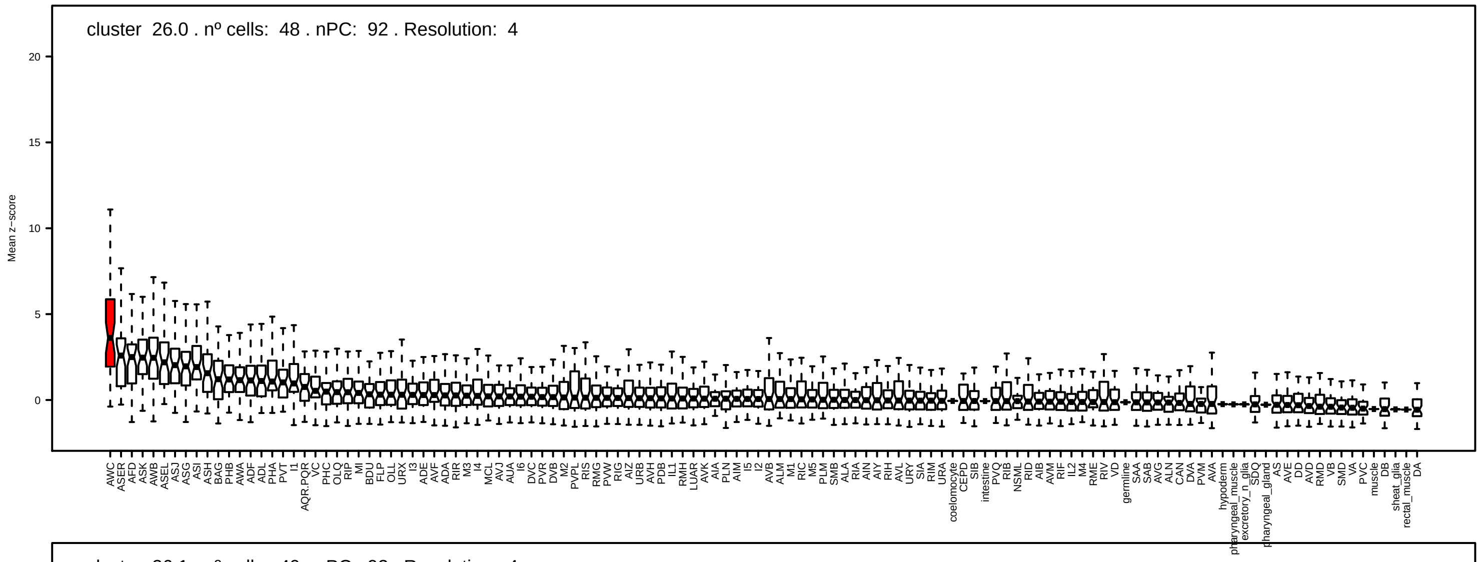
cluster 26.0 . nº cells: 48 . nPC: 92 . Resolution: 4
Mean z−score
20
15
10
5
0

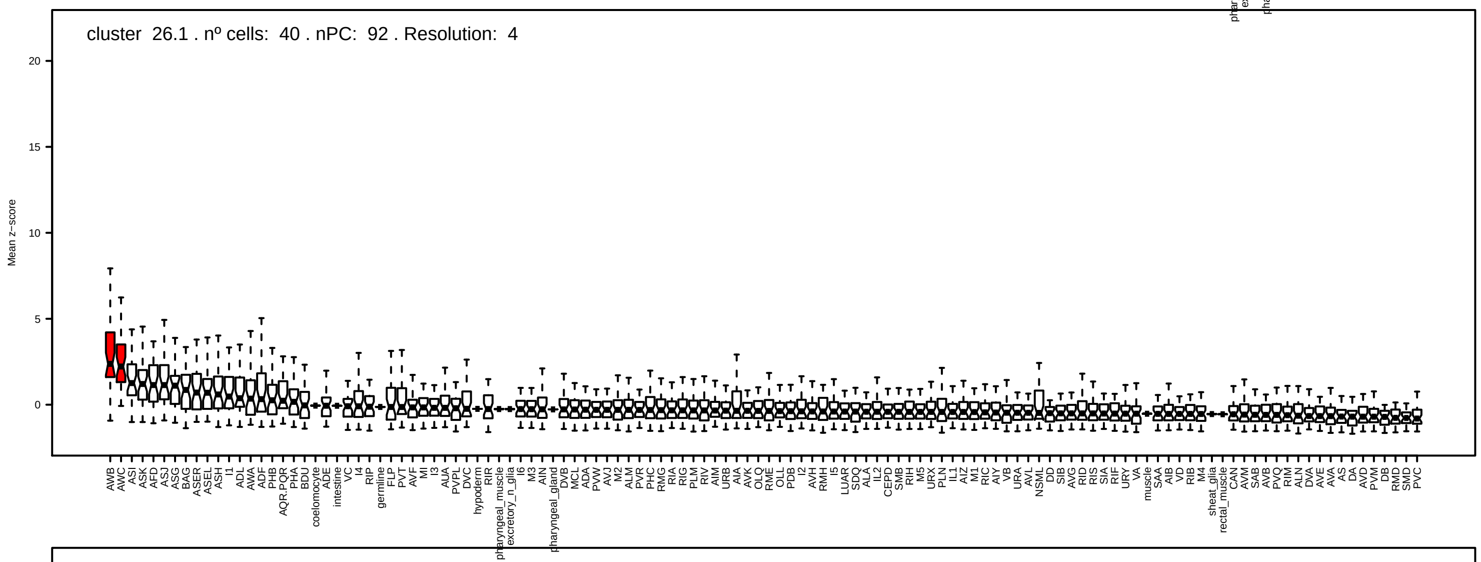
cluster 26.1 . nº cells: 40 . nPC: 92 . Resolution: 4
Mean z−score
20
15
10
5
0

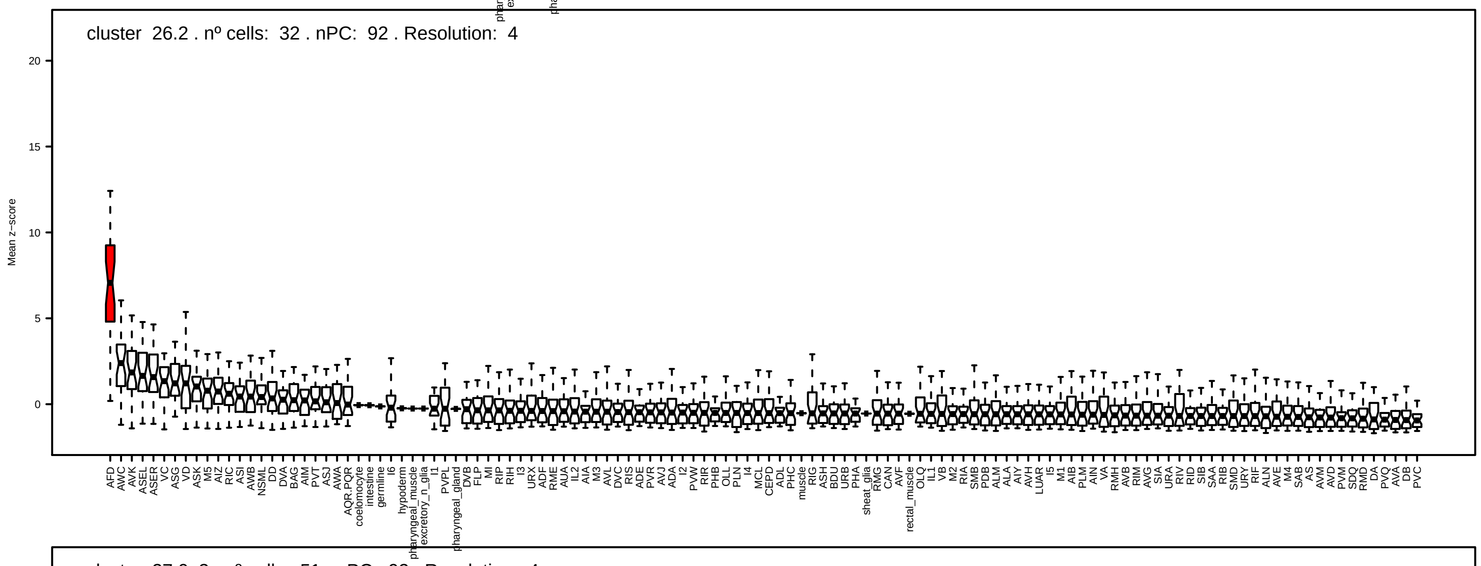
cluster 26.2 . nº cells: 32 . nPC: 92 . Resolution: 4
Mean z−score
20
15
10
5
0

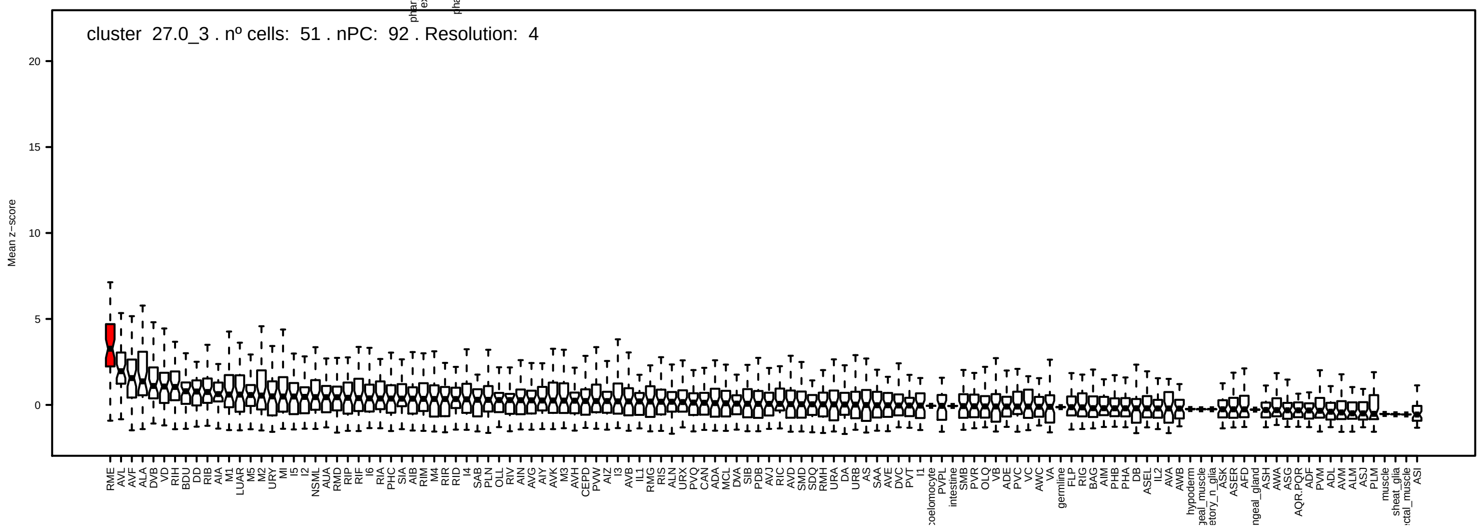
cluster 27.0_3 . nº cells: 51 . nPC: 92 . Resolution: 4
Mean z−score
20
15
10
5
0

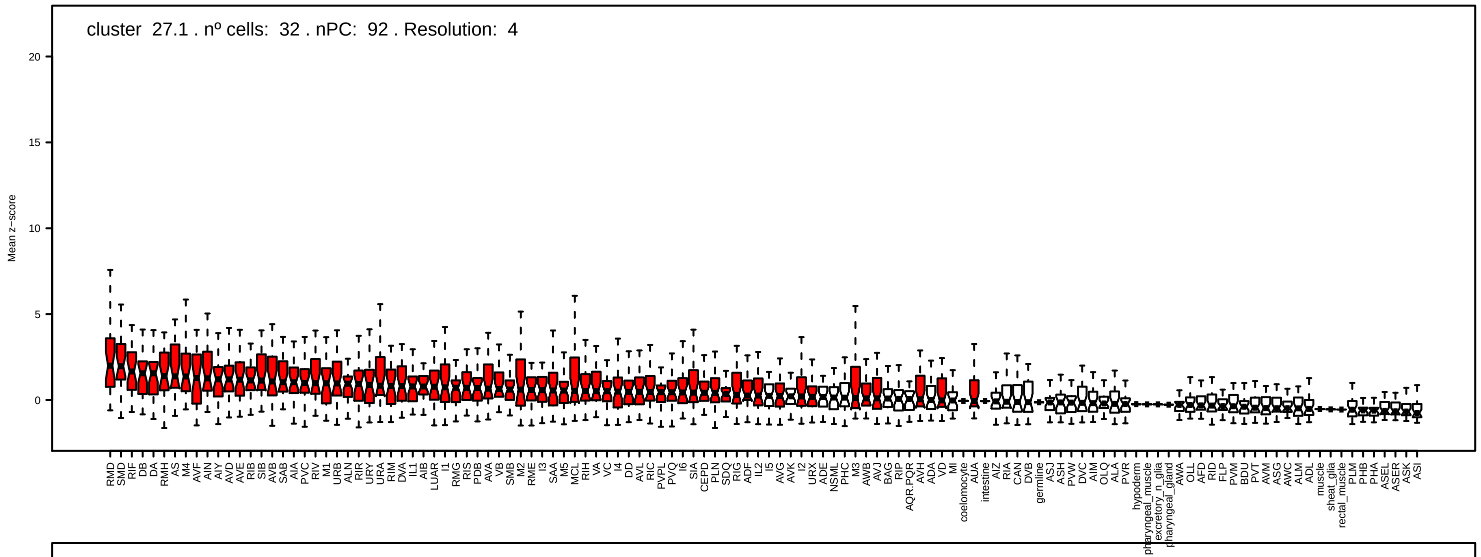
cluster 27.1 . nº cells: 32 . nPC: 92 . Resolution: 4
Mean z−score

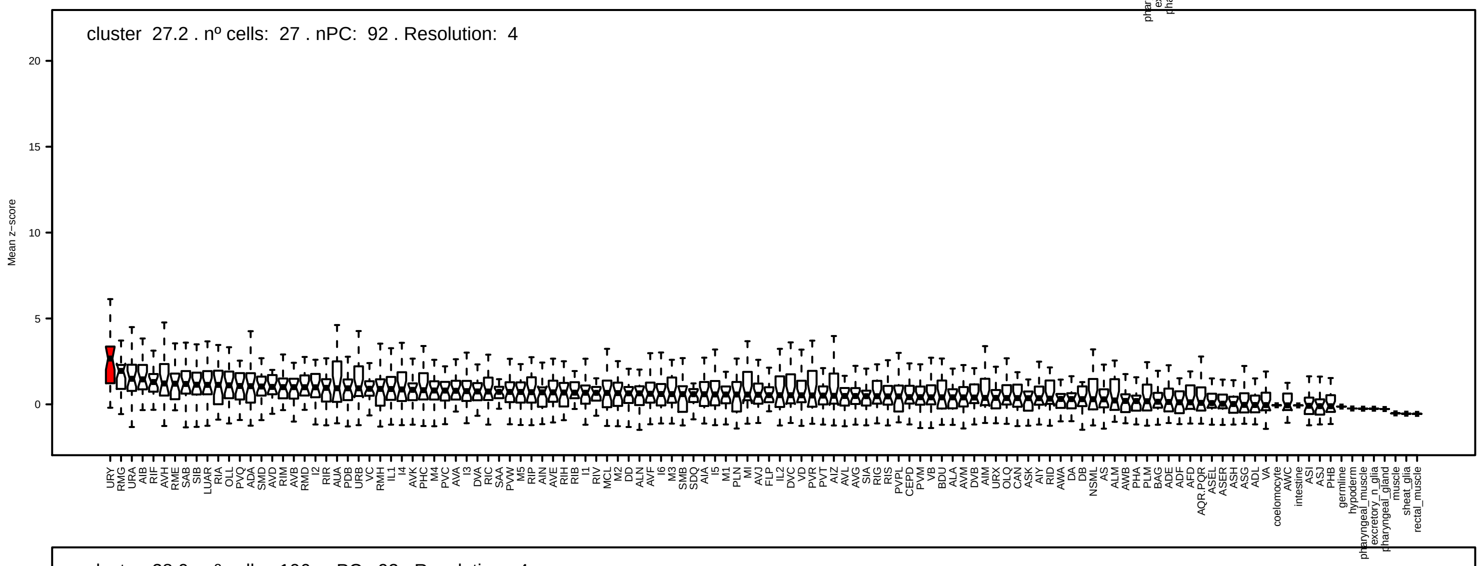
cluster 27.2 . nº cells: 27 . nPC: 92 . Resolution: 4
Mean z−score

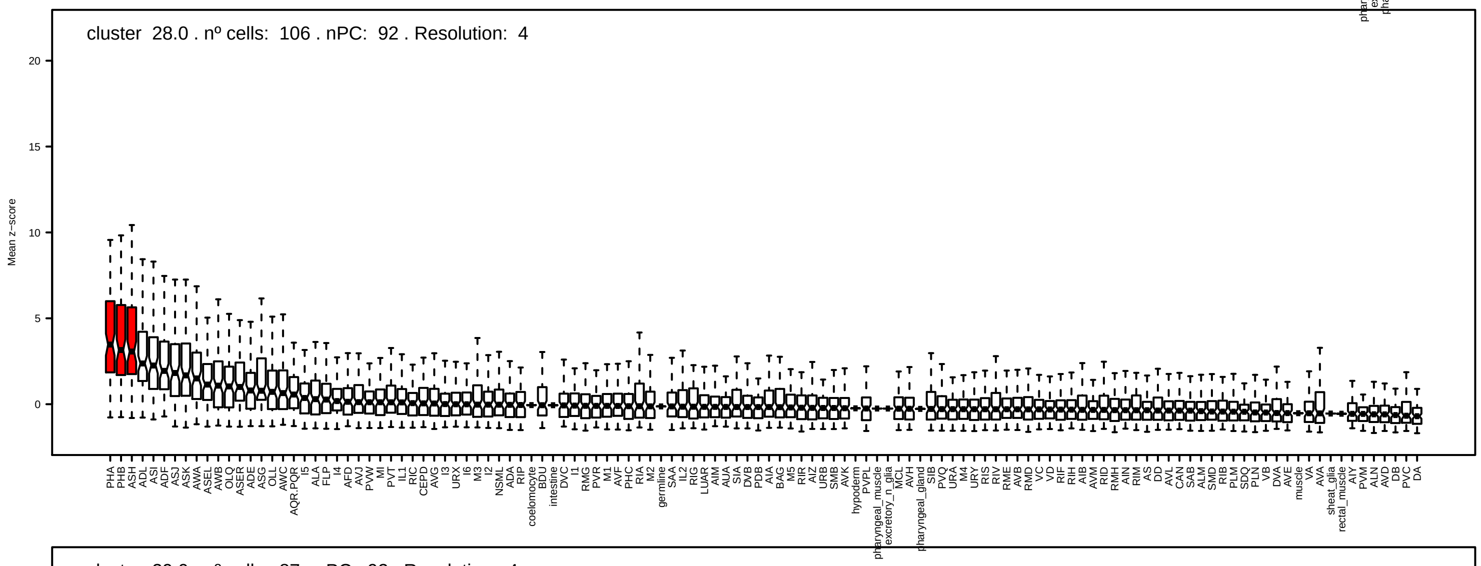
cluster 28.0 . nº cells: 106 . nPC: 92 . Resolution: 4
Mean z−score

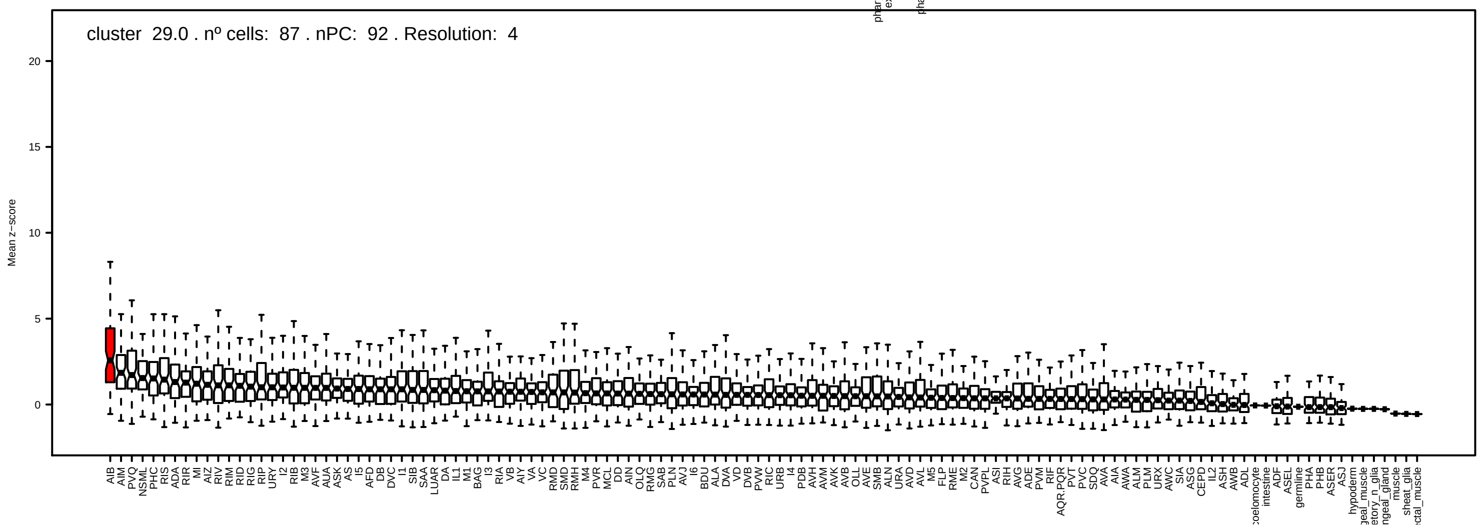
cluster 29.0 . nº cells: 87 . nPC: 92 . Resolution: 4
Mean z−score

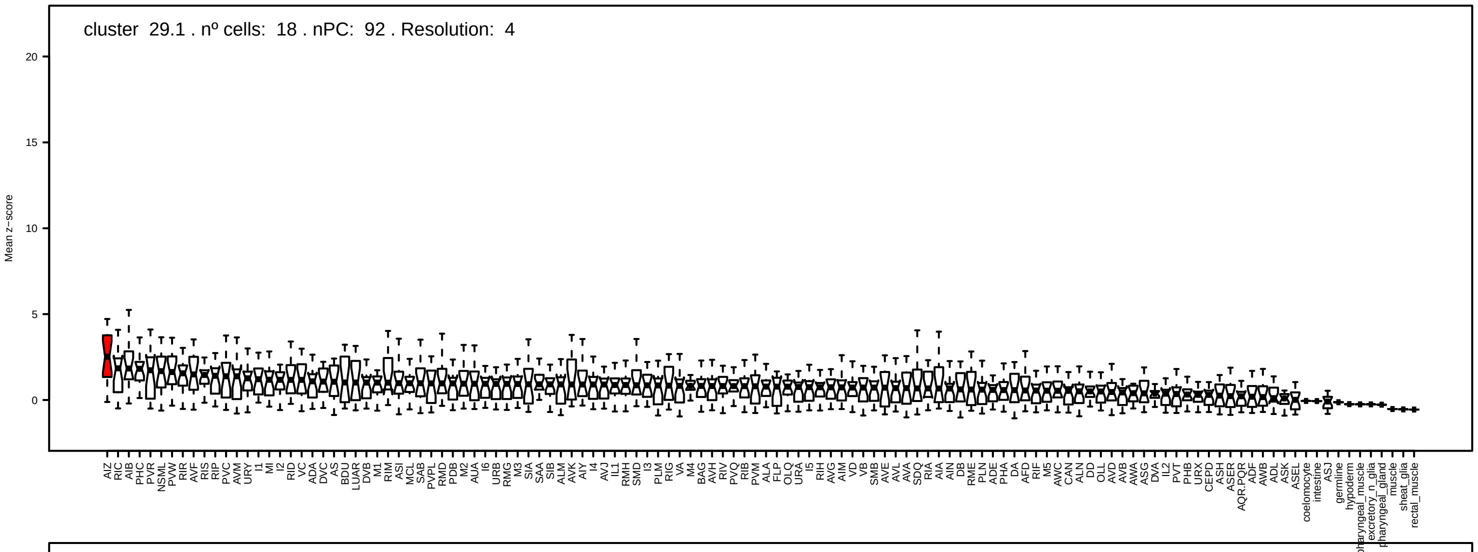

cluster 29.1 . nº cells: 18 . nPC: 92 . Resolution: 4
Mean z−score


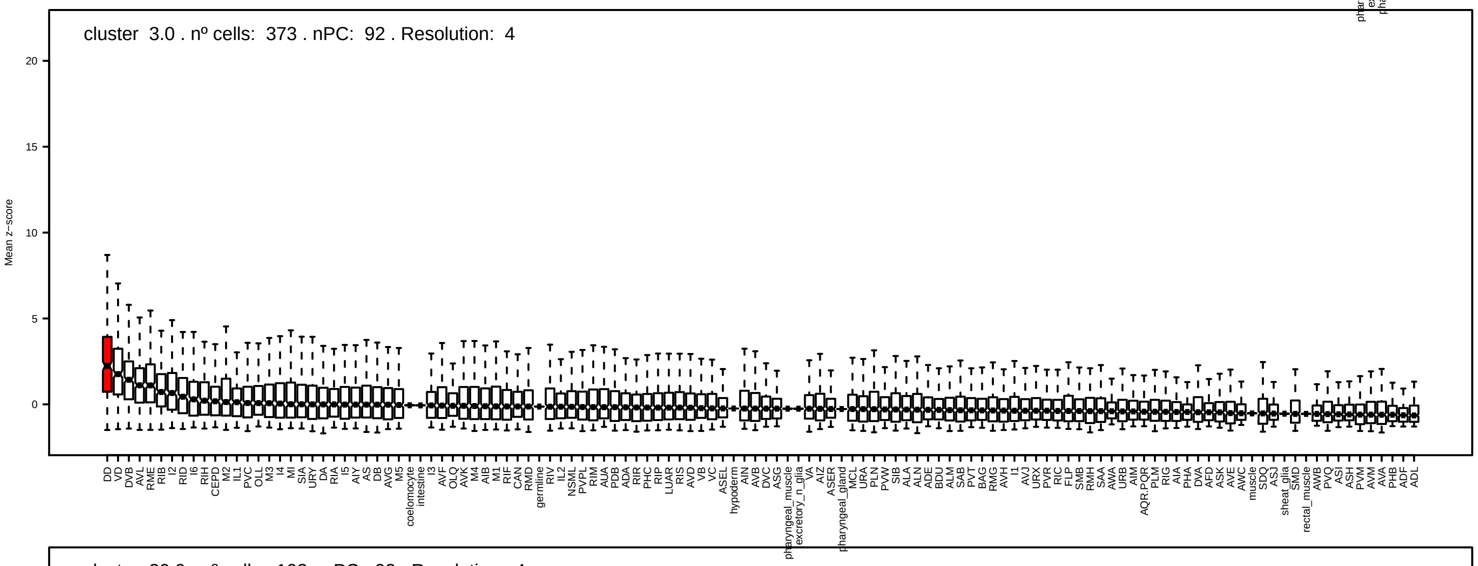

cluster 3.0 . nº cells: 373 . nPC: 92 . Resolution: 4
Mean z−score


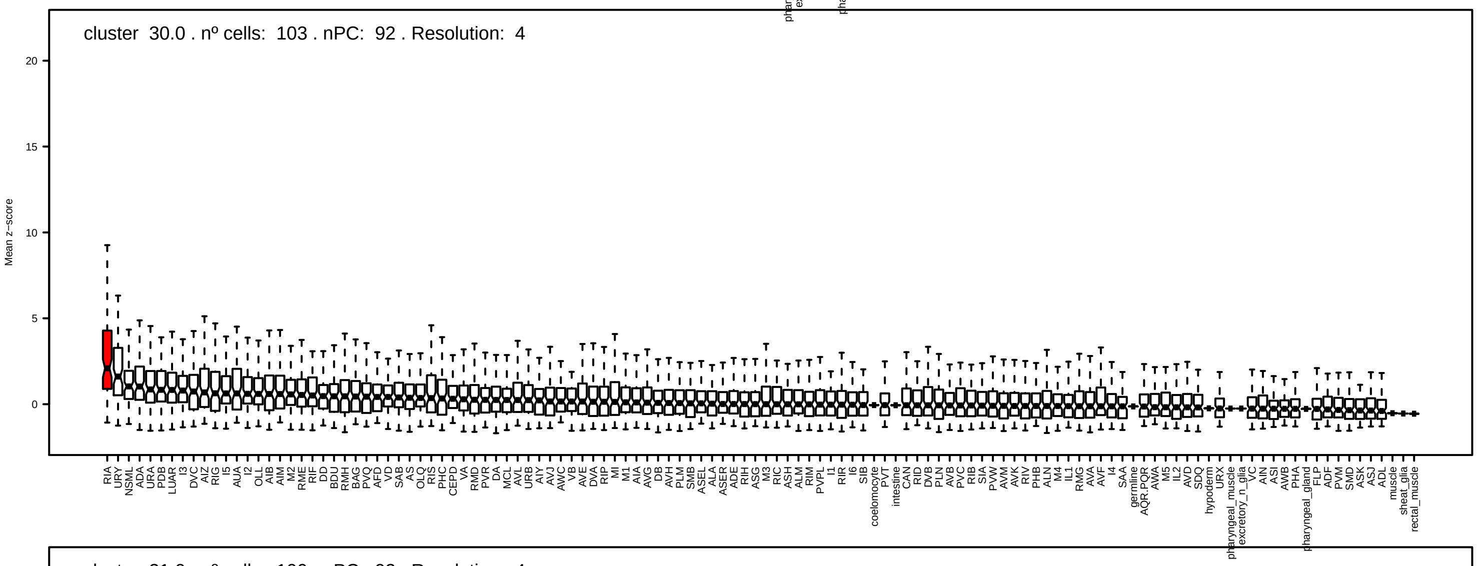

cluster 30.0 . nº cells: 103 . nPC: 92 . Resolution: 4
Mean z−score


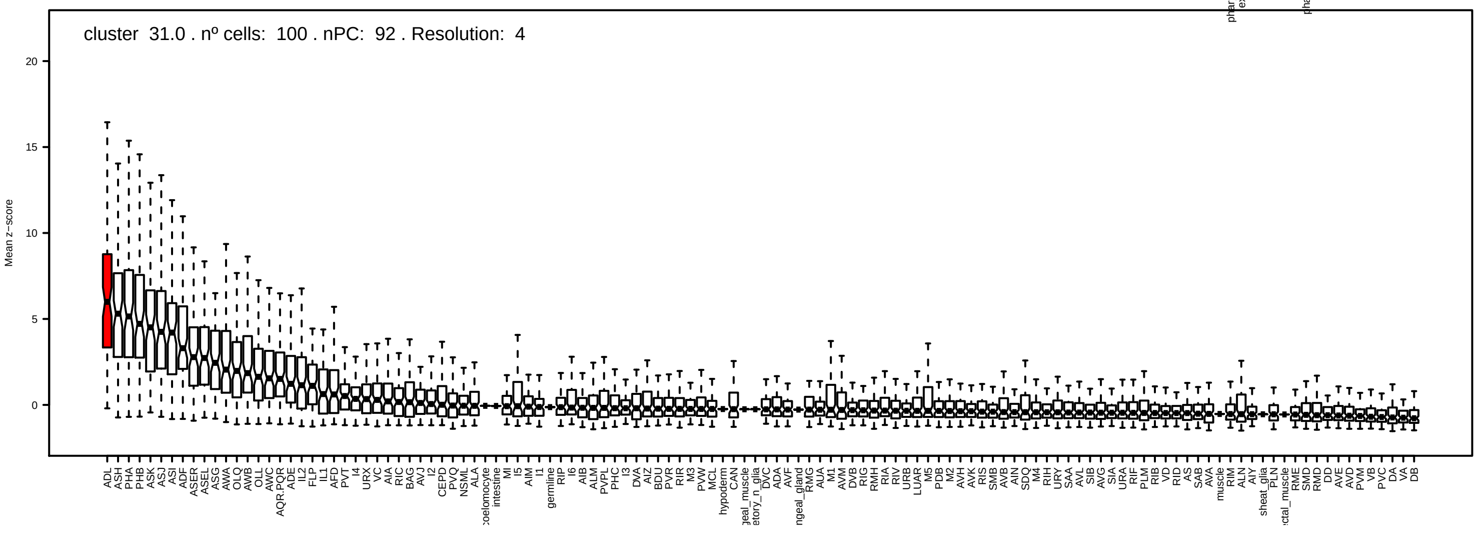

cluster 31.0 . nº cells: 100 . nPC: 92 . Resolution: 4
Mean z−score

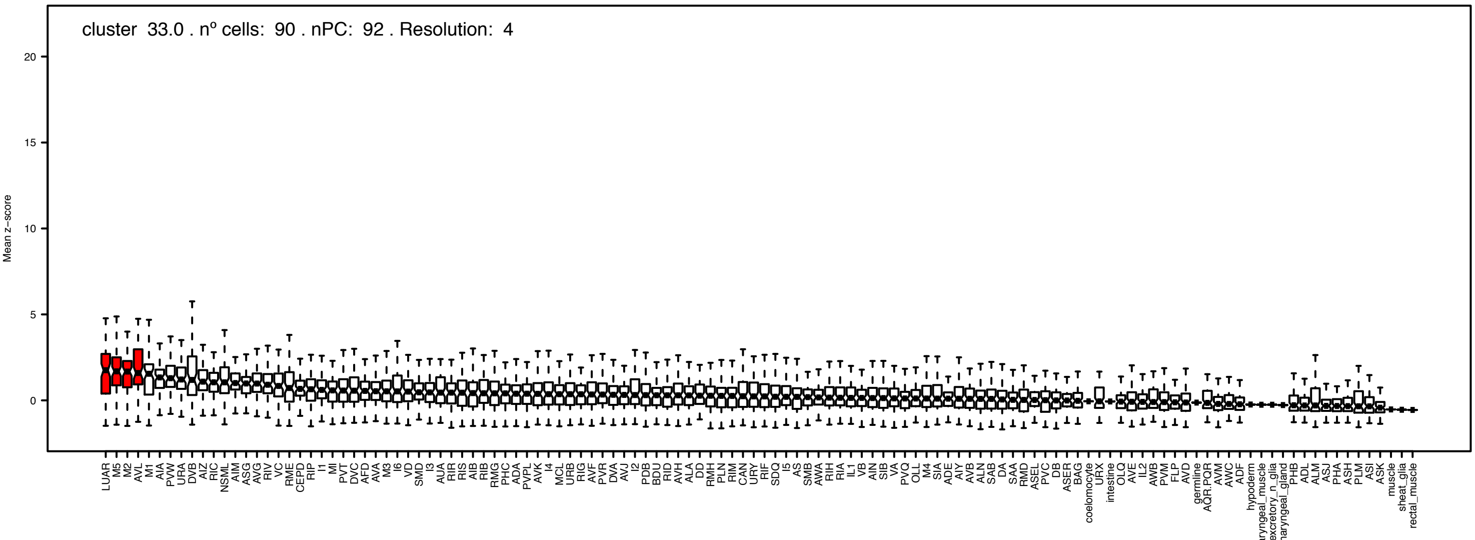
cluster 33.0 . nº cells: 90 . nPC: 92 . Resolution: 4
Mean z−score

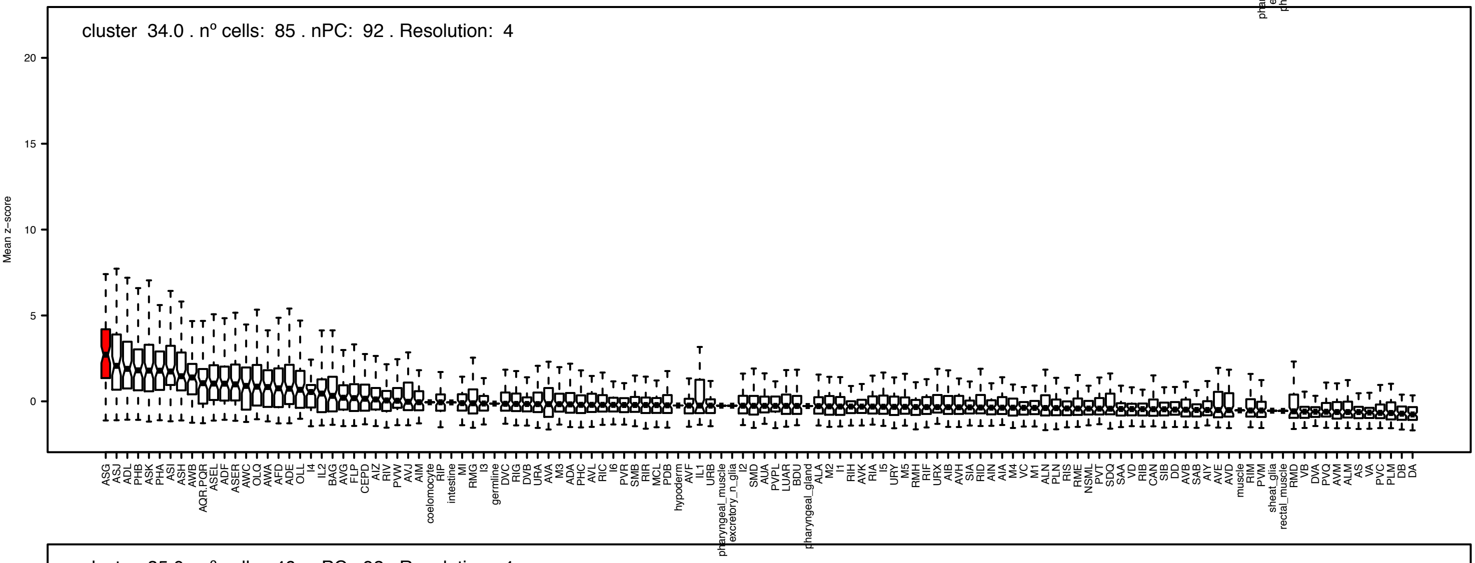
cluster 34.0 . nº cells: 85 . nPC: 92 . Resolution: 4
Mean z−score

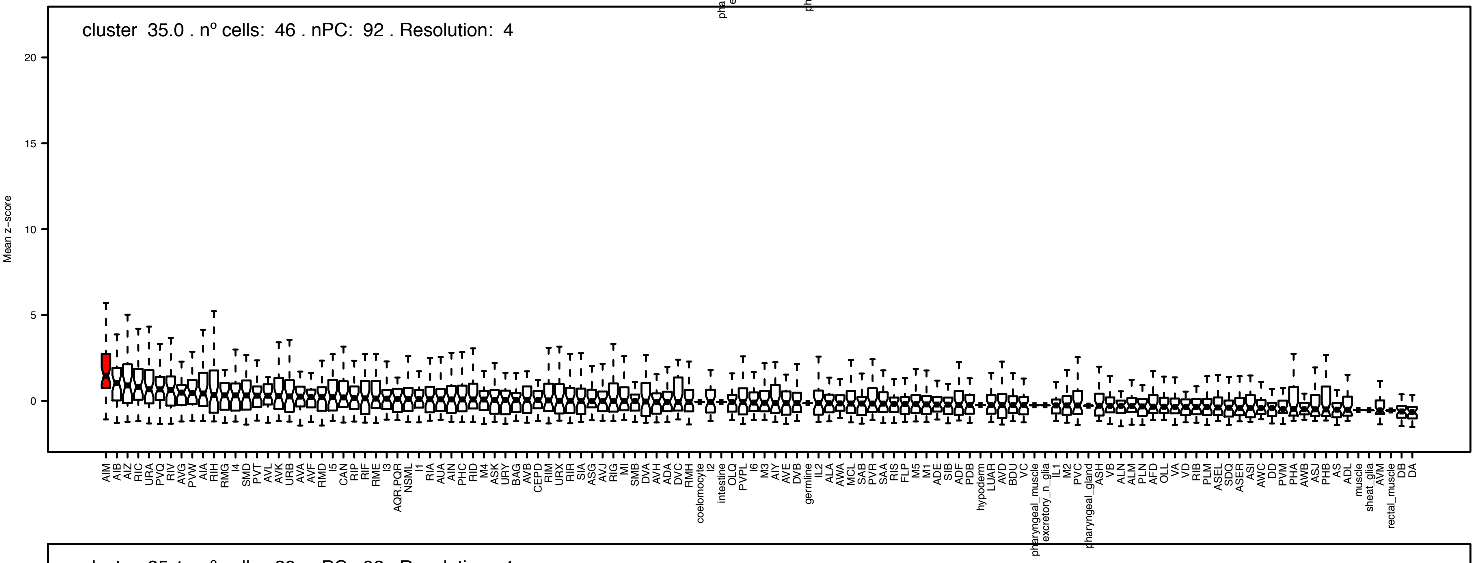
cluster 35.0 . nº cells: 46 . nPC: 92 . Resolution: 4
Mean z−score

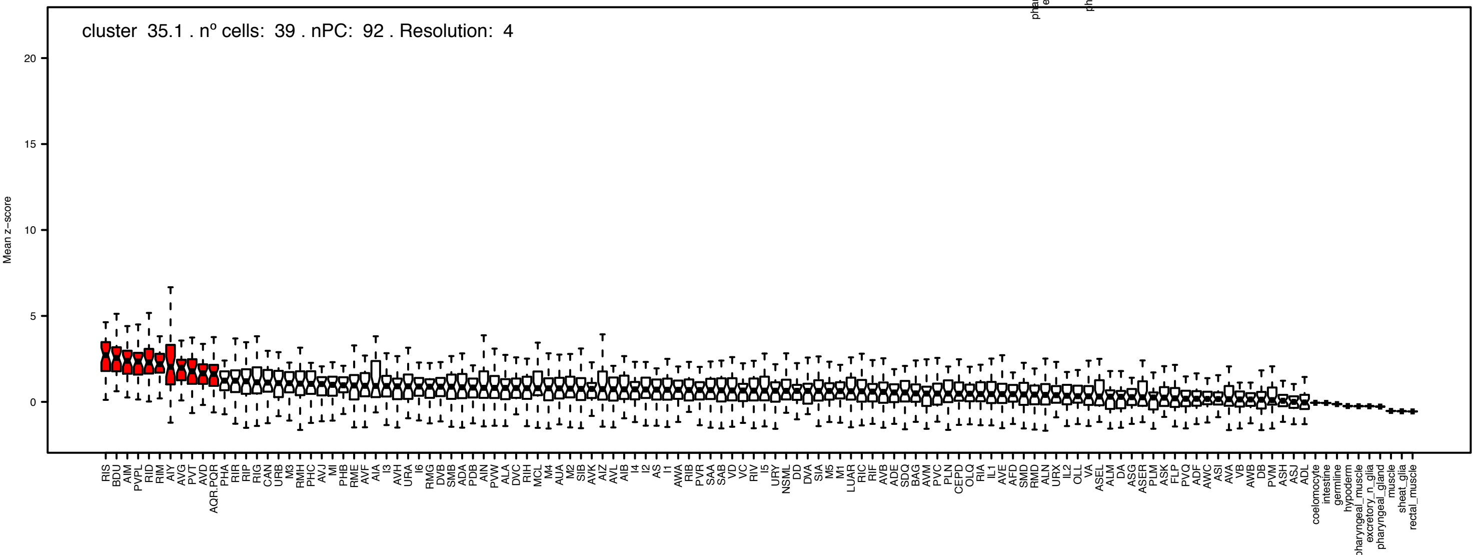
cluster 35.1 . nº cells: 39 . nPC: 92 . Resolution: 4
Mean z−score

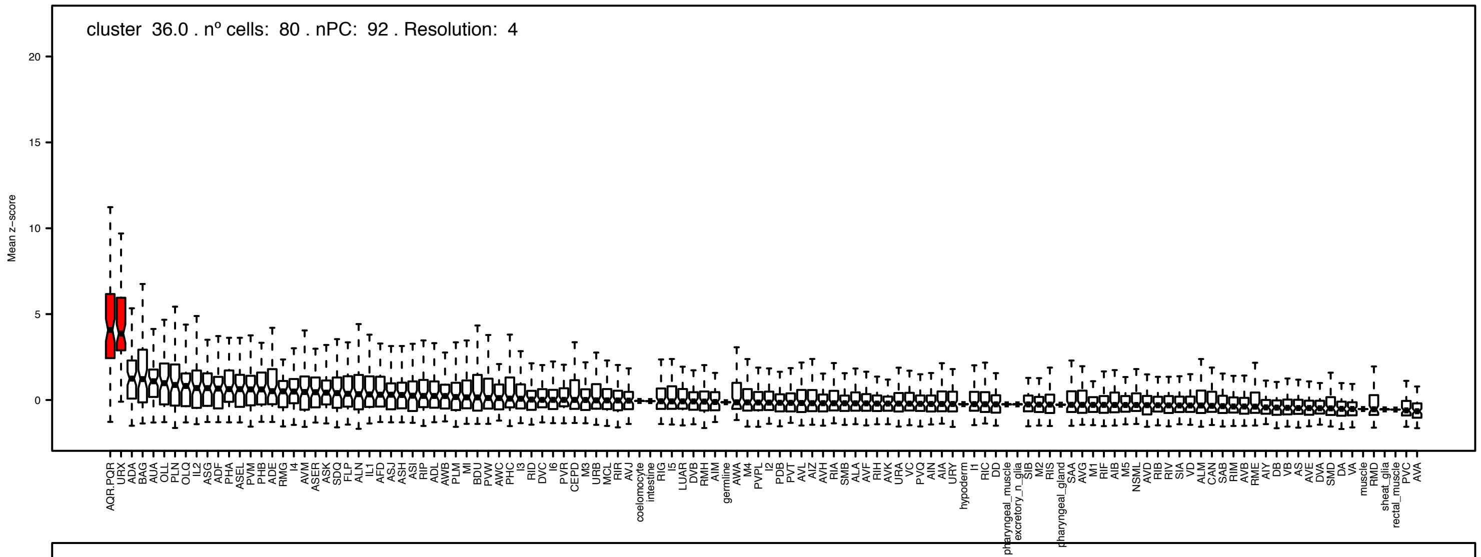
cluster 36.0 . nº cells: 80 . nPC: 92 . Resolution: 4
Mean z−score

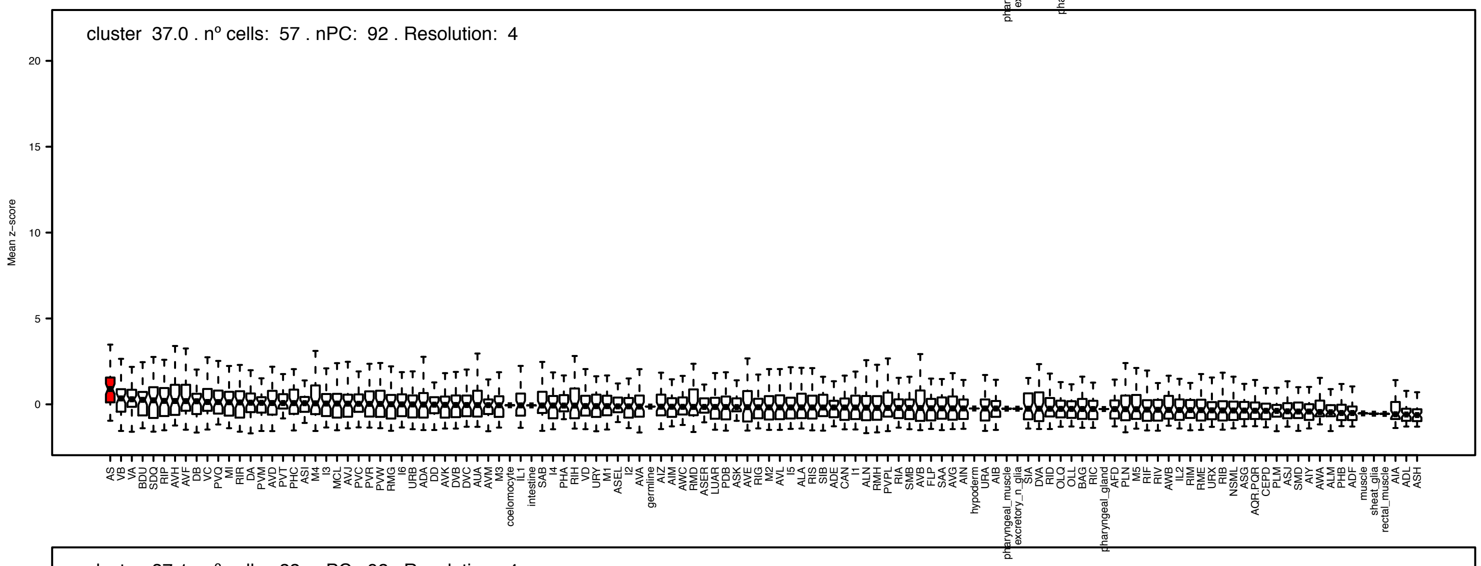
cluster 37.0 . nº cells: 57 . nPC: 92 . Resolution: 4
Mean z−score

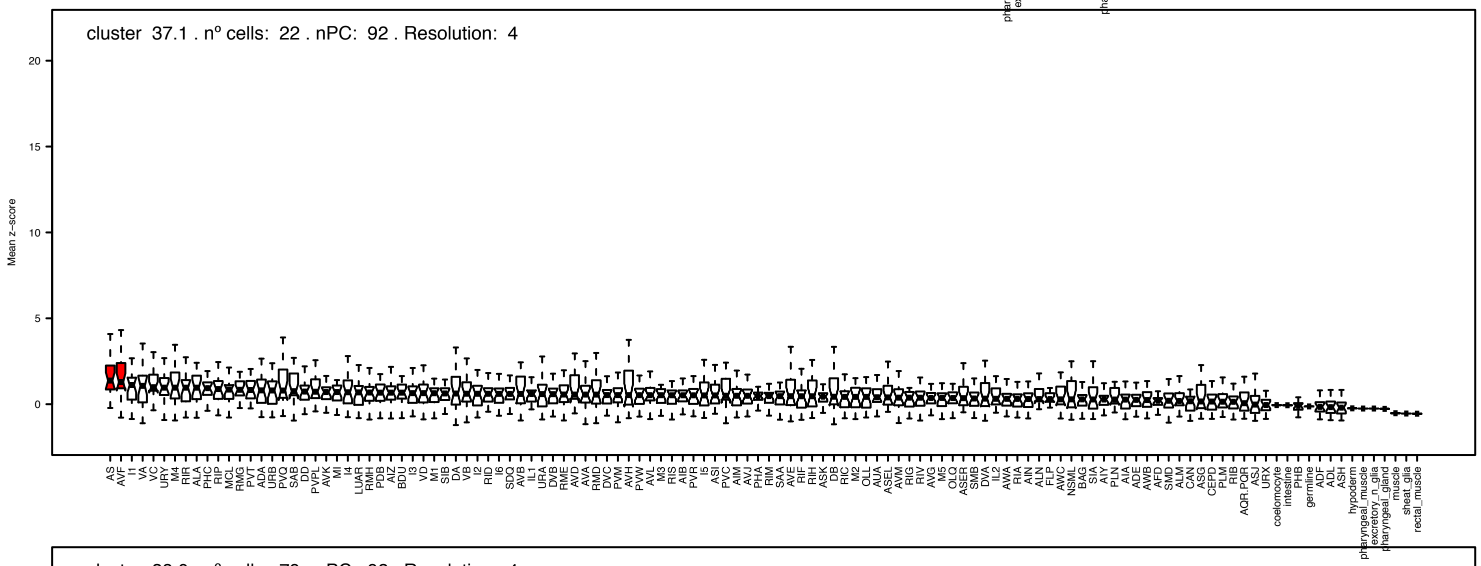
cluster 37.1 . nº cells: 22 . nPC: 92 . Resolution: 4
Mean z−score

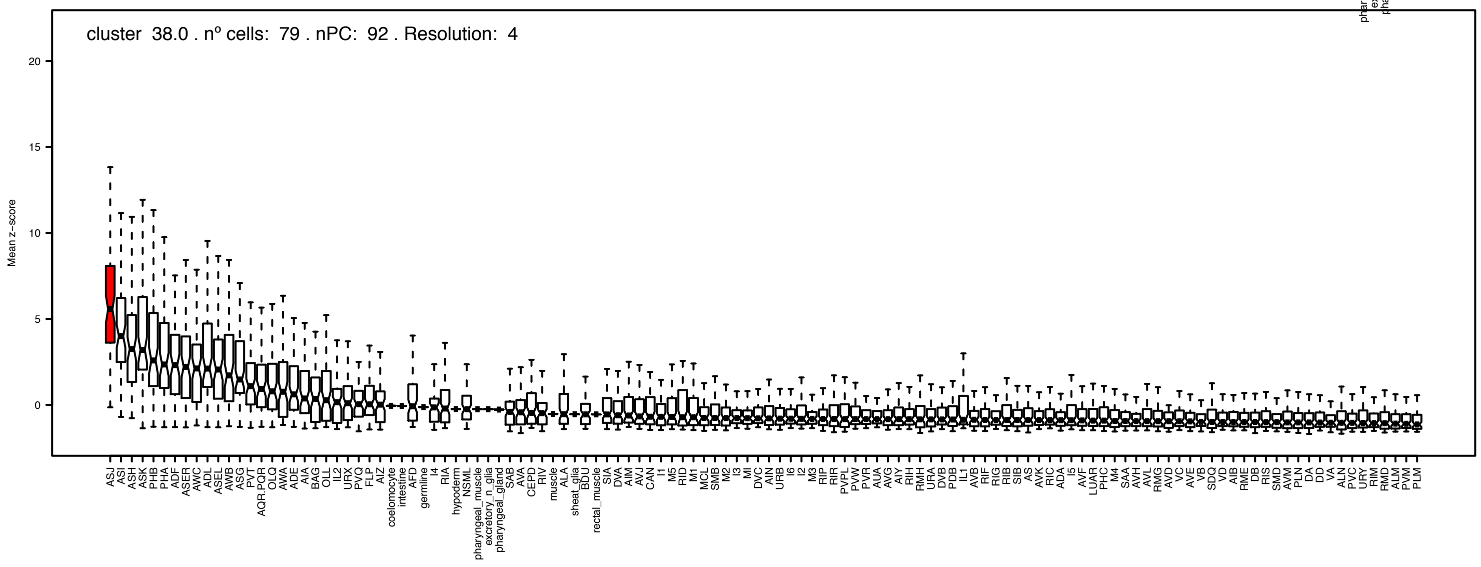
cluster 38.0 . nº cells: 79 . nPC: 92 . Resolution: 4
Mean z−score

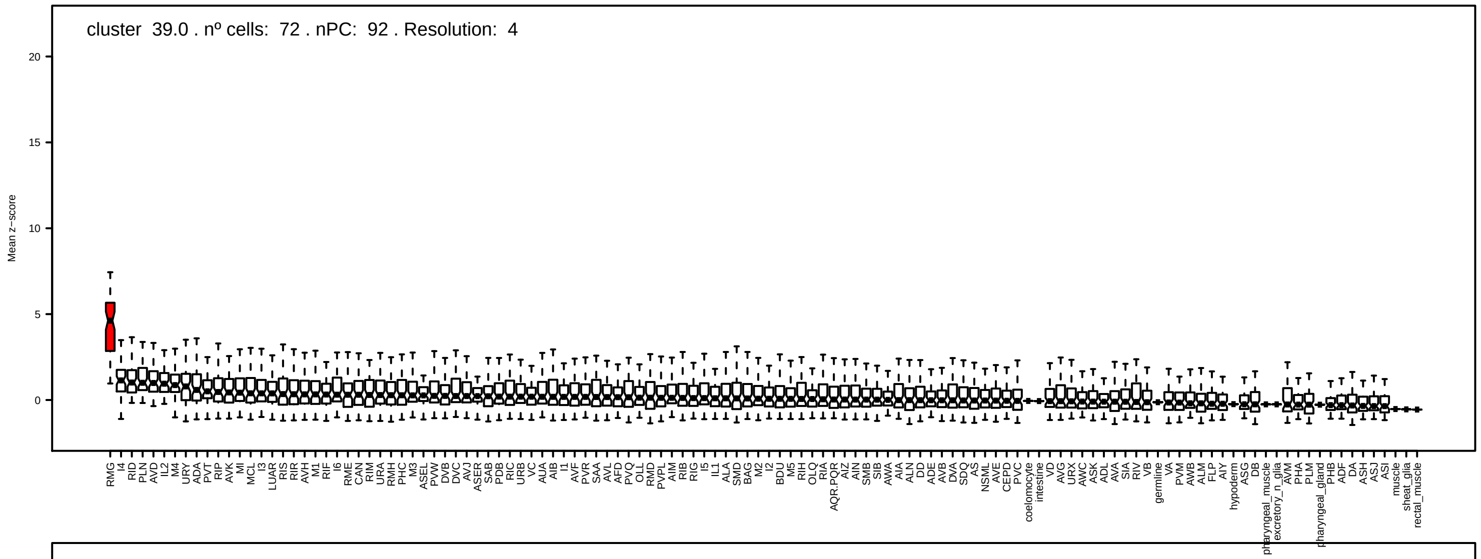
cluster 39.0 . nº cells: 72 . nPC: 92 . Resolution: 4
Mean z−score

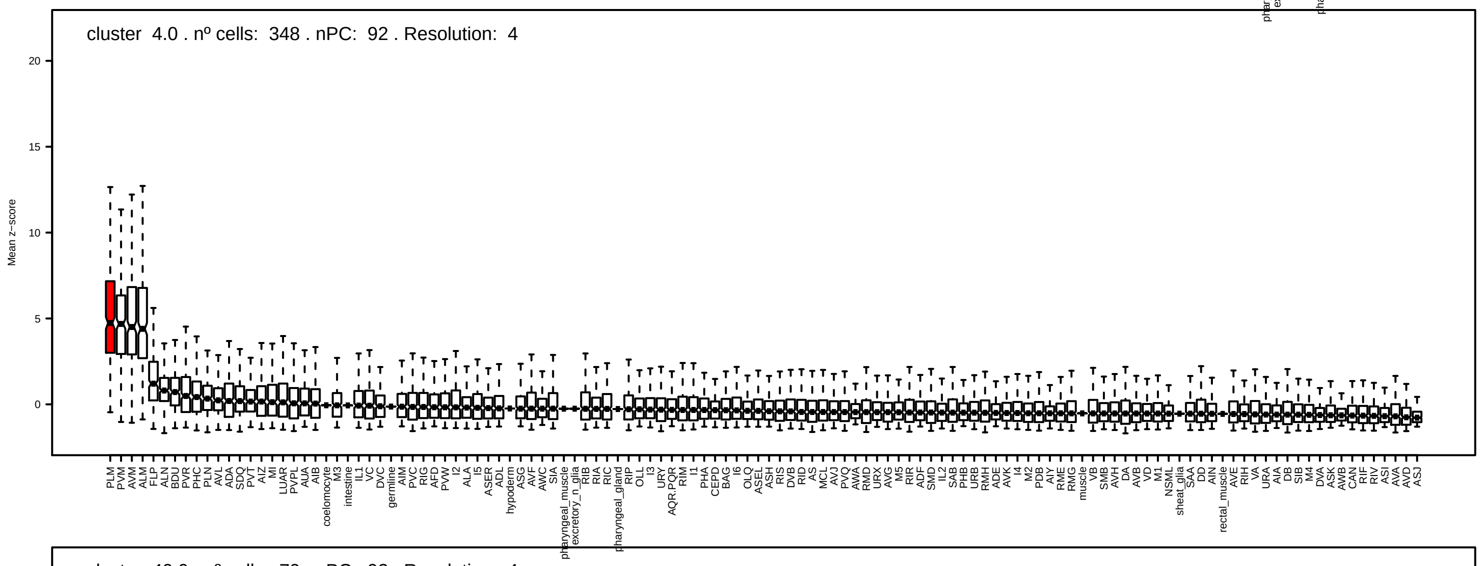
cluster 4.0 . nº cells: 348 . nPC: 92 . Resolution: 4
Mean z−score

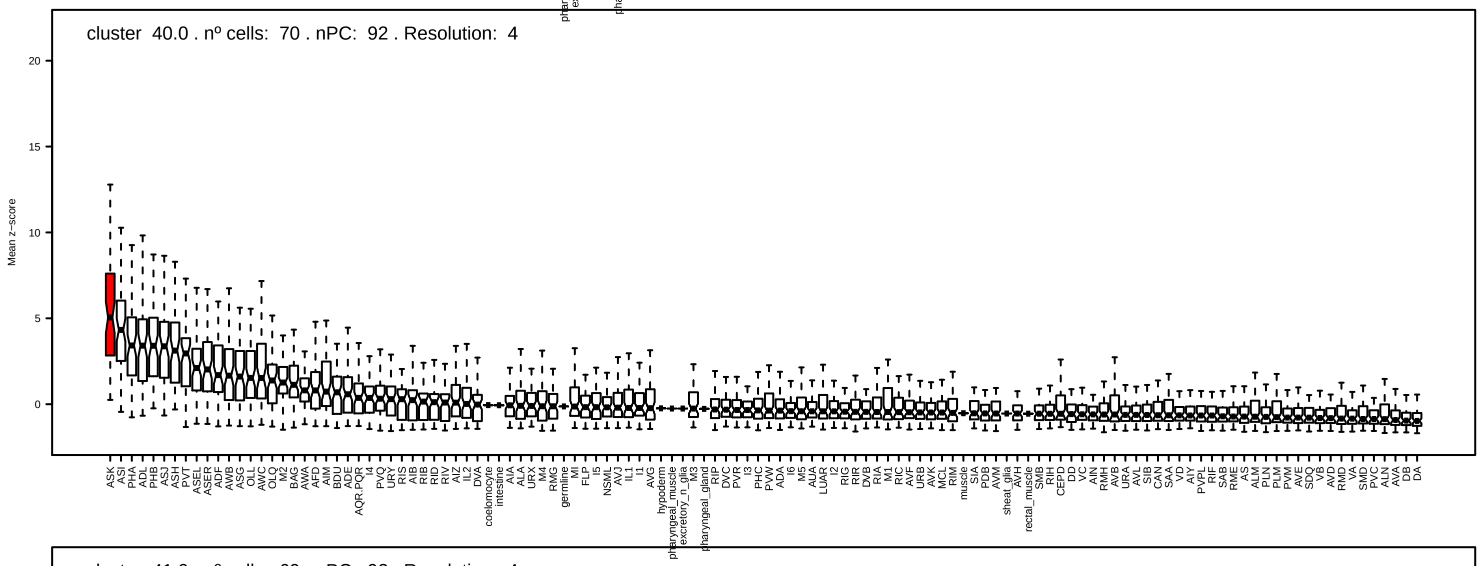
cluster 40.0 . nº cells: 70 . nPC: 92 . Resolution: 4
Mean z−score

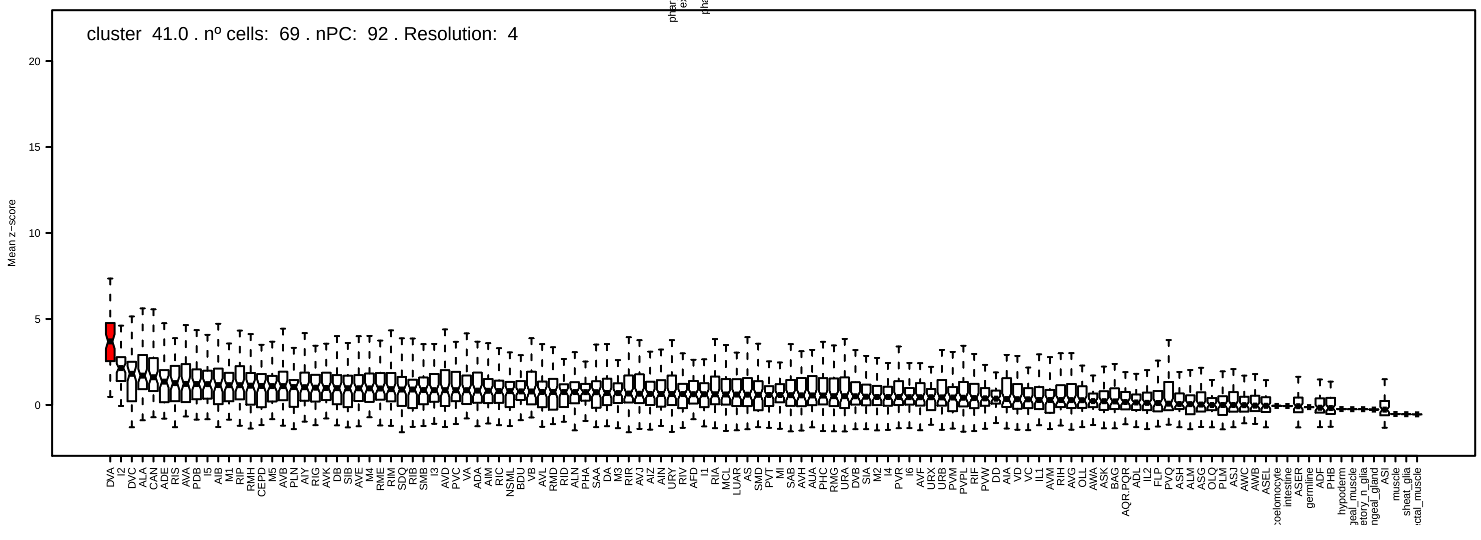
cluster 41.0 . nº cells: 69 . nPC: 92 . Resolution: 4
Mean z−score

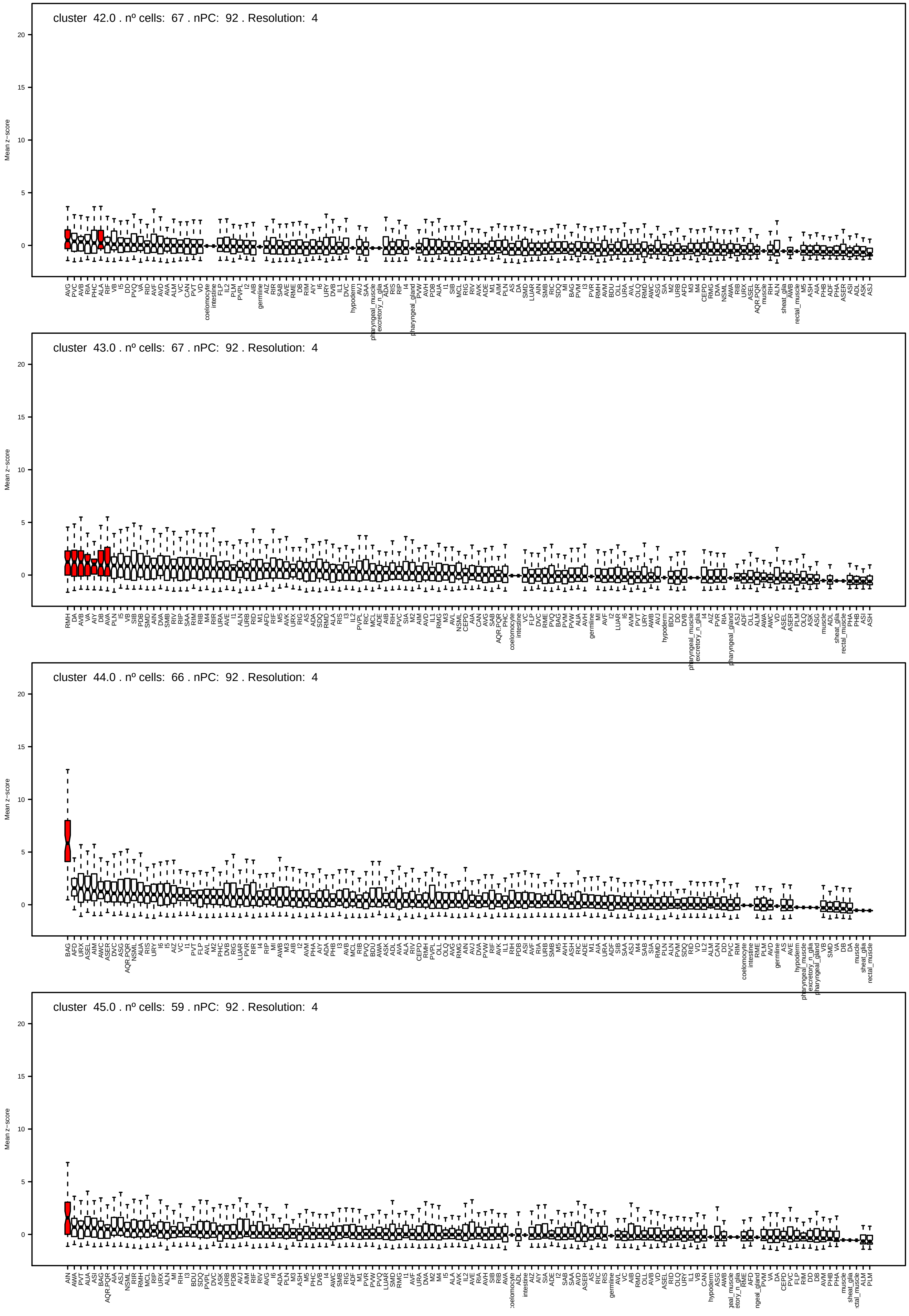

cluster 42.0 . nº cells: 67 . nPC: 92 . Resolution: 4
cluster 43.0 . nº cells: 67 . nPC: 92 . Resolution: 4
cluster 44.0 . nº cells: 66 . nPC: 92 . Resolution: 4
cluster 45.0 . nº cells: 59 . nPC: 92 . Resolution: 4
Mean z−score

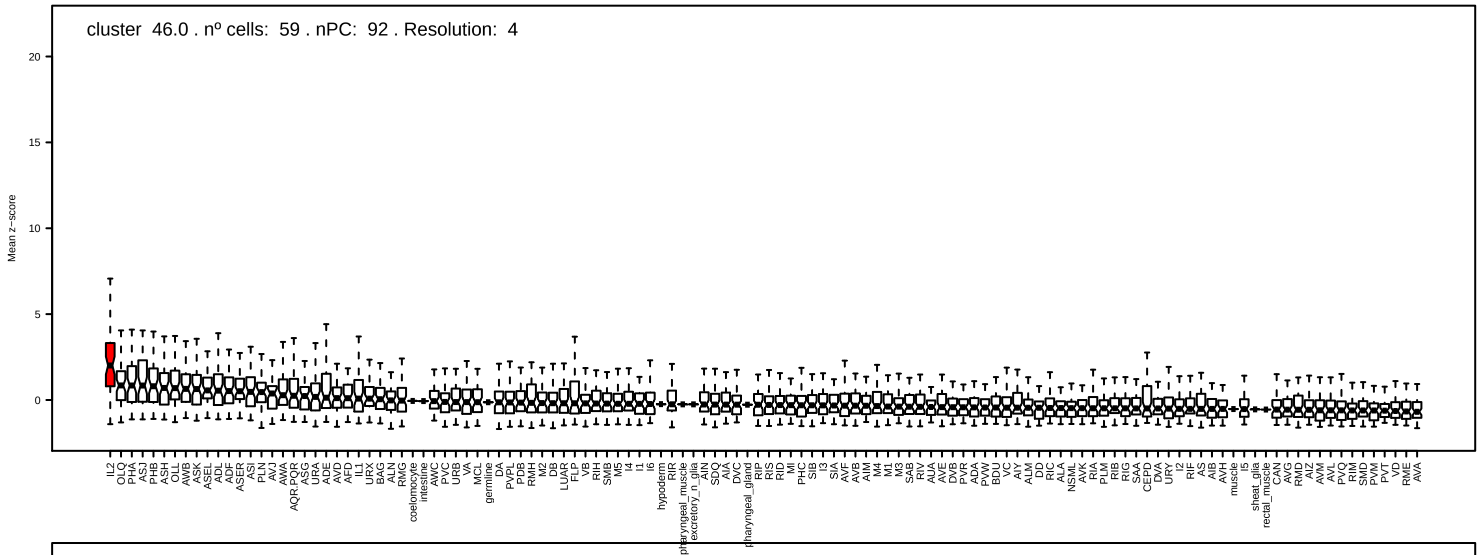
cluster 46.0 . nº cells: 59 . nPC: 92 . Resolution: 4
Mean z−score

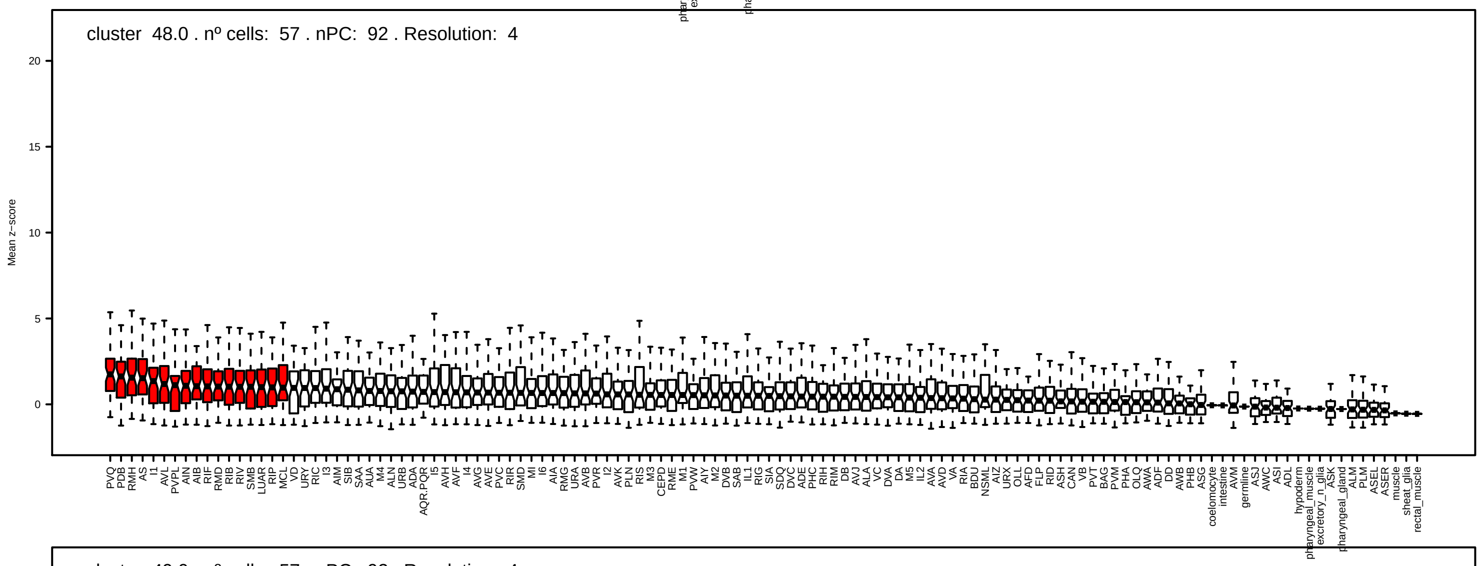
cluster 48.0 . nº cells: 57 . nPC: 92 . Resolution: 4
Mean z−score

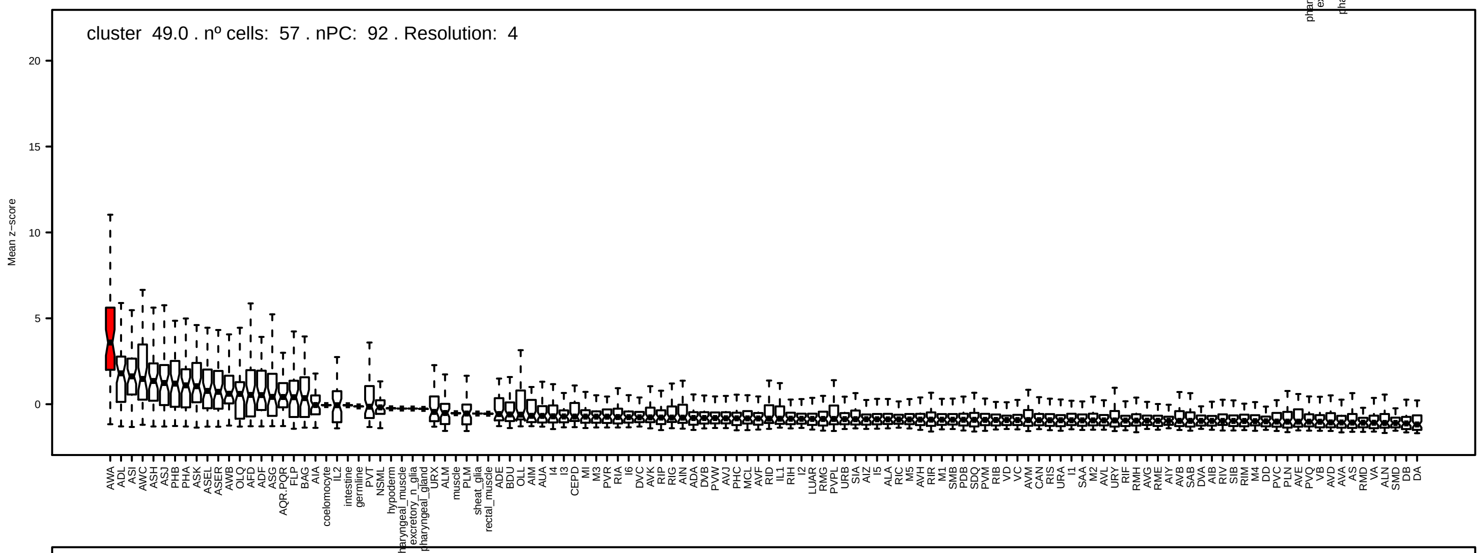
cluster 49.0 . nº cells: 57 . nPC: 92 . Resolution: 4
Mean z−score

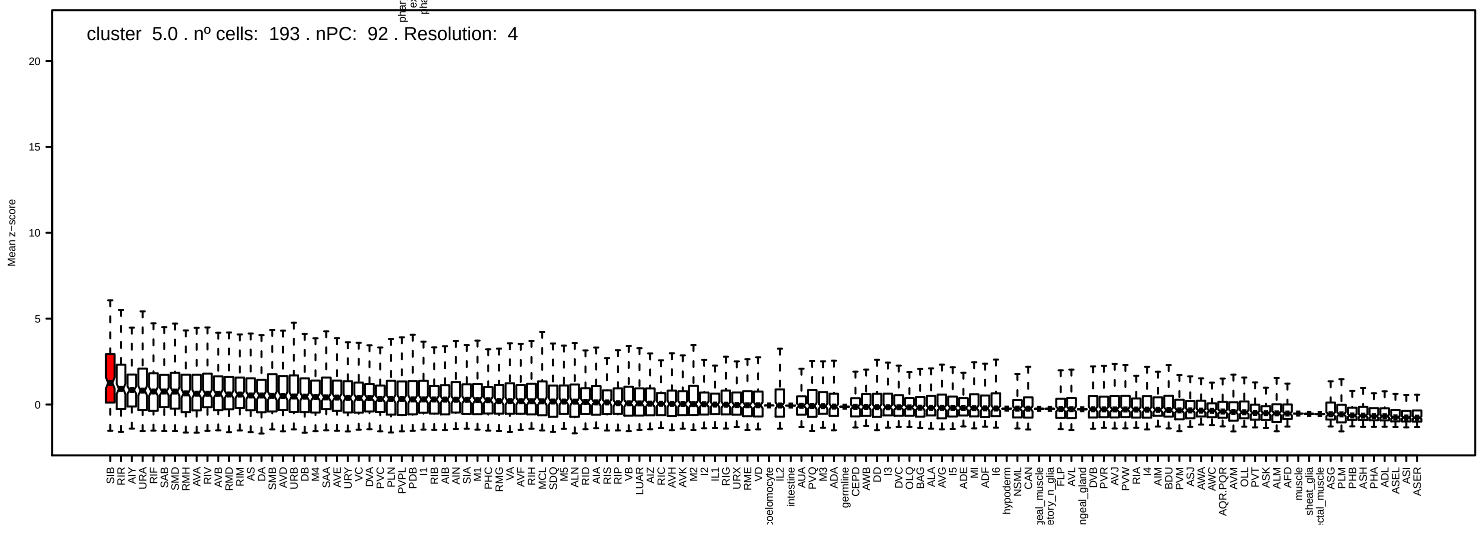
cluster 5.0 . nº cells: 193 . nPC: 92 . Resolution: 4
Mean z−score

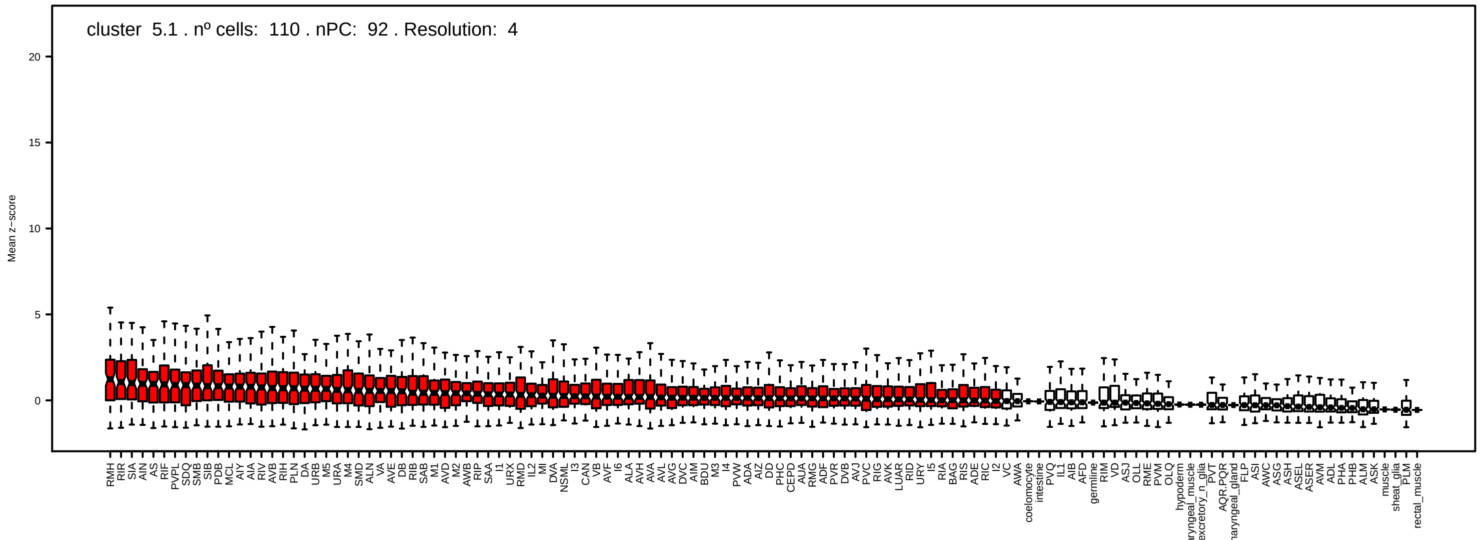
cluster 5.1 . nº cells: 110 . nPC: 92 . Resolution: 4
Mean z−score
0
5
10
15
20

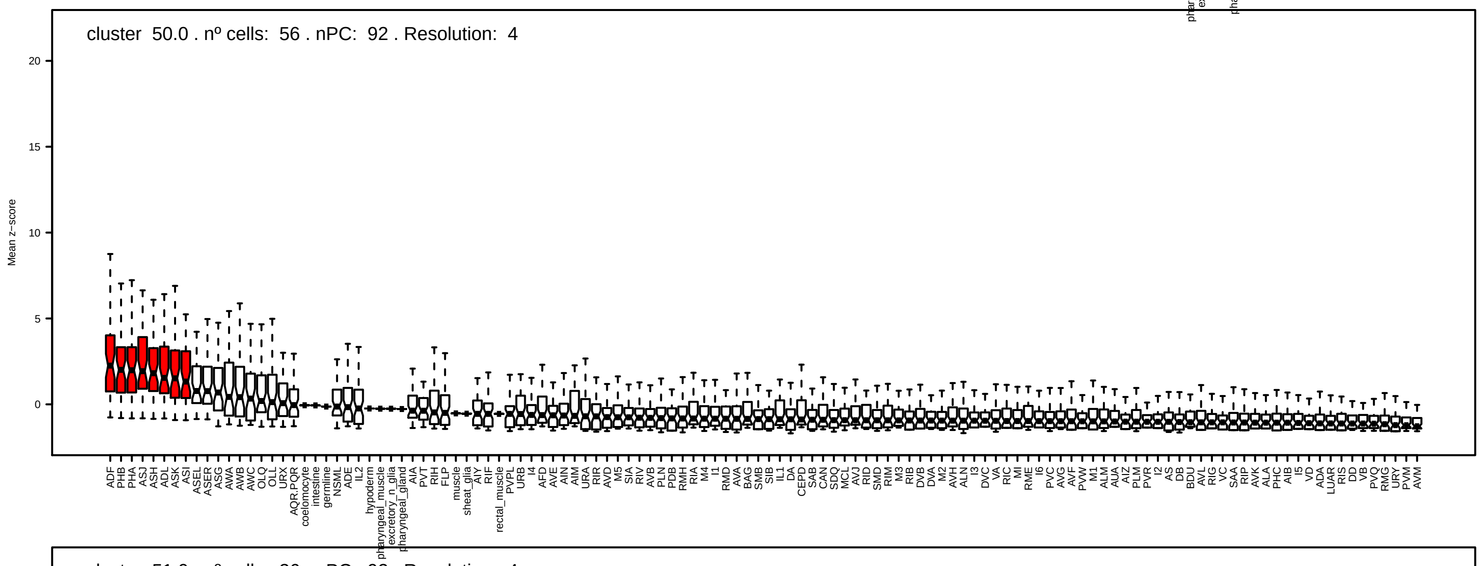
cluster 50.0 . nº cells: 56 . nPC: 92 . Resolution: 4
Mean z−score
0
5
10
15
20

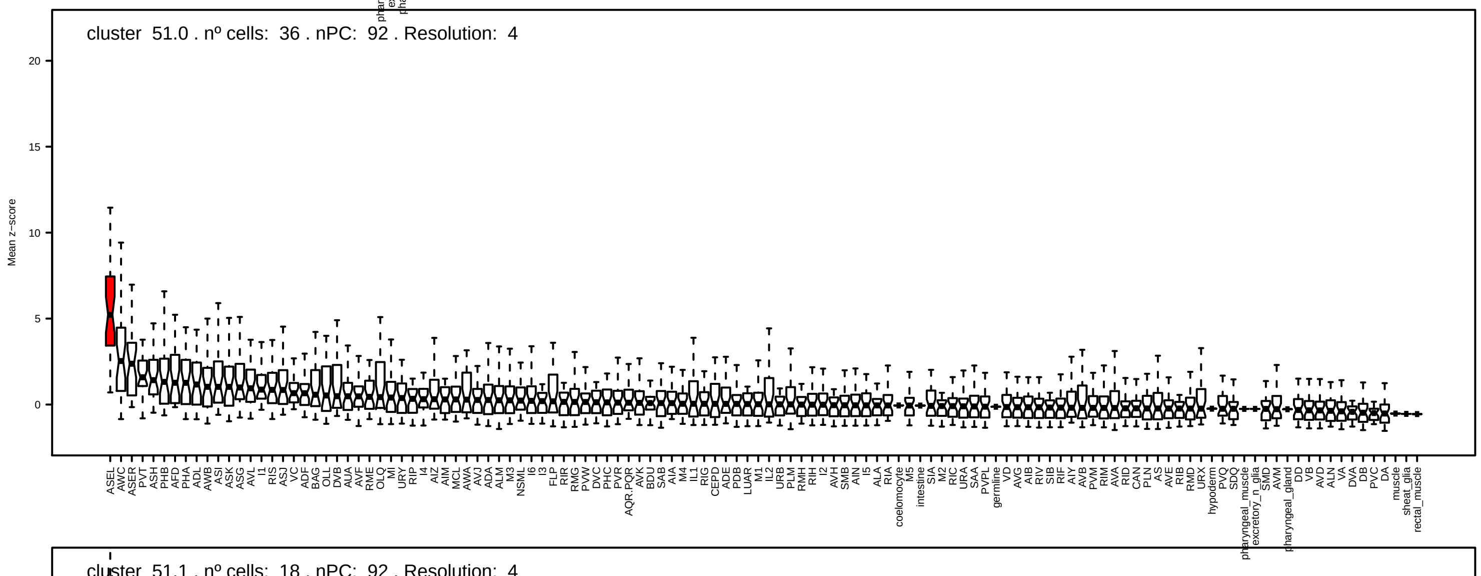
cluster 51.0 . nº cells: 36 . nPC: 92 . Resolution: 4
Mean z−score
0
5
10
15
20

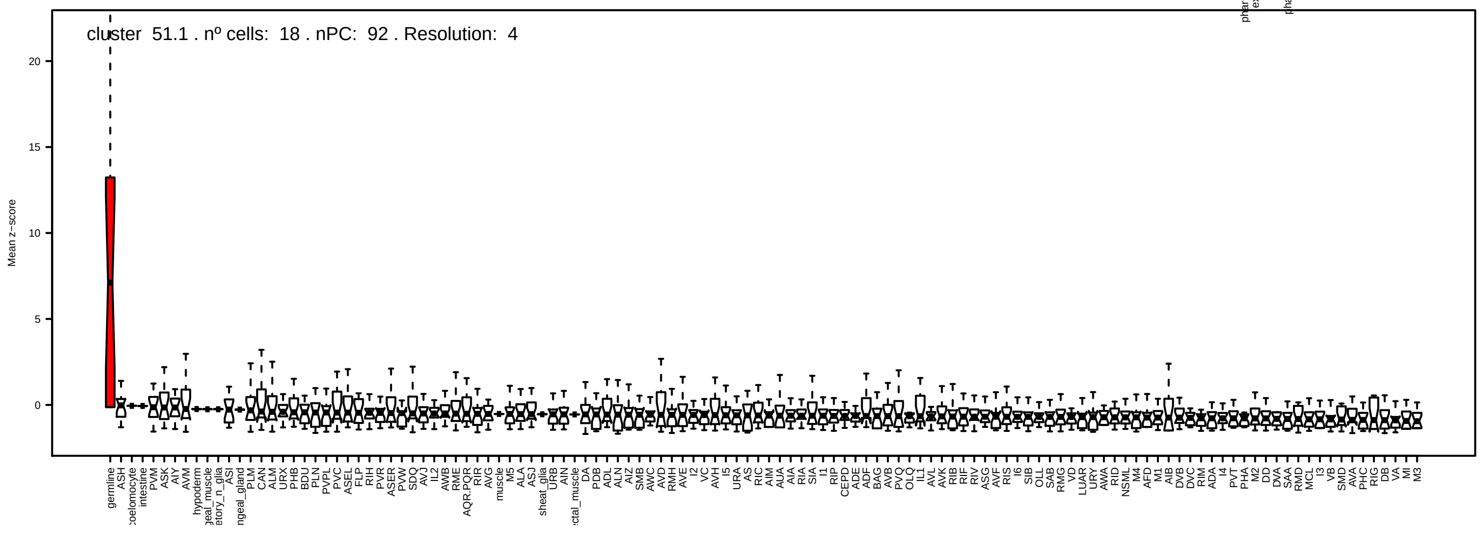
cluster 51.1 . nº cells: 18 . nPC: 92 . Resolution: 4
Mean z−score
0
5
10
15
20

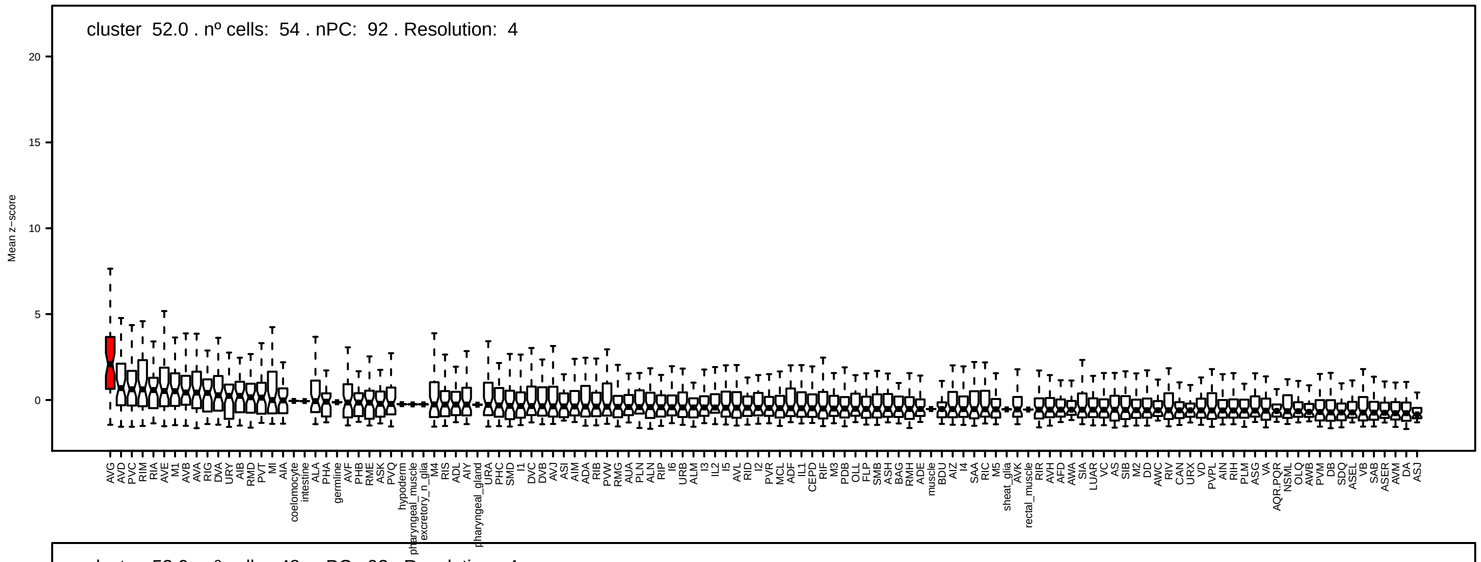
cluster 52.0 . nº cells: 54 . nPC: 92 . Resolution: 4
Mean z-score

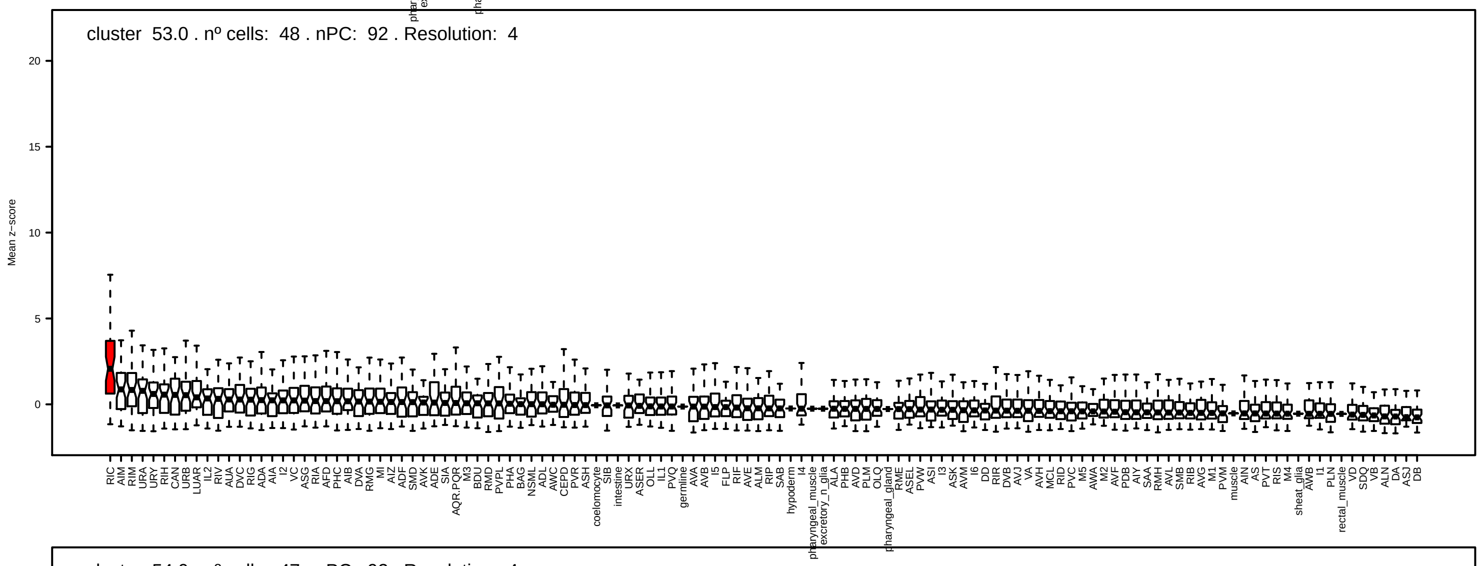
cluster 53.0 . nº cells: 48 . nPC: 92 . Resolution: 4
Mean z-score

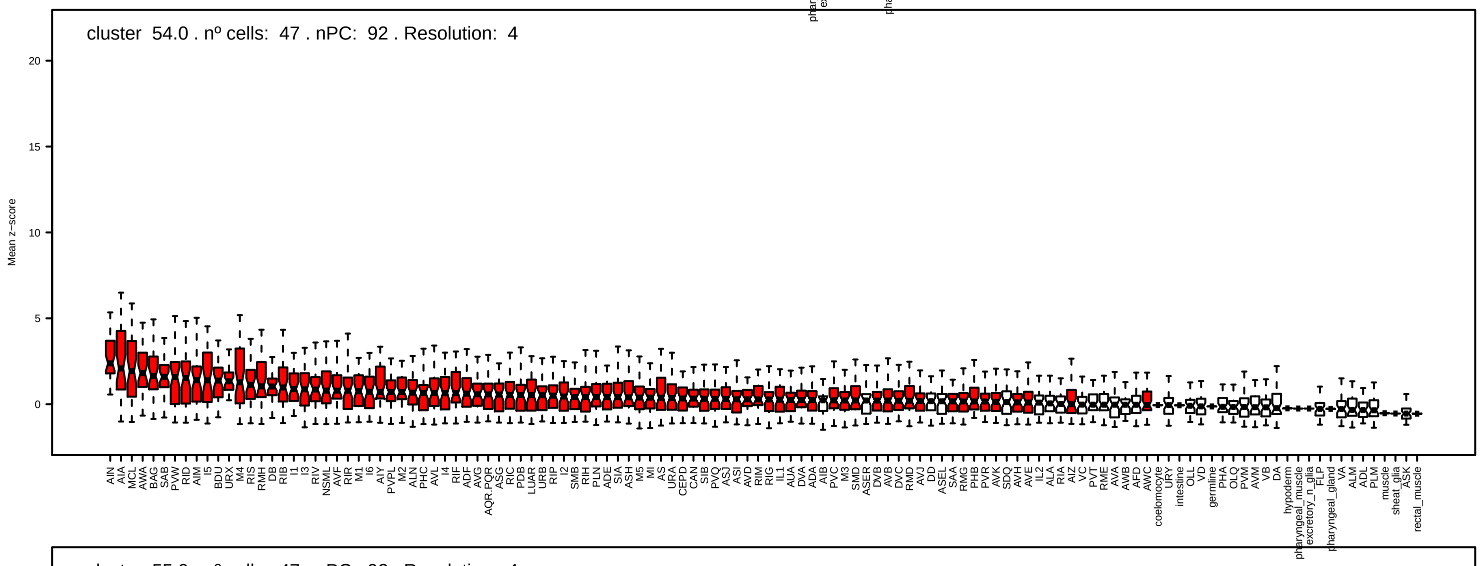
cluster 54.0 . nº cells: 47 . nPC: 92 . Resolution: 4
Mean z-score

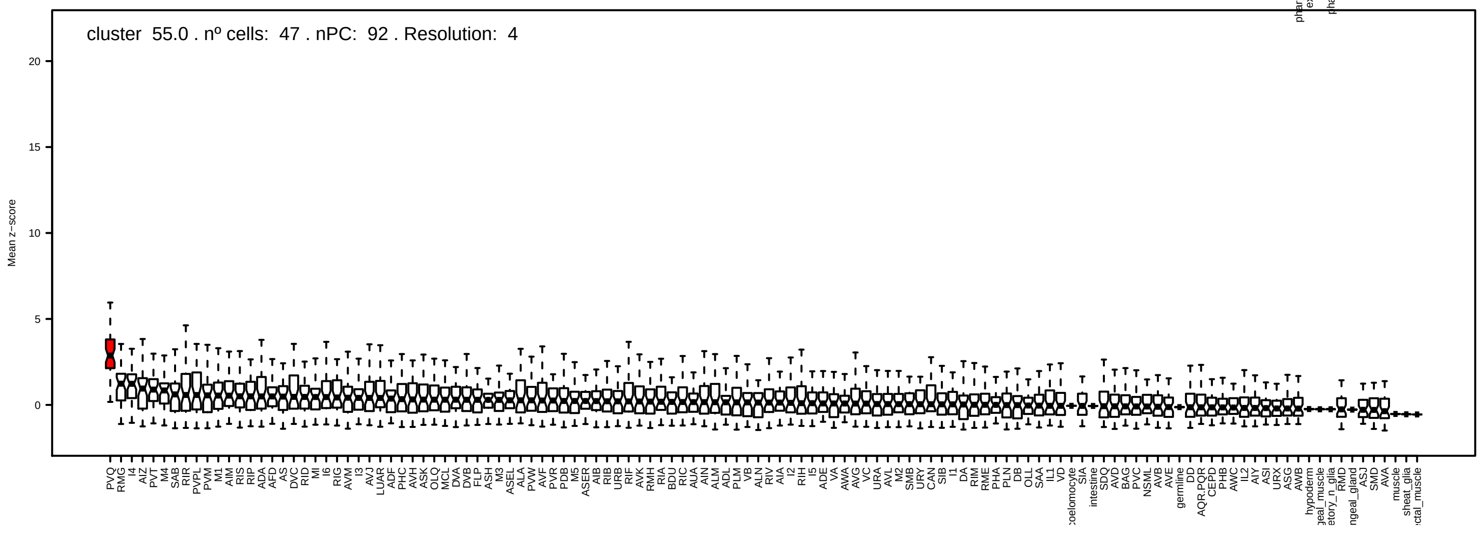
cluster 55.0 . nº cells: 47 . nPC: 92 . Resolution: 4
Mean z-score

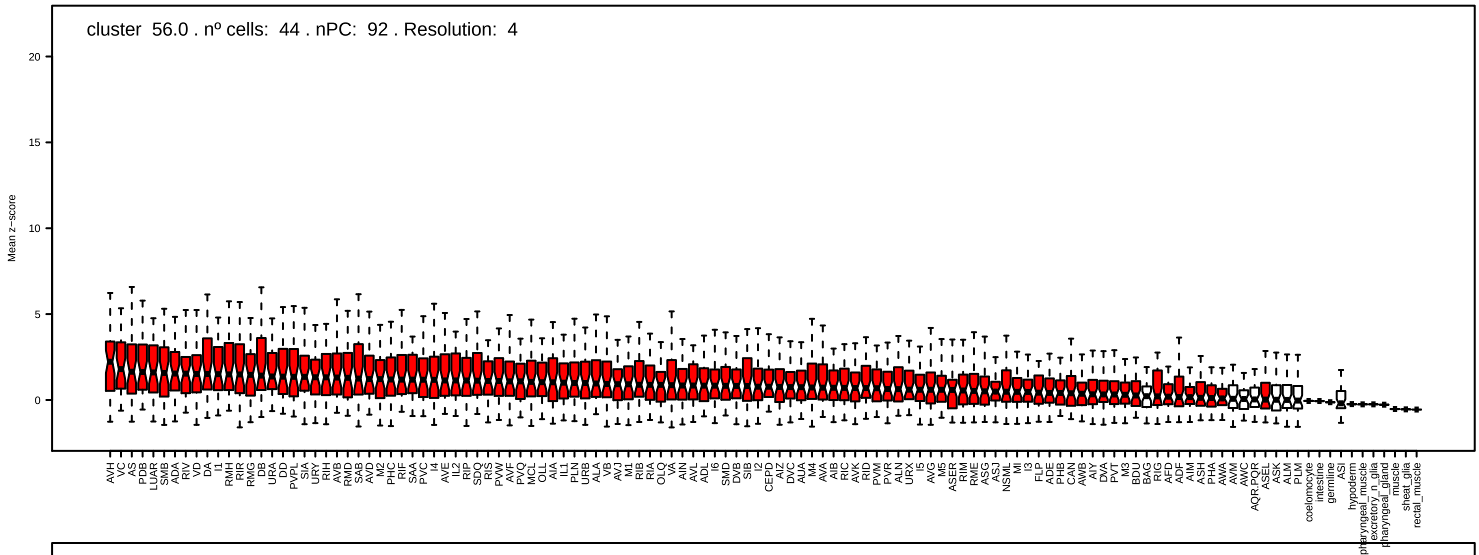
cluster 56.0 . nº cells: 44 . nPC: 92 . Resolution: 4
Mean z−score
0
5
10
15
20

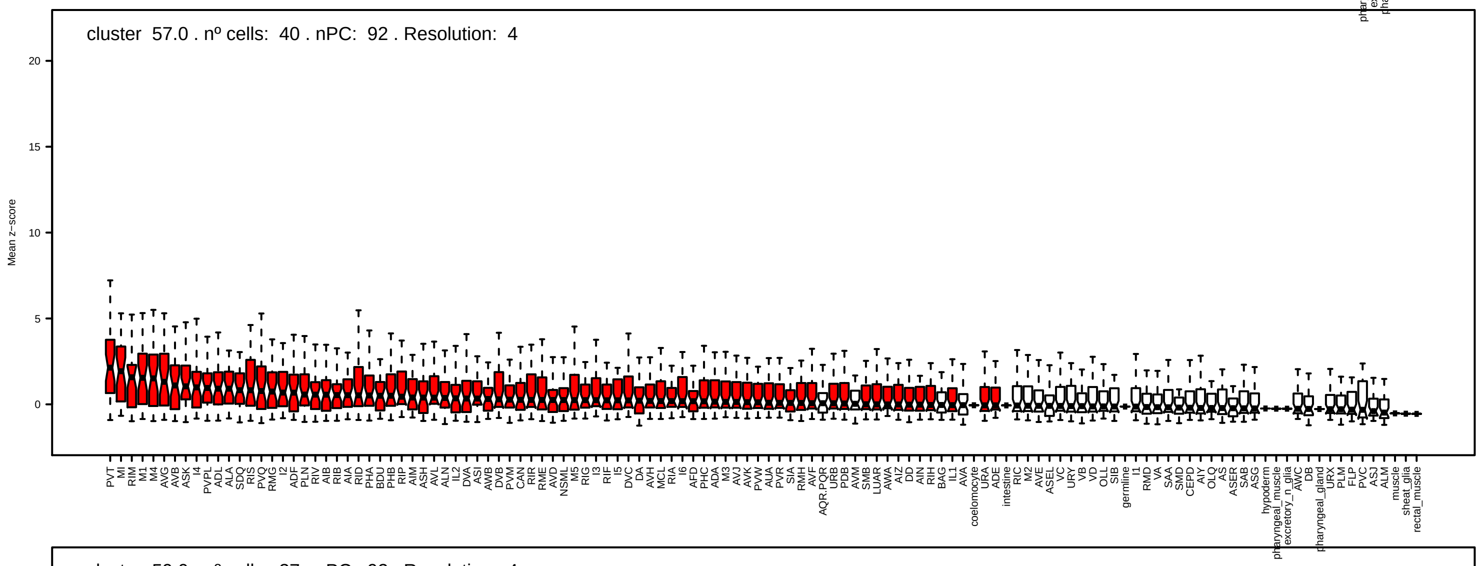
cluster 57.0 . nº cells: 40 . nPC: 92 . Resolution: 4
Mean z−score
0
5
10
15
20

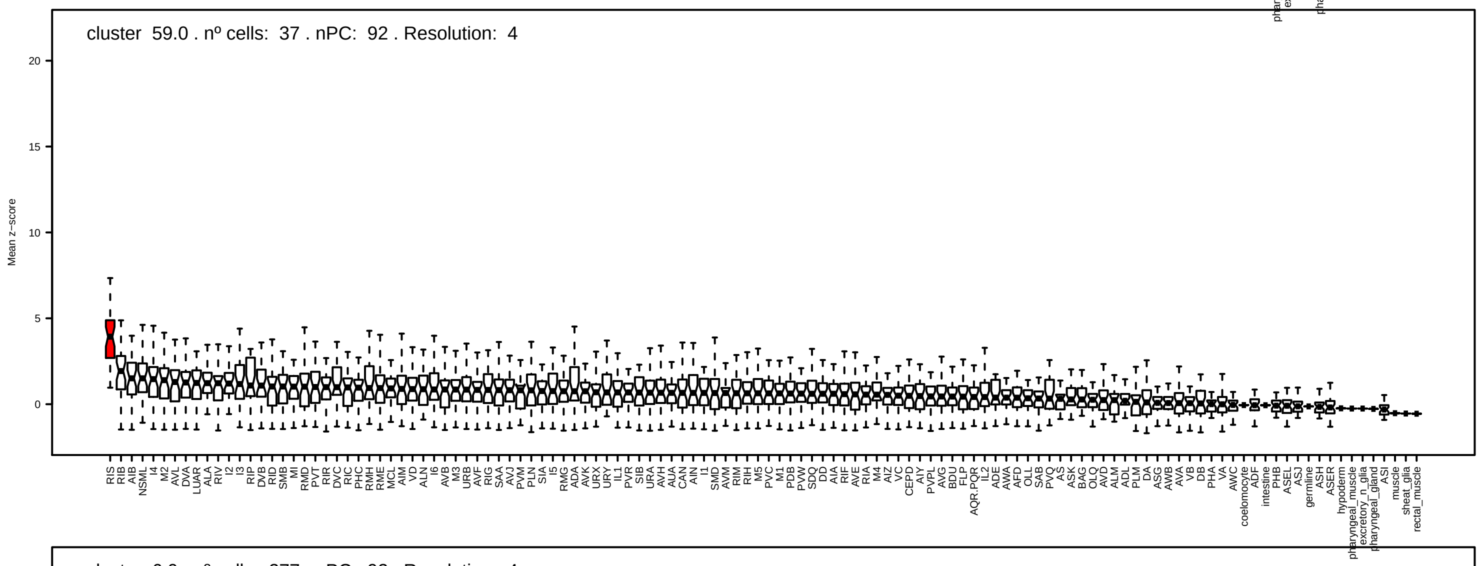
cluster 59.0 . nº cells: 37 . nPC: 92 . Resolution: 4
Mean z−score
0
5
10
15
20

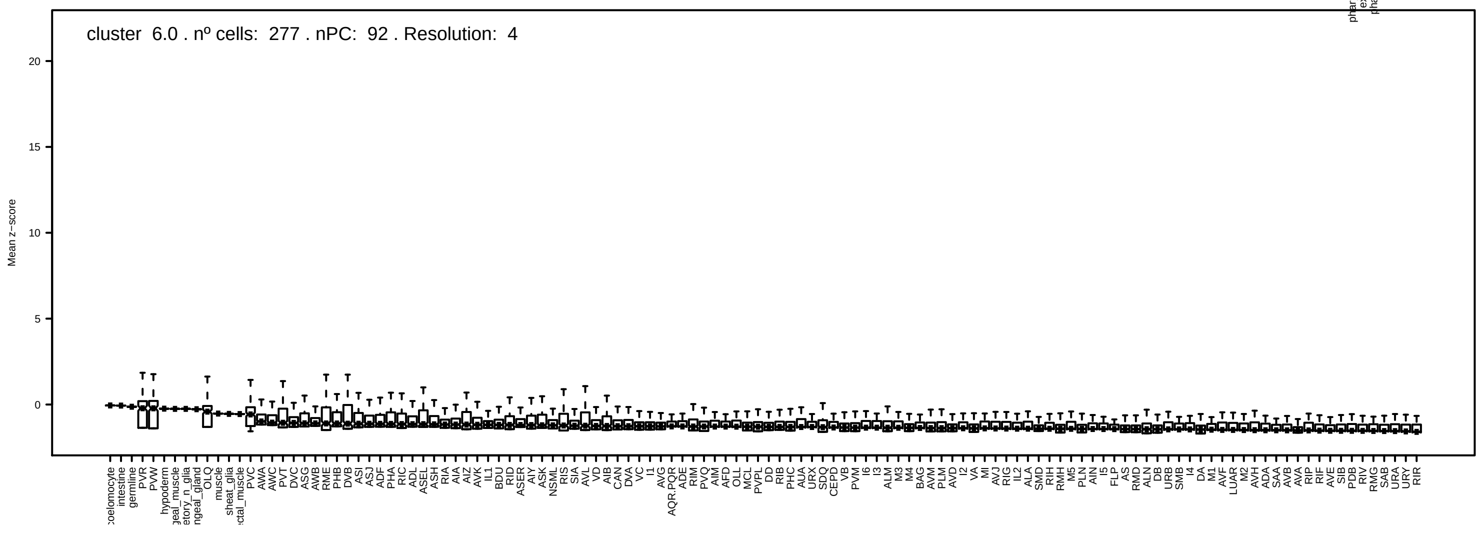
cluster 6.0 . nº cells: 277 . nPC: 92 . Resolution: 4
Mean z−score
0
5
10
15
20

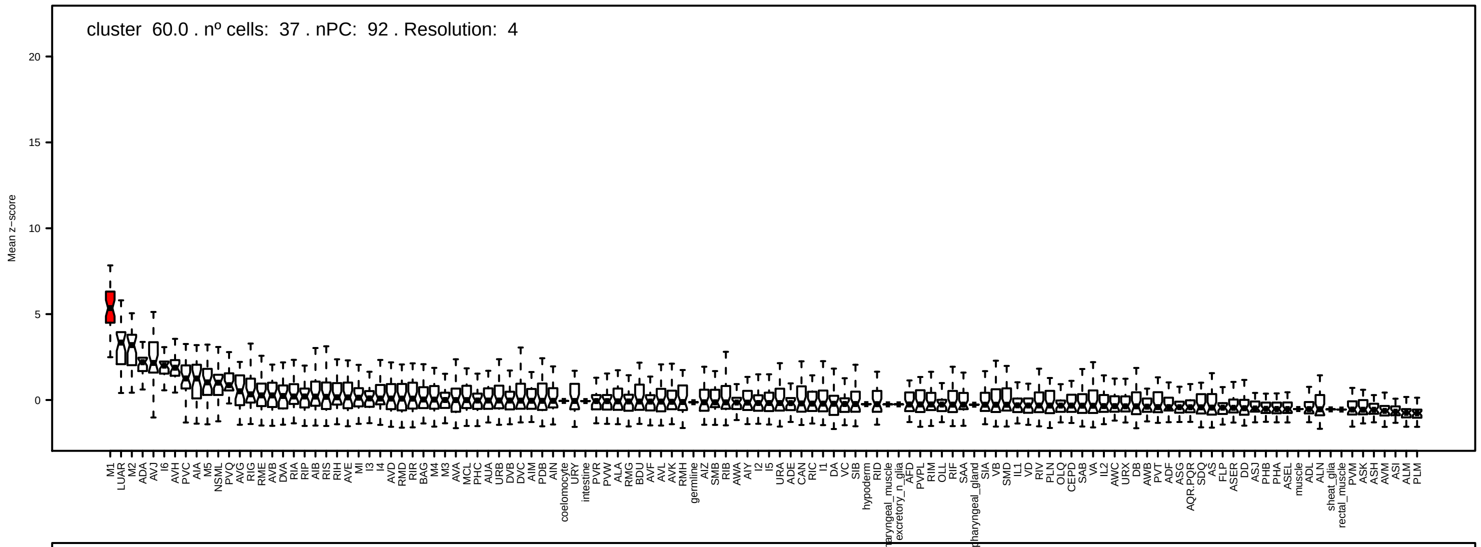
cluster 60.0 . nº cells: 37 . nPC: 92 . Resolution: 4
Mean z−score

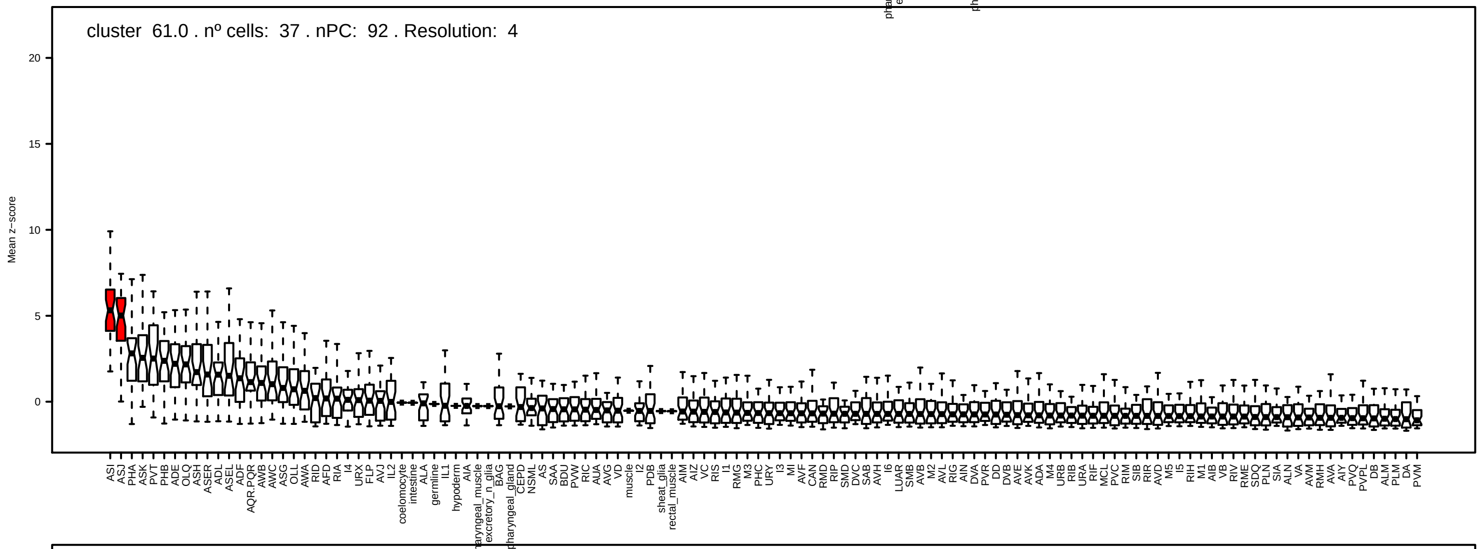
cluster 61.0 . nº cells: 37 . nPC: 92 . Resolution: 4
Mean z−score

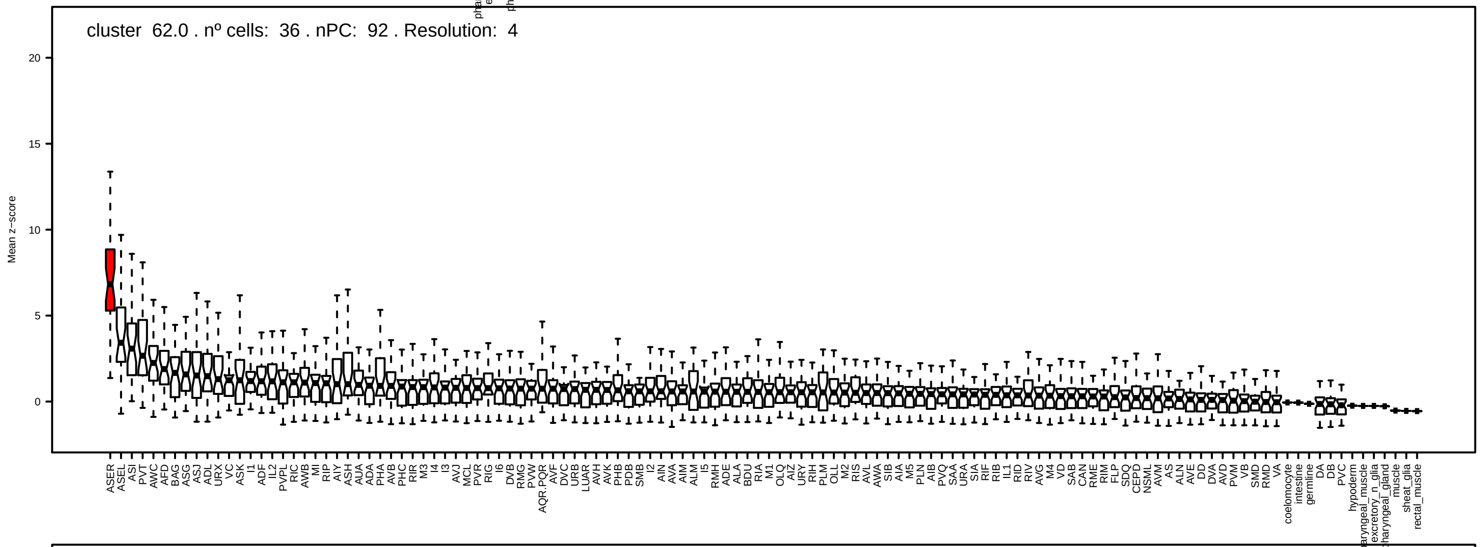
cluster 62.0 . nº cells: 36 . nPC: 92 . Resolution: 4
Mean z−score

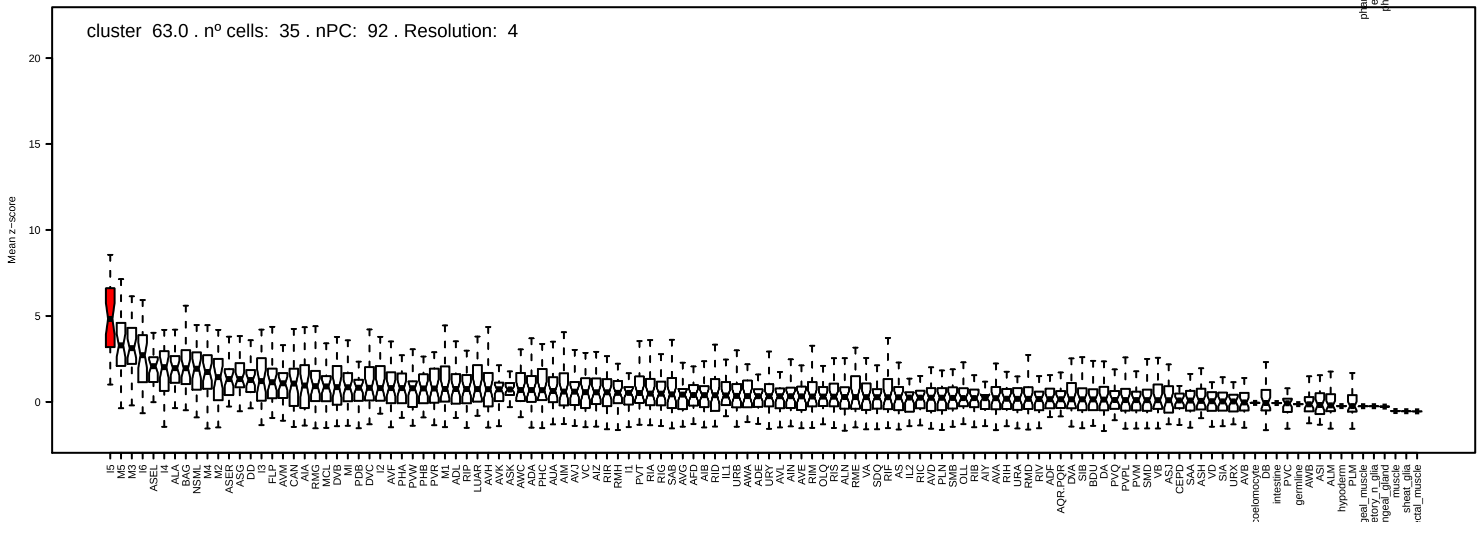
cluster 63.0 . nº cells: 35 . nPC: 92 . Resolution: 4
Mean z−score

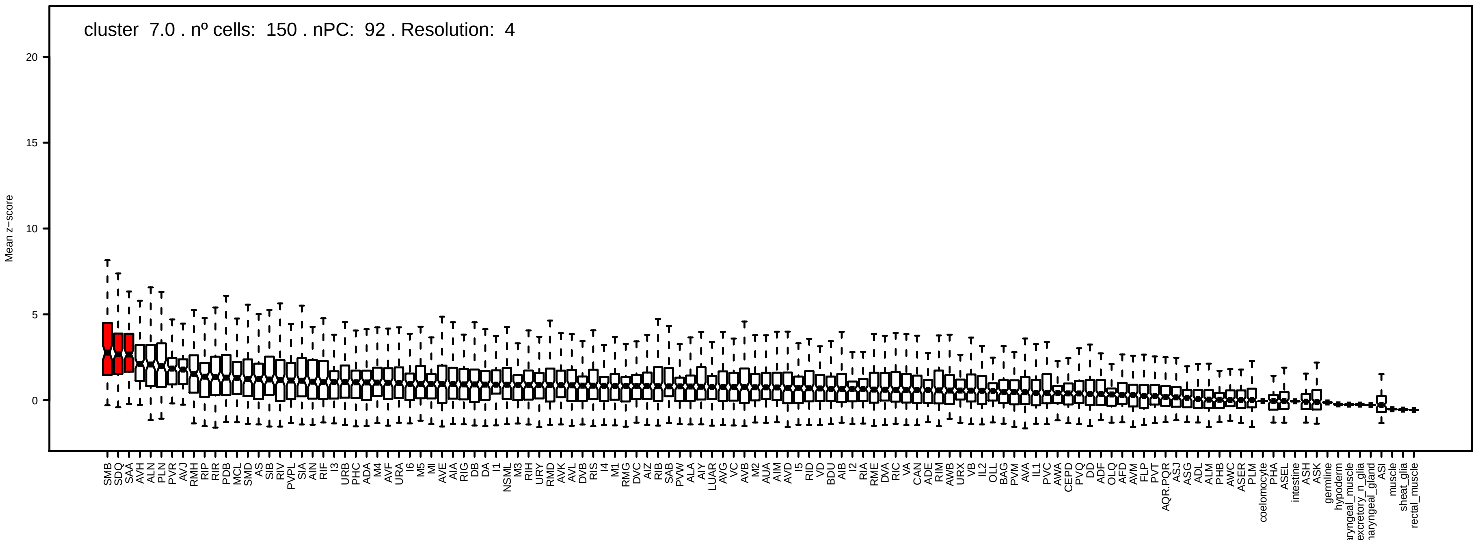
cluster 7.0 . nº cells: 150 . nPC: 92 . Resolution: 4
Mean z−score

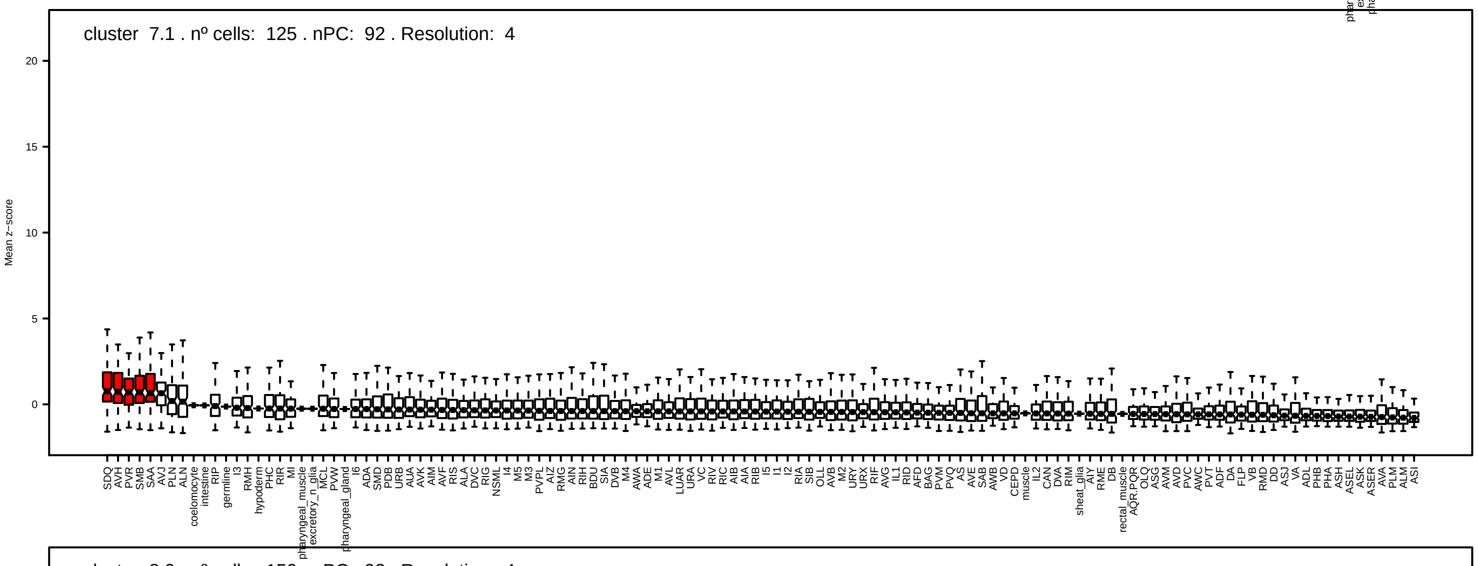
cluster 7.1 . nº cells: 125 . nPC: 92 . Resolution: 4
Mean z−score

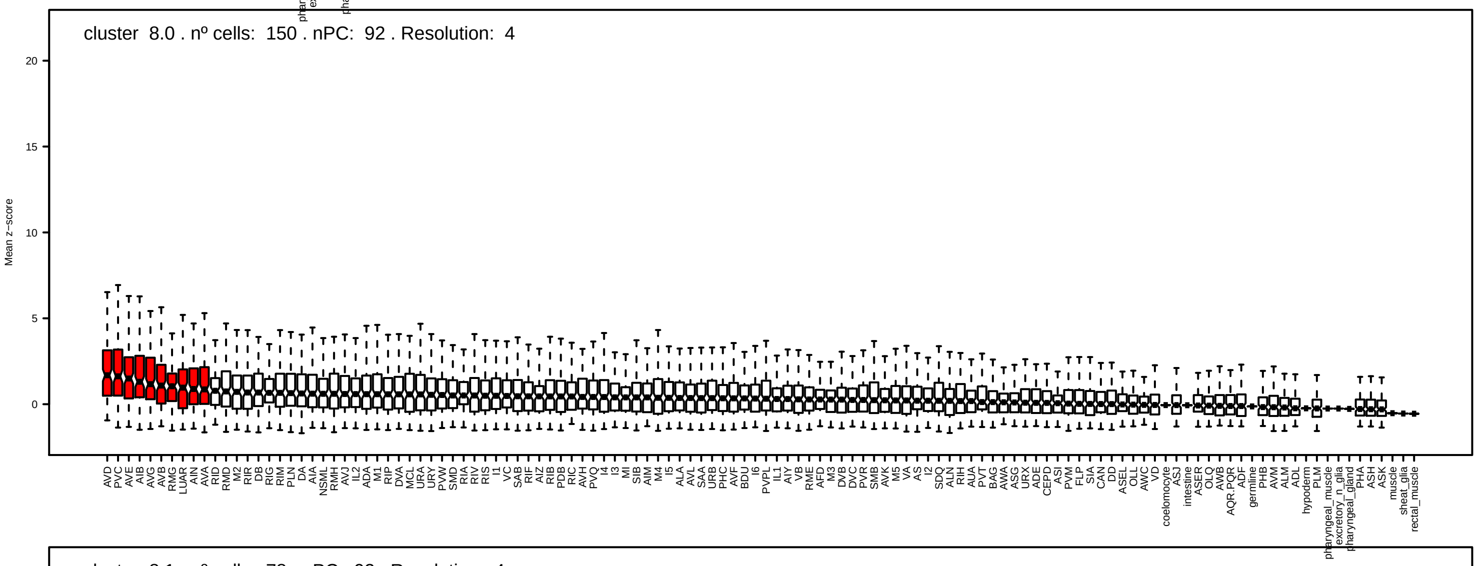
cluster 8.0 . nº cells: 150 . nPC: 92 . Resolution: 4
Mean z−score

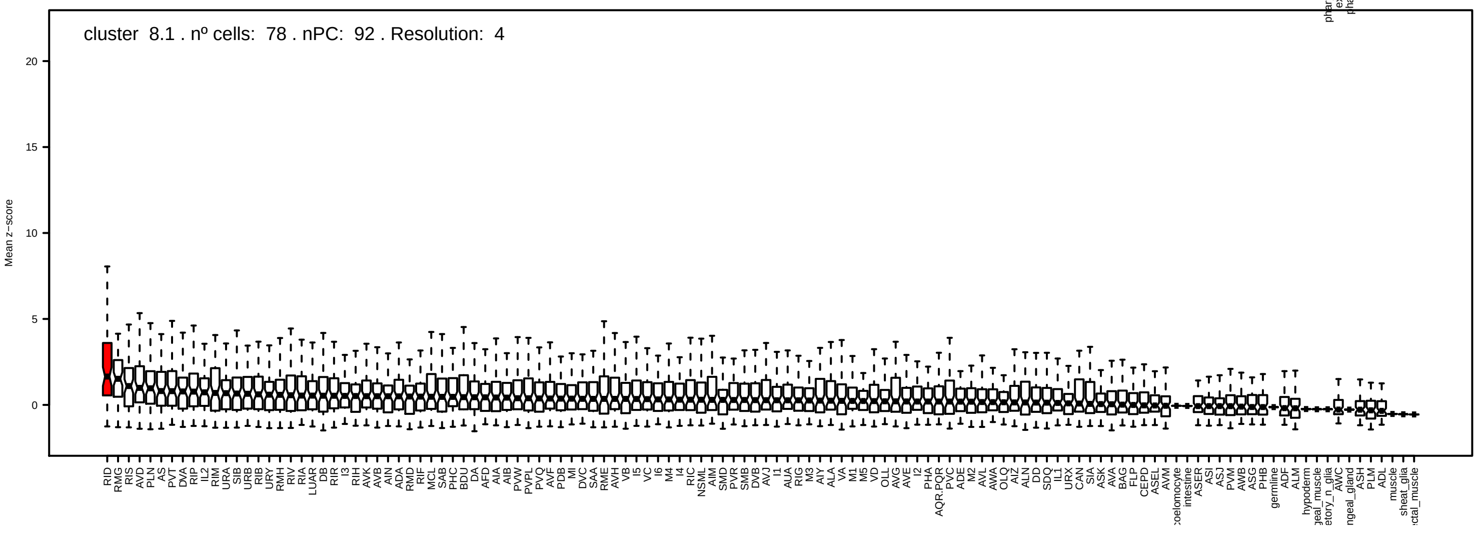
cluster 8.1 . nº cells: 78 . nPC: 92 . Resolution: 4
Mean z−score

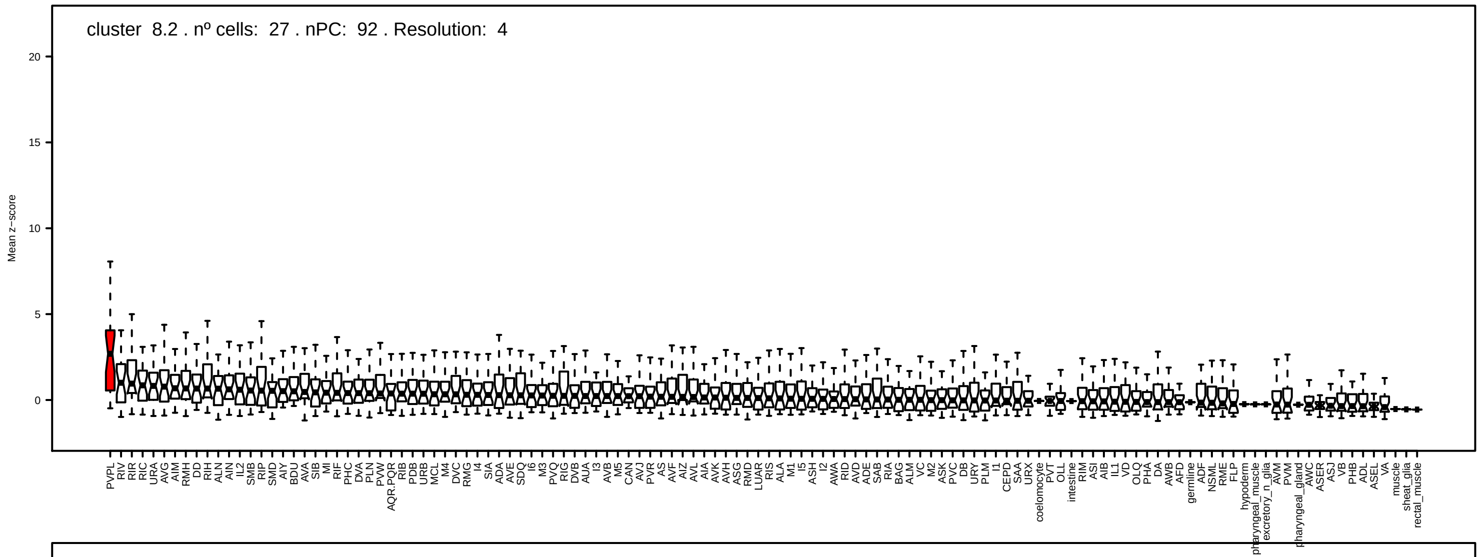
cluster 8.2 . nº cells: 27 . nPC: 92 . Resolution: 4
Mean z−score

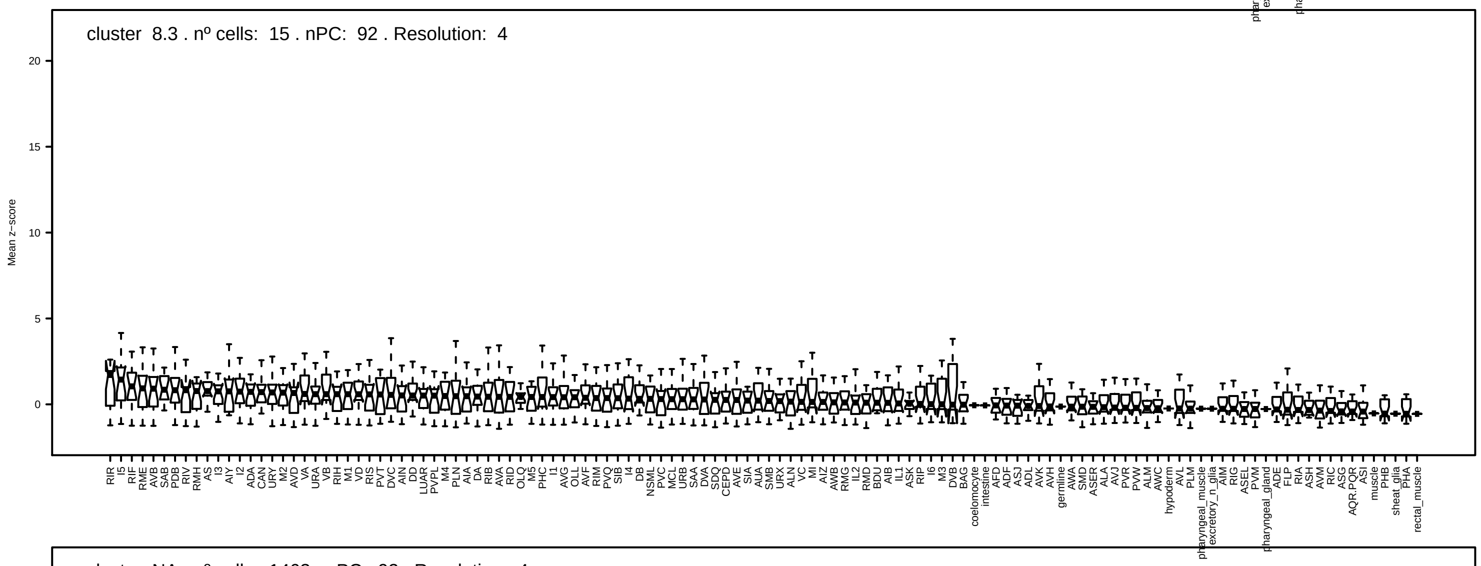
cluster 8.3 . nº cells: 15 . nPC: 92 . Resolution: 4
Mean z−score

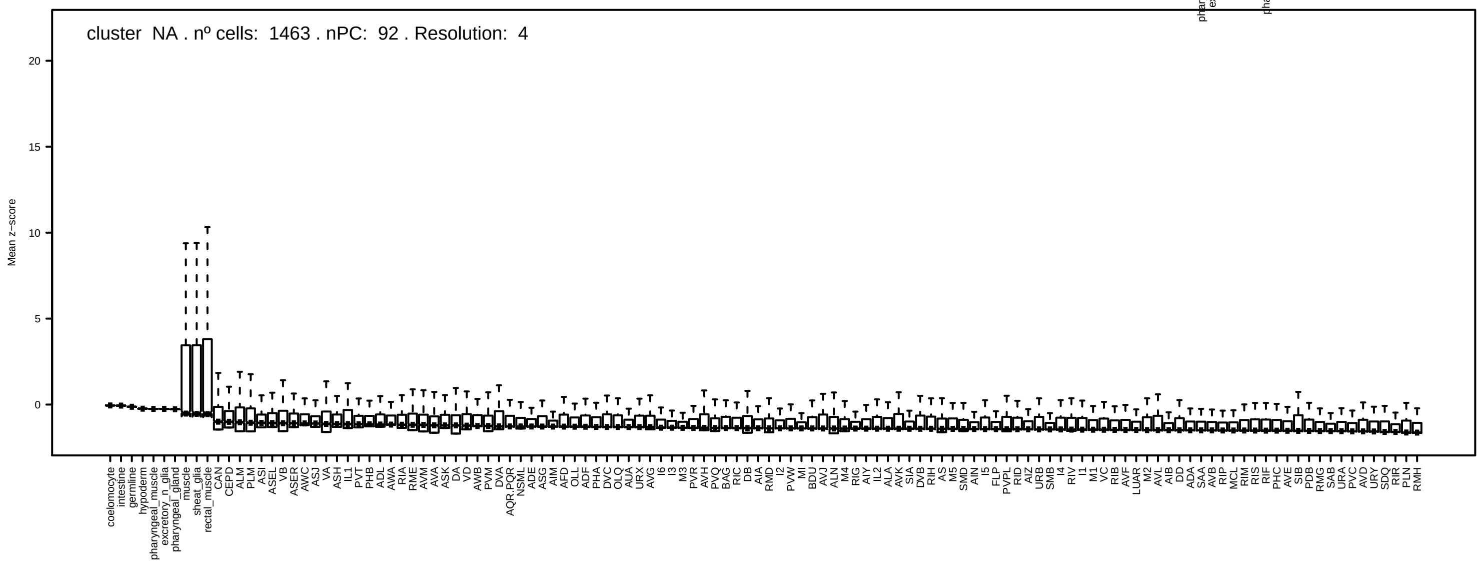
cluster NA . nº cells: 1463 . nPC: 92 . Resolution: 4
Mean z−score

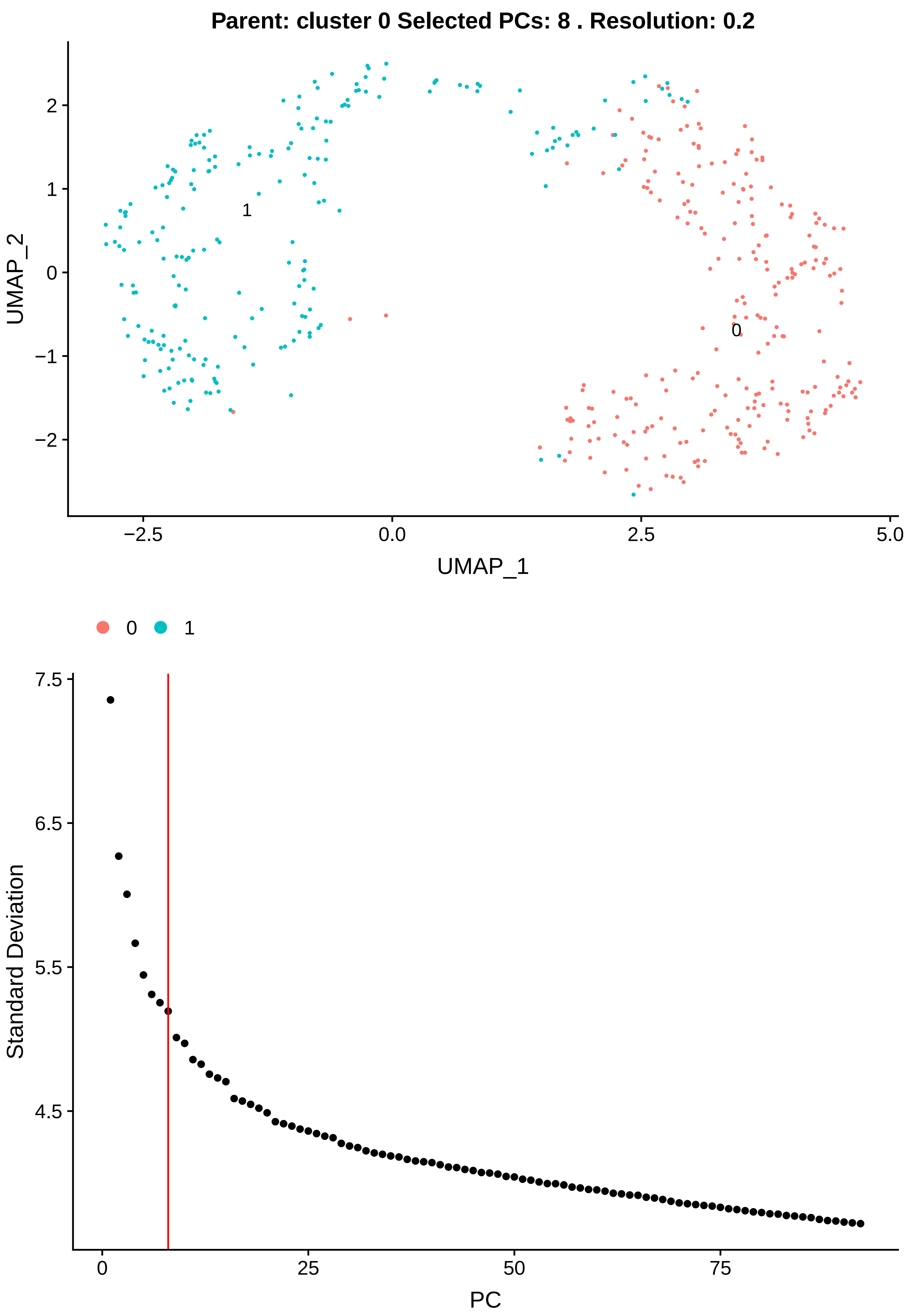

Parent: cluster 0 Selected PCs: 8 . Resolution: 0.2
UMAP_2
UMAP_1
−2.5
0.0
2.5
5.0
2
1
0
−1
−2
1
0
0
1
Standard Deviation
PC
7.5
6.5
5.5
4.5
0
25
50
75

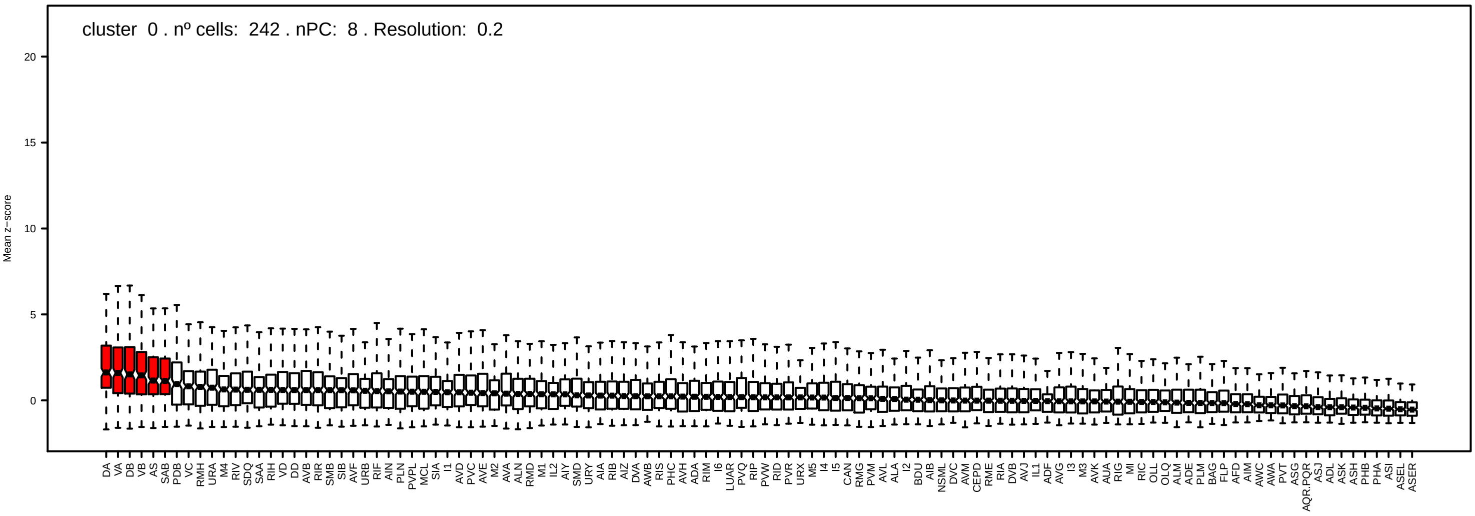
cluster 0 . nº cells: 242 . nPC: 8 . Resolution: 0.2
Mean z−score
20
15
10
5
0

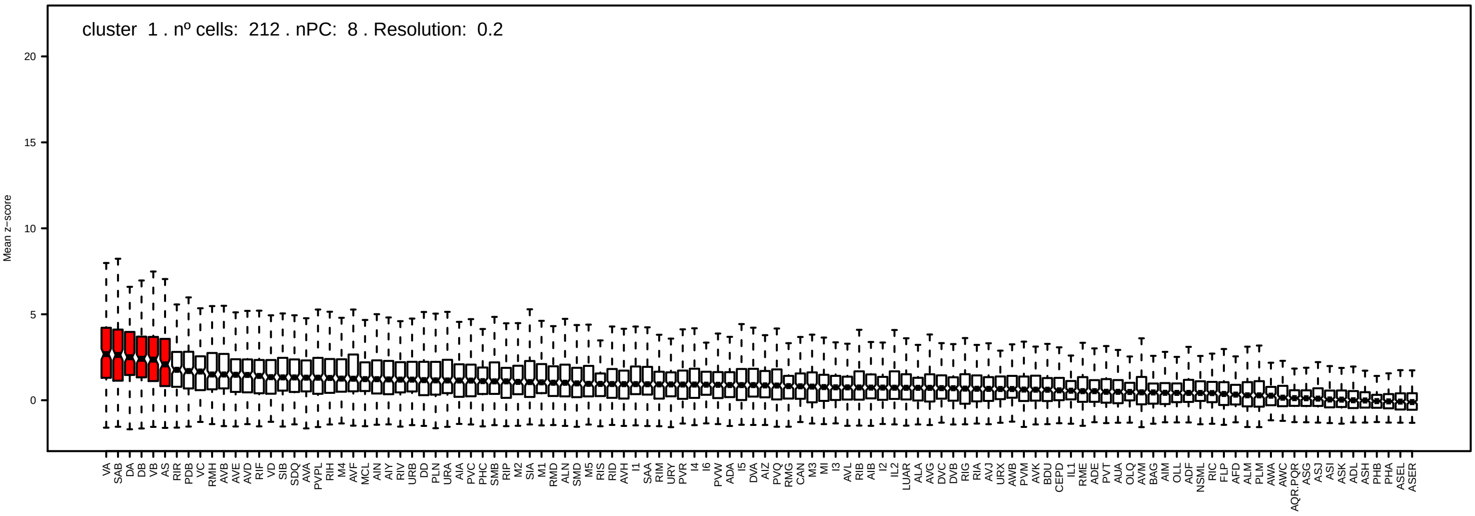
cluster 1 . nº cells: 212 . nPC: 8 . Resolution: 0.2
Mean z−score
20
15
10
5
0

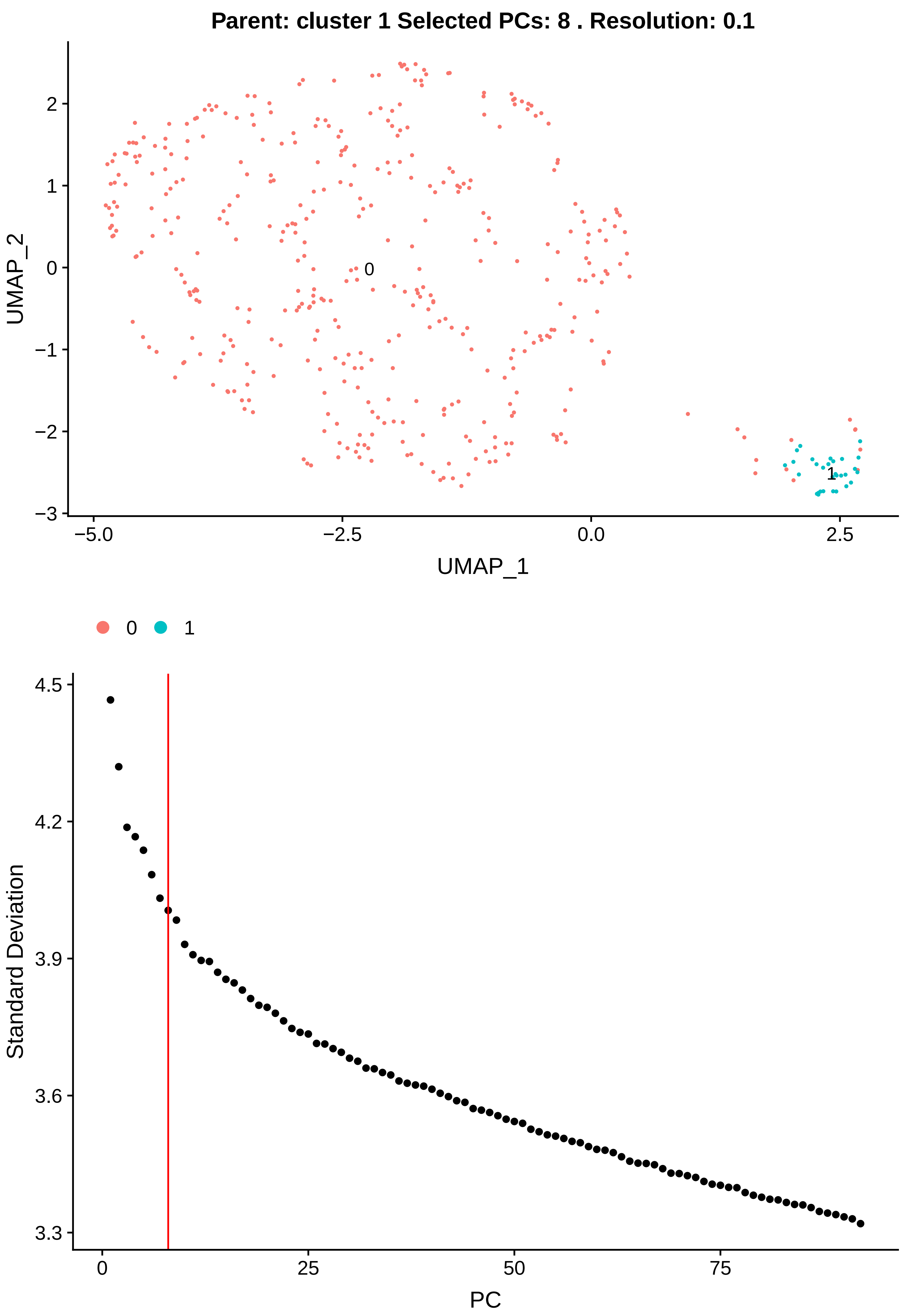

Parent: cluster 1 Selected PCs: 8 . Resolution: 0.1
UMAP_2
UMAP_1
0
1
2
−1
−2
−3
−5.0
−2.5
0.0
2.5
0
1
Standard Deviation
PC
4.5
4.2
3.9
3.6
3.3
0
25
50
75

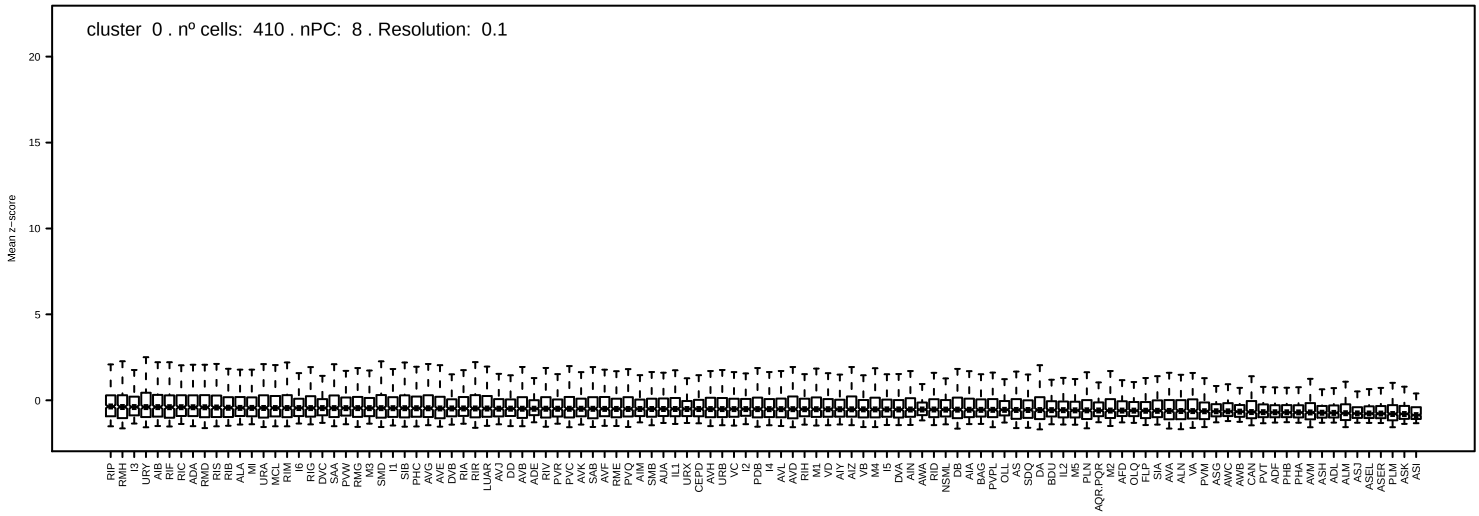
cluster 0 . nº cells: 410 . nPC: 8 . Resolution: 0.1
Mean z−score
20
15
10
5
0

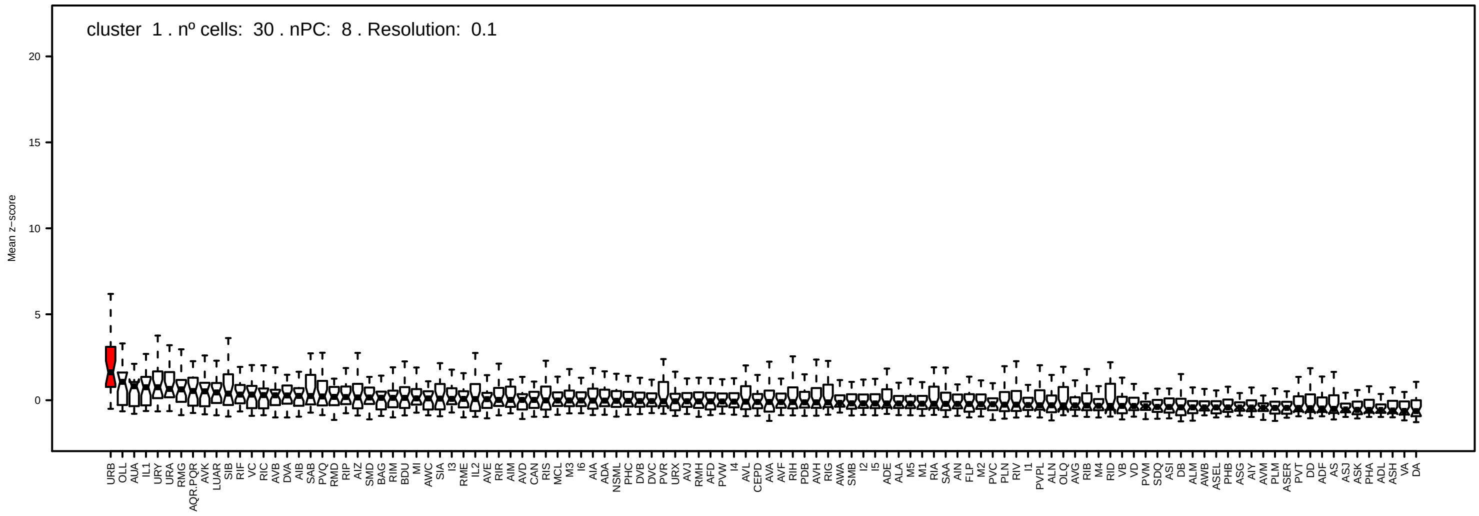
cluster 1 . nº cells: 30 . nPC: 8 . Resolution: 0.1
Mean z−score
20
15
10
5
0

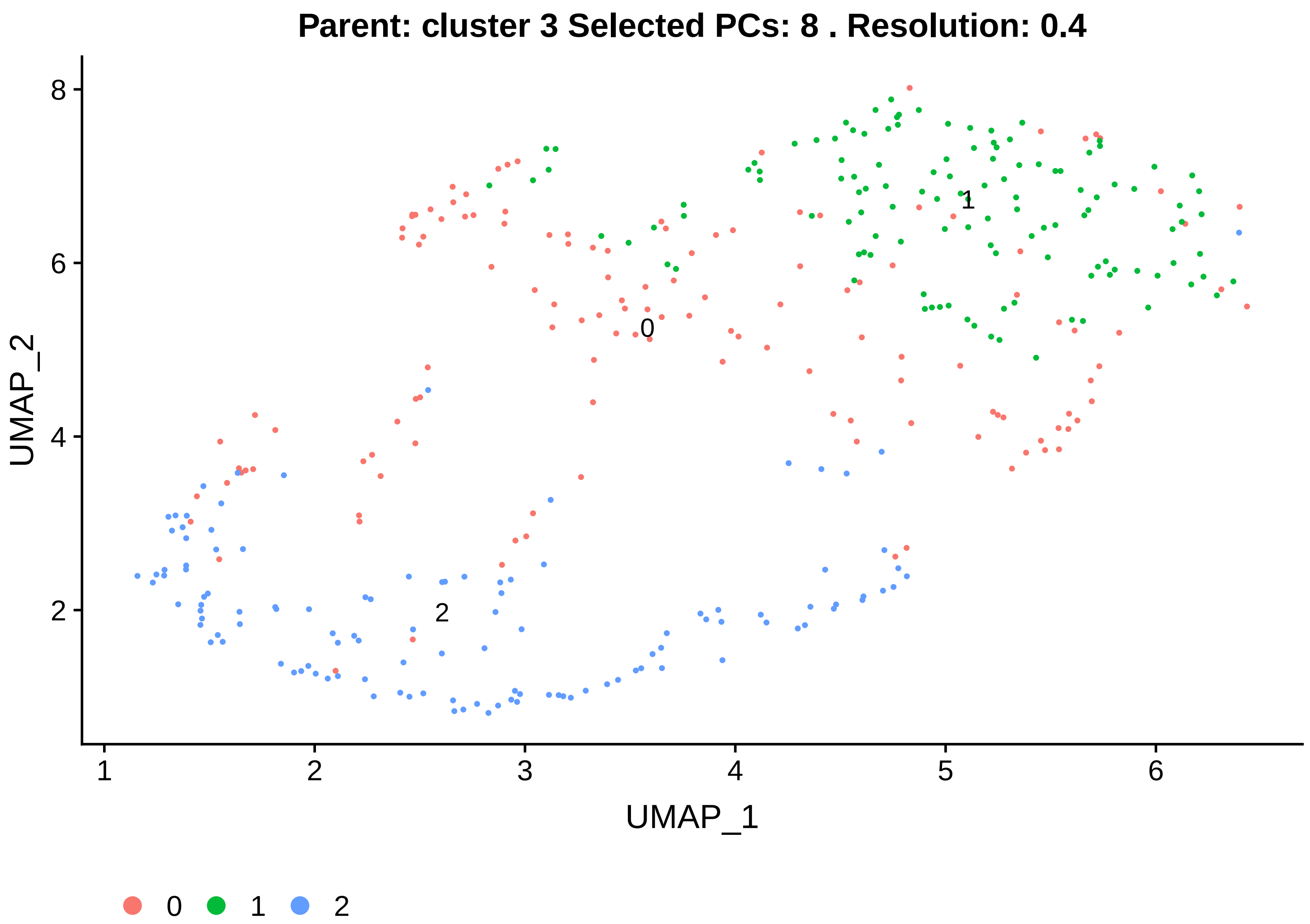

Parent: cluster 3 Selected PCs: 8 . Resolution: 0.4
UMAP_2
UMAP_1
0
1
2


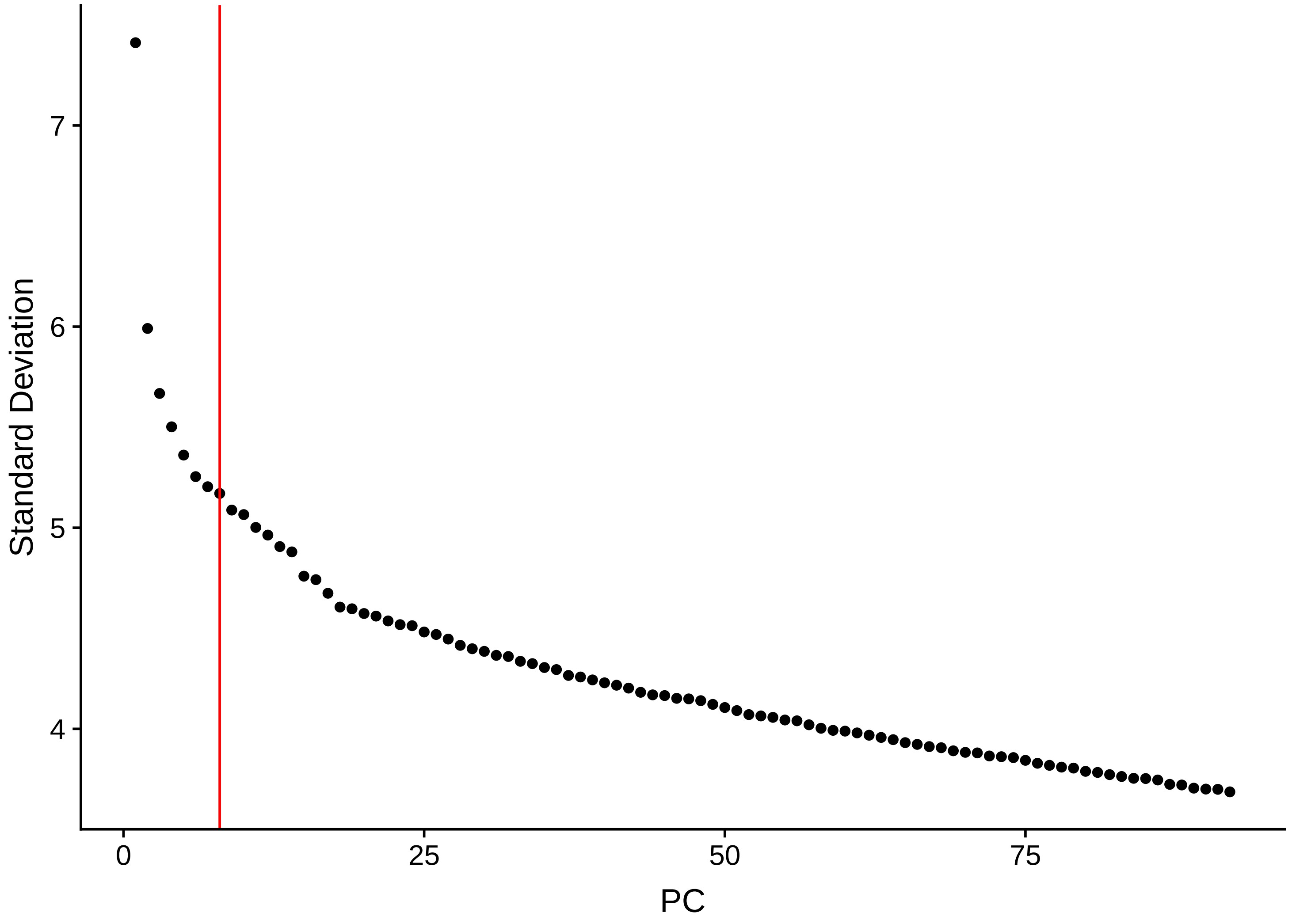

Standard Deviation
PC

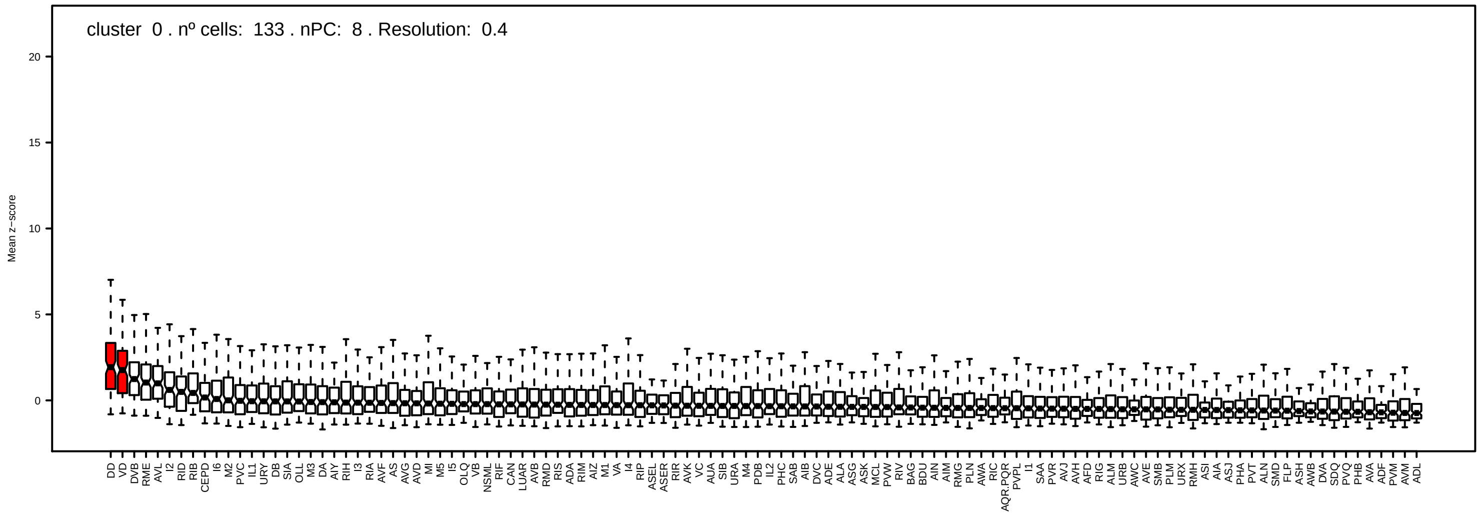
cluster 0 . nº cells: 133 . nPC: 8 . Resolution: 0.4
Mean z−score

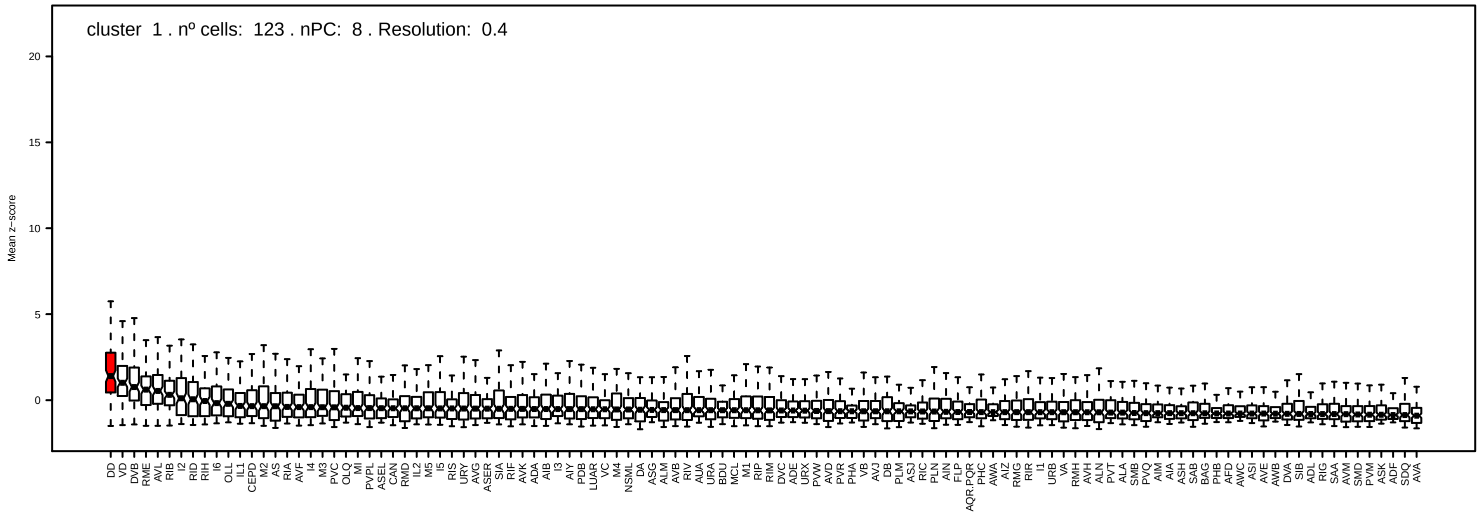
cluster 1 . nº cells: 123 . nPC: 8 . Resolution: 0.4
Mean z−score

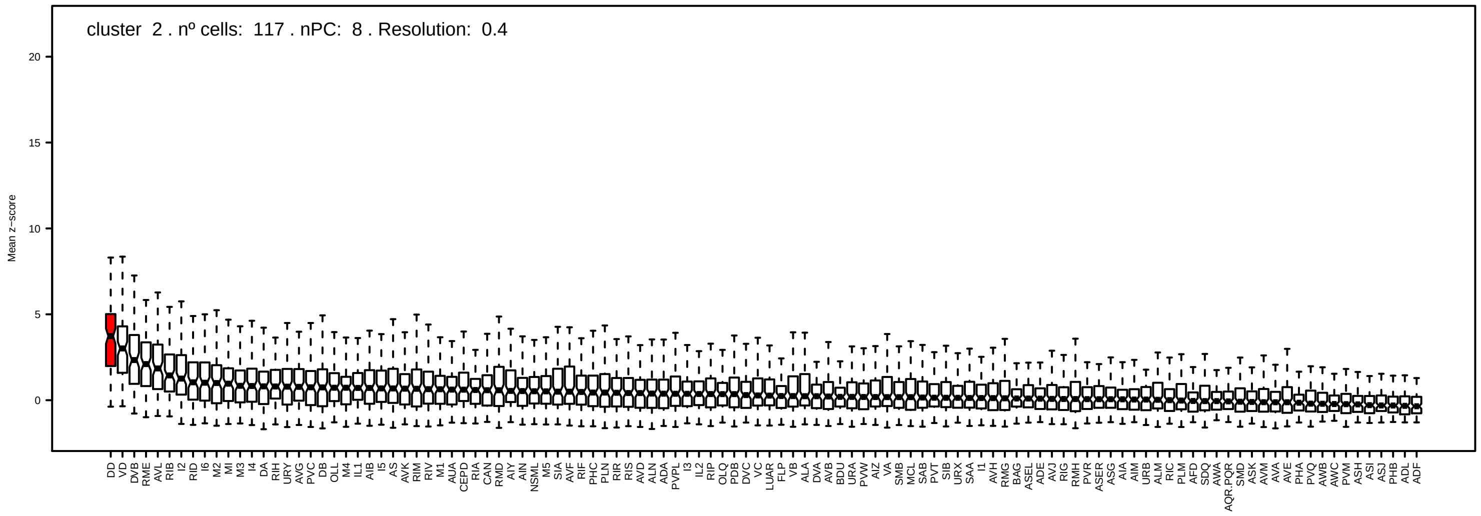
cluster 2 . nº cells: 117 . nPC: 8 . Resolution: 0.4
Mean z−score

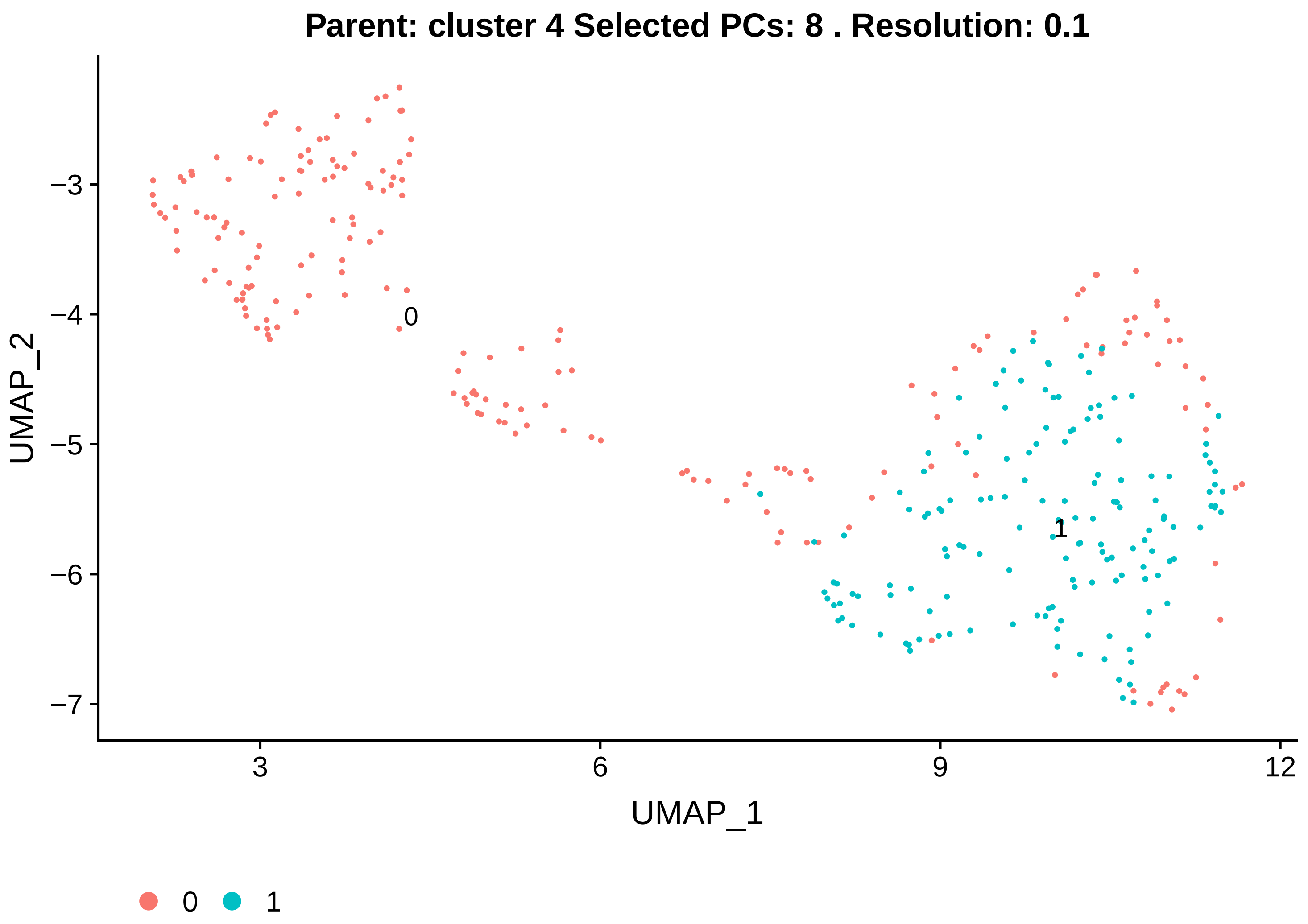
Parent: cluster 4 Selected PCs: 8 . Resolution: 0.1
UMAP_2
−3
−4
−5
−6
−7
0
1
3
6
9
12
UMAP_1
0
1

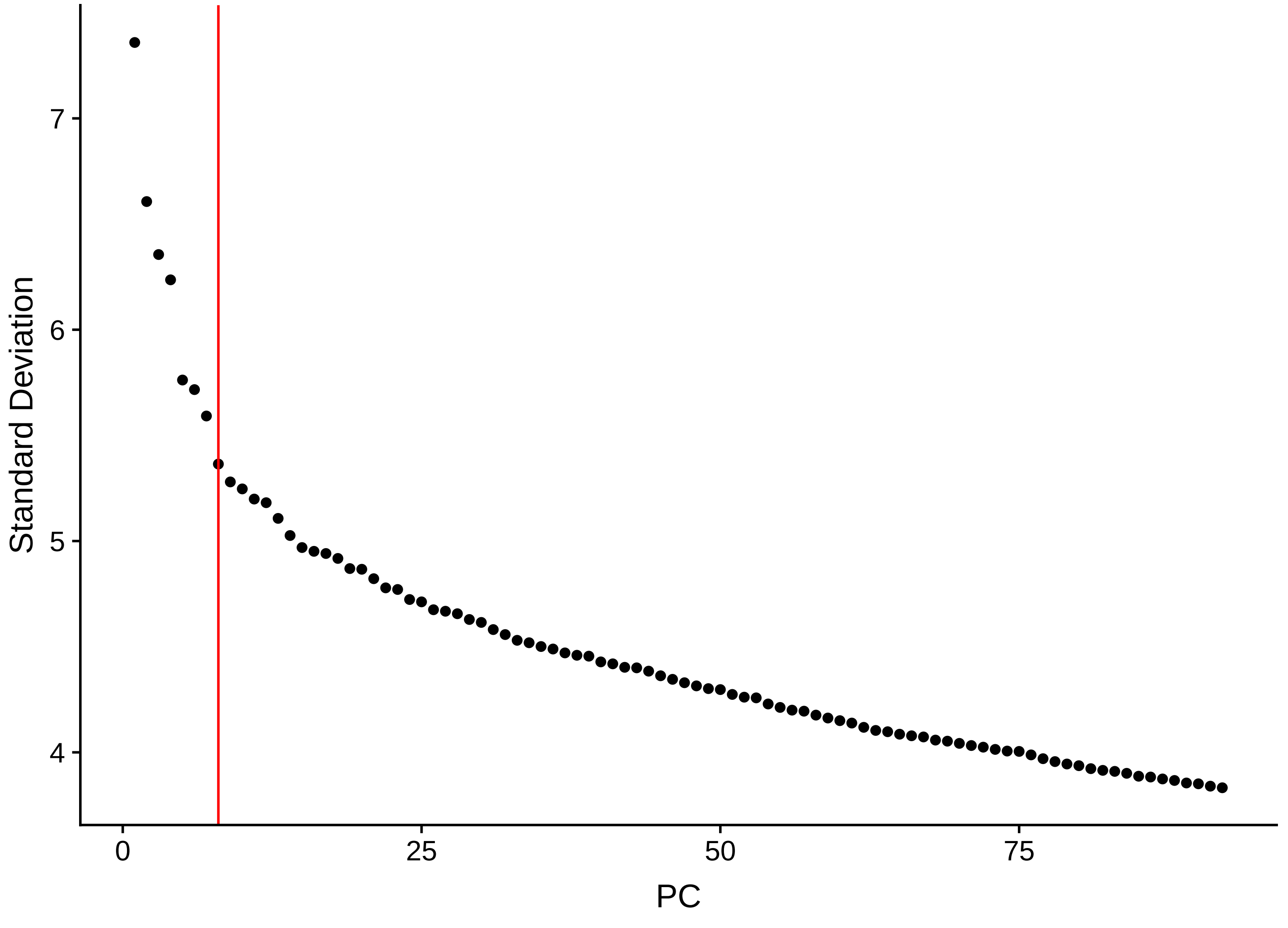
Standard Deviation
7
6
5
4
0
25
50
75
PC

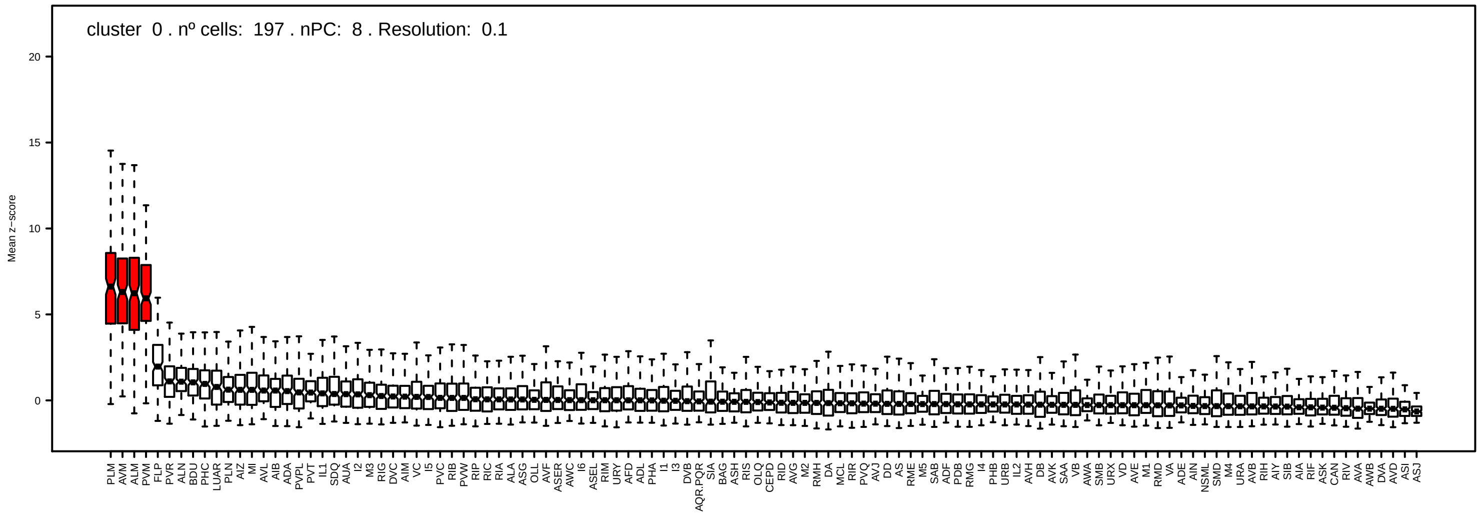
cluster 0 . nº cells: 197 . nPC: 8 . Resolution: 0.1
Mean z−score
20
15
10
5
0

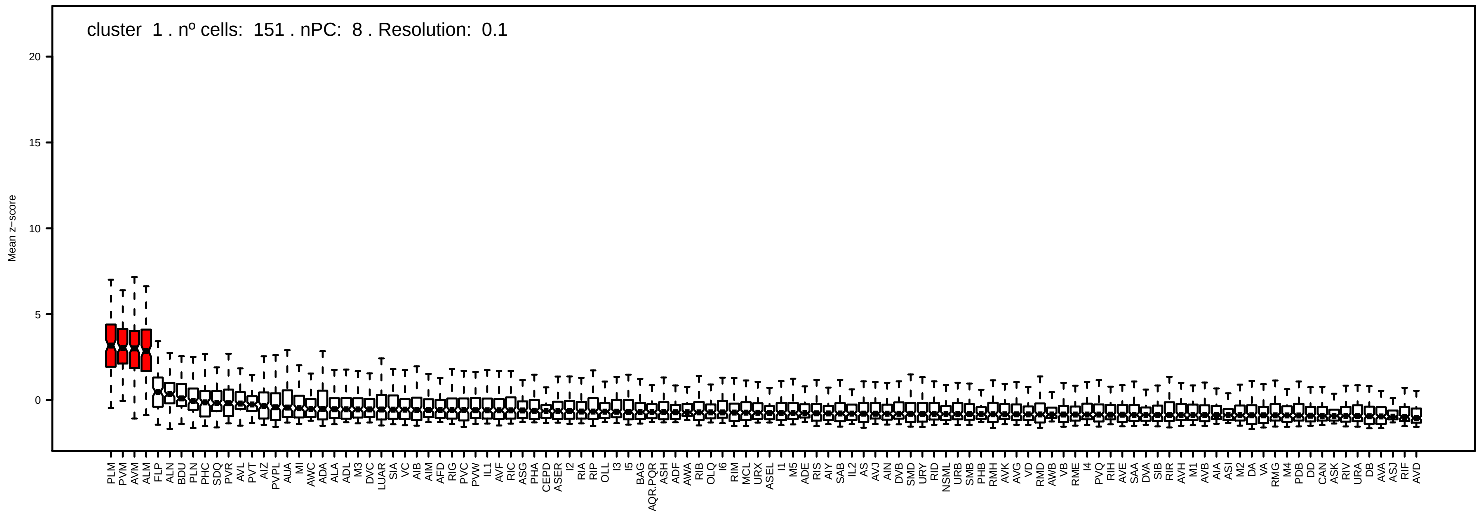
cluster 1 . nº cells: 151 . nPC: 8 . Resolution: 0.1
Mean z−score
20
15
10
5
0

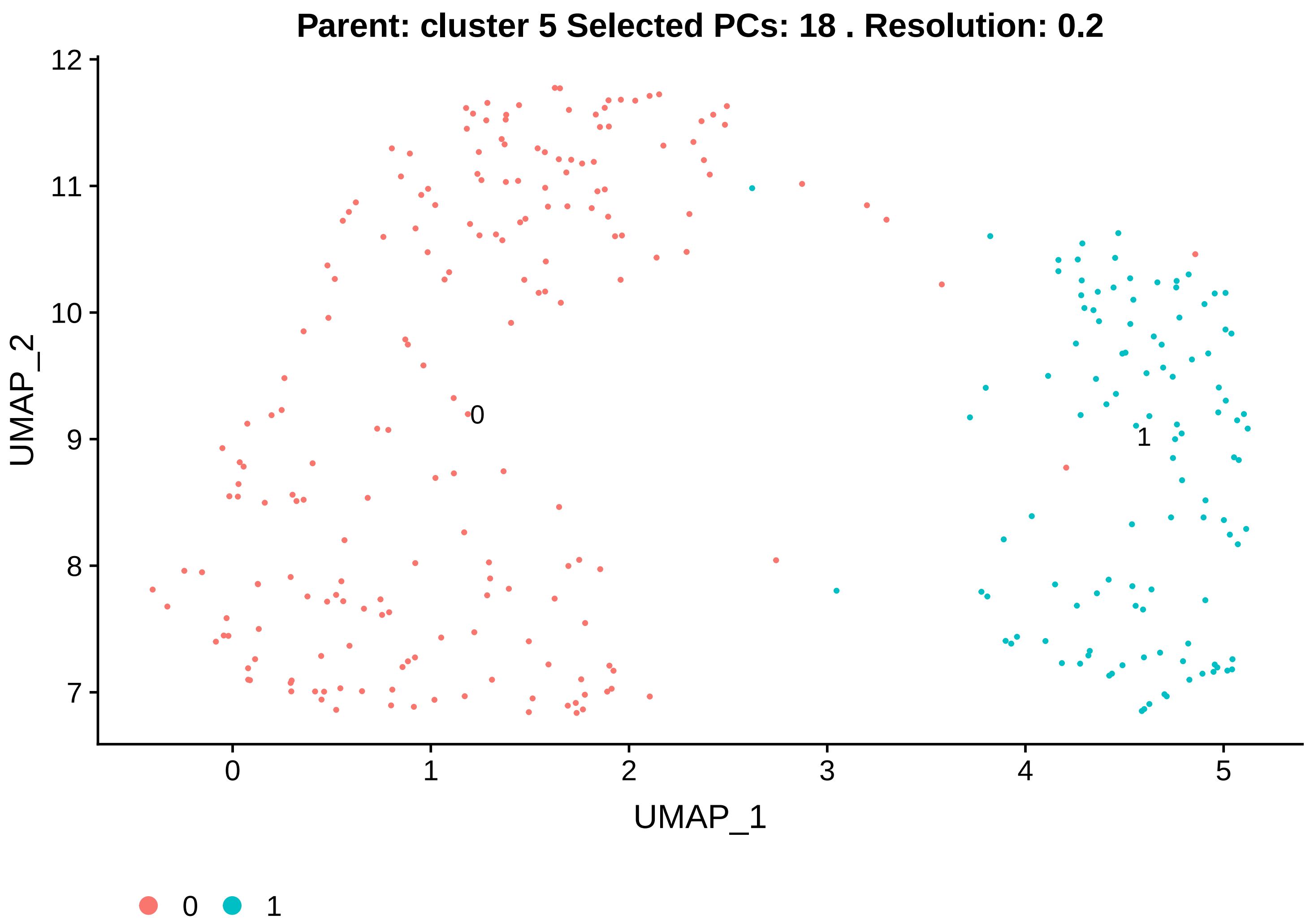

Parent: cluster 5 Selected PCs: 18 . Resolution: 0.2
UMAP_2
UMAP_1
0
1


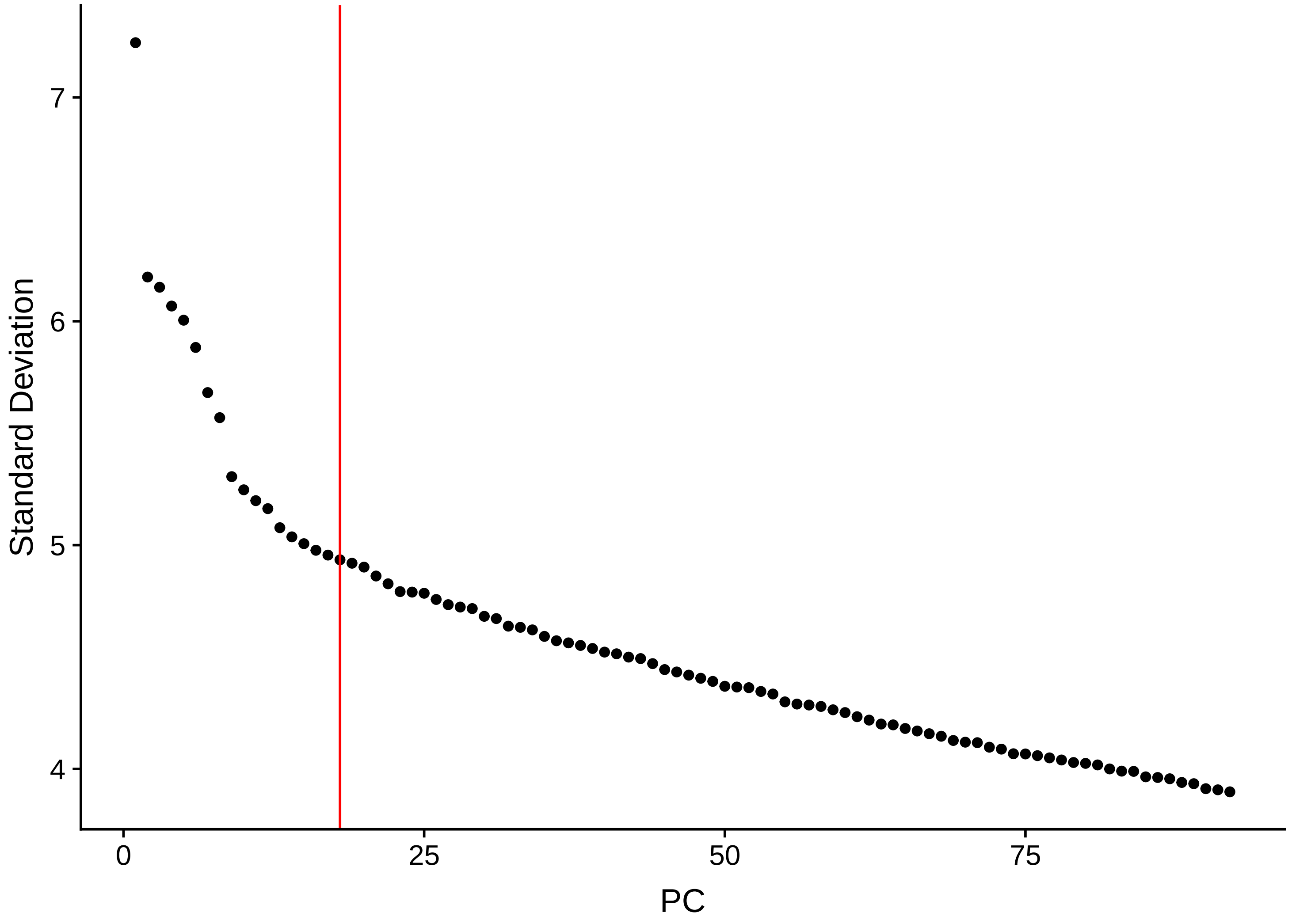

Standard Deviation
PC

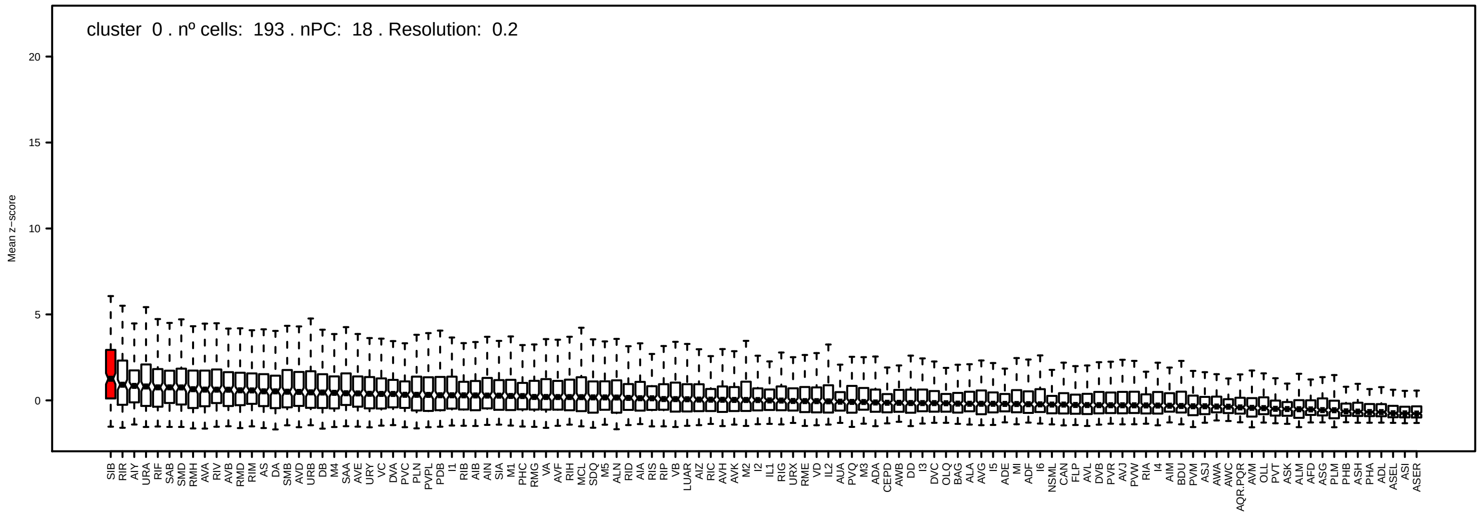
cluster 0 . nº cells: 193 . nPC: 18 . Resolution: 0.2
Mean z−score

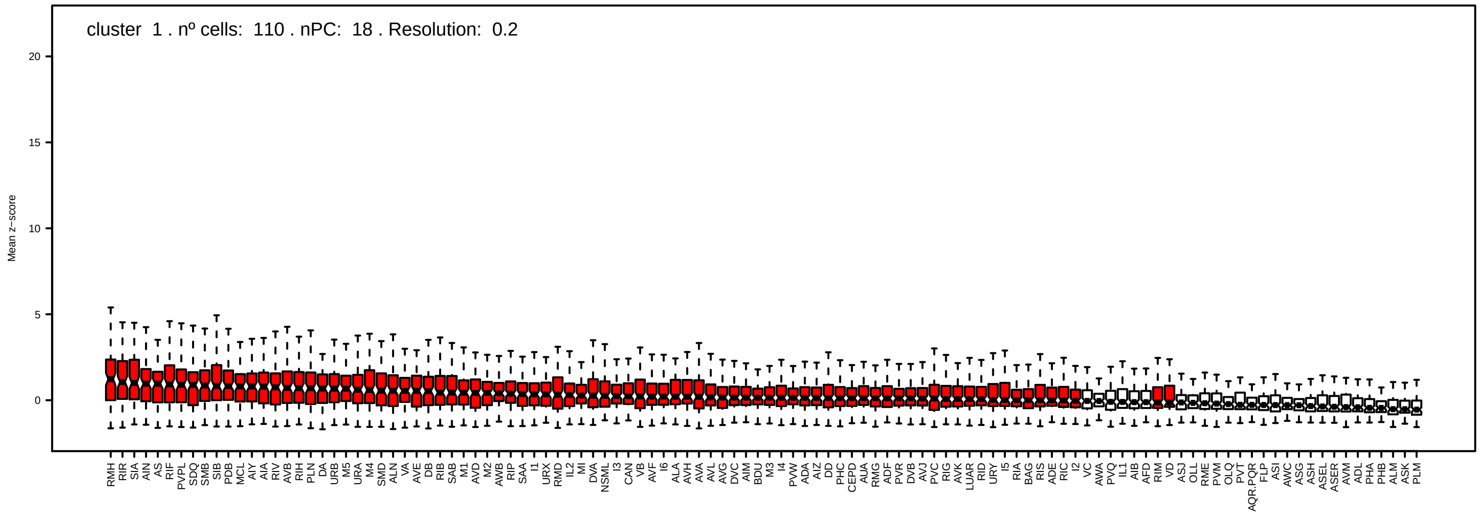
cluster 1 . nº cells: 110 . nPC: 18 . Resolution: 0.2
Mean z−score

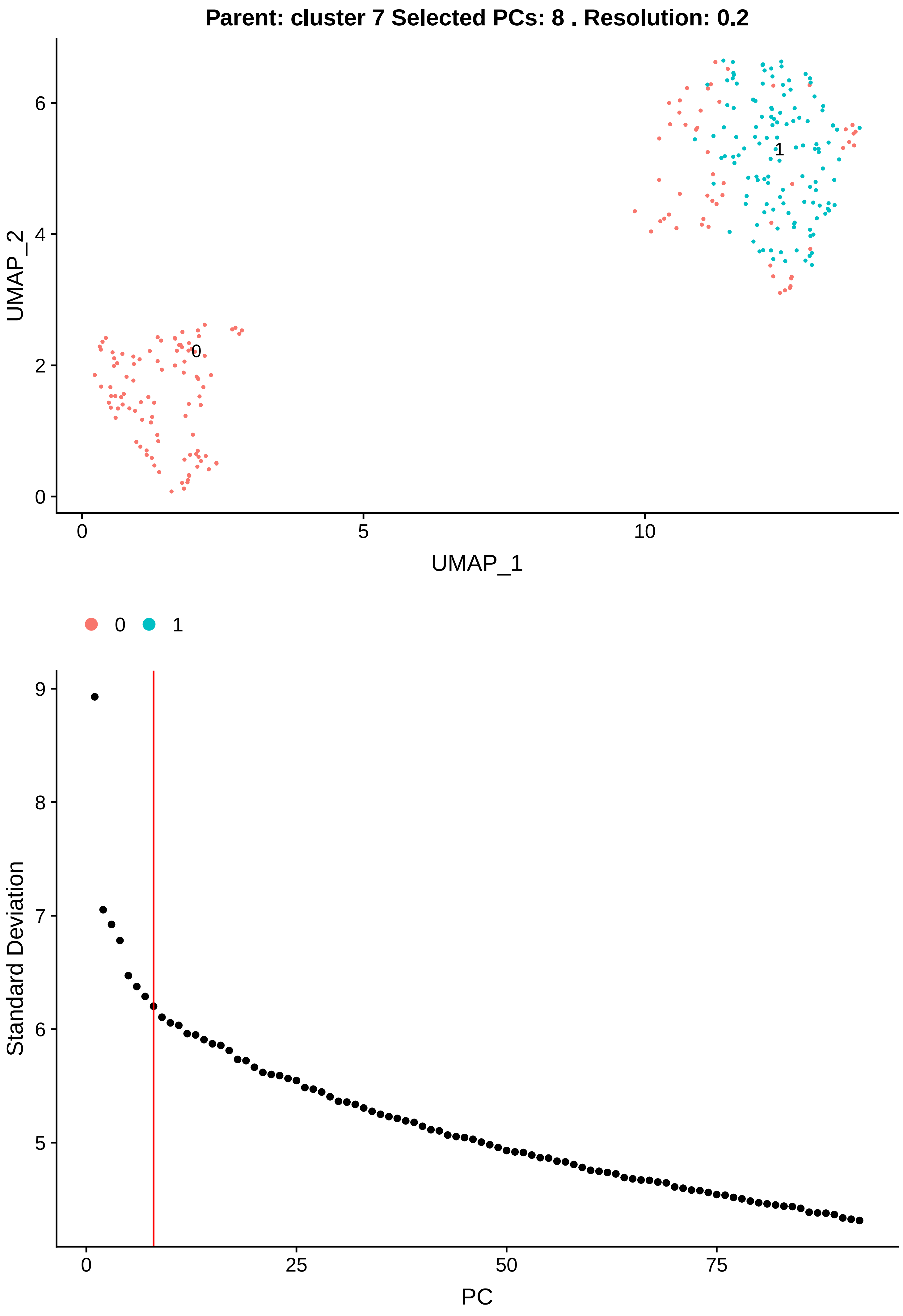

Parent: cluster 7 Selected PCs: 8 . Resolution: 0.2
UMAP_2
UMAP_1
0
1
Standard Deviation
PC

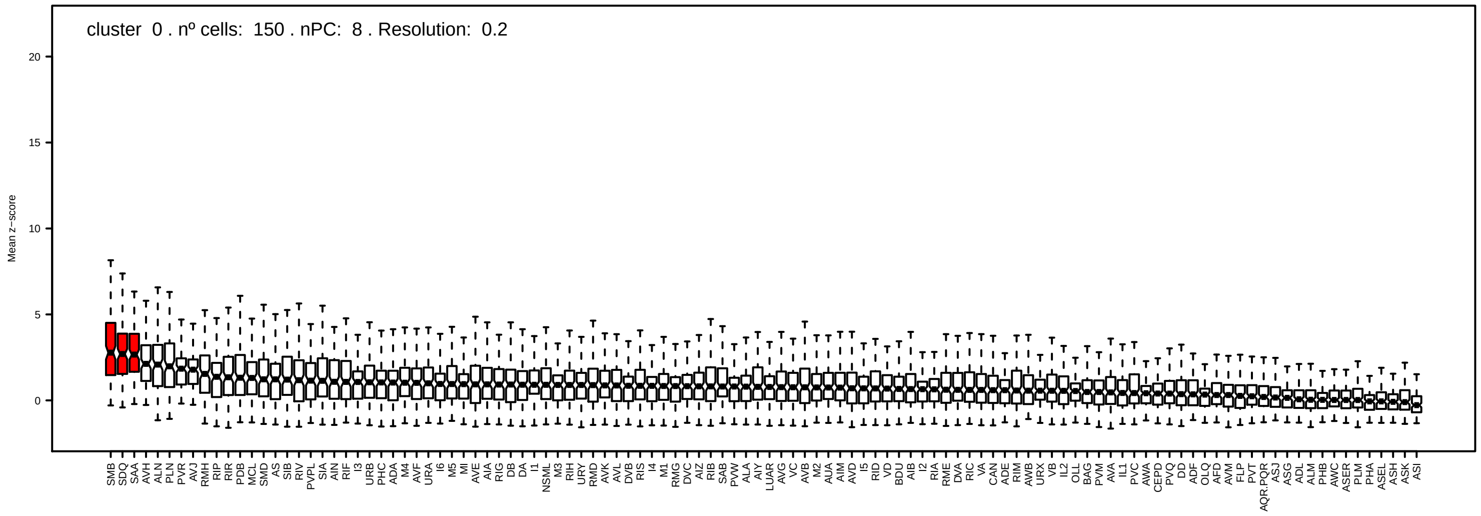
cluster 0 . nº cells: 150 . nPC: 8 . Resolution: 0.2
Mean z−score

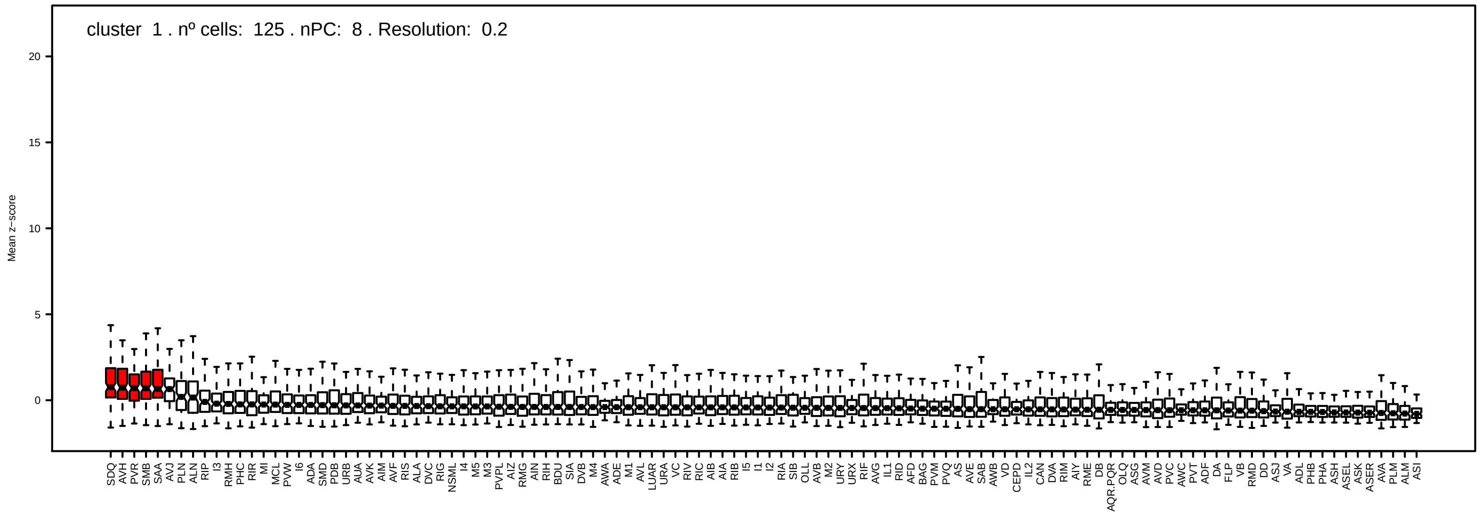
cluster 1 . nº cells: 125 . nPC: 8 . Resolution: 0.2
Mean z−score

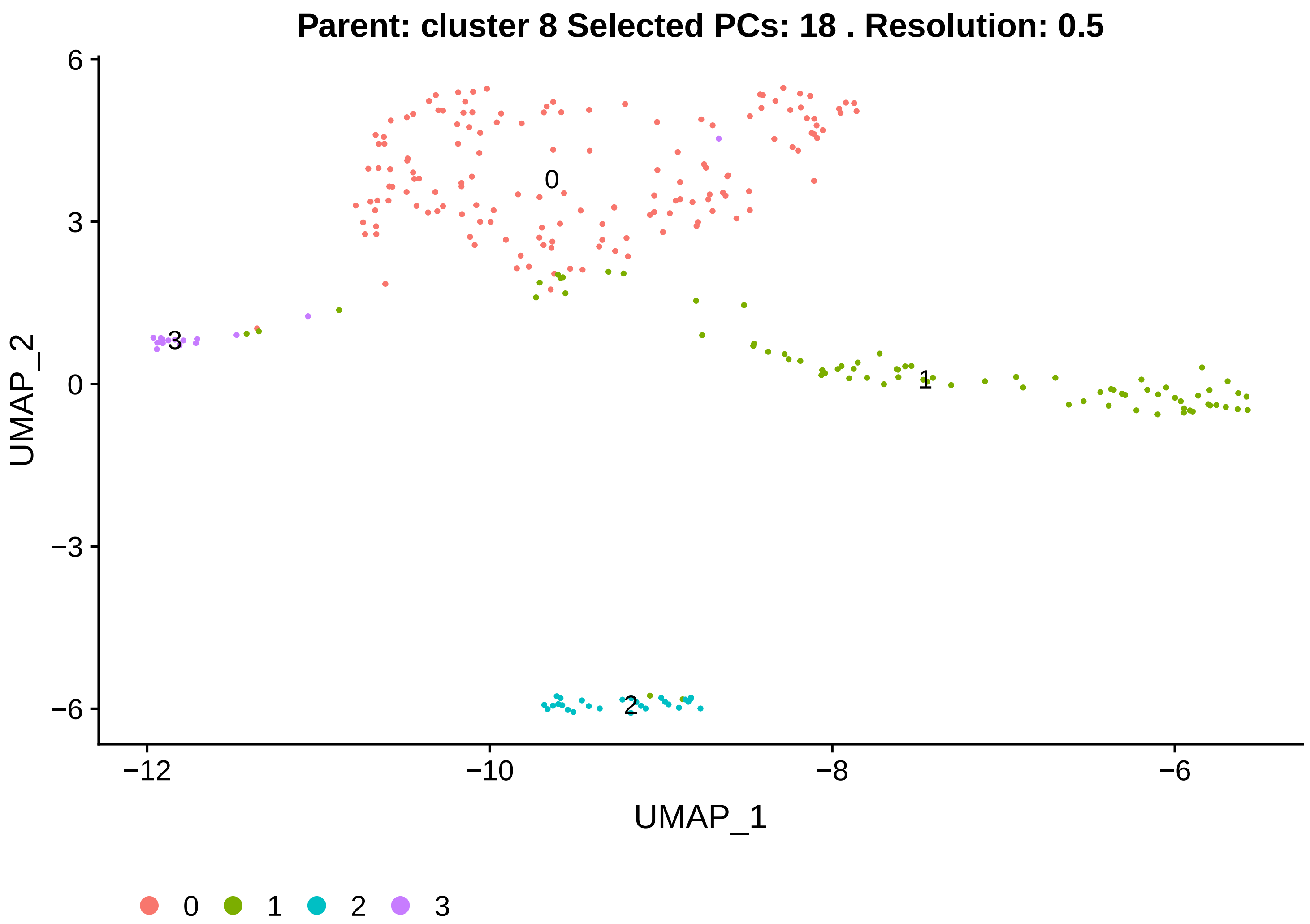

Parent: cluster 8 Selected PCs: 18 . Resolution: 0.5
UMAP_2
UMAP_1
0
1
2
3
6
3
0
−3
−6
−12
−10
−8
−6
0
1
2
3


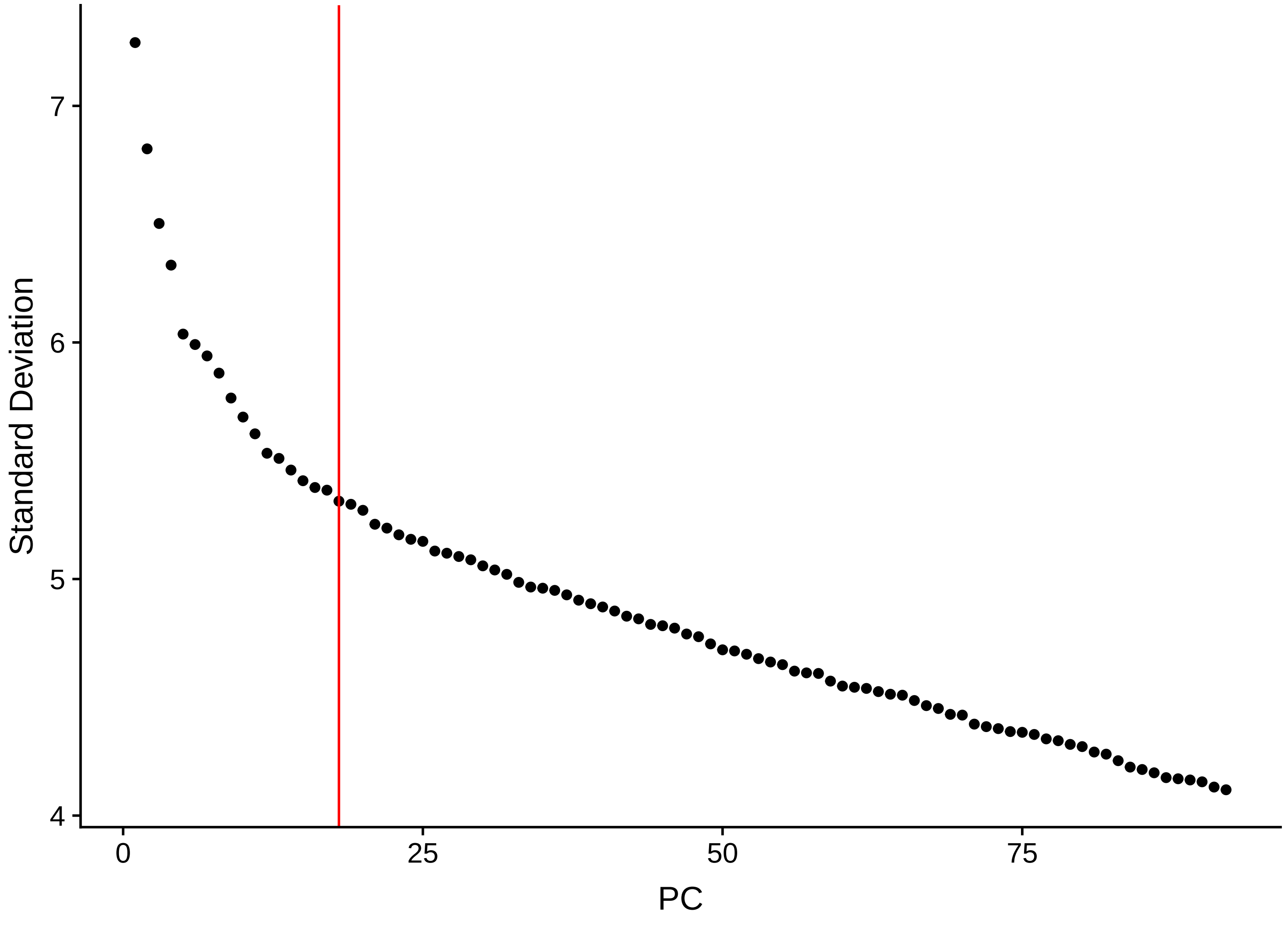

Standard Deviation
PC
7
6
5
4
0
25
50
75

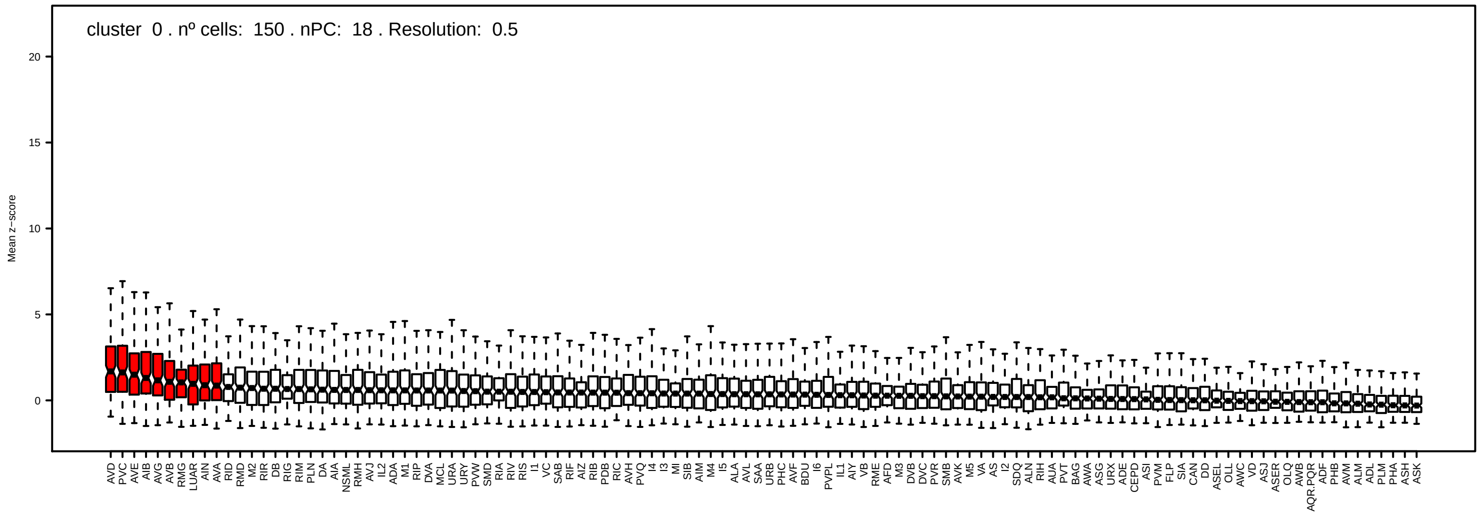
cluster 0 . nº cells: 150 . nPC: 18 . Resolution: 0.5
Mean z−score

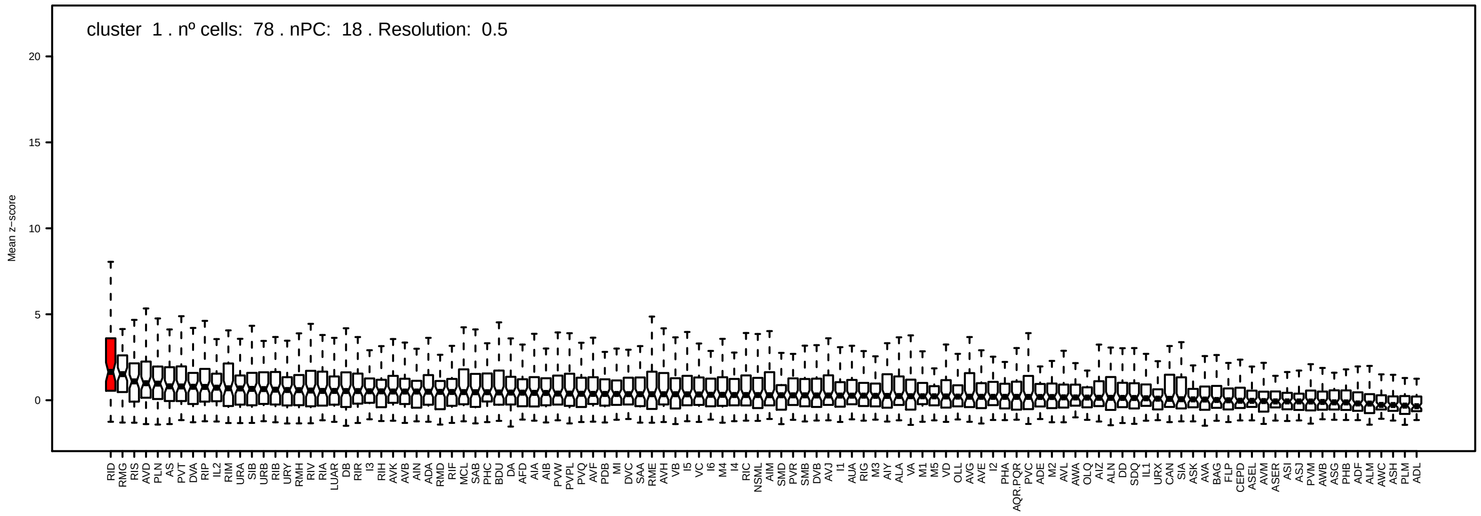
cluster 1 . nº cells: 78 . nPC: 18 . Resolution: 0.5
Mean z−score

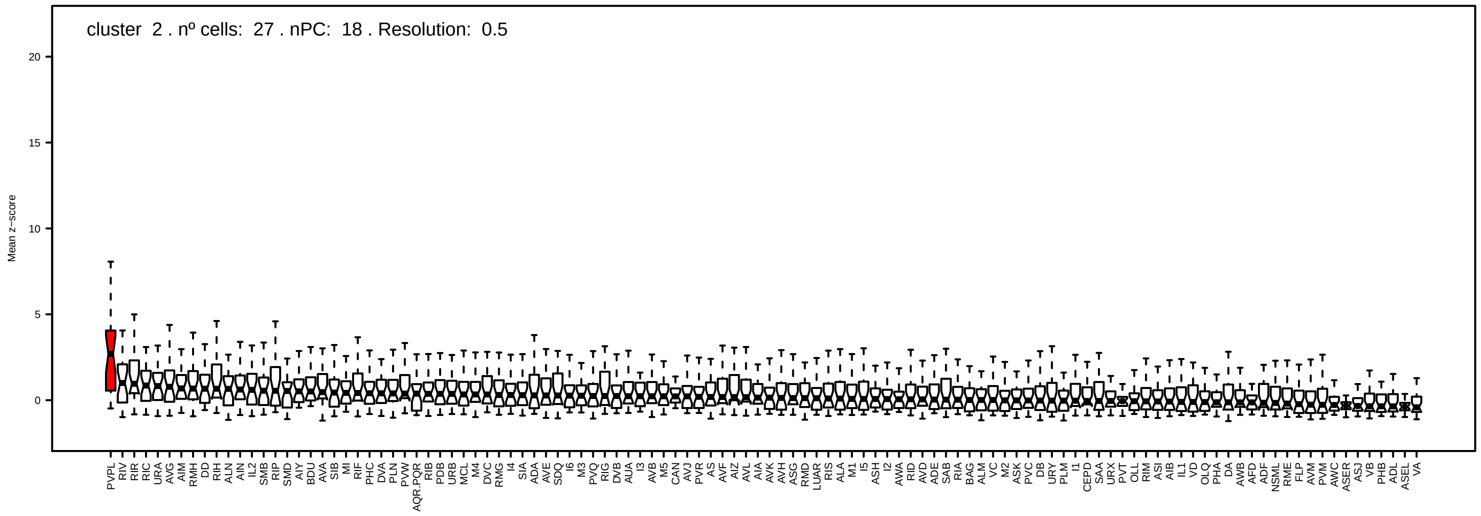
cluster 2 . nº cells: 27 . nPC: 18 . Resolution: 0.5
Mean z−score

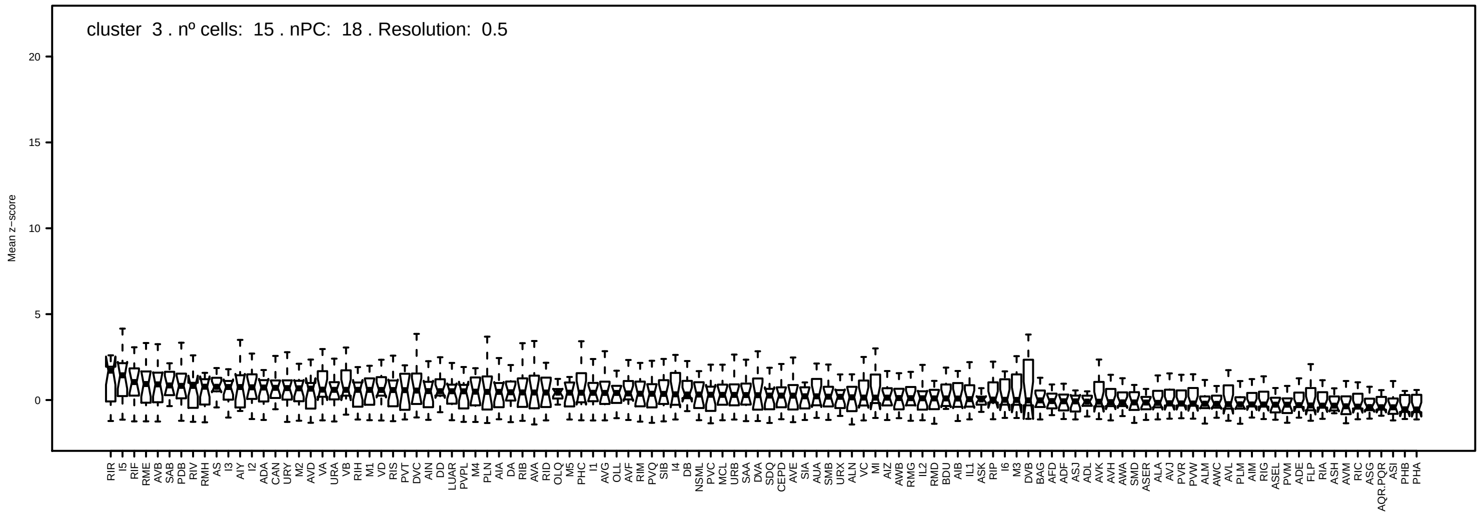
cluster 3 . nº cells: 15 . nPC: 18 . Resolution: 0.5
Mean z−score

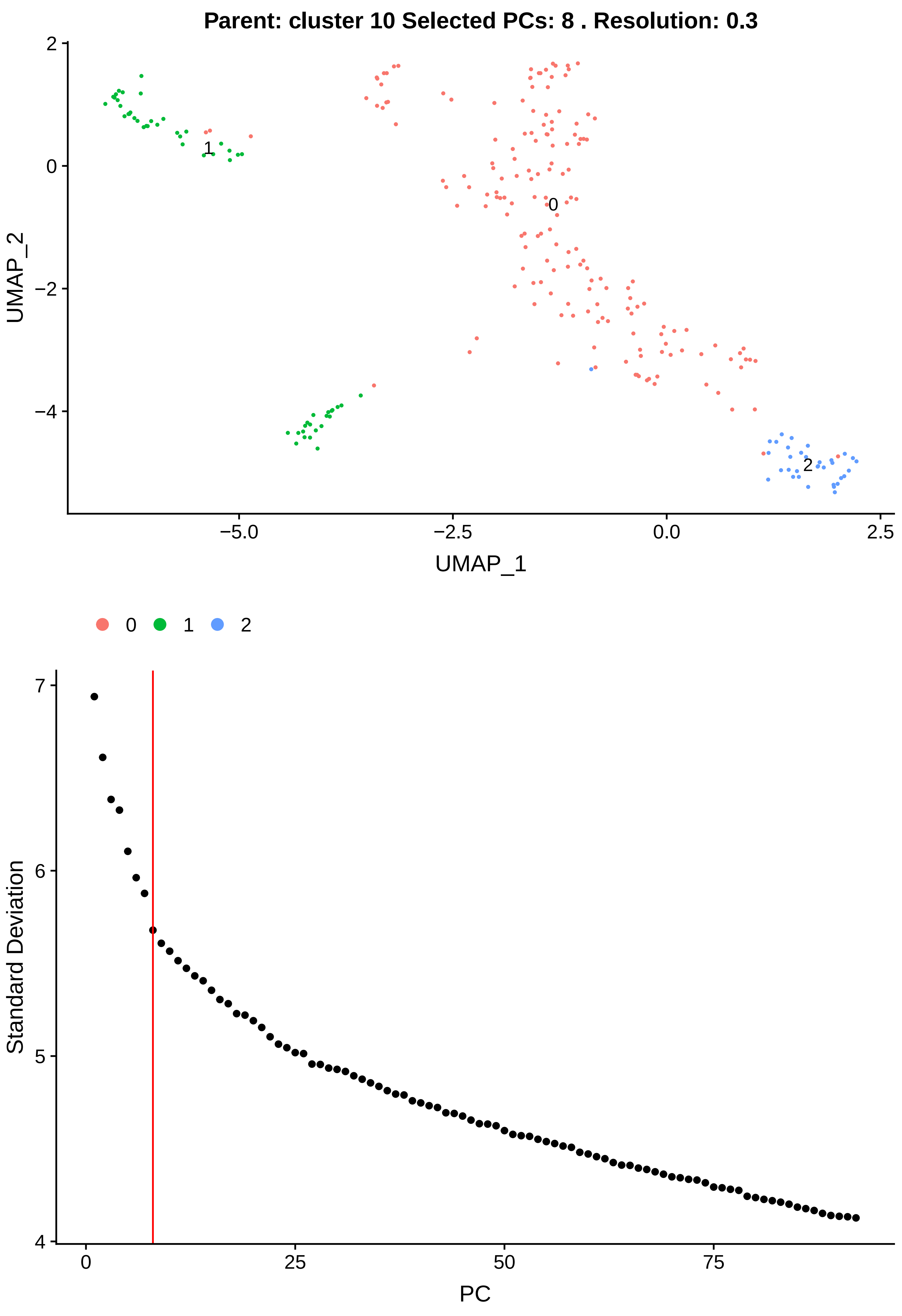

Parent: cluster 10 Selected PCs: 8 . Resolution: 0.3
UMAP_2
UMAP_1
0
1
2
Standard Deviation
PC

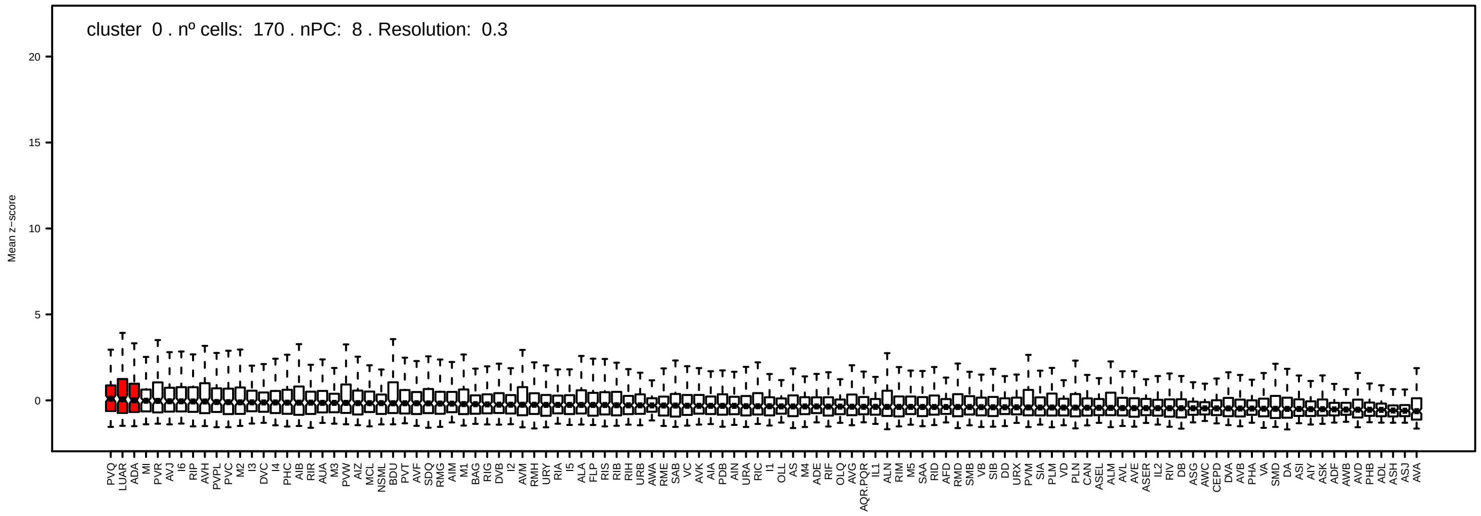
cluster 0 . nº cells: 170 . nPC: 8 . Resolution: 0.3
Mean z-score

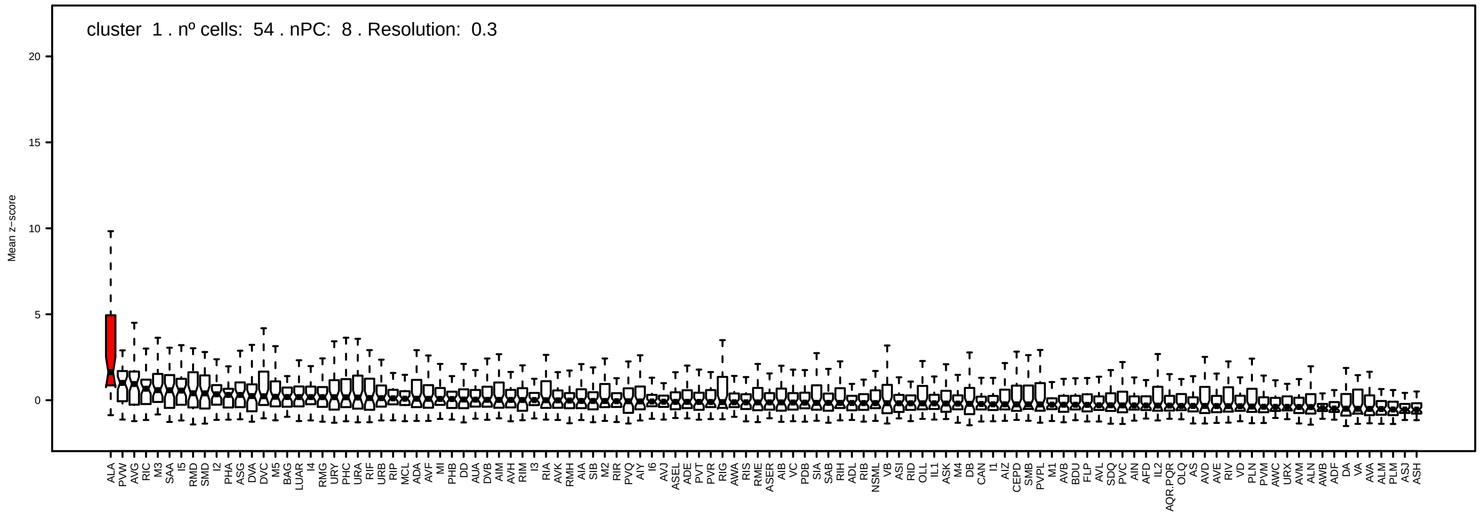
cluster 1 . nº cells: 54 . nPC: 8 . Resolution: 0.3
Mean z-score

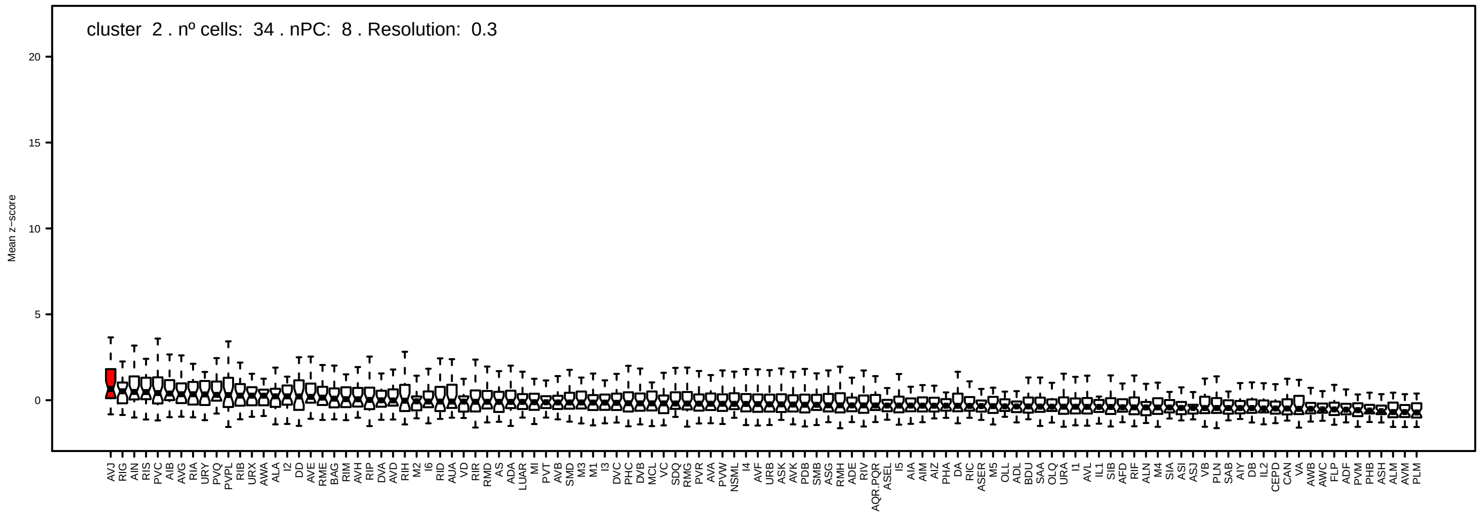
cluster 2 . nº cells: 34 . nPC: 8 . Resolution: 0.3
Mean z-score

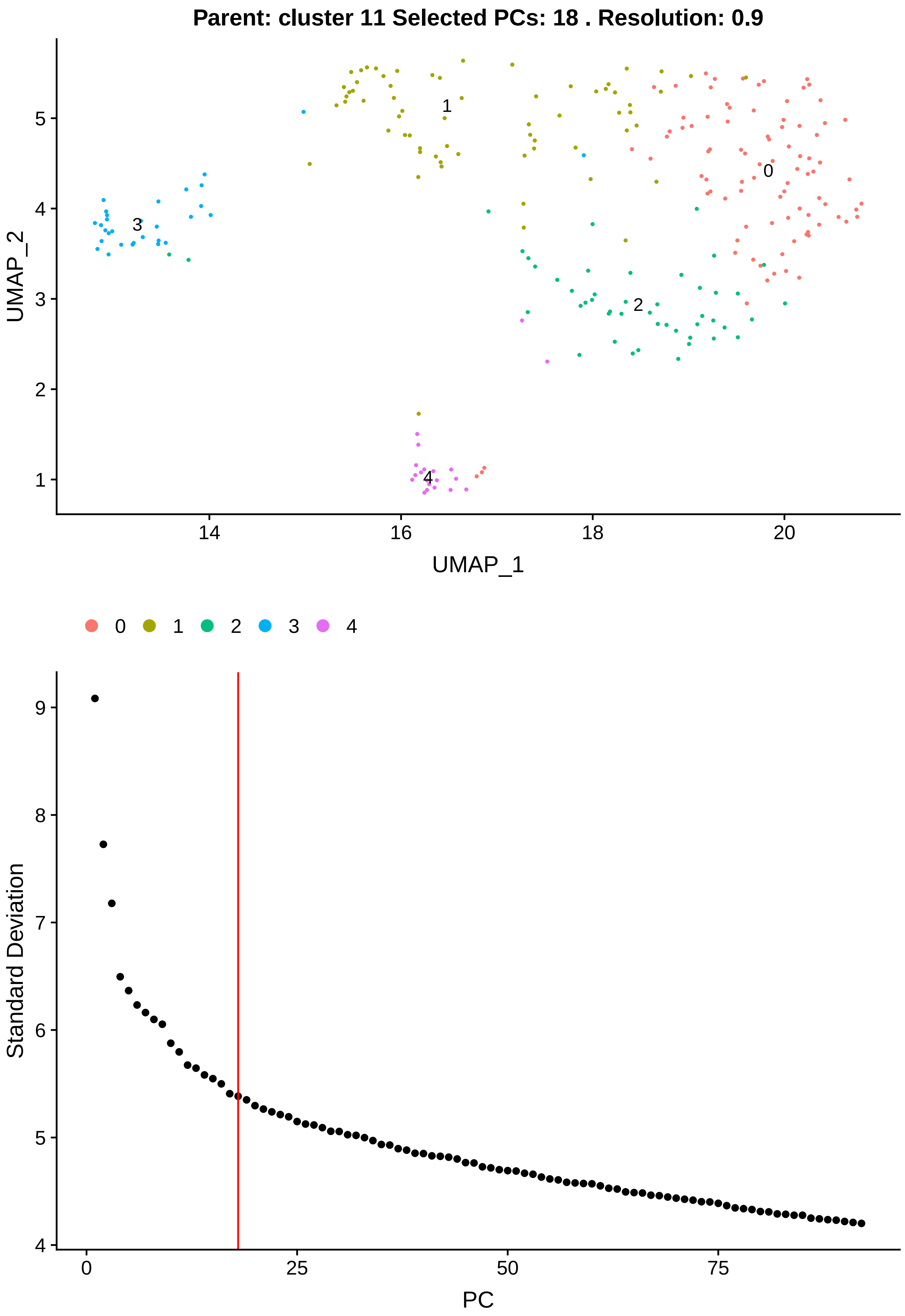
Parent: cluster 11 Selected PCs: 18 . Resolution: 0.9
UMAP_2
UMAP_1
0
1
2
3
4
Standard Deviation
PC

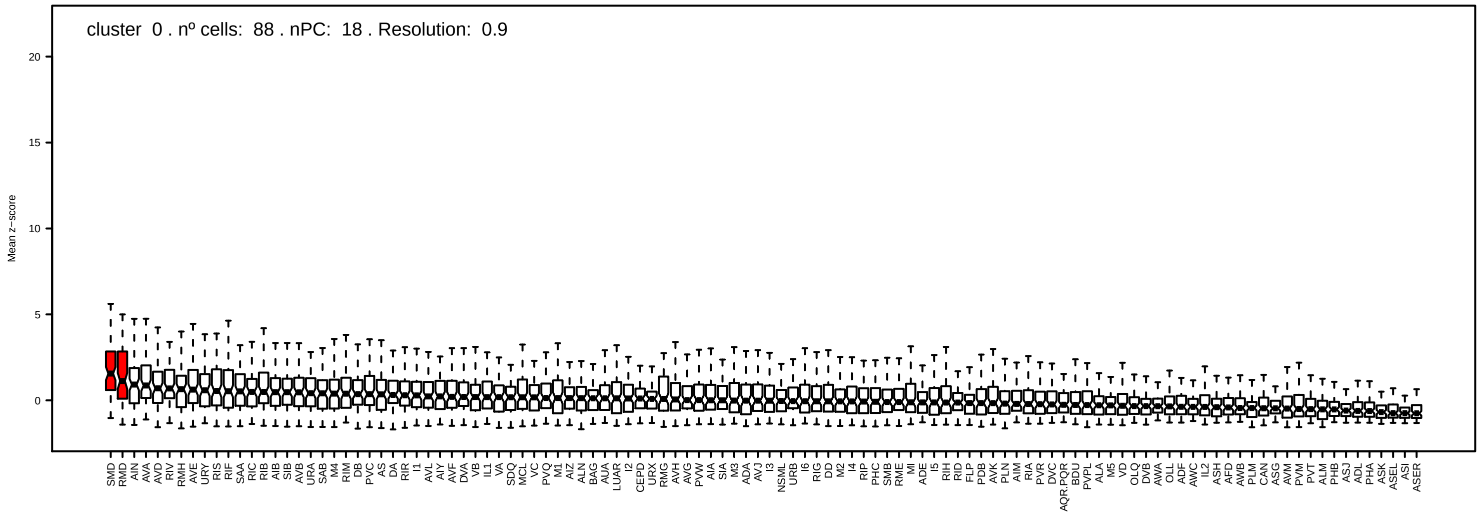
cluster 0 . nº cells: 88 . nPC: 18 . Resolution: 0.9
Mean z−score
0
5
10
15
20

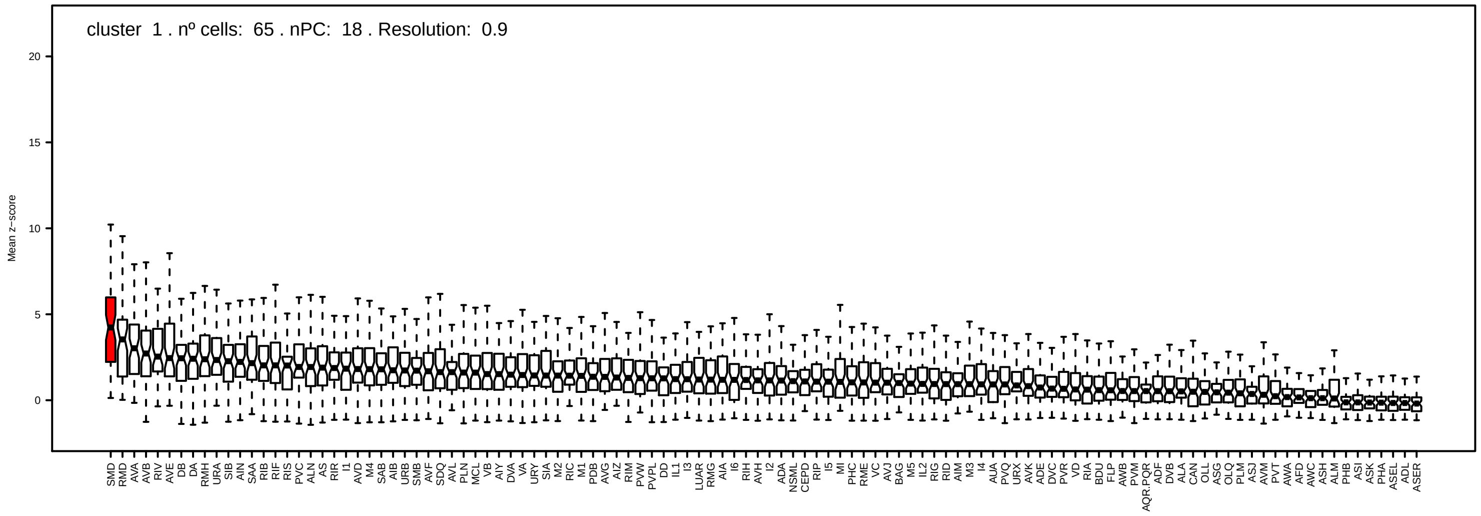
cluster 1 . nº cells: 65 . nPC: 18 . Resolution: 0.9
Mean z−score
0
5
10
15
20

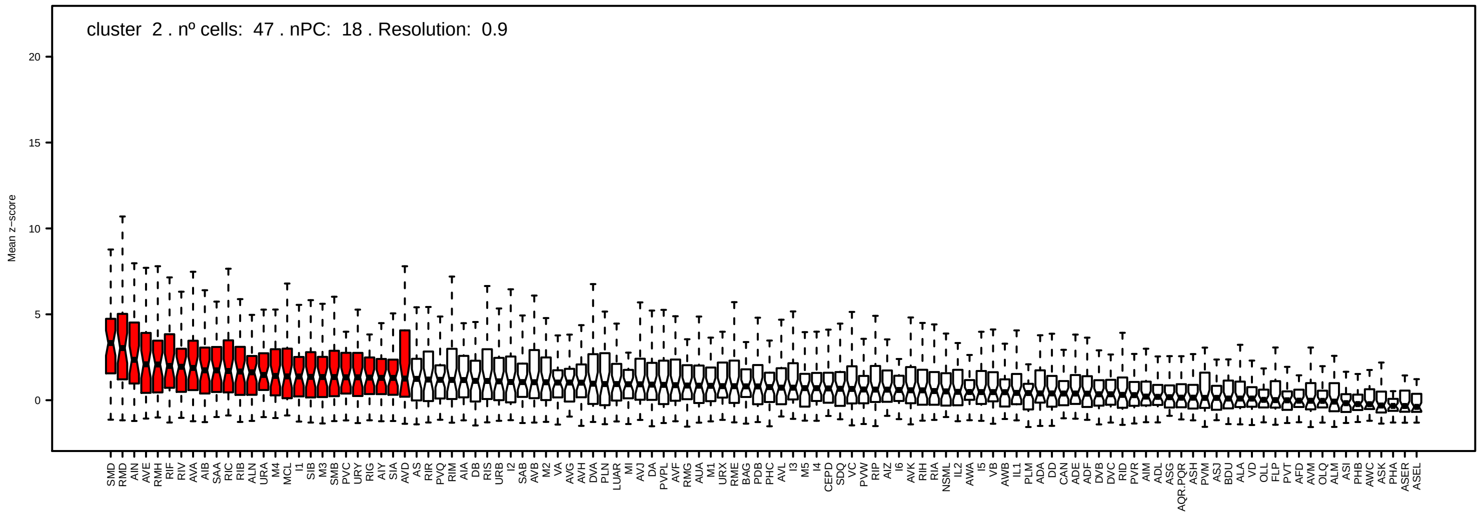
cluster 2 . nº cells: 47 . nPC: 18 . Resolution: 0.9
Mean z−score
0
5
10
15
20

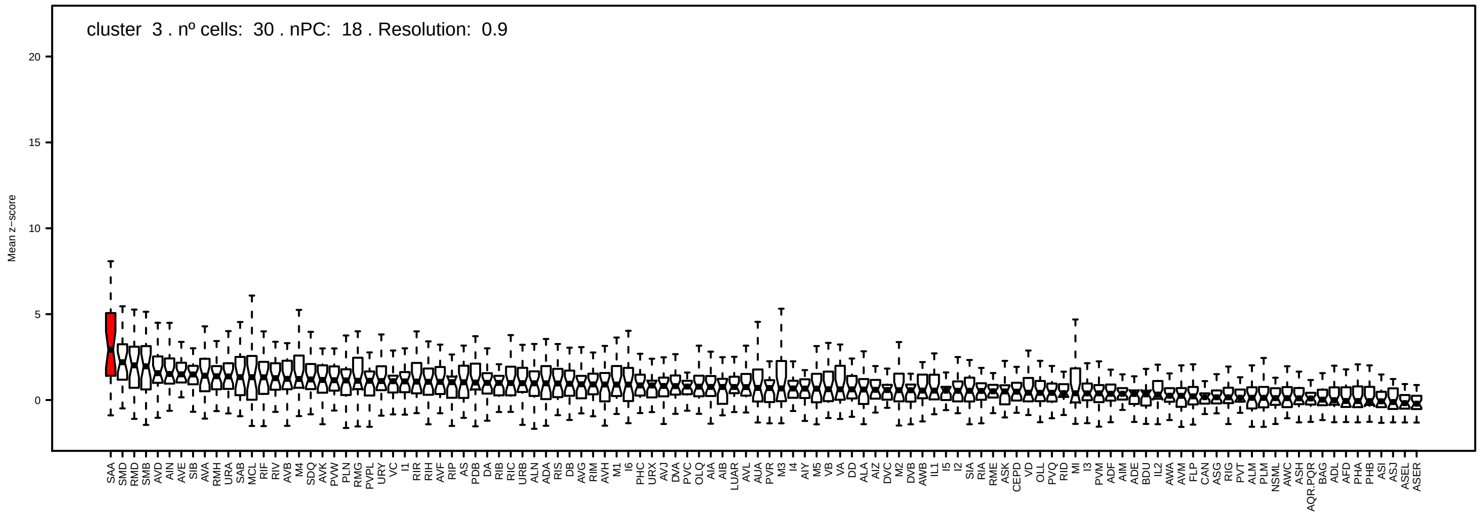
cluster 3 . nº cells: 30 . nPC: 18 . Resolution: 0.9
Mean z−score
0
5
10
15
20

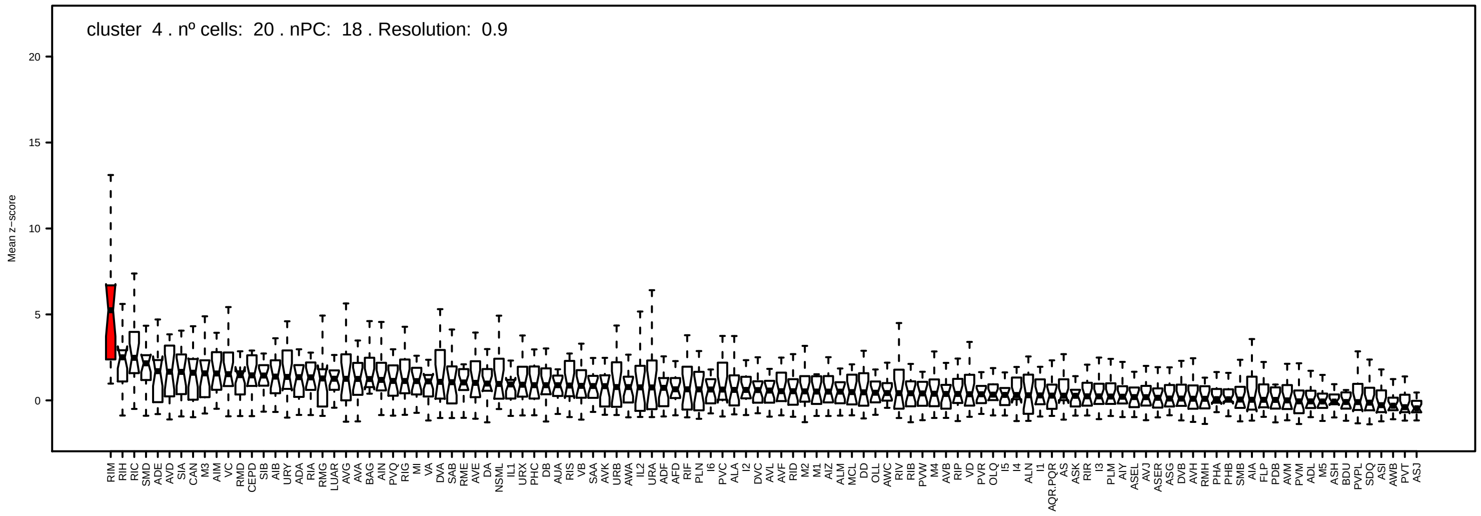
cluster 4 . nº cells: 20 . nPC: 18 . Resolution: 0.9
Mean z−score
0
5
10
15
20
RIM RIH RIC SMD ADE AVD SIA CAN M3 AIM VC RMD CEPD SIB AIB URY ADA RIA RMG LUAR AVG AVA BAG AIN PVQ RIG MI VA DVA SAB RME AVE DA NSML IL1 URX PHC DB AUA RIS VB SAA AVK URB AWA IL2 URA ADF AFD RIF PLN I6 PVC ALA I2 DVC AVL AVF RID M2 M1 AIZ ALM MCL DD OLL AWC RIV RIB PVW M4 AVB RIP VD PVR OLQ I5 I4 ALN I1 AQR.PQR AS ASK RIR I3 PLM AIY ASEL AVJ ASER ASG DVB AVH RMH PHA PHB SMB AIA FLP PDB AVM PVM ADL M5 ASH BDU PVPL SDQ ASI AWB PVT ASJ

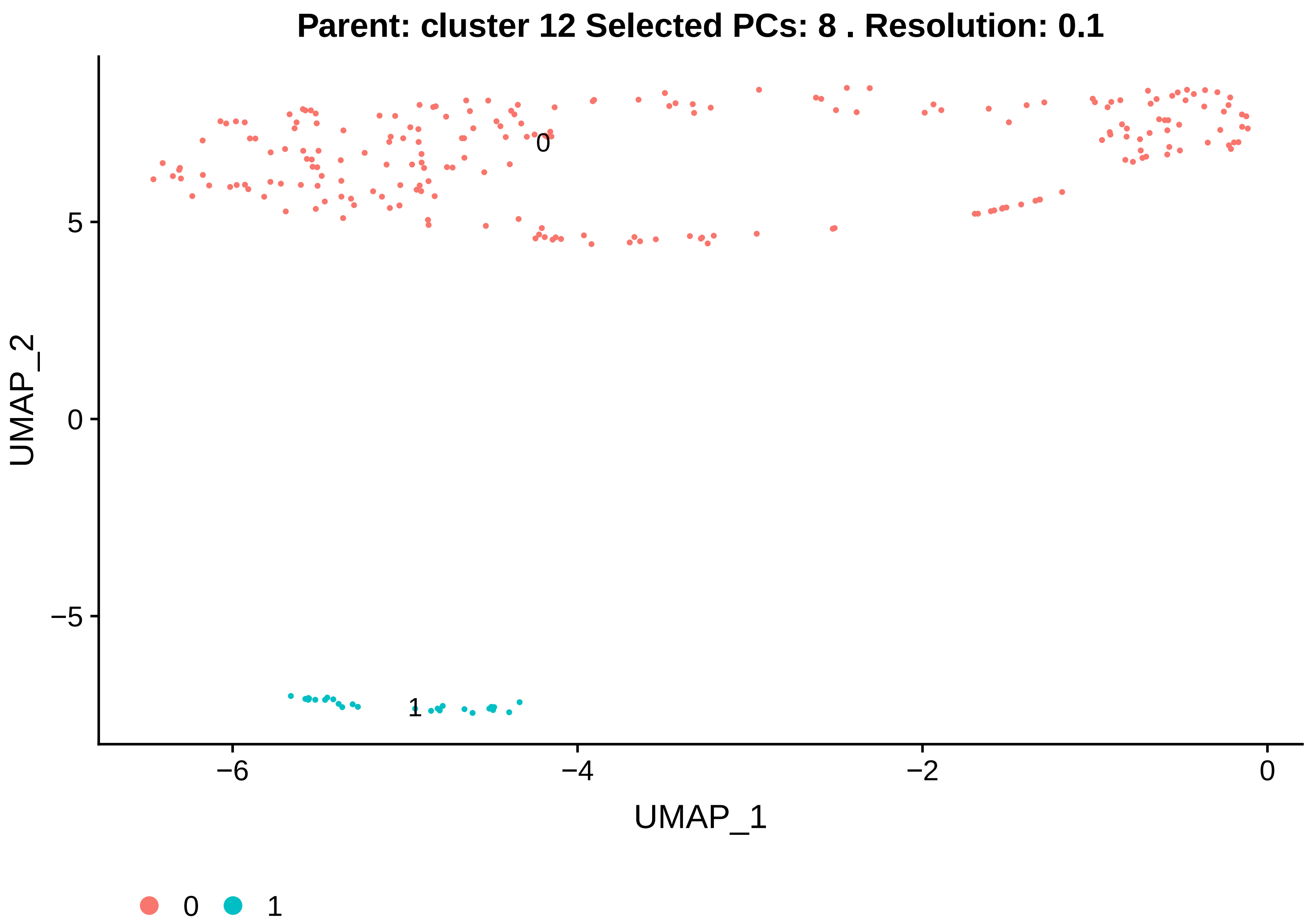

Parent: cluster 12 Selected PCs: 8 . Resolution: 0.1
UMAP_2
UMAP_1
0
1
5
0
−5
−6
−4
−2
0


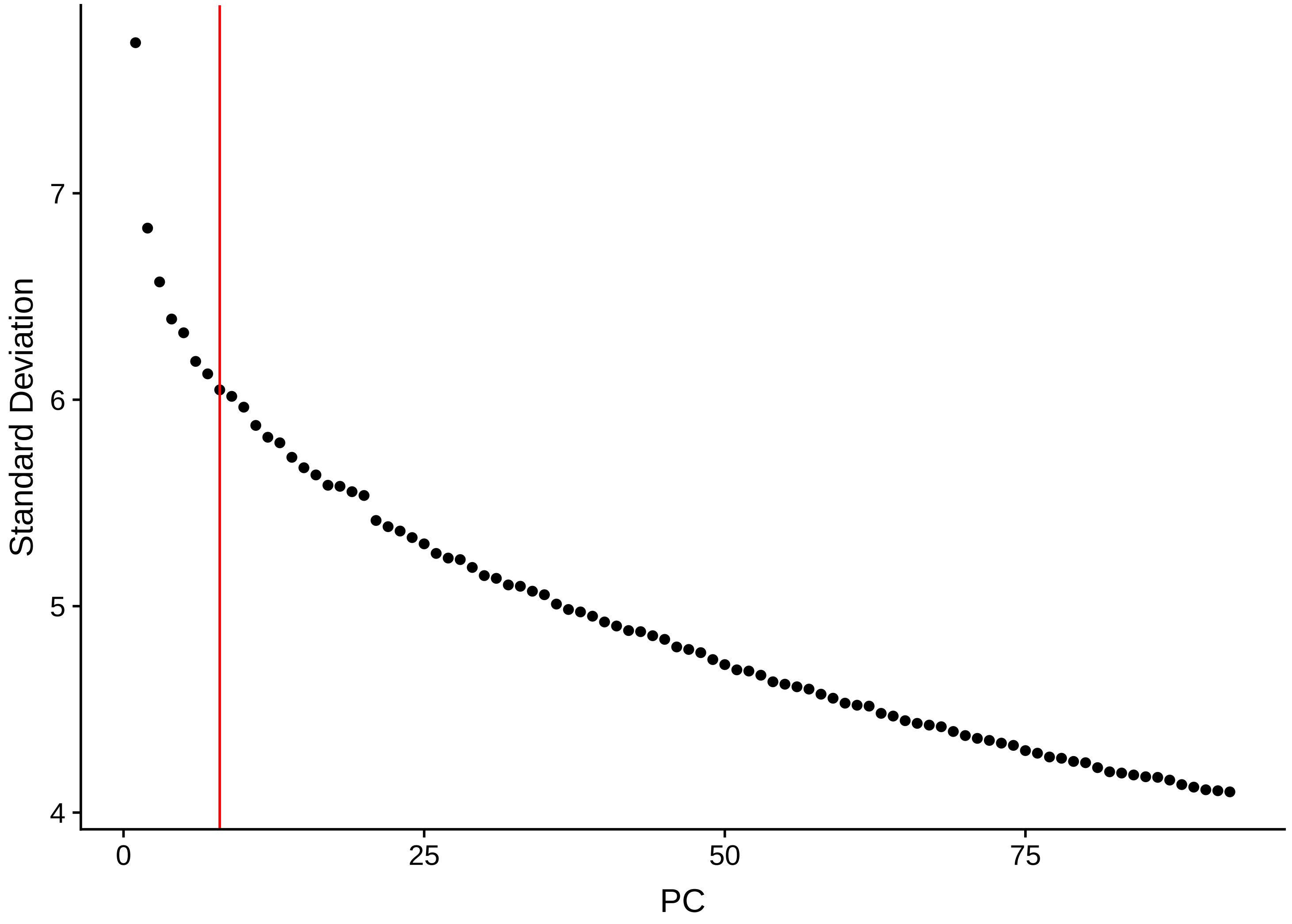

Standard Deviation
PC
7
6
5
4
0
25
50
75

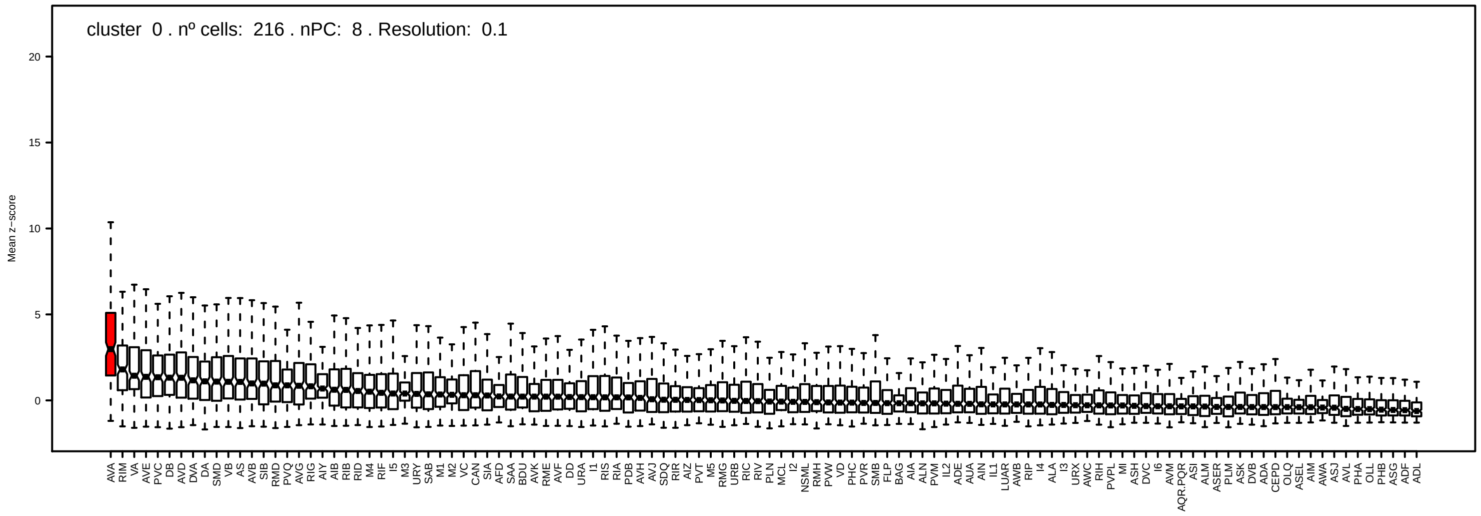
cluster 0 . nº cells: 216 . nPC: 8 . Resolution: 0.1
Mean z−score
20
15
10
5
0

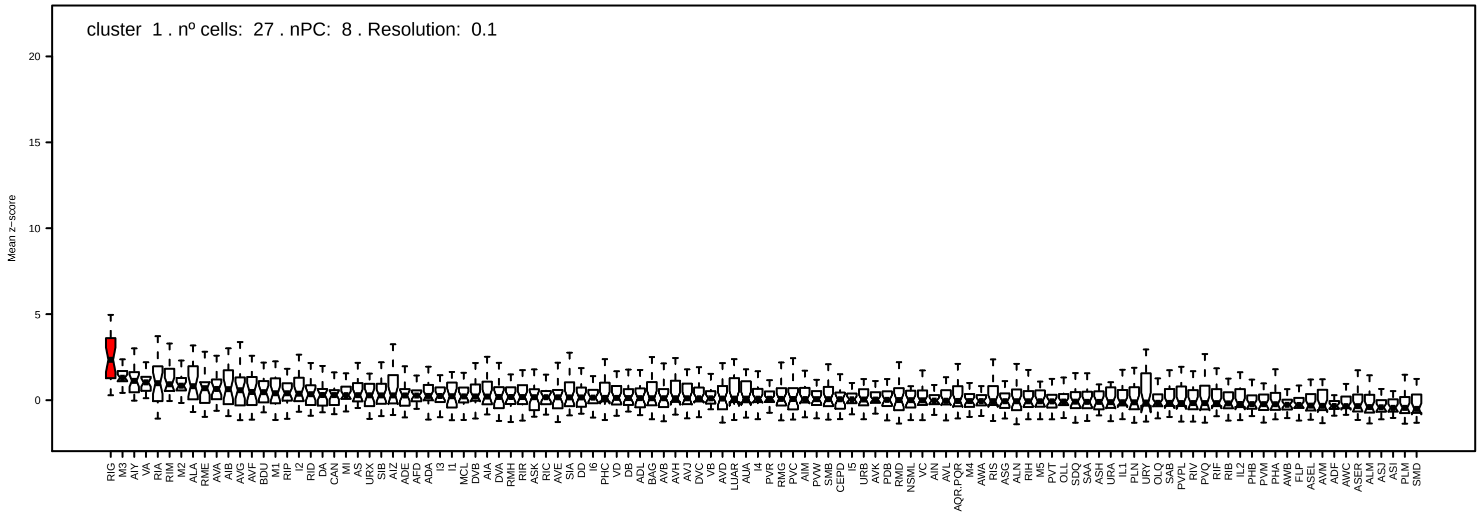
cluster 1 . nº cells: 27 . nPC: 8 . Resolution: 0.1
Mean z−score
20
15
10
5
0

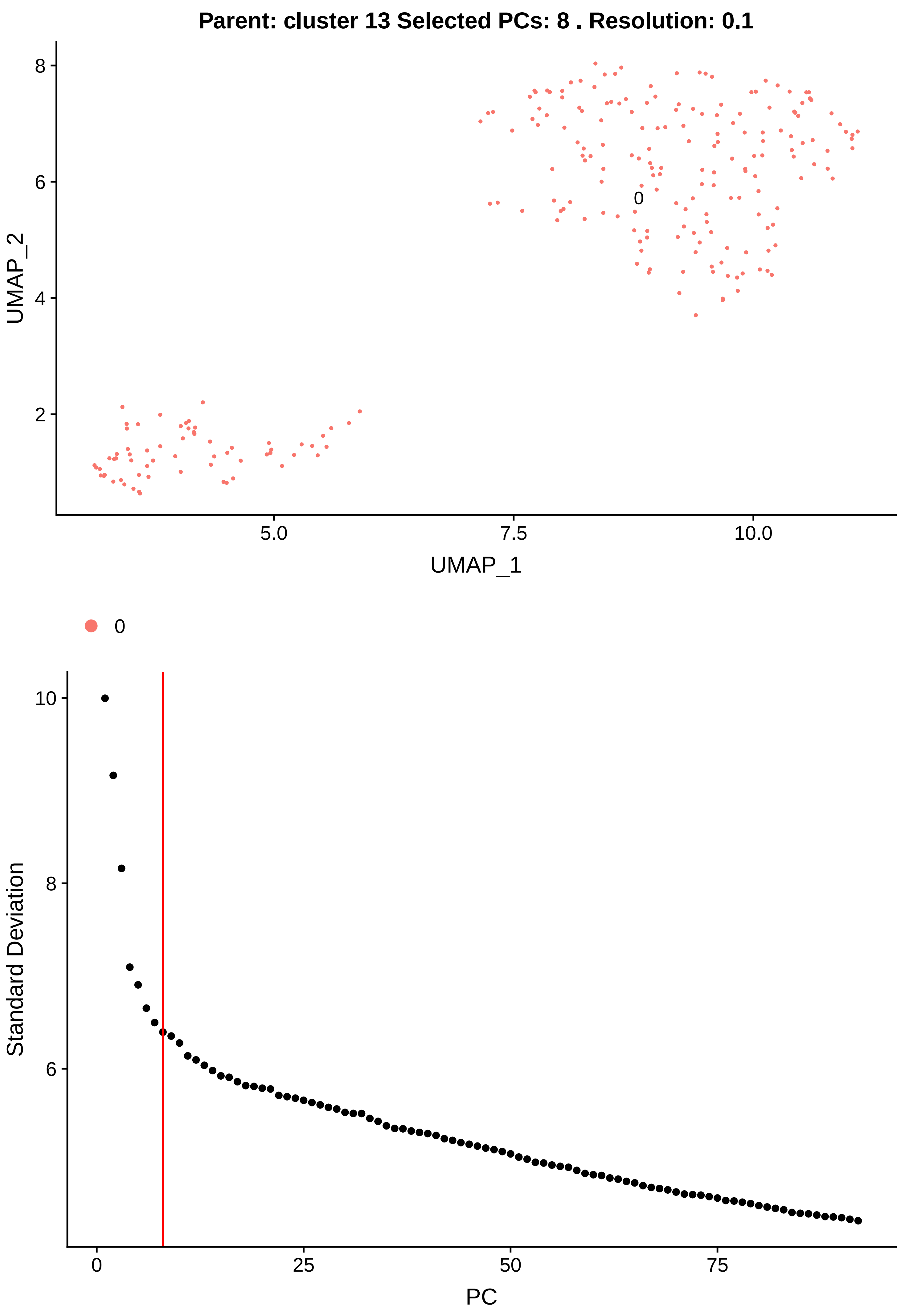

Parent: cluster 13 Selected PCs: 8 . Resolution: 0.1
UMAP_2
UMAP_1
0
Standard Deviation
PC

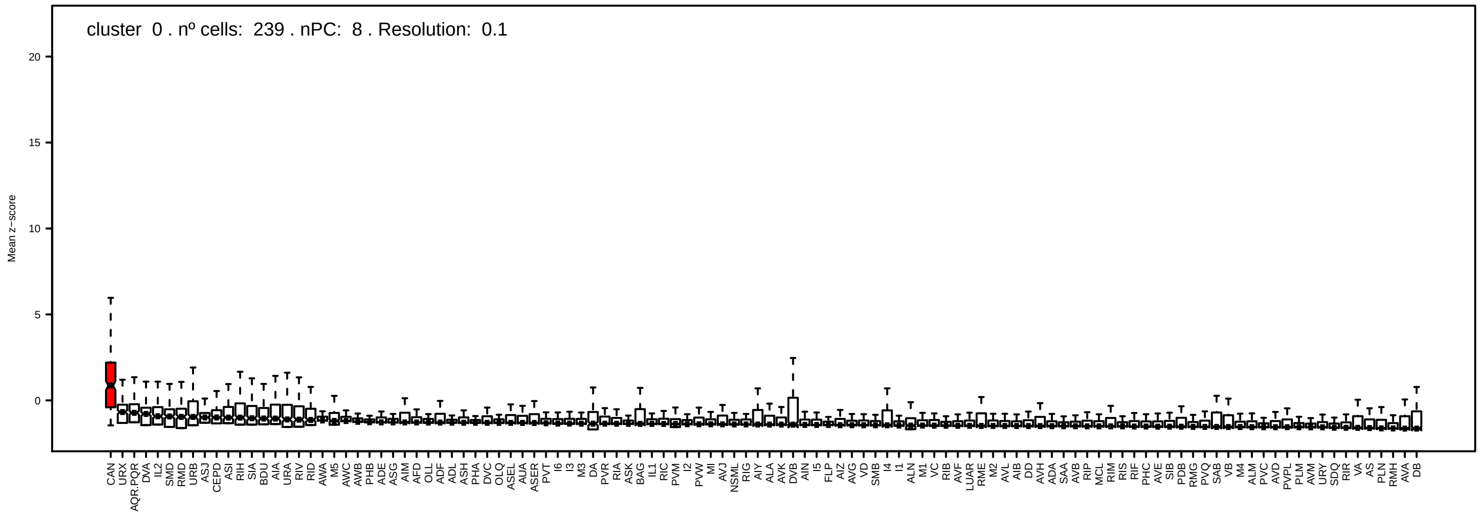
cluster 0 . nº cells: 239 . nPC: 8 . Resolution: 0.1
Mean z−score
20
15
10
5
0
CAN URX AQR.PQR DVA IL2 SMD RMD URB ASJ CEPD ASI RIH SIA BDU AIA URA RIV RID AWA M5 AWC AWB PHB ADE ASG AIM AFD OLL ADF ADL ASH PHA DVC OLQ ASEL AUA ASER PVT I6 I3 M3 DA PVR RIA ASK BAG IL1 RIC PVM I2 PVW MI AVJ NSML RIG AIY ALA AVK DVB AIN I5 FLP AIZ AVG VD SMB I4 I1 ALN M1 VC RIB AVF LUAR RME M2 AVL AIB DD AVH ADA SAA AVB RIP MCL RIM RIS RIF PHC AVE SIB PDB RMG PVQ SAB VB M4 ALM PVC AVD PVPL PLM AVM URY SDQ RIR VA AS PLN RMH AVA DB

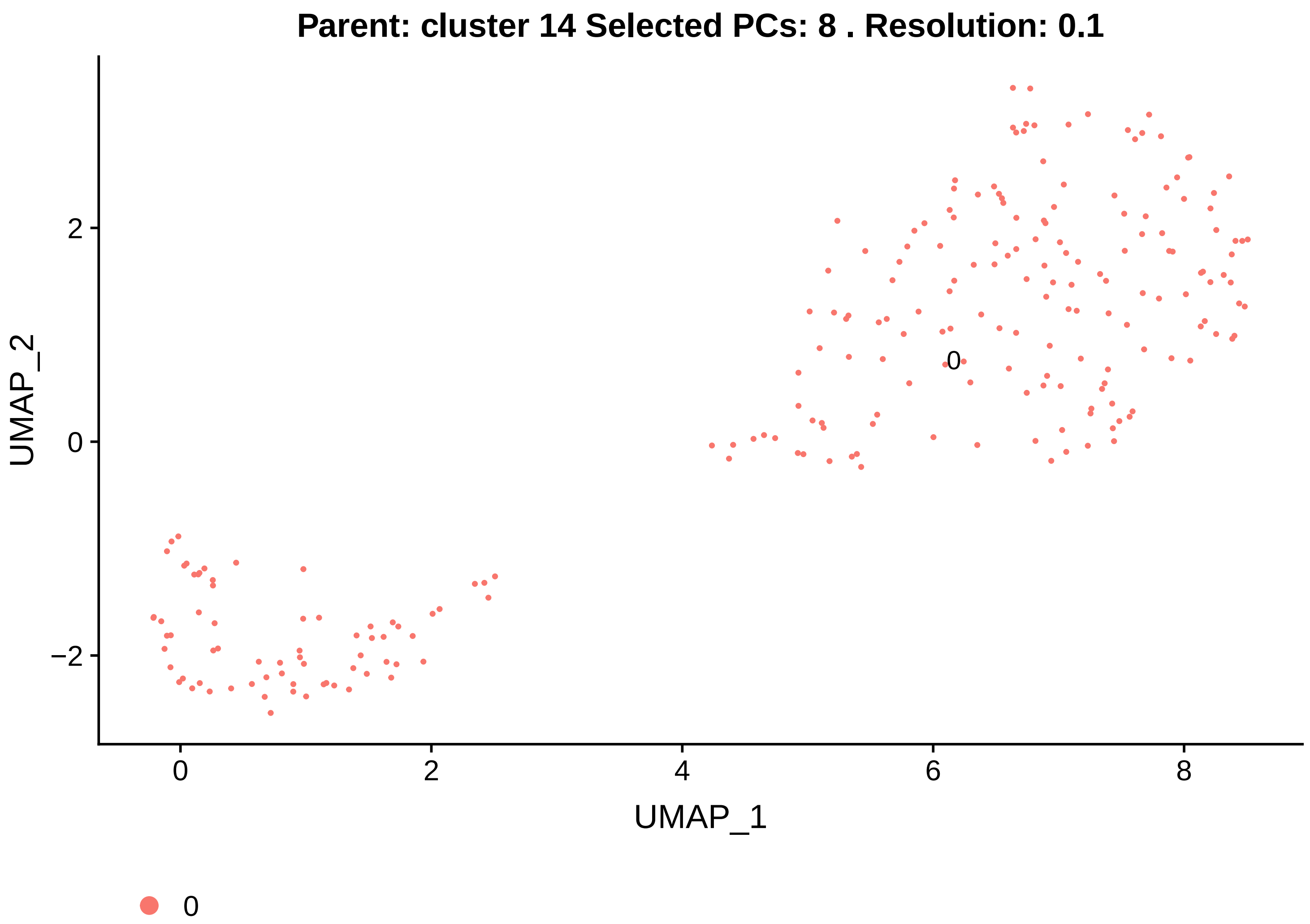

Parent: cluster 14 Selected PCs: 8 . Resolution: 0.1
UMAP_2
UMAP_1
0
2
−2
4
6
8
0


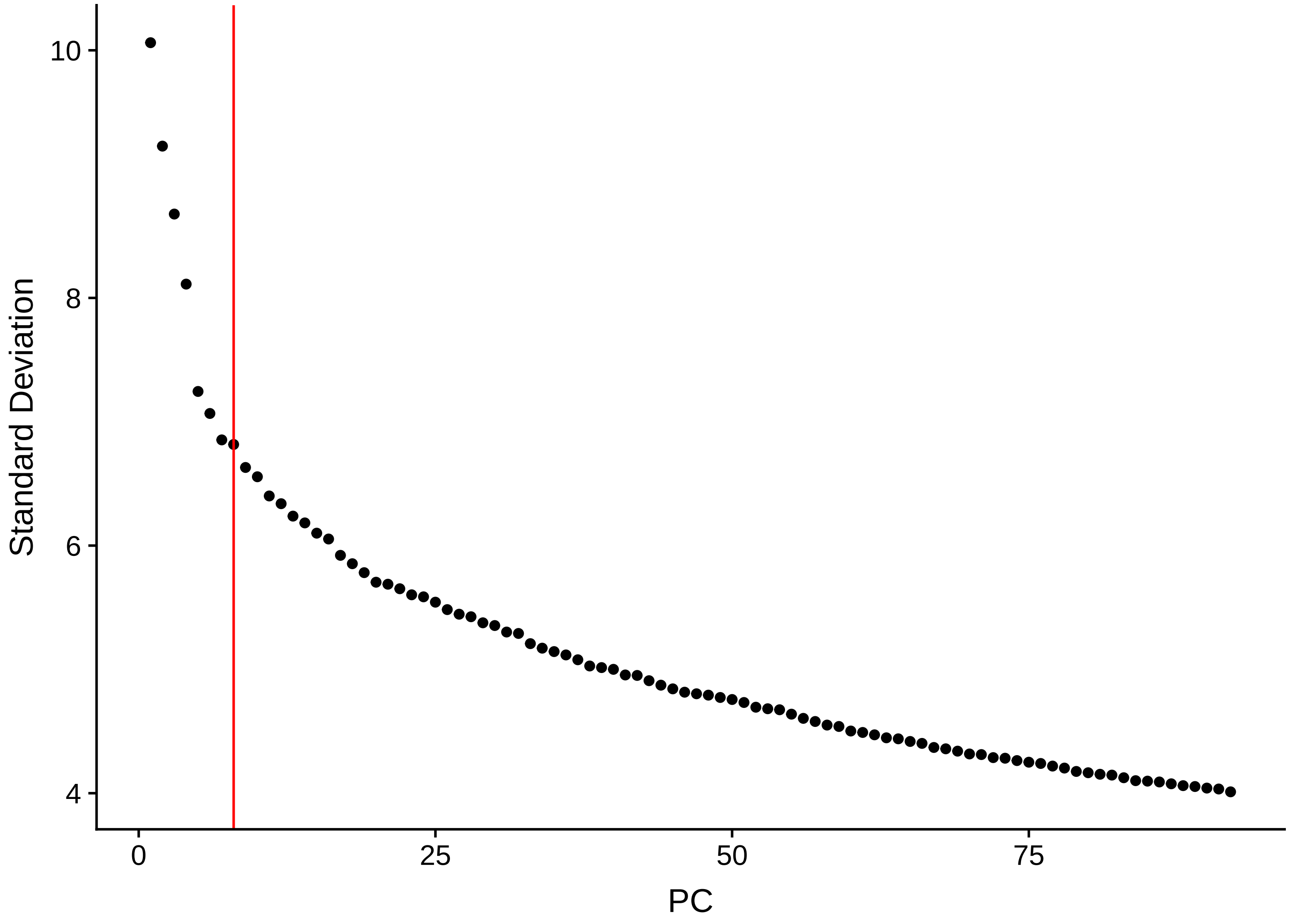

Standard Deviation
PC
10
8
6
4
0
25
50
75

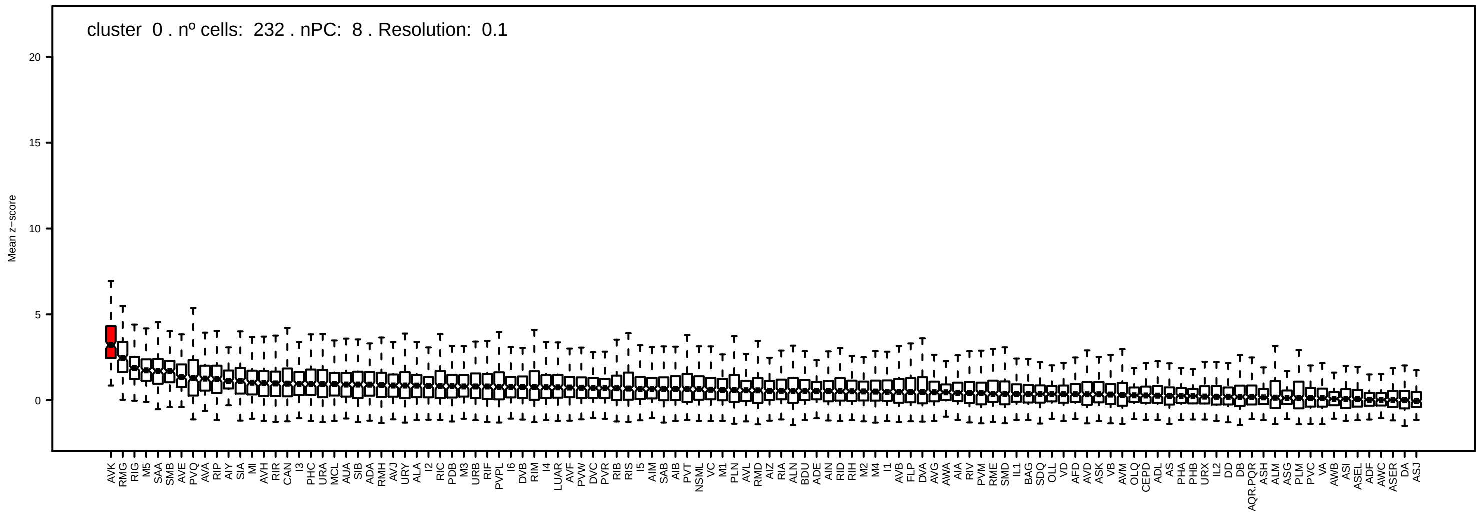

cluster 0 . nº cells: 232 . nPC: 8 . Resolution: 0.1
Mean z−score

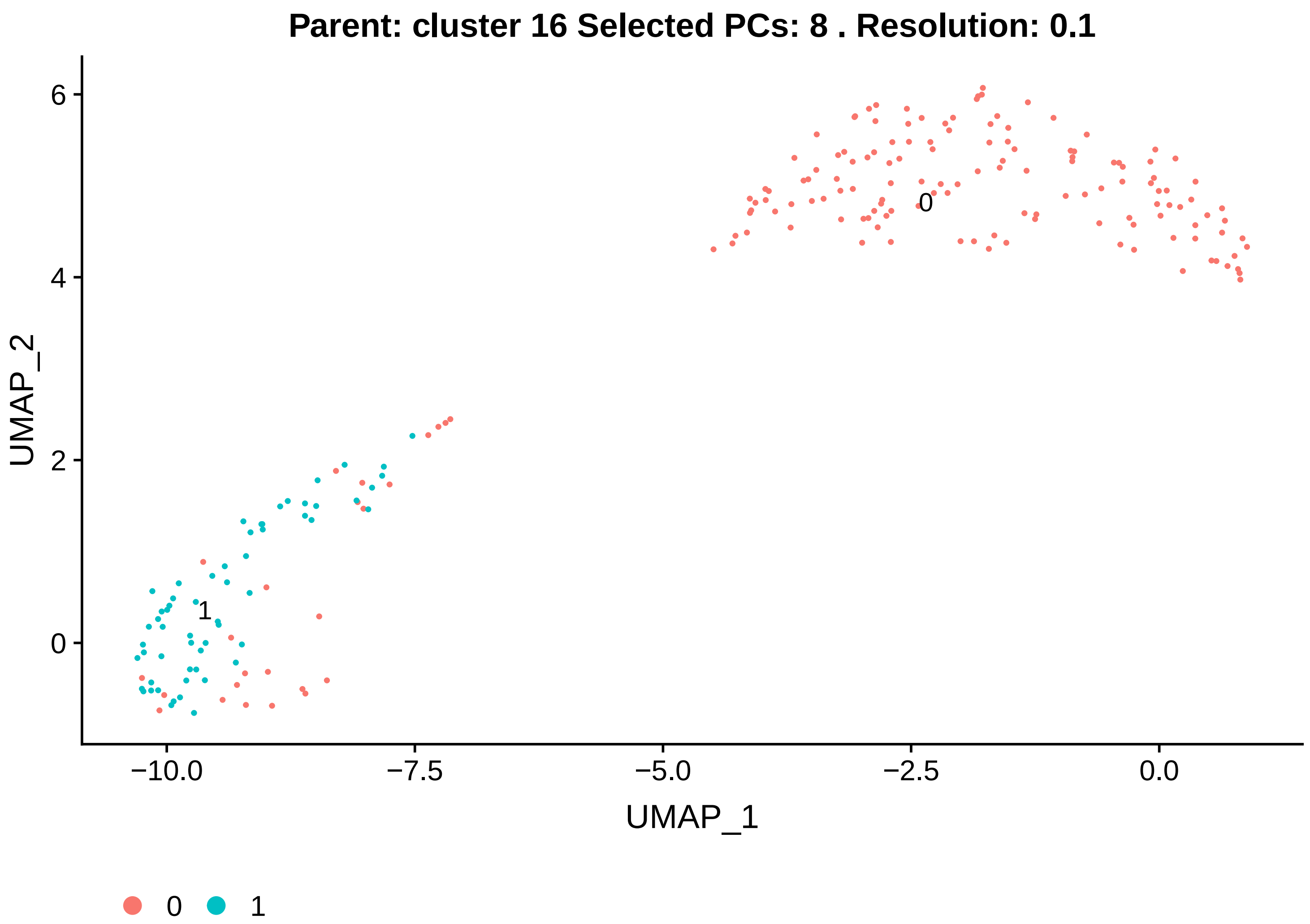

Parent: cluster 16 Selected PCs: 8 . Resolution: 0.1
UMAP_2
UMAP_1
6
4
2
0
−10.0
−7.5
−5.0
−2.5
0.0
0
1


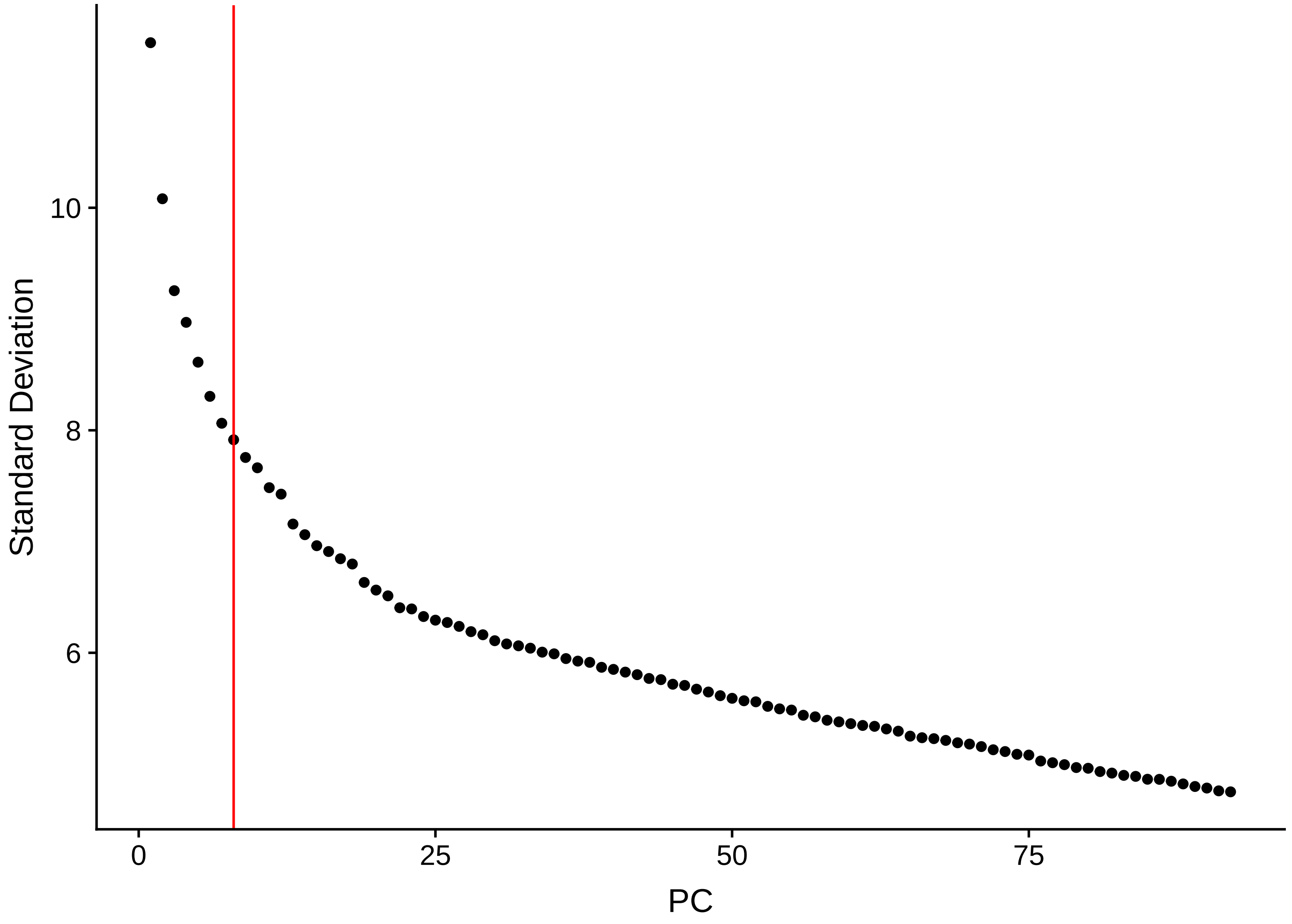

Standard Deviation
PC
10
8
6
0
25
50
75

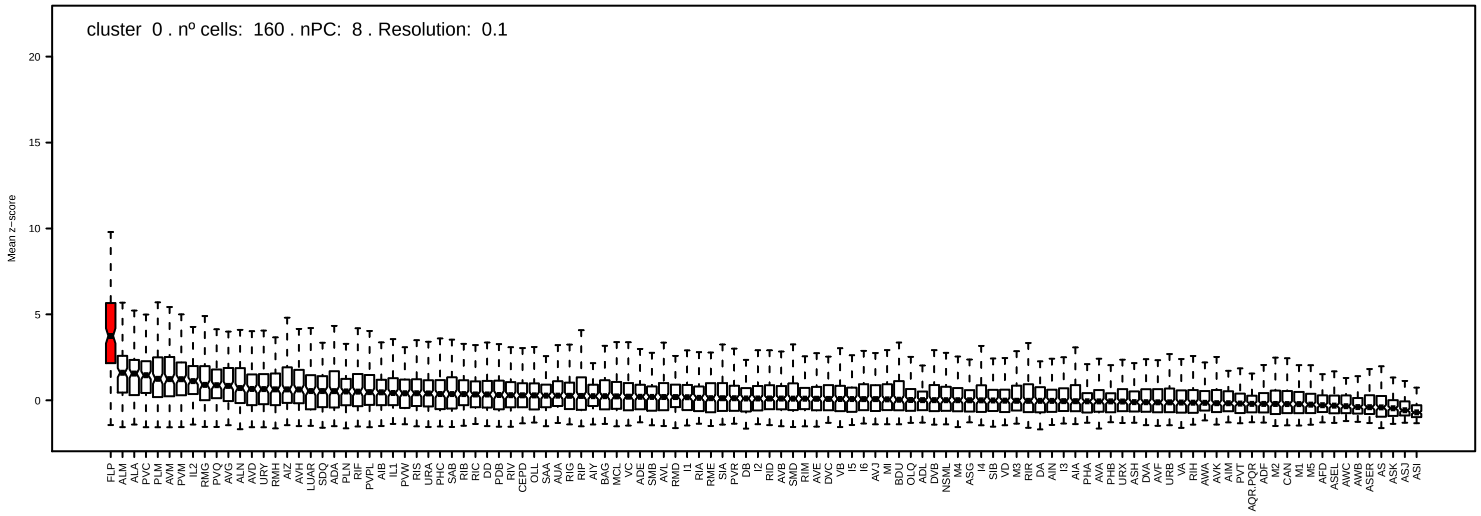
cluster 0 . nº cells: 160 . nPC: 8 . Resolution: 0.1
Mean z−score
20
15
10
5
0

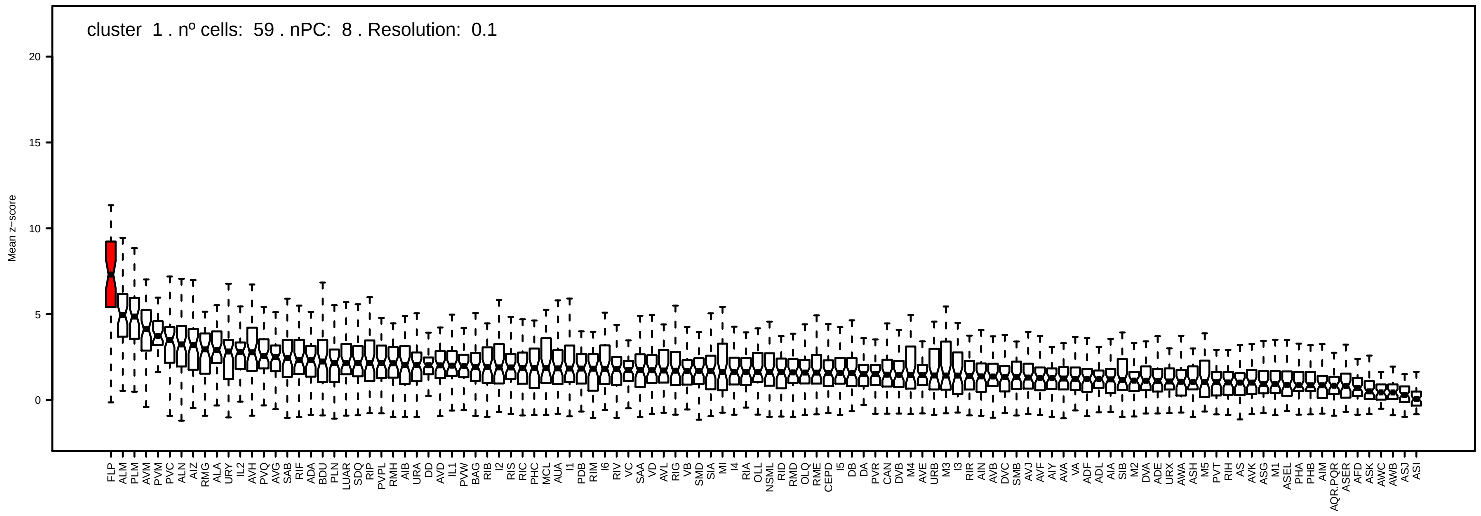
cluster 1 . nº cells: 59 . nPC: 8 . Resolution: 0.1
Mean z−score
20
15
10
5
0

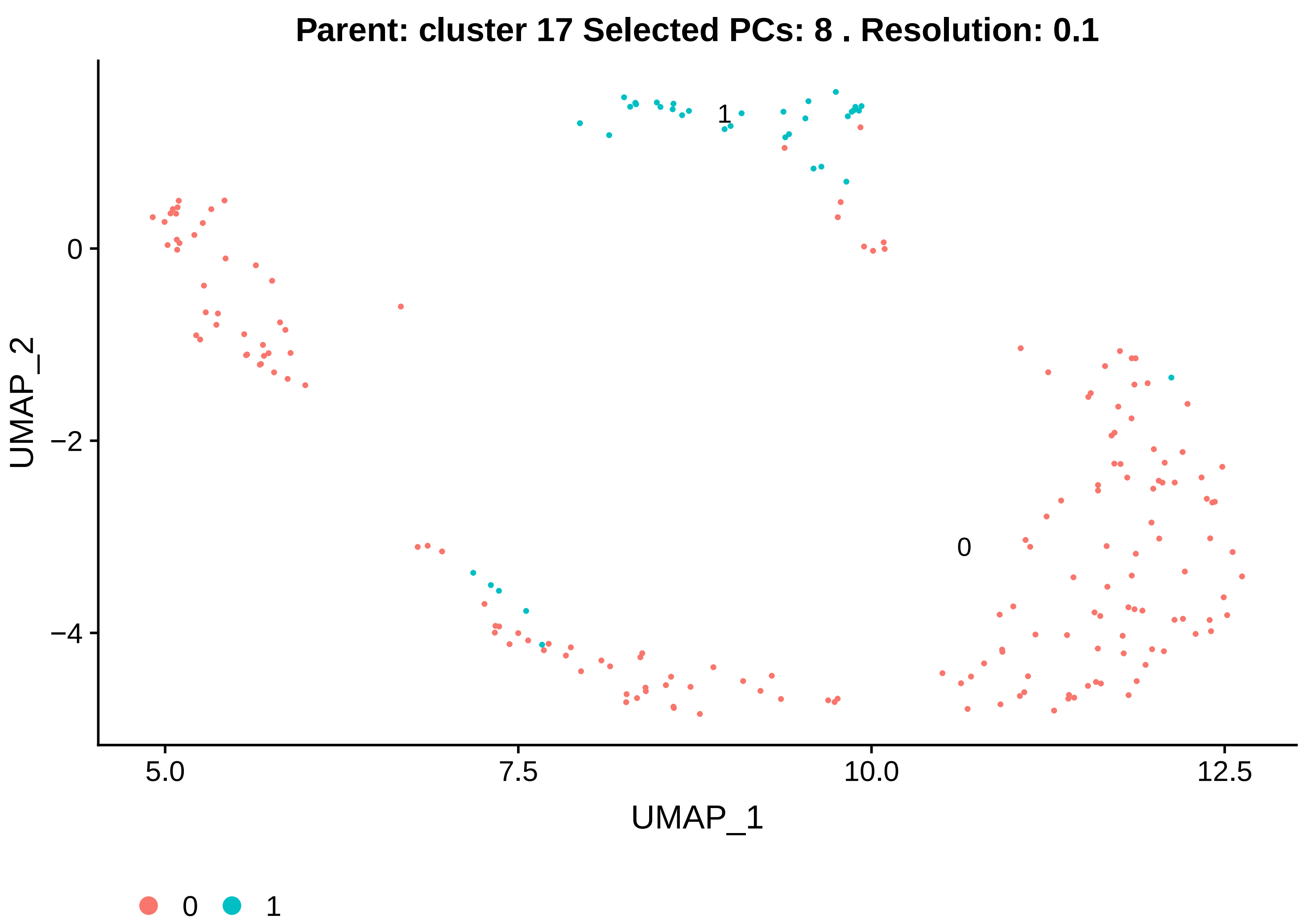

Parent: cluster 17 Selected PCs: 8 . Resolution: 0.1
UMAP_2
UMAP_1
0
1
0
−2
−4
5.0
7.5
10.0
12.5
0
1


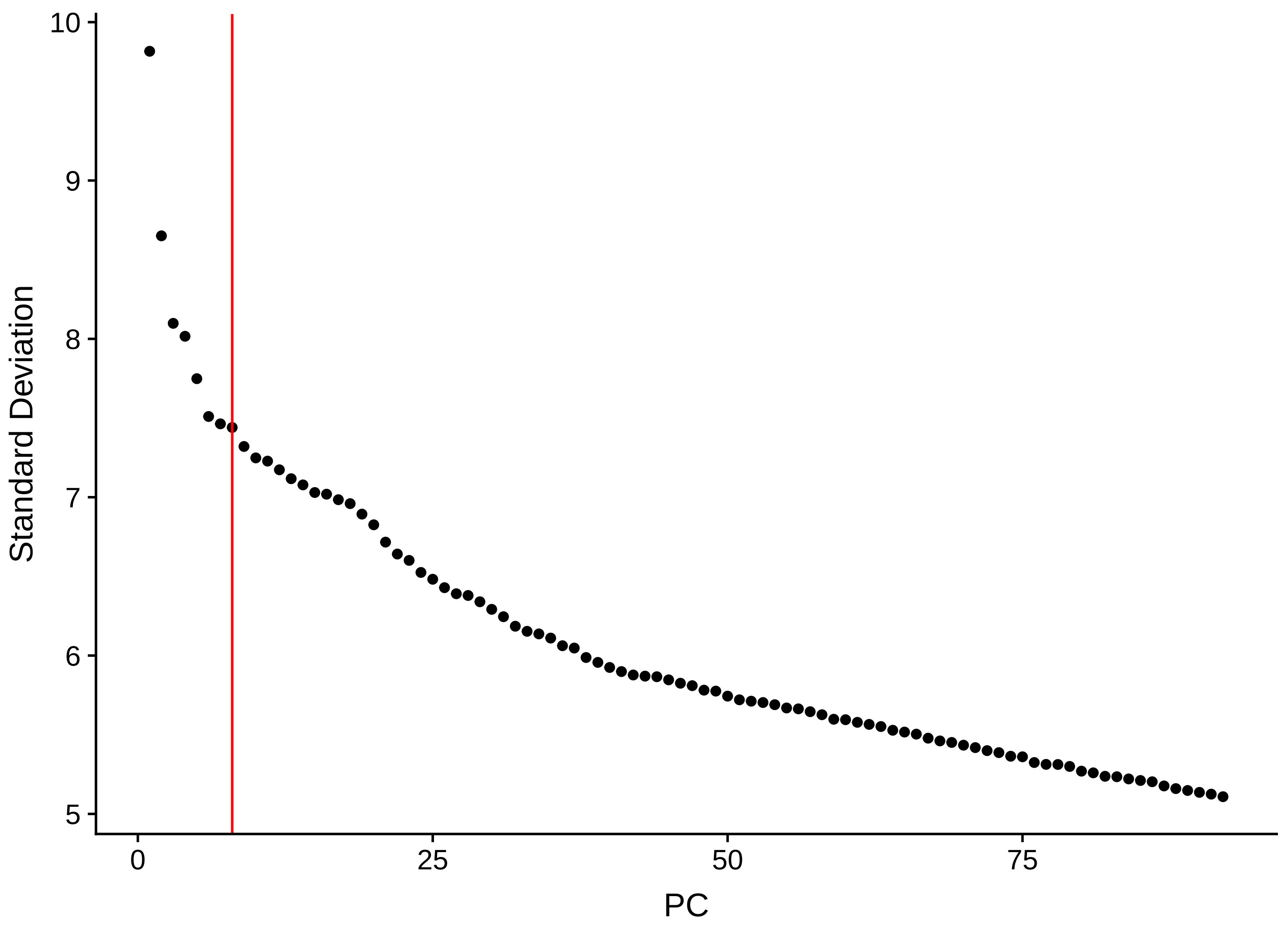

Standard Deviation
PC
10
9
8
7
6
5
0
25
50
75

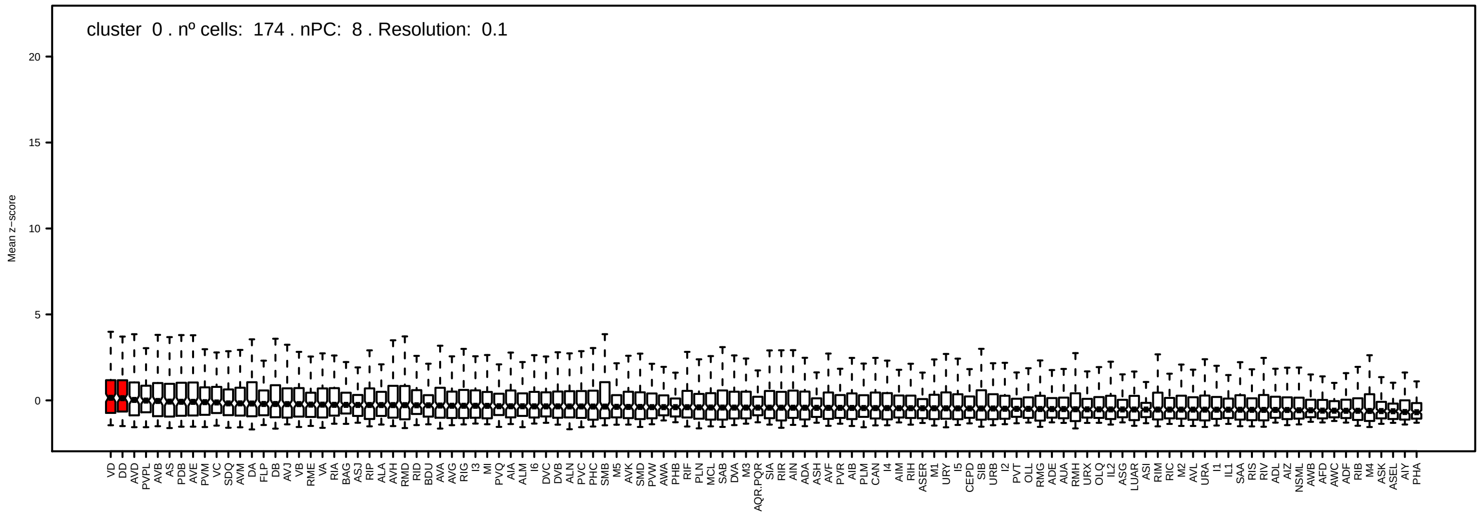
cluster 0 . nº cells: 174 . nPC: 8 . Resolution: 0.1
Mean z−score

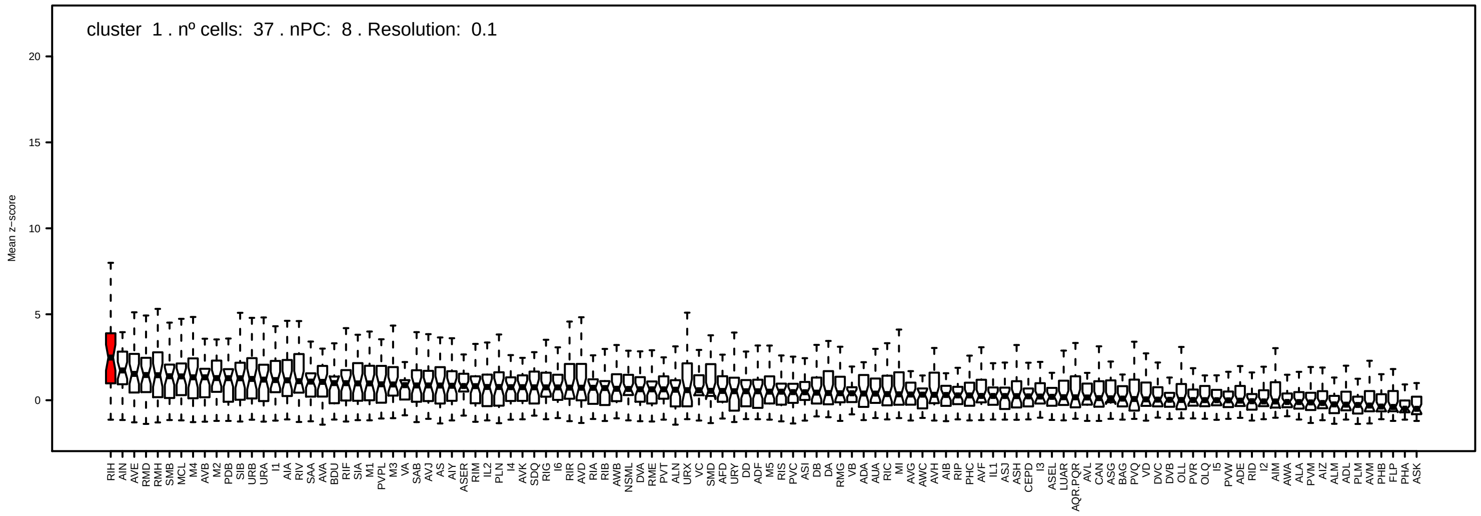
cluster 1 . nº cells: 37 . nPC: 8 . Resolution: 0.1
Mean z−score

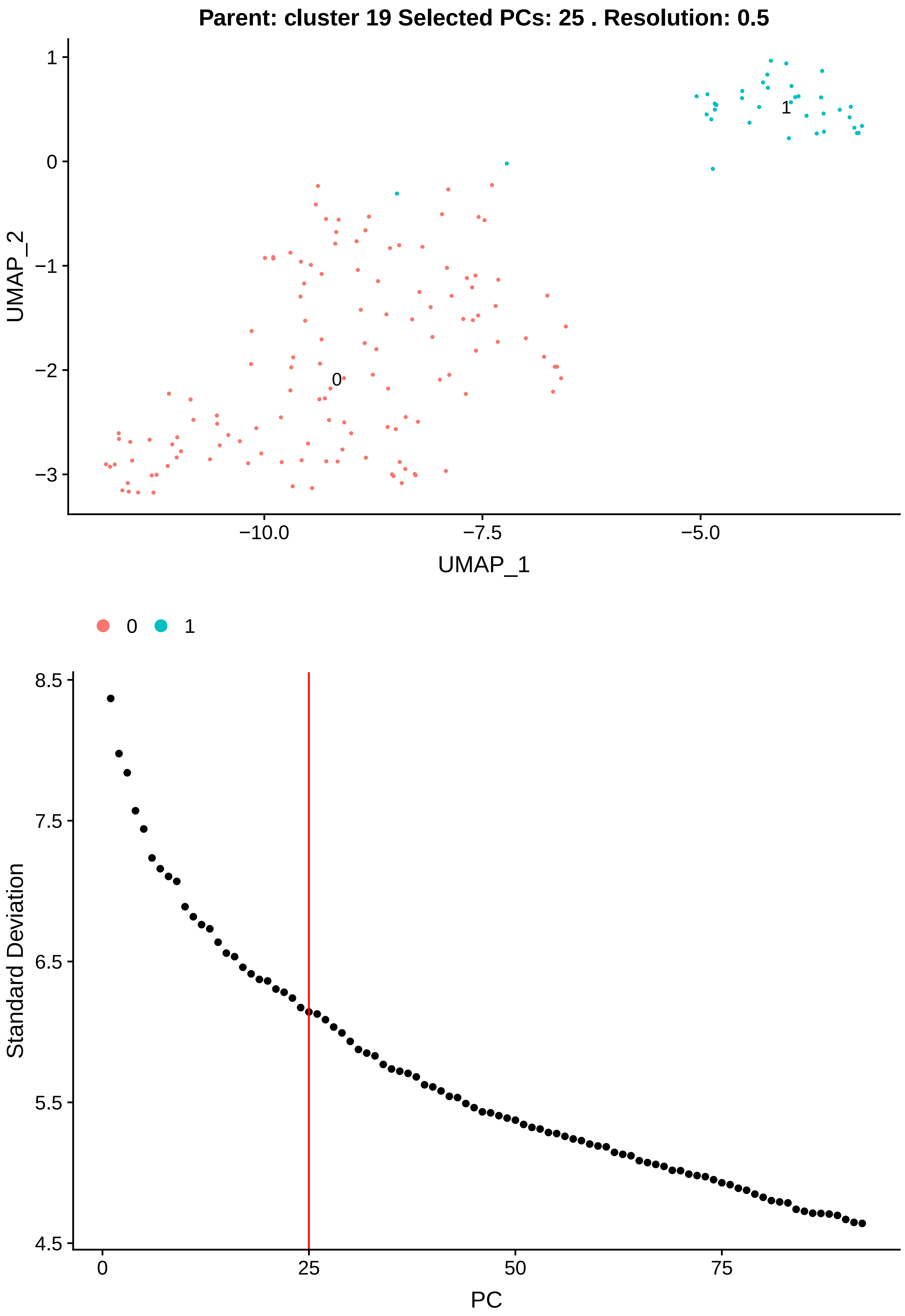

Parent: cluster 19 Selected PCs: 25 . Resolution: 0.5
1
0
−1
−2
−3
UMAP_2
0
1
−10.0
−7.5
−5.0
UMAP_1
0
1
8.5
7.5
6.5
5.5
4.5
Standard Deviation
0
25
50
75
PC

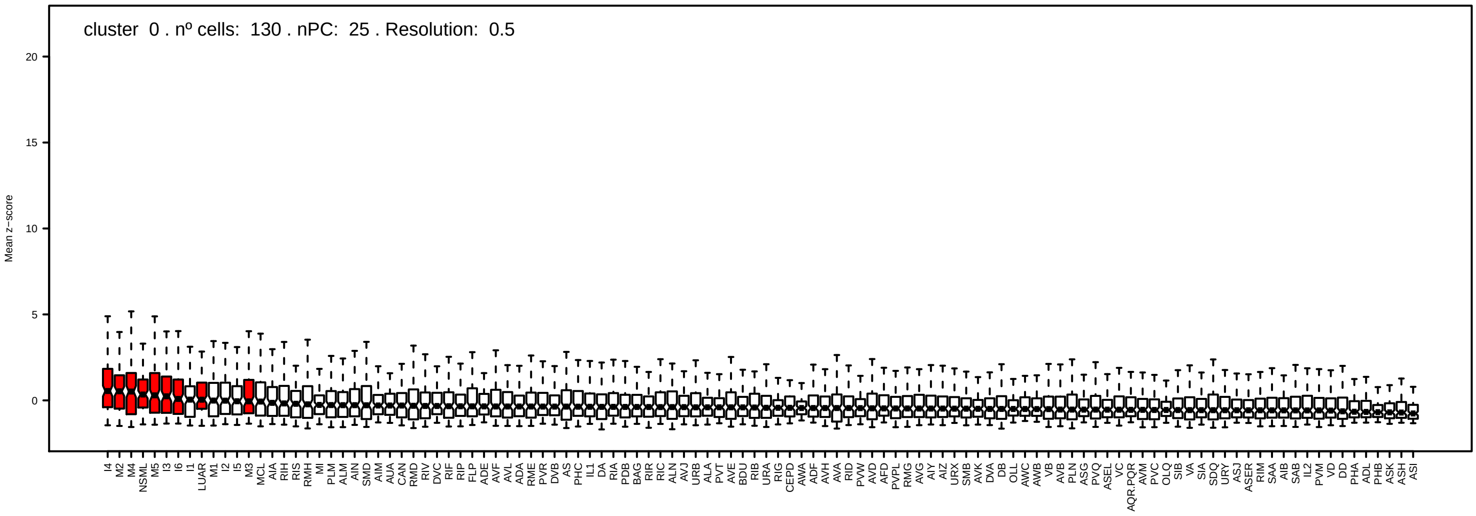
cluster 0 . nº cells: 130 . nPC: 25 . Resolution: 0.5
Mean z−score
20
15
10
5
0

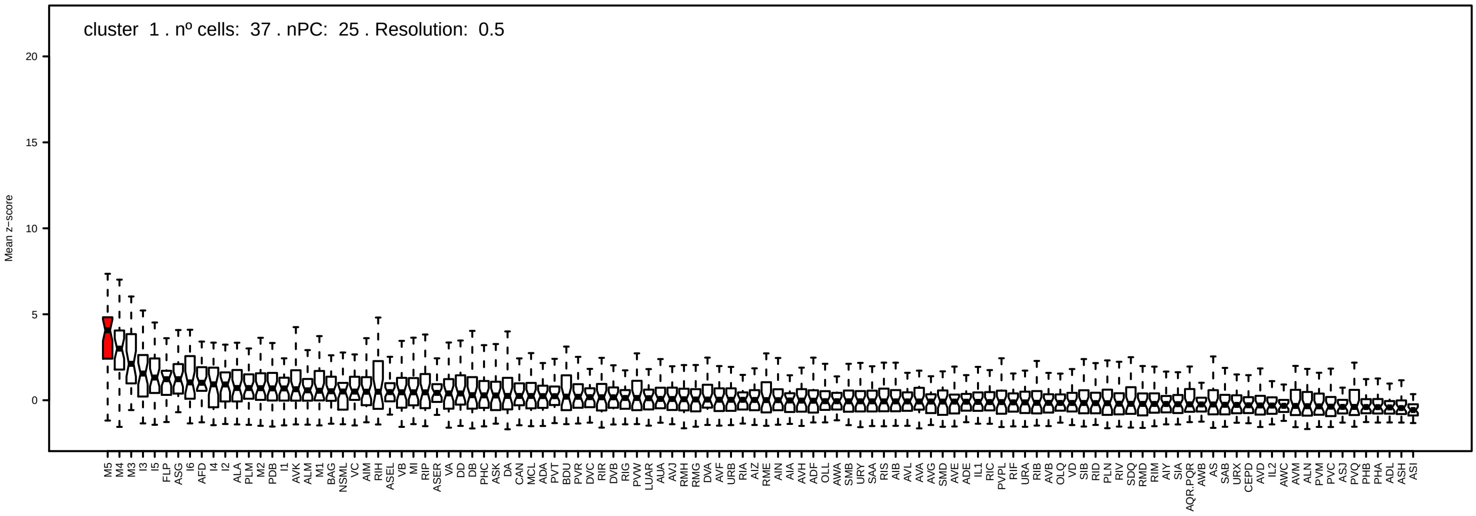
cluster 1 . nº cells: 37 . nPC: 25 . Resolution: 0.5
Mean z−score
20
15
10
5
0

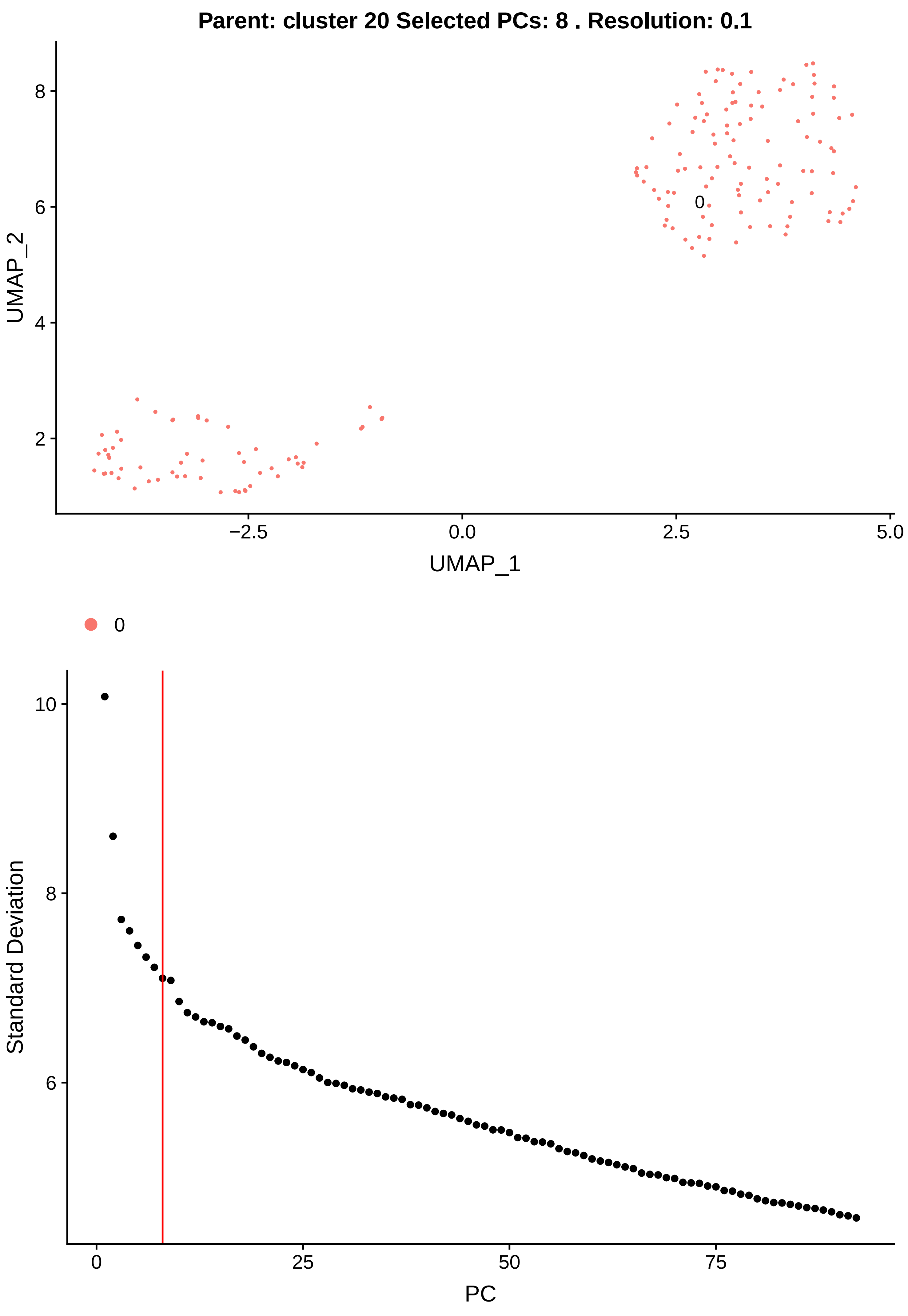

Parent: cluster 20 Selected PCs: 8 . Resolution: 0.1
UMAP_2
UMAP_1
0
8
6
4
2
−2.5
0.0
2.5
5.0
0
Standard Deviation
PC
10
8
6
0
25
50
75

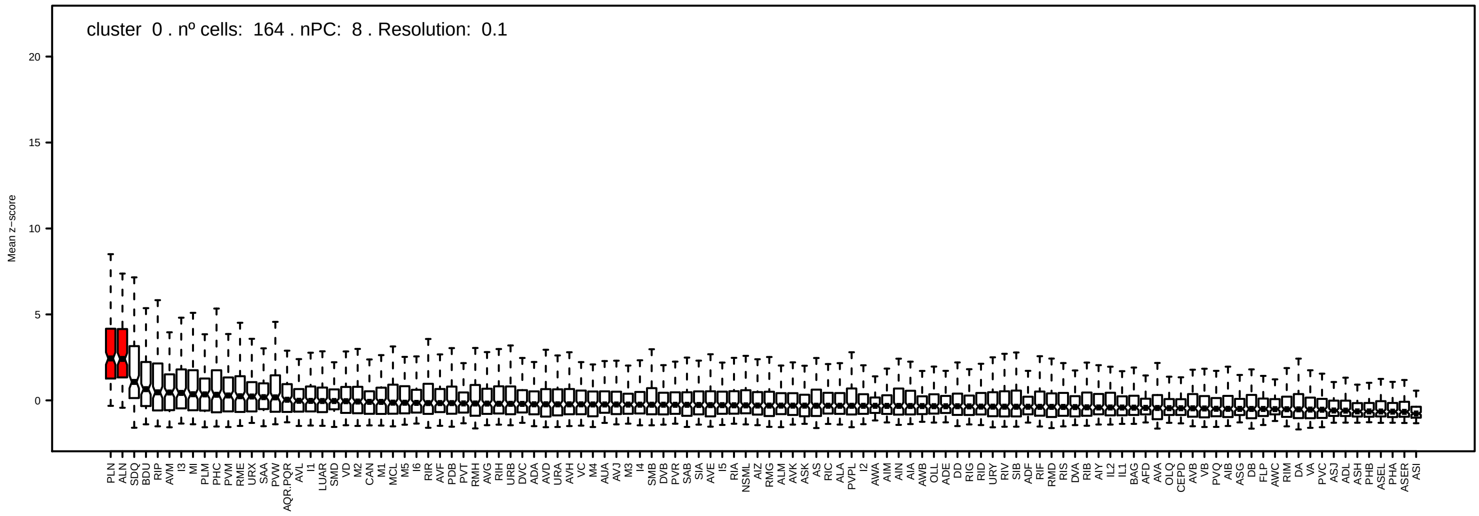

cluster 0 . nº cells: 164 . nPC: 8 . Resolution: 0.1
Mean z−score
20
15
10
5
0
PLN ALN SDQ BDU RIP AVM I3 MI PLM PHC PVM RME URX SAA PVW AQR.PQR AVL I1 LUAR SMD VD M2 CAN M1 MCL M5 I6 RIR AVF PDB PVT RMH AVG RIH URB DVC ADA AVD URA AVH VC M4 AUA AVJ M3 I4 SMB DVB PVR SAB SIA AVE I5 RIA NSML AIZ RMG ALM AVK ASK AS RIC ALA PVPL I2 AWA AIM AIN AIA AWB OLL ADE DD RIG RID URY RIV SIB ADF RIF RMD RIS DVA RIB AIY IL2 IL1 BAG AFD AVA OLQ CEPD AVB VB PVQ AIB ASG DB FLP AWC RIM DA VA PVC ASJ ADL ASH PHB ASEL PHA ASER ASI

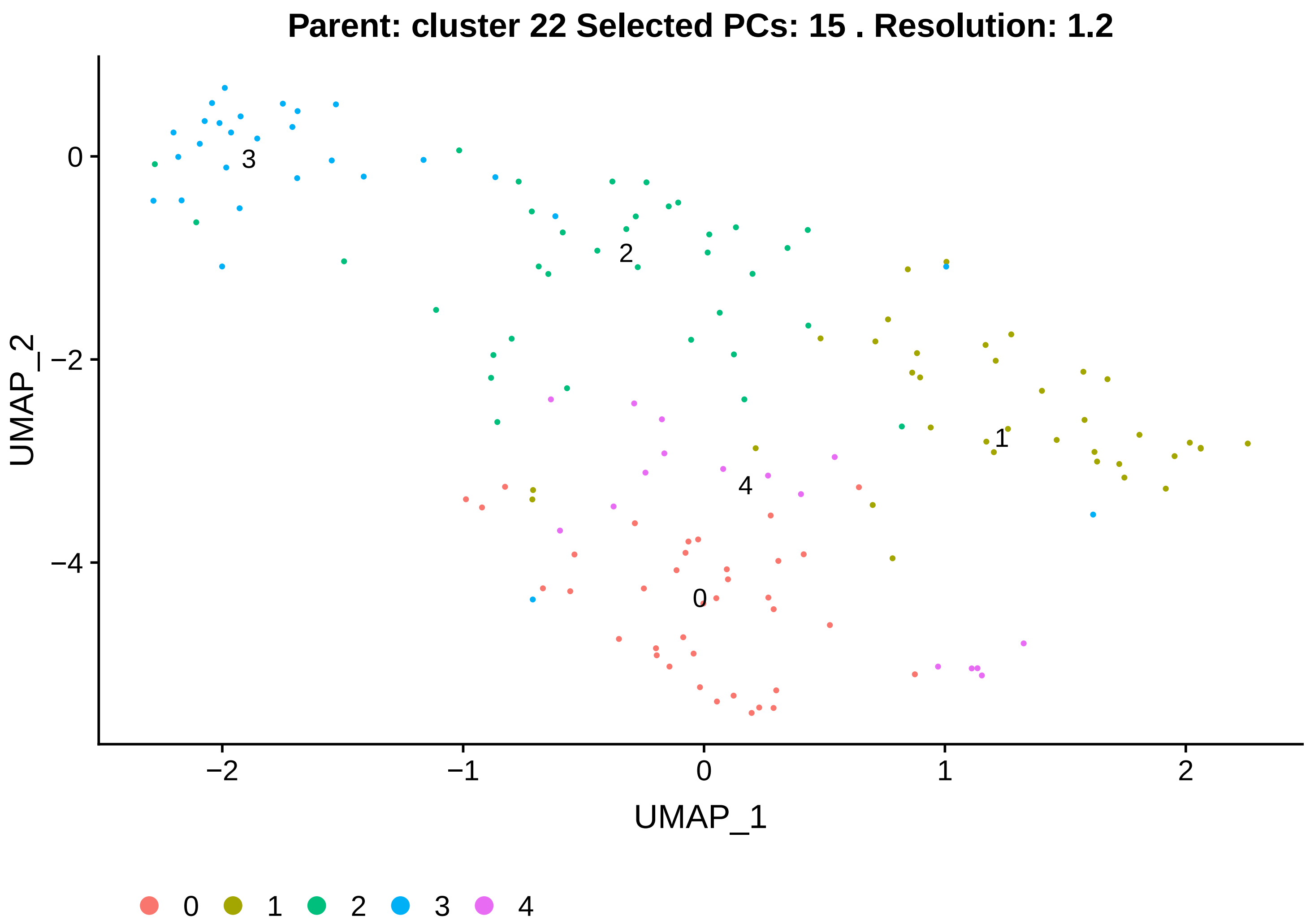
Parent: cluster 22 Selected PCs: 15 . Resolution: 1.2
UMAP_2
UMAP_1
0
1
2
3
4
−2
−1
0
1
2
0
−2
−4

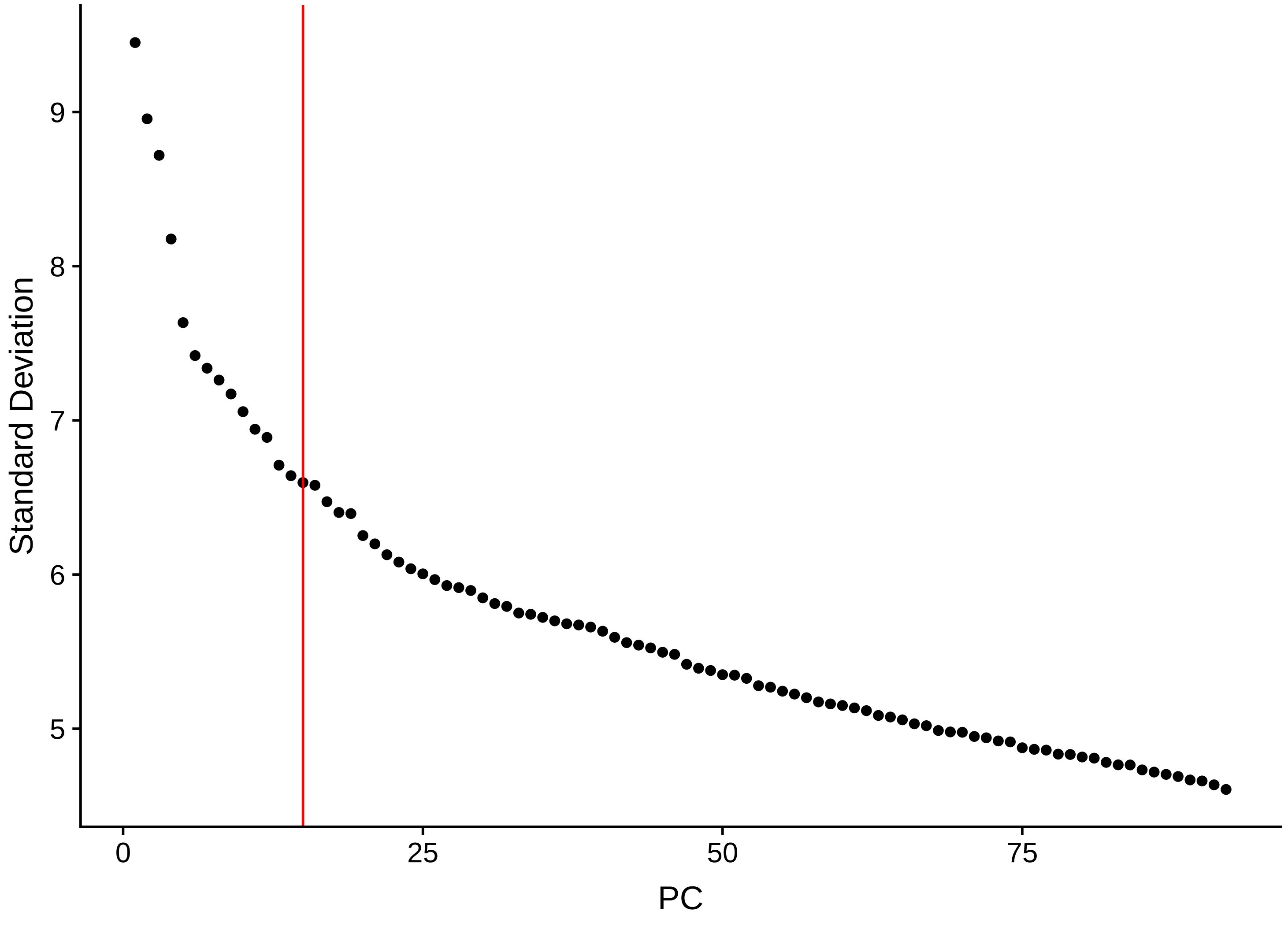
Standard Deviation
PC
9
8
7
6
5
0
25
50
75

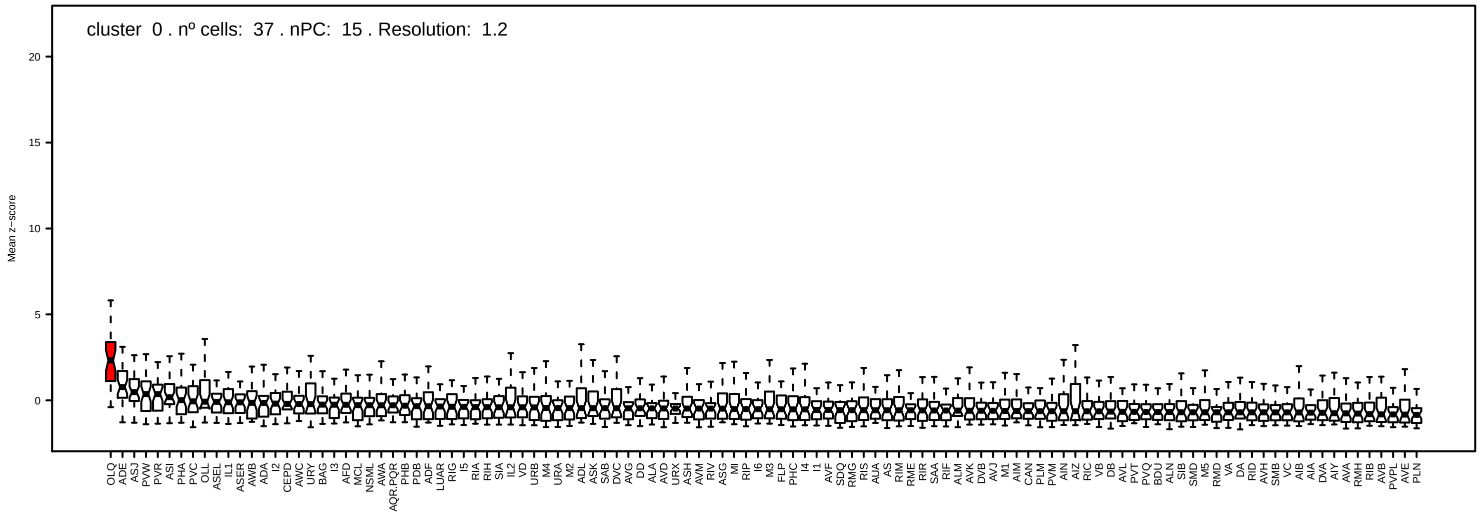
cluster 0 . nº cells: 37 . nPC: 15 . Resolution: 1.2
Mean z−score

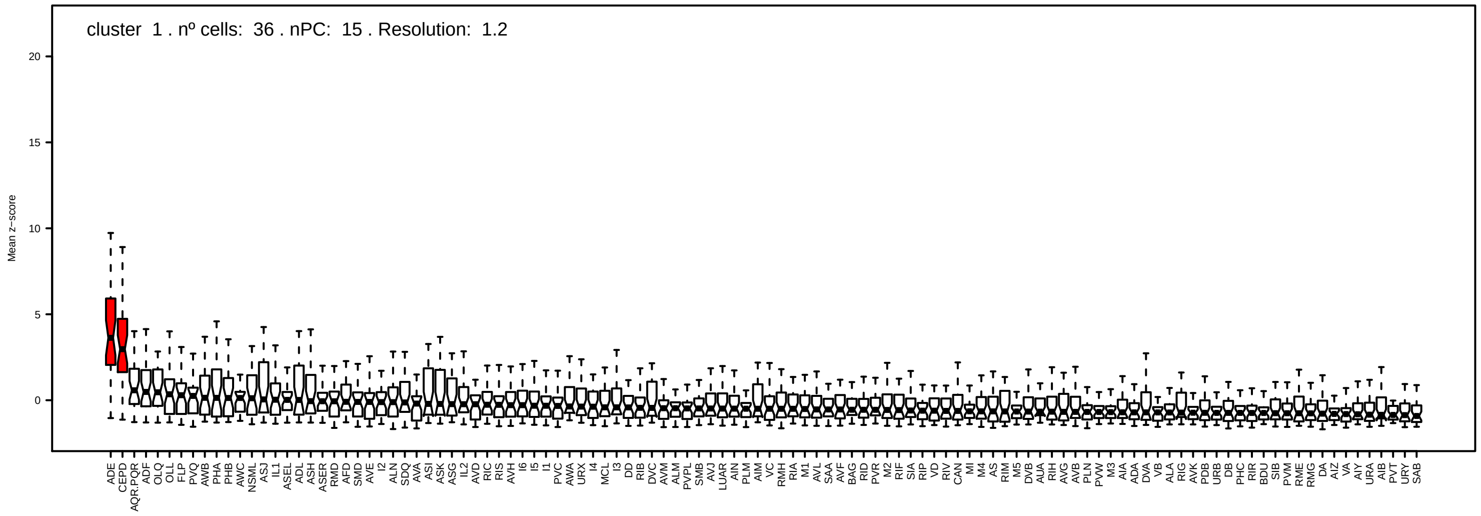
cluster 1 . nº cells: 36 . nPC: 15 . Resolution: 1.2
Mean z−score

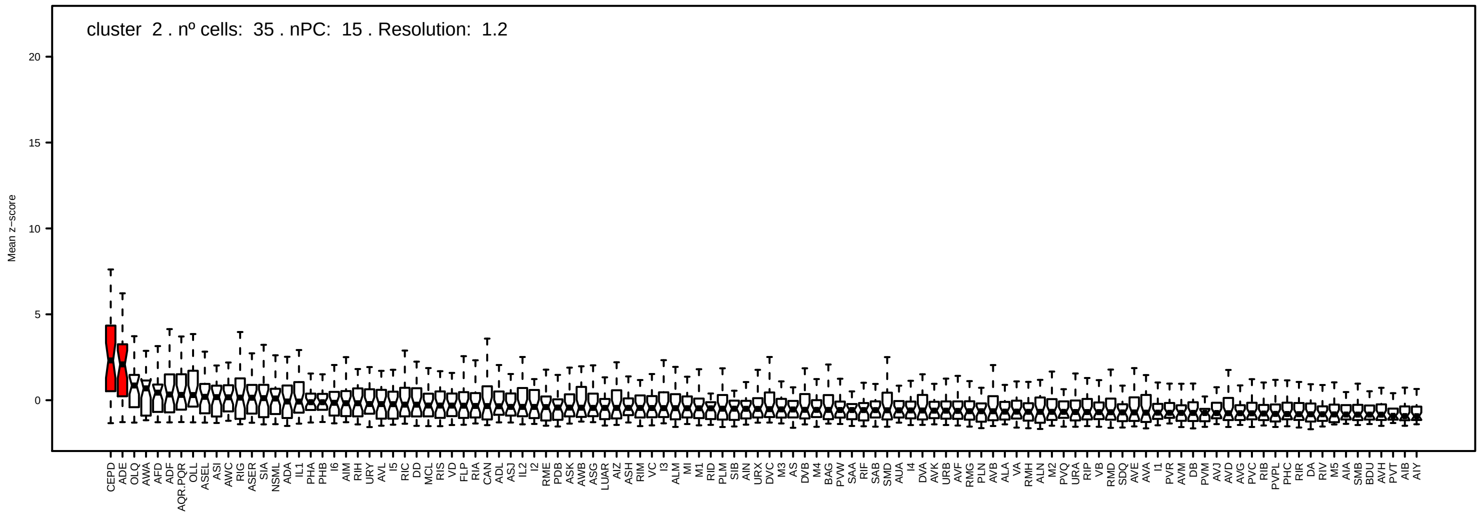
cluster 2 . nº cells: 35 . nPC: 15 . Resolution: 1.2
Mean z−score

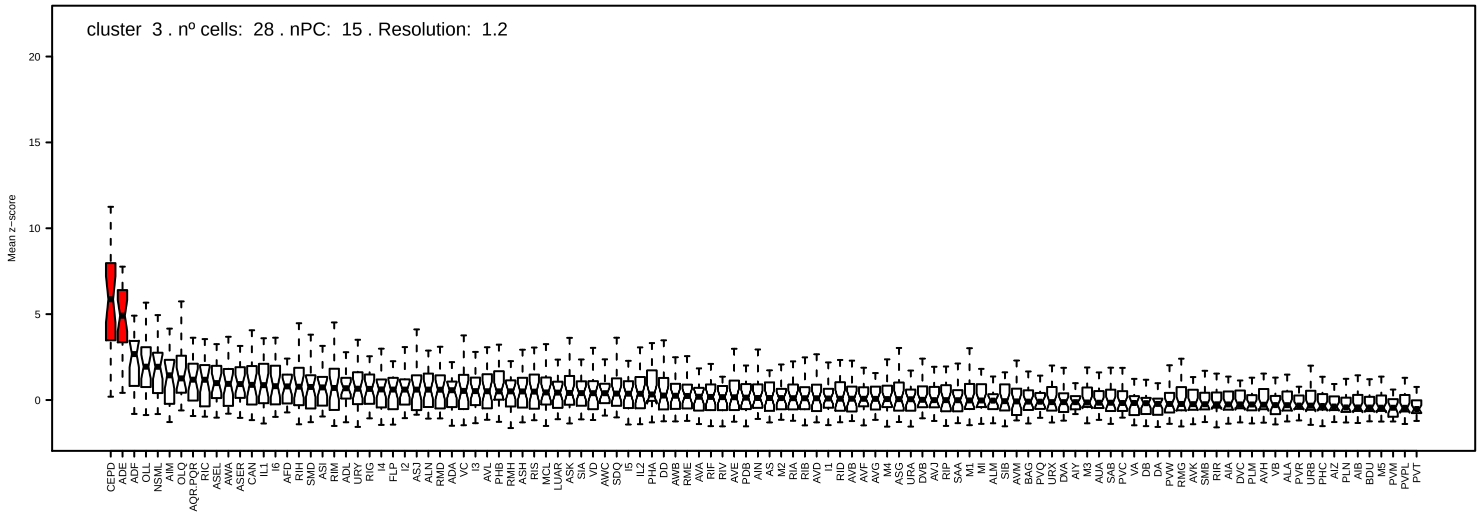
cluster 3 . nº cells: 28 . nPC: 15 . Resolution: 1.2
Mean z−score

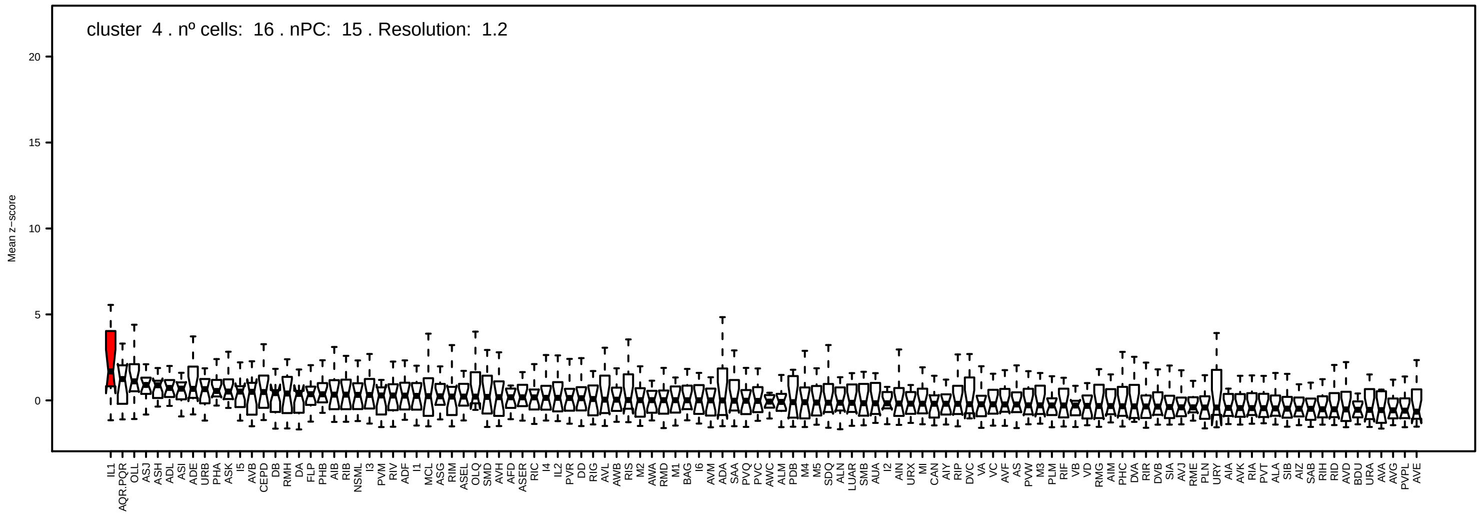
cluster 4 . nº cells: 16 . nPC: 15 . Resolution: 1.2
Mean z−score

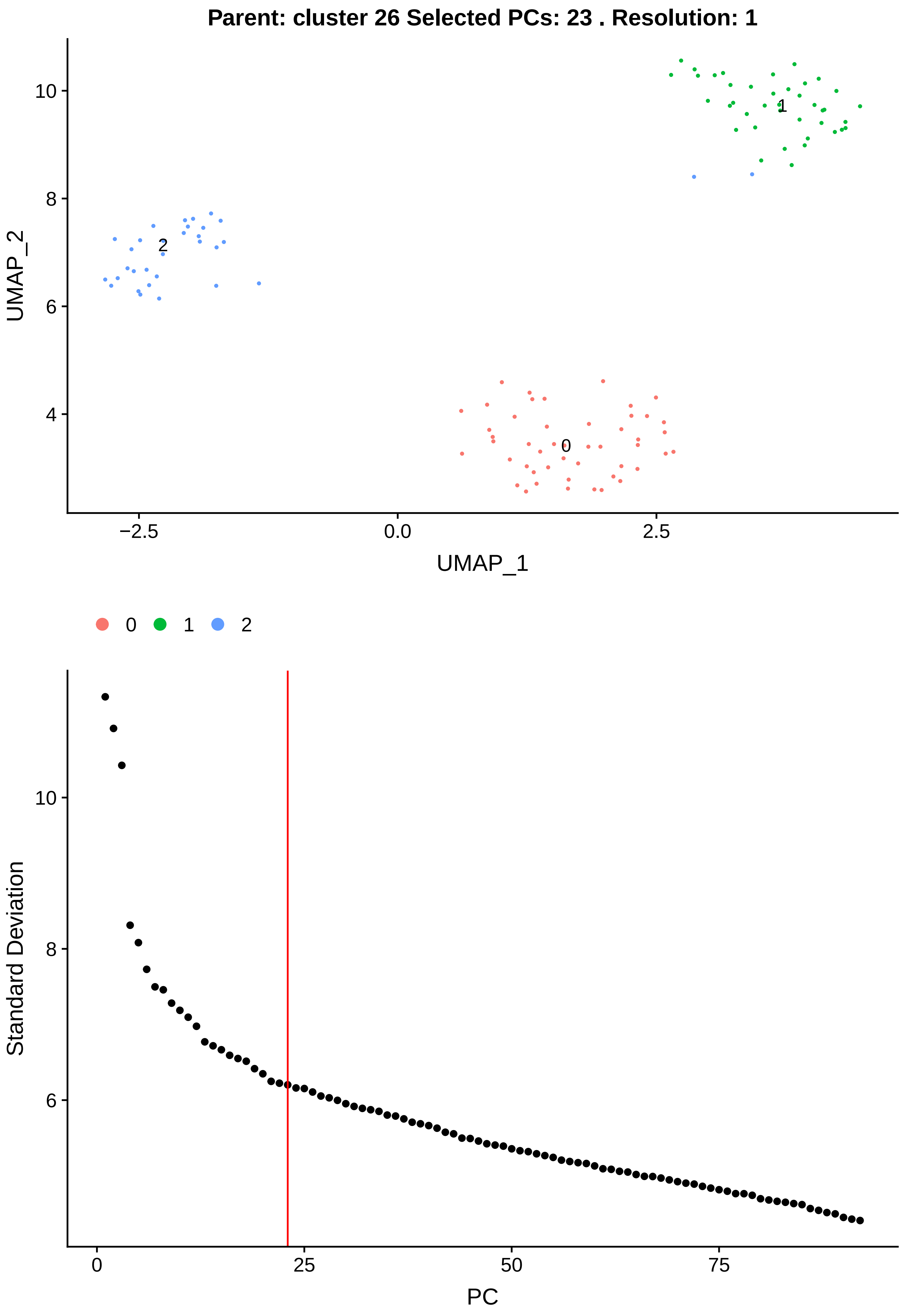

Parent: cluster 26 Selected PCs: 23 . Resolution: 1
UMAP_2
UMAP_1
−2.5
0.0
2.5
10
8
6
4
0
1
2
Standard Deviation
PC
0
25
50
75

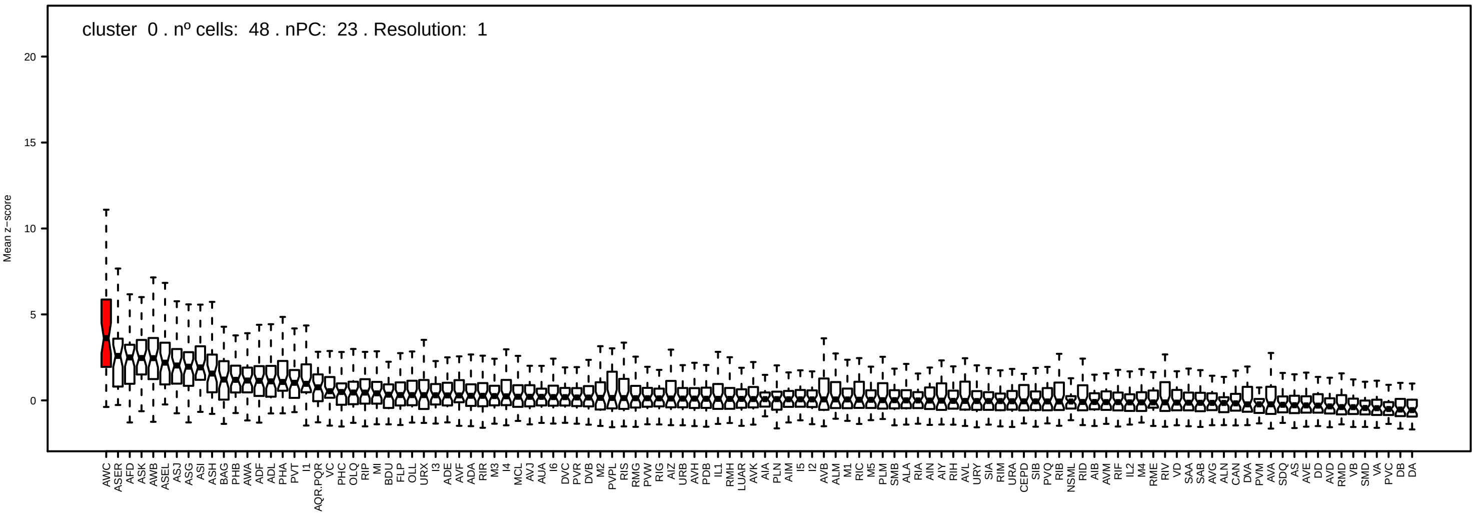
cluster 0 . nº cells: 48 . nPC: 23 . Resolution: 1
Mean z-score
0
5
10
15
20

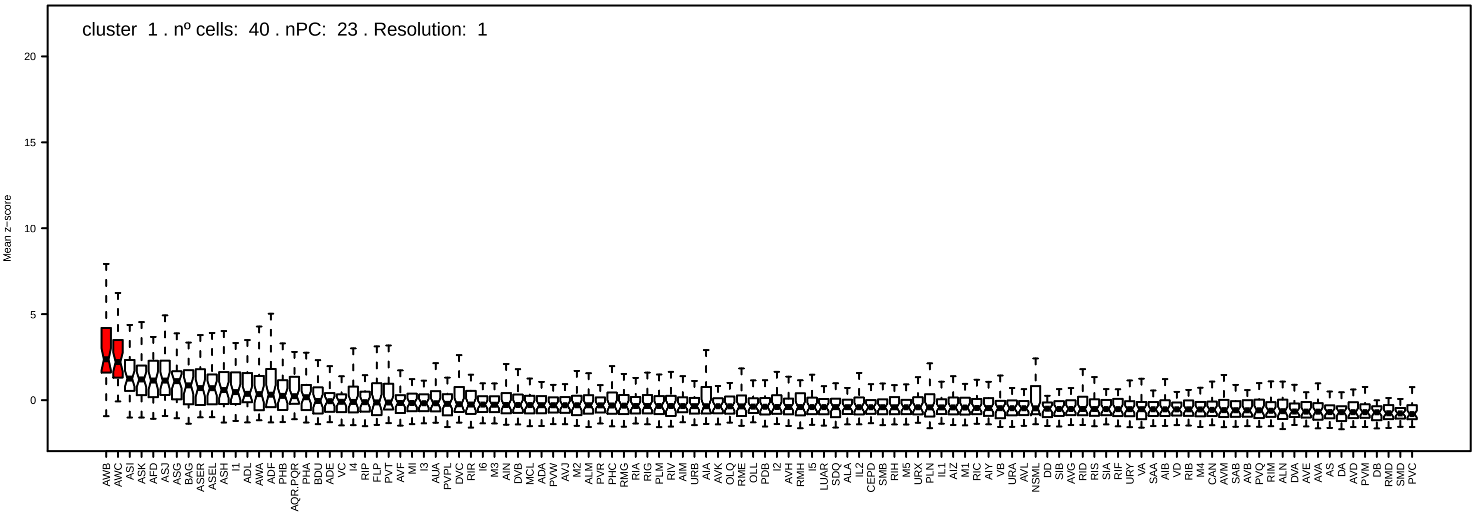
cluster 1 . nº cells: 40 . nPC: 23 . Resolution: 1
Mean z-score
0
5
10
15
20

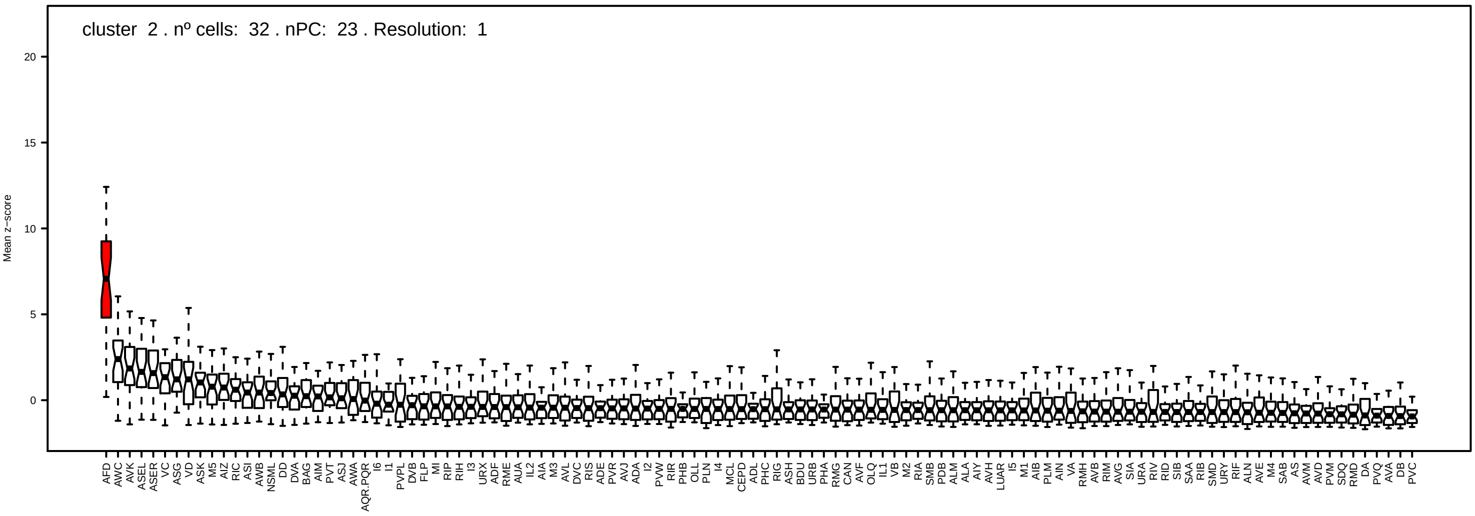
cluster 2 . nº cells: 32 . nPC: 23 . Resolution: 1
Mean z-score
0
5
10
15
20

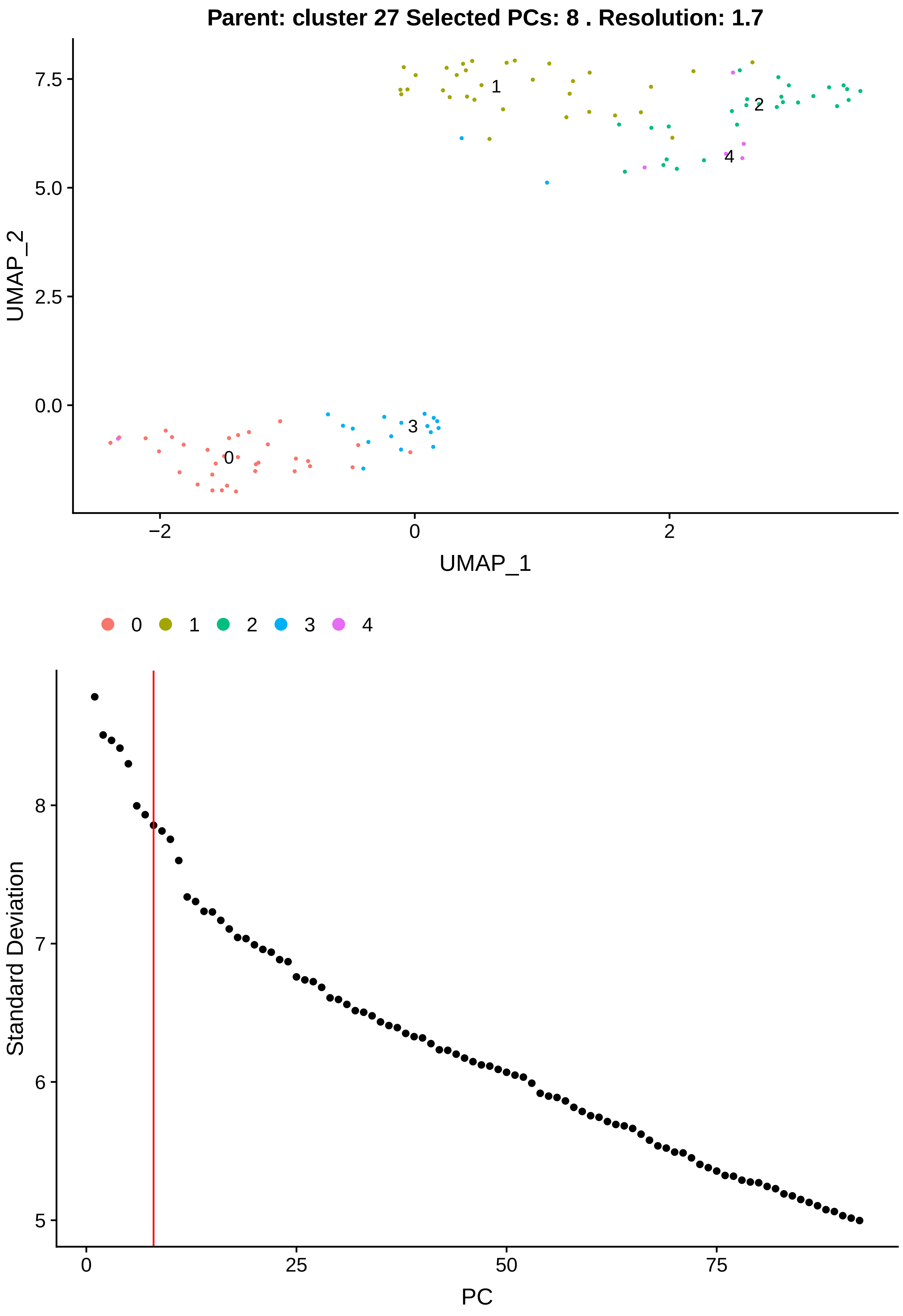
Parent: cluster 27 Selected PCs: 8 . Resolution: 1.7
UMAP_2
UMAP_1
0
1
2
3
4
Standard Deviation
PC

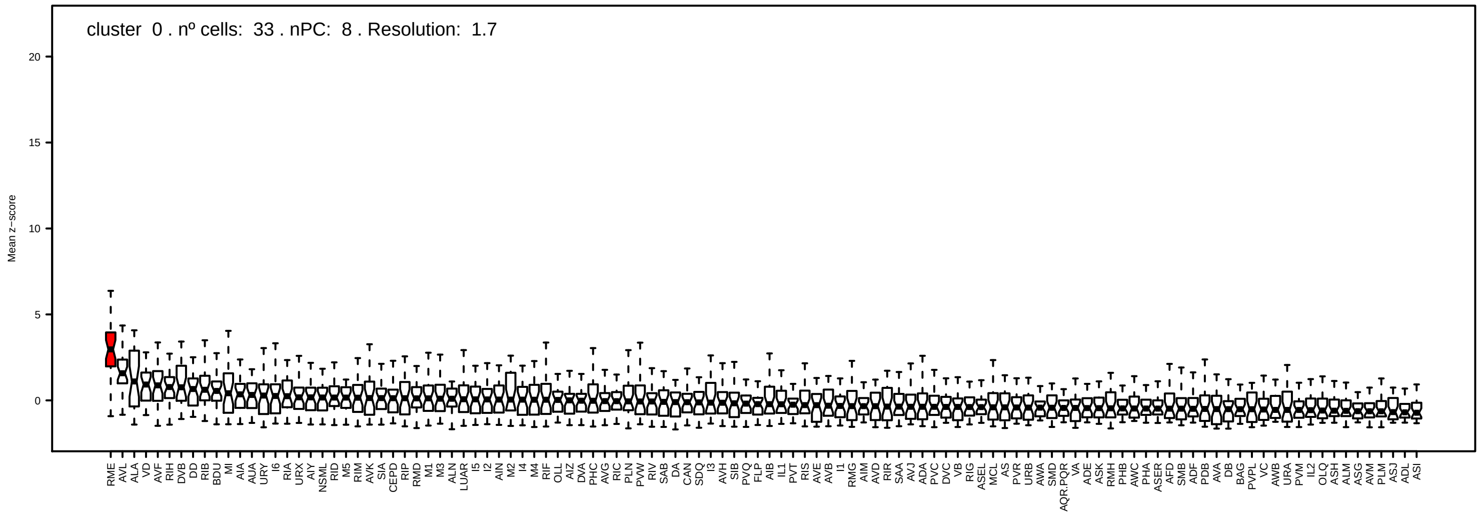
cluster 0 . nº cells: 33 . nPC: 8 . Resolution: 1.7
Mean z−score
20
15
10
5
0

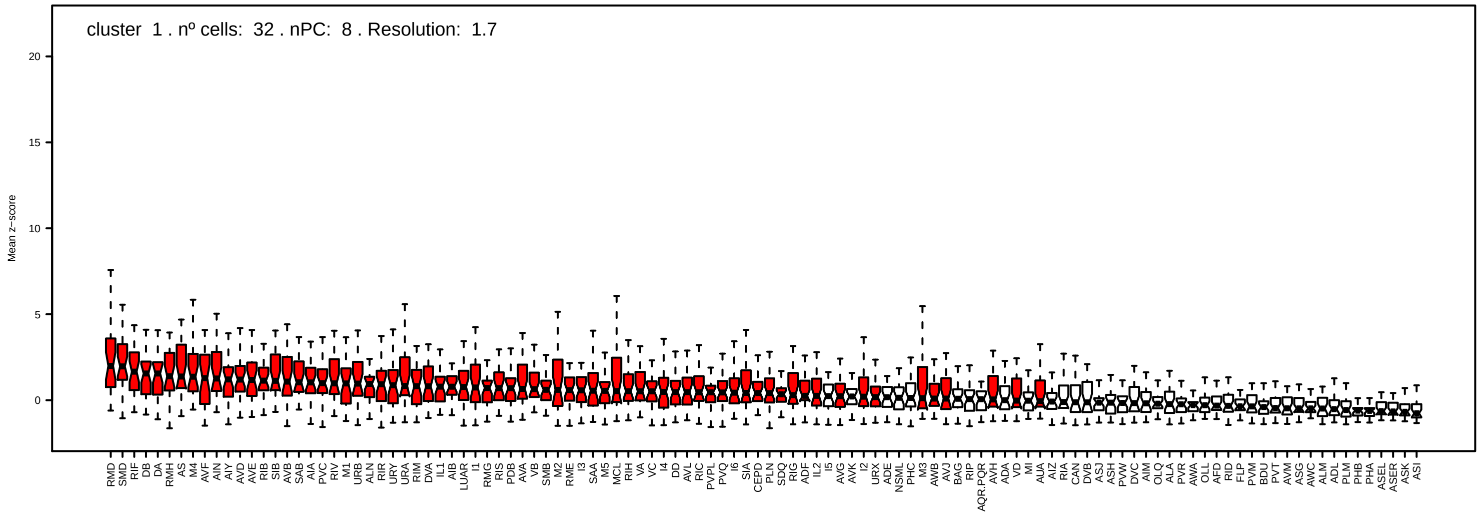
cluster 1 . nº cells: 32 . nPC: 8 . Resolution: 1.7
Mean z−score
20
15
10
5
0

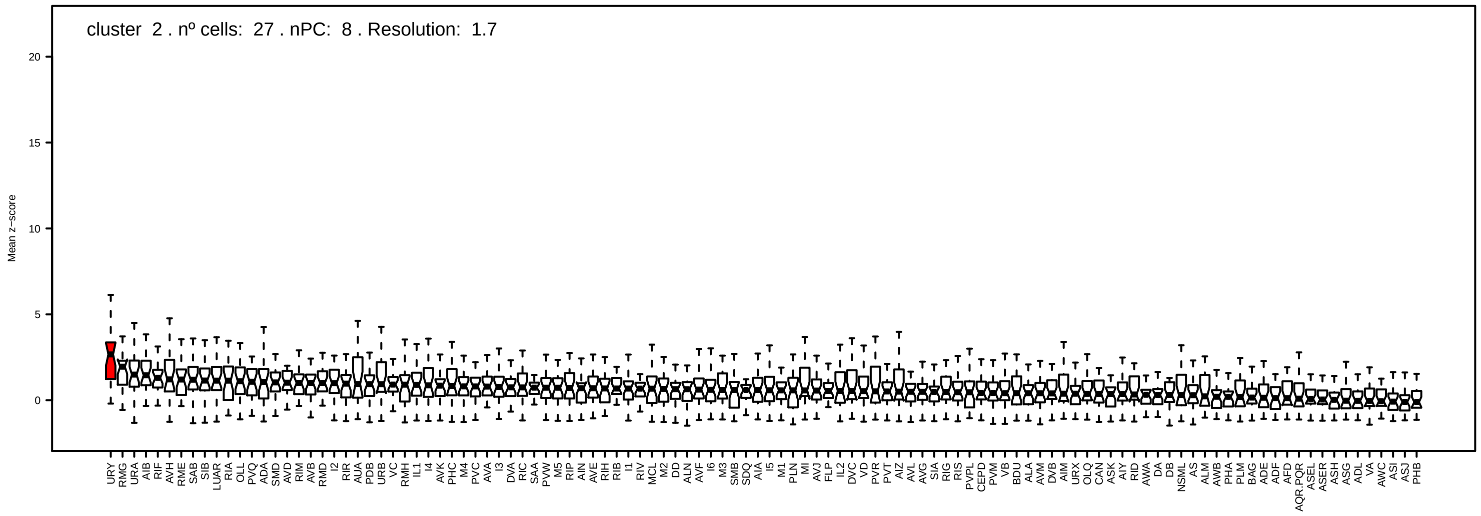
cluster 2 . nº cells: 27 . nPC: 8 . Resolution: 1.7
Mean z−score
20
15
10
5
0

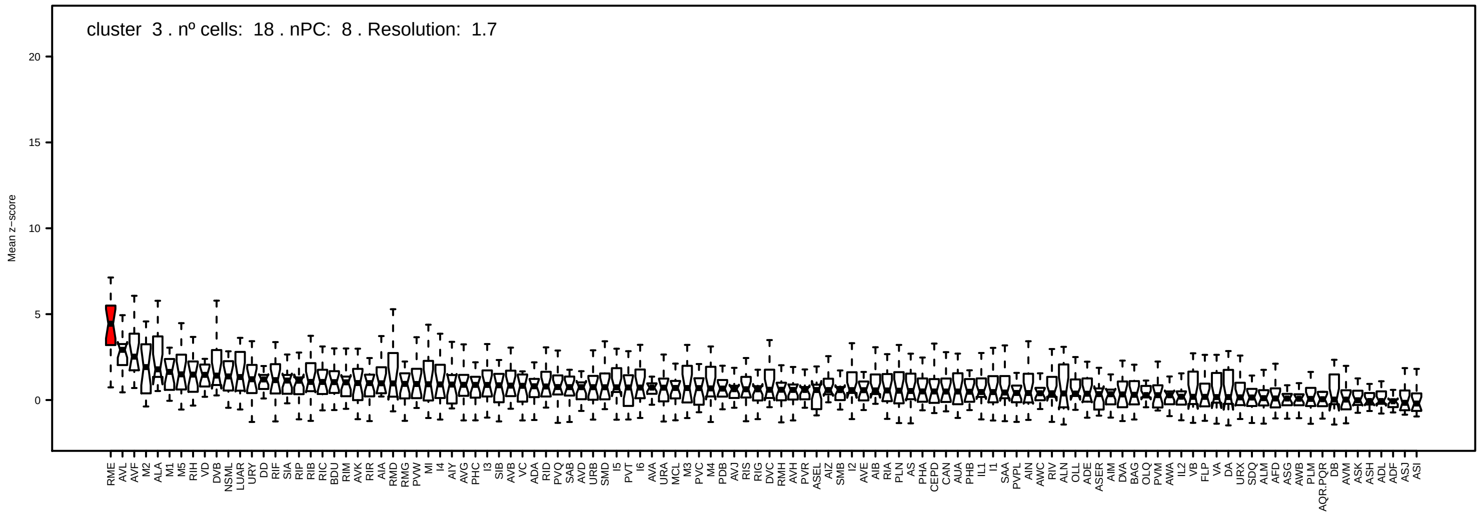
cluster 3 . nº cells: 18 . nPC: 8 . Resolution: 1.7
Mean z−score
20
15
10
5
0

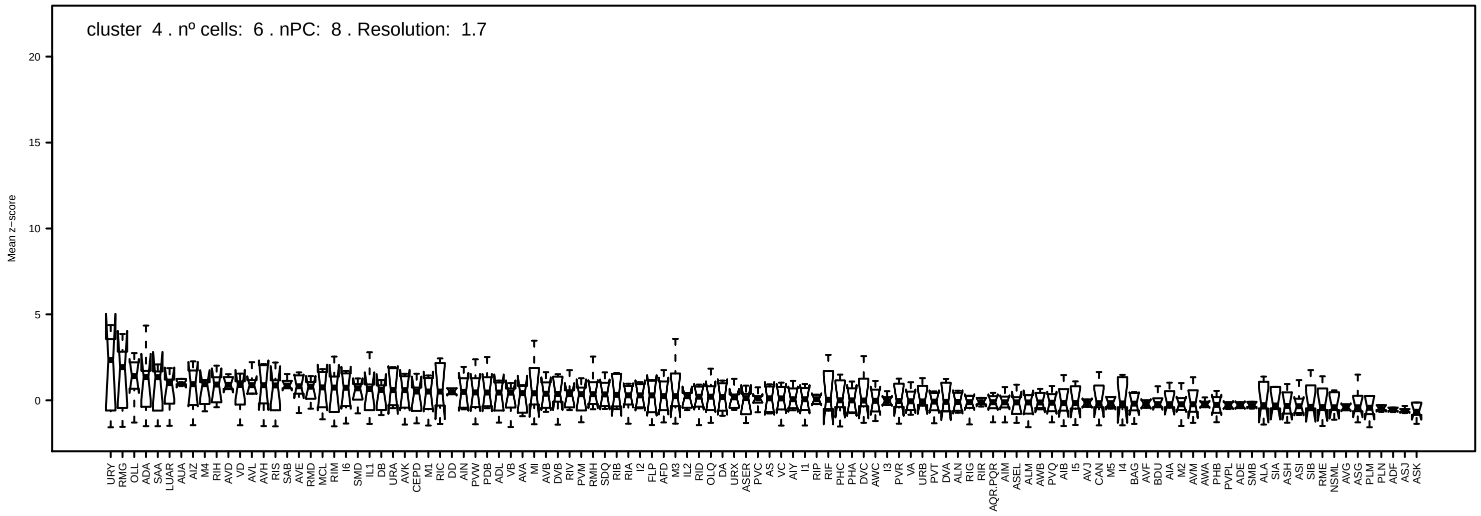
cluster 4 . nº cells: 6 . nPC: 8 . Resolution: 1.7
Mean z−score
20
15
10
5
0

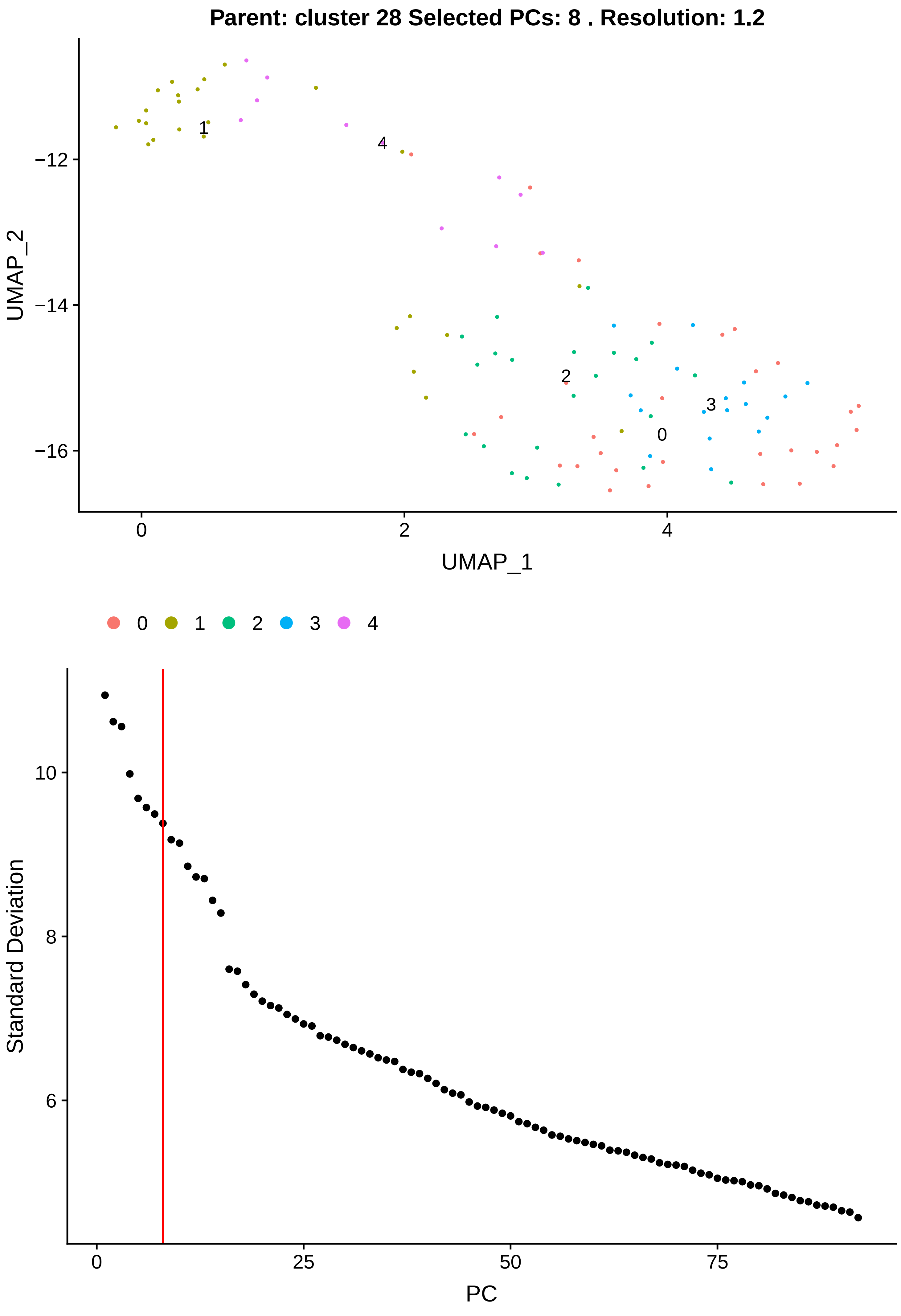

Parent: cluster 28 Selected PCs: 8 . Resolution: 1.2
UMAP_2
UMAP_1
−12
−14
−16
0
2
4
0
1
2
3
4
Standard Deviation
PC
10
8
6
0
25
50
75

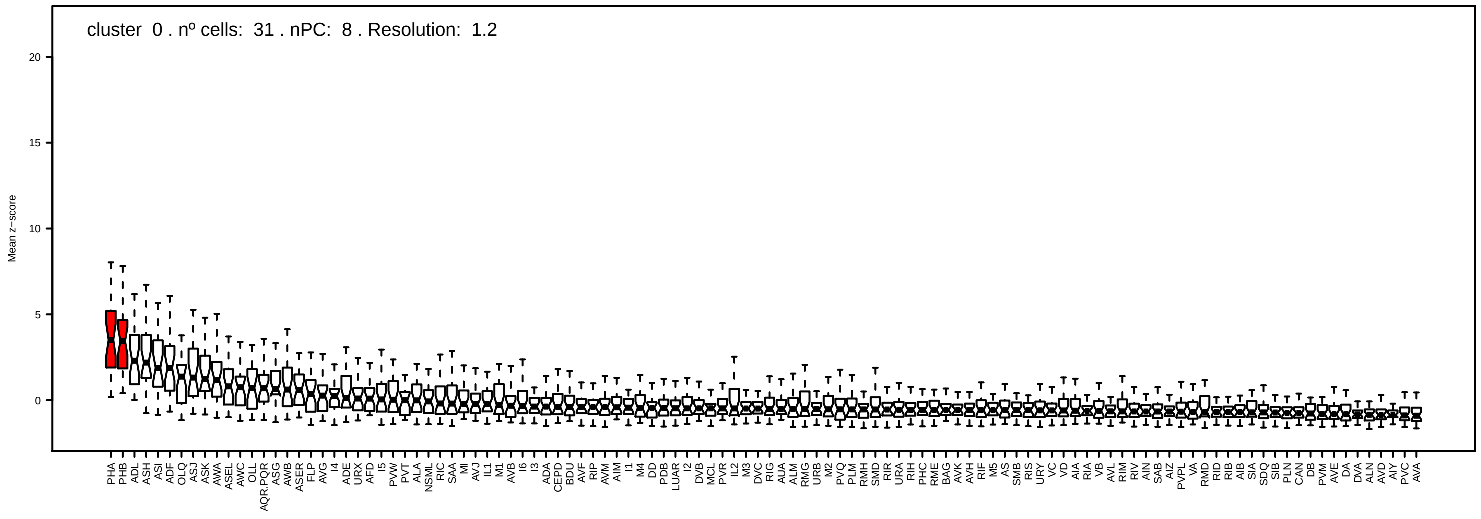
cluster 0 . nº cells: 31 . nPC: 8 . Resolution: 1.2
Mean z−score
20
15
10
5
0

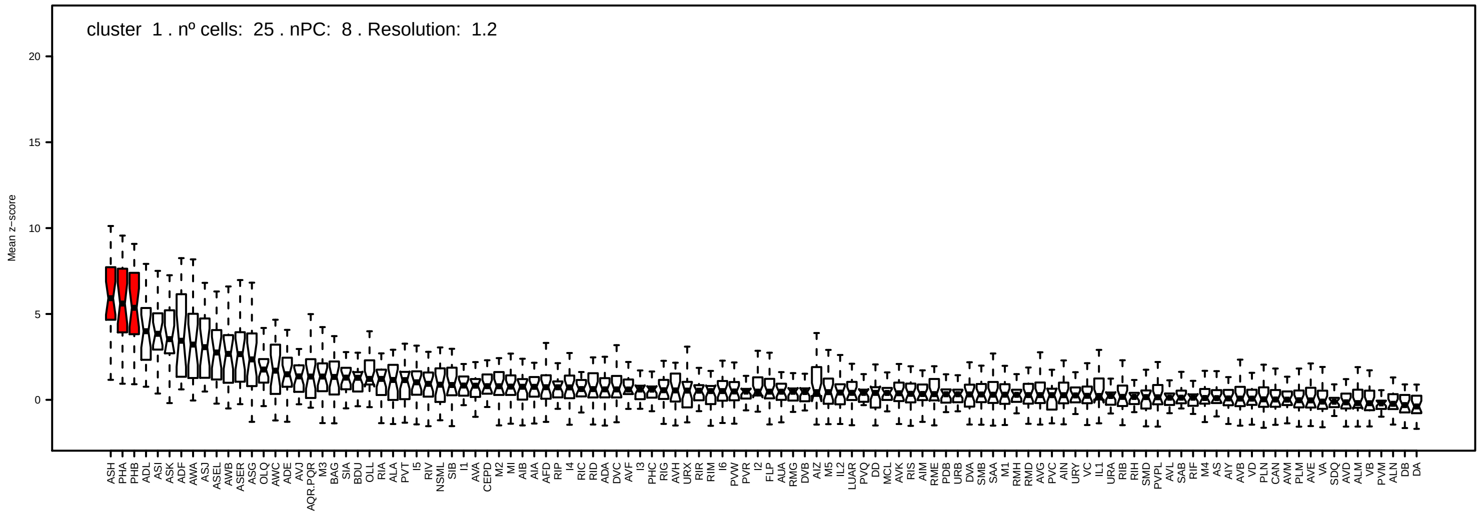
cluster 1 . nº cells: 25 . nPC: 8 . Resolution: 1.2
Mean z−score
20
15
10
5
0

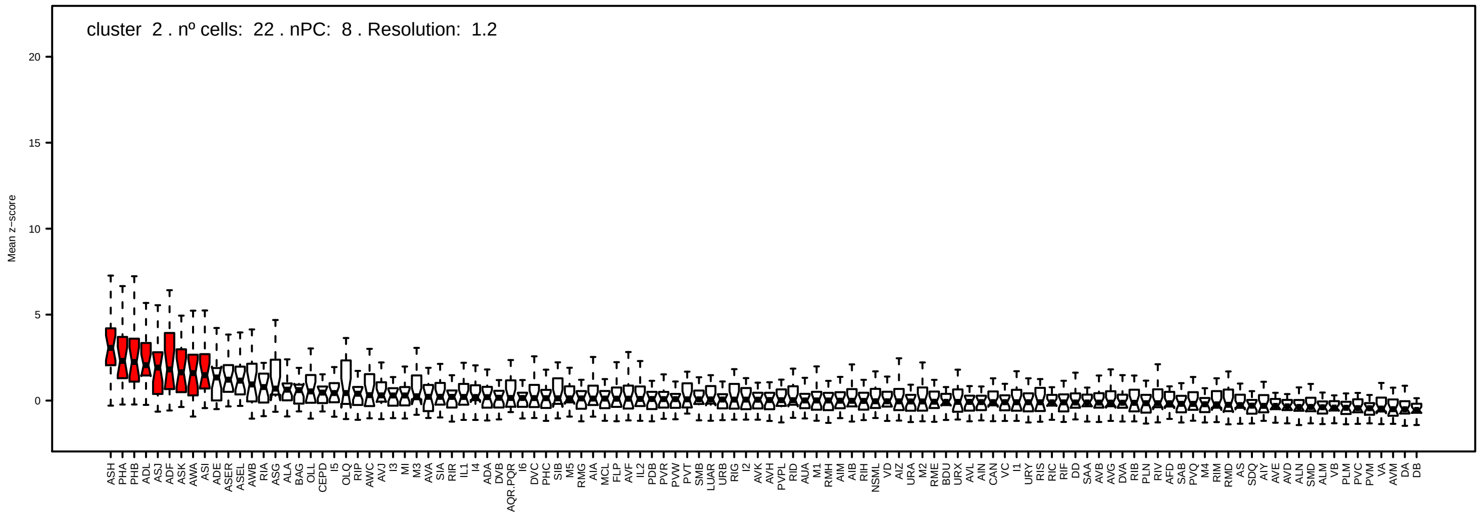
cluster 2 . nº cells: 22 . nPC: 8 . Resolution: 1.2
Mean z−score
20
15
10
5
0

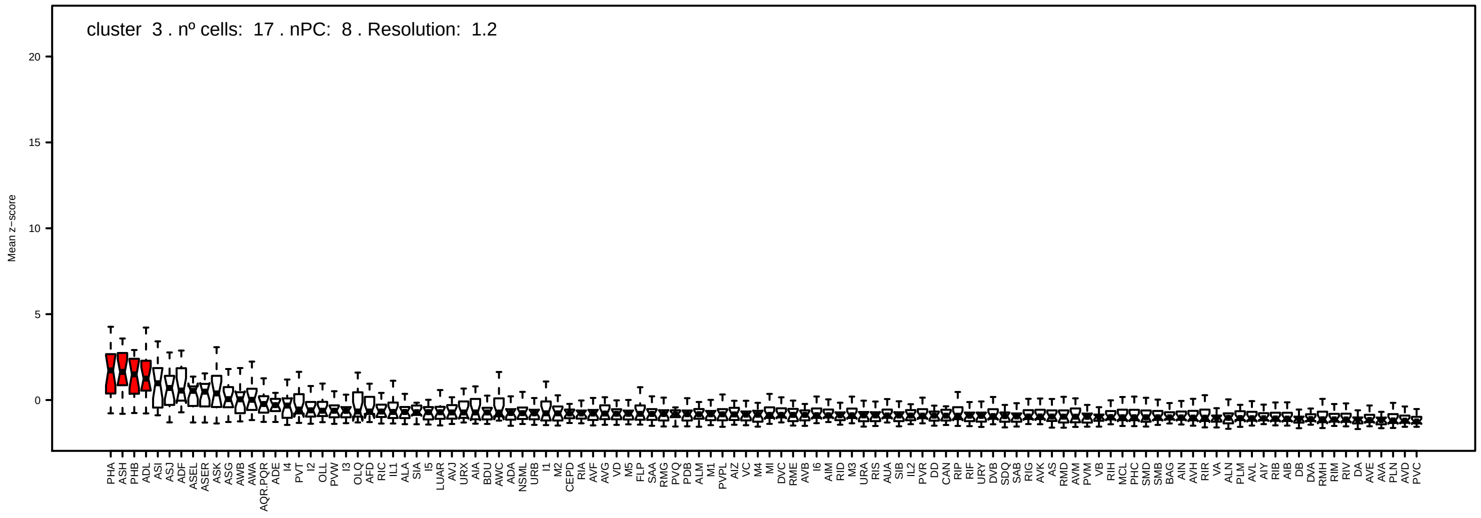
cluster 3 . nº cells: 17 . nPC: 8 . Resolution: 1.2
Mean z−score
20
15
10
5
0

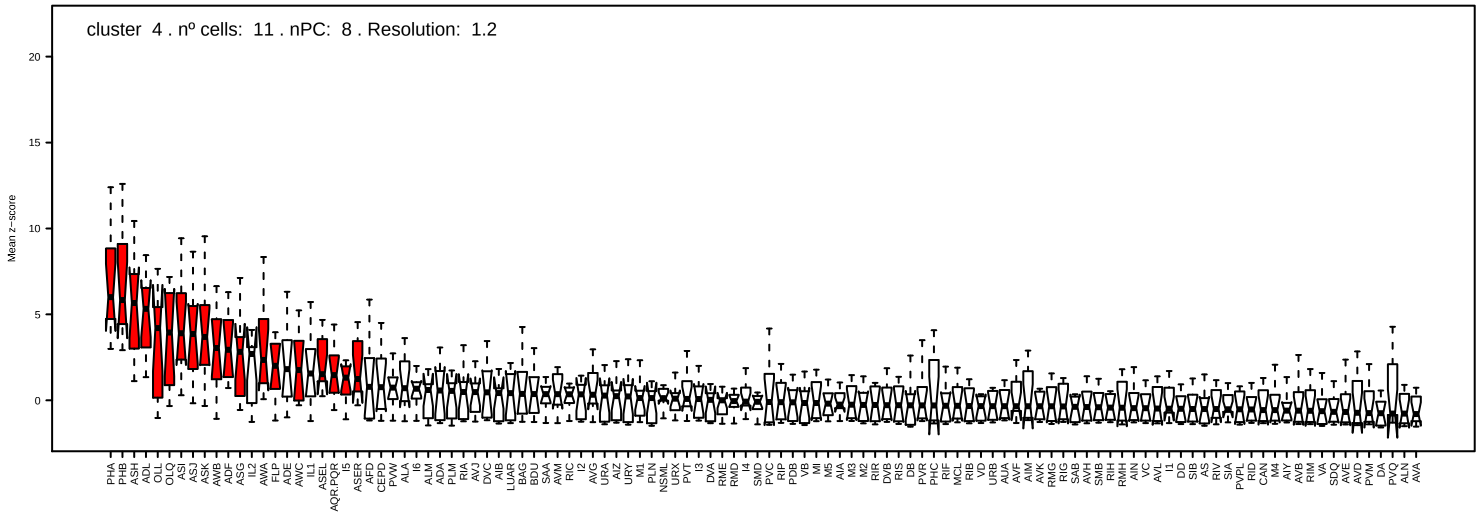

cluster 4 . nº cells: 11 . nPC: 8 . Resolution: 1.2
Mean z−score

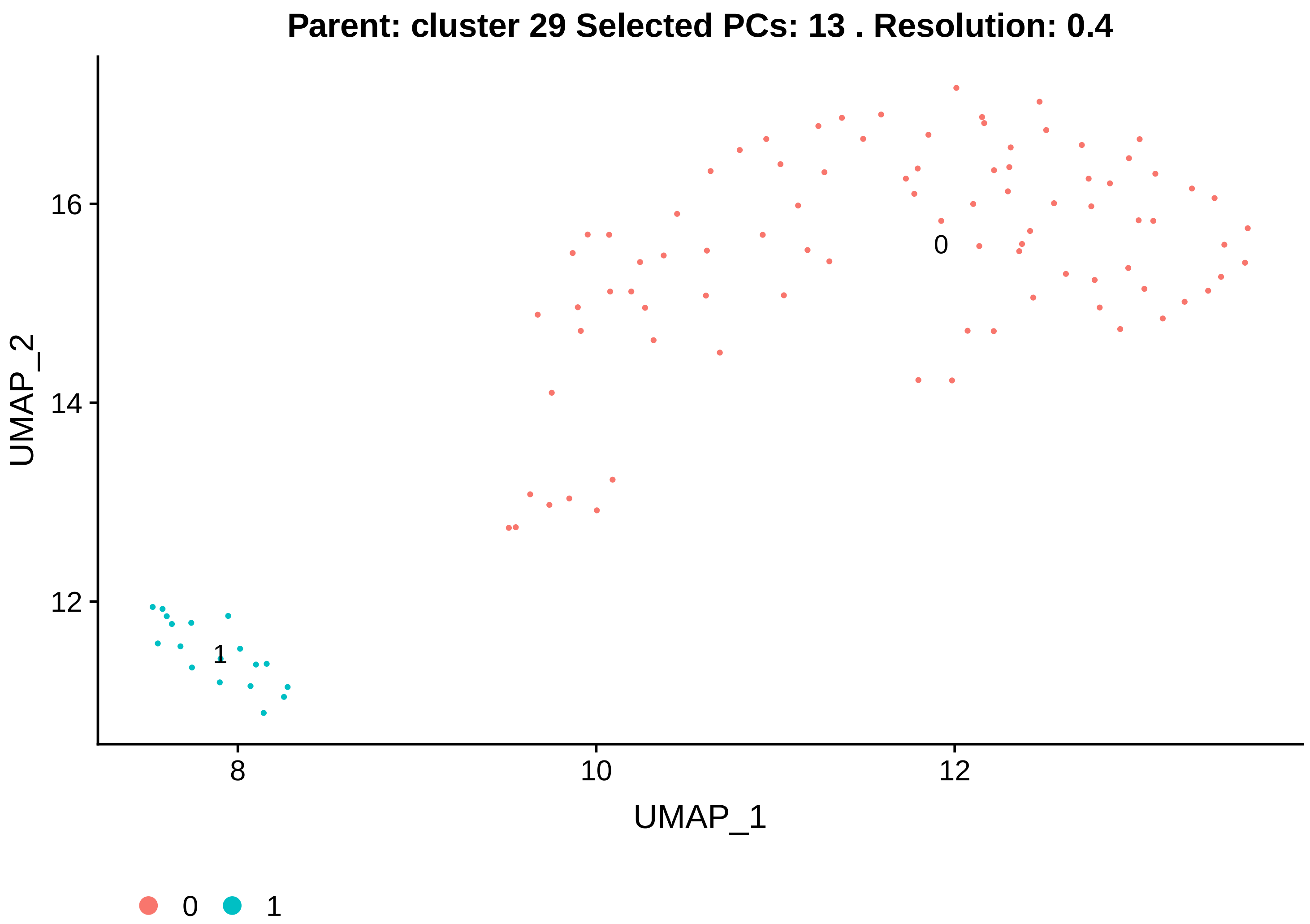
Parent: cluster 29 Selected PCs: 13 . Resolution: 0.4
UMAP_2
UMAP_1
16
14
12
8
10
12
0
1
0
1

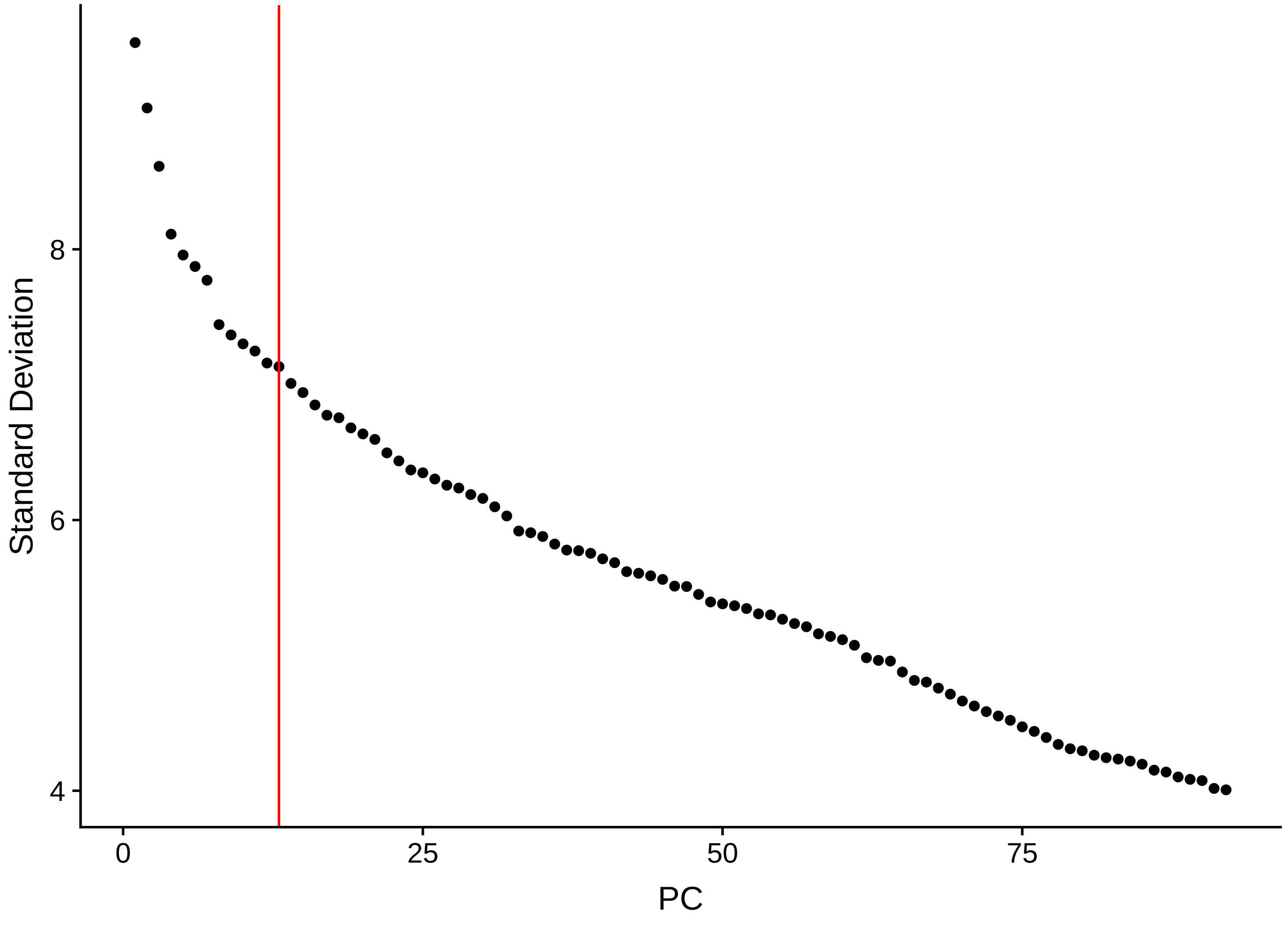
Standard Deviation
PC
8
6
4
0
25
50
75

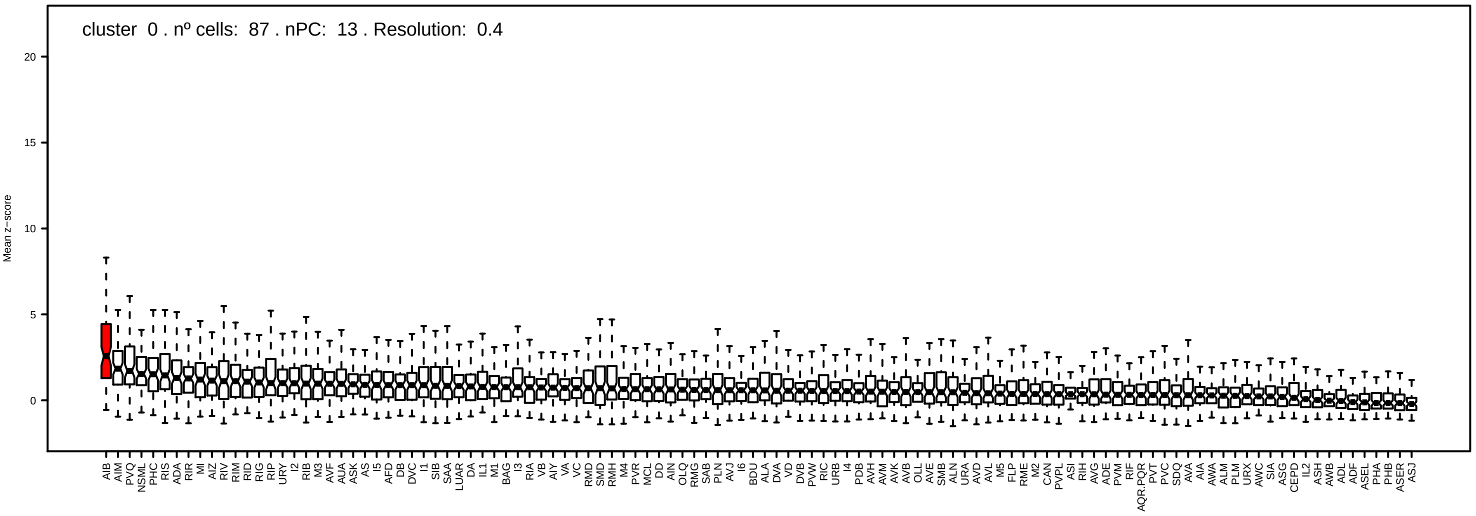
cluster 0 . nº cells: 87 . nPC: 13 . Resolution: 0.4
Mean z−score
20
15
10
5
0

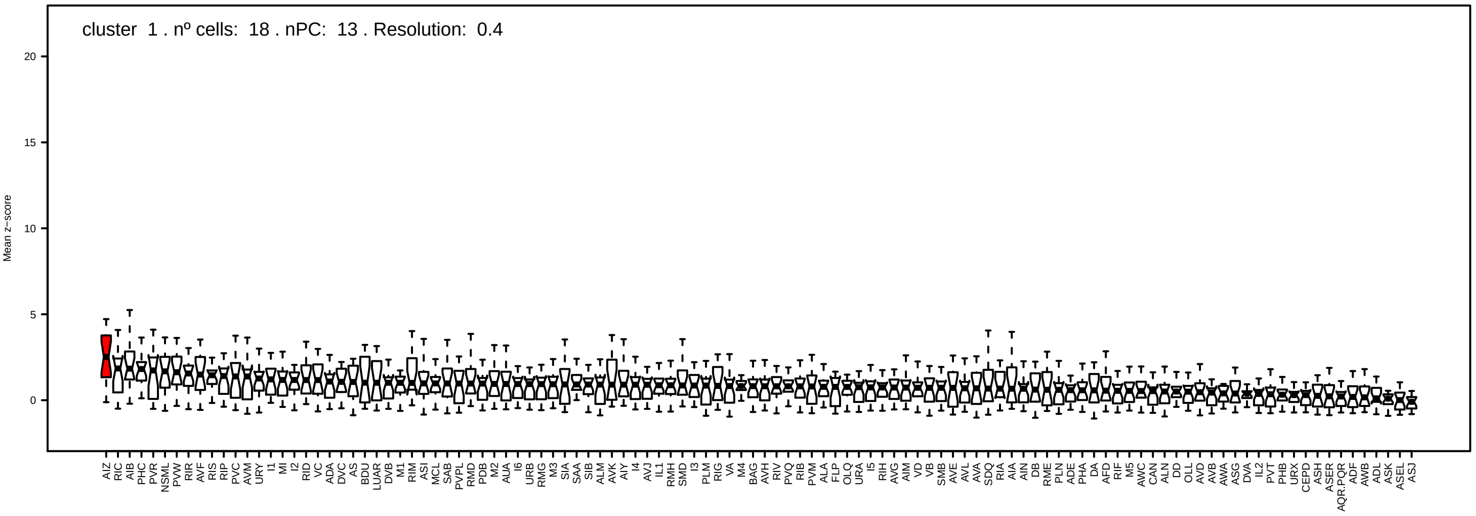
cluster 1 . nº cells: 18 . nPC: 13 . Resolution: 0.4
Mean z−score
20
15
10
5
0

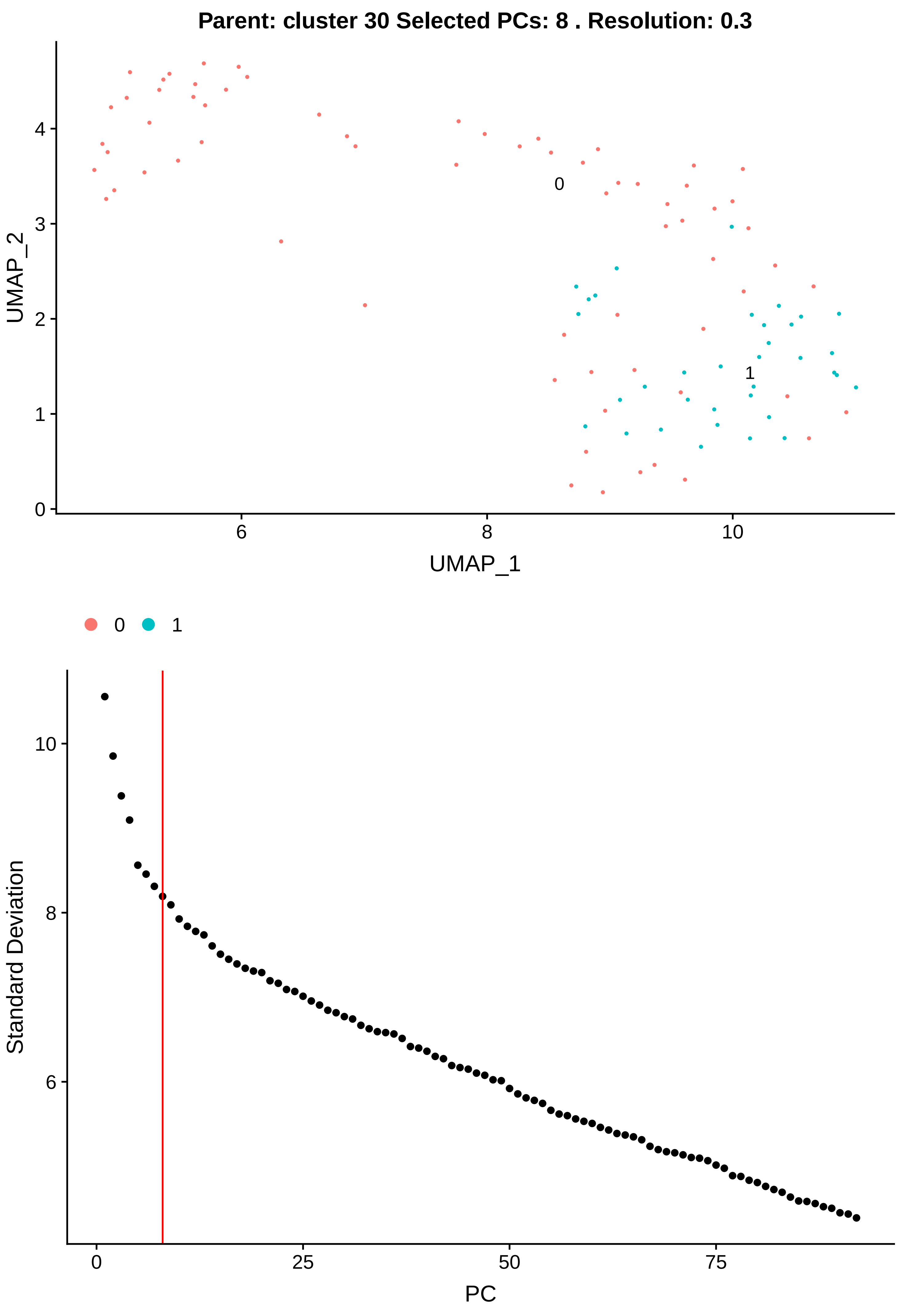

Parent: cluster 30 Selected PCs: 8 . Resolution: 0.3
UMAP_2
UMAP_1
0
1
6
8
10
0
1
2
3
4
0
1
Standard Deviation
PC
0
25
50
75
6
8
10

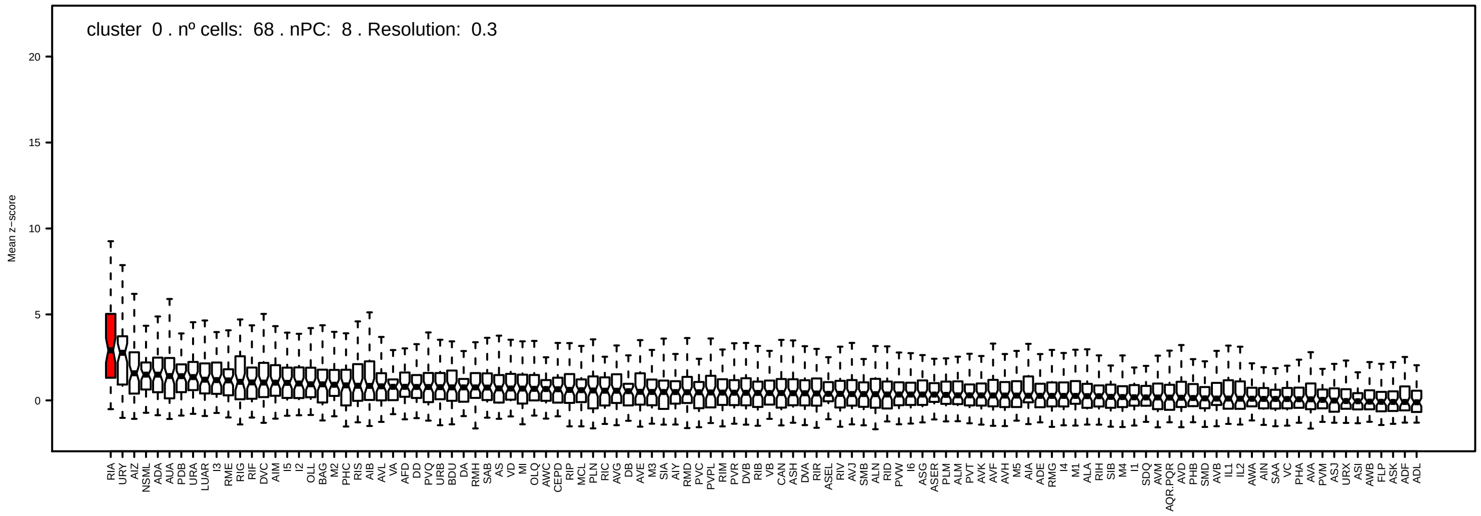
cluster 0 . nº cells: 68 . nPC: 8 . Resolution: 0.3
Mean z−score
20
15
10
5
0

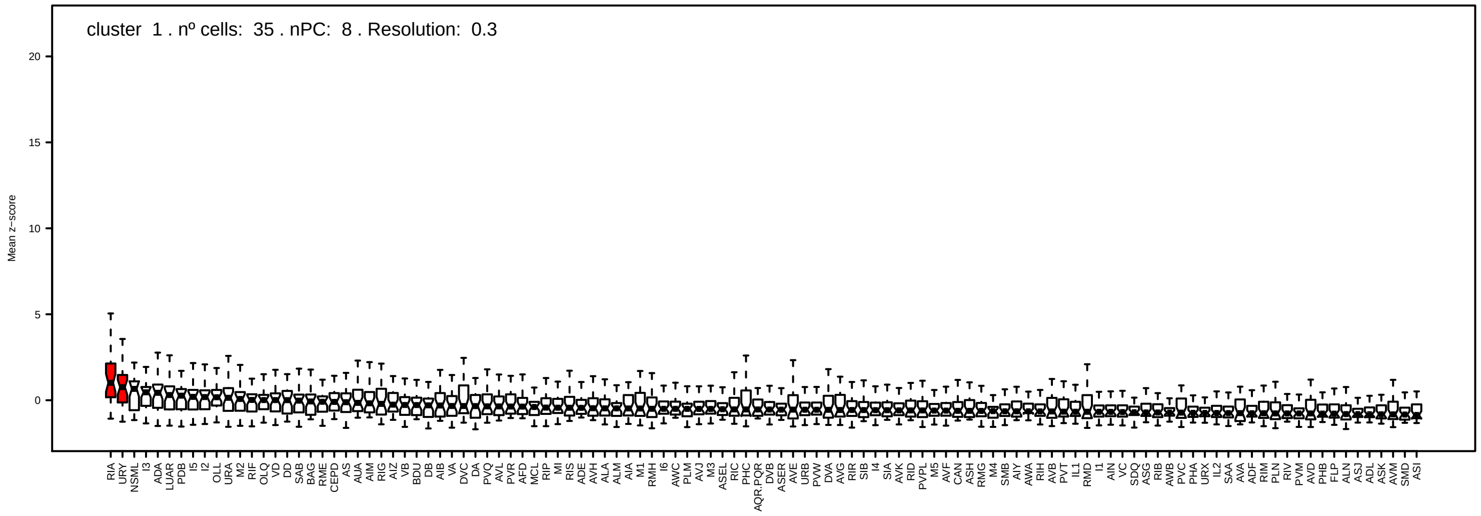
cluster 1 . nº cells: 35 . nPC: 8 . Resolution: 0.3
Mean z−score
20
15
10
5
0

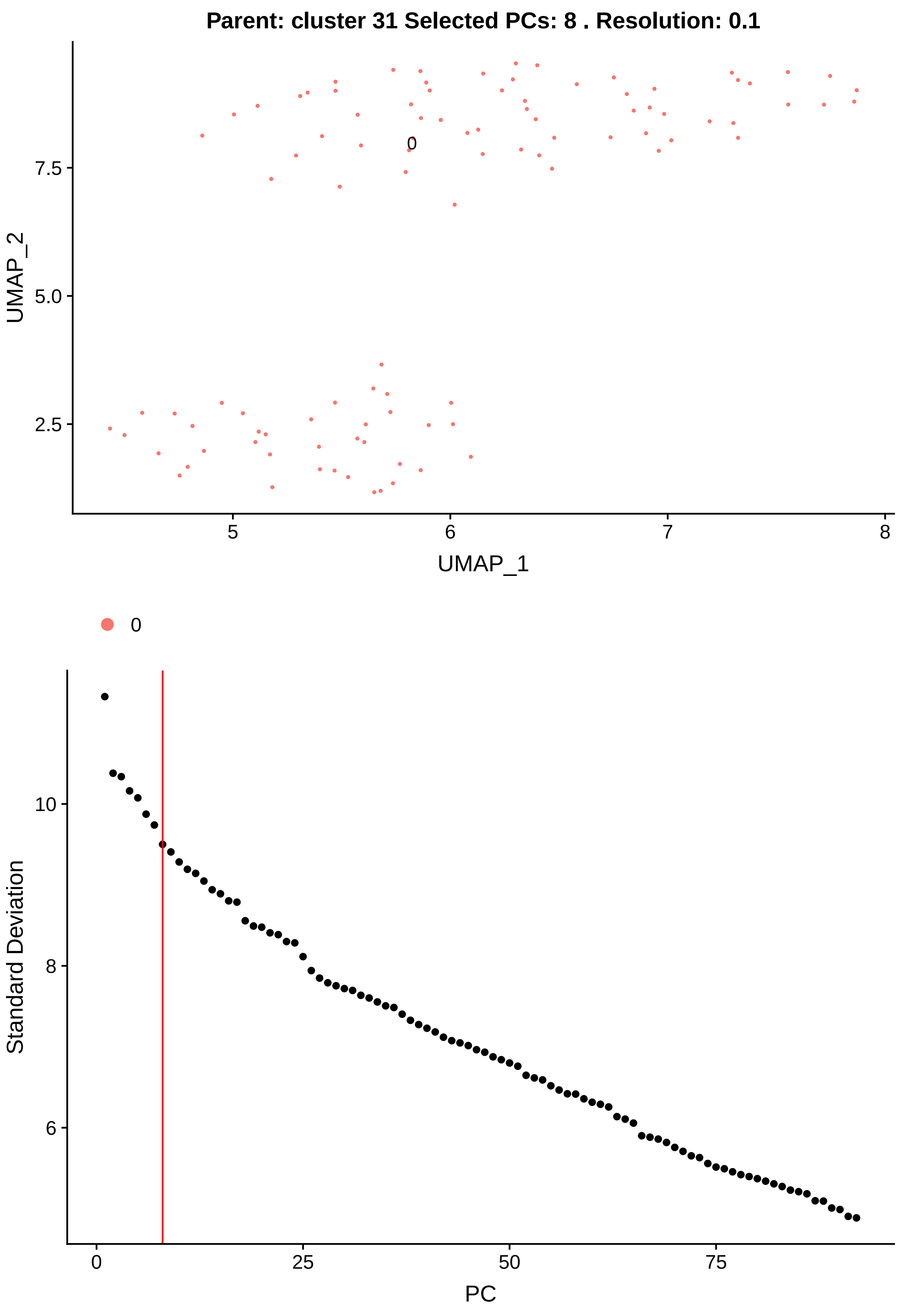

Parent: cluster 31 Selected PCs: 8 . Resolution: 0.1
0
UMAP_2
7.5
5.0
2.5
5
6
7
8
UMAP_1
0
Standard Deviation
10
8
6
0
25
50
75
PC

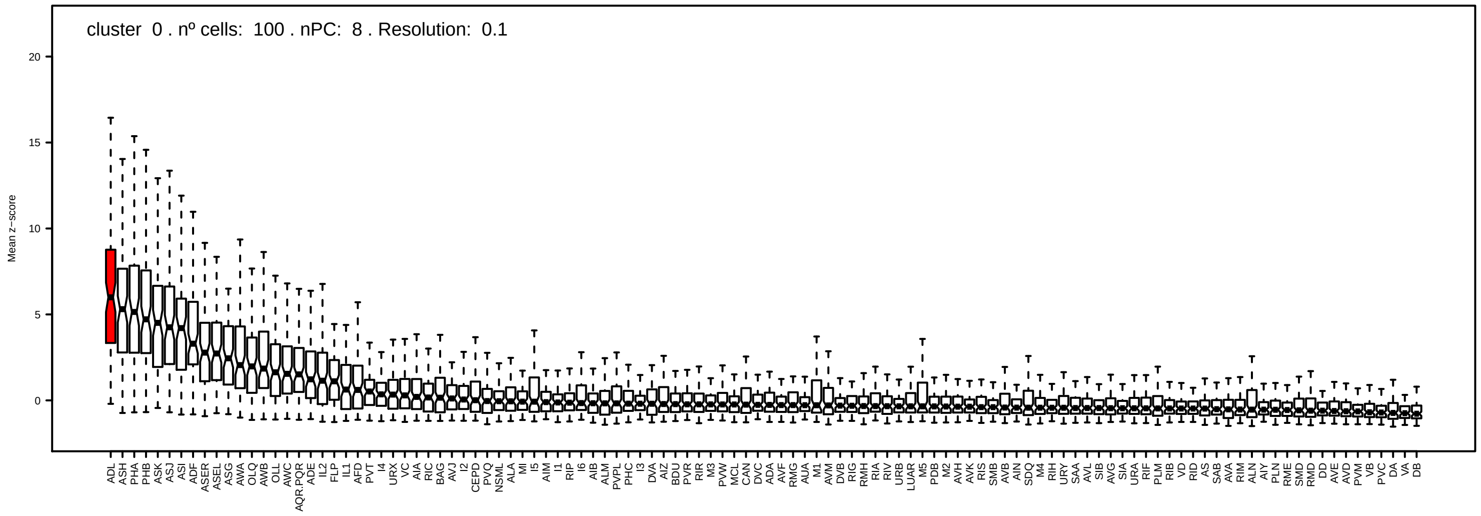
cluster 0 . nº cells: 100 . nPC: 8 . Resolution: 0.1
Mean z−score
20
15
10
5
0
ADL ASH PHA PHB ASK ASJ ASI ADF ASER ASEL ASG AWA OLQ AWB OLL AWC AQR.PQR ADE IL2 FLP IL1 AFD PVT I4 URX VC AIA RIC BAG AVJ I2 CEPD PVQ NSML ALA MI I5 AIM I1 RIP I6 AIB ALM PVPL PHC I3 DVA AIZ BDU PVR RIR M3 PVW MCL CAN DVC ADA AVF RMG AUA M1 AVM DVB RIG RMH RIA RIV URB LUAR M5 PDB M2 AVH AVK RIS SMB AVB AIN SDQ M4 RIH URY SAA AVL SIB AVG SIA URA RIF PLM RIB VD RID AS SAB AVA RIM ALN AIY PLN RME SMD RMD DD AVE AVD PVM VB PVC DA VA DB

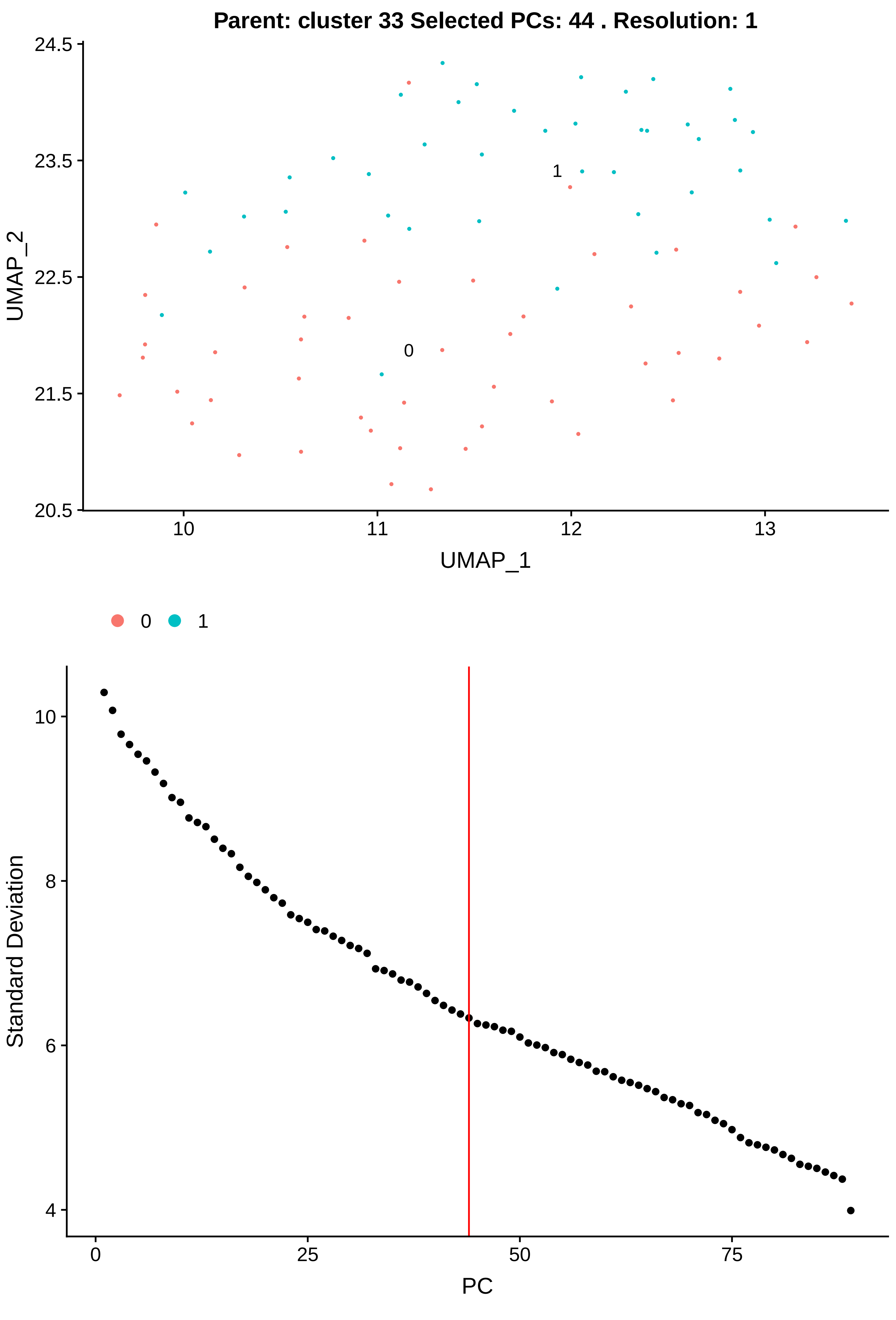

Parent: cluster 33 Selected PCs: 44 . Resolution: 1
24.5
23.5
22.5
21.5
20.5
UMAP_2
10
11
12
13
UMAP_1
0
1
0
1
10
8
6
4
Standard Deviation
0
25
50
75
PC

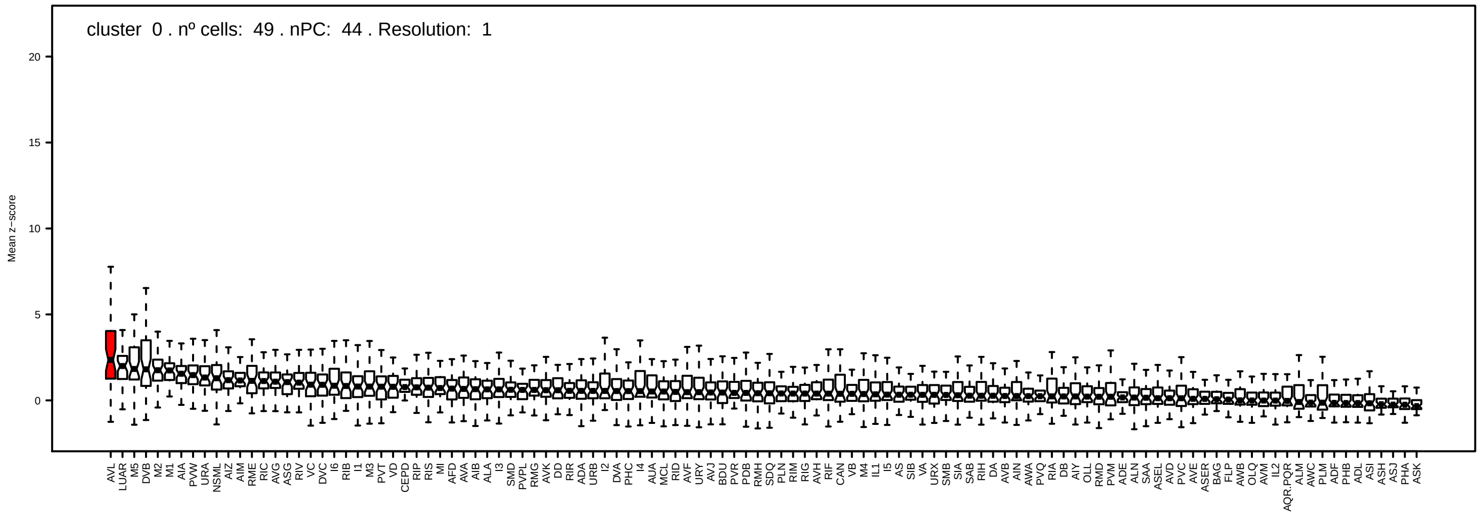
cluster 0 . nº cells: 49 . nPC: 44 . Resolution: 1
Mean z−score
20
15
10
5
0

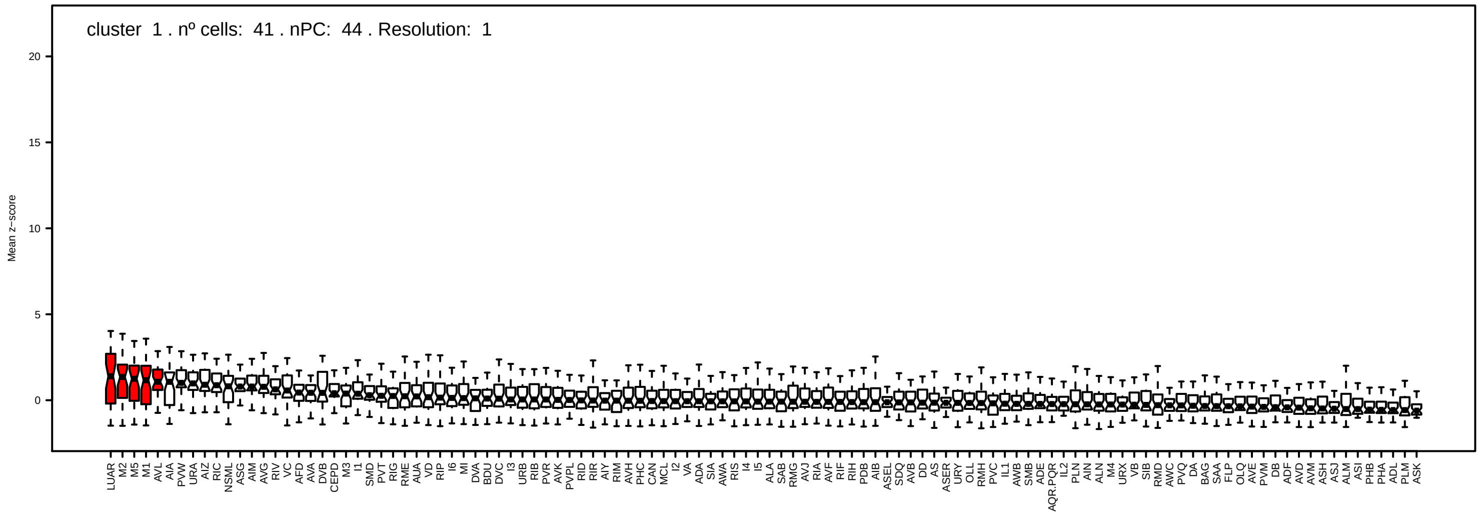
cluster 1 . nº cells: 41 . nPC: 44 . Resolution: 1
Mean z−score
20
15
10
5
0

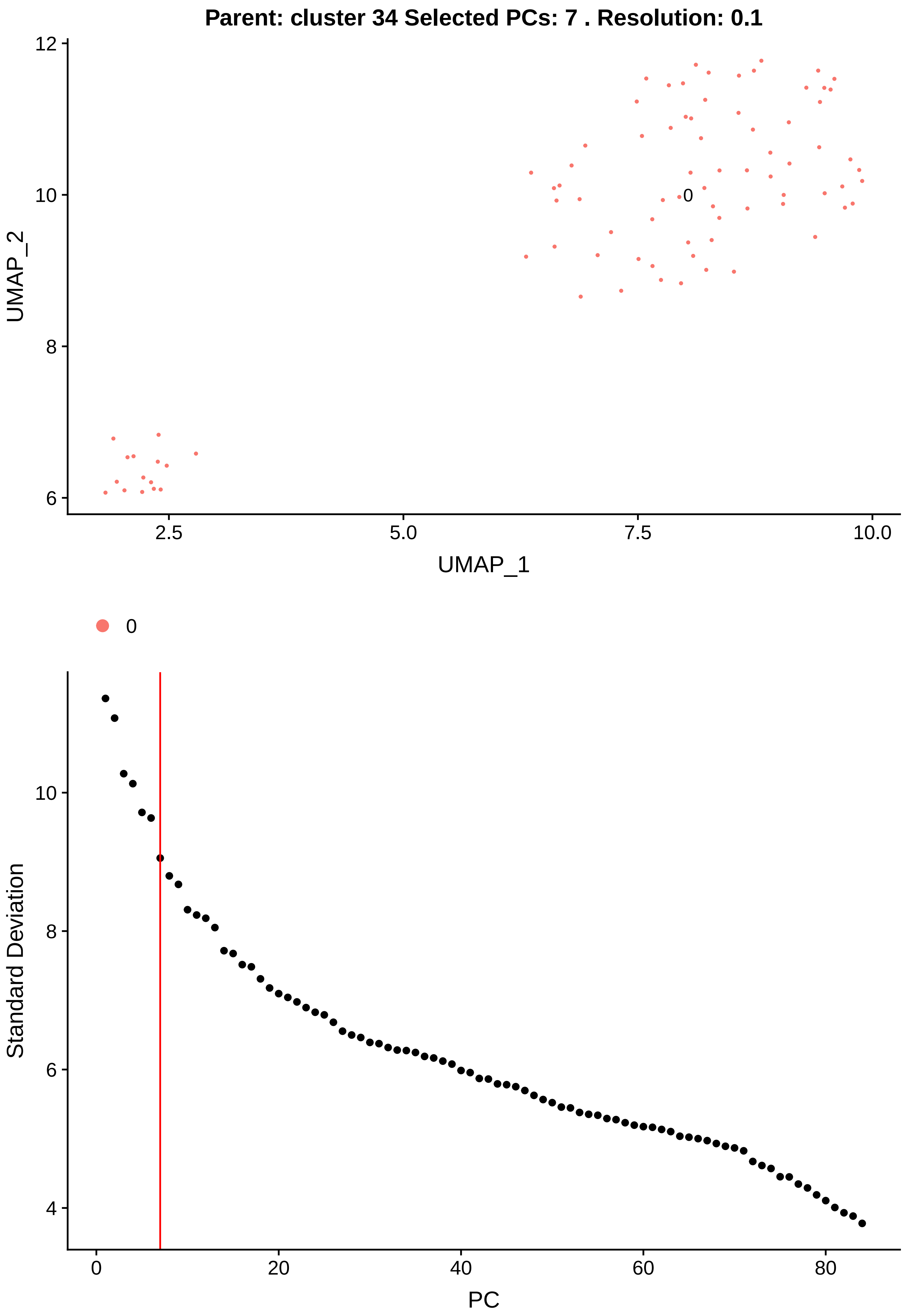

Parent: cluster 34 Selected PCs: 7 . Resolution: 0.1
UMAP_2
UMAP_1
0
Standard Deviation
PC

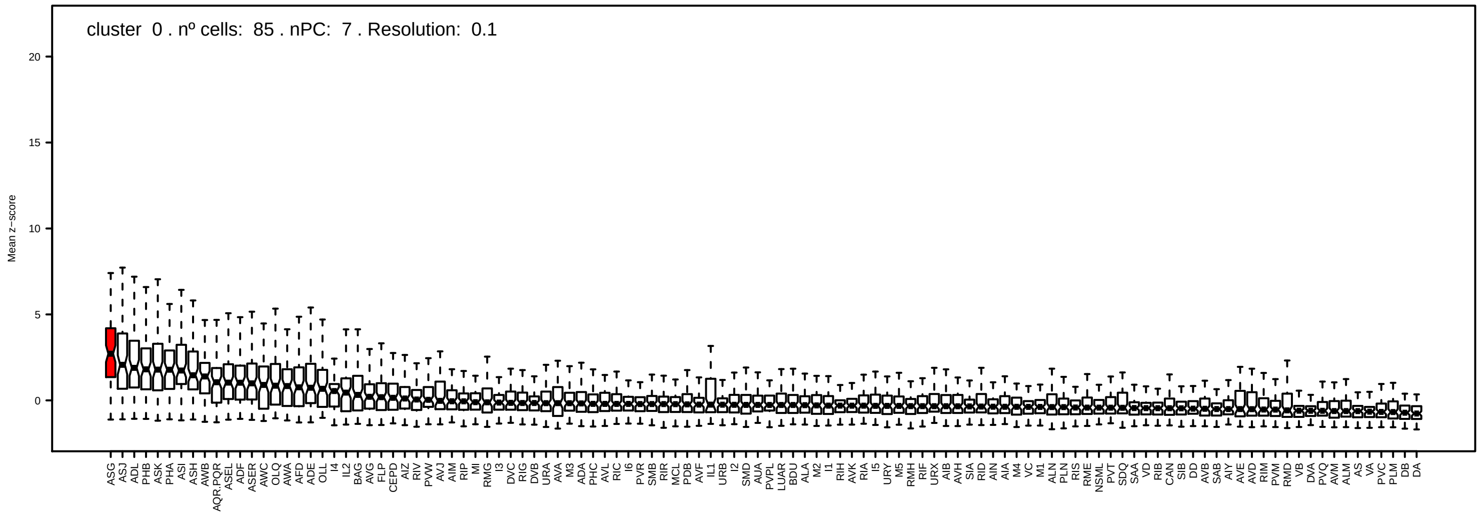
cluster 0 . nº cells: 85 . nPC: 7 . Resolution: 0.1
Mean z−score

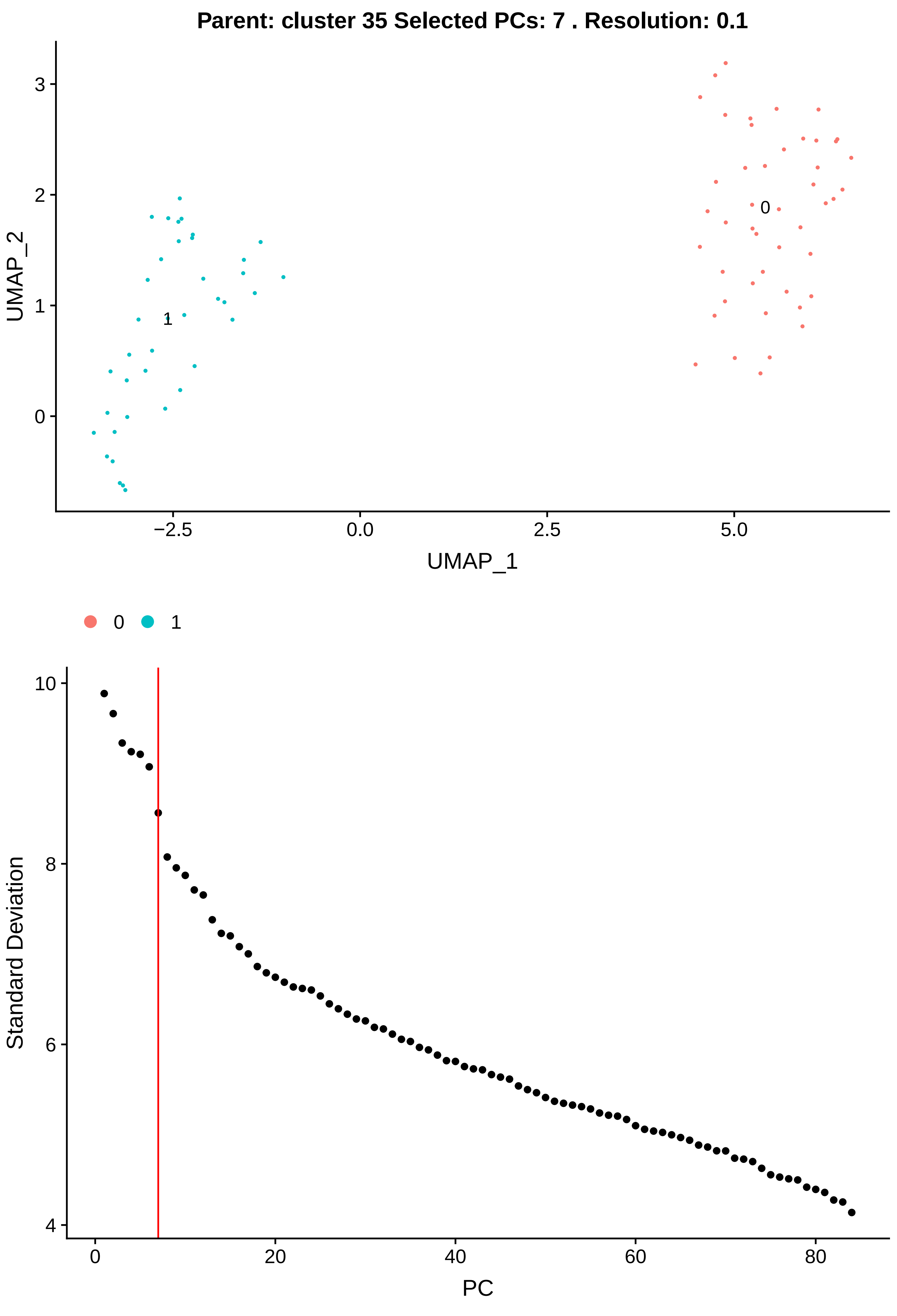

Parent: cluster 35 Selected PCs: 7 . Resolution: 0.1
UMAP_2
UMAP_1
3
2
1
0
−2.5
0.0
2.5
5.0
0
1
0
1
Standard Deviation
PC
10
8
6
4
0
20
40
60
80

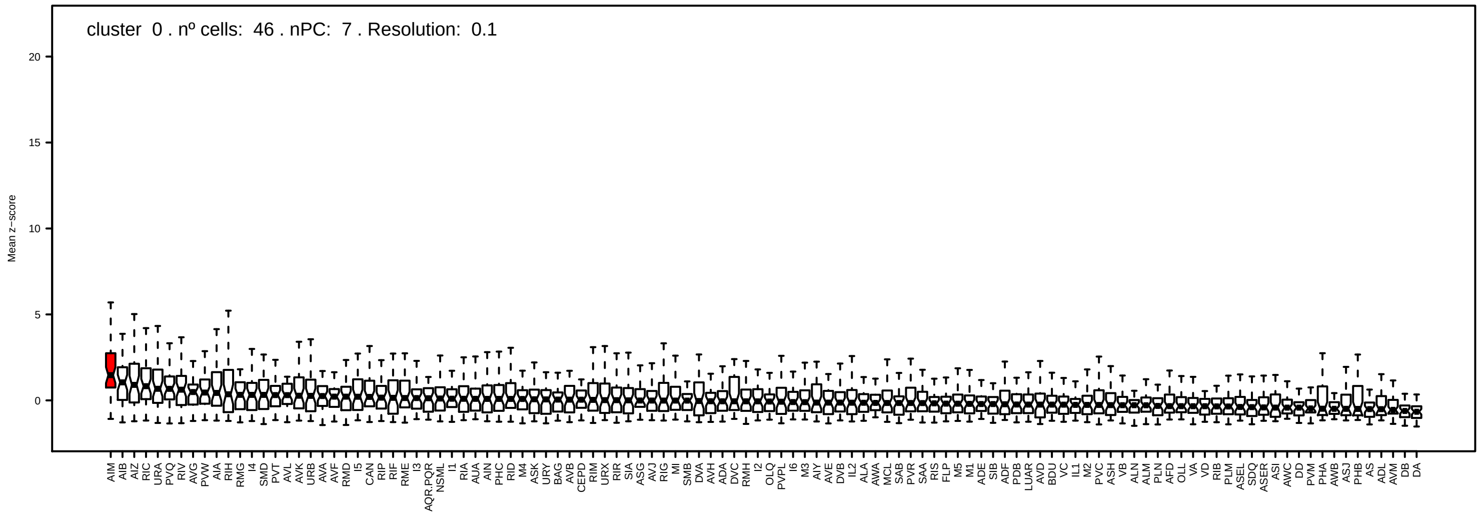
cluster 0 . nº cells: 46 . nPC: 7 . Resolution: 0.1
Mean z−score

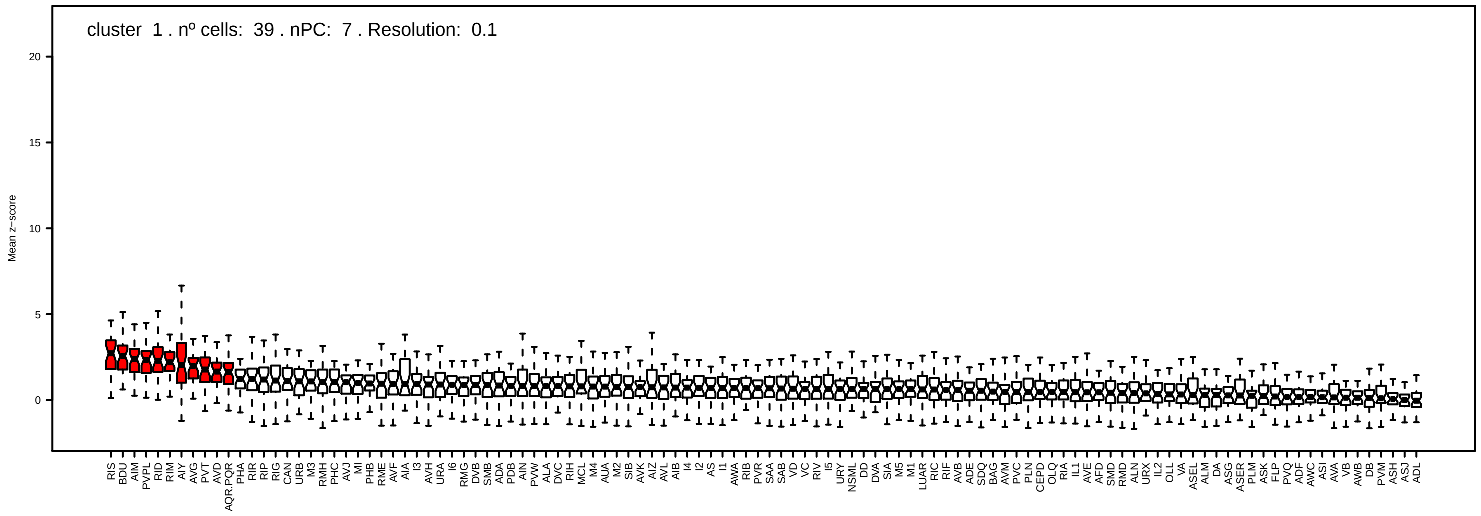
cluster 1 . nº cells: 39 . nPC: 7 . Resolution: 0.1
Mean z−score

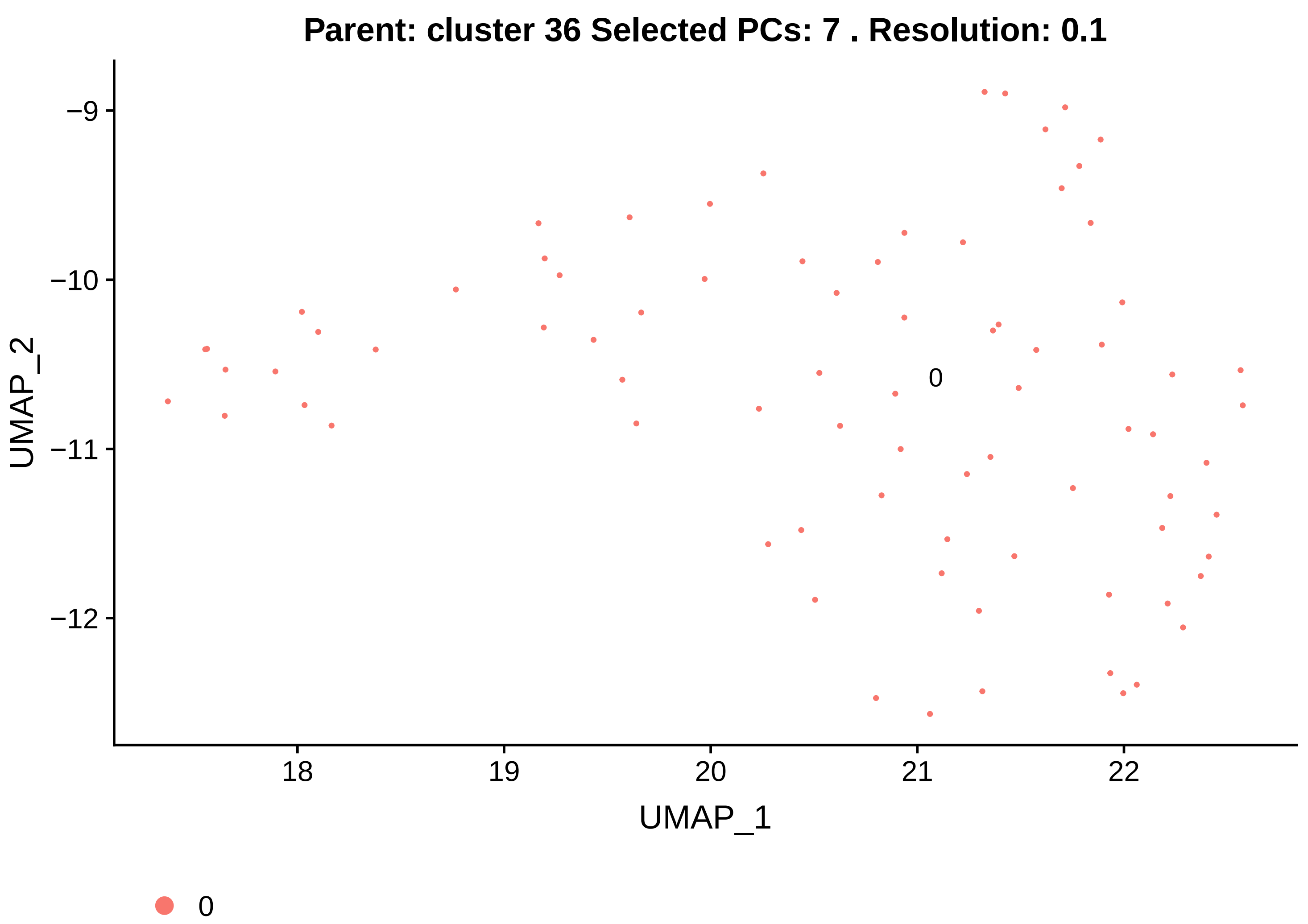
Parent: cluster 36 Selected PCs: 7 . Resolution: 0.1
−9
−10
−11
−12
UMAP_2
18
19
20
21
22
UMAP_1
0
0

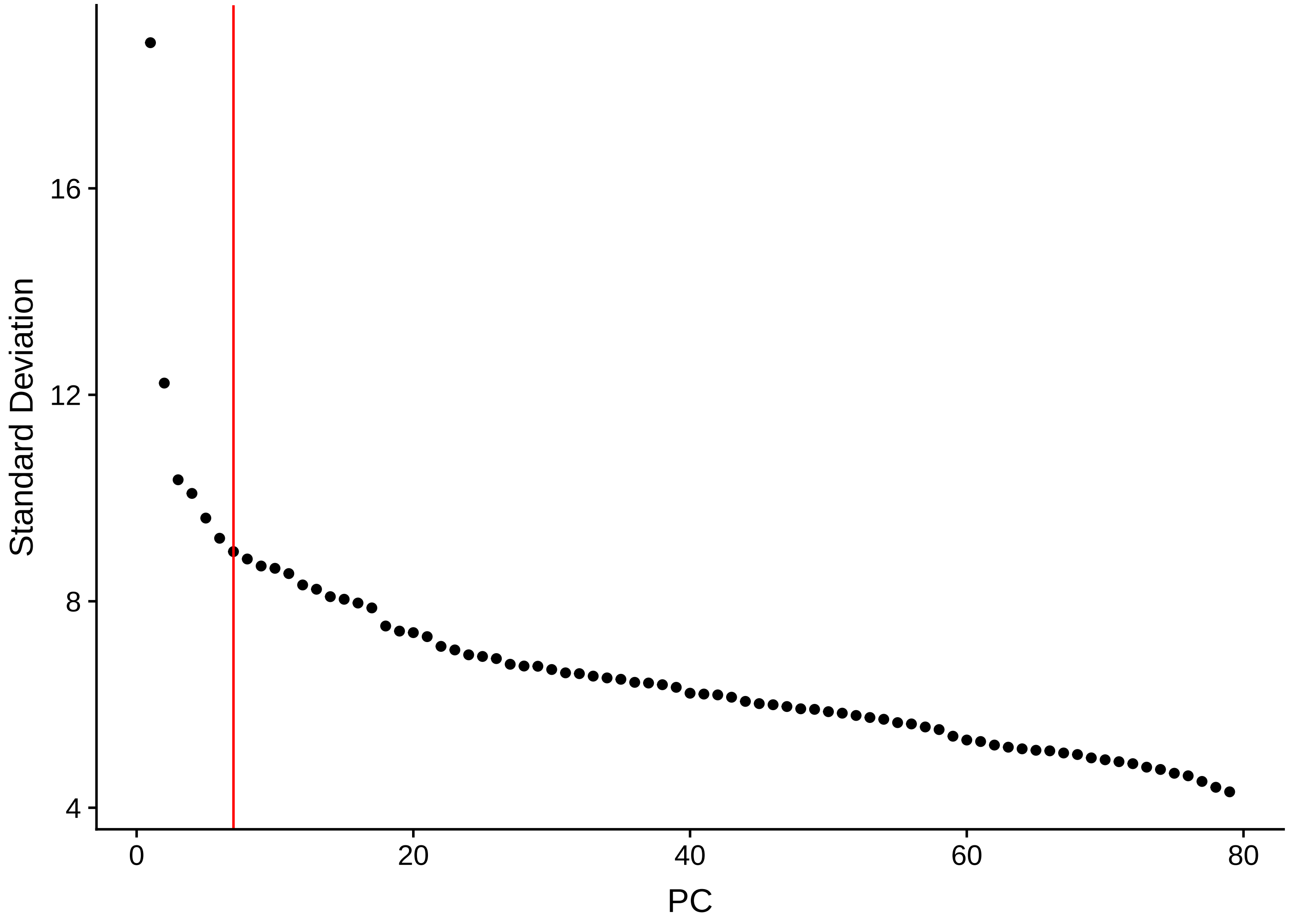
16
12
8
4
Standard Deviation
0
20
40
60
80
PC

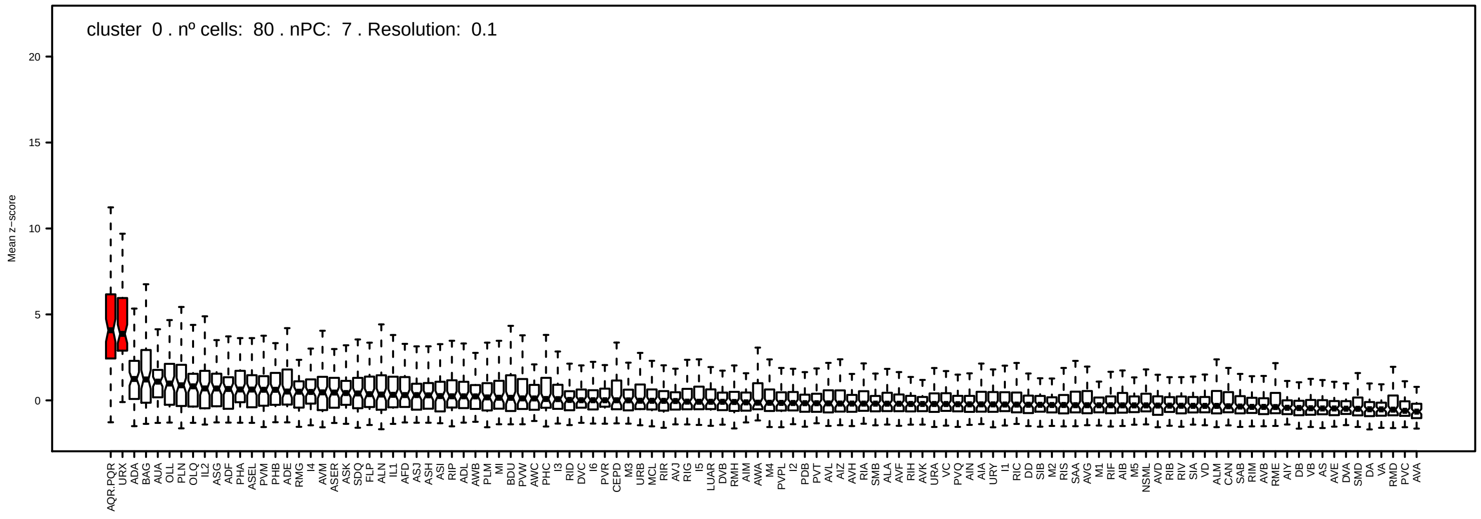
cluster 0 . nº cells: 80 . nPC: 7 . Resolution: 0.1
Mean z−score
20
15
10
5
0
AQR.PQR URX ADA BAG AUA OLL PLN OLQ IL2 ASG ADF PHA ASEL PVM PHB ADE RMG I4 AVM ASER ASK SDQ FLP ALN IL1 AFD ASJ ASH ASI RIP ADL AWB PLM MI BDU PVW AWC PHC I3 RID DVC I6 PVR CEPD M3 URB MCL RIR AVJ RIG I5 LUAR DVB RMH AIM AWA M4 PVPL I2 PDB PVT AVL AIZ AVH RIA SMB ALA AVF RIH AVK URA VC PVQ AIN AIA URY I1 RIC DD SIB M2 RIS SAA AVG M1 RIF AIB M5 NSML AVD RIB RIV SIA VD ALM CAN SAB RIM AVB RME AIY DB VB AS AVE DVA SMD DA VA RMD PVC AVA

Parent: cluster 37 Selected PCs: 5 . Resolution: 0.4

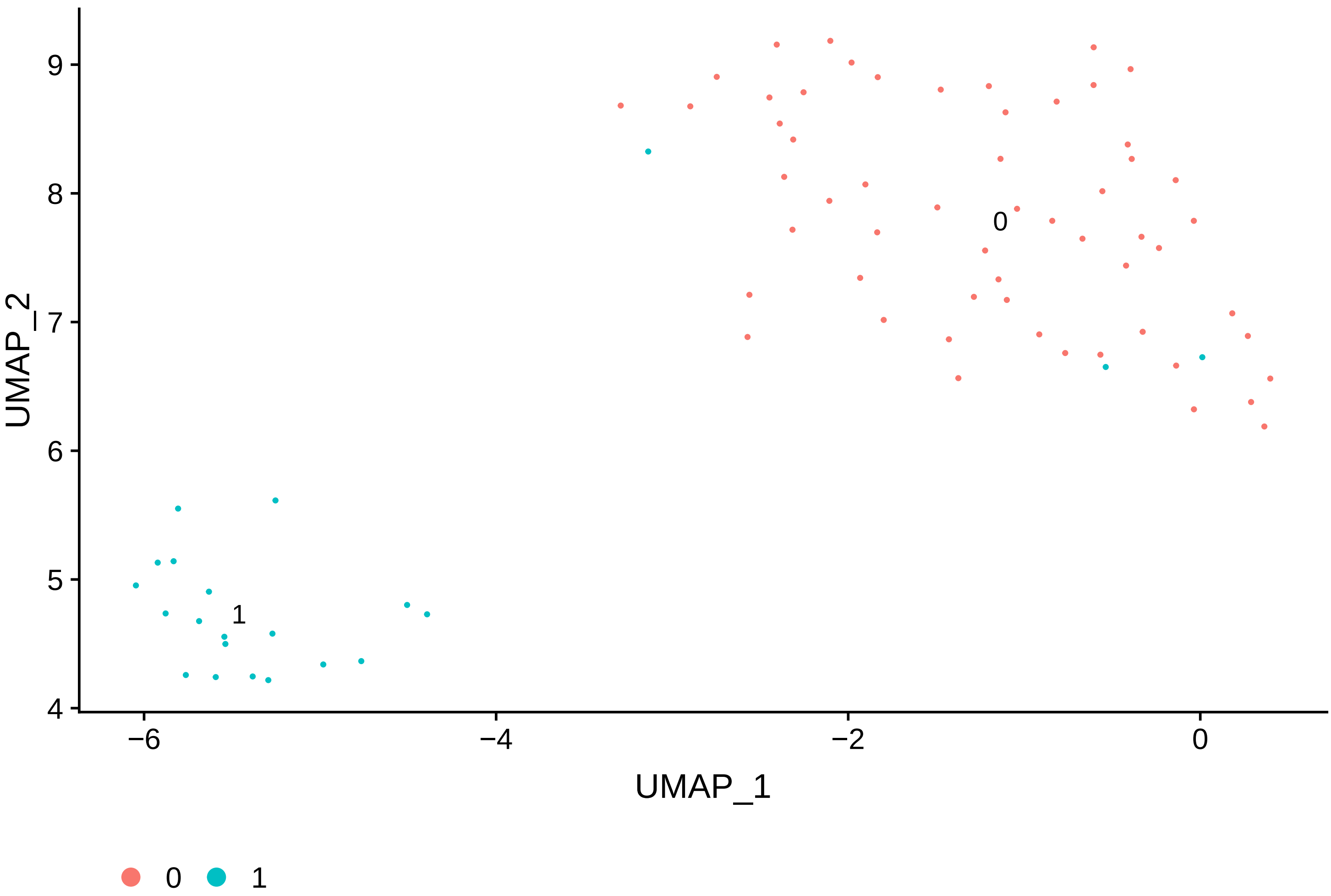


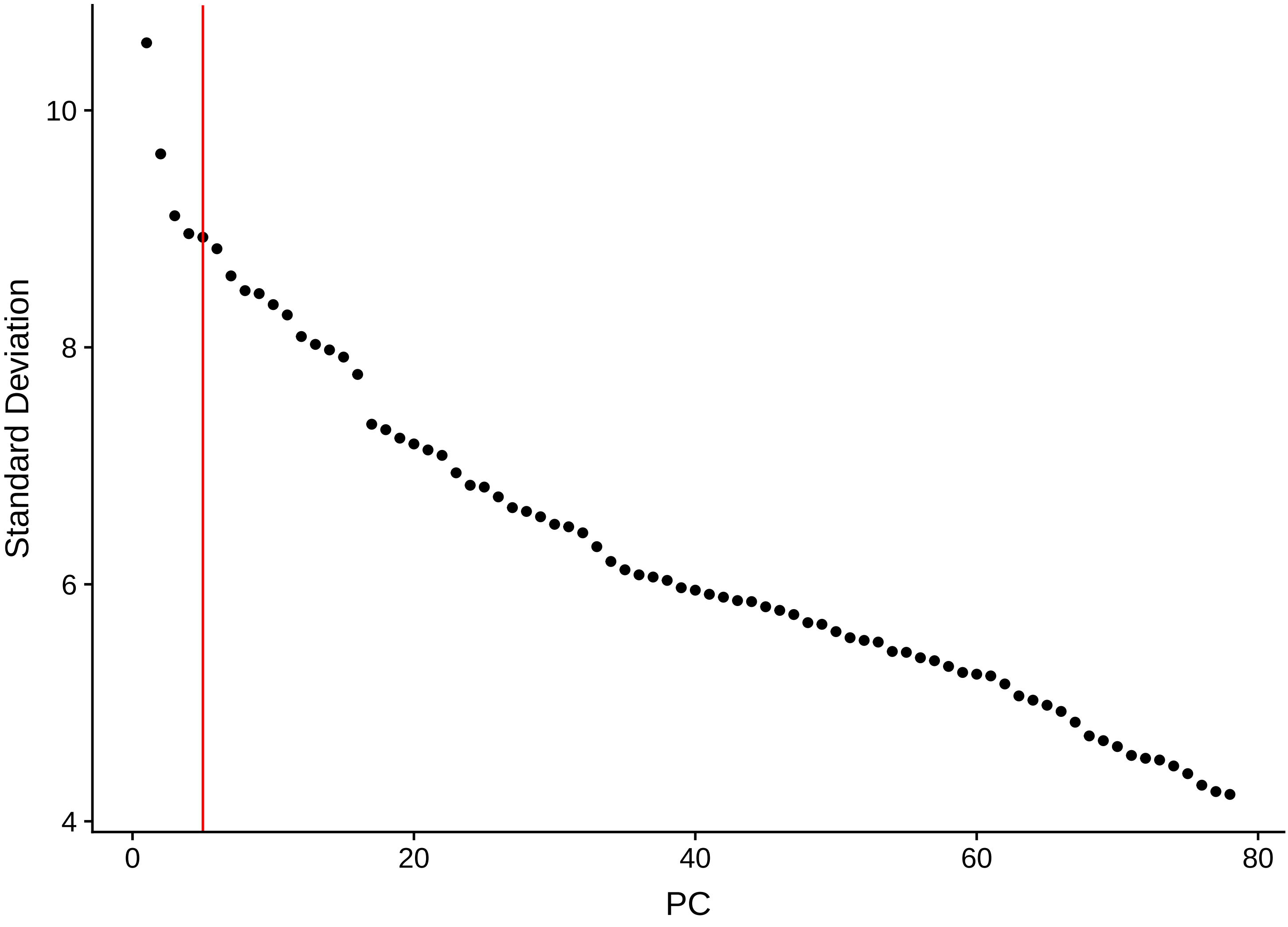

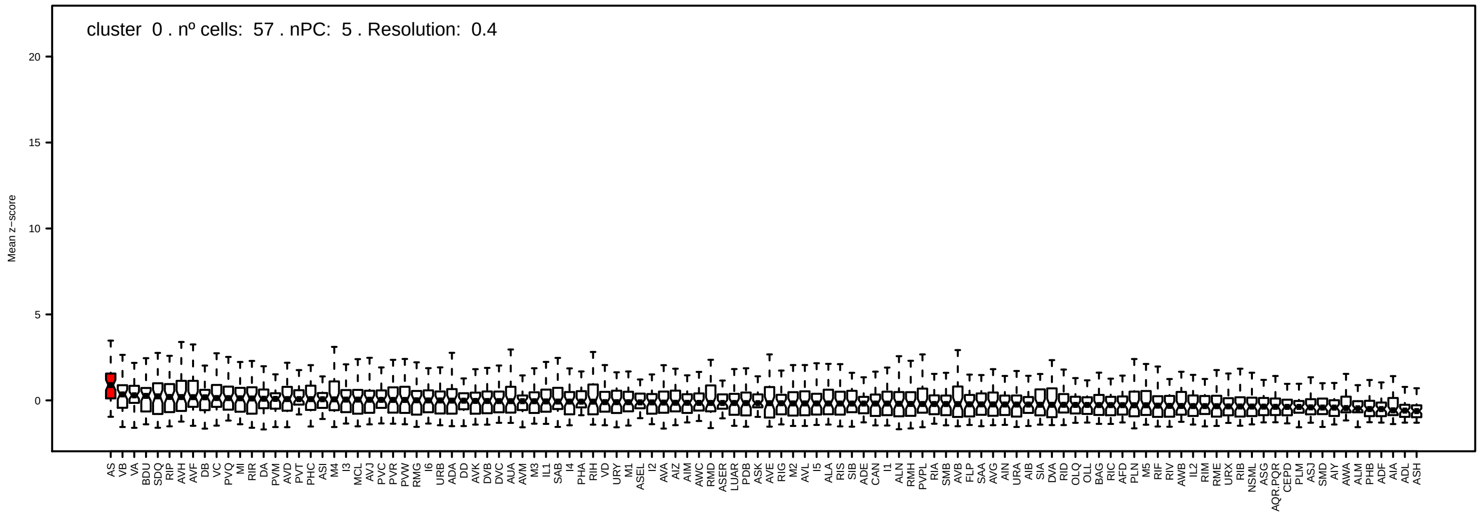

cluster 0 . nº cells: 57 . nPC: 5 . Resolution: 0.4
Mean z−score


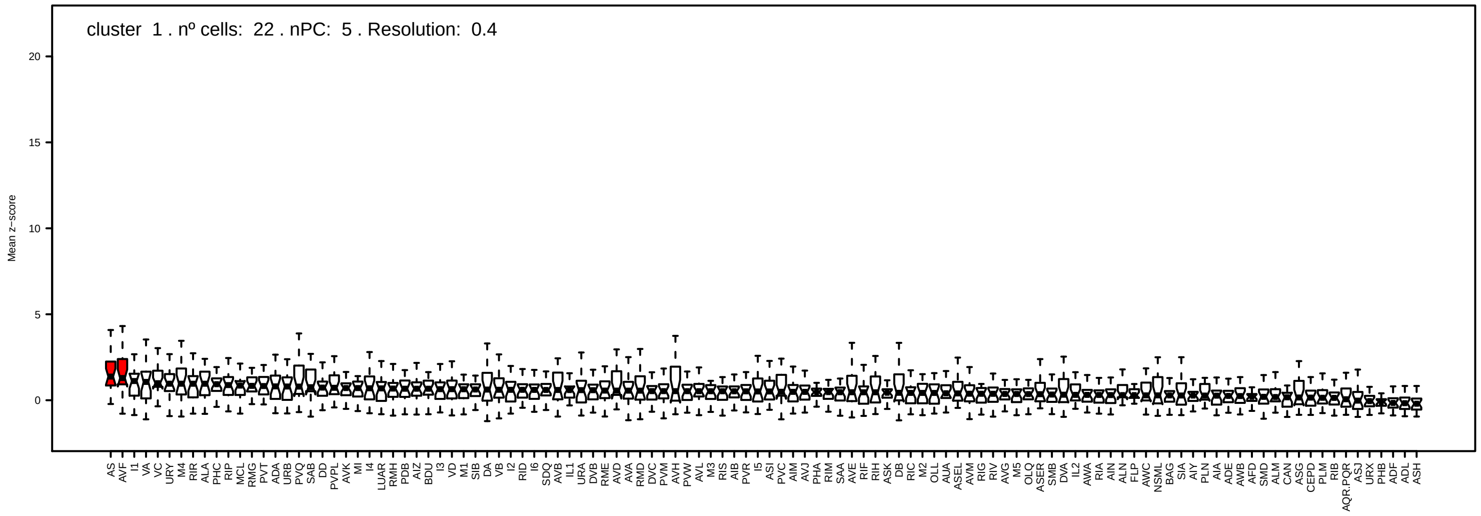

cluster 1 . nº cells: 22 . nPC: 5 . Resolution: 0.4
Mean z−score

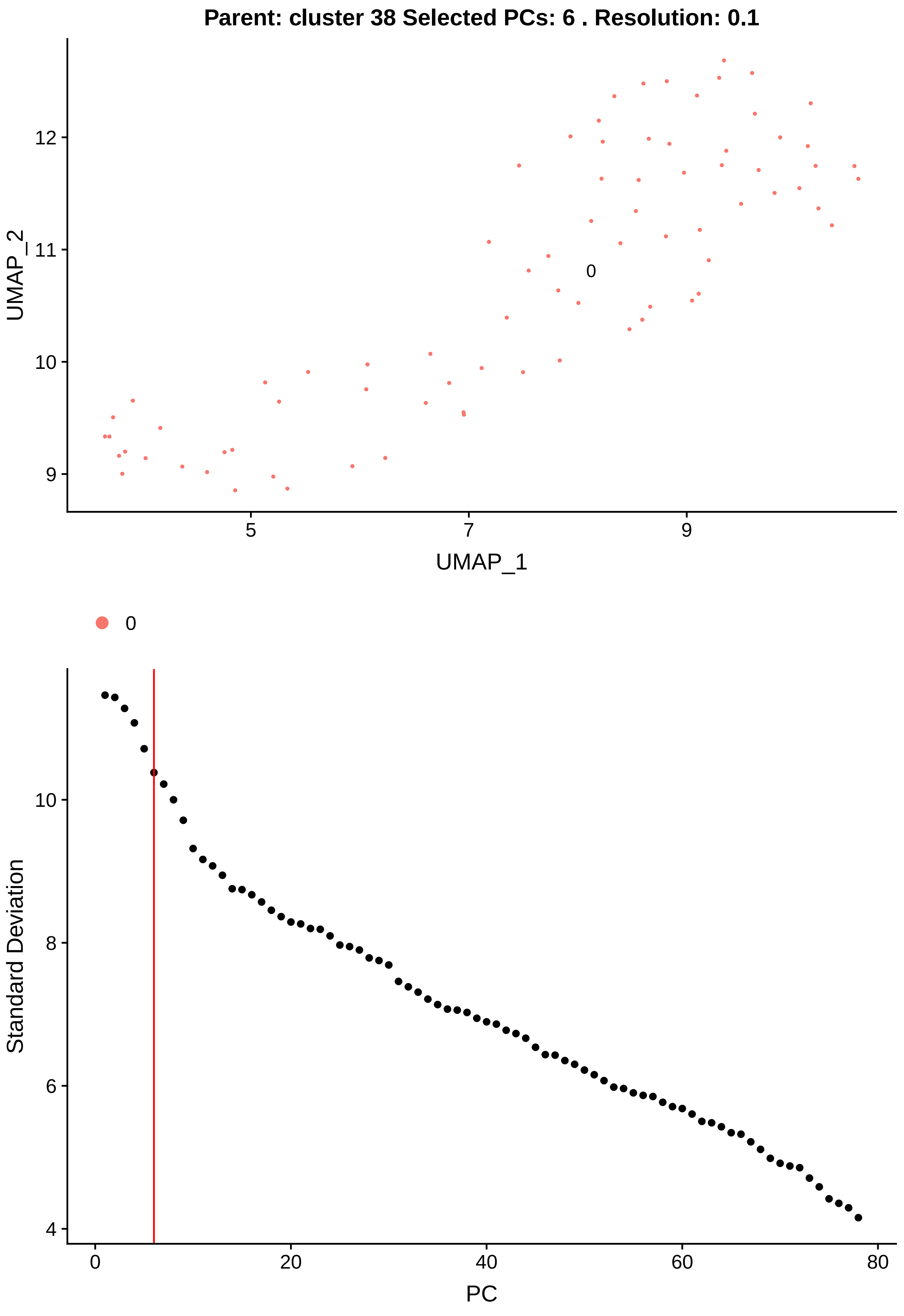

Parent: cluster 38 Selected PCs: 6 . Resolution: 0.1
UMAP_2
UMAP_1
12
11
10
9
5
7
9
0
0
Standard Deviation
PC
10
8
6
4
0
20
40
60
80

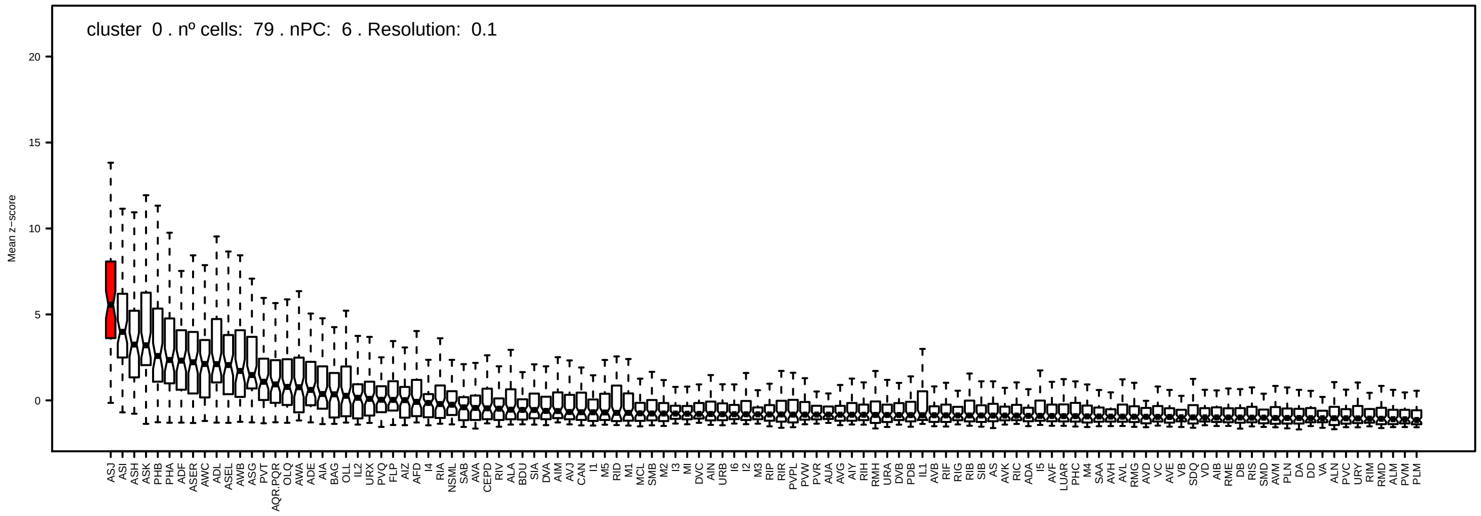
cluster 0 . nº cells: 79 . nPC: 6 . Resolution: 0.1
Mean z−score
0
5
10
15
20
ASJ ASI ASH ASK PHB PHA ADF ASER AWC ADL ASEL AWB ASG PVT AQR.PQR OLQ AWA ADE AIA BAG OLL IL2 URX PVQ FLP AIZ AFD I4 RIA NSML SAB AVA CEPD RIV ALA BDU SIA DVA AIM AVJ CAN I1 M5 RID M1 MCL SMB M2 I3 MI DVC AIN URB I6 I2 M3 RIP RIR PVPL PVW PVR AUA AVG AIY RIH RMH URA DVB PDB IL1 AVB RIF RIG RIB SIB AS AVK RIC ADA I5 AVF LUAR PHC M4 SAA AVH AVL RMG AVD VC AVE VB SDQ VD AIB RME DB RIS SMD AVM PLN DA DD VA ALN PVC URY RIM RMD ALM PVM PLM

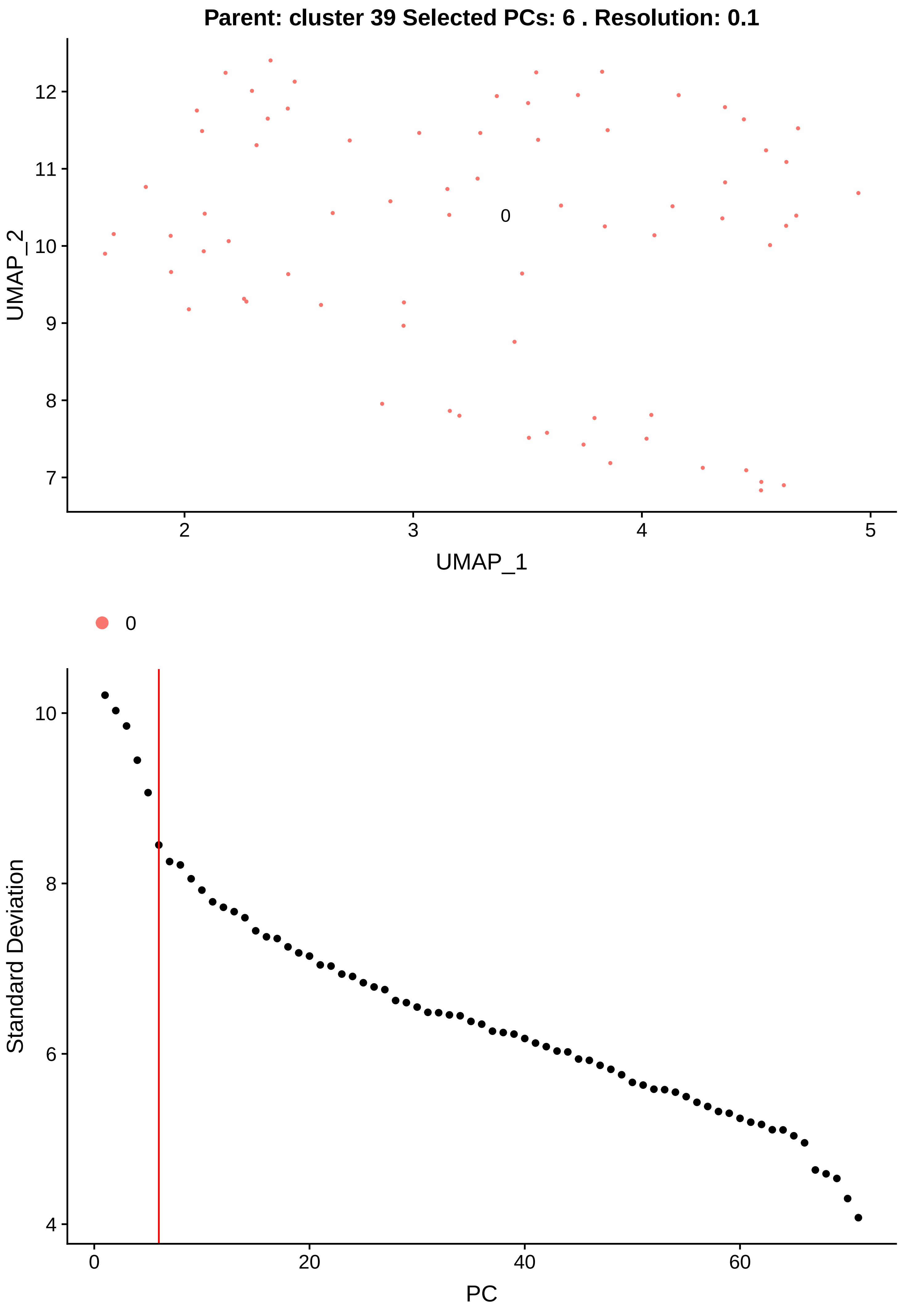

Parent: cluster 39 Selected PCs: 6 . Resolution: 0.1
UMAP_2
UMAP_1
0
Standard Deviation
PC

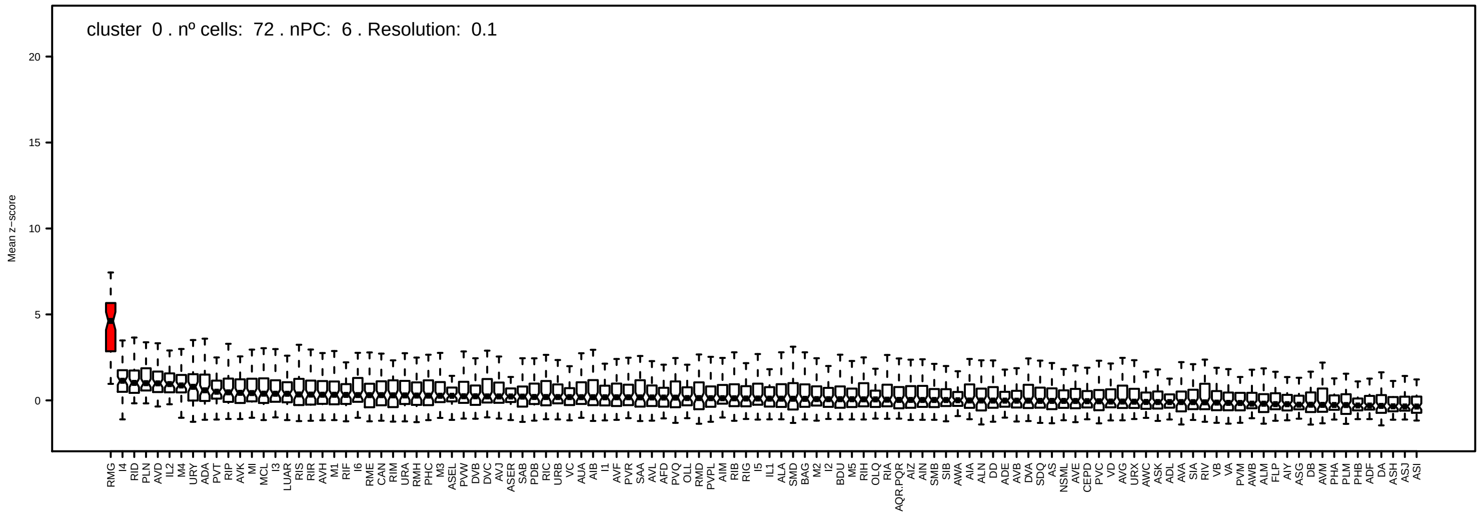
cluster 0 . nº cells: 72 . nPC: 6 . Resolution: 0.1
Mean z−score
RMG
I4
RID
PLN
AVD
IL2
M4
URY
ADA
PVT
RIP
AVK
MI
MCL
I3
LUAR
RIS
RIR
AVH
M1
RIF
I6
RME
CAN
RIM
URA
RMH
PHC
M3
ASEL
PVW
DVB
DVC
AVJ
ASER
SAB
PDB
RIC
URB
VC
AUA
AIB
I1
AVF
PVR
SAA
AVL
AFD
PVQ
OLL
RMD
PVPL
AIM
RIB
RIG
I5
IL1
ALA
SMD
BAG
M2
I2
BDU
M5
RIH
OLQ
RIA
AQR.PQR
AIZ
AIN
SMB
SIB
AWA
AIA
ALN
DD
ADE
AVB
DVA
SDQ
AS
NSML
AVE
CEPD
PVC
VD
AVG
URX
AWC
ASK
ADL
AVA
SIA
RIV
VB
VA
PVM
AWB
ALM
FLP
AIY
ASG
DB
AVM
PHA
PLM
PHB
ADF
DA
ASH
ASJ
ASI

Parent: cluster 40 Selected PCs: 6 . Resolution: 0.1

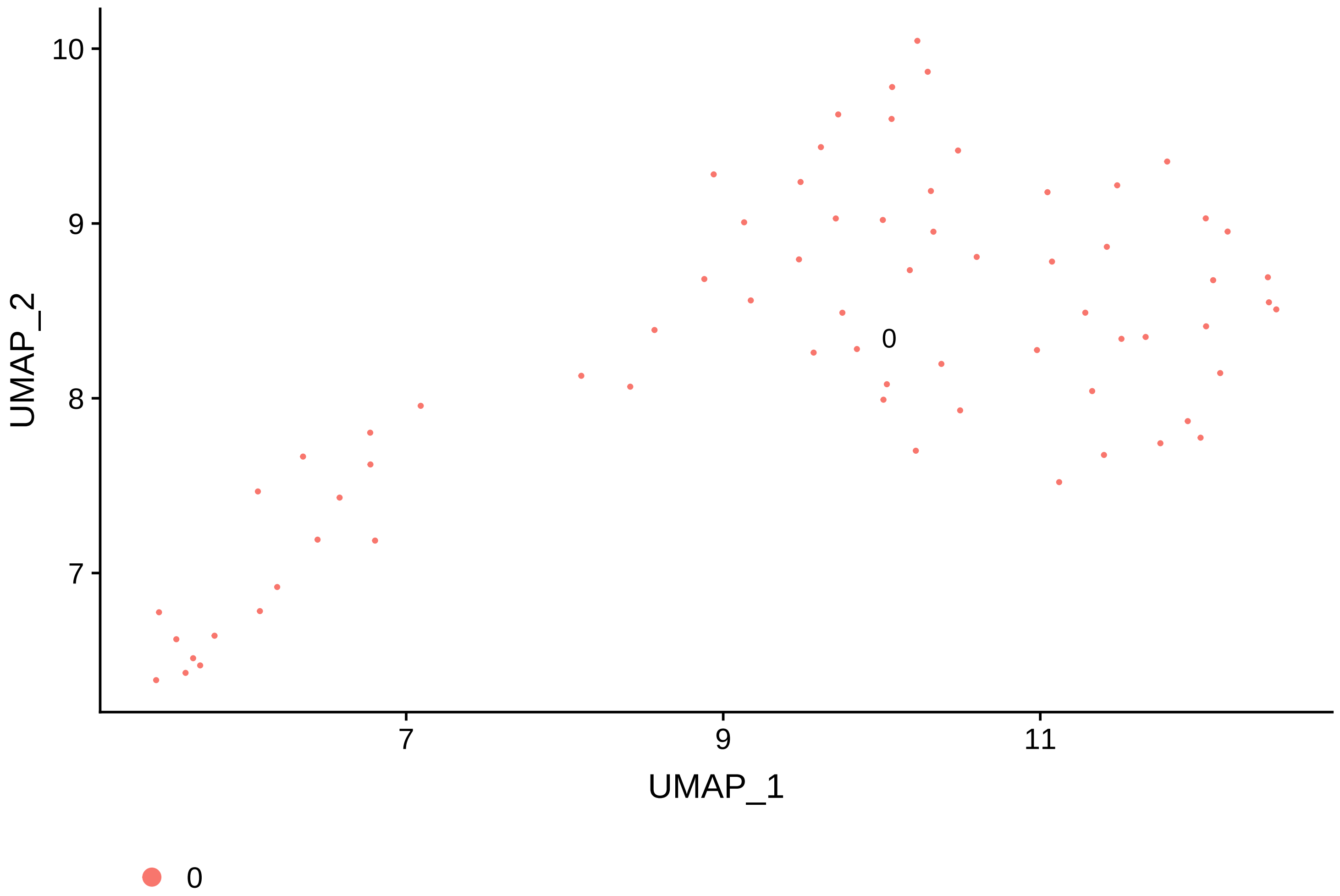


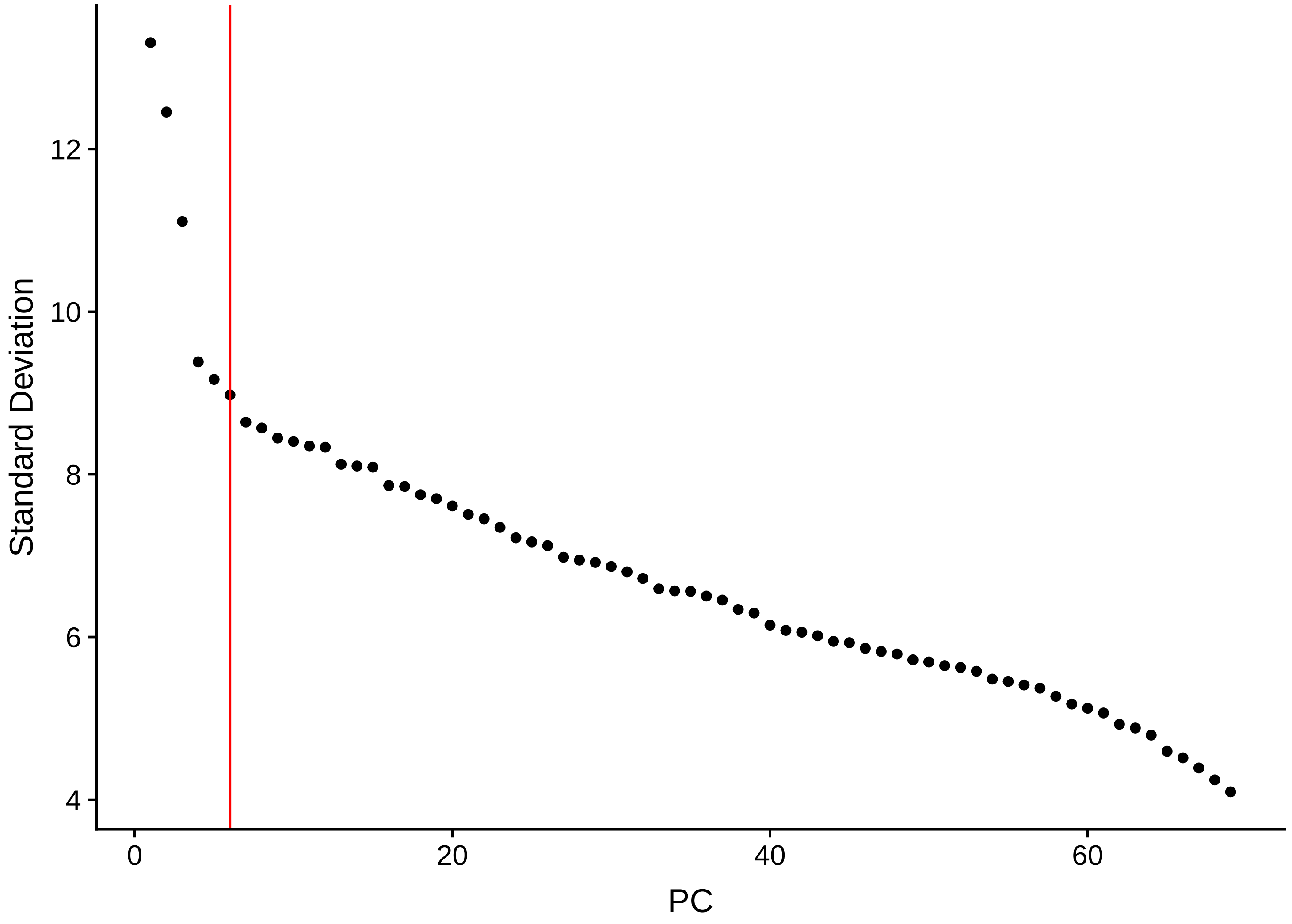

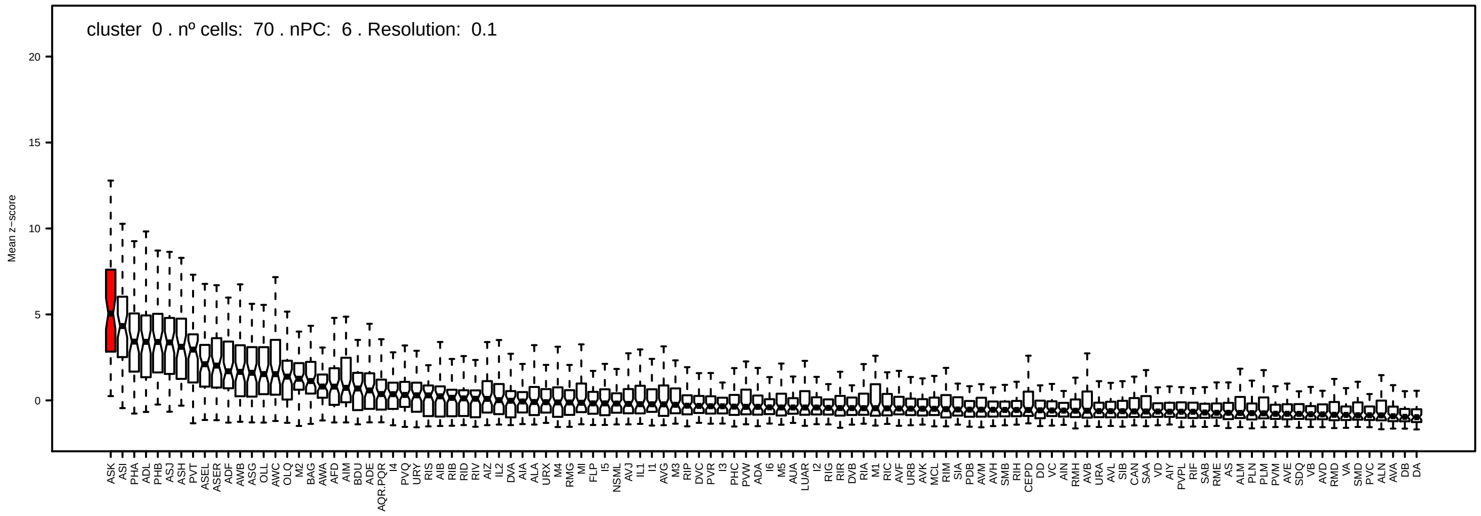
cluster 0 . nº cells: 70 . nPC: 6 . Resolution: 0.1
Mean z−score
20
15
10
5
0
ASK ASI PHA ADL PHB ASJ ASH PVT ASEL ASER ADF AWB ASG OLL AWC OLQ M2 BAG AWA AFD AIM BDU ADE AQR.PQR I4 PVQ URY RIS AIB RIB RID RIV AIZ IL2 DVA AIA ALA URX M4 RMG MI FLP I5 NSML AVJ IL1 I1 AVG M3 RIP DVC PVR I3 PHC PVW ADA I6 M5 AUA LUAR I2 RIG RIR DVB RIA M1 RIC AVF URB AVK MCL RIM SIA PDB AVM AVH SMB RIH CEPD DD VC AIN RMH AVB URA AVL SIB CAN SAA VD AIY PVPL RIF SAB RME AS ALM PLN PLM PVM AVE SDQ VB AVD RMD VA SMD PVC ALN AVA DB DA

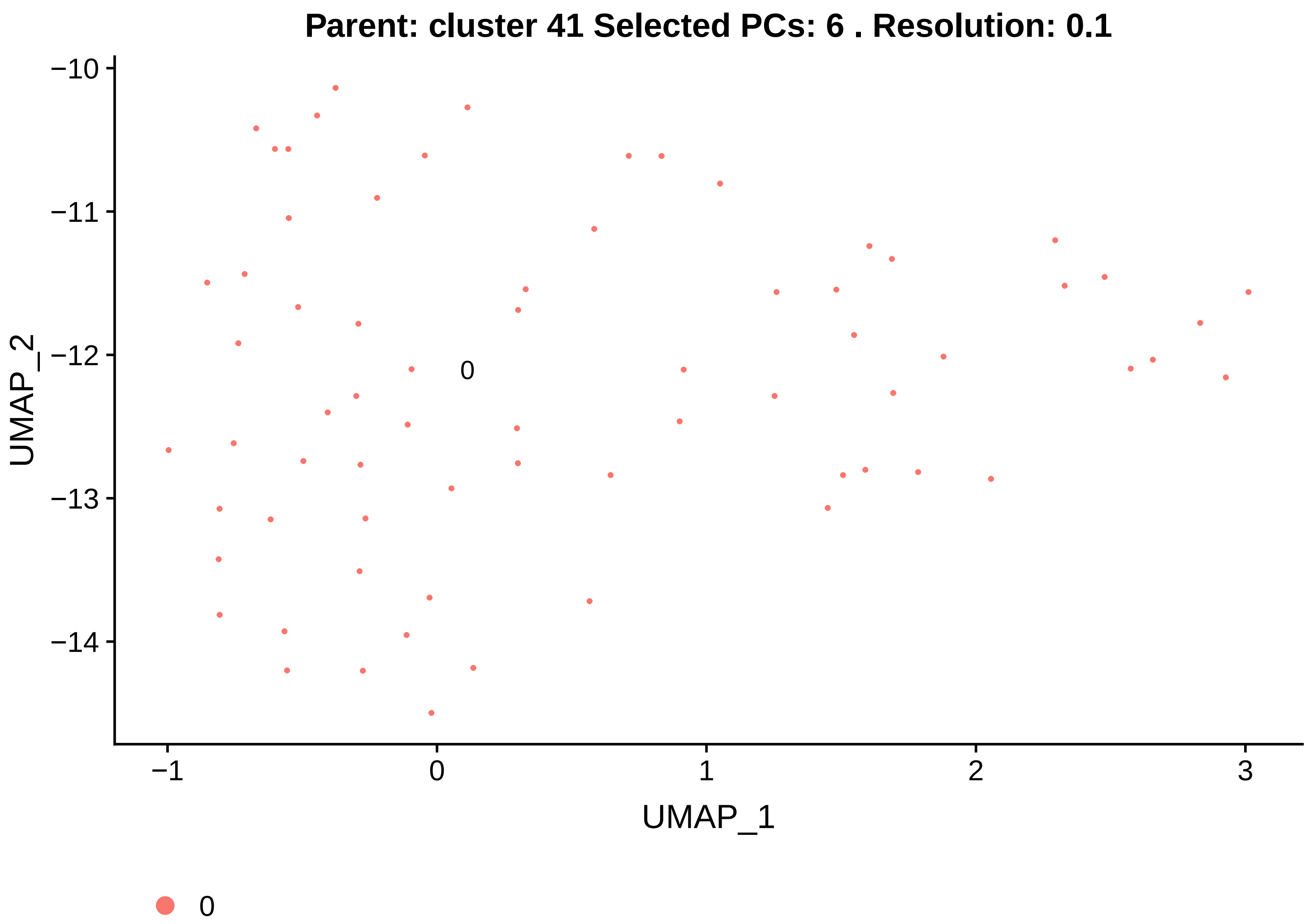

Parent: cluster 41 Selected PCs: 6 . Resolution: 0.1
UMAP_2
UMAP_1
−10
−11
−12
−13
−14
−1
0
1
2
3
0
0


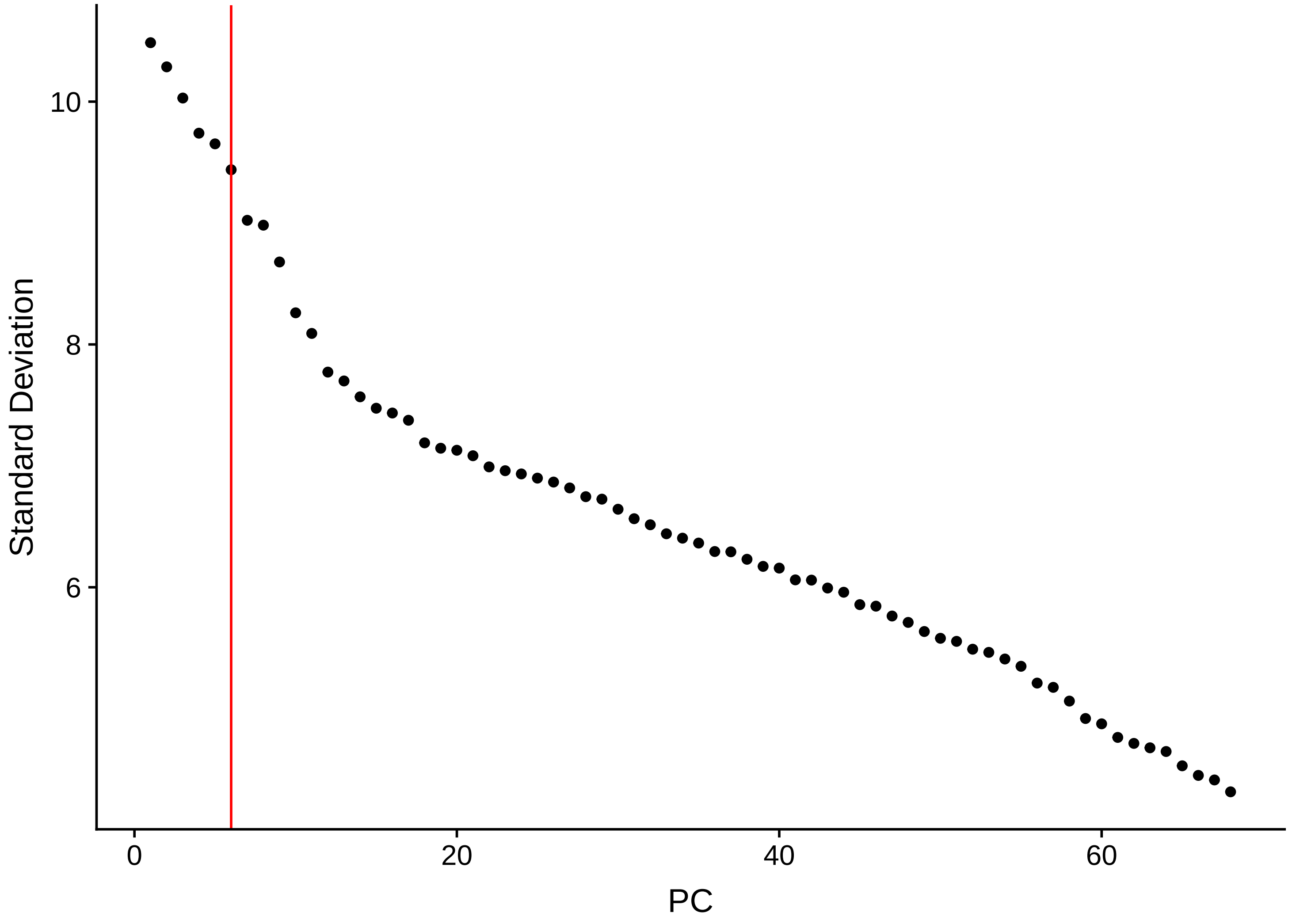

Standard Deviation
PC
10
8
6
0
20
40
60

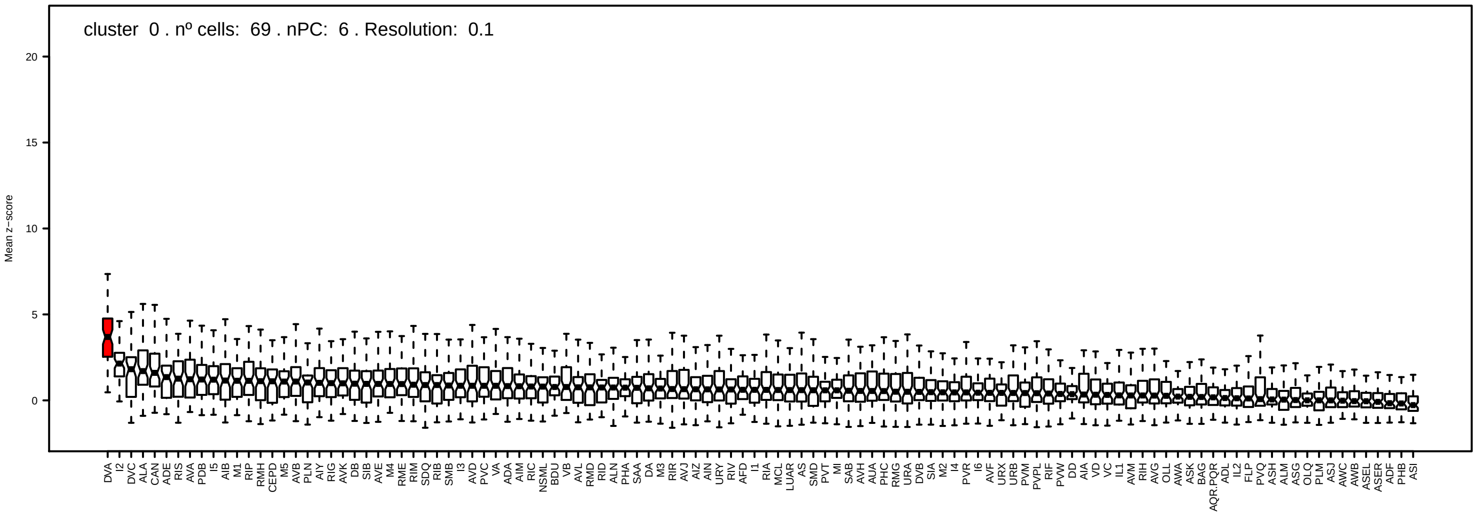
cluster 0 . nº cells: 69 . nPC: 6 . Resolution: 0.1
Mean z−score
20
15
10
5
0
DVA I2 DVC ALA CAN ADE RIS AVA PDB I5 AIB M1 RIP RMH CEPD M5 AVB PLN AIY RIG AVK DB SIB AVE M4 RME RIM SDQ RIB SMB I3 AVD PVC VA ADA AIM RIC NSML BDU VB AVL RMD RID ALN PHA SAA DA M3 RIR AVJ AIZ AIN URY RIV AFD I1 RIA MCL LUAR AS SMD PVT MI SAB AVH AUA PHC RMG URA DVB SIA M2 I4 PVR I6 AVF URX URB PVM PVPL RIF PVW DD AIA VD VC IL1 AVM RIH AVG OLL AWA ASK BAG AQR.PQR ADL IL2 FLP PVQ ASH ALM ASG OLQ PLM ASJ AWC AWB ASEL ASER ADF PHB ASI

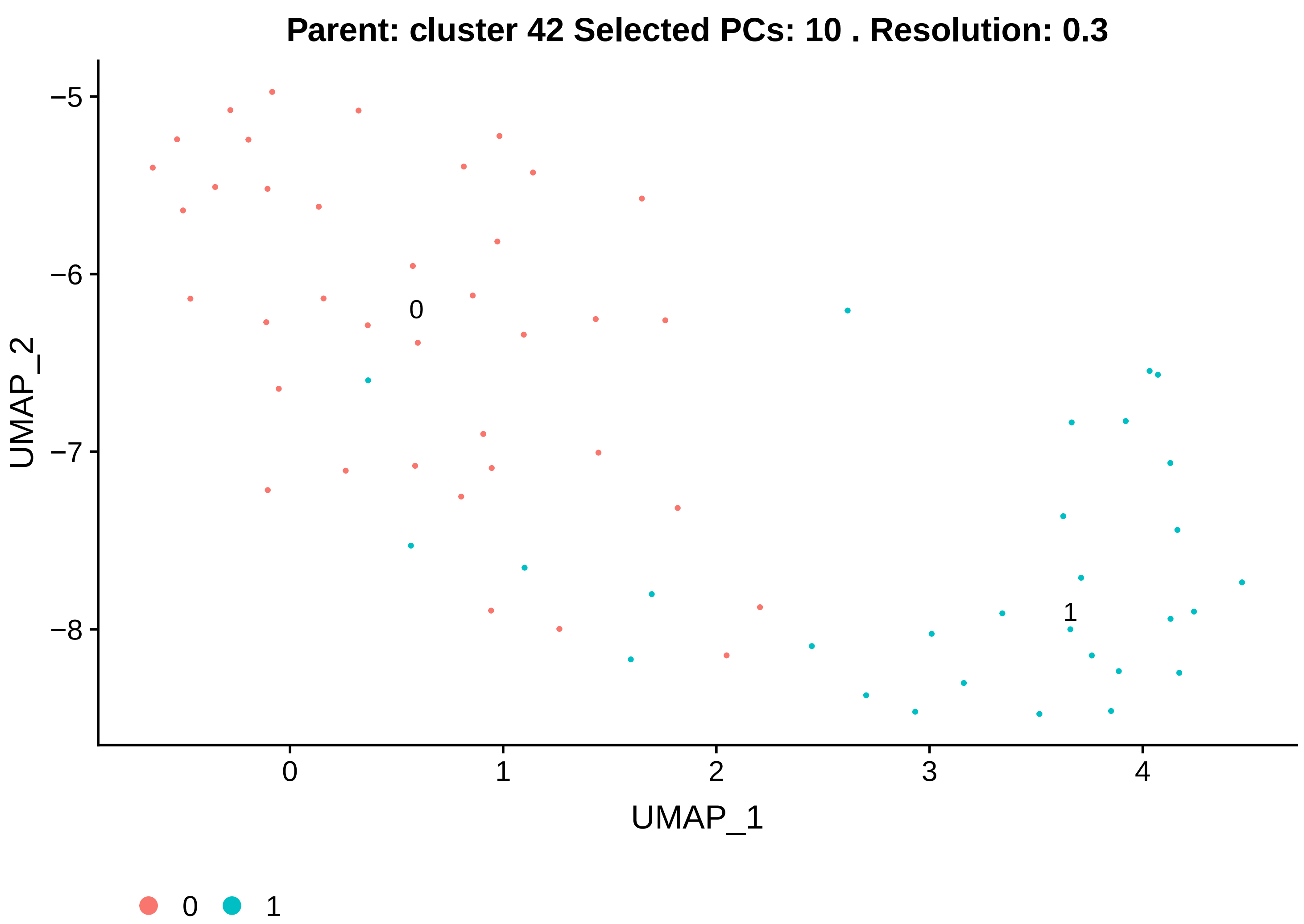
Parent: cluster 42 Selected PCs: 10 . Resolution: 0.3
−5
−6
−7
−8
UMAP_2
0
1
0
1
2
3
4
UMAP_1
0
1

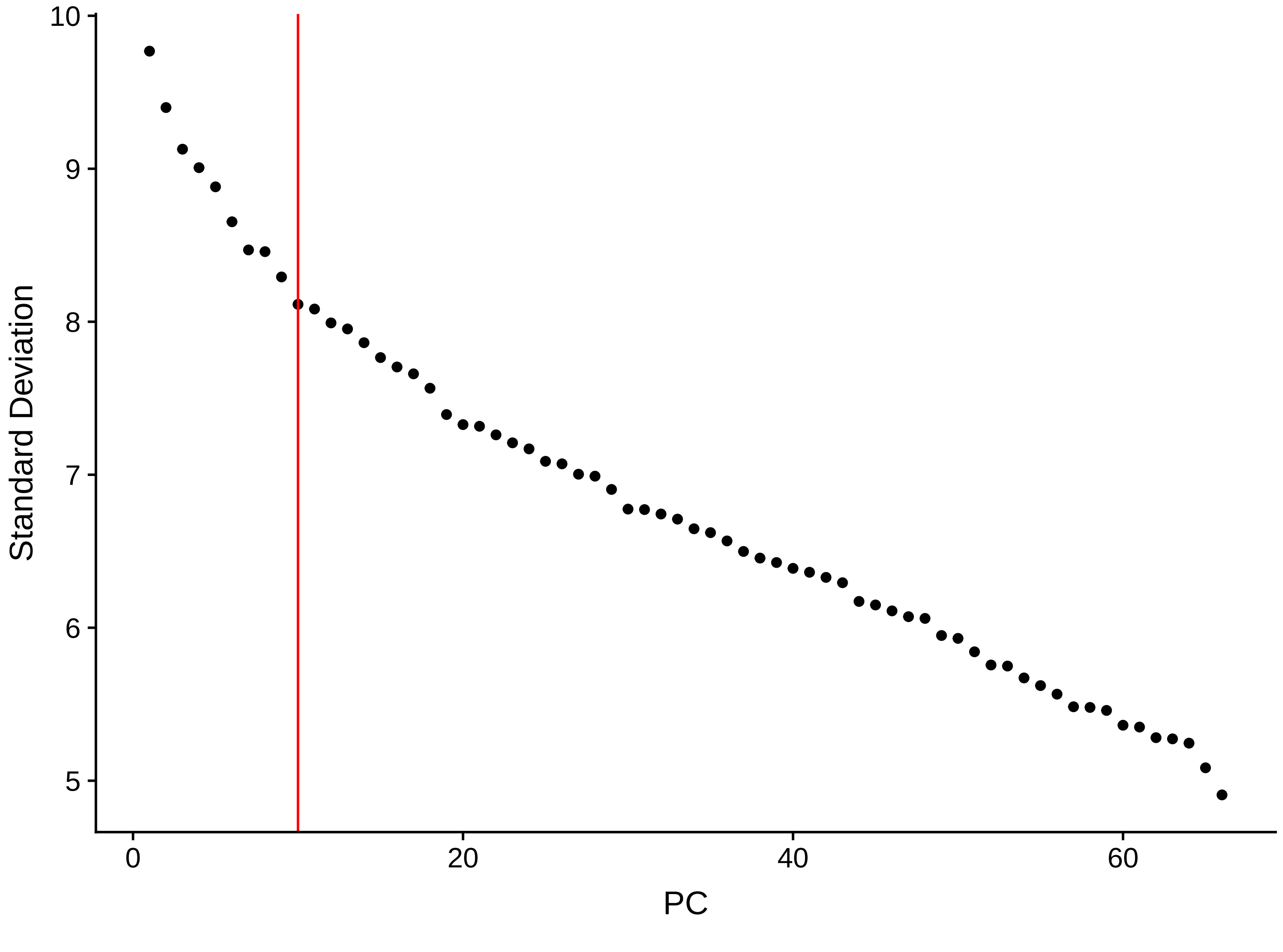
10
9
8
7
6
5
Standard Deviation
0
20
40
60
PC

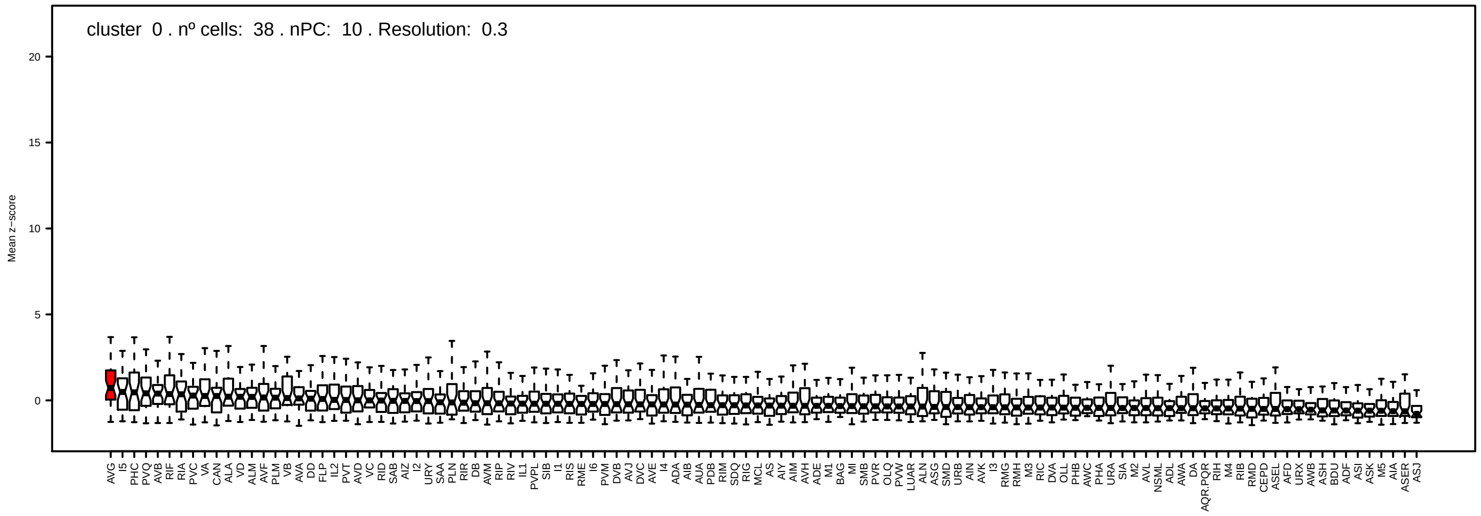
cluster 0 . nº cells: 38 . nPC: 10 . Resolution: 0.3
Mean z−score

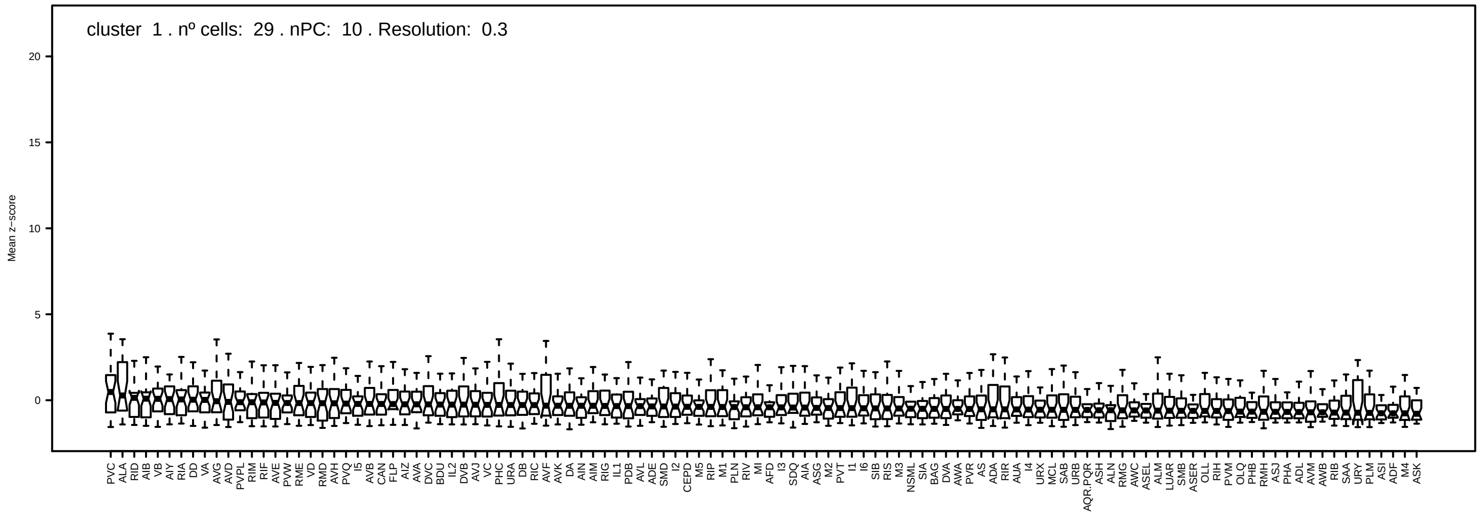
cluster 1 . nº cells: 29 . nPC: 10 . Resolution: 0.3
Mean z−score

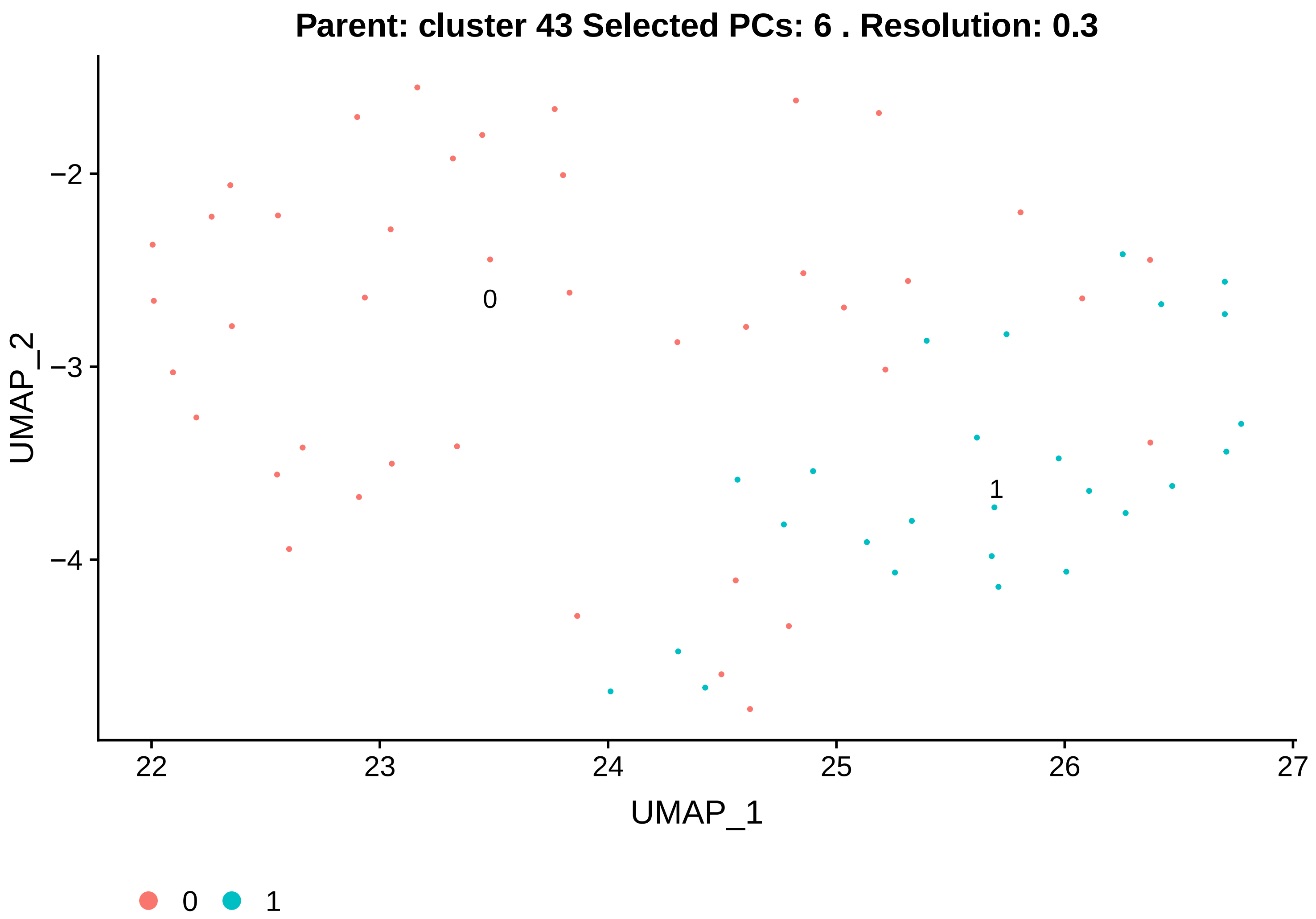

Parent: cluster 43 Selected PCs: 6 . Resolution: 0.3
UMAP_2
UMAP_1
0
1
−2
−3
−4
22
23
24
25
26
27


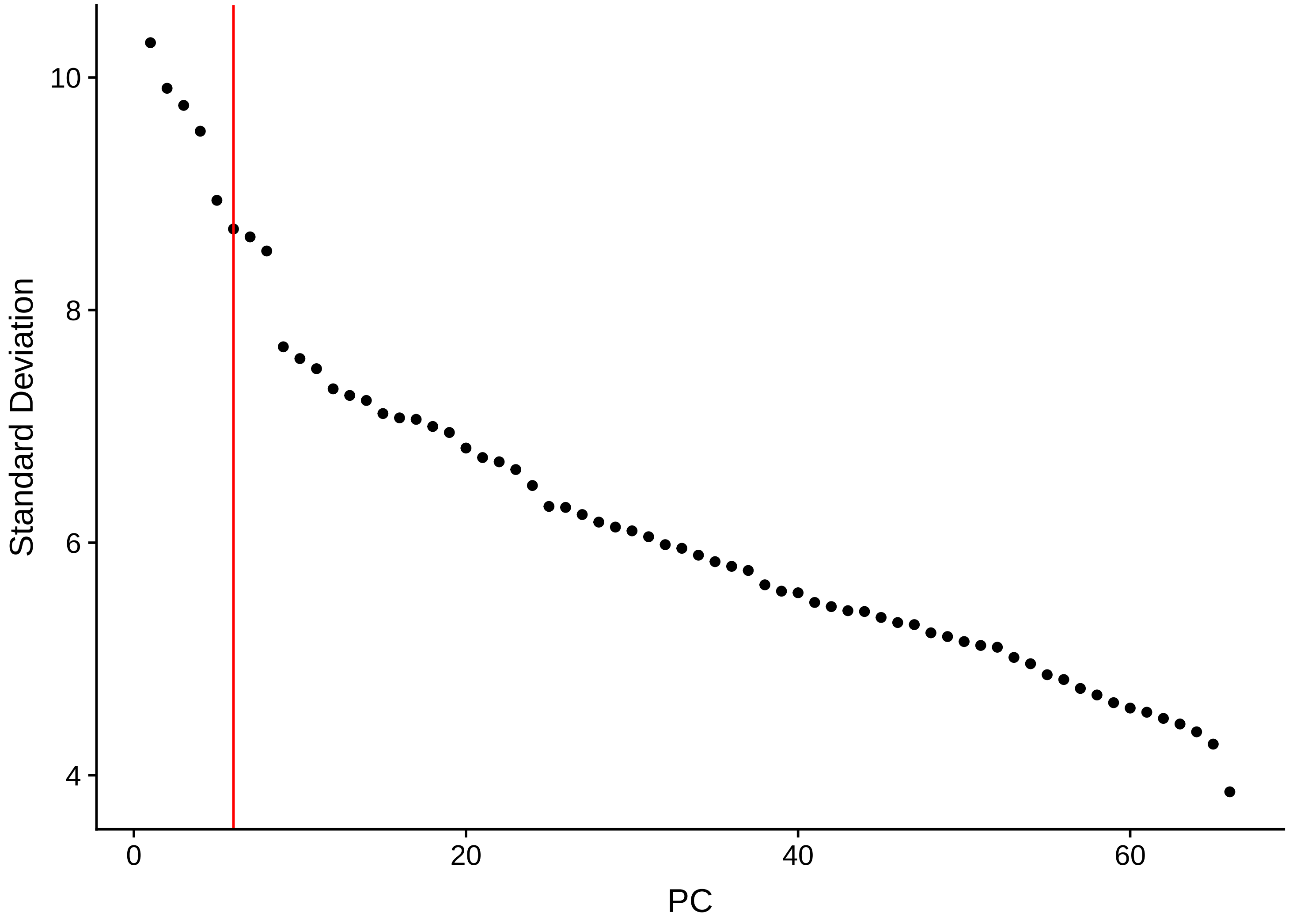

Standard Deviation
PC
10
8
6
4
0
20
40
60

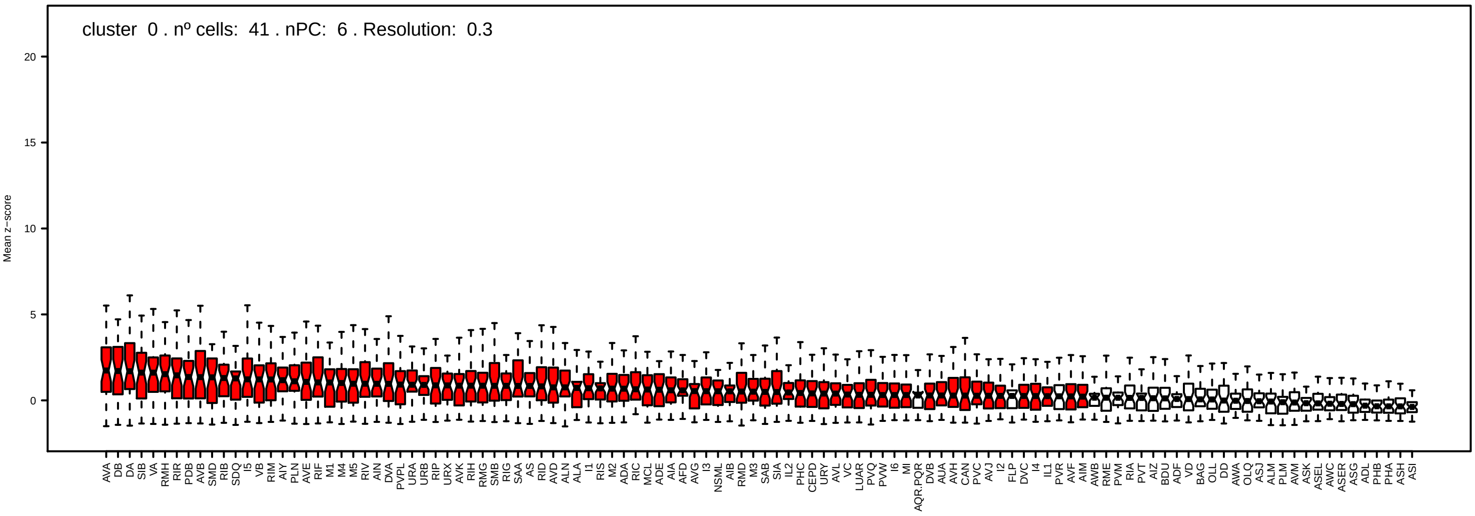
cluster 0 . nº cells: 41 . nPC: 6 . Resolution: 0.3
Mean z−score
20
15
10
5
0
AVA DB DA SIB VA RMH RIR PDB AVB SMD RIB SDQ I5 VB RIM AIY PLN AVE RIF M1 M4 M5 RIV AIN DVA PVPL URA URB RIP URX AVK RIH RMG SMB RIG SAA AS RID AVD ALN ALA I1 RIS M2 ADA RIC MCL ADE AIA AFD AVG I3 NSML AIB RMD M3 SAB SIA IL2 PHC CEPD URY AVL VC LUAR PVQ PVW I6 MI AQR.PQR DVB AUA AVH CAN PVC AVJ I2 FLP DVC I4 IL1 PVR AVF AIM AWB RME PVM RIA PVT AIZ BDU ADF VD BAG OLL DD AWA OLQ ASJ ALM PLM AVM ASK ASEL AWC ASER ASG ADL PHB PHA ASH ASI

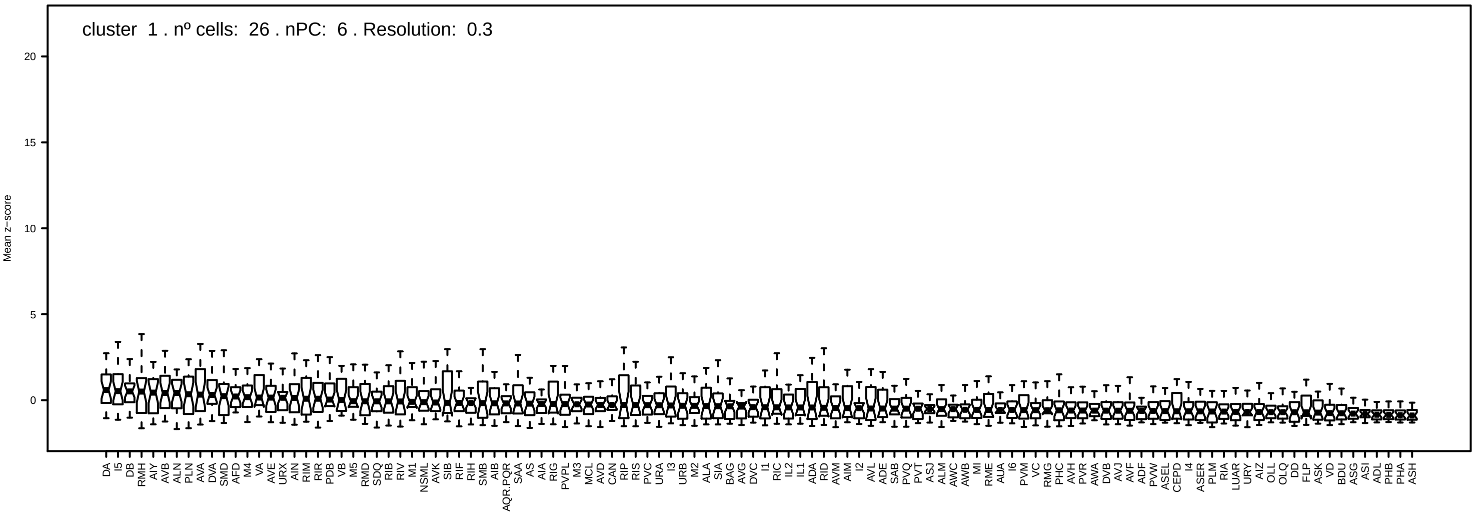
cluster 1 . nº cells: 26 . nPC: 6 . Resolution: 0.3
Mean z−score
20
15
10
5
0
DA I5 DB RMH AIY AVB ALN PLN AVA DVA SMD AFD M4 VA AVE URX AIN RIM RIR PDB VB M5 RMD SDQ RIB RIV M1 NSML AVK SIB RIF RIH SMB AIB AQR.PQR SAA AS AIA RIG PVPL M3 MCL AVD CAN RIP RIS PVC URA I3 URB M2 ALA SIA BAG AVG DVC I1 RIC IL2 IL1 ADA RID AVM AIM I2 AVL ADE SAB PVQ PVT ASJ ALM AWC AWB MI RME AUA I6 PVM VC RMG PHC AVH PVR AWA DVB AVJ AVF PVW ASEL CEPD I4 ASER PLM RIA LUAR URY AIZ OLL OLQ DD FLP ASK VD BDU ASG ASI ADL PHB PHA ASH

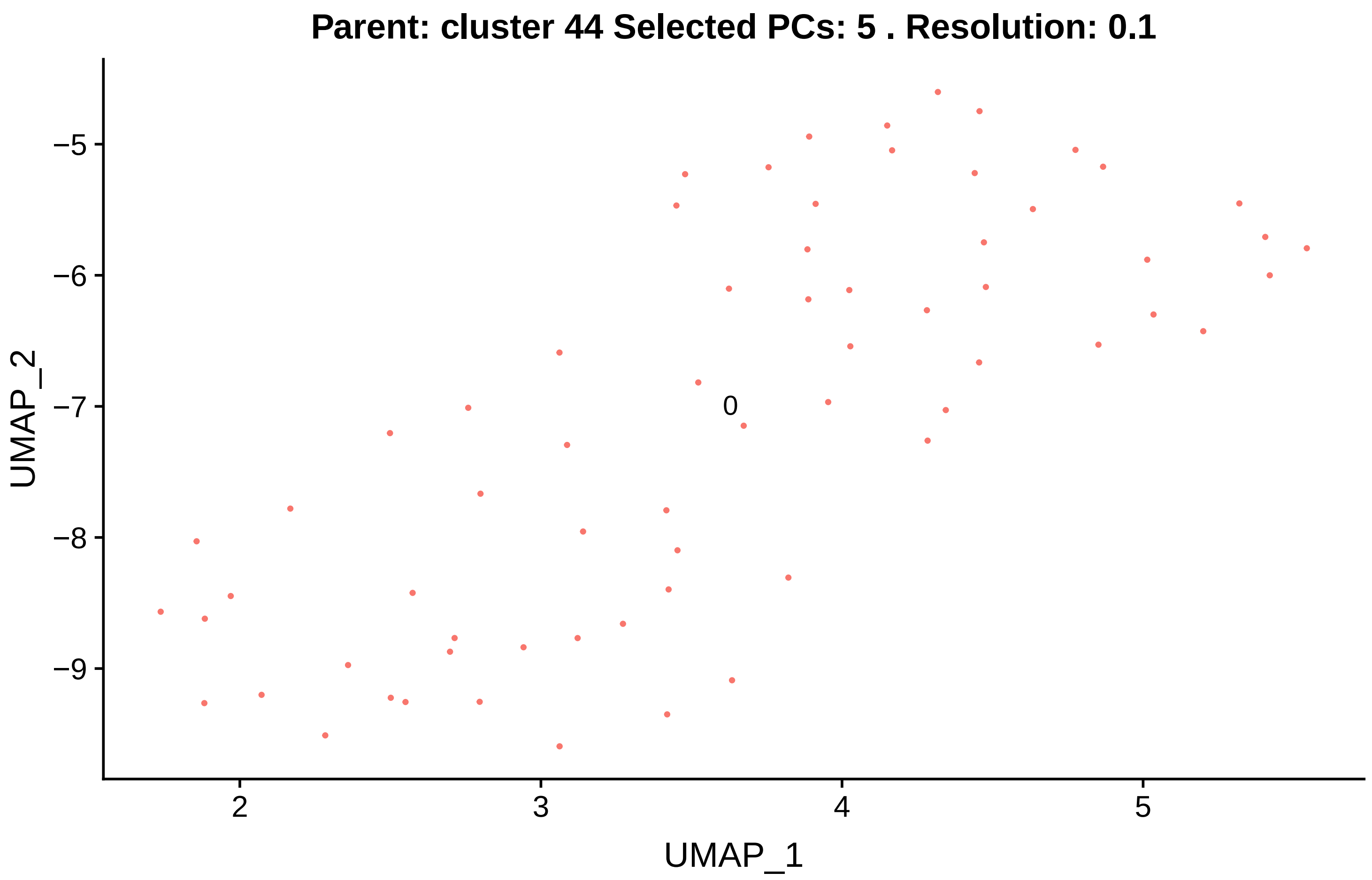

Parent: cluster 44 Selected PCs: 5 . Resolution: 0.1
UMAP_2
UMAP_1
−5
−6
−7
−8
−9
2
3
4
5
0


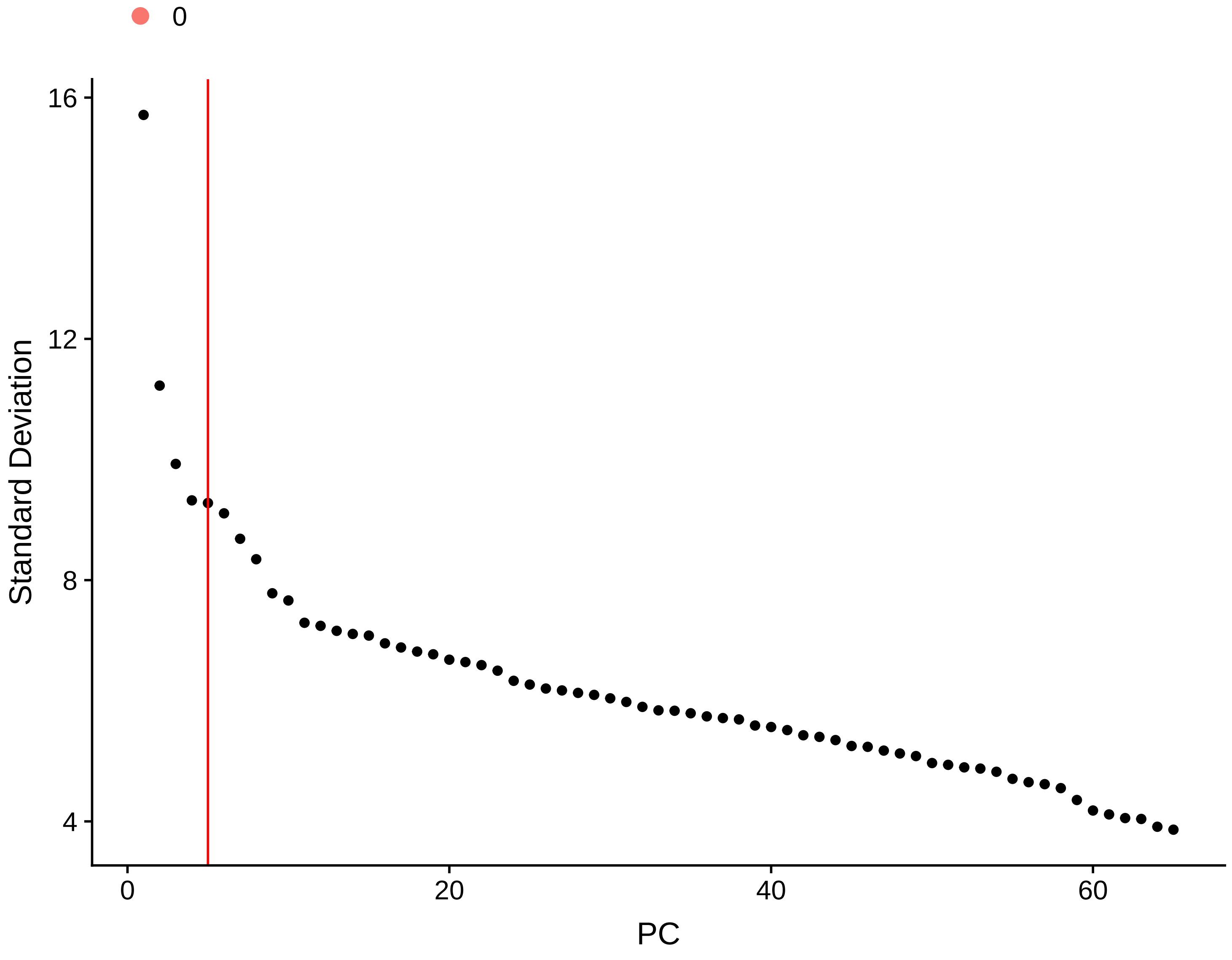

0
Standard Deviation
PC
16
12
8
4
0
20
40
60

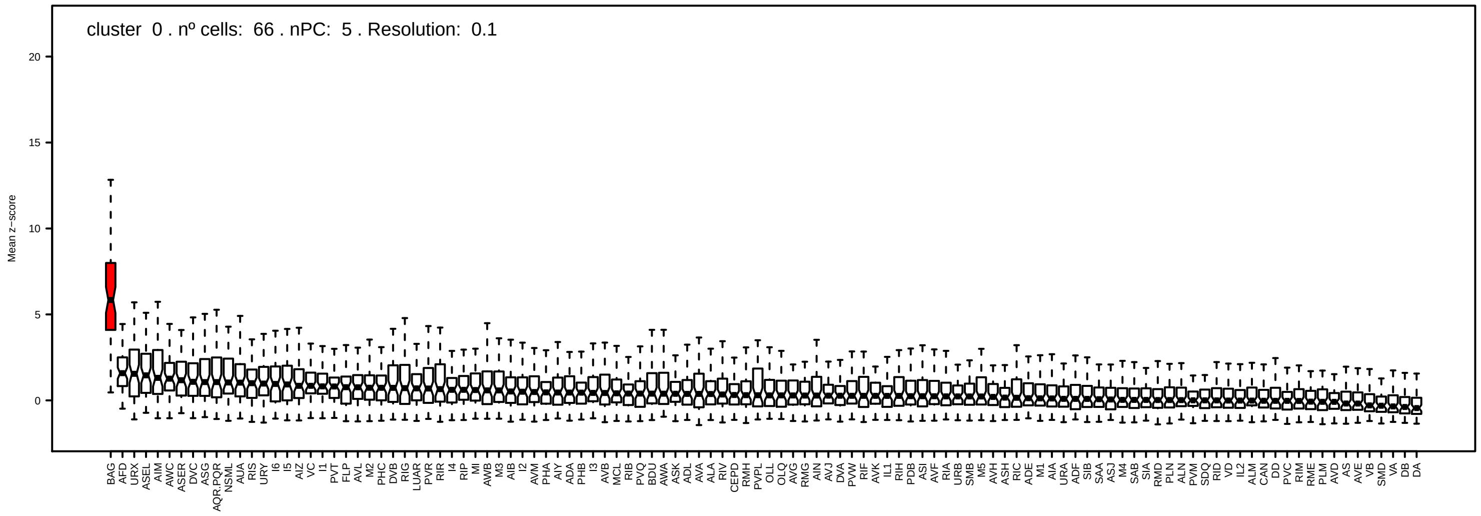
cluster 0 . nº cells: 66 . nPC: 5 . Resolution: 0.1
Mean z−score
20
15
10
5
0
BAG
AFD
URX
ASEL
AIM
AWC
ASER
DVC
ASG
AQR.PQR
NSML
AUA
RIS
URY
I6
I5
AIZ
VC
I1
PVT
FLP
AVL
M2
PHC
DVB
RIG
LUAR
PVR
RIR
I4
RIP
MI
AWB
M3
AIB
I2
AVM
PHA
AIY
ADA
PHB
I3
AVB
MCL
RIB
PVQ
BDU
AWA
ASK
ADL
AVA
ALA
RIV
CEPD
RMH
PVPL
OLL
OLQ
AVG
RMG
AIN
AVJ
DVA
PVW
RIF
AVK
IL1
RIH
PDB
ASI
AVF
RIA
URB
SMB
M5
AVH
ASH
RIC
ADE
M1
AIA
URA
ADF
SIB
SAA
ASJ
M4
SAB
SIA
RMD
PLN
ALN
PVM
SDQ
RID
VD
IL2
ALM
CAN
DD
PVC
RIM
RME
PLM
AVD
AS
AVE
VB
SMD
VA
DB
DA

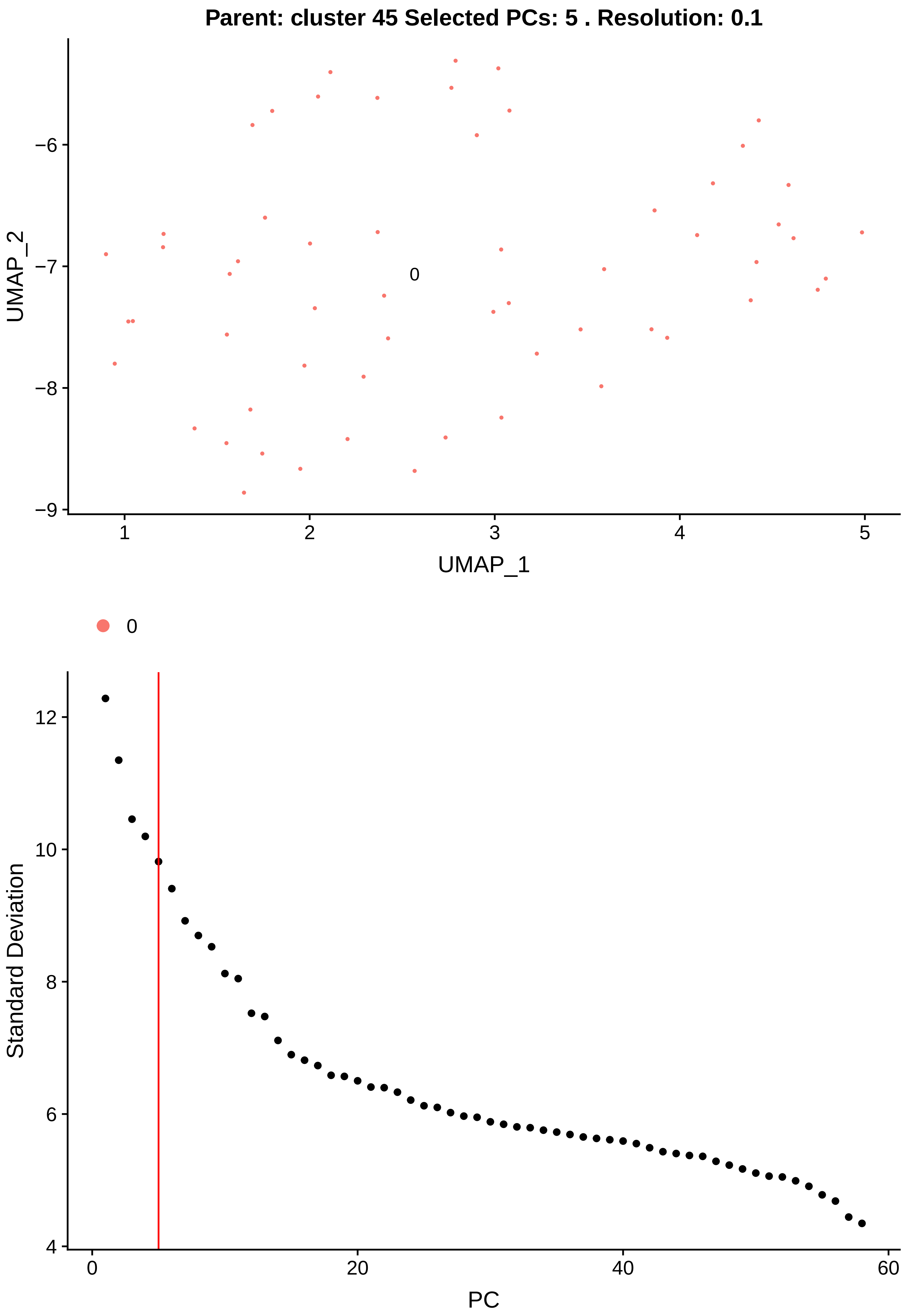

Parent: cluster 45 Selected PCs: 5 . Resolution: 0.1
UMAP_2
UMAP_1
0
Standard Deviation
PC

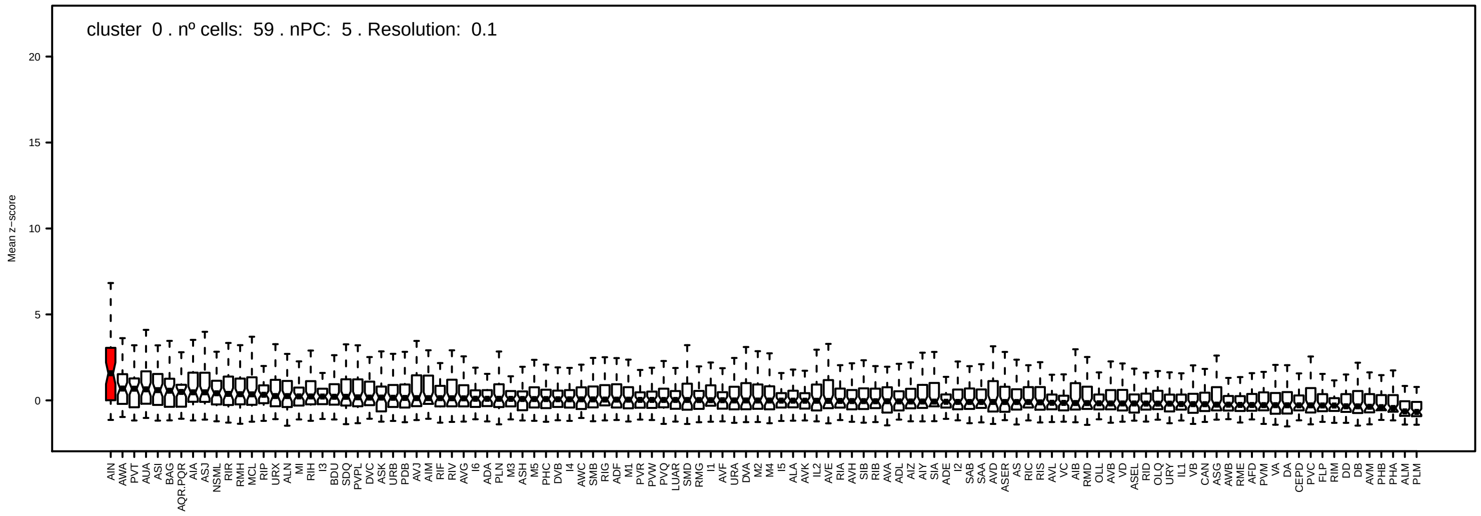

cluster 0 . nº cells: 59 . nPC: 5 . Resolution: 0.1
Mean z−score
20
15
10
5
0
AIN AWA PVT AUA ASI BAG AQR.PQR AIA ASJ NSML RIR RMH MCL RIP URX ALN MI RIH I3 BDU SDQ PVPL DVC ASK URB PDB AVJ AIM RIF RIV AVG I6 ADA PLN M3 ASH M5 PHC DVB I4 AWC SMB RIG ADF M1 PVR PVW PVQ LUAR SMD RMG I1 AVF URA DVA M2 M4 I5 ALA AVK IL2 AVE RIA AVH SIB RIB AVA ADL AIZ AIY SIA ADE I2 SAB SAA AVD ASER AS RIC RIS AVL VC AIB RMD OLL AVB VD ASEL RID OLQ URY IL1 VB CAN ASG AWB RME AFD PVM VA DA CEPD PVC FLP RIM DD DB AVM PHB PHA ALM PLM

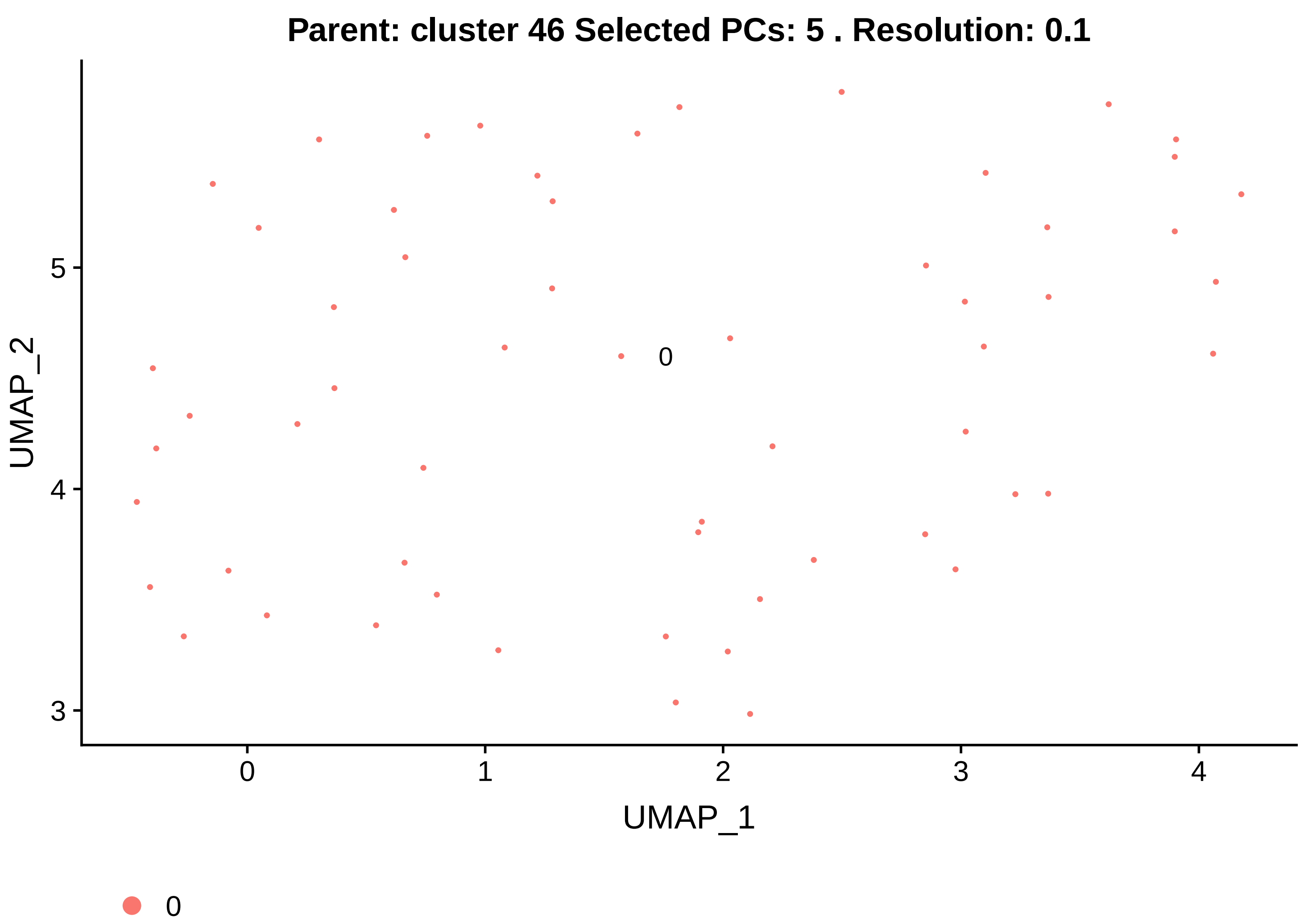

Parent: cluster 46 Selected PCs: 5 . Resolution: 0.1
UMAP_2
UMAP_1
0
3
4
5
0
1
2
3
4


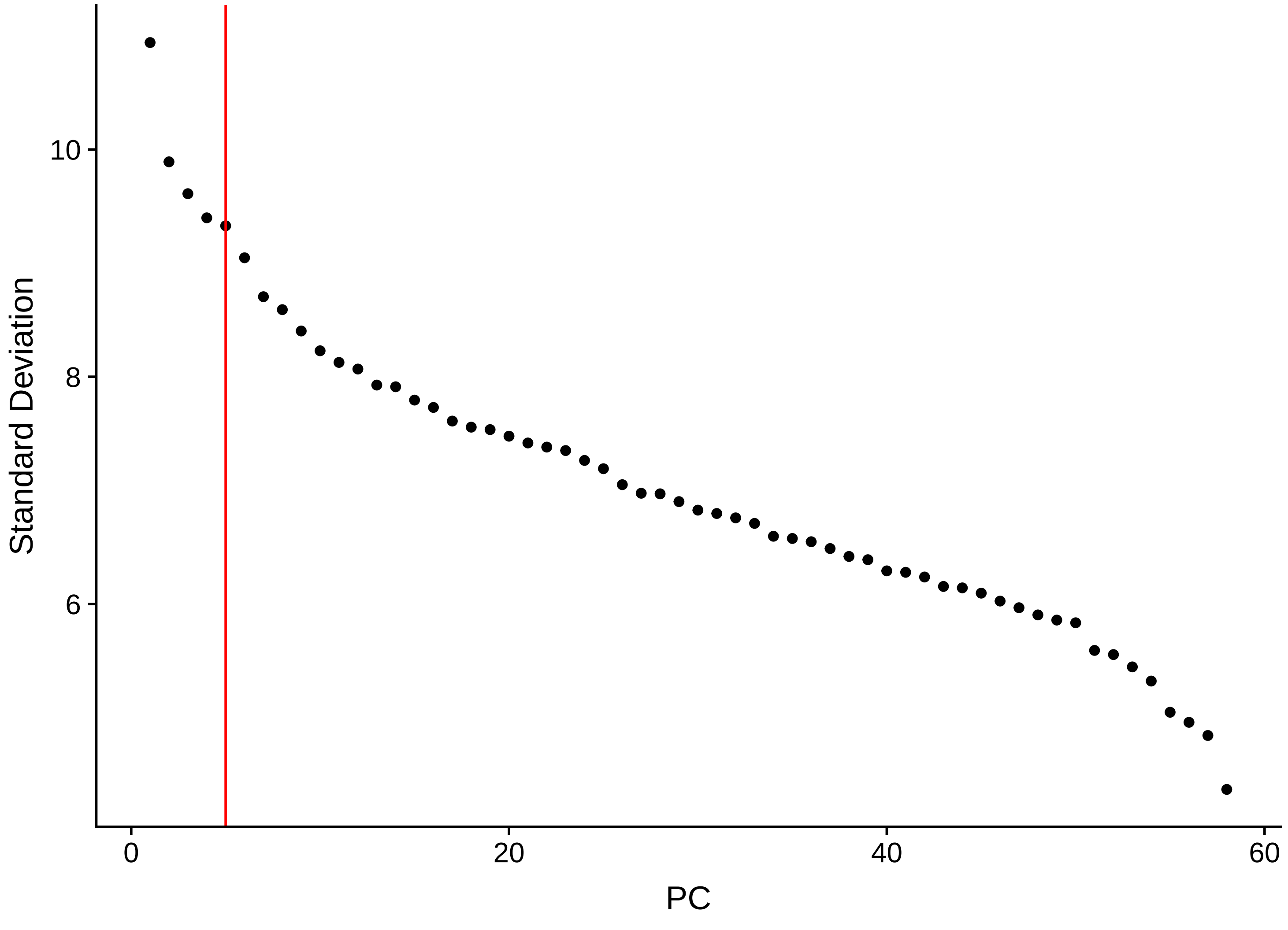

Standard Deviation
PC
6
8
10
0
20
40
60

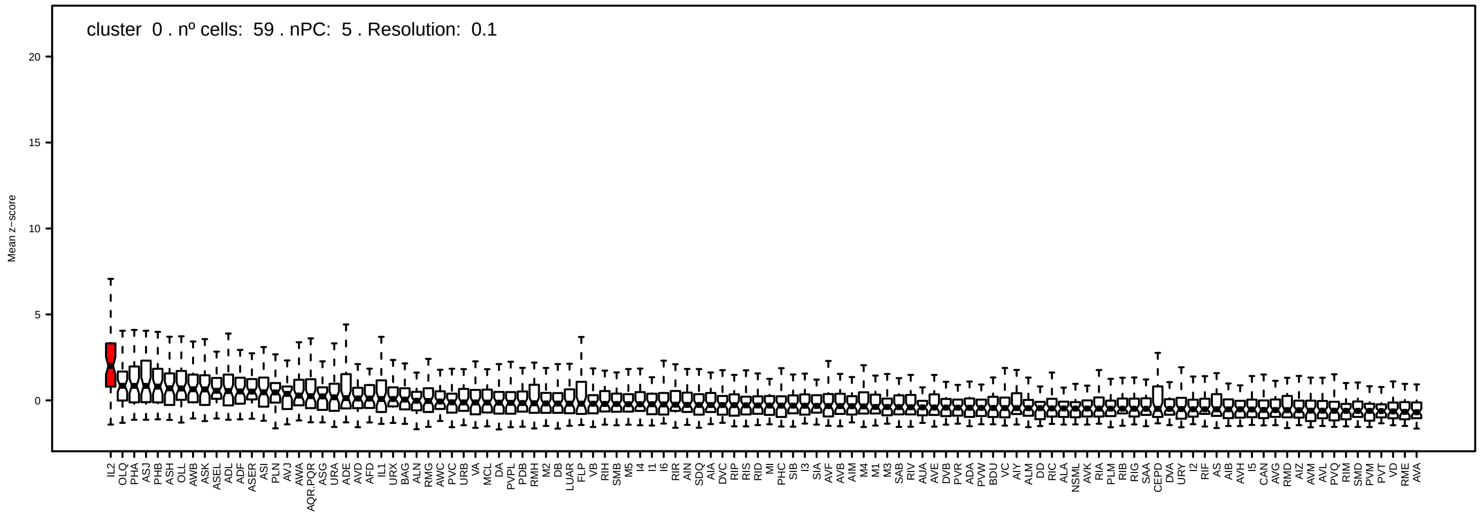
cluster 0 . nº cells: 59 . nPC: 5 . Resolution: 0.1
Mean z−score
0
5
10
15
20
IL2 OLQ PHA ASJ PHB ASH OLL AWB ASK ASEL ADL ADF ASER ASI PLN AVJ AWA AQR.PQR ASG URA ADE AVD AFD IL1 URX BAG ALN RMG AWC PVC URB VA MCL DA PVPL PDB RMH M2 DB LUAR FLP VB RIH SMB M5 I4 I1 I6 RIR AIN SDQ AIA DVC RIP RIS RID MI PHC SIB I3 SIA AVF AVB AIM M4 M1 M3 SAB RIV AUA AVE DVB PVR ADA PVW BDU VC AIY ALM DD RIC ALA NSML AVK RIA PLM RIB RIG SAA CEPD DVA URY I2 RIF AS AIB AVH I5 CAN AVG RMD AIZ AVM AVL PVQ RIM SMD PVM PVT VD RME AVA

Parent: cluster 48 Selected PCs: 4 . Resolution: 0.3

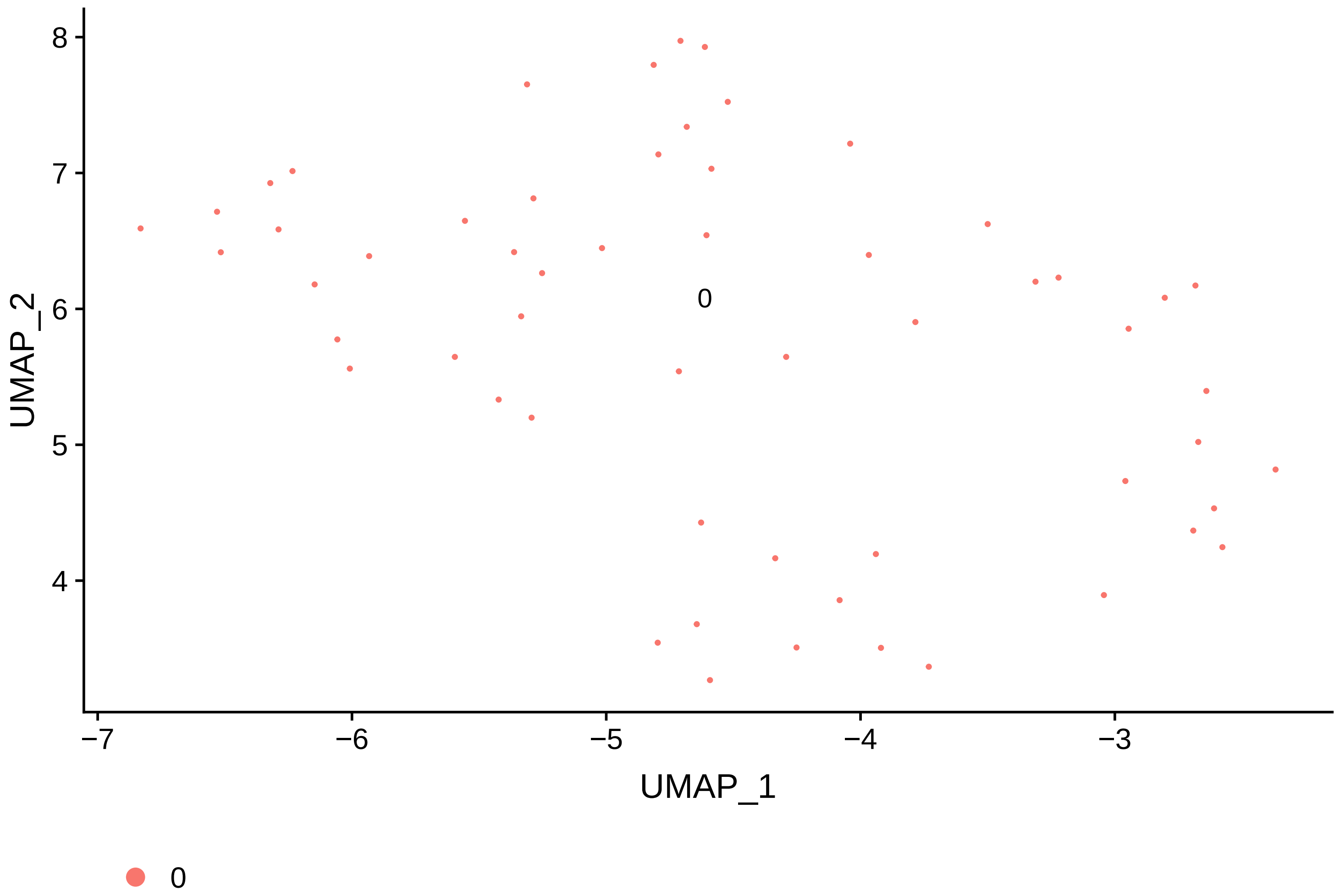


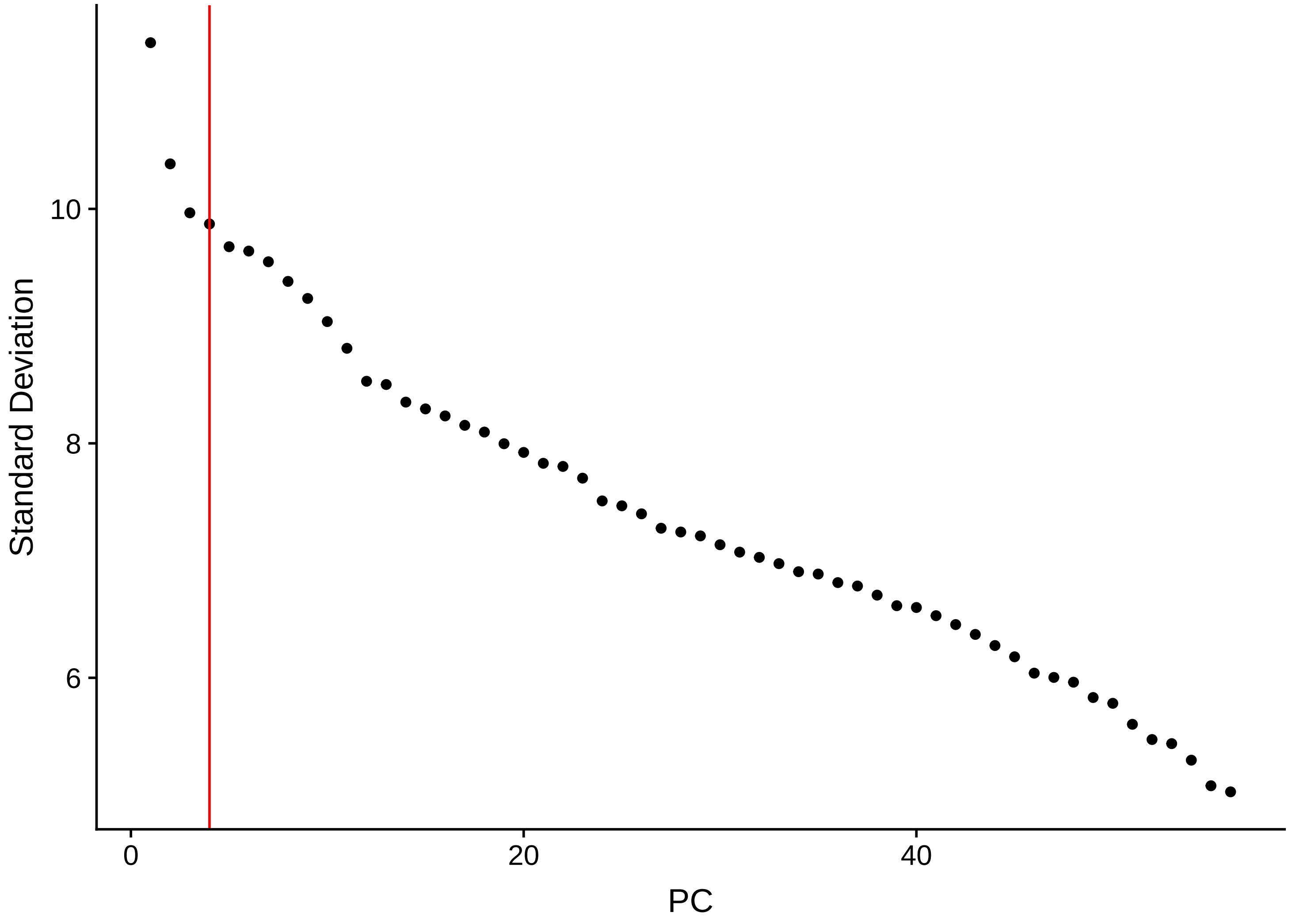

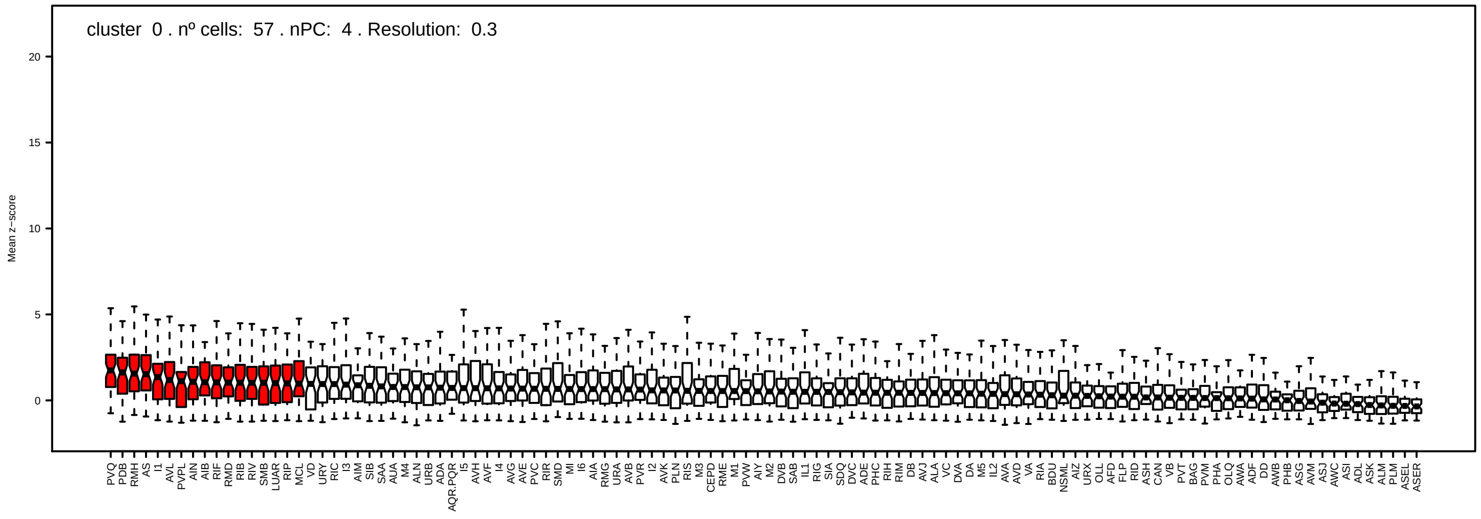
cluster 0 . nº cells: 57 . nPC: 4 . Resolution: 0.3
Mean z−score

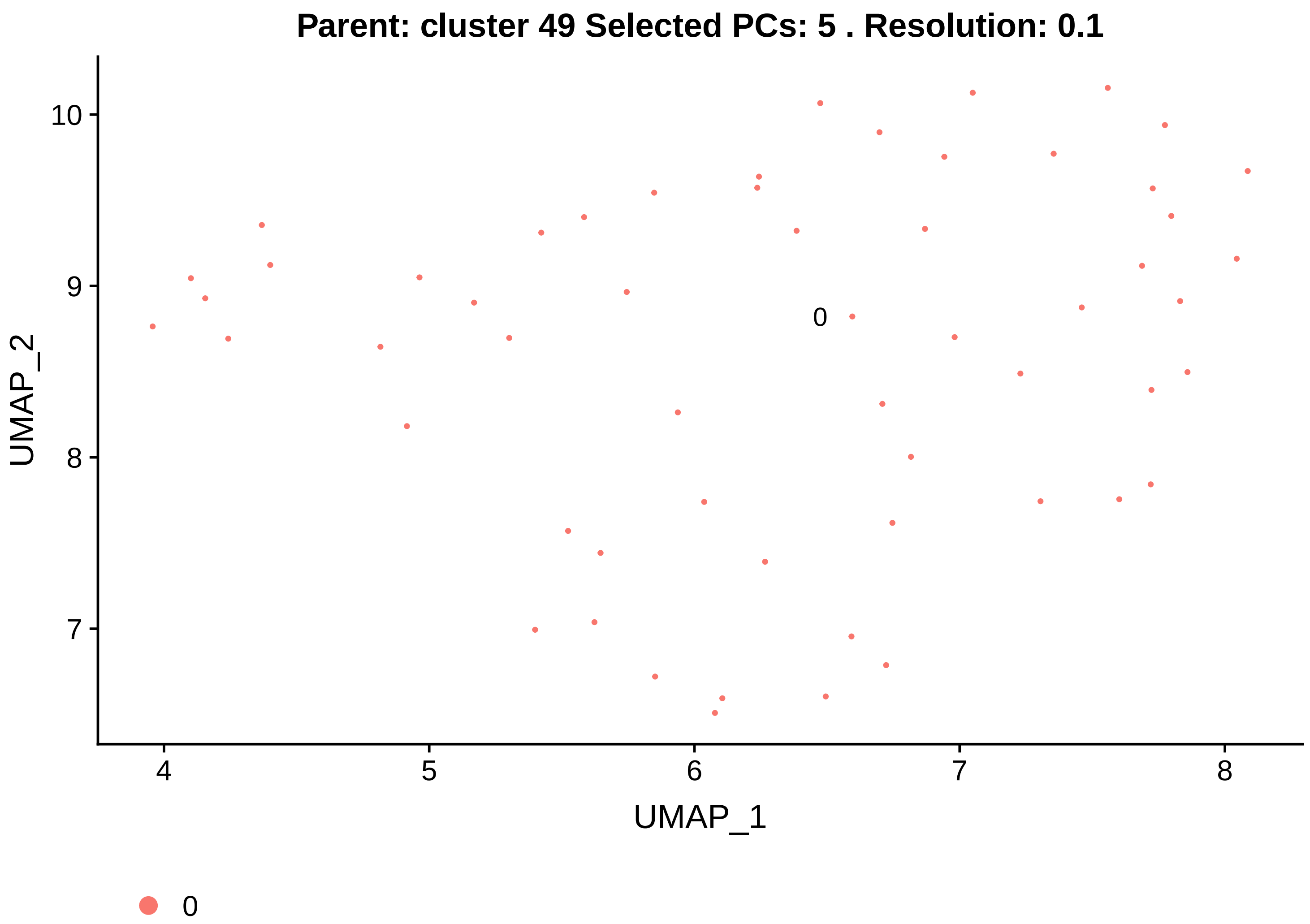
Parent: cluster 49 Selected PCs: 5 . Resolution: 0.1
UMAP_2
10
9
8
7
0
4
5
6
7
8
UMAP_1
0

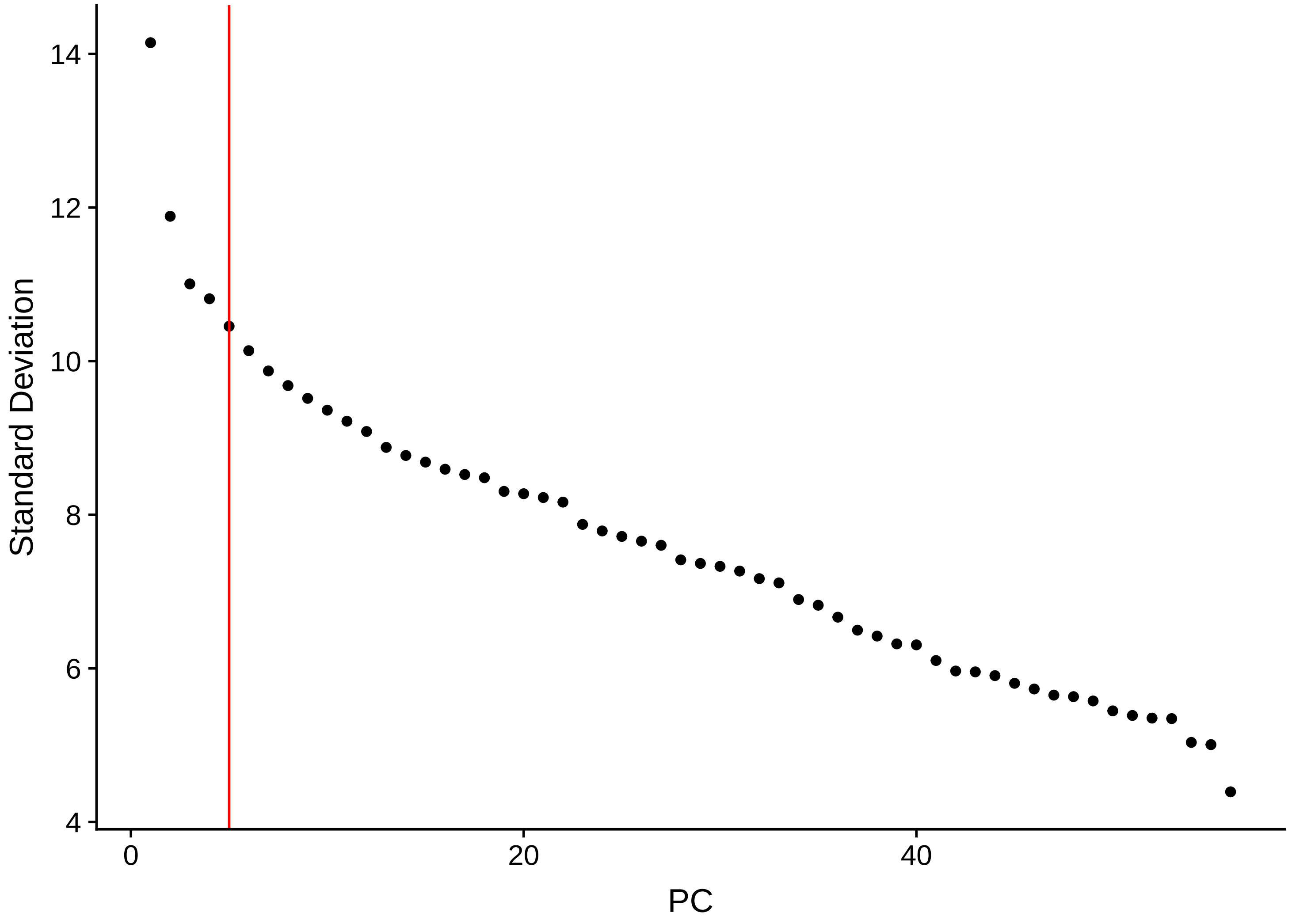
Standard Deviation
14
12
10
8
6
4
0
20
40
PC

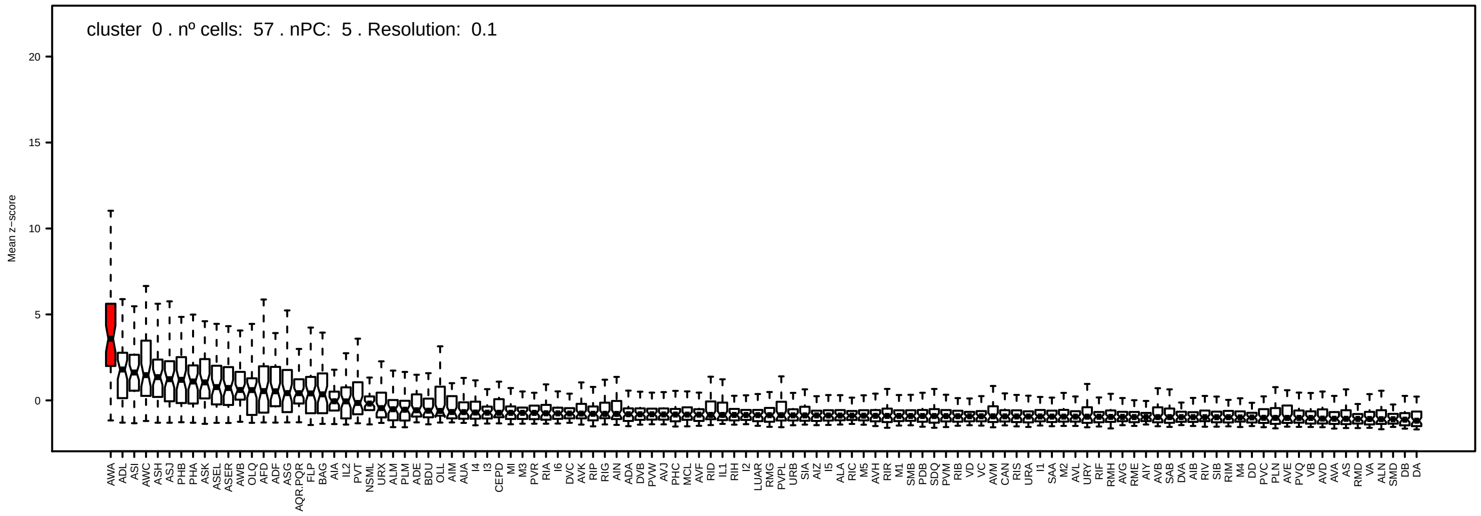
cluster 0 . nº cells: 57 . nPC: 5 . Resolution: 0.1
Mean z−score
0
5
10
15
20
AWA ADL ASI AWC ASH ASJ PHB PHA ASK ASEL ASER AWB OLQ AFD ADF ASG AQR.PQR FLP BAG AIA IL2 PVT NSML URX ALM PLM ADE BDU OLL AIM AUA I4 I3 CEPD MI M3 PVR RIA I6 DVC AVK RIP RIG AIN ADA DVB PVW AVJ PHC MCL AVF RID IL1 RIH I2 LUAR RMG PVPL URB SIA AIZ I5 ALA RIC M5 AVH RIR M1 SMB PDB SDQ PVM RIB VD VC AVM CAN RIS URA I1 SAA M2 AVL URY RIF RMH AVG RME AIY AVB SAB DVA AIB RIV SIB RIM M4 DD PVC PLN AVE PVQ VB AVD AVA AS RMD VA ALN SMD DB DA

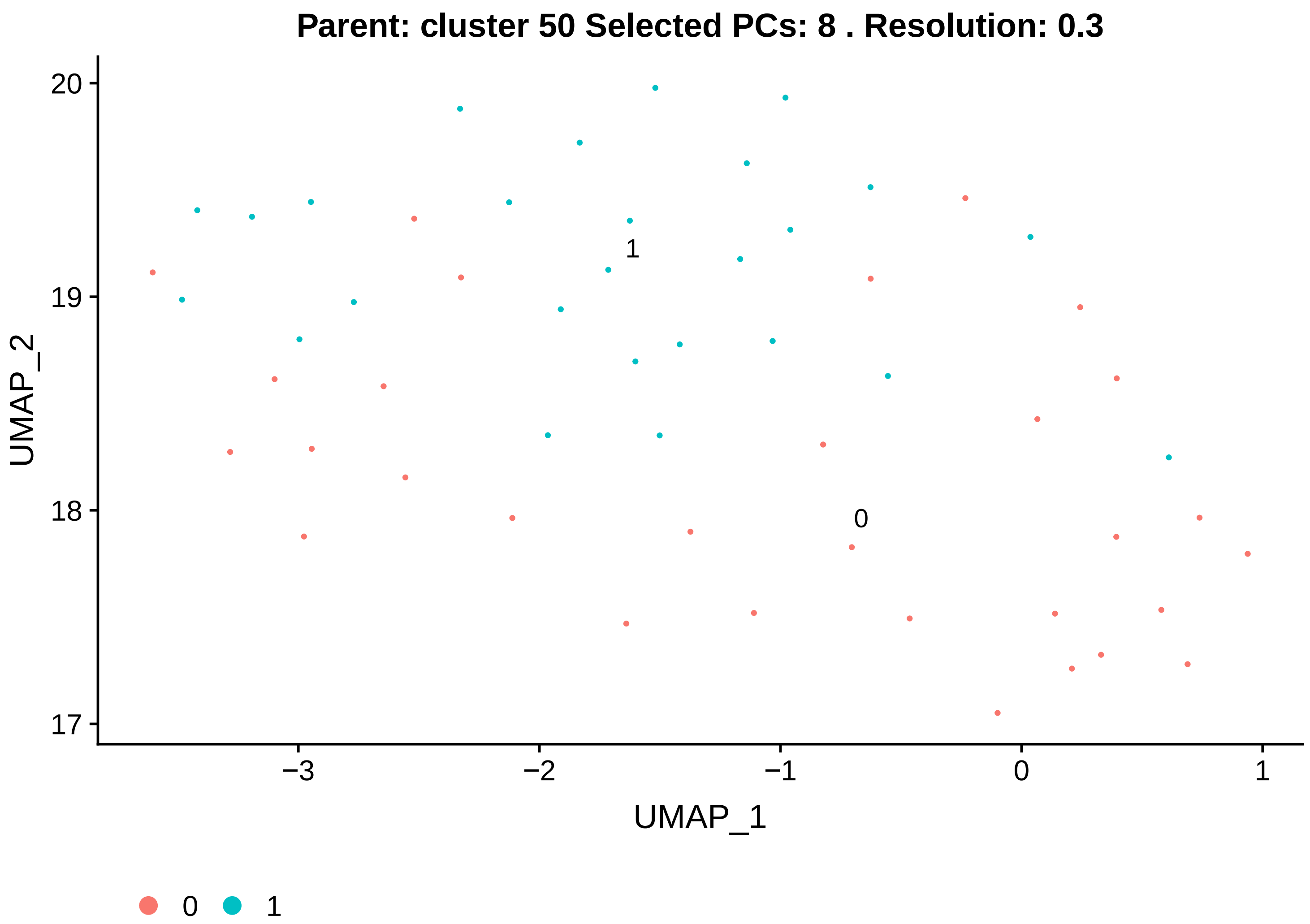
Parent: cluster 50 Selected PCs: 8 . Resolution: 0.3
20
19
18
17
UMAP_2
−3
−2
−1
0
1
UMAP_1
1
0
0
1

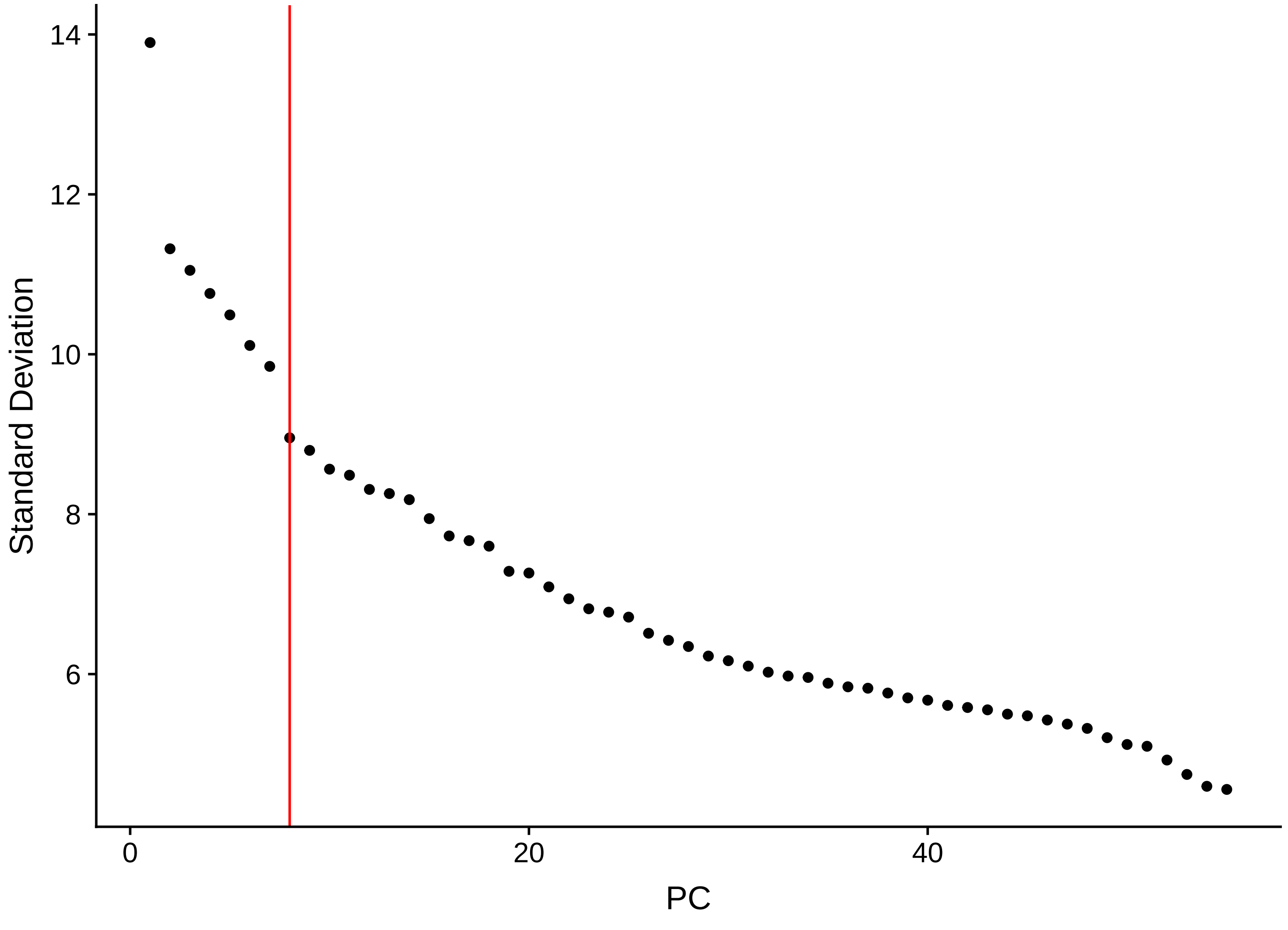
14
12
10
8
6
Standard Deviation
0
20
40
PC

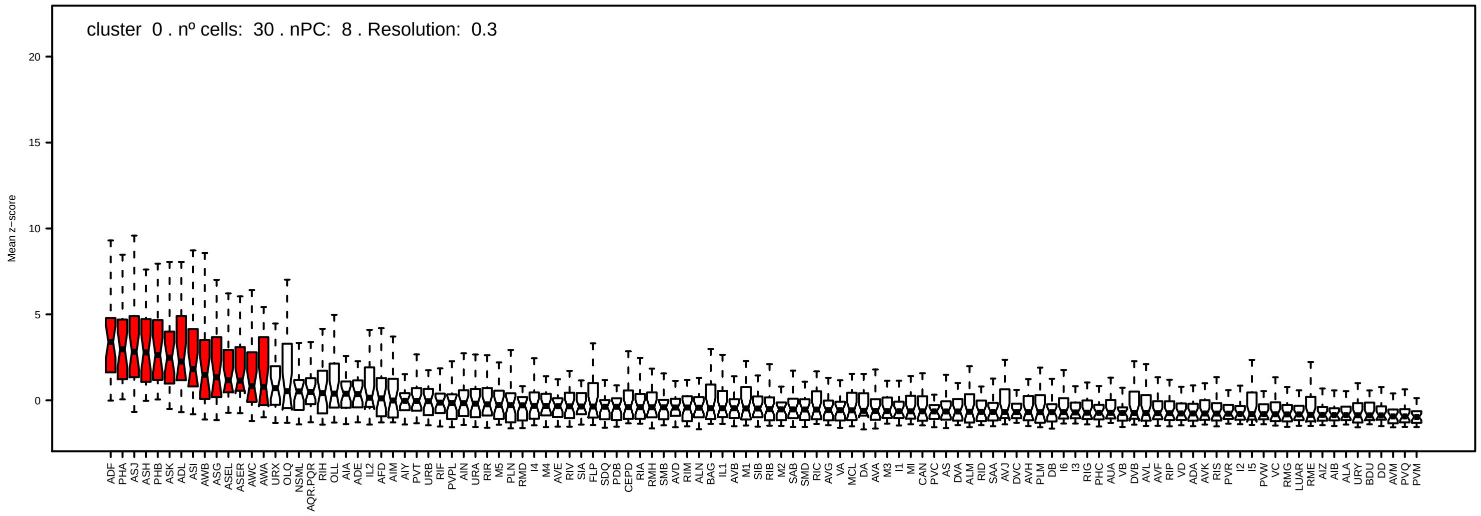
cluster 0 . nº cells: 30 . nPC: 8 . Resolution: 0.3
Mean z−score

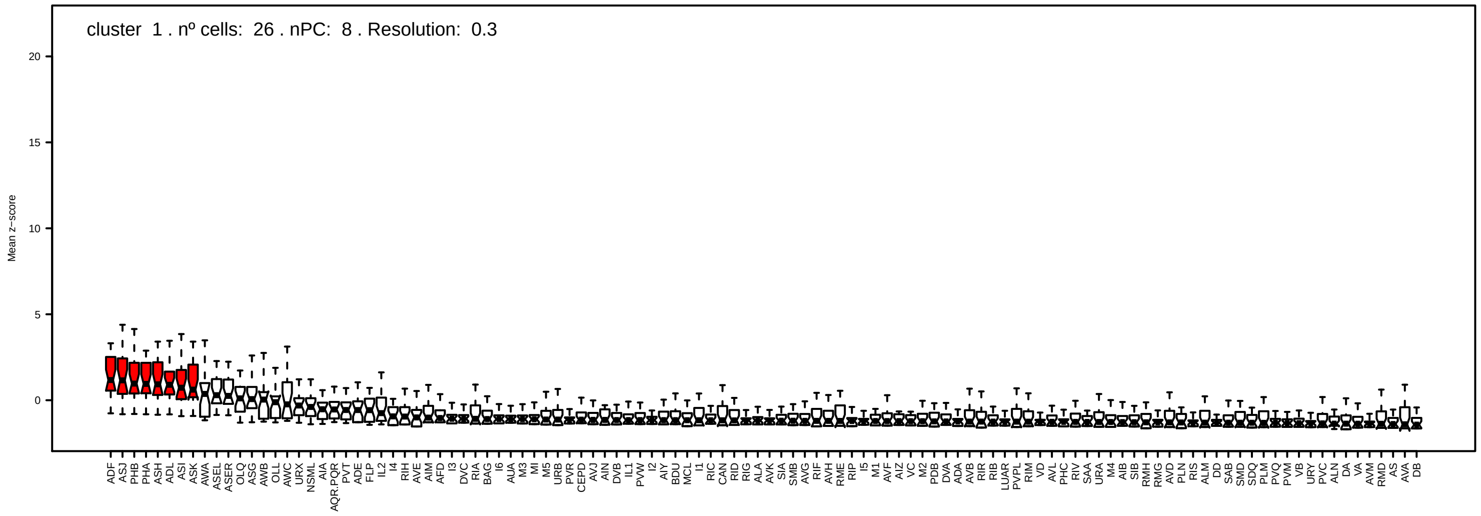
cluster 1 . nº cells: 26 . nPC: 8 . Resolution: 0.3
Mean z−score

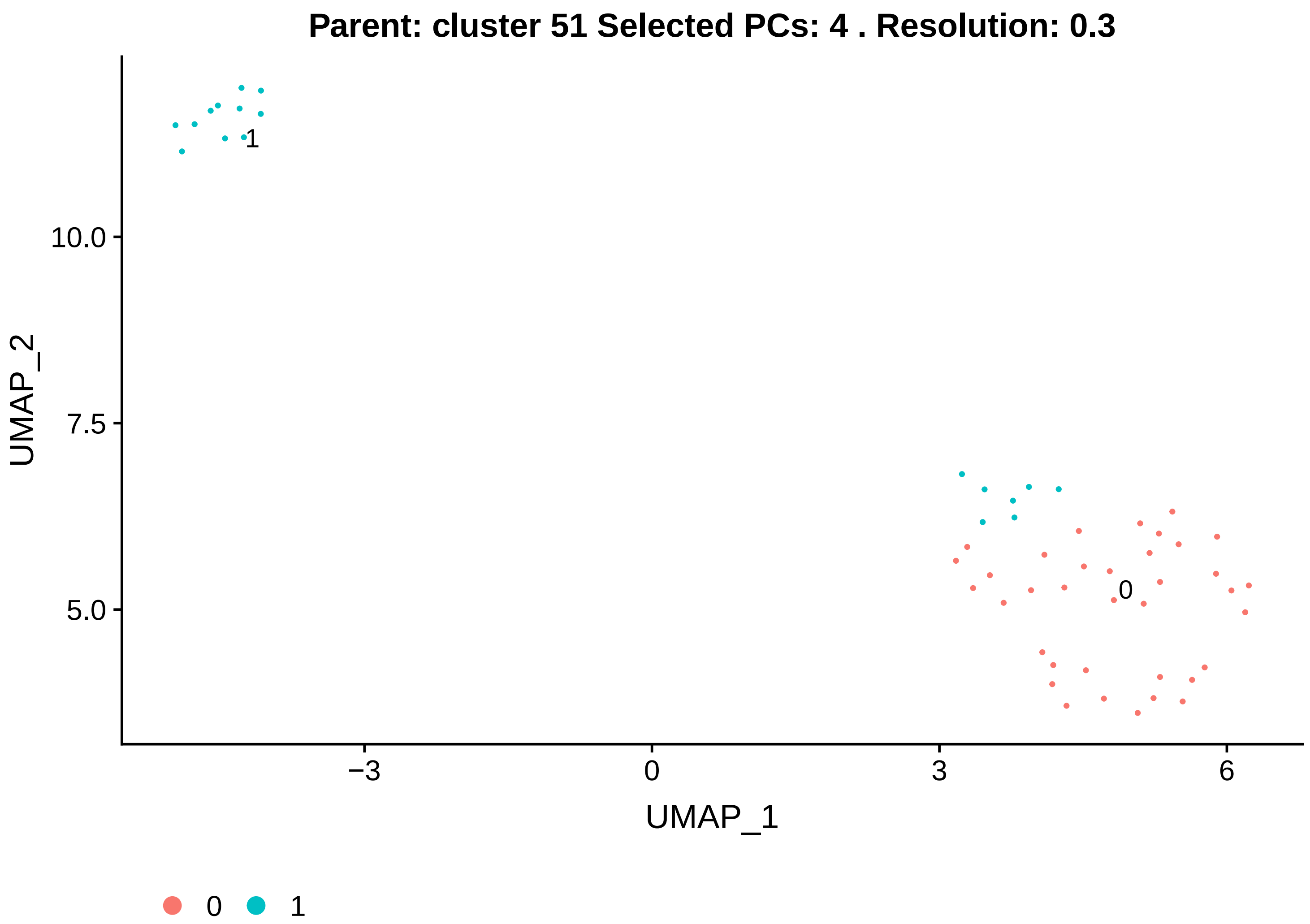
Parent: cluster 51 Selected PCs: 4 . Resolution: 0.3
UMAP_2
UMAP_1
10.0
7.5
5.0
−3
0
3
6
1
0
0
1

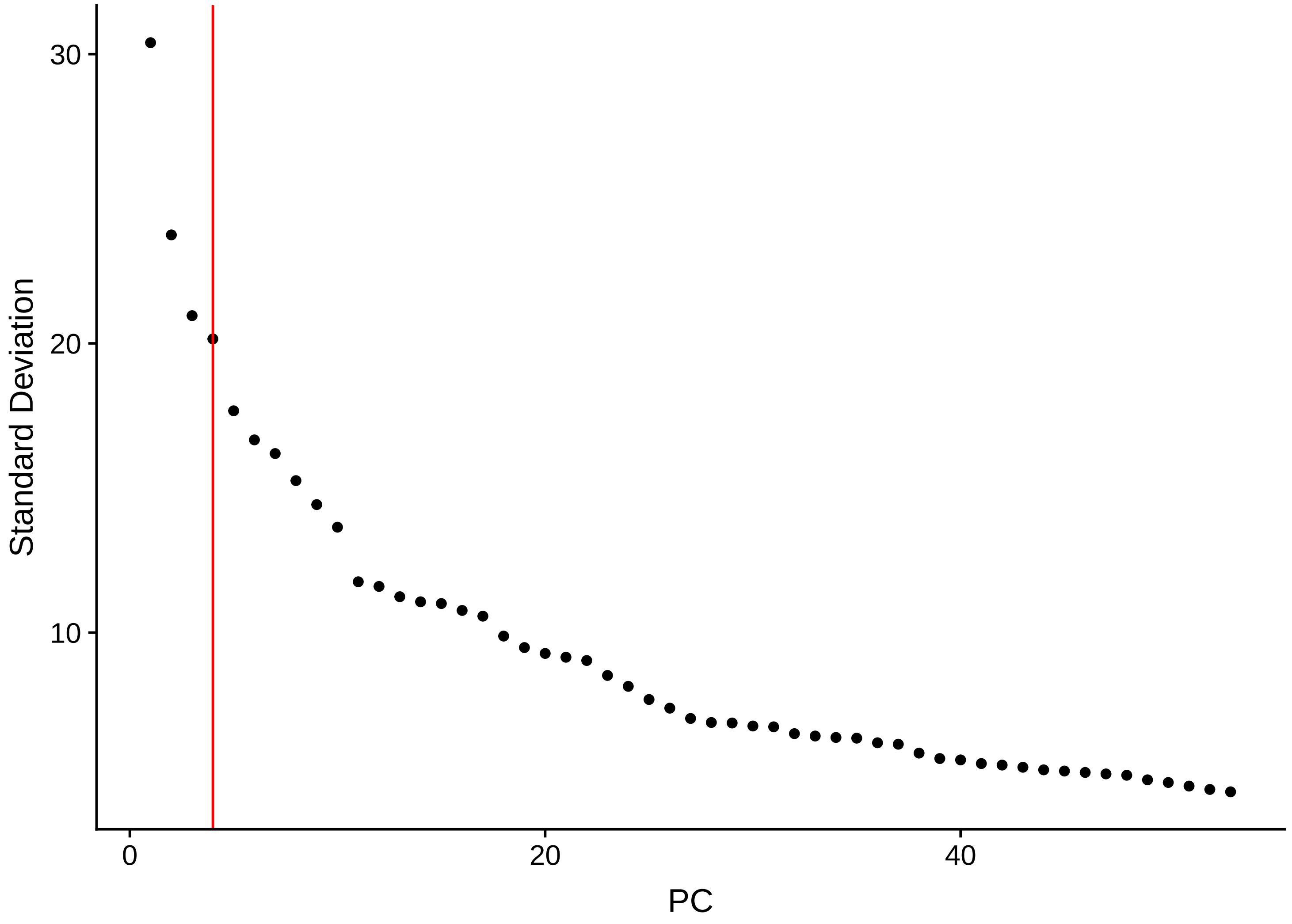
Standard Deviation
PC
30
20
10
0
20
40

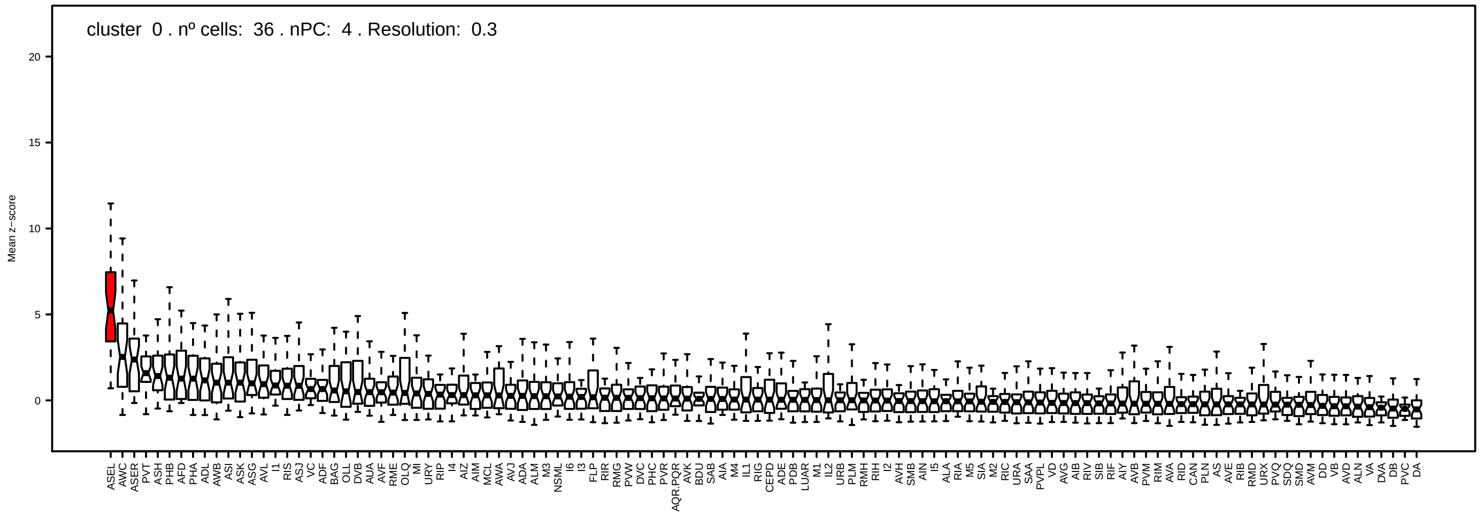
cluster 0 . nº cells: 36 . nPC: 4 . Resolution: 0.3
Mean z−score

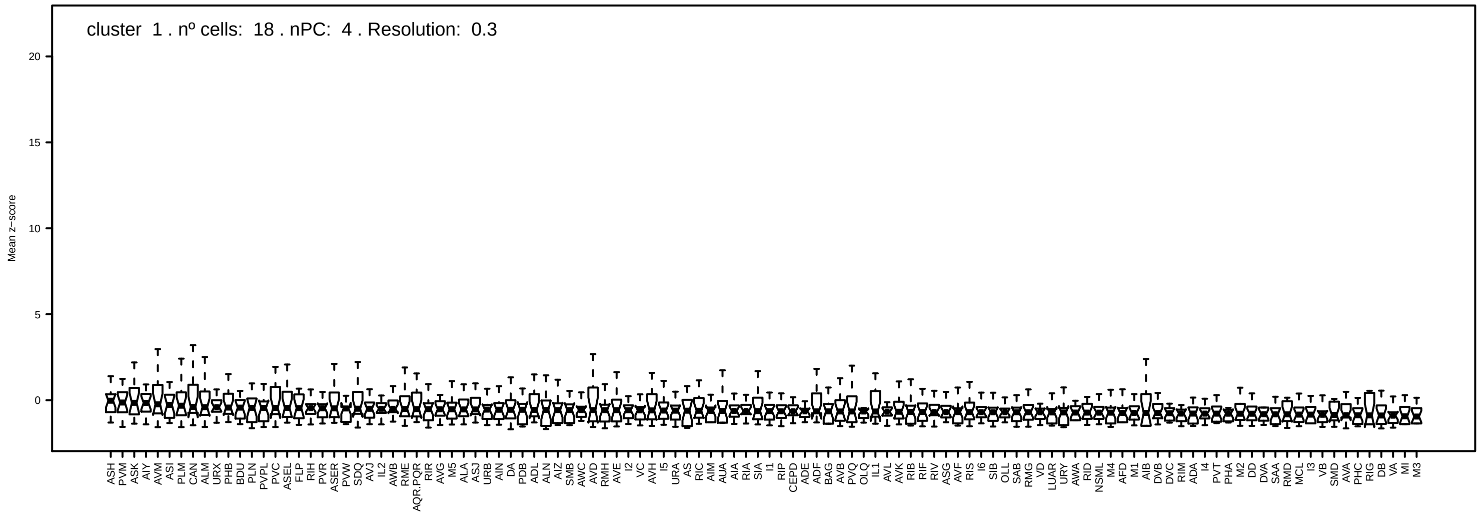
cluster 1 . nº cells: 18 . nPC: 4 . Resolution: 0.3
Mean z−score

### Parent: cluster 52 Selected PCs: 4 . Resolution: 0.1

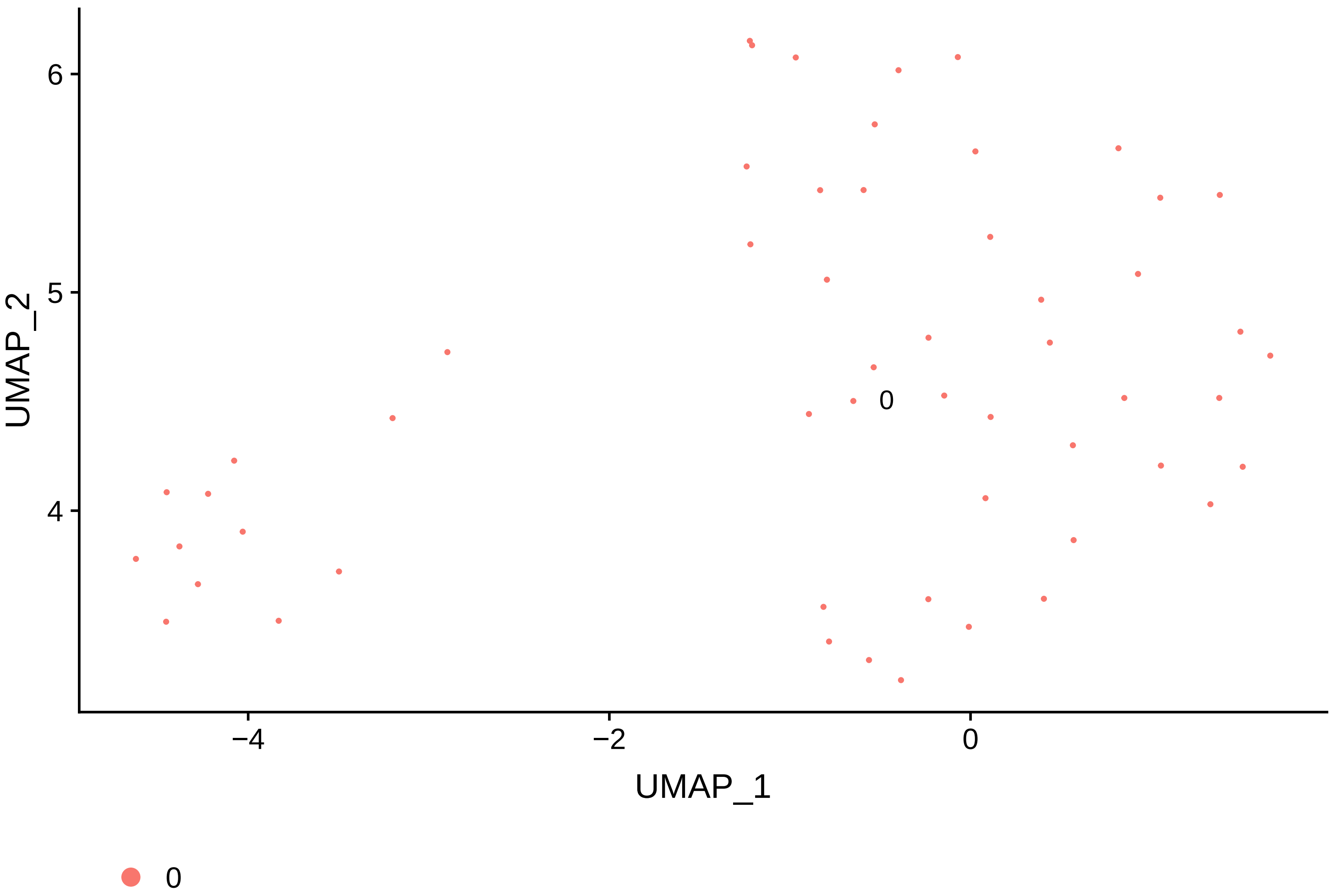


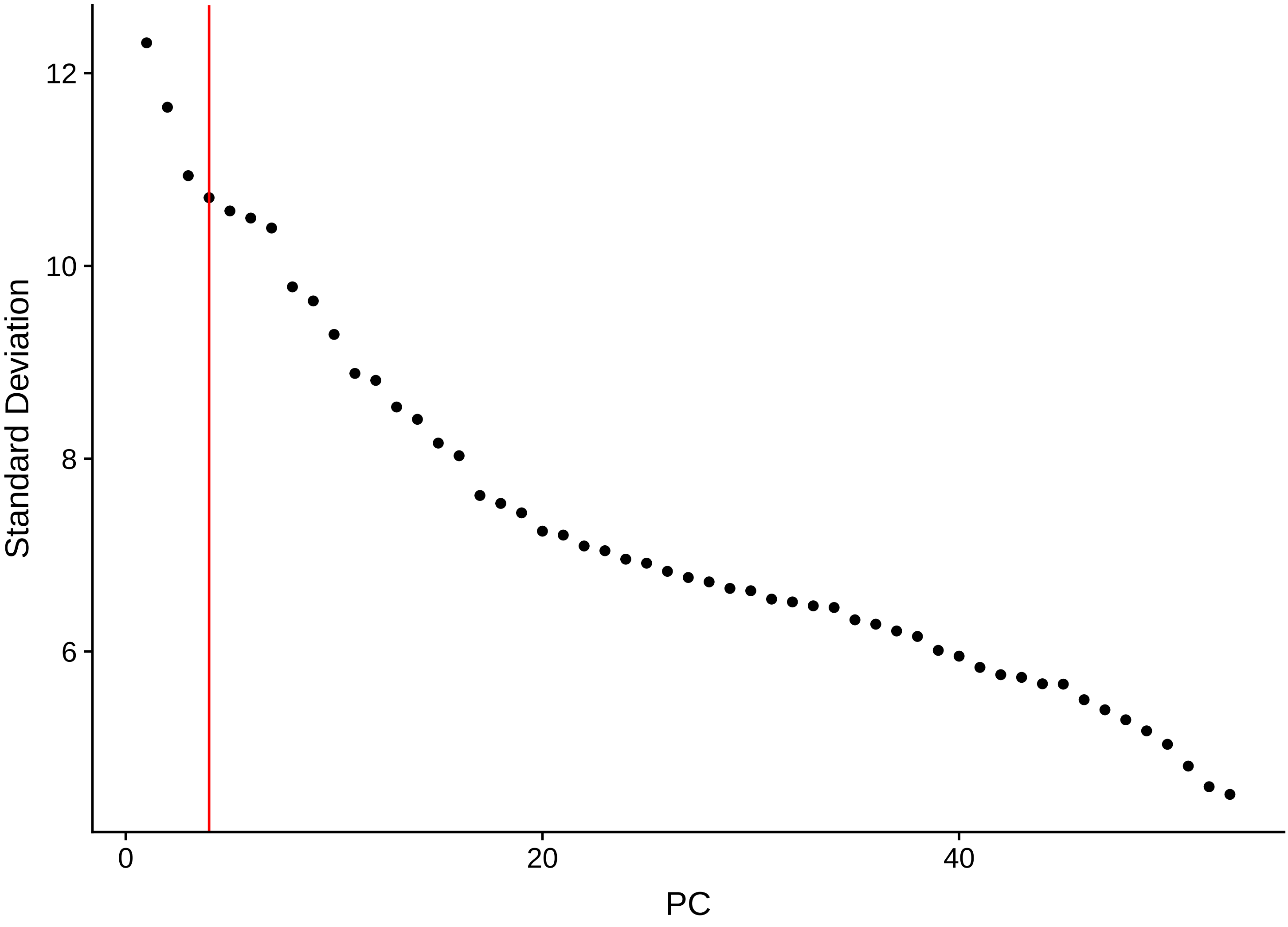

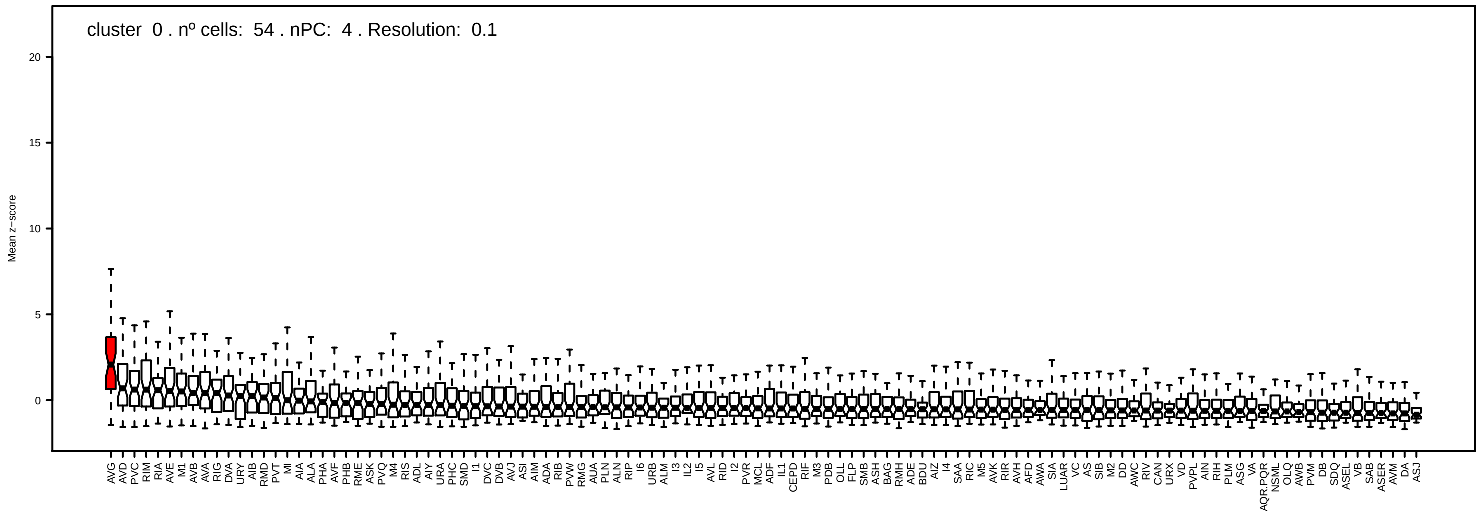

cluster 0 . nº cells: 54 . nPC: 4 . Resolution: 0.1
Mean z−score
20
15
10
5
0
AVG AVD PVC RIM RIA AVE M1 AVB AVA RIG DVA URY AIB RMD PVT MI AIA ALA PHA AVF PHB RME ASK PVQ M4 RIS ADL AIY URA PHC SMD I1 DVC DVB AVJ ASI AIM ADA RIB PVW RMG AUA PLN ALN RIP I6 URB ALM I3 IL2 I5 AVL RID I2 PVR MCL ADF IL1 CEPD RIF M3 PDB OLL FLP SMB ASH BAG RMH ADE BDU AIZ I4 SAA RIC M5 AVK RIR AVH AFD AWA SIA LUAR VC AS SIB M2 DD AWC RIV CAN URX VD PVPL AIN RIH PLM ASG VA AQR.PQR NSML OLQ AWB PVM DB SDQ ASEL VB SAB ASER AVM DA ASJ

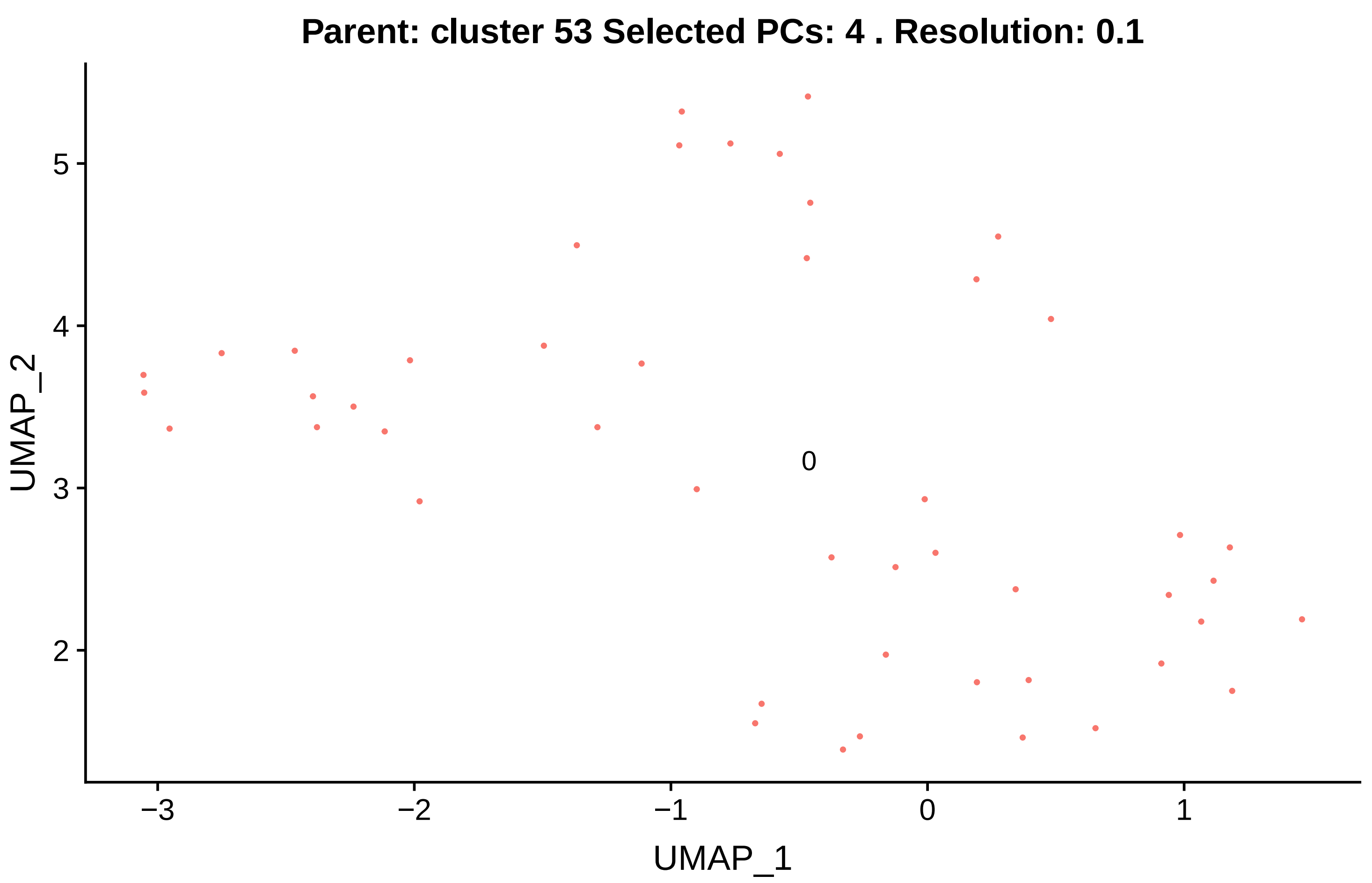

Parent: cluster 53 Selected PCs: 4 . Resolution: 0.1
UMAP_2
UMAP_1
0
−3
−2
−1
0
1
2
3
4
5


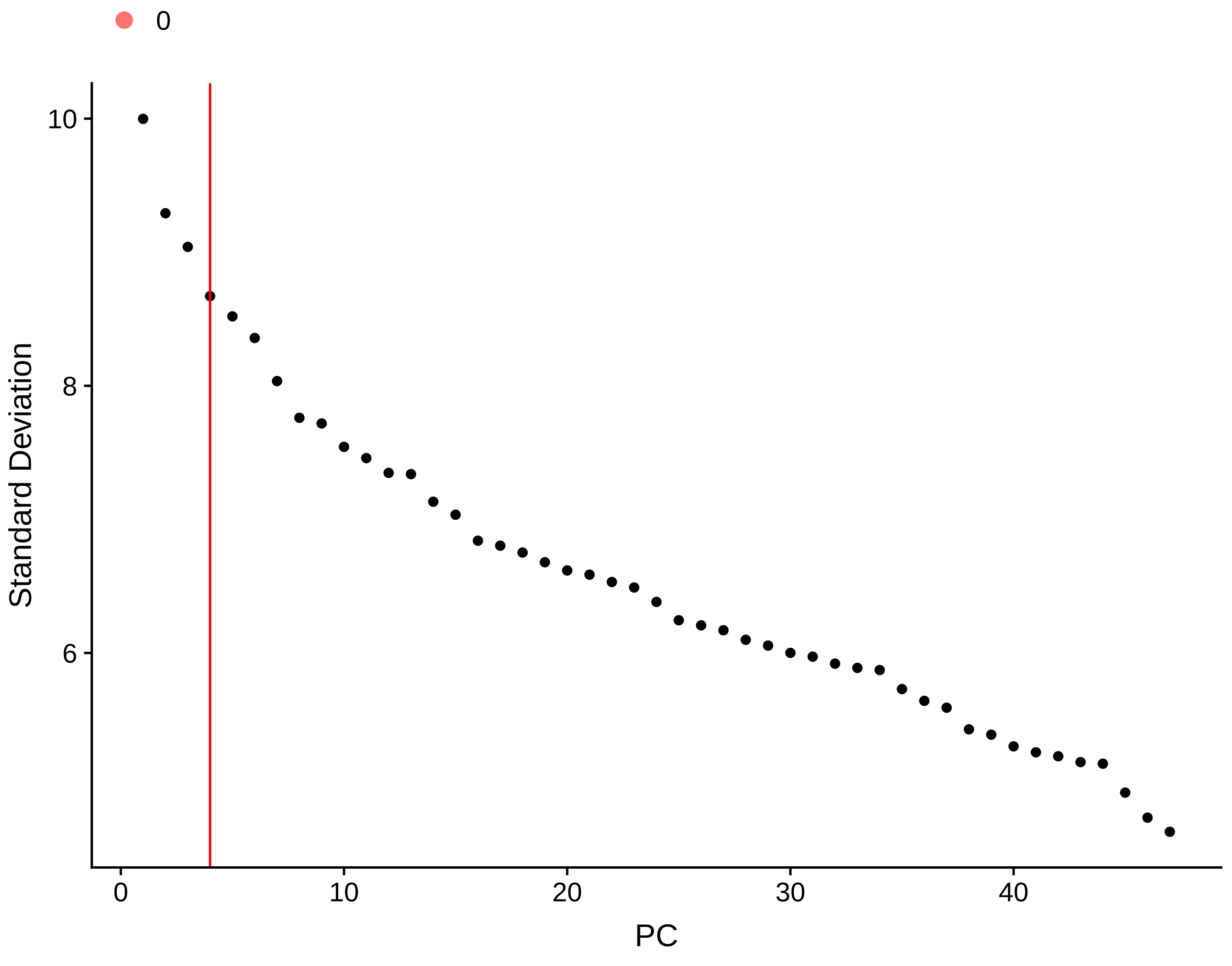

0
Standard Deviation
PC
10
8
6
0
10
20
30
40

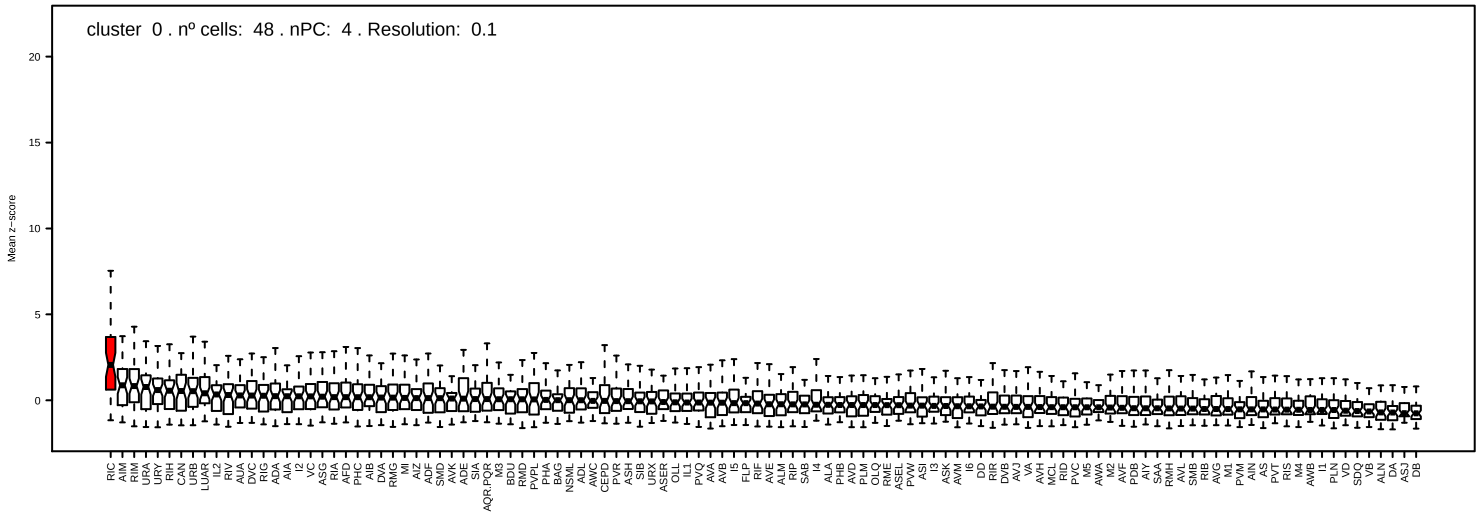
cluster 0 . nº cells: 48 . nPC: 4 . Resolution: 0.1
Mean z−score

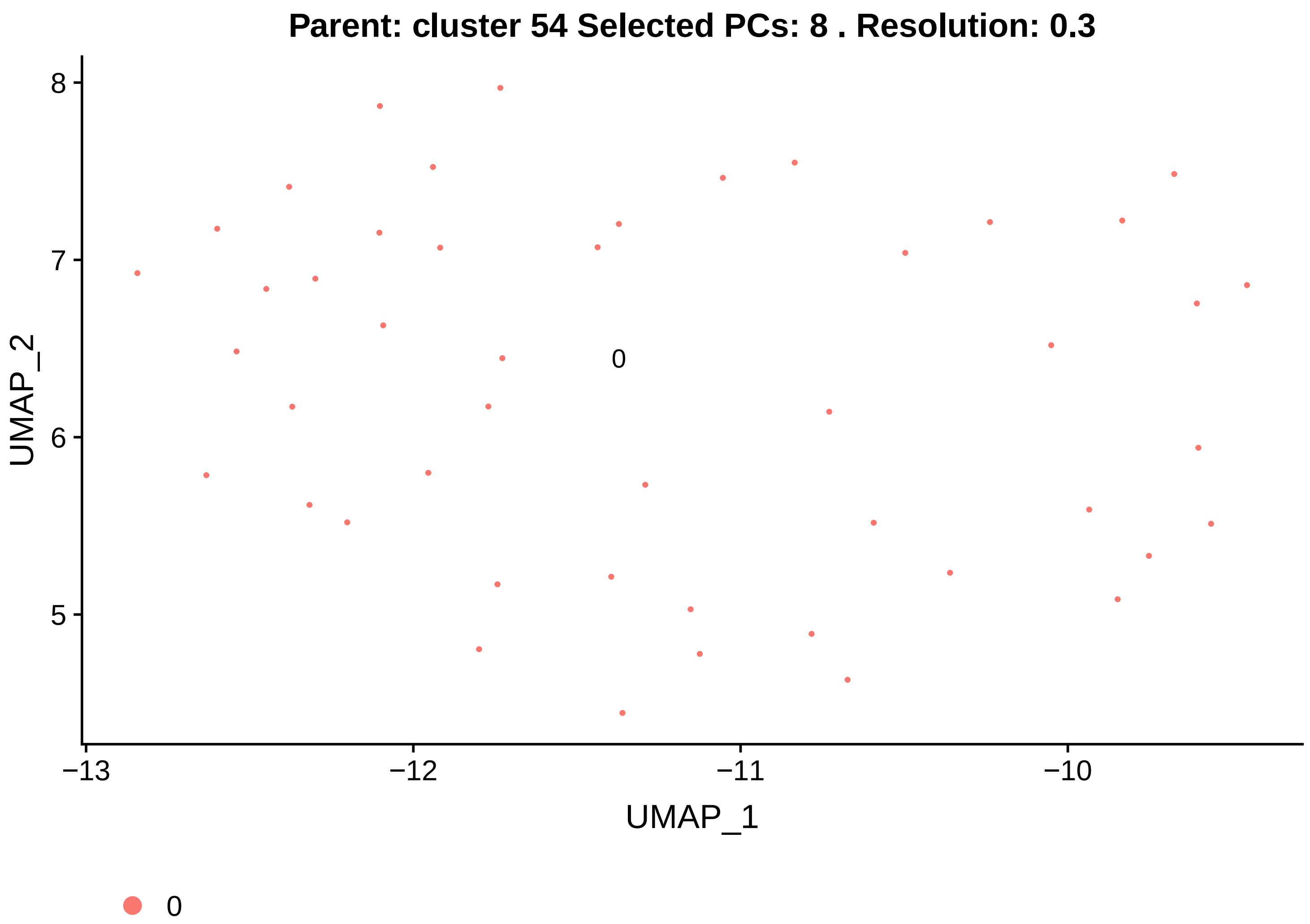

Parent: cluster 54 Selected PCs: 8 . Resolution: 0.3
UMAP_2
UMAP_1
0


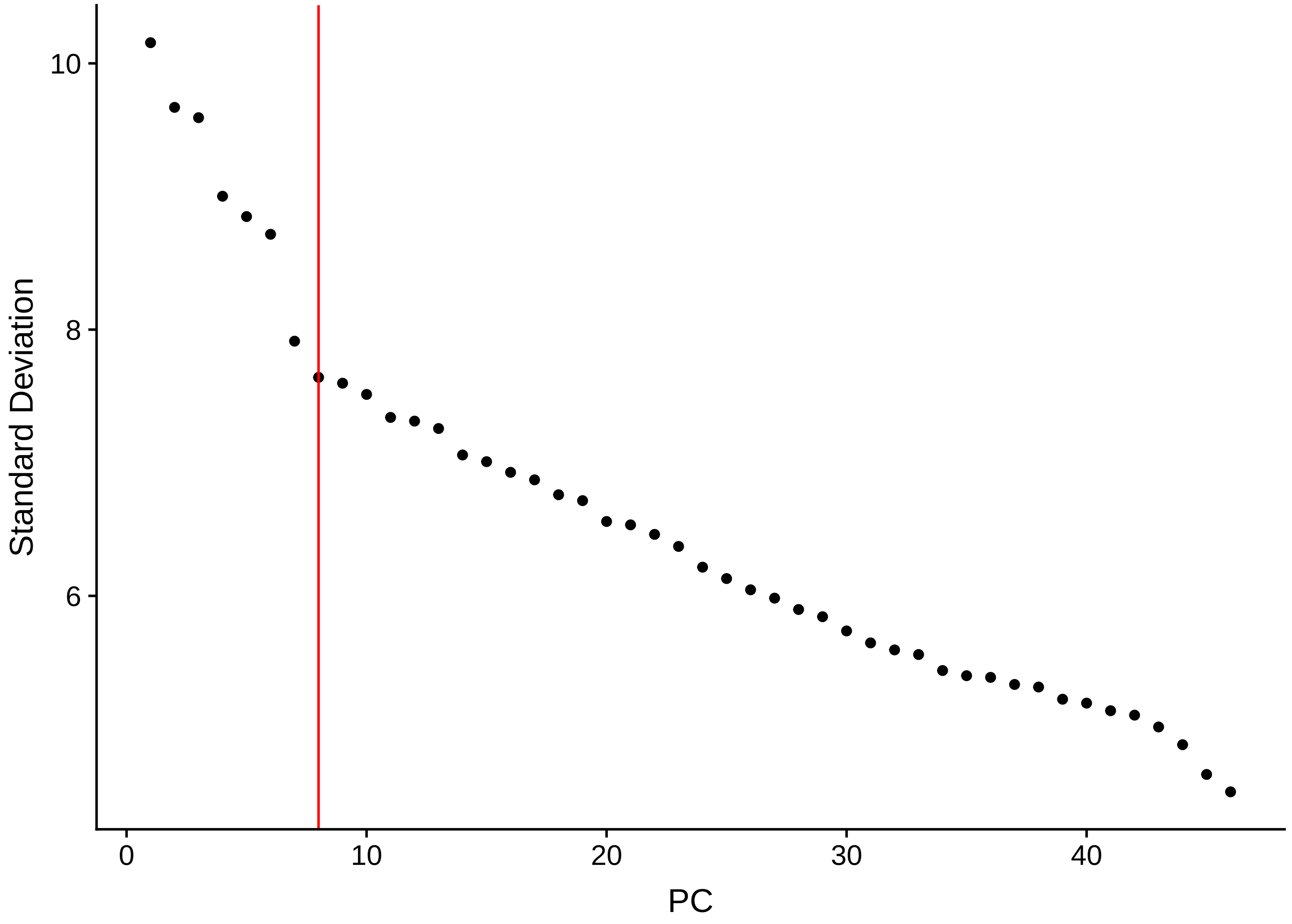

Standard Deviation
PC

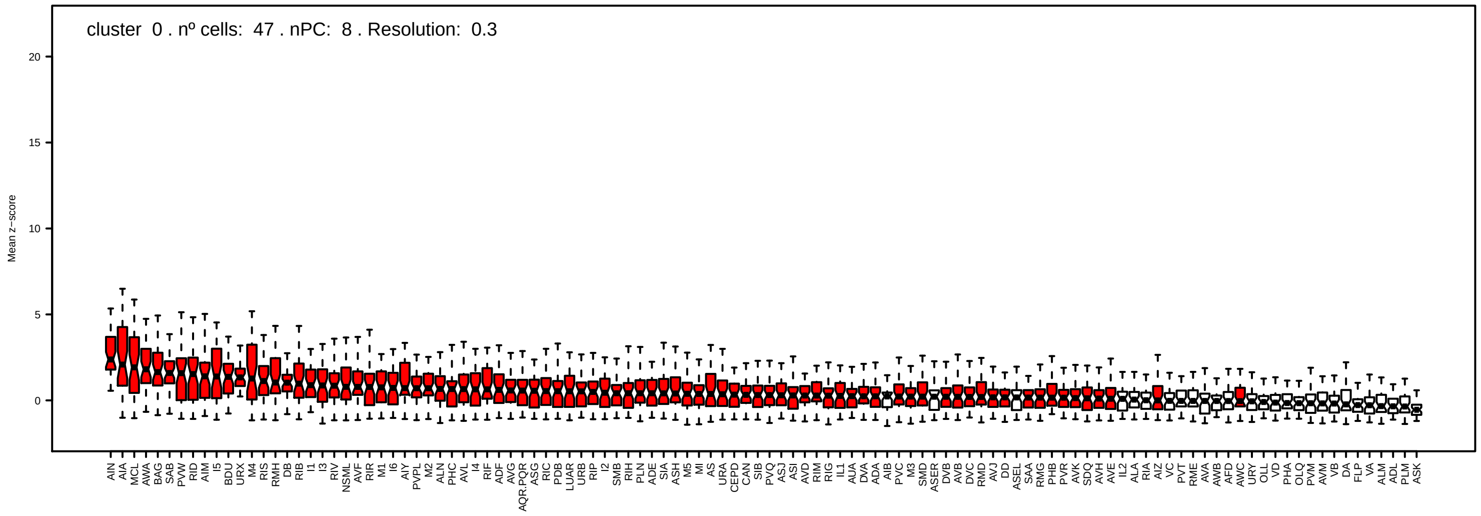
cluster 0 . nº cells: 47 . nPC: 8 . Resolution: 0.3
Mean z−score
AIN AIA MCL AWA BAG SAB PVW RID AIM I5 BDU URX M4 RIS RMH DB RIB I1 I3 RIV NSML AVF RIR M1 I6 AIY PVPL M2 ALN PHC AVL I4 RIF ADF AVG AQR.PQR ASG RIC PDB LUAR URB RIP I2 SMB RIH PLN ADE SIA ASH M5 MI AS URA CEPD CAN SIB PVQ ASJ ASI AVD RIM RIG IL1 AUA DVA ADA AIB PVC M3 SMD ASER DVB AVB DVC RMD AVJ DD ASEL SAA RMG PHB PVR AVK SDQ AVH AVE IL2 ALA RIA AIZ VC PVT RME AVA AWB AFD AWC URY OLL VD PHA OLQ PVM AVM VB DA FLP VA ALM ADL PLM ASK

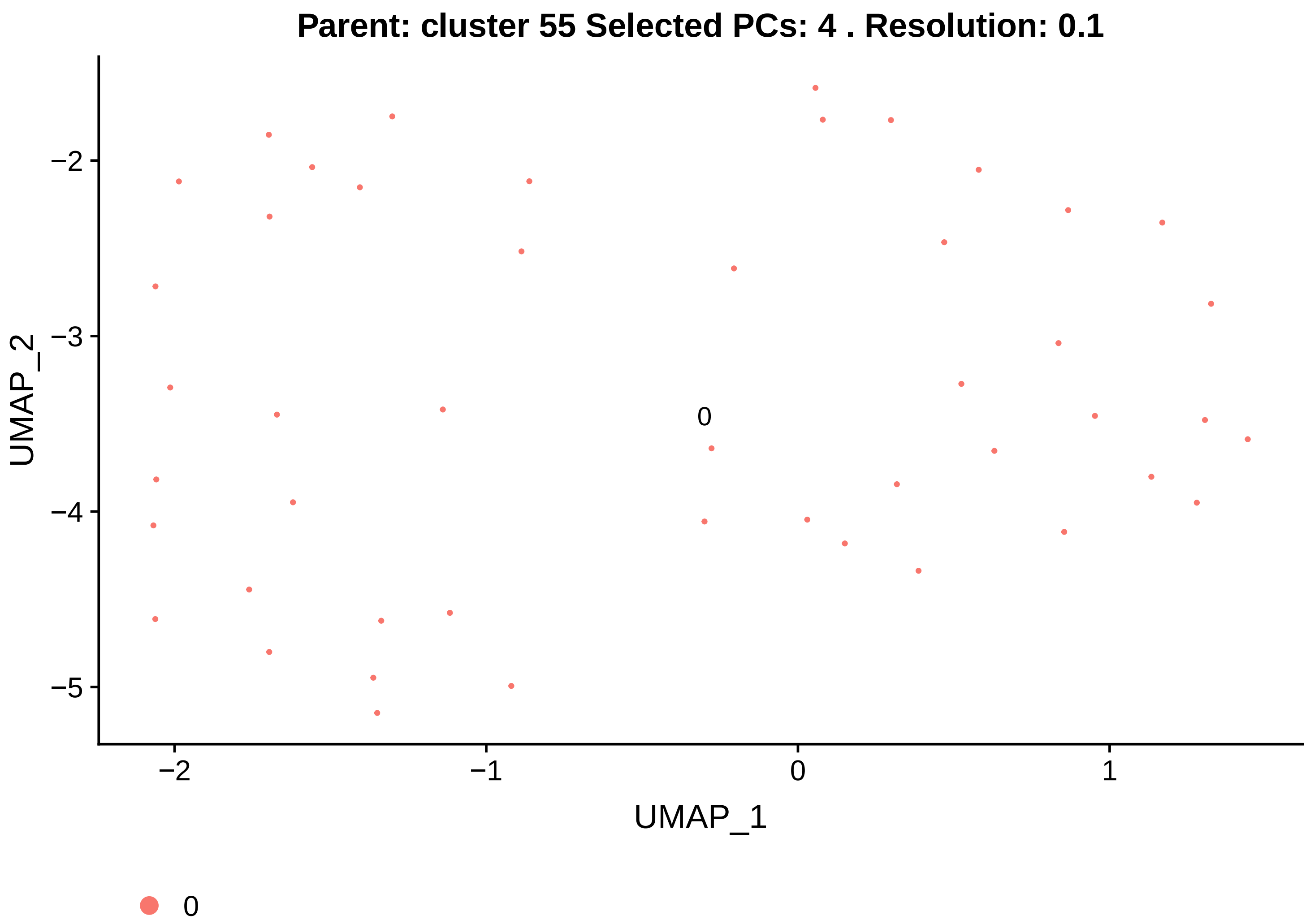
Parent: cluster 55 Selected PCs: 4 . Resolution: 0.1
UMAP_2
−2
−3
−4
−5
0
−2
−1
0
1
UMAP_1
0

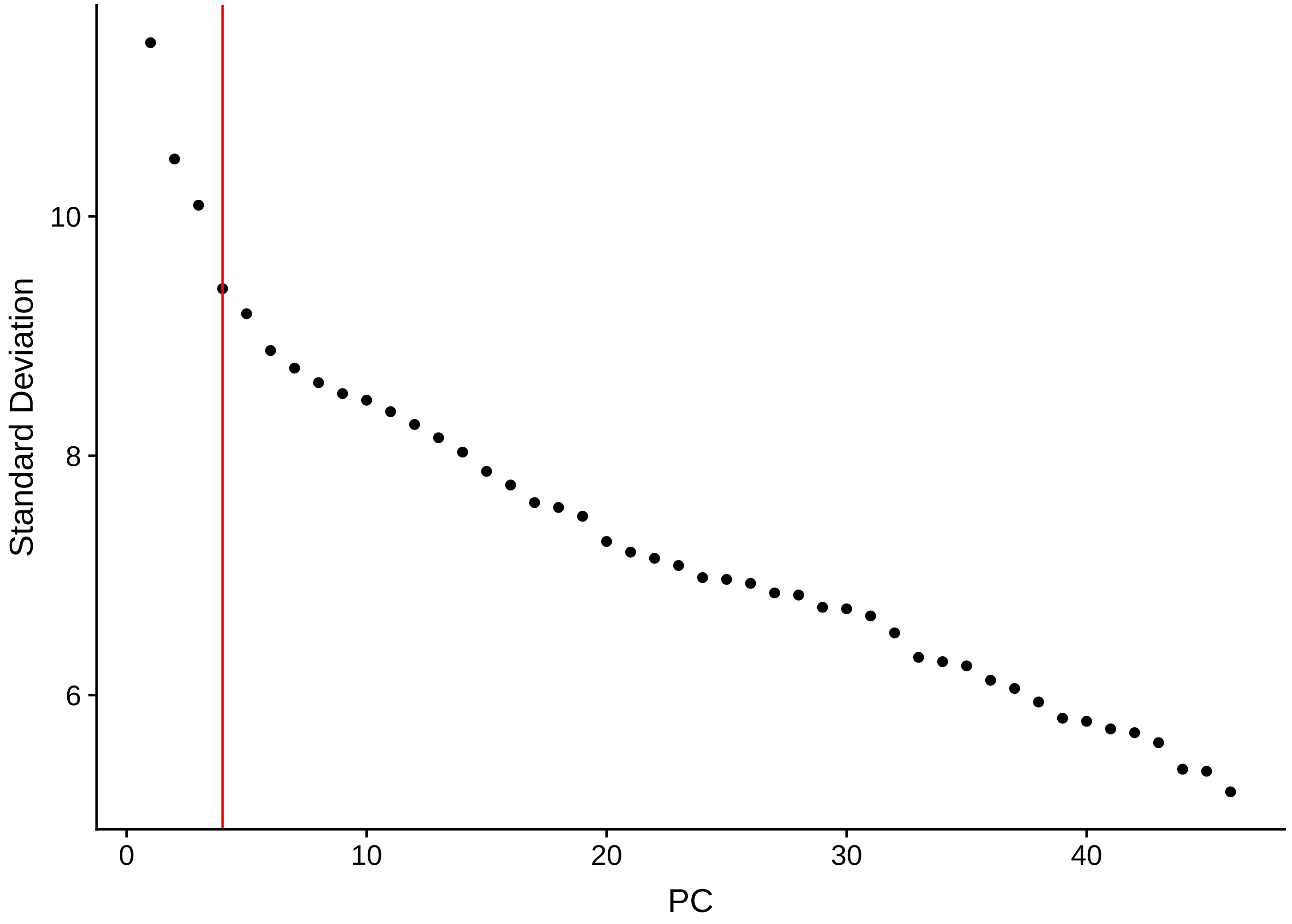
Standard Deviation
10
8
6
0
10
20
30
40
PC

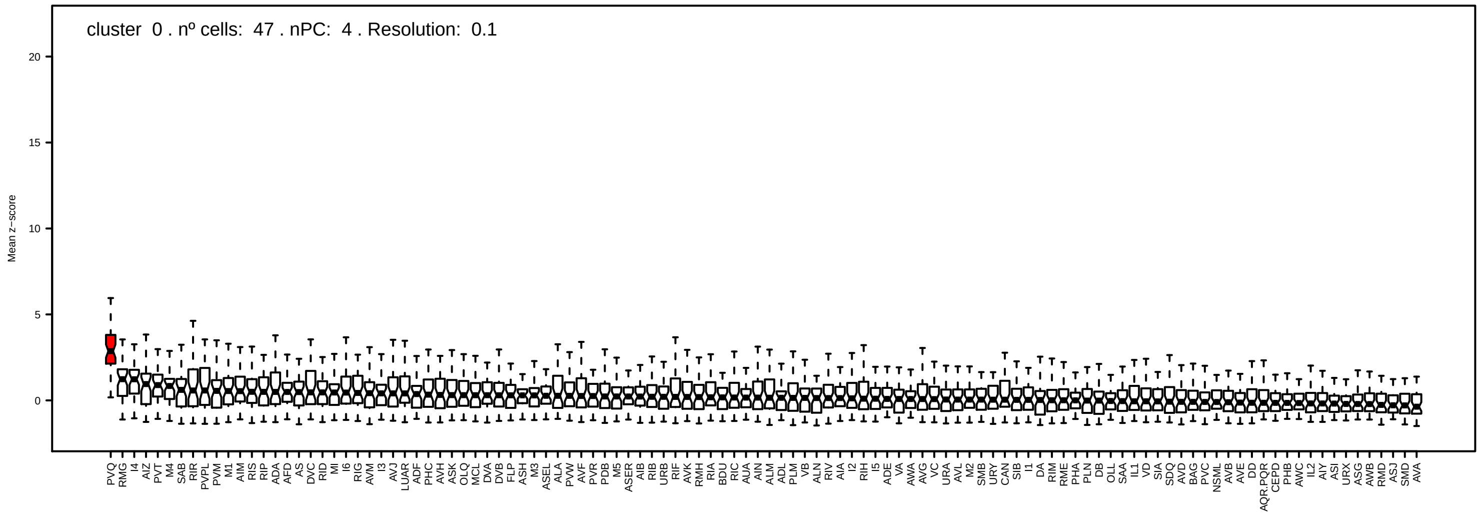
cluster 0 . nº cells: 47 . nPC: 4 . Resolution: 0.1
Mean z−score
20
15
10
5
0
PVQ RMG I4 AIZ PVT M4 SAB RIR PVPL PVM M1 AIM RIS RIP ADA AFD AS DVC RID MI I6 RIG AVM I3 AVJ LUAR ADF PHC AVH ASK OLQ MCL DVA DVB FLP ASH M3 ASEL ALA PVW AVF PVR PDB M5 ASER AIB RIB URB RIF AVK RMH RIA BDU RIC AUA AIN ALM ADL PLM VB ALN RIV AIA I2 RIH I5 ADE VA AWA AVG VC URA AVL M2 SMB URY CAN SIB I1 DA RIM RME PHA PLN DB OLL SAA IL1 VD SIA SDQ AVD BAG PVC NSML AVB AVE DD AQR.PQR CEPD PHB AWC IL2 AIY ASI URX ASG AWB RMD ASJ SMD AVA

Parent: cluster 56 Selected PCs: 10 . Resolution: 1

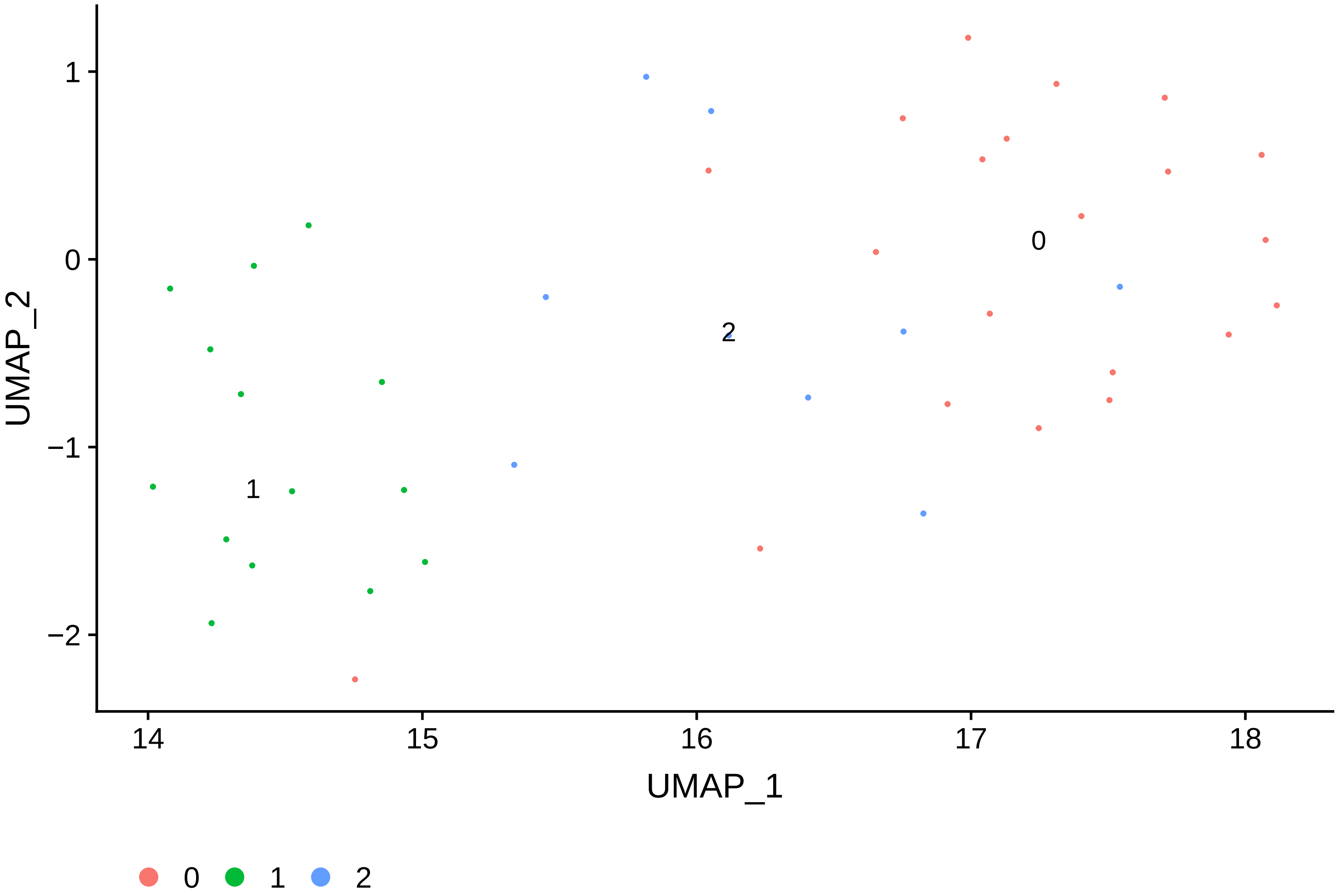


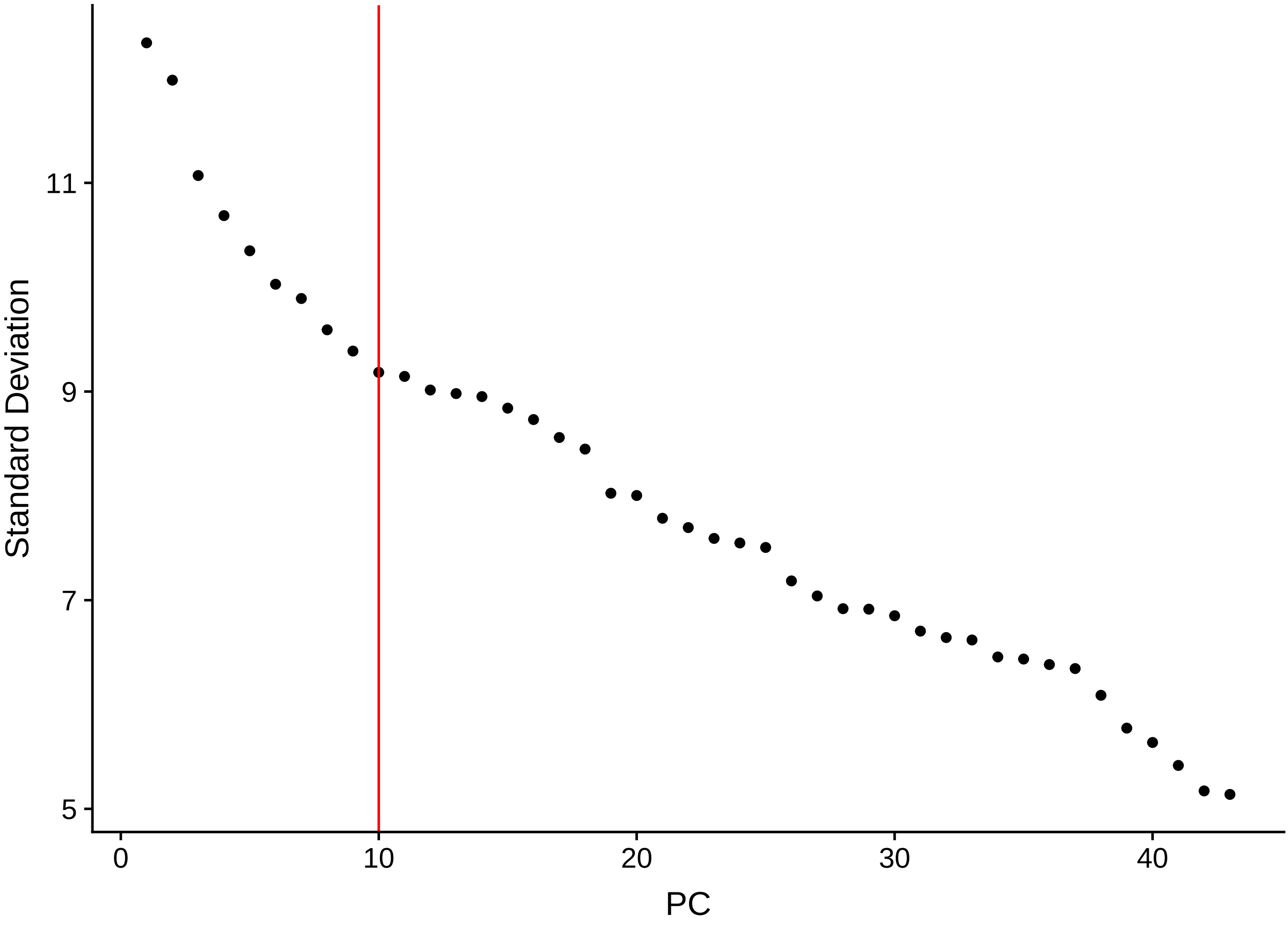

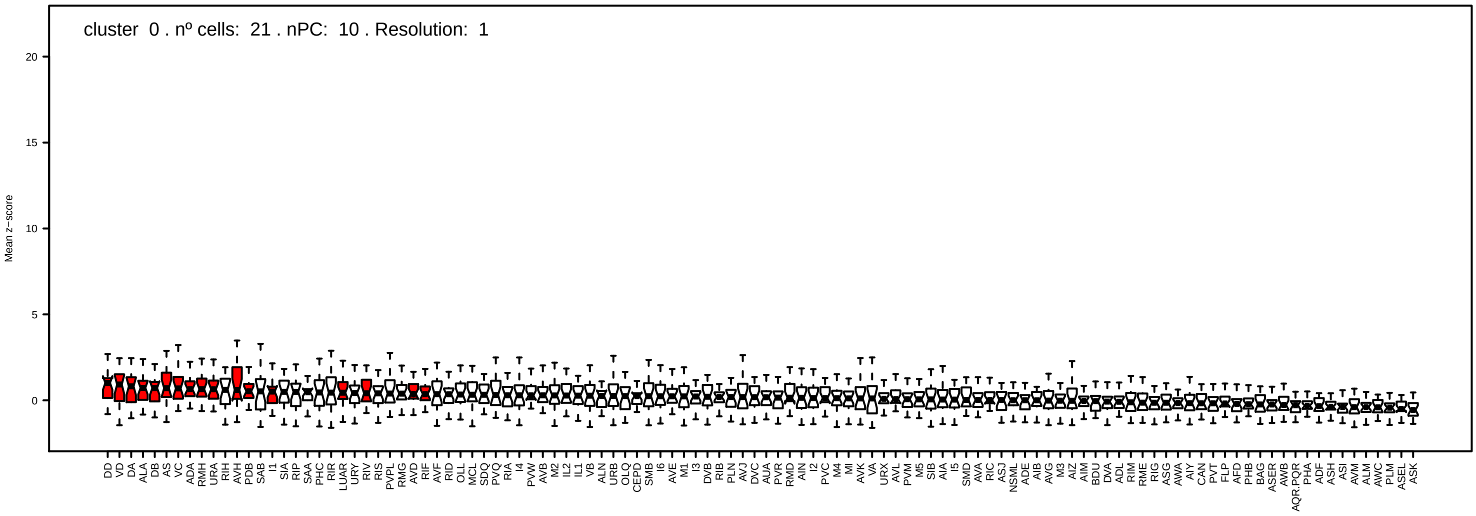
cluster 0 . nº cells: 21 . nPC: 10 . Resolution: 1
Mean z−score

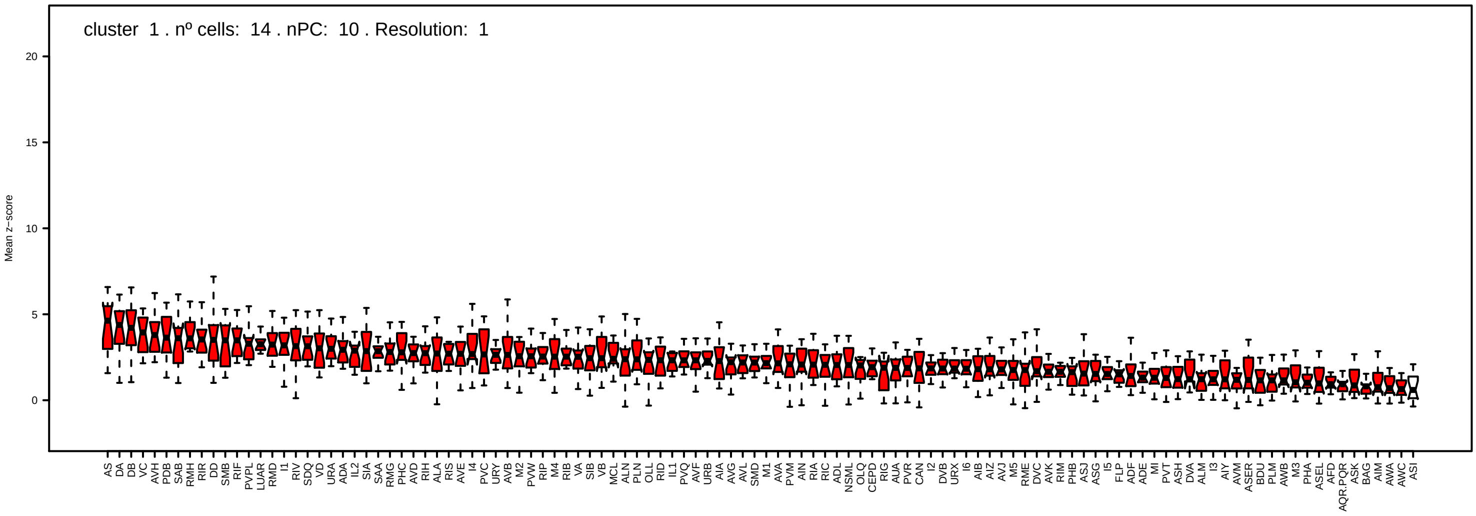
cluster 1 . nº cells: 14 . nPC: 10 . Resolution: 1
Mean z−score

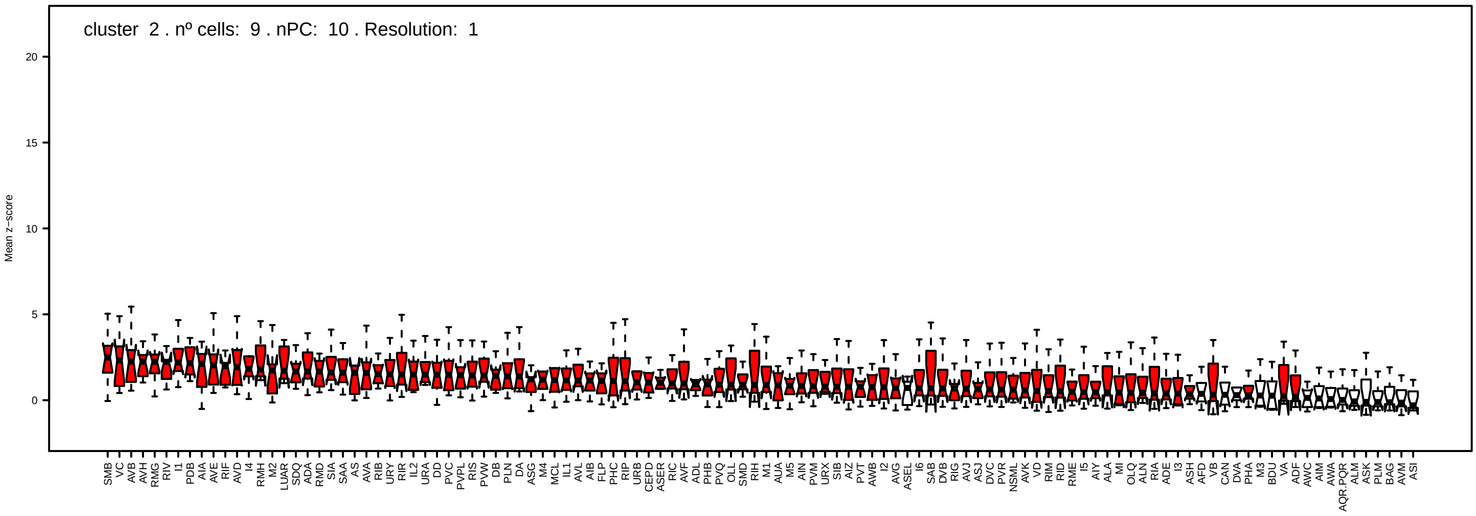
cluster 2 . nº cells: 9 . nPC: 10 . Resolution: 1
Mean z−score

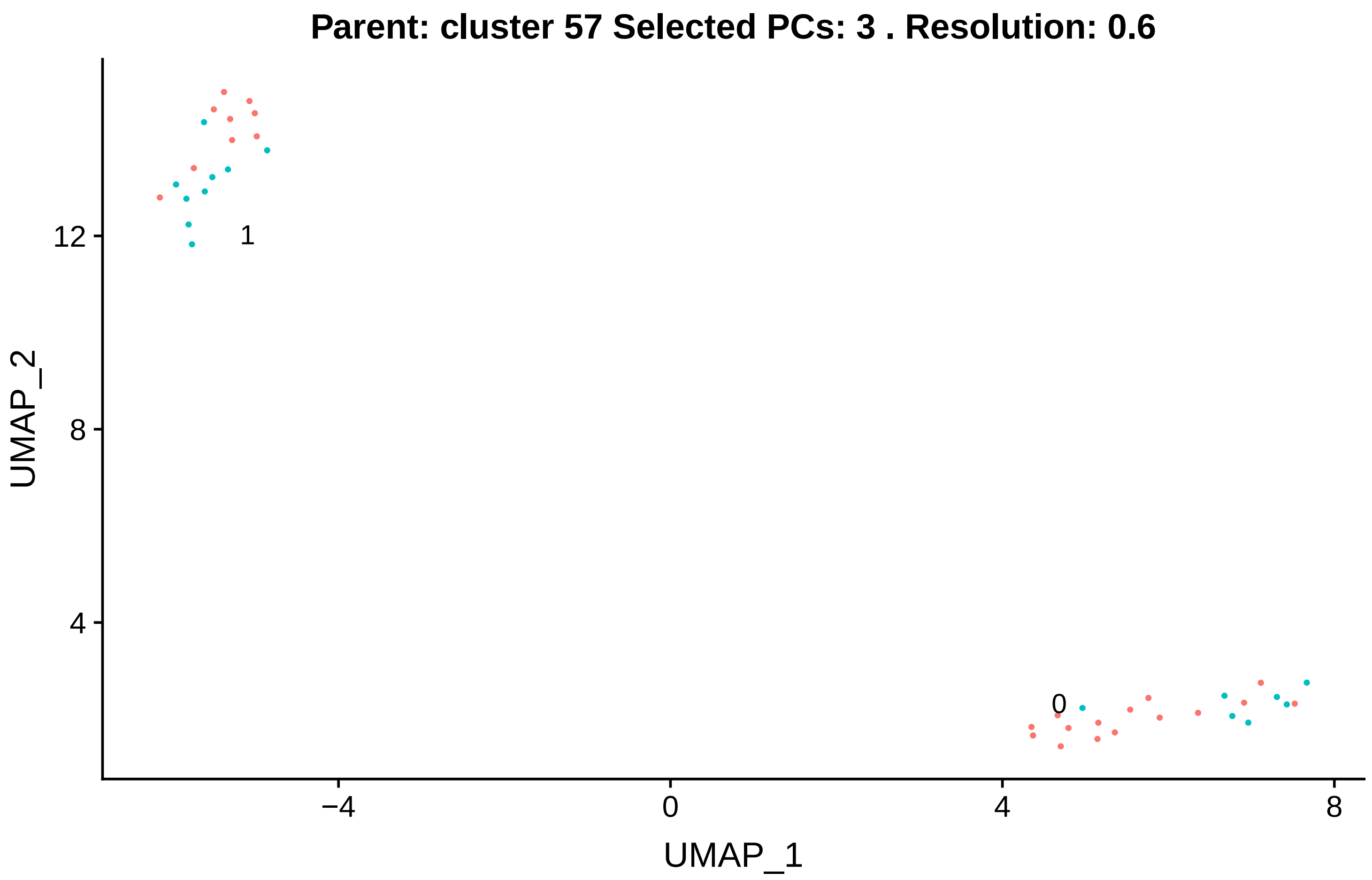
Parent: cluster 57 Selected PCs: 3 . Resolution: 0.6
UMAP_2
UMAP_1
12
8
4
−4
0
4
8
1
0

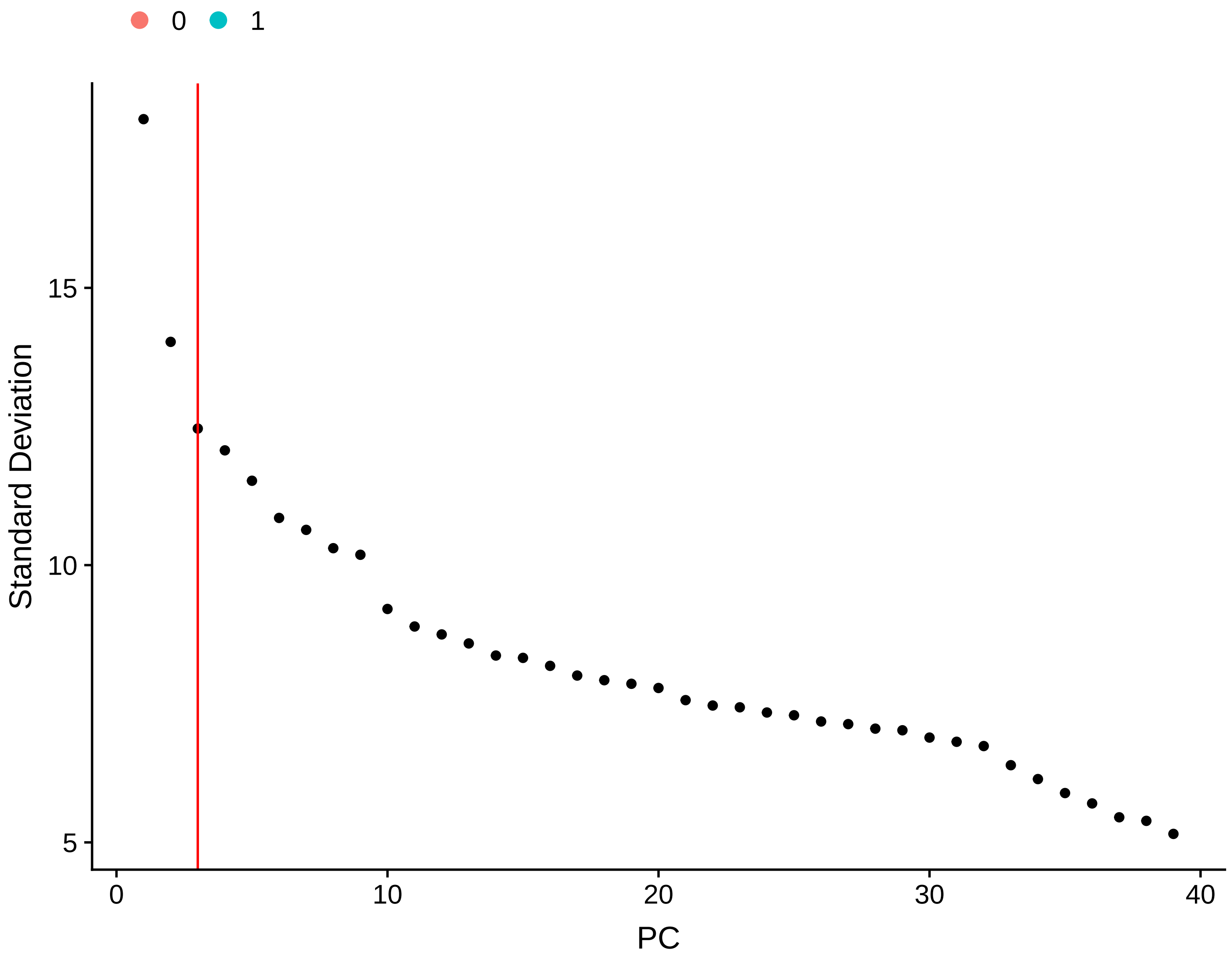
0
1
Standard Deviation
PC
15
10
5
0
10
20
30
40

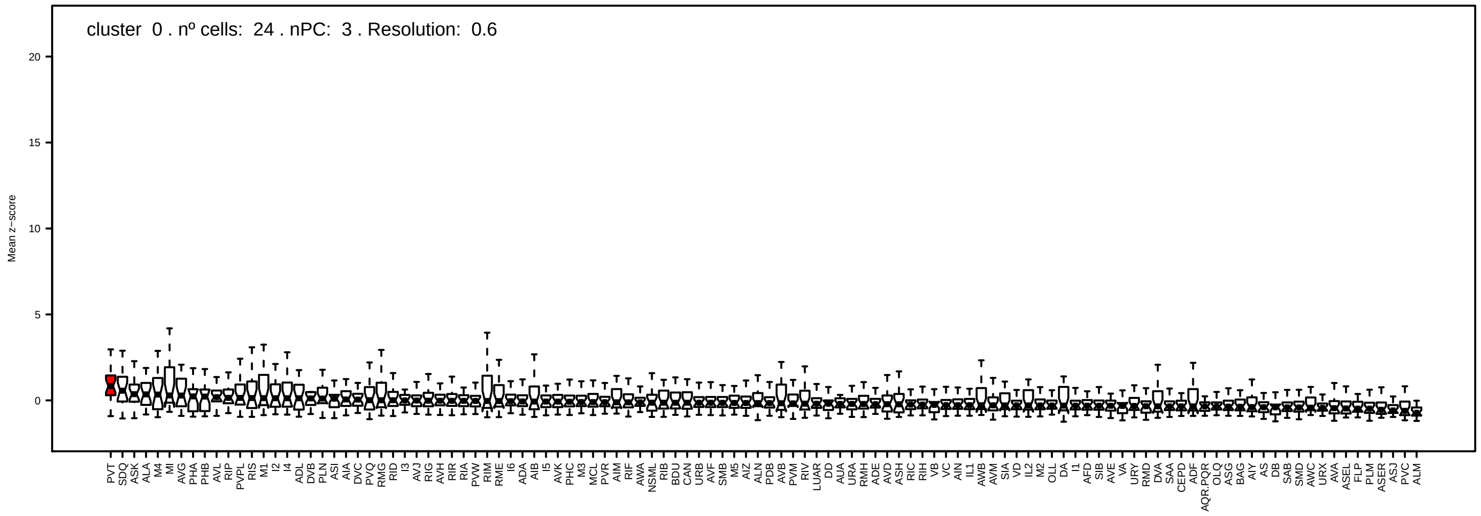
cluster 0 . nº cells: 24 . nPC: 3 . Resolution: 0.6
Mean z−score
0
5
10
15
20
PVT SDQ ASK ALA M4 MI AVG PHA PHB AVL RIP PVPL RIS M1 I2 I4 ADL DVB PLN ASI AIA DVC PVQ RMG RID I3 AVJ RIG AVH RIR RIA PVW RIM RME I6 ADA AIB I5 AVK PHC M3 MCL PVR AIM RIF AWA NSML RIB BDU CAN URB AVF SMB M5 AIZ ALN PDB AVB PVM RIV LUAR DD AUA URA RMH ADE AVD ASH RIC RIH VB VC AIN IL1 AWB AVM SIA VD IL2 M2 OLL DA I1 AFD SIB AVE VA URY RMD DVA SAA CEPD ADF AQR.PQR OLQ ASG BAG AIY AS DB SAB SMD AWC URX AVA ASEL FLP PLM ASER ASJ PVC ALM

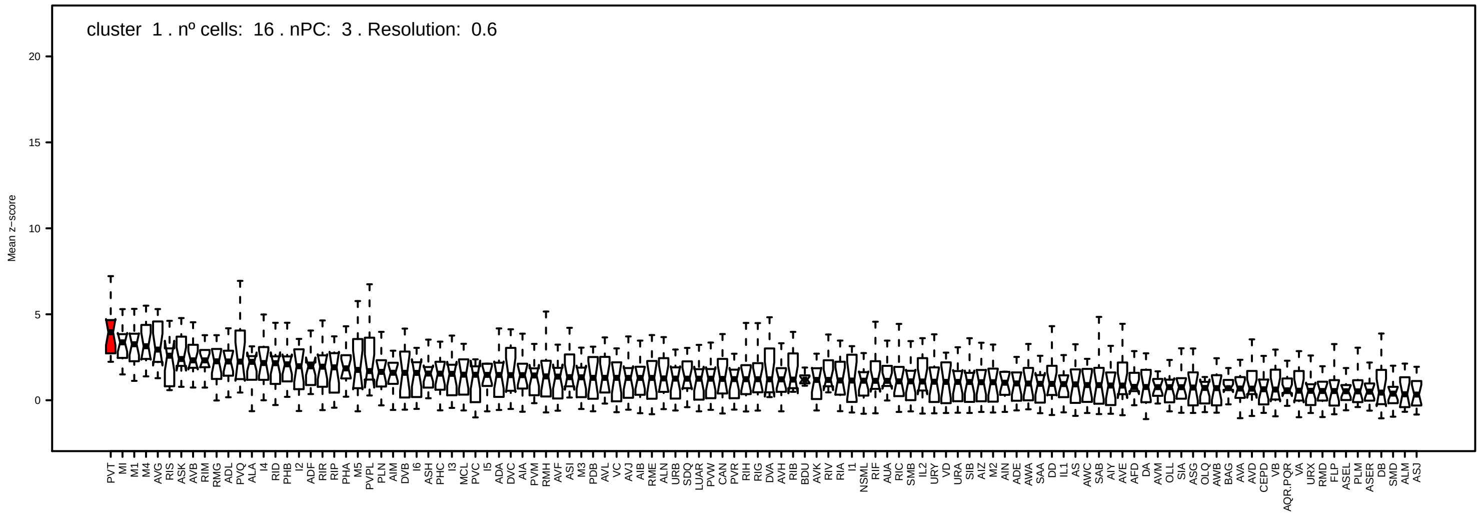
cluster 1 . nº cells: 16 . nPC: 3 . Resolution: 0.6
Mean z−score
0
5
10
15
20
PVT MI M1 M4 AVG RIS ASK AVB RIM RMG ADL PVQ ALA I4 RID PHB I2 ADF RIR RIP PHA M5 PVPL PLN AIM DVB I6 ASH PHC I3 MCL PVC I5 ADA DVC AIA PVM RMH AVF ASI M3 PDB AVL VC AVJ AIB RME ALN URB SDQ LUAR PVW CAN PVR RIH RIG DVA AVH RIB BDU AVK RIV RIA I1 NSML RIF AUA RIC SMB IL2 URY VD URA SIB AIZ M2 AIN ADE AWA SAA DD IL1 AS AWC SAB AIY AVE AFD DA AVM OLL SIA ASG OLQ AWB BAG AVA AVD CEPD VB AQR.PQR VA URX RMD FLP ASEL PLM ASER DB SMD ALM ASJ

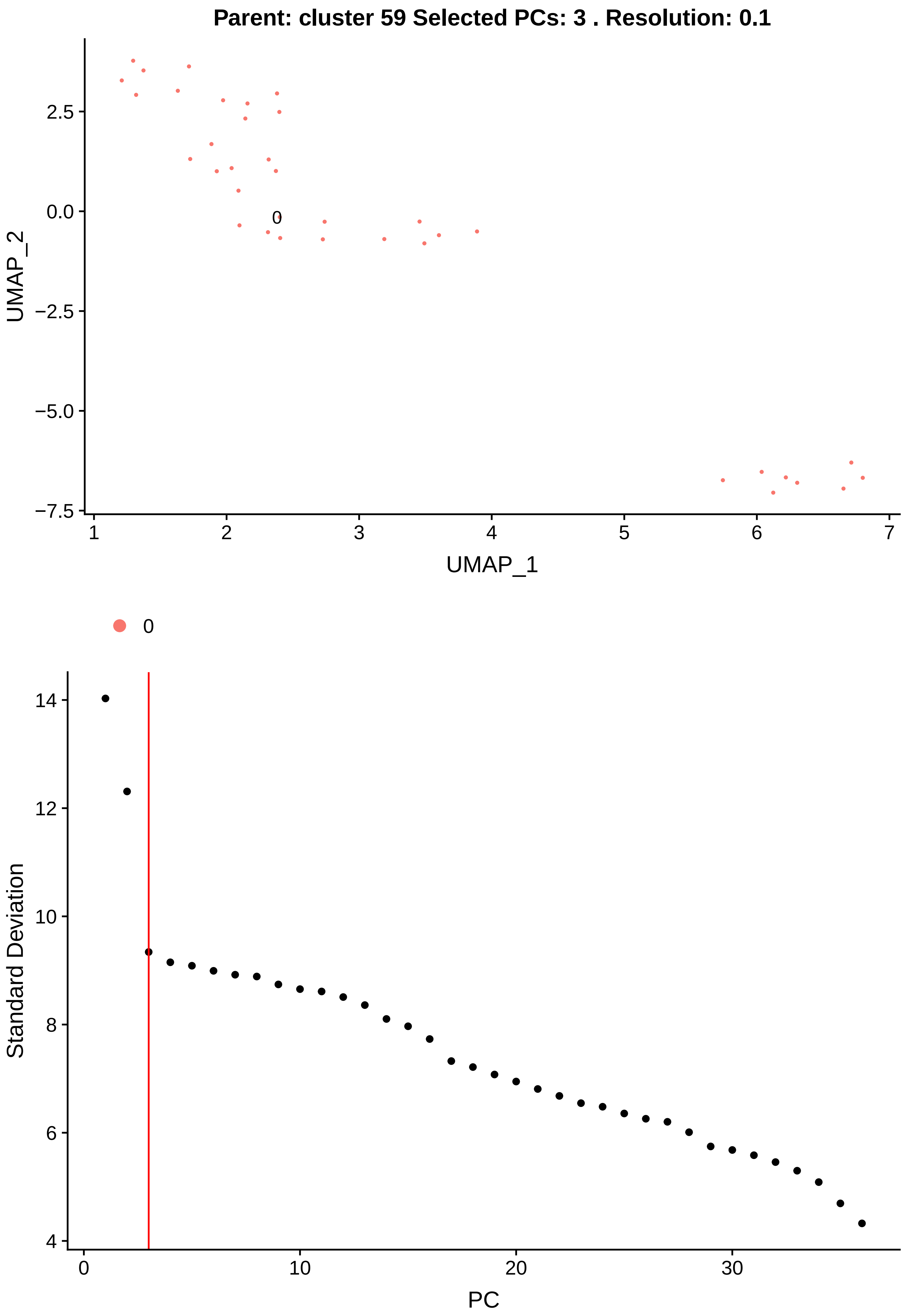

Parent: cluster 59 Selected PCs: 3 . Resolution: 0.1
UMAP_2
2.5
0.0
−2.5
−5.0
−7.5
0
1
2
3
4
5
6
7
UMAP_1
0
Standard Deviation
14
12
10
8
6
4
0
10
20
30
PC

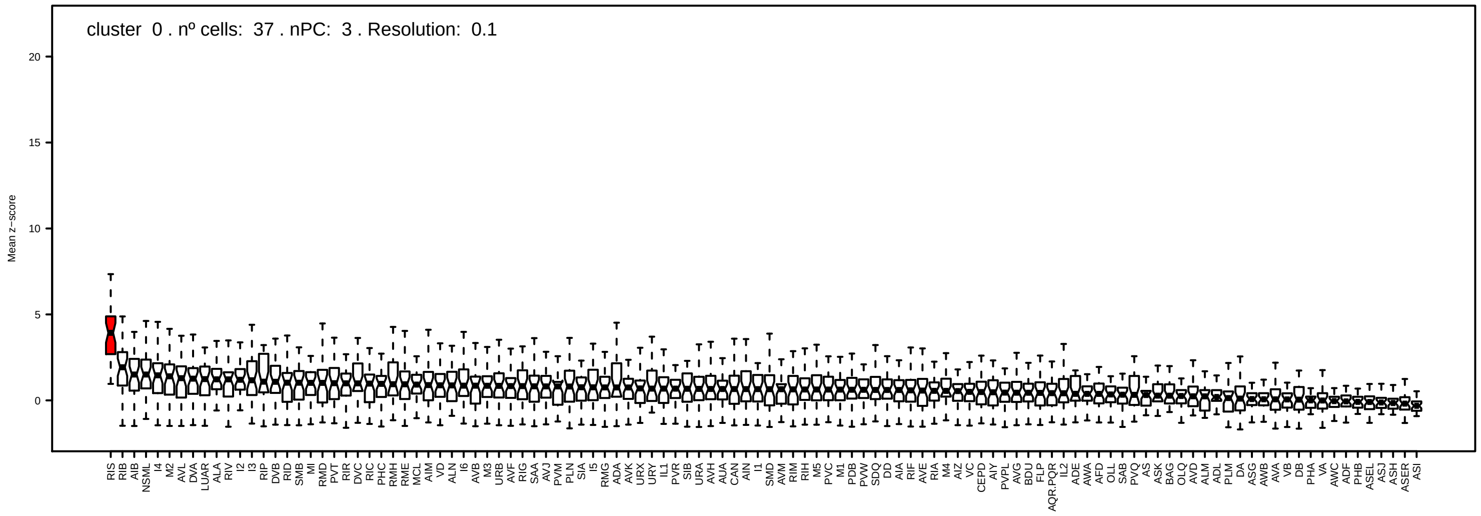

cluster 0 . nº cells: 37 . nPC: 3 . Resolution: 0.1
Mean z−score
20
15
10
5
0
RIS RIB AIB NSML I4 M2 AVL DVA LUAR ALA RIV I2 I3 RIP DVB RID SMB MI RMD PVT RIR DVC RIC PHC RMH RME MCL AIM VD ALN I6 AVB M3 URB AVF RIG SAA AVJ PVM PLN SIA I5 RMG ADA AVK URX URY IL1 PVR SIB URA AVH AUA CAN AIN I1 SMD AVM RIM RIH M5 PVC M1 PDB PVW SDQ DD AIA RIF AVE RIA M4 AIZ VC CEPD AIY PVPL AVG BDU FLP AQR.PQR IL2 ADE AWA AFD OLL SAB PVQ AS ASK BAG OLQ AVD ALM ADL PLM DA ASG AWB AVA VB DB PHA VA AWC ADF PHB ASEL ASJ ASH ASER ASI

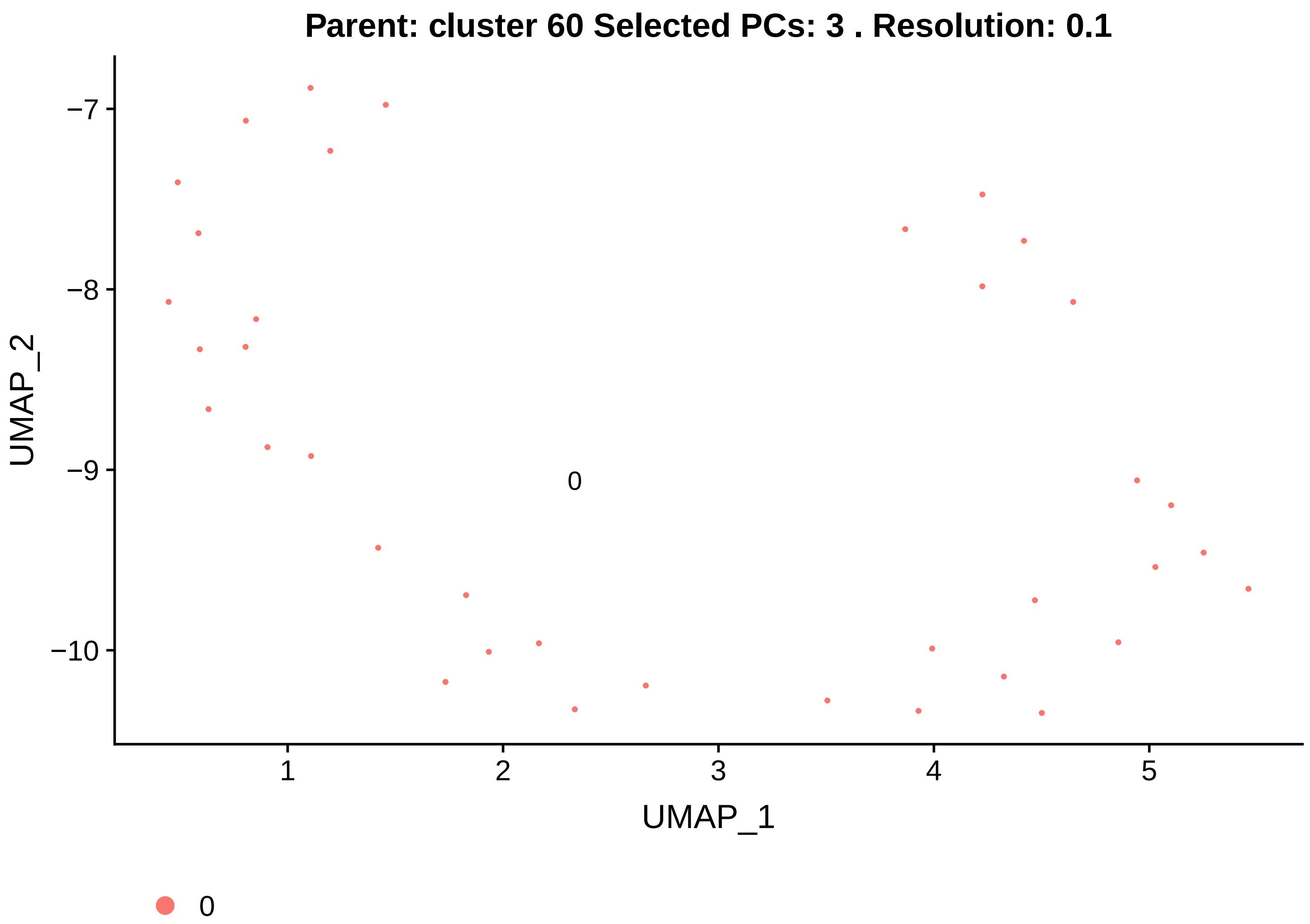
Parent: cluster 60 Selected PCs: 3 . Resolution: 0.1
−7
−8
−9
−10
UMAP_2
0
1
2
3
4
5
UMAP_1
0

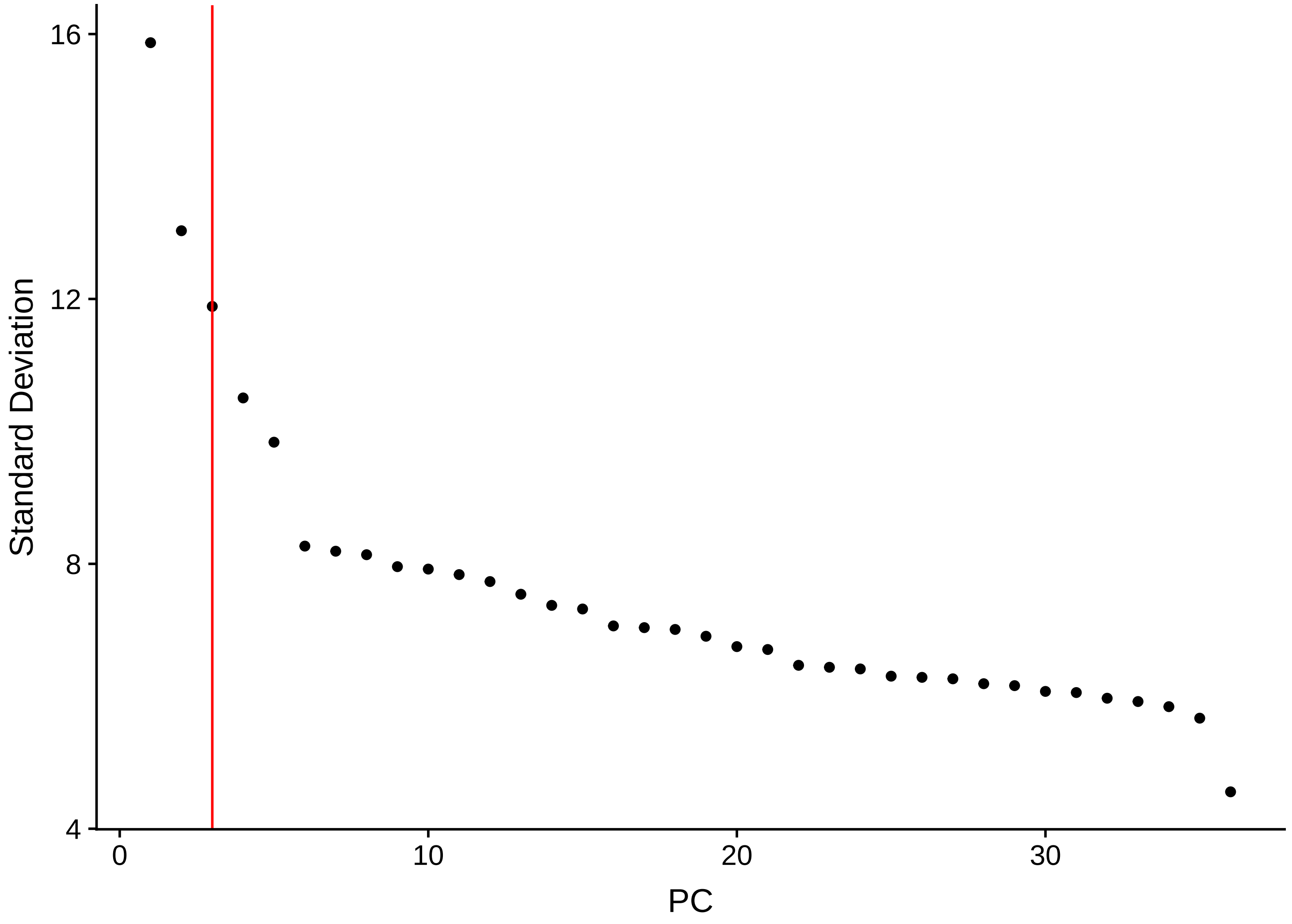
16
12
8
4
Standard Deviation
0
10
20
30
PC

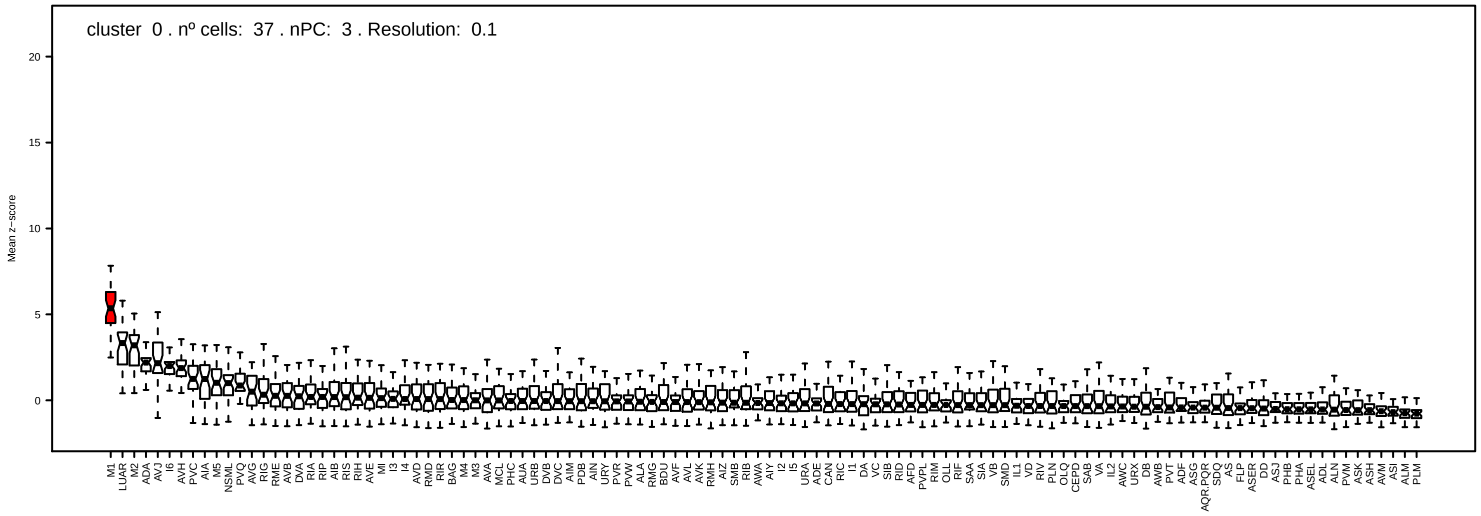
cluster 0 . nº cells: 37 . nPC: 3 . Resolution: 0.1
Mean z−score

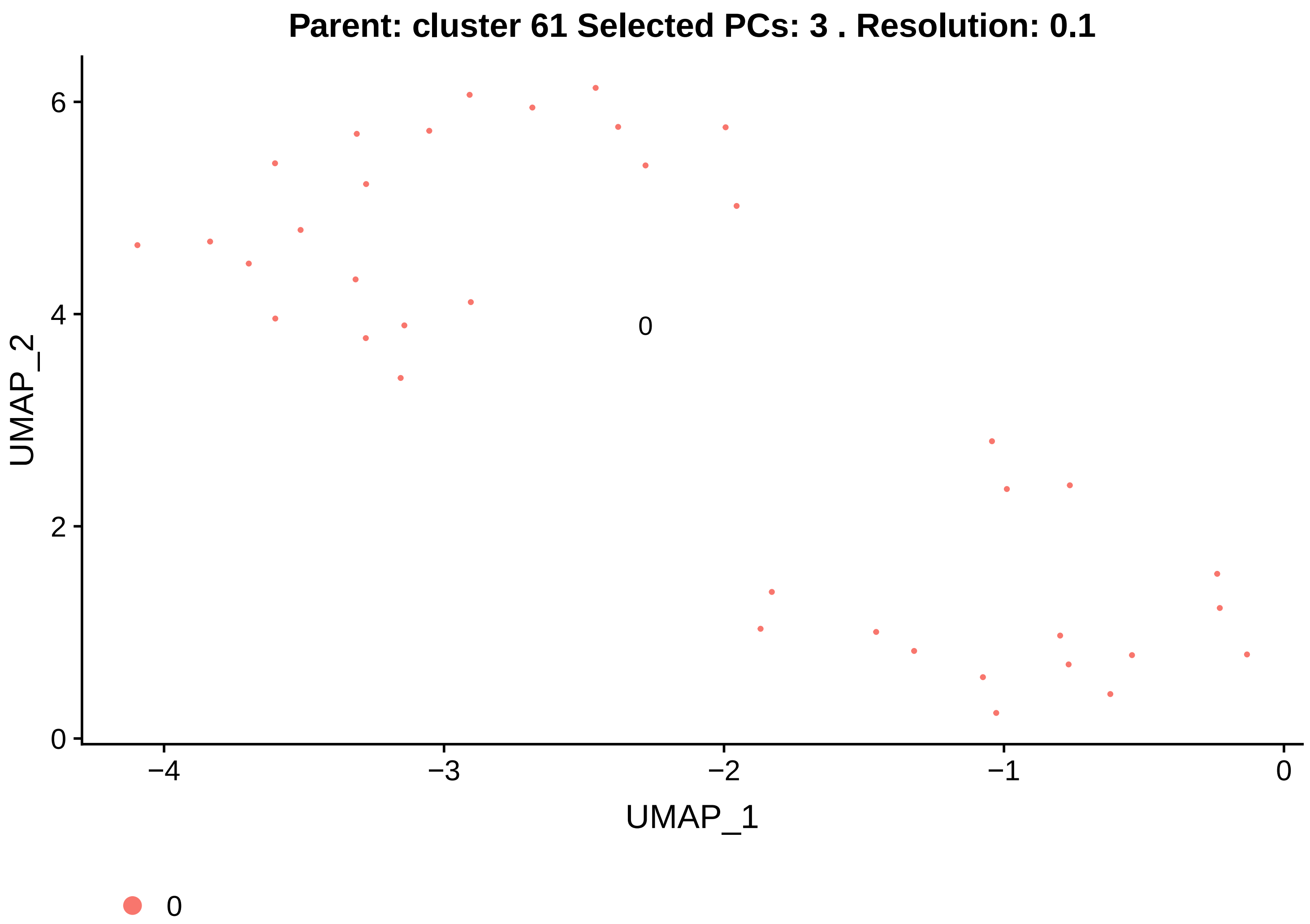
Parent: cluster 61 Selected PCs: 3 . Resolution: 0.1
UMAP_2
UMAP_1
0
−4
−3
−2
−1
0
2
4
6
0

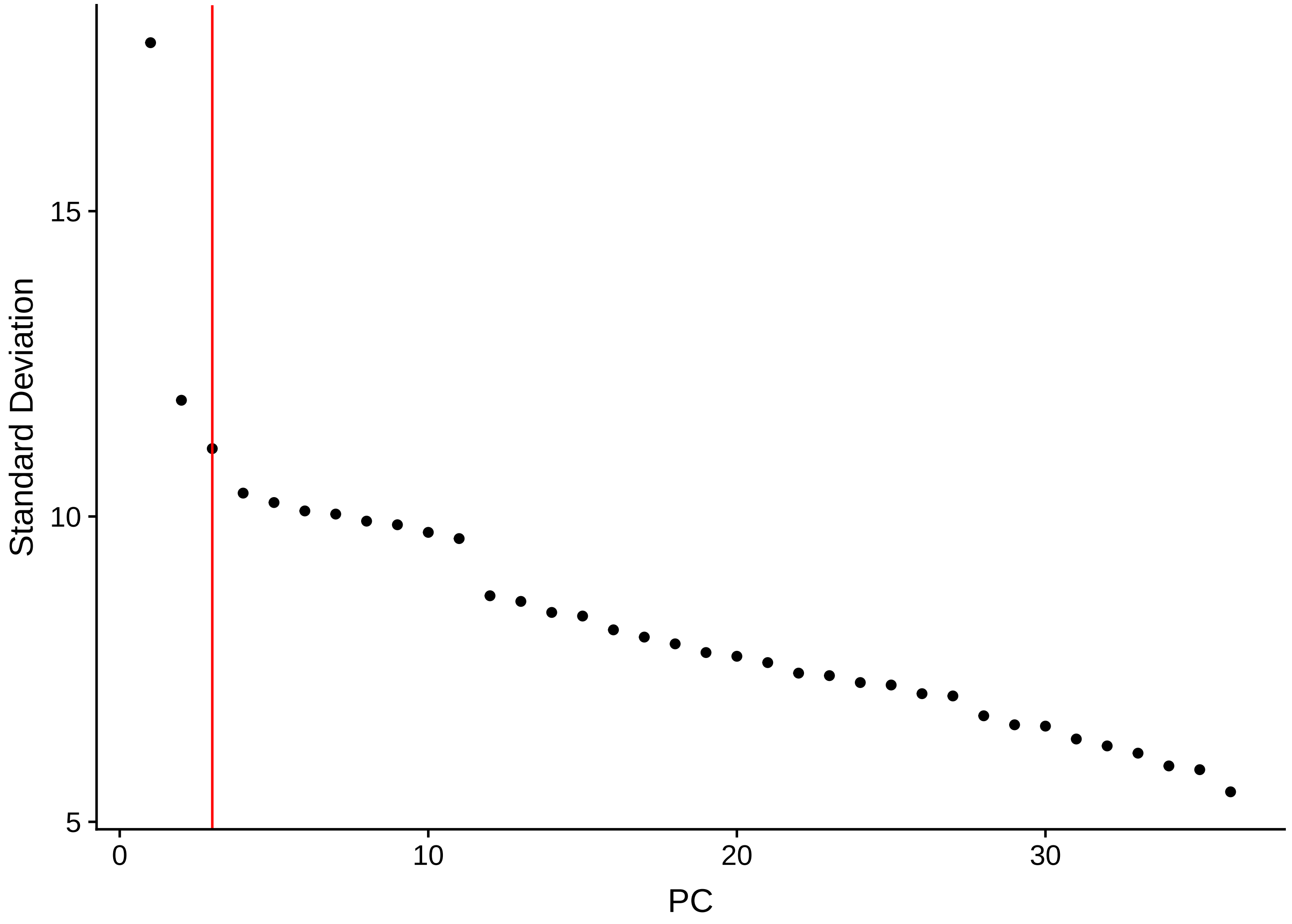
Standard Deviation
PC
5
10
15
0
10
20
30

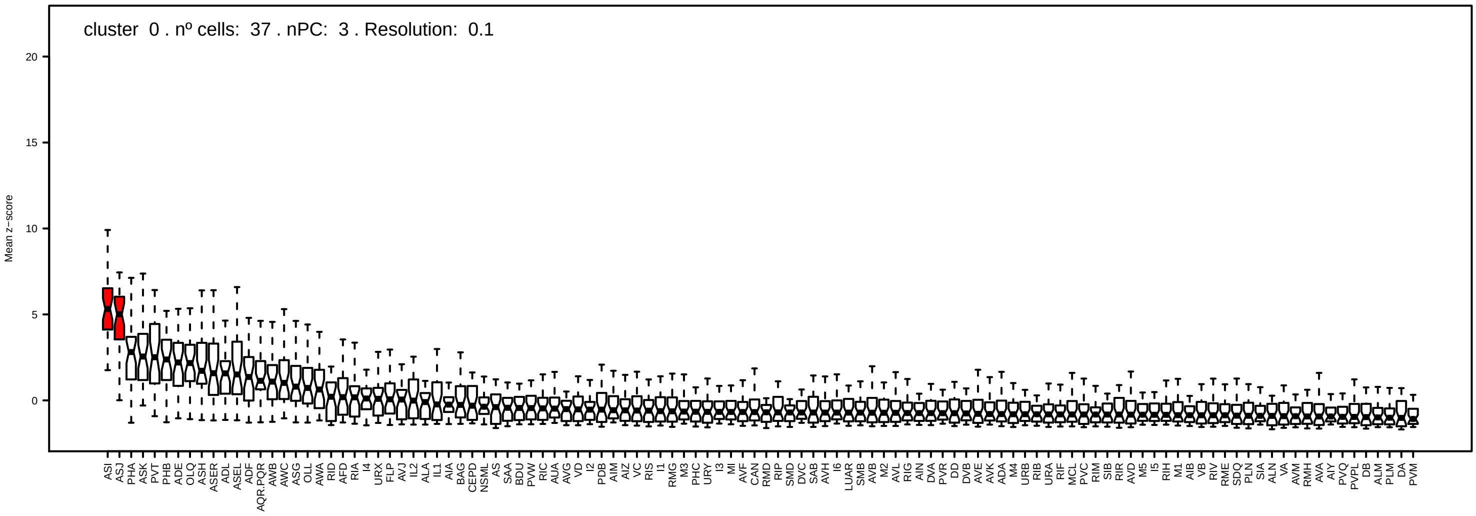
cluster 0 . nº cells: 37 . nPC: 3 . Resolution: 0.1
Mean z−score
20
15
10
5
0
ASI
ASJ
PHA
ASK
PVT
PHB
ADE
OLQ
ASH
ASER
ADL
ASEL
ADF
AQR.PQR
AWB
AWC
ASG
OLL
AWA
RID
AFD
RIA
I4
URX
FLP
AVJ
IL2
ALA
IL1
AIA
BAG
CEPD
NSML
AS
SAA
BDU
PVW
RIC
AUA
AVG
VD
I2
PDB
AIM
AIZ
VC
RIS
I1
RMG
M3
PHC
URY
I3
MI
AVF
CAN
RMD
RIP
SMD
DVC
SAB
AVH
I6
LUAR
SMB
AVB
M2
AVL
RIG
AIN
DVA
PVR
DD
DVB
AVE
AVK
ADA
M4
URB
RIB
URA
RIF
MCL
PVC
RIM
SIB
RIR
AVD
M5
I5
RIH
M1
AIB
VB
RIV
RME
SDQ
PLN
SIA
ALN
VA
AVM
RMH
AVA
AIY
PVQ
PVPL
DB
ALM
PLM
DA
PVM

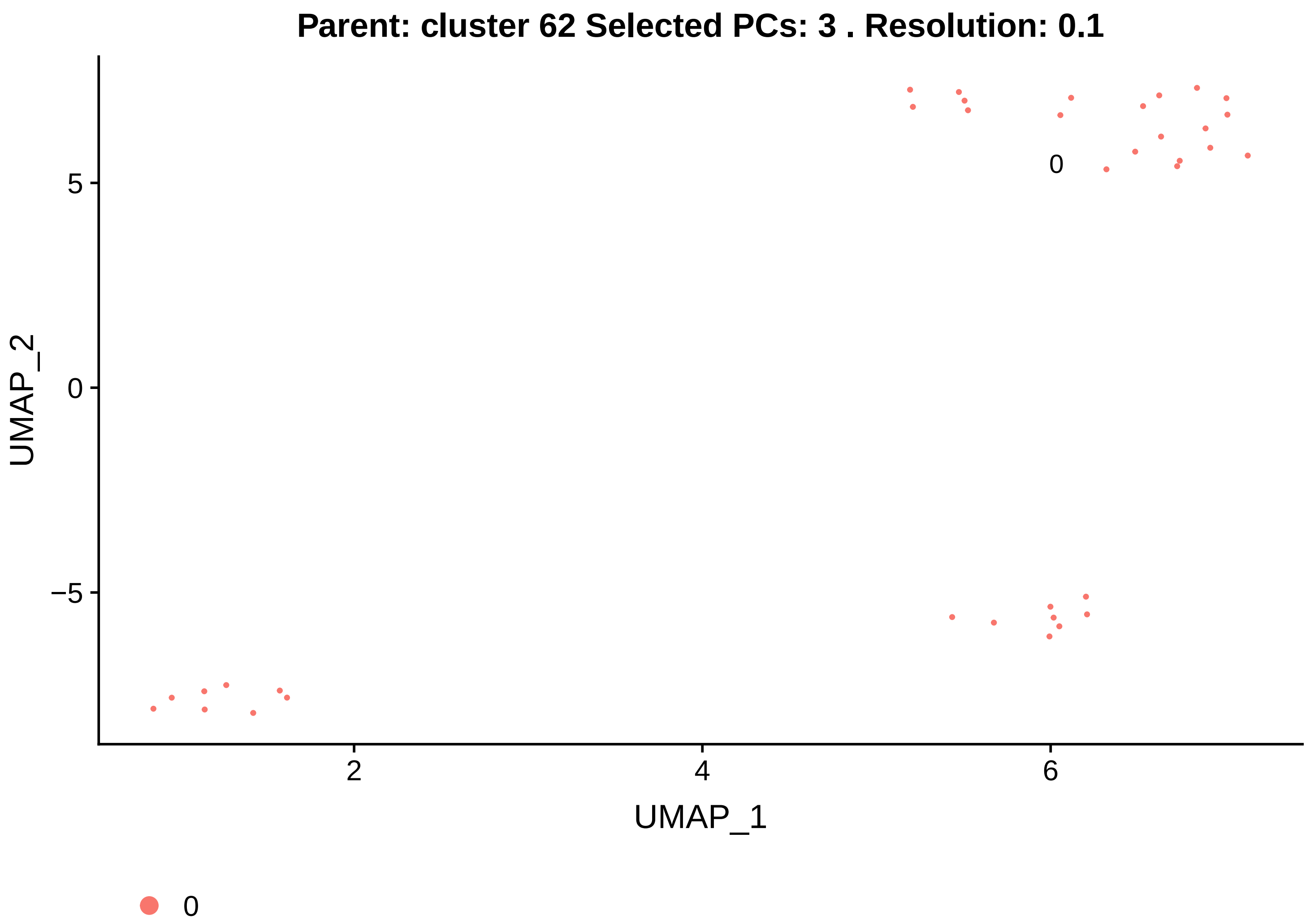
Parent: cluster 62 Selected PCs: 3 . Resolution: 0.1
UMAP_2
UMAP_1
0
5
0
−5
2
4
6
0

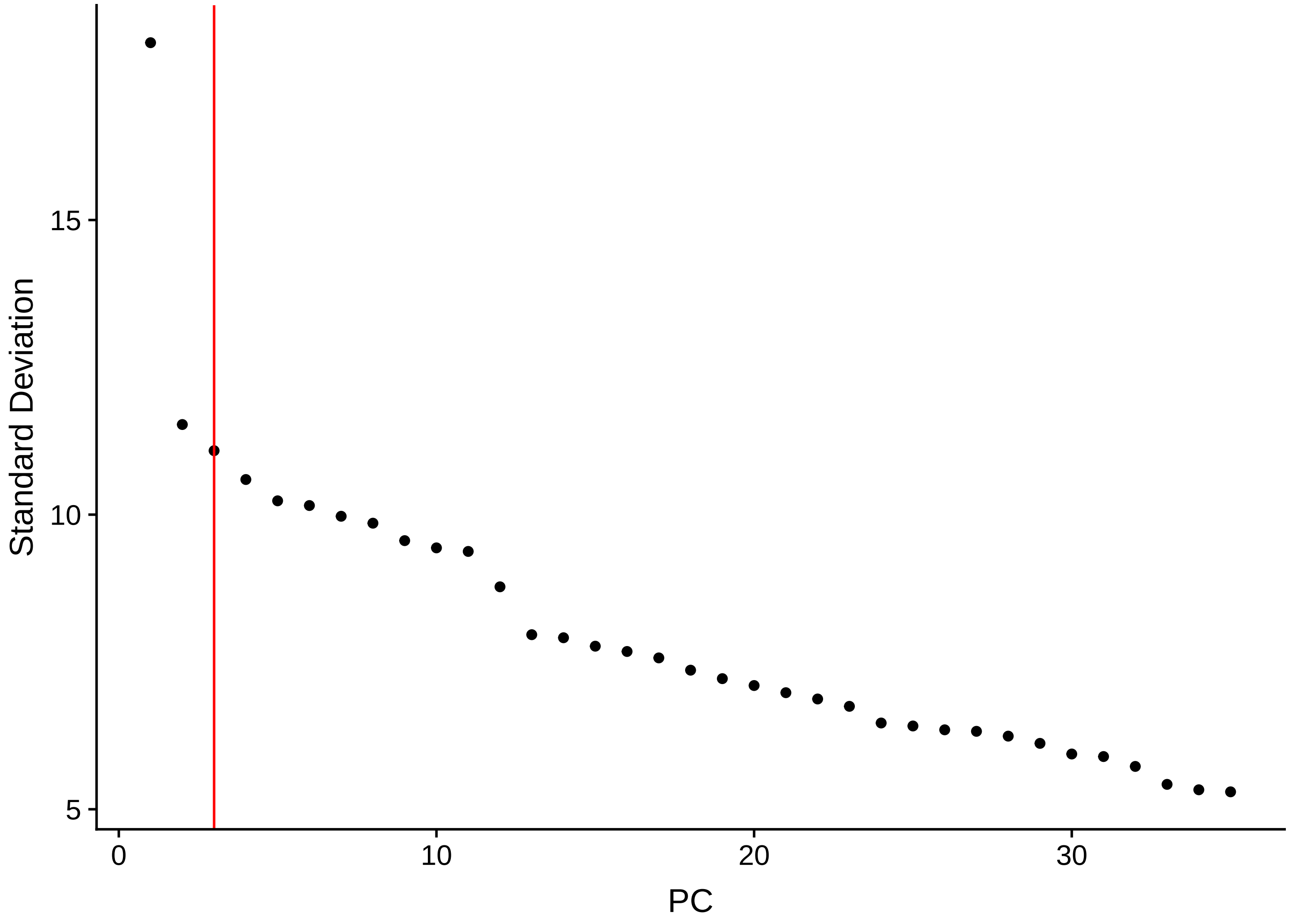
Standard Deviation
PC
15
10
5
0
10
20
30

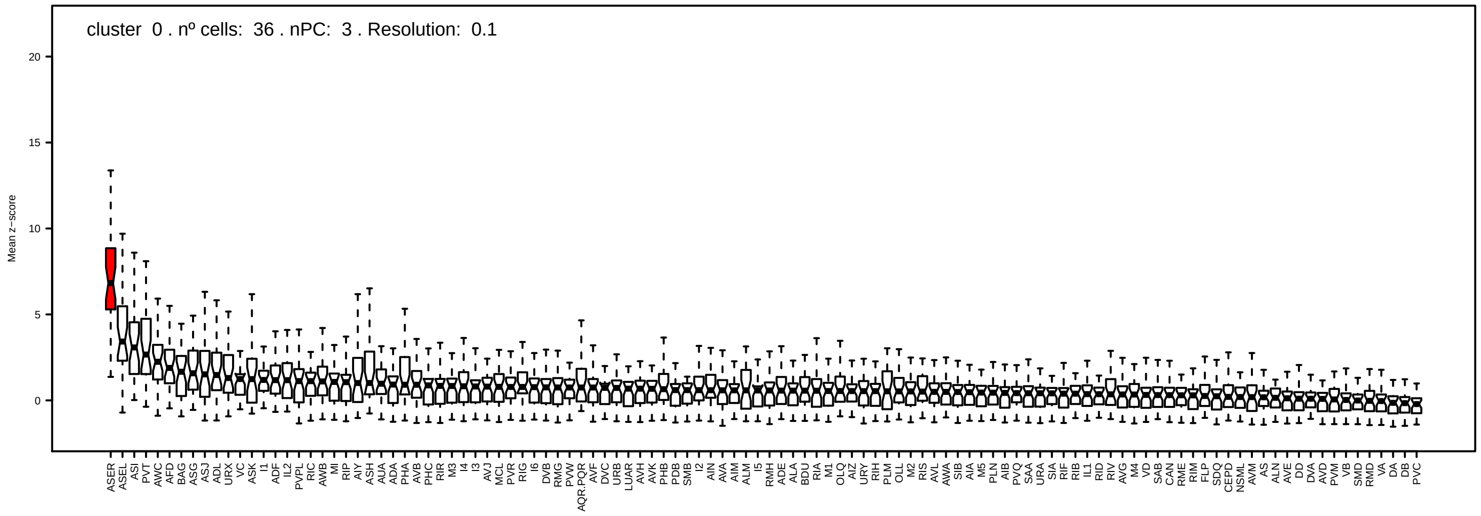

cluster 0 . nº cells: 36 . nPC: 3 . Resolution: 0.1
Mean z−score
20
15
10
5
0
ASER ASEL ASI PVT AWC AFD BAG ASG ASJ ADL URX VC ASK I1 ADF IL2 PVPL RIC AWB MI RIP AIY ASH AUA ADA PHA AVB PHC RIR M3 I4 I3 AVJ MCL PVR RIG I6 DVB RMG PVW AQR.PQR AVF DVC URB LUAR AVH AVK PHB PDB SMB I2 AIN AVA AIM ALM I5 RMH ADE ALA BDU RIA M1 OLQ AIZ URY RIH PLM OLL M2 RIS AVL AWA SIB AIA M5 PLN AIB PVQ SAA URA SIA RIF RIB IL1 RID RIV AVG M4 VD SAB CAN RME RIM FLP SDQ CEPD NSML AVM AS ALN AVE DD DVA AVD PVM VB SMD RMD VA DA DB PVC

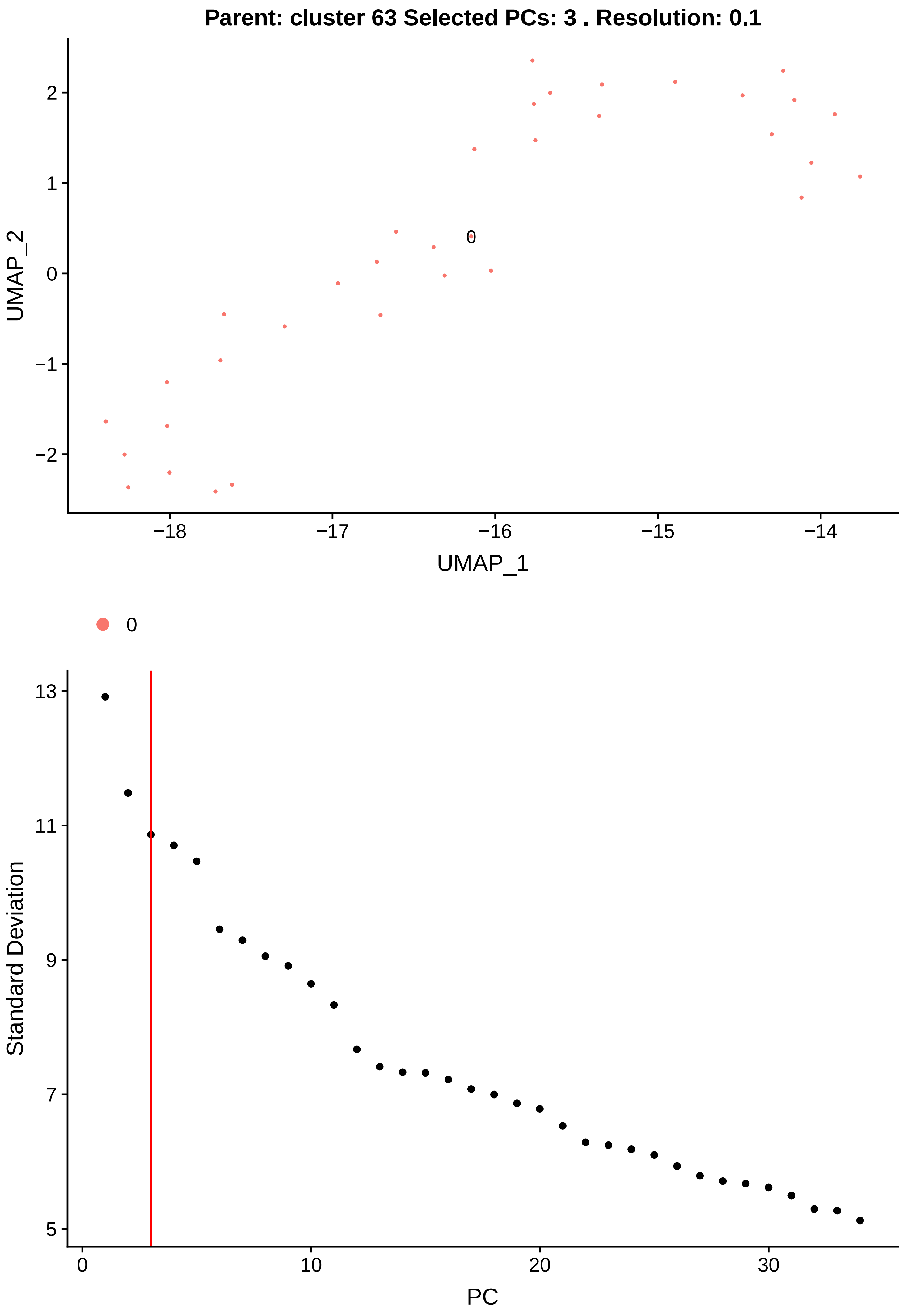
Parent: cluster 63 Selected PCs: 3 . Resolution: 0.1
UMAP_2
UMAP_1
0
Standard Deviation
PC

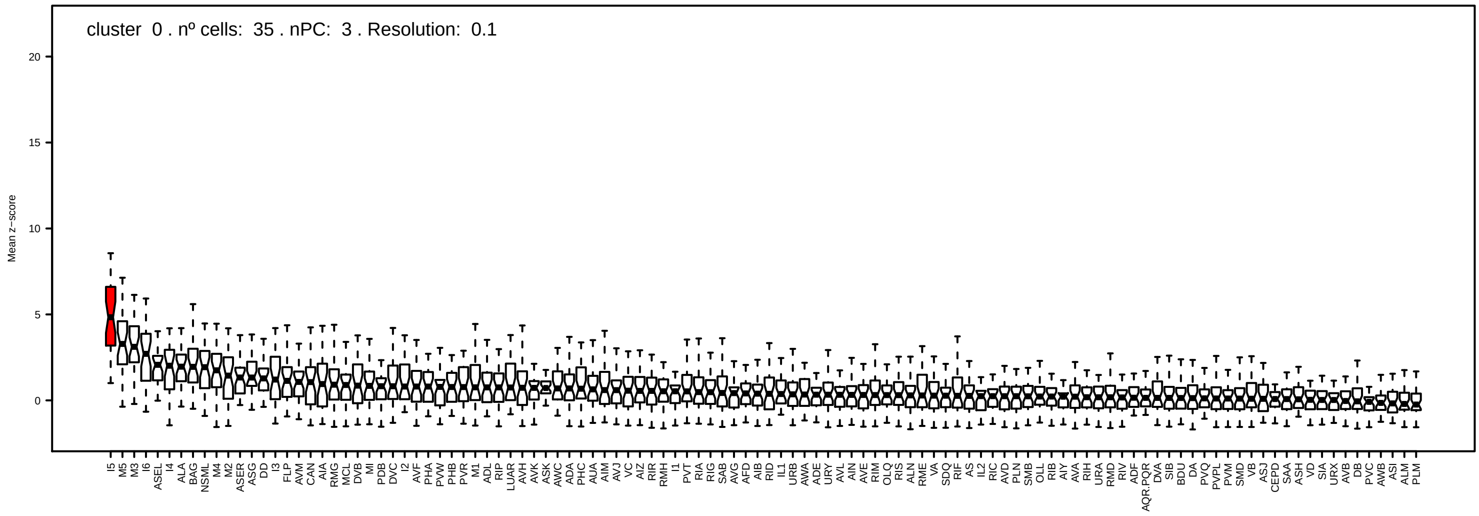
cluster 0 . nº cells: 35 . nPC: 3 . Resolution: 0.1
Mean z−score
20
15
10
5
0
I5 M5 M3 I6 ASEL I4 ALA BAG NSML M4 M2 ASER ASG DD I3 FLP AVM CAN AIA RMG MCL DVB MI PDB DVC I2 AVF PHA PVW PHB PVR M1 ADL RIP LUAR AVH AVK ASK AWC ADA PHC AUA AIM AVJ VC AIZ RIR RMH I1 PVT RIA RIG SAB AVG AFD AIB RID IL1 URB AWA ADE URY AVL AIN AVE RIM OLQ RIS ALN RME VA SDQ RIF AS IL2 RIC AVD PLN SMB OLL RIB AIY AVA RIH URA RMD RIV ADF AQR.PQR DVA SIB BDU DA PVQ PVPL PVM SMD VB ASJ CEPD SAA ASH VD SIA URX AVB DB PVC AWB ASI ALM PLM
