## Supplementary Folder 2 for "Combining single-cell RNA-sequencing with a molecular atlas unveils new markers for *C. elegans* neuron classes"

# 0.0 vs DA

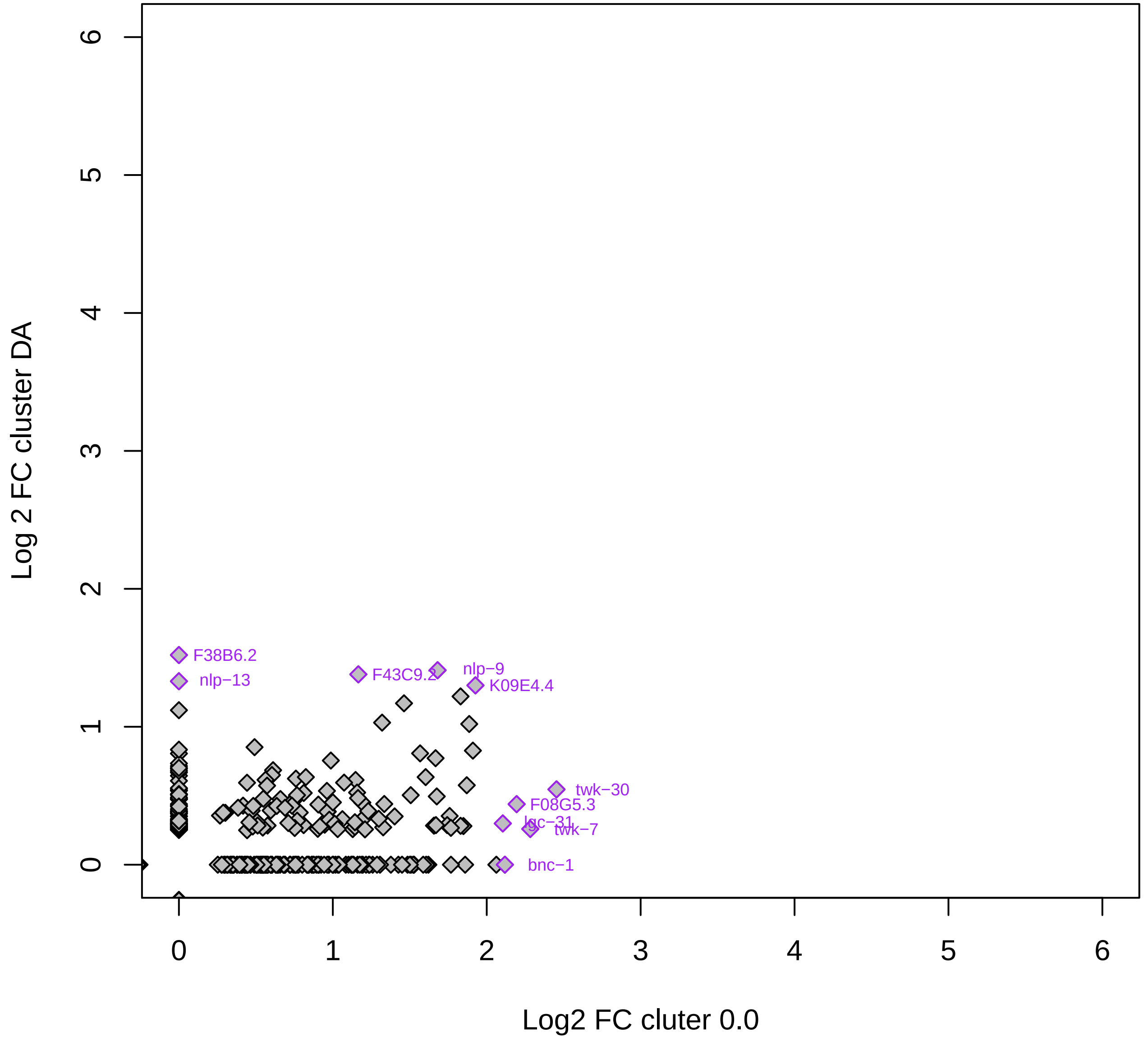

Log 2 FC cluster DA
Log2 FC cluter 0.0
0
1
2
3
4
5
6
F38B6.2
nlp−13
F43C9.2
nlp−9
K09E4.4
twk−30
F08G5.3
lgc−31
twk−7
bnc−1

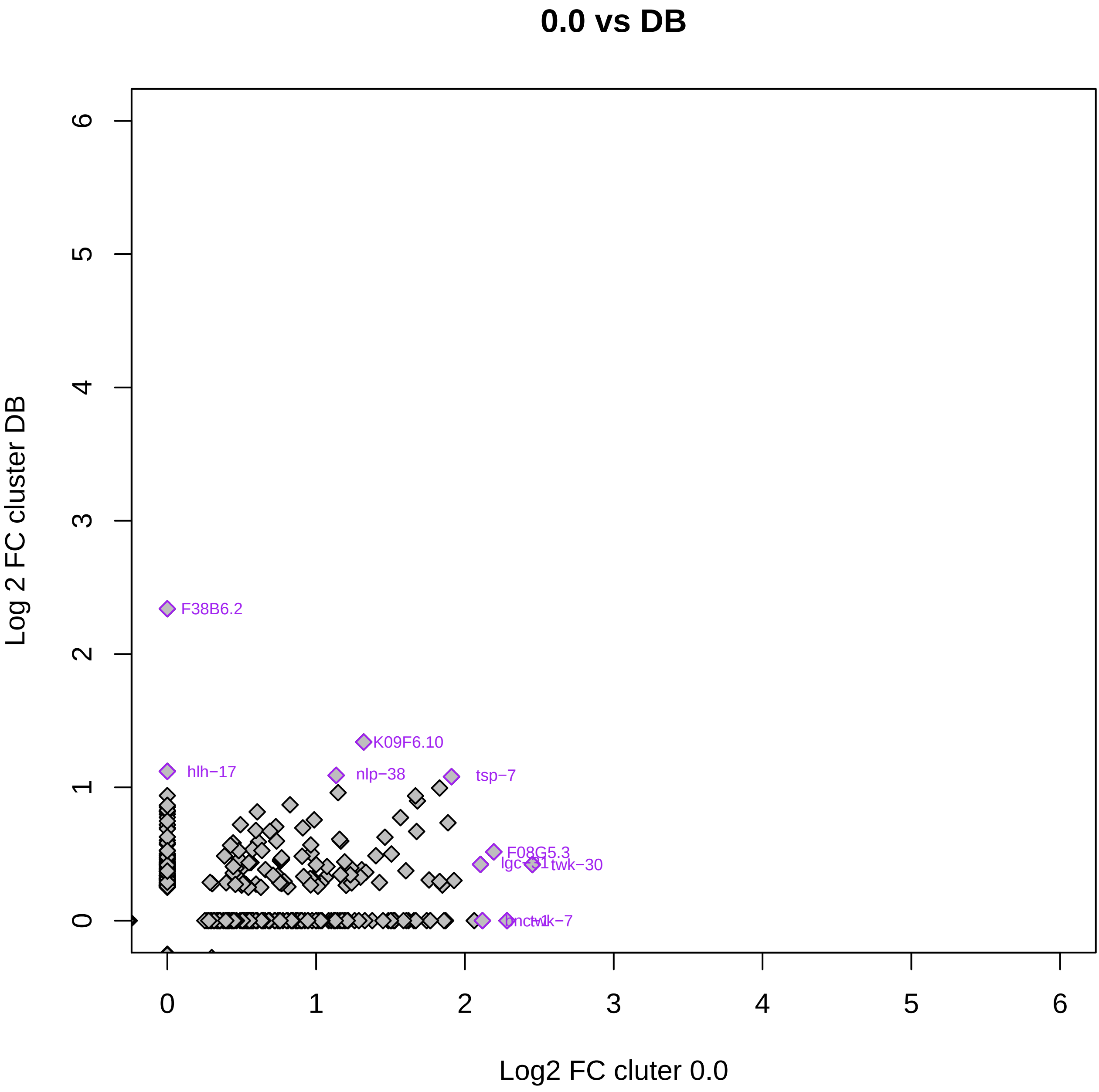
0.0 vs DB
Log 2 FC cluster DB
Log2 FC cluter 0.0
F38B6.2
K09F6.10
hlh−17
nlp−38
tsp−7
F08G5.3
lgc−31
twk−30
twk−7
0
1
2
3
4
5
6

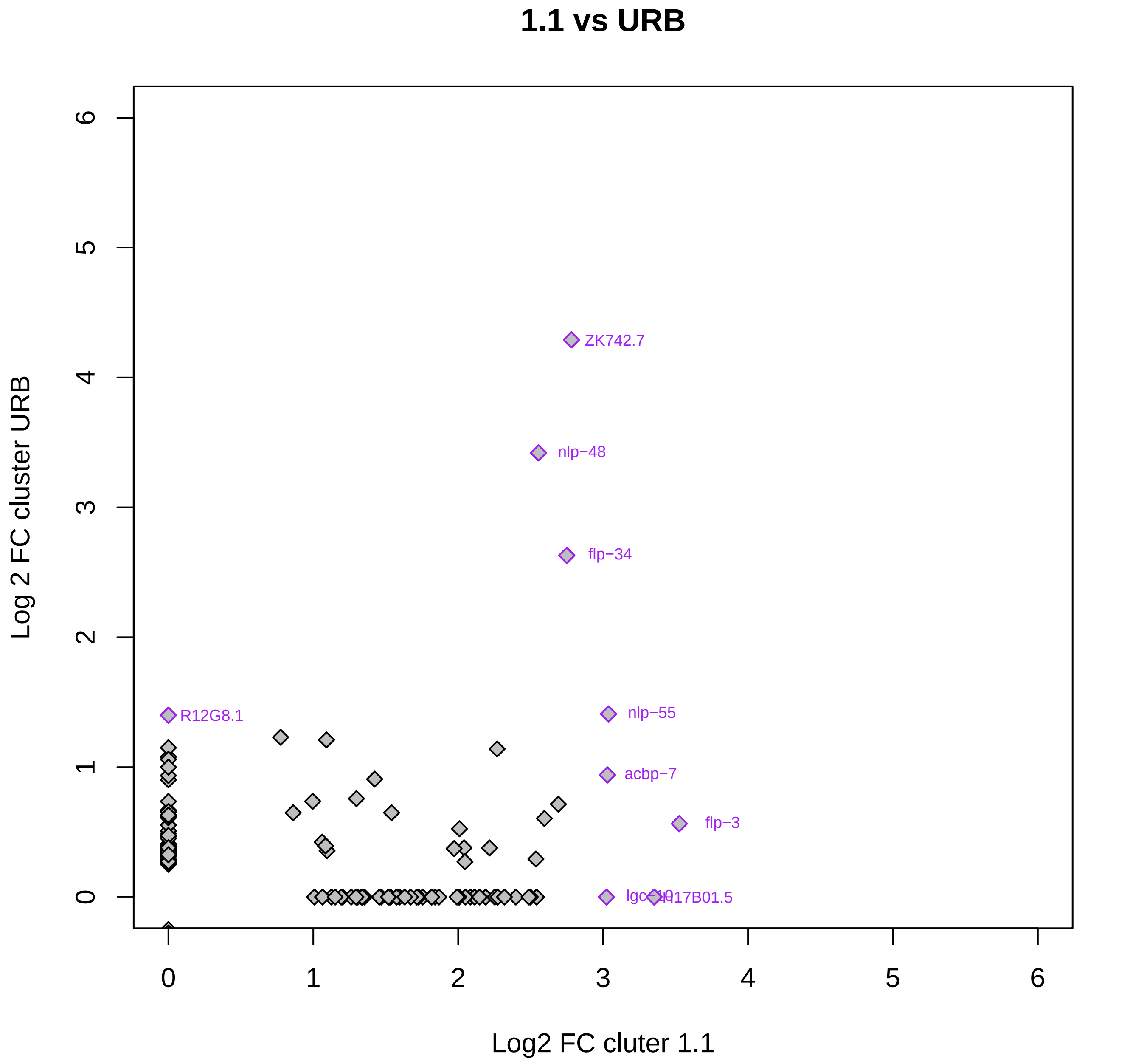
1.1 vs URB
Log 2 FC cluster URB
Log2 FC cluter 1.1
0
1
2
3
4
5
6
ZK742.7
nlp−48
flp−34
R12G8.1
nlp−55
acbp−7
flp−3
lgc−10
H17B01.5

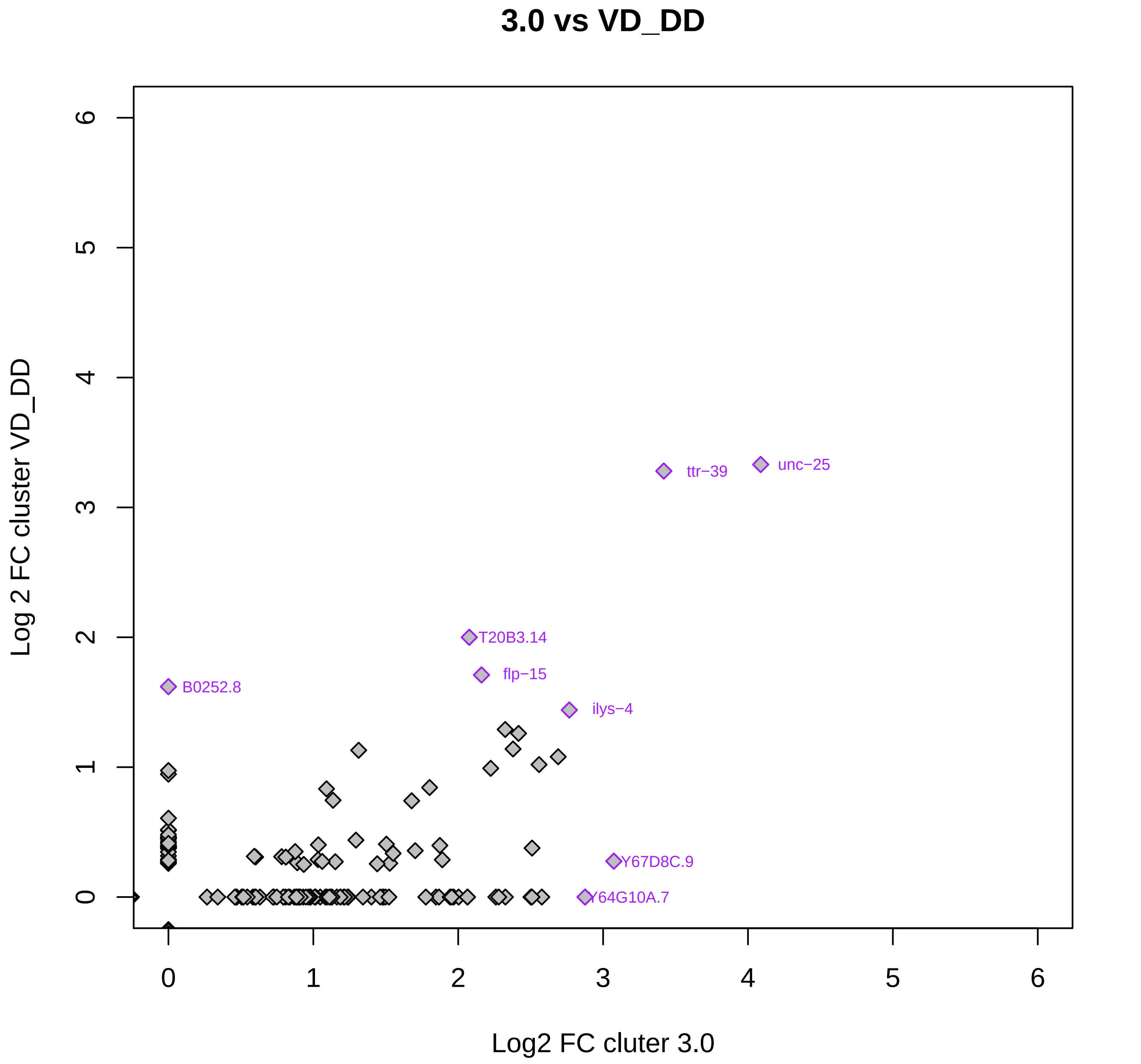

3.0 vs VD_DD
Log 2 FC cluster VD_DD
Log2 FC cluter 3.0
0
1
2
3
4
5
6
ttr−39
unc−25
T20B3.14
flp−15
B0252.8
ilys−4
Y67D8C.9
Y64G10A.7

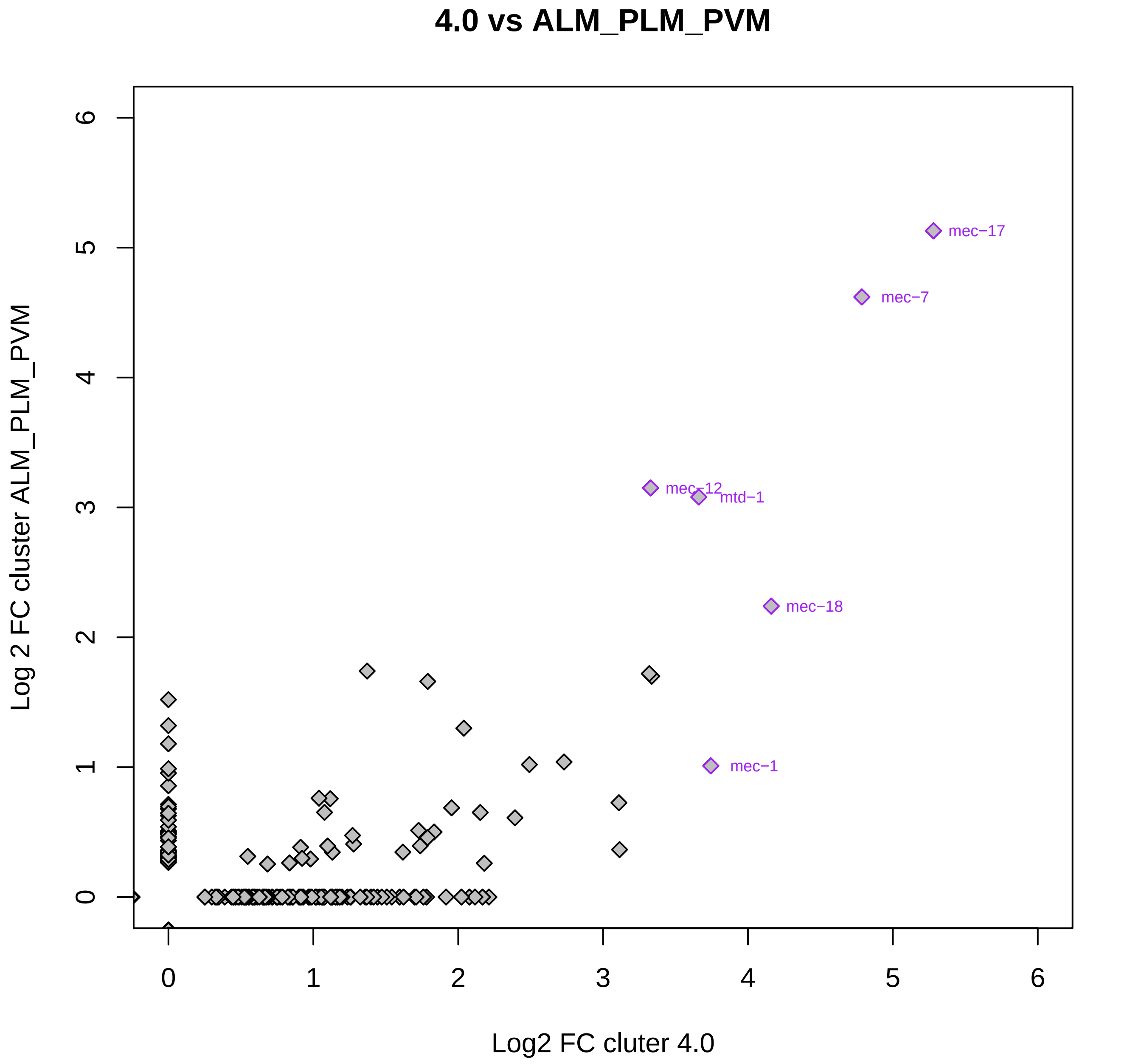

4.0 vs ALM_PLM_PVM
Log 2 FC cluster ALM_PLM_PVM
Log2 FC cluter 4.0
mec−17
mec−7
mec−12
mtd−1
mec−18
mec−1
0
1
2
3
4
5
6

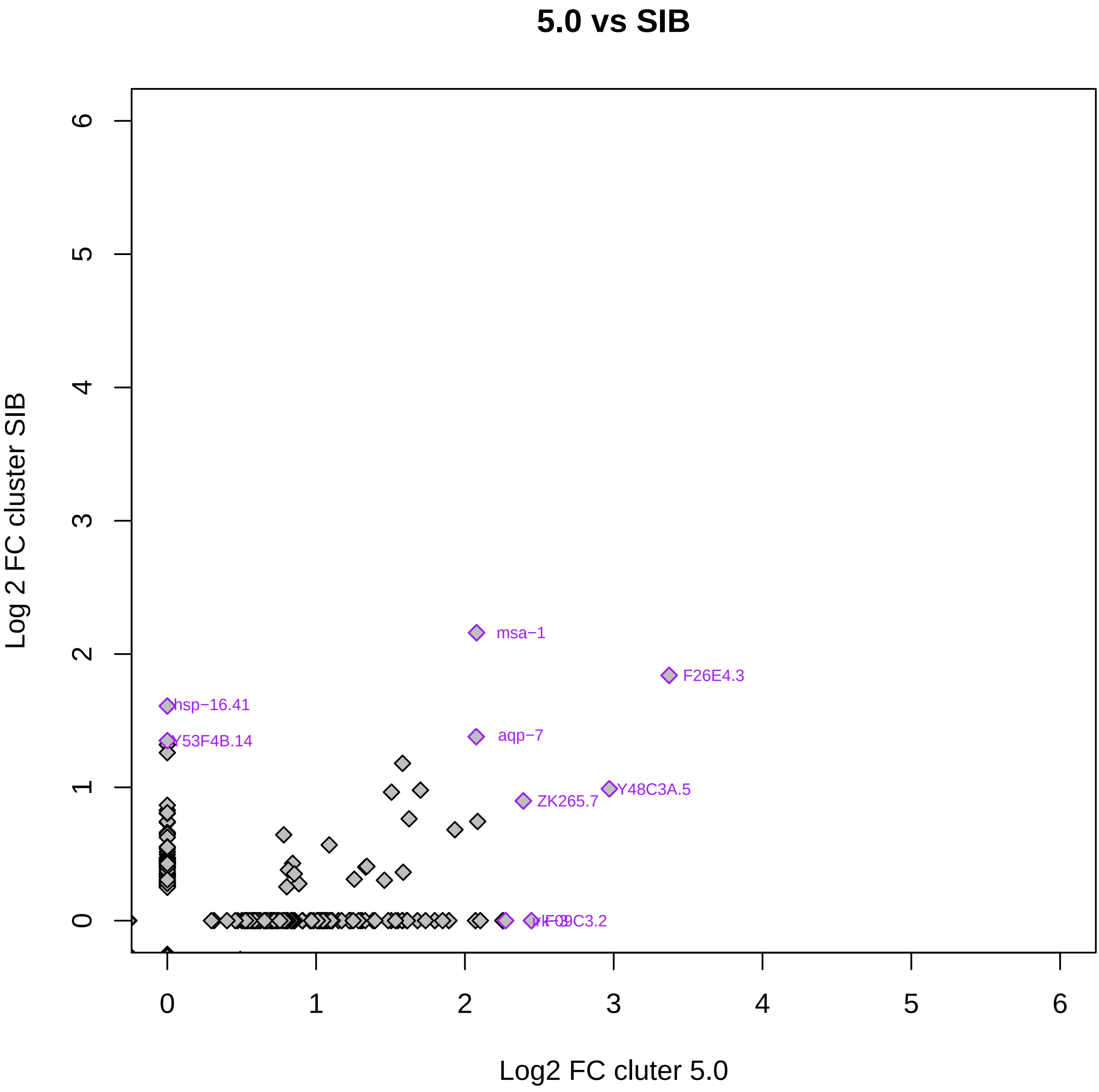

5.0 vs SIB
Log 2 FC cluster SIB
Log2 FC cluter 5.0
0
1
2
3
4
5
6
msa−1
F26E4.3
hsp−16.41
Y53F4B.14
aqp−7
Y48C3A.5
ZK265.7

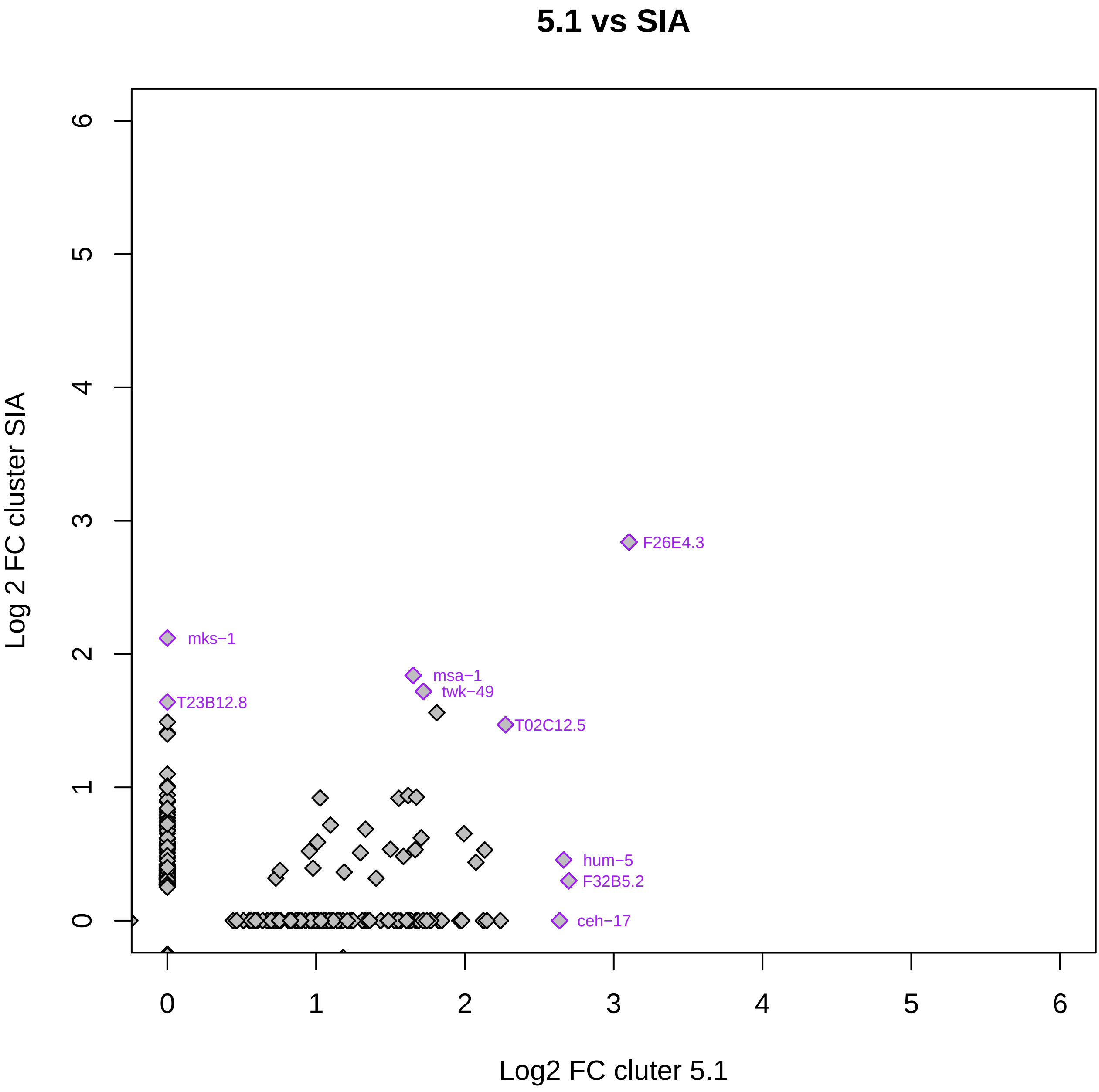
5.1 vs SIA
Log 2 FC cluster SIA
Log2 FC cluter 5.1
0
1
2
3
4
5
6
F26E4.3
mks−1
msa−1
twk−49
T23B12.8
T02C12.5
hum−5
F32B5.2
ceh−17

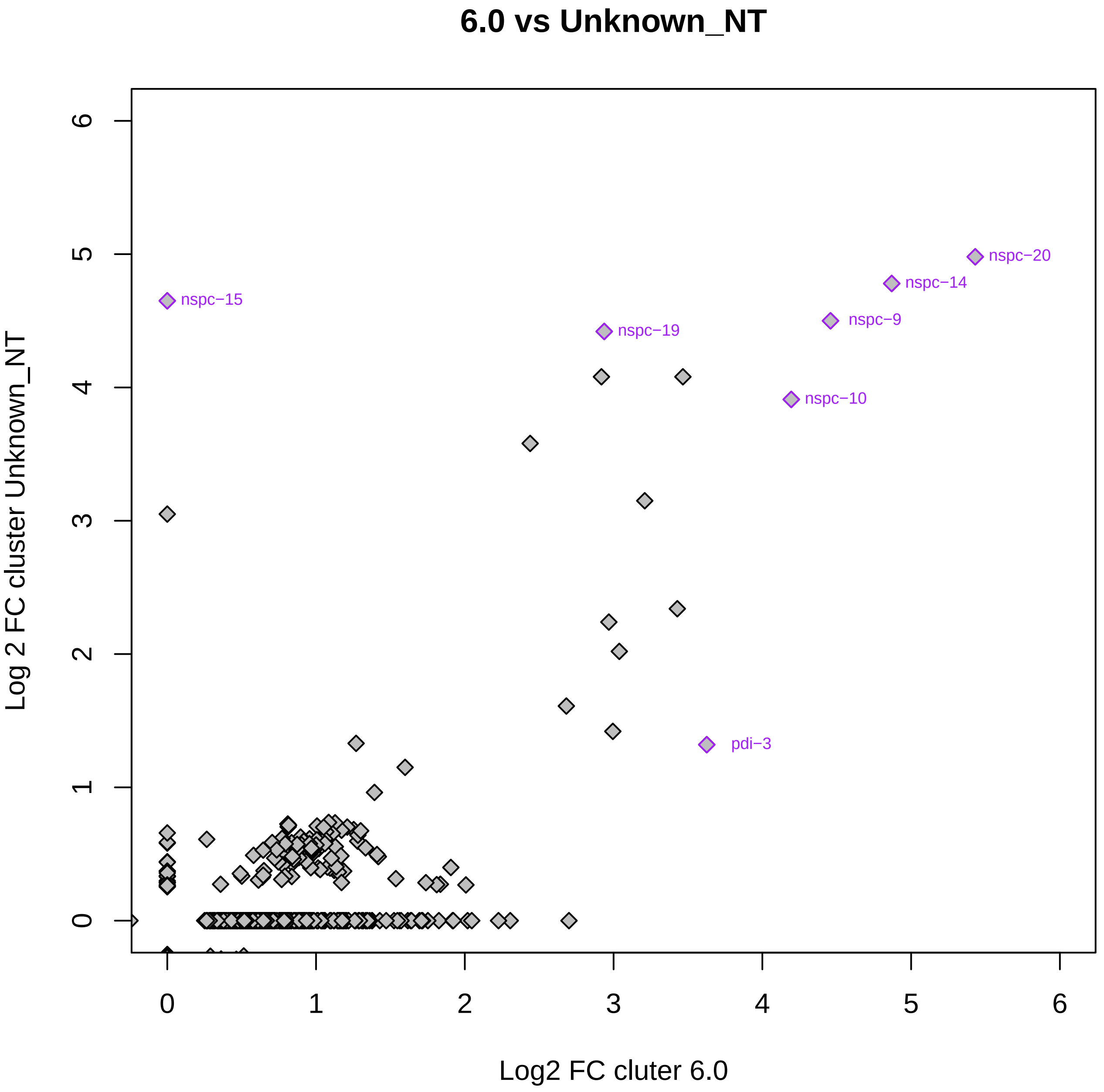
6.0 vs Unknown_NT
Log 2 FC cluster Unknown_NT
Log2 FC cluter 6.0
0
1
2
3
4
5
6
nspc−20
nspc−14
nspc−9
nspc−15
nspc−19
nspc−10
pdi−3

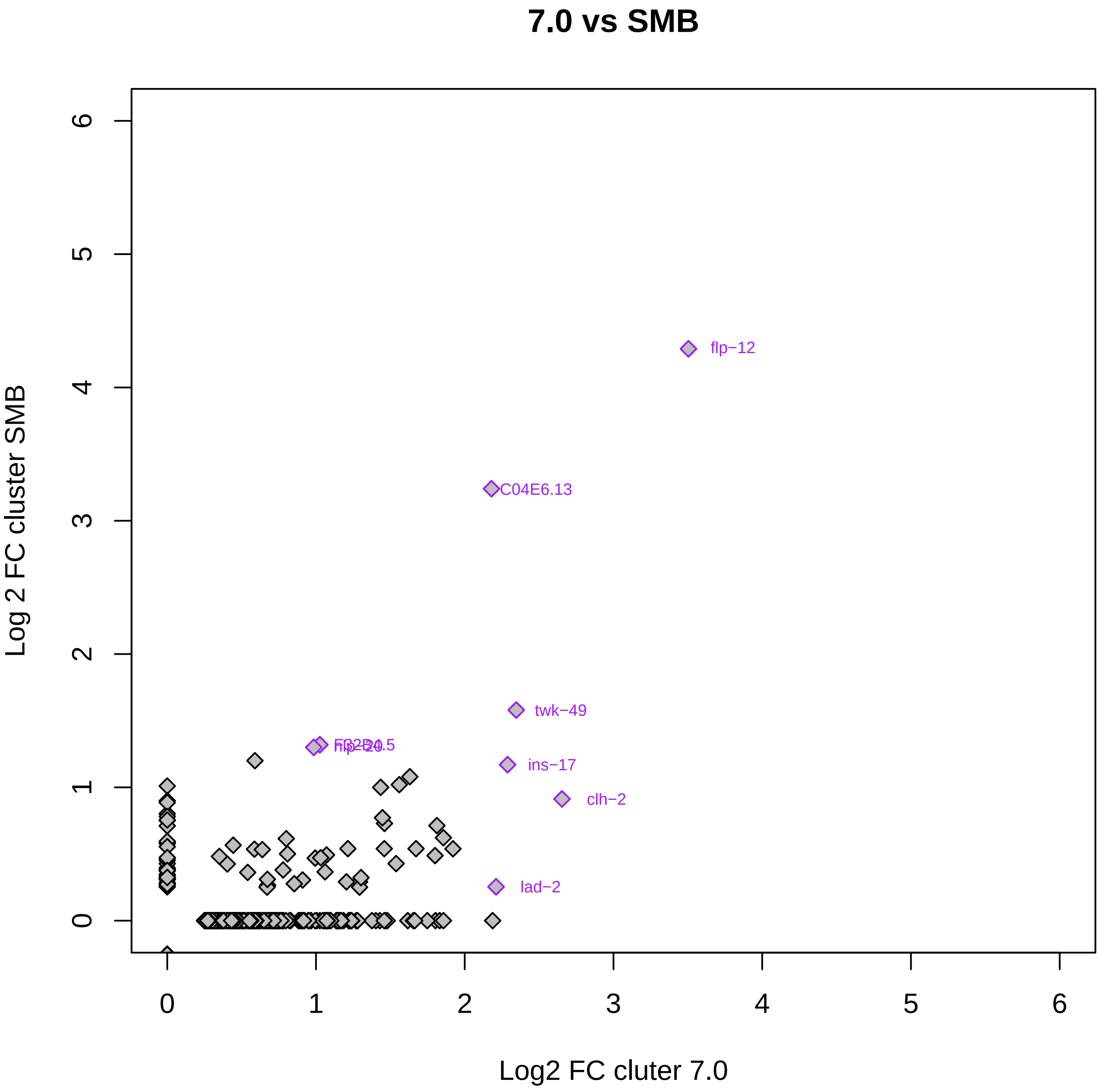

7.0 vs SMB
Log 2 FC cluster SMB
Log2 FC cluter 7.0
0
1
2
3
4
5
6
flp-12
C04E6.13
twk-49
ins-17
clh-2
lad-2
F32B4.5
flp-20

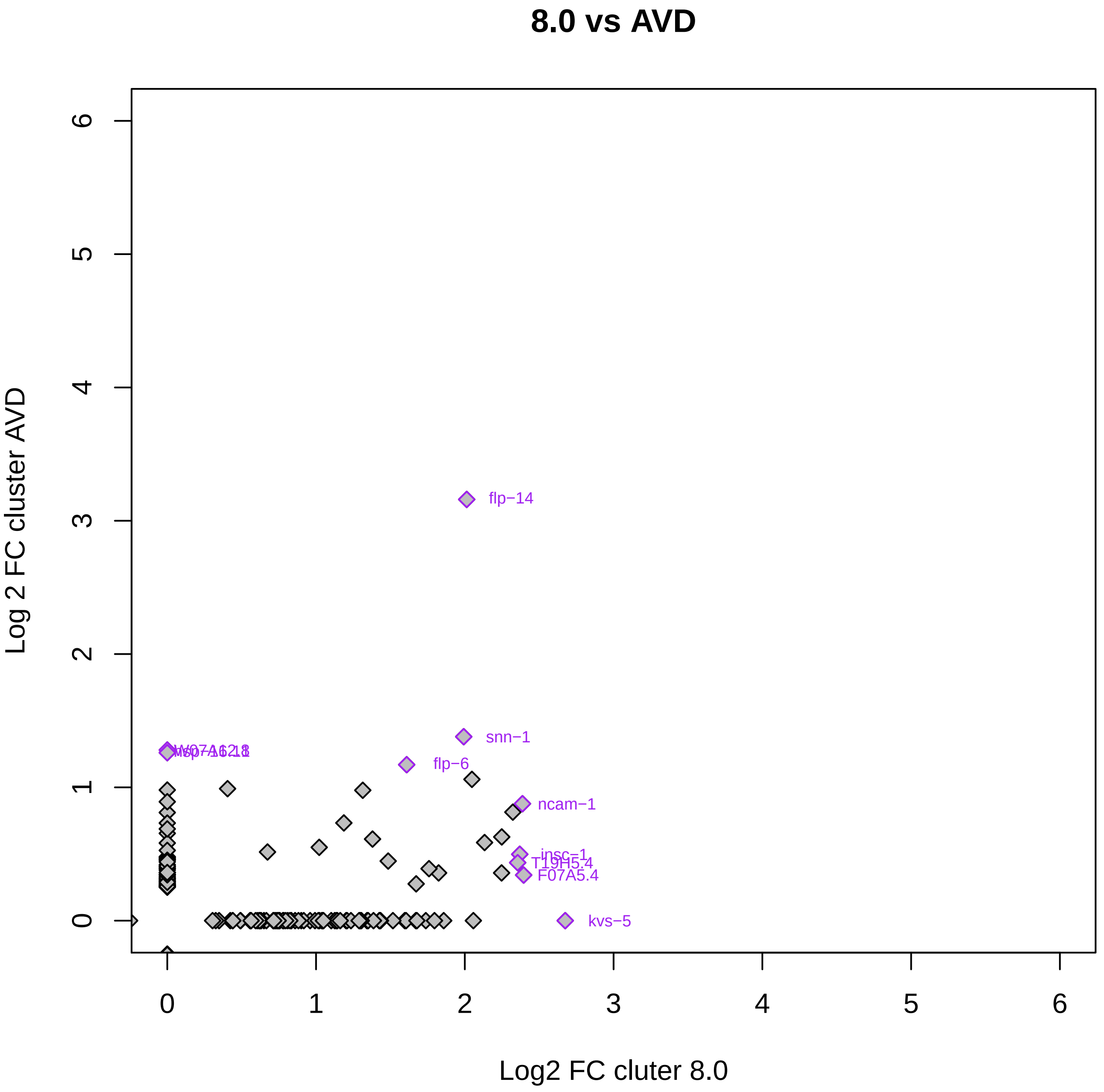

8.0 vs AVD
Log 2 FC cluster AVD
Log2 FC cluter 8.0
0
1
2
3
4
5
6
flp−14
snn−1
flp−6
ncam−1
insc−1
T19H5.4
F07A5.4
kvs−5

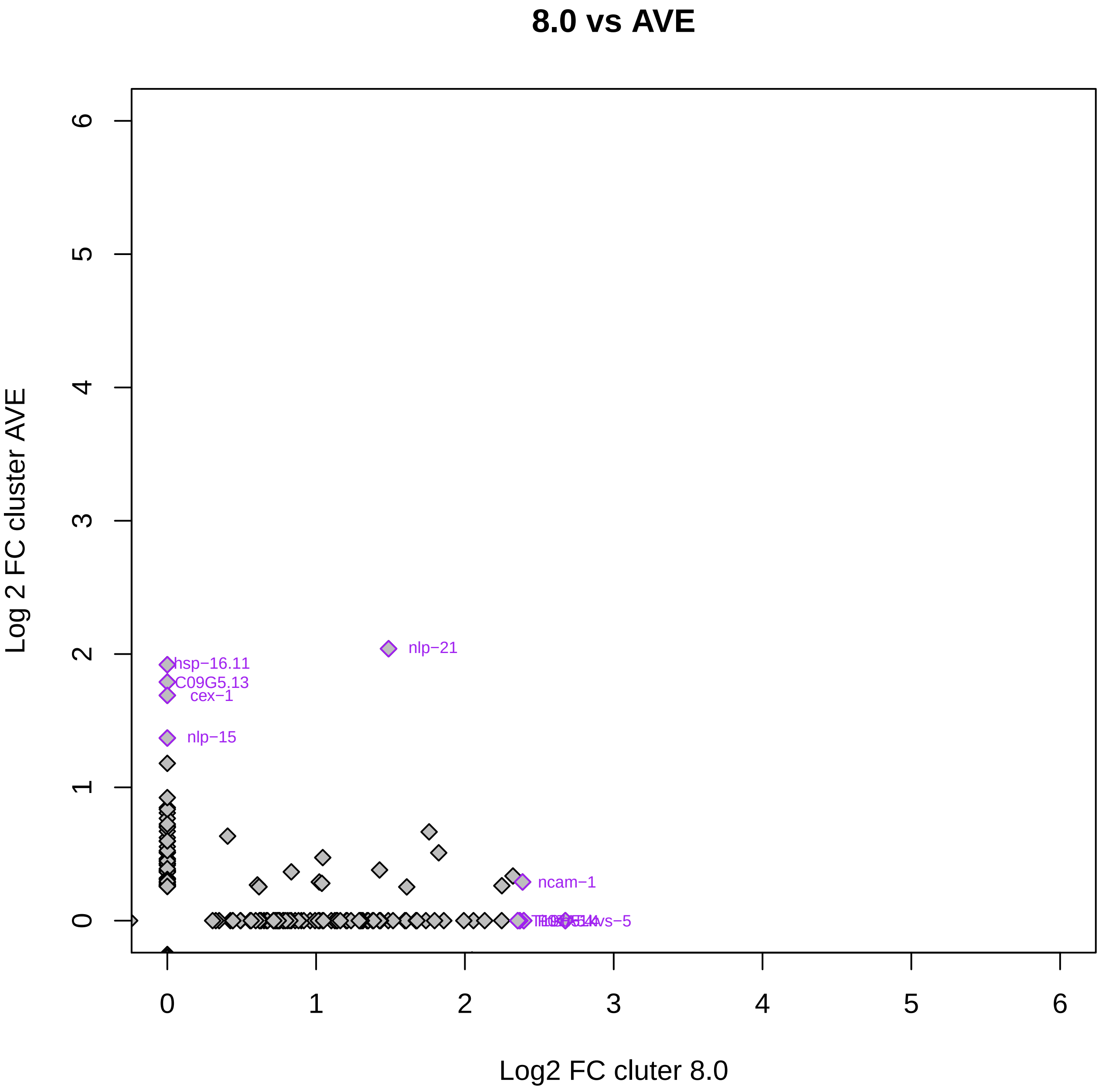

8.0 vs AVE
Log 2 FC cluster AVE
Log2 FC cluter 8.0
0
1
2
3
4
5
6
nlp−21
hsp−16.11
C09G5.13
cex−1
nlp−15
ncam−1

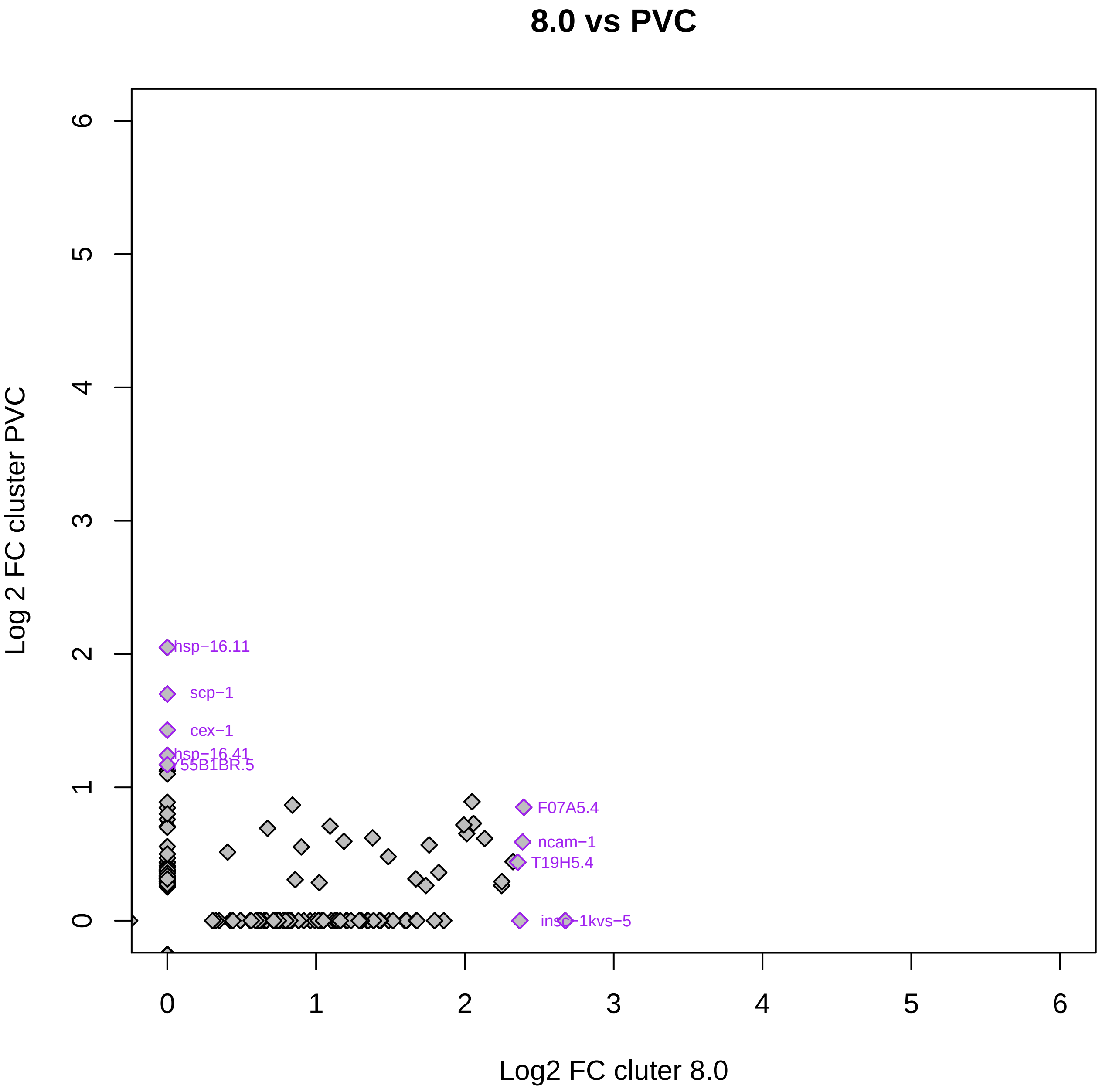
8.0 vs PVC
Log 2 FC cluster PVC
Log2 FC cluter 8.0
0
1
2
3
4
5
6
hsp−16.11
scp−1
cex−1
hsp−16.41
Y55B1BR.5
F07A5.4
ncam−1
T19H5.4
insc−1kvs−5

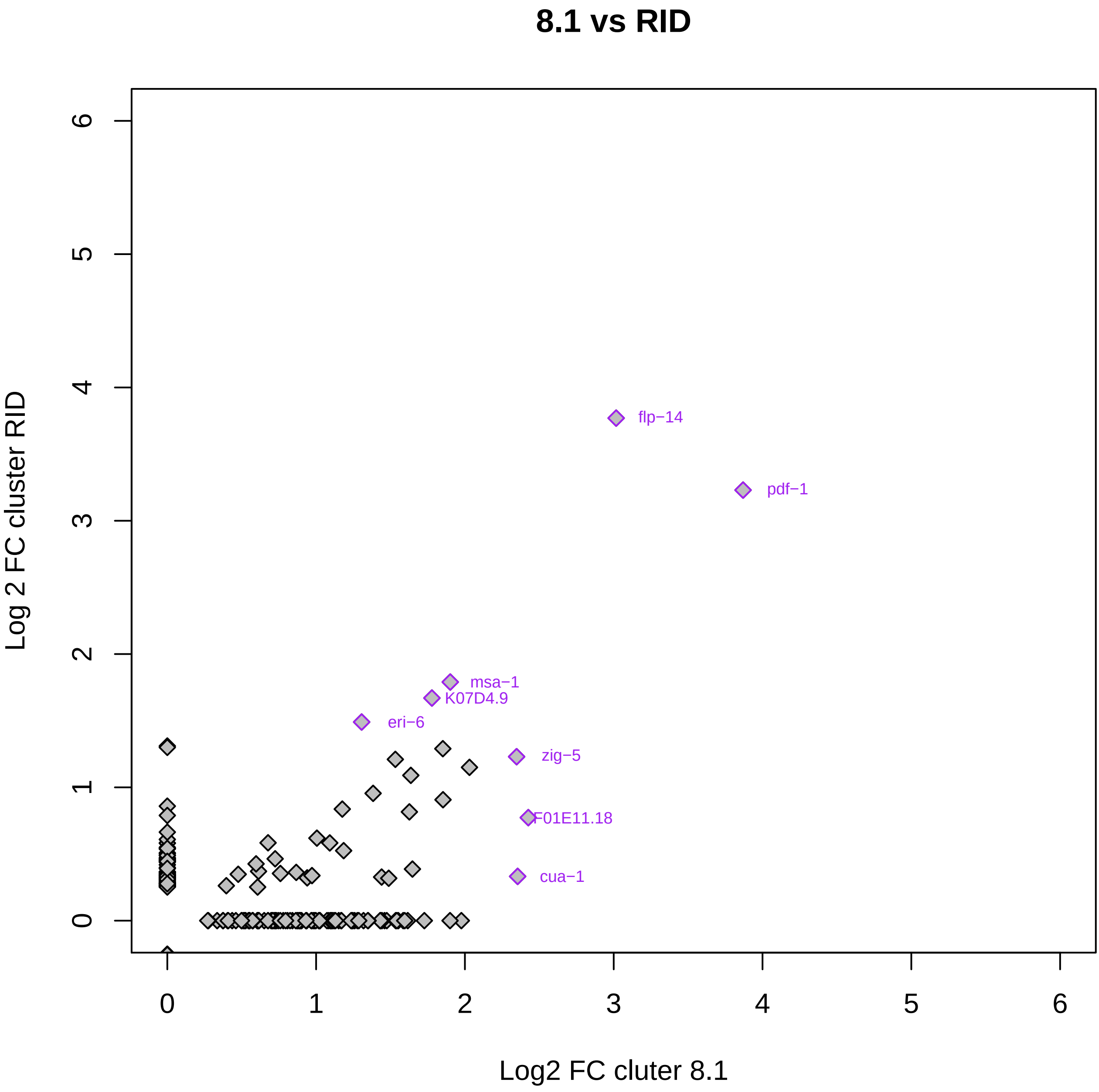
8.1 vs RID
Log 2 FC cluster RID
Log2 FC cluter 8.1
flp−14
pdf−1
msa−1
K07D4.9
eri−6
zig−5
F01E11.18
cua−1
0
1
2
3
4
5
6

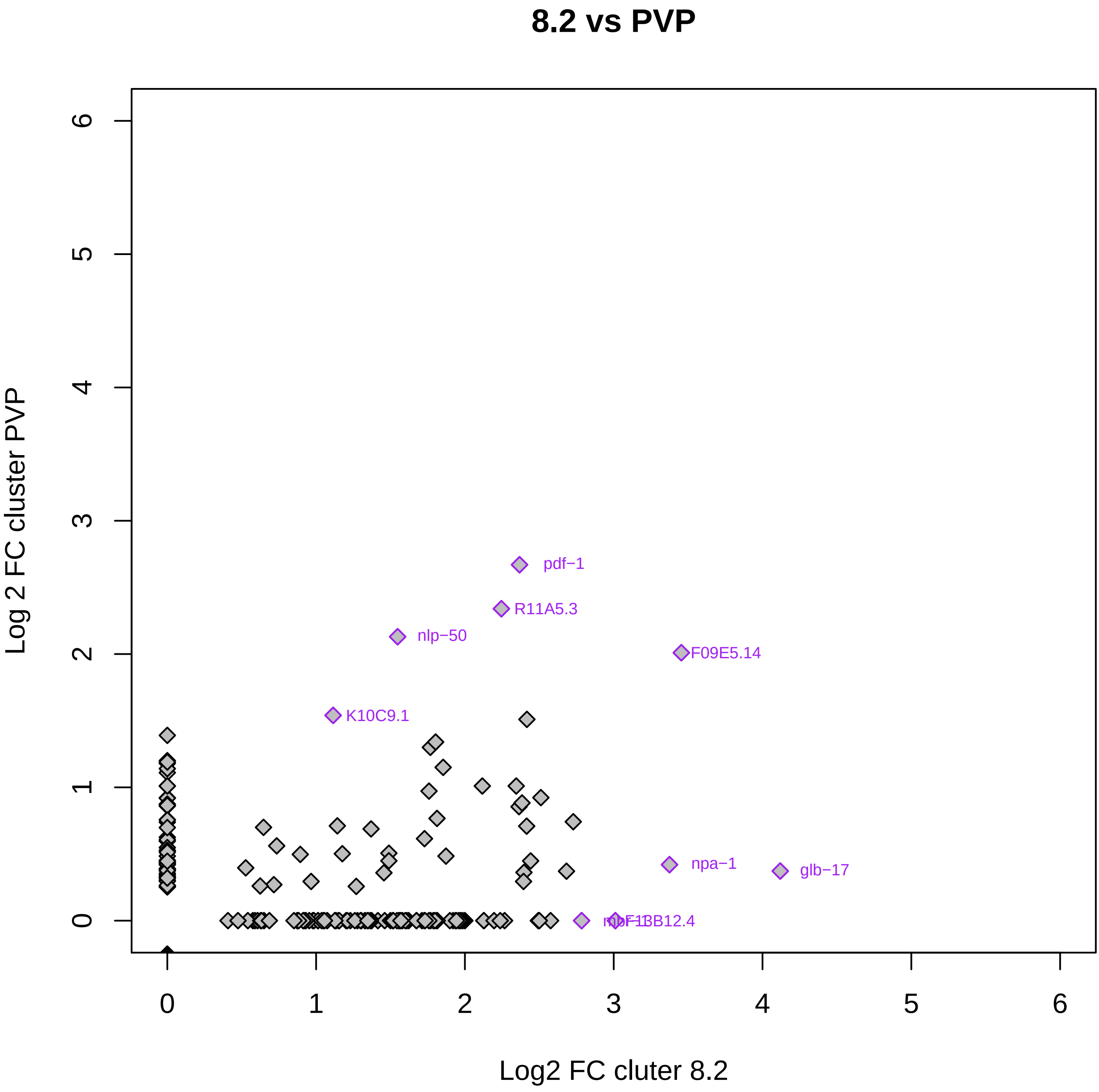

8.2 vs PVP
Log 2 FC cluster PVP
Log2 FC cluter 8.2
0
1
2
3
4
5
6
pdf−1
R11A5.3
nlp−50
F09E5.14
K10C9.1
npa−1
glb−17

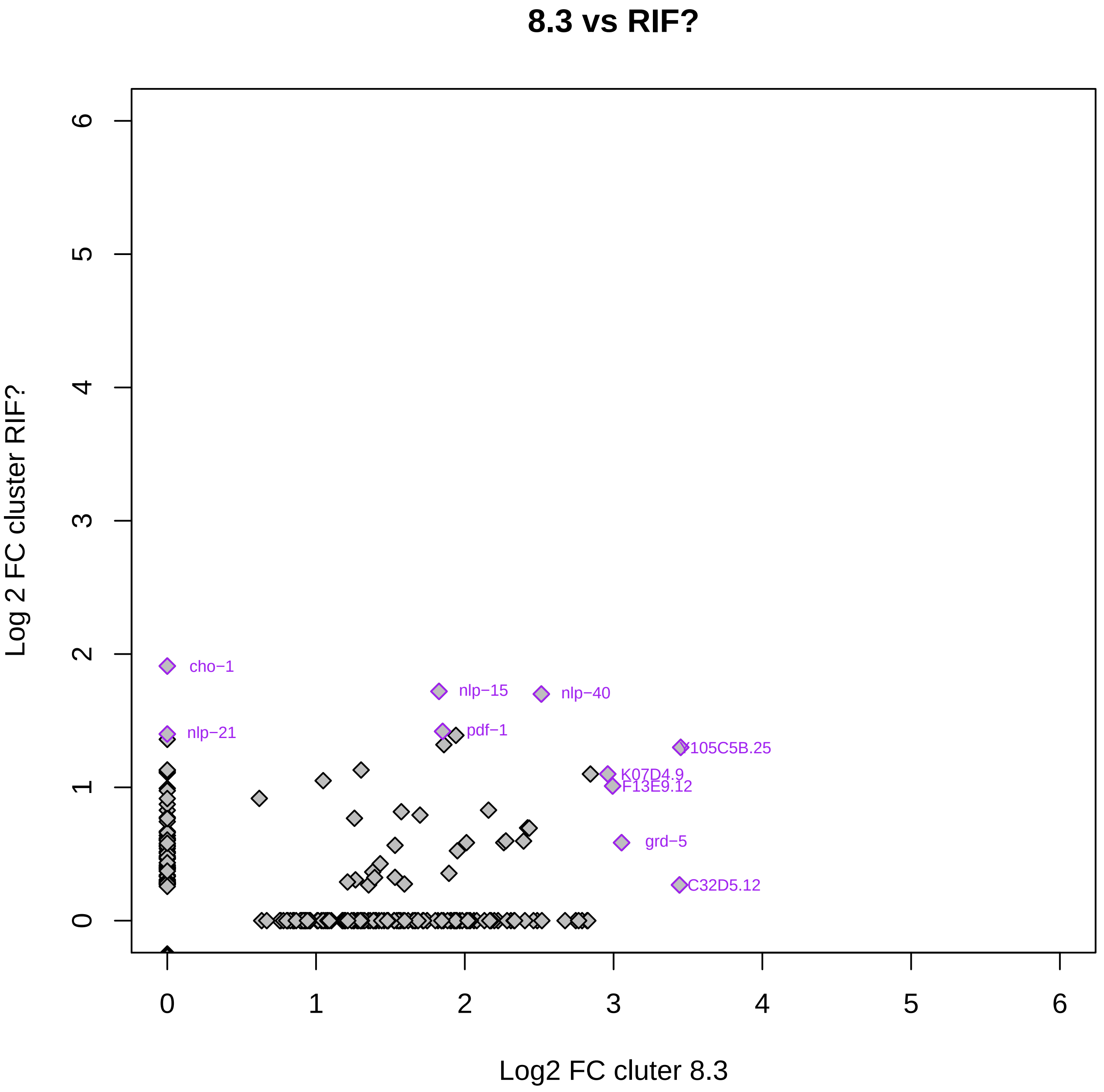

8.3 vs RIF?
Log 2 FC cluster RIF?
Log2 FC cluter 8.3
0
1
2
3
4
5
6
cho−1
nlp−21
nlp−15
pdf−1
nlp−40
Y105C5B.25
K07D4.9
F13E9.12
grd−5
C32D5.12

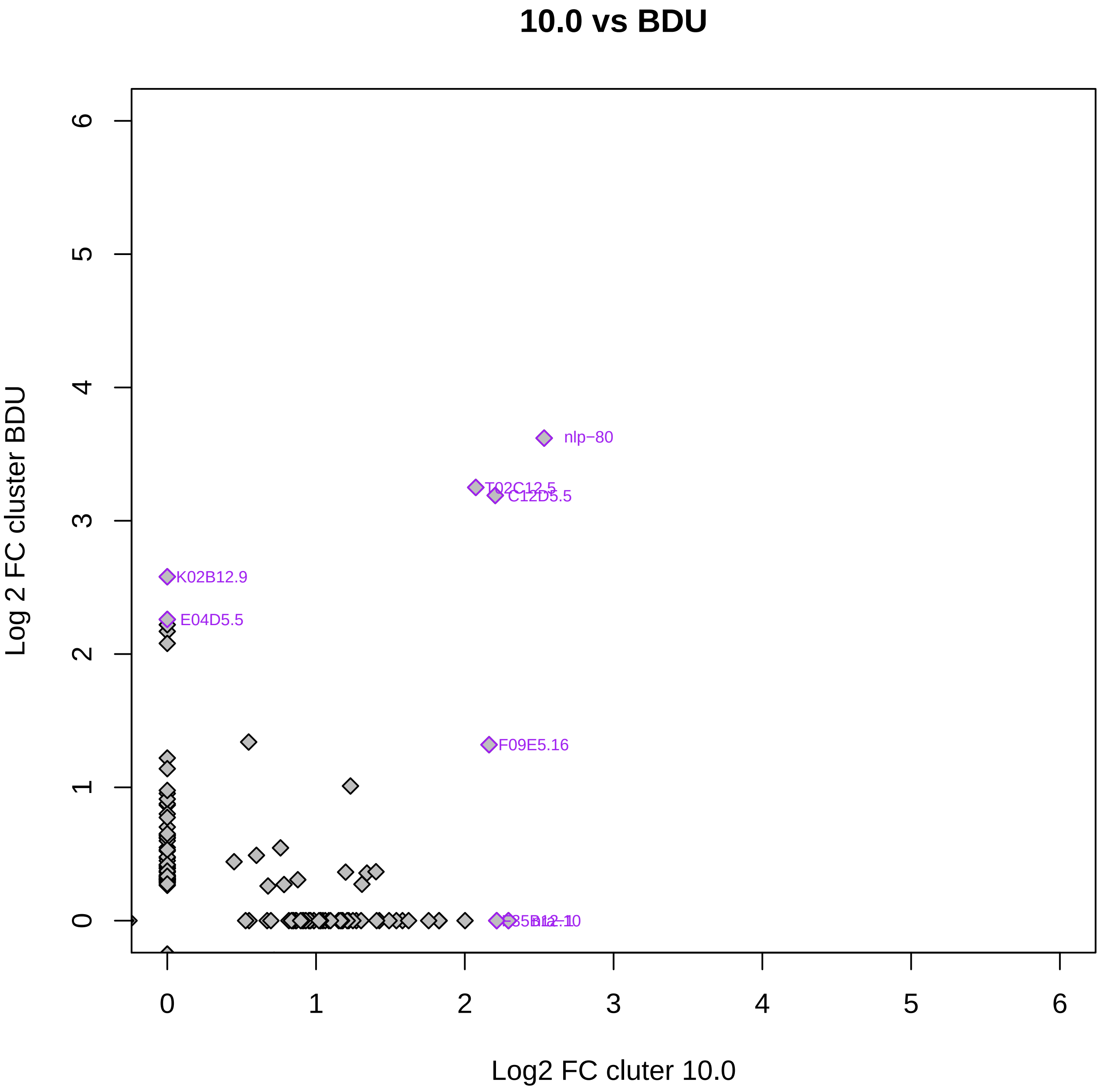

10.0 vs BDU
nlp−80
T02C12.5
C12D5.5
K02B12.9
E04D5.5
F09E5.16
0
1
2
3
4
5
6
0
1
2
3
4
5
6
Log 2 FC cluster BDU
Log2 FC cluter 10.0

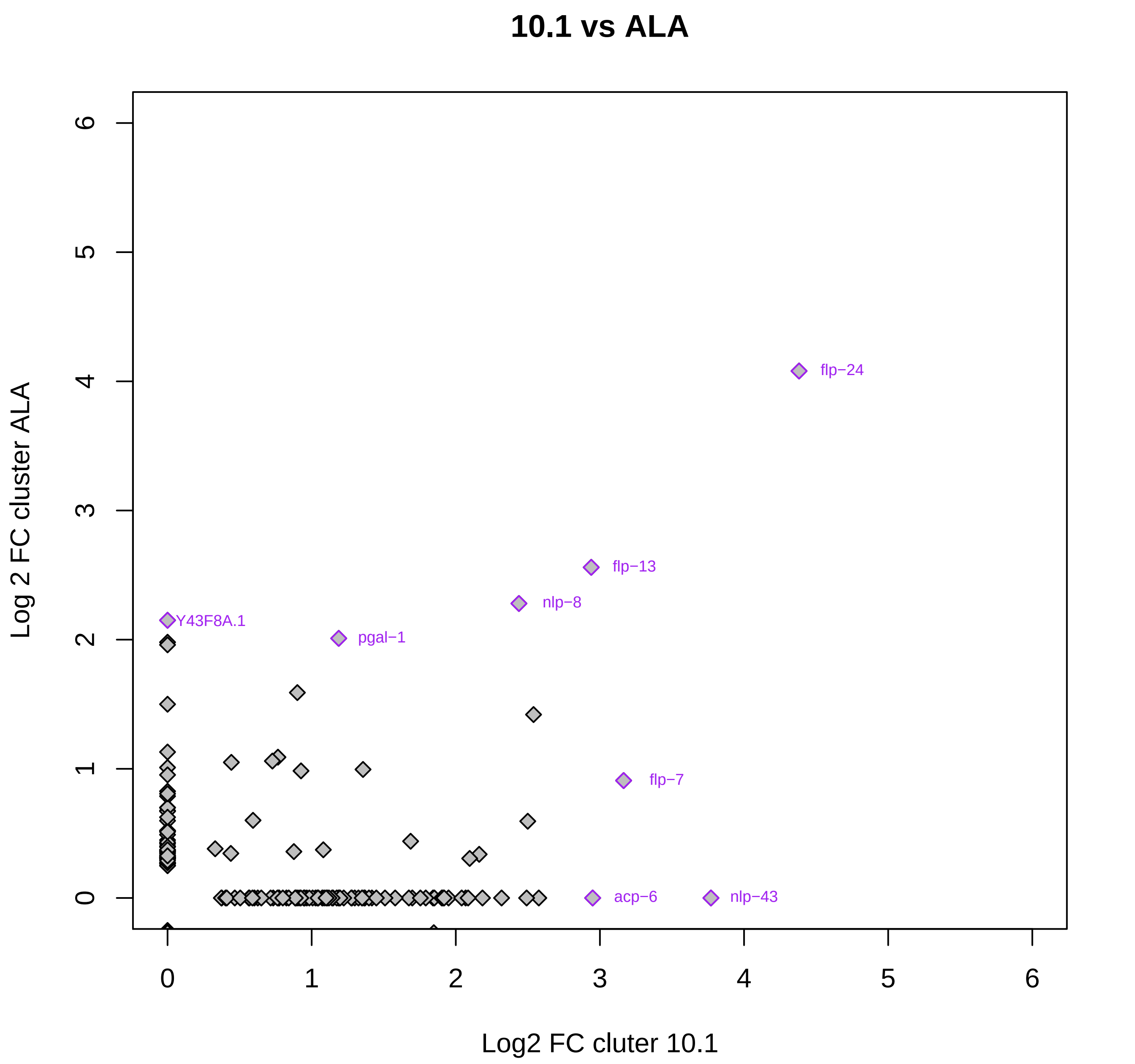

10.1 vs ALA
Log 2 FC cluster ALA
Log2 FC cluter 10.1
0
1
2
3
4
5
6
flp−24
flp−13
nlp−8
Y43F8A.1
pgal−1
flp−7
acp−6
nlp−43

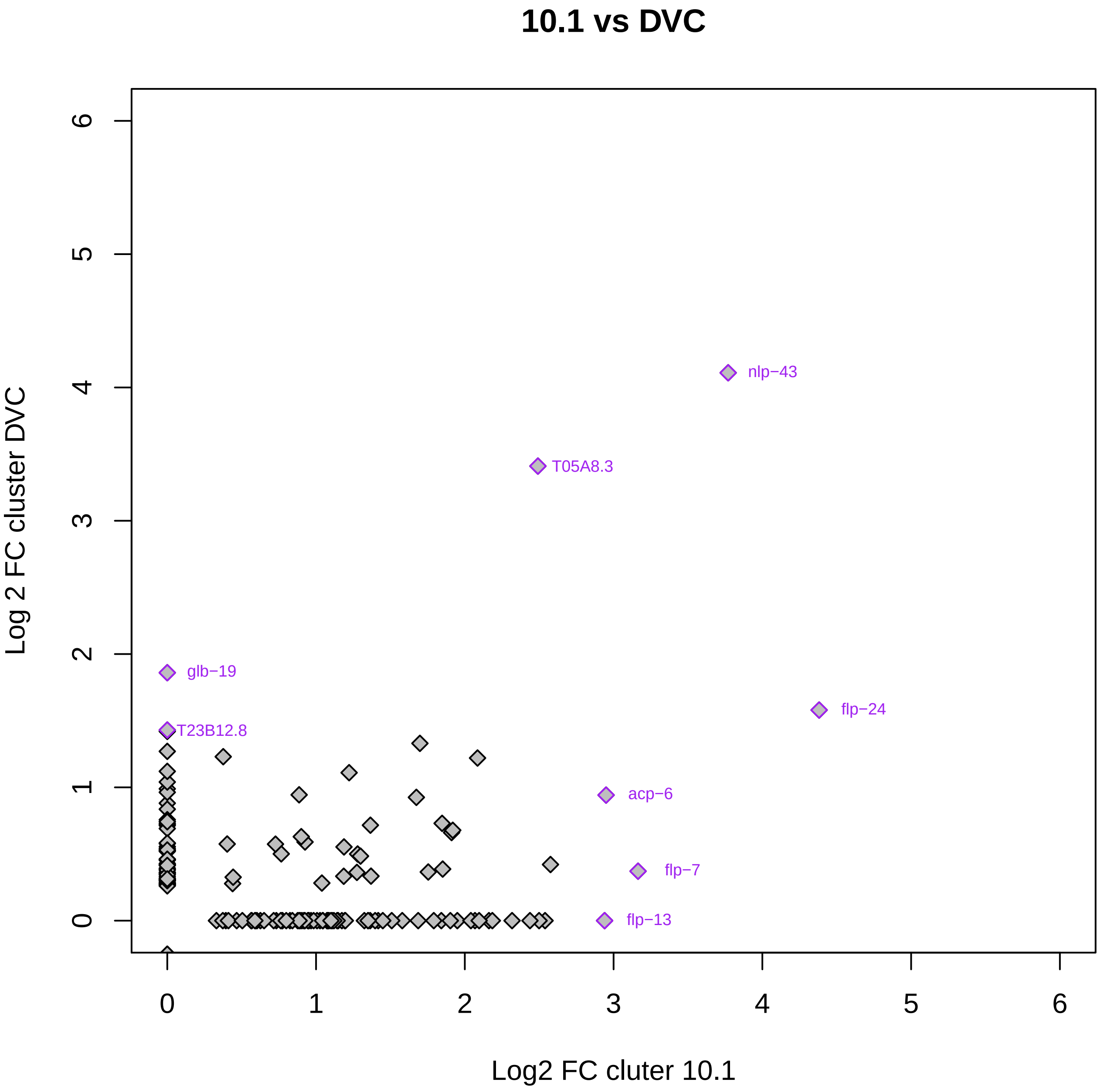

10.1 vs DVC
Log 2 FC cluster DVC
Log2 FC cluter 10.1
0
1
2
3
4
5
6
nlp−43
T05A8.3
glb−19
T23B12.8
flp−24
acp−6
flp−7
flp−13

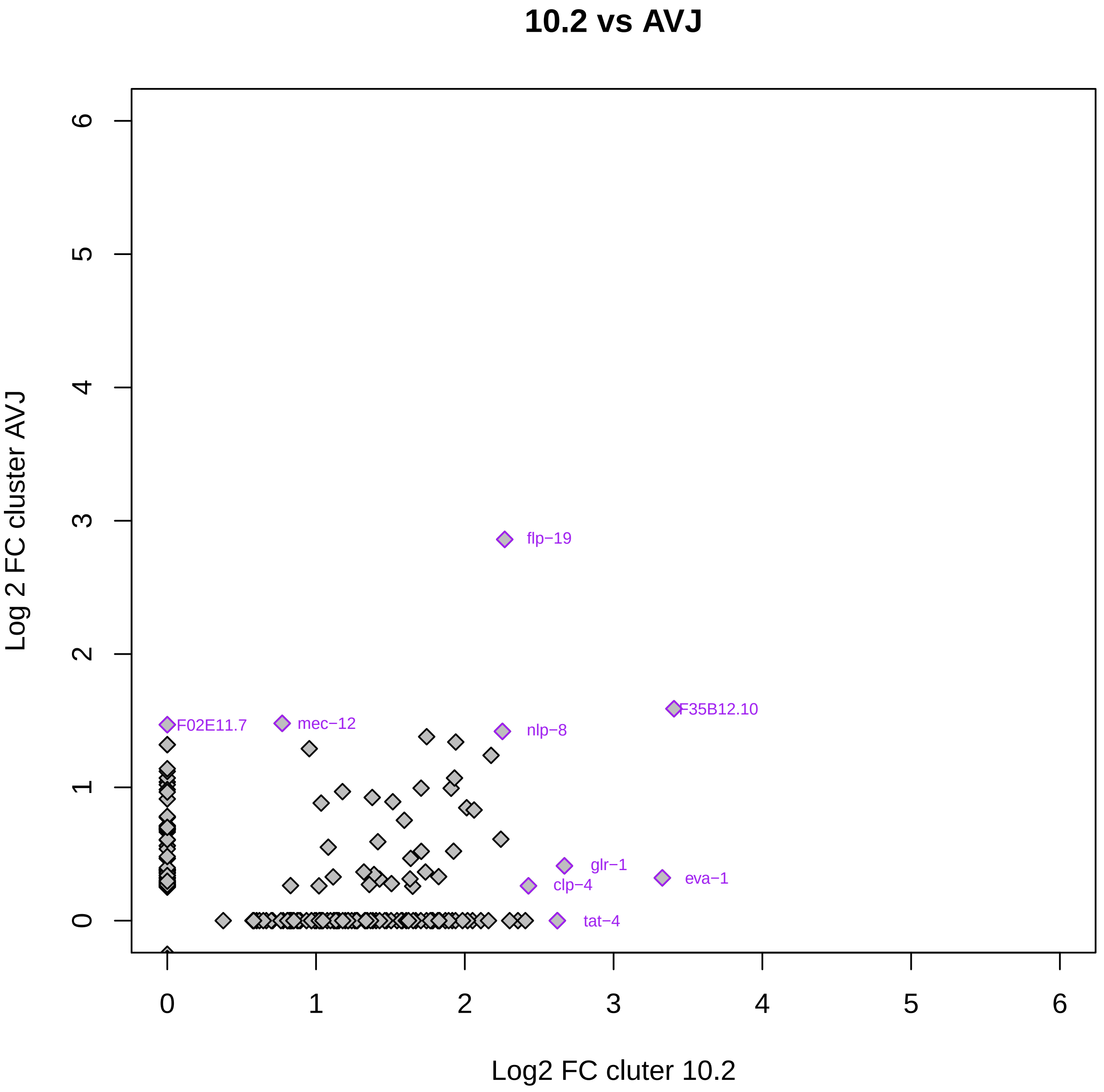

10.2 vs AVJ
Log 2 FC cluster AVJ
Log2 FC cluter 10.2
0
1
2
3
4
5
6
flp−19
F35B12.10
F02E11.7
mec−12
nlp−8
glr−1
eva−1
clp−4
tat−4

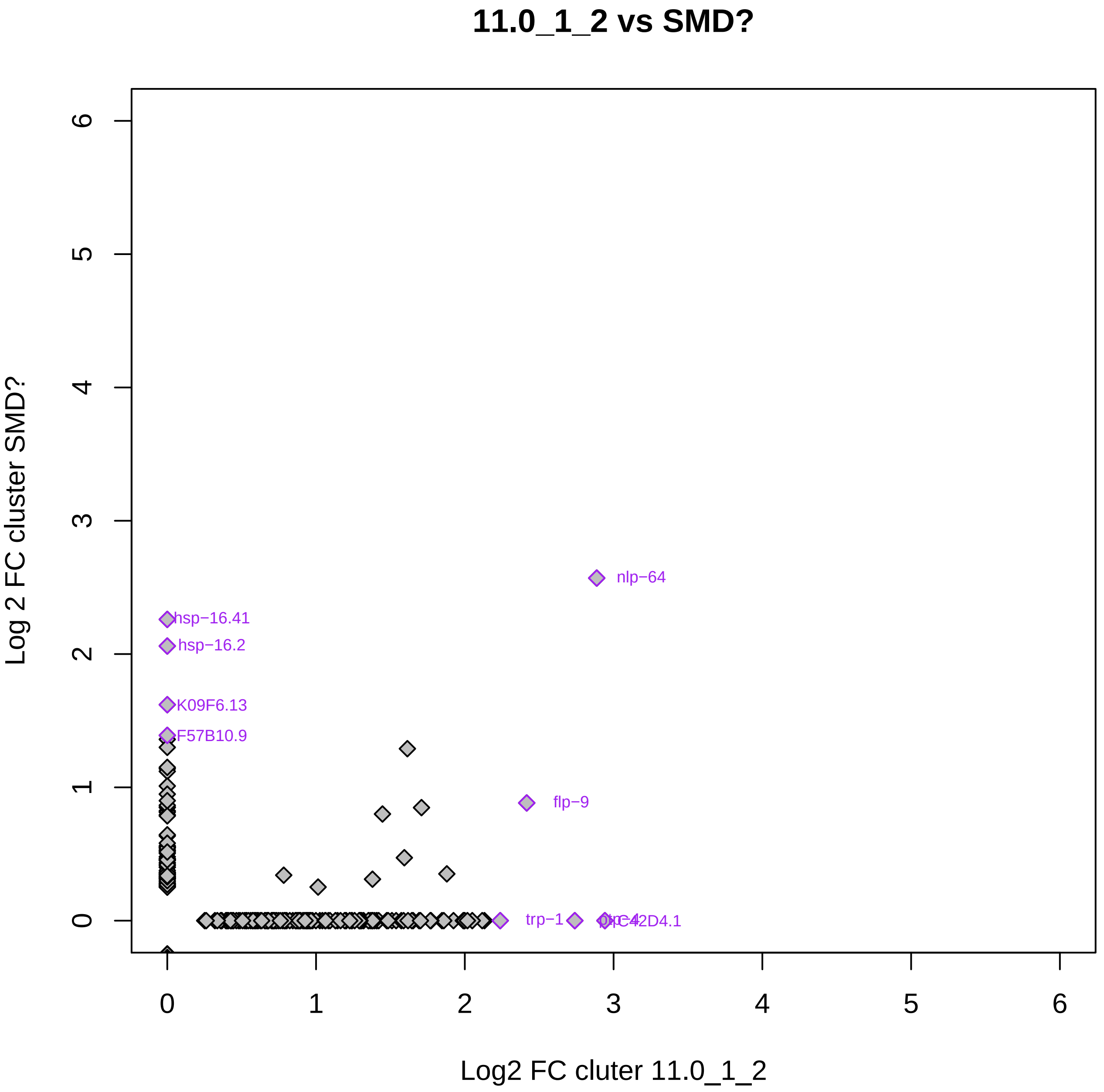

11.0_1_2 vs SMD?
Log 2 FC cluster SMD?
Log2 FC cluter 11.0_1_2
0
1
2
3
4
5
6
0
1
2
3
4
5
6
nlp−64
hsp−16.41
hsp−16.2
K09F6.13
F57B10.9
flp−9
trp−1

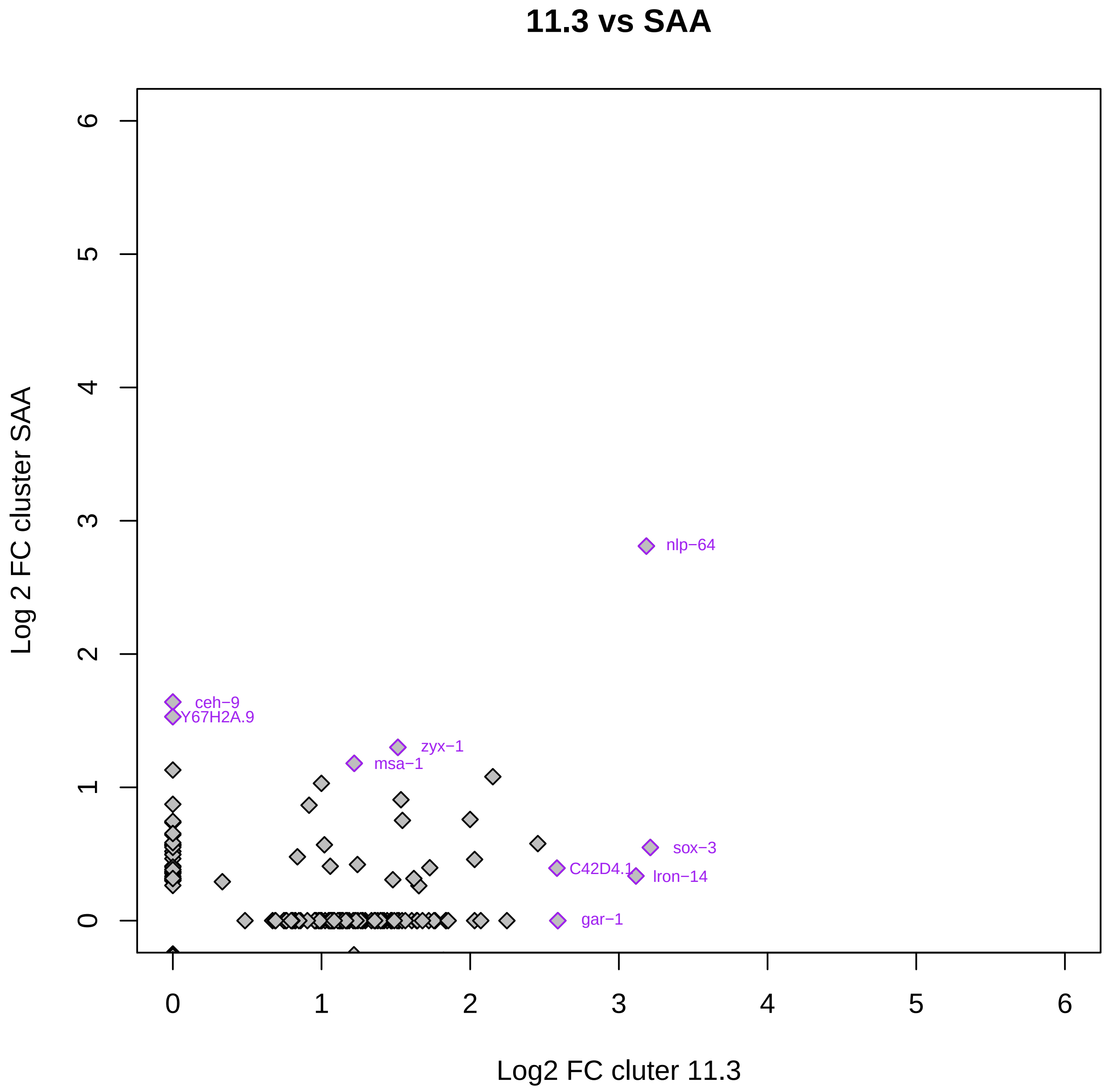

11.3 vs SAA
Log 2 FC cluster SAA
Log2 FC cluter 11.3
0
1
2
3
4
5
6
nlp−64
ceh−9
Y67H2A.9
zyx−1
msa−1
sox−3
C42D4.1
lron−14
gar−1

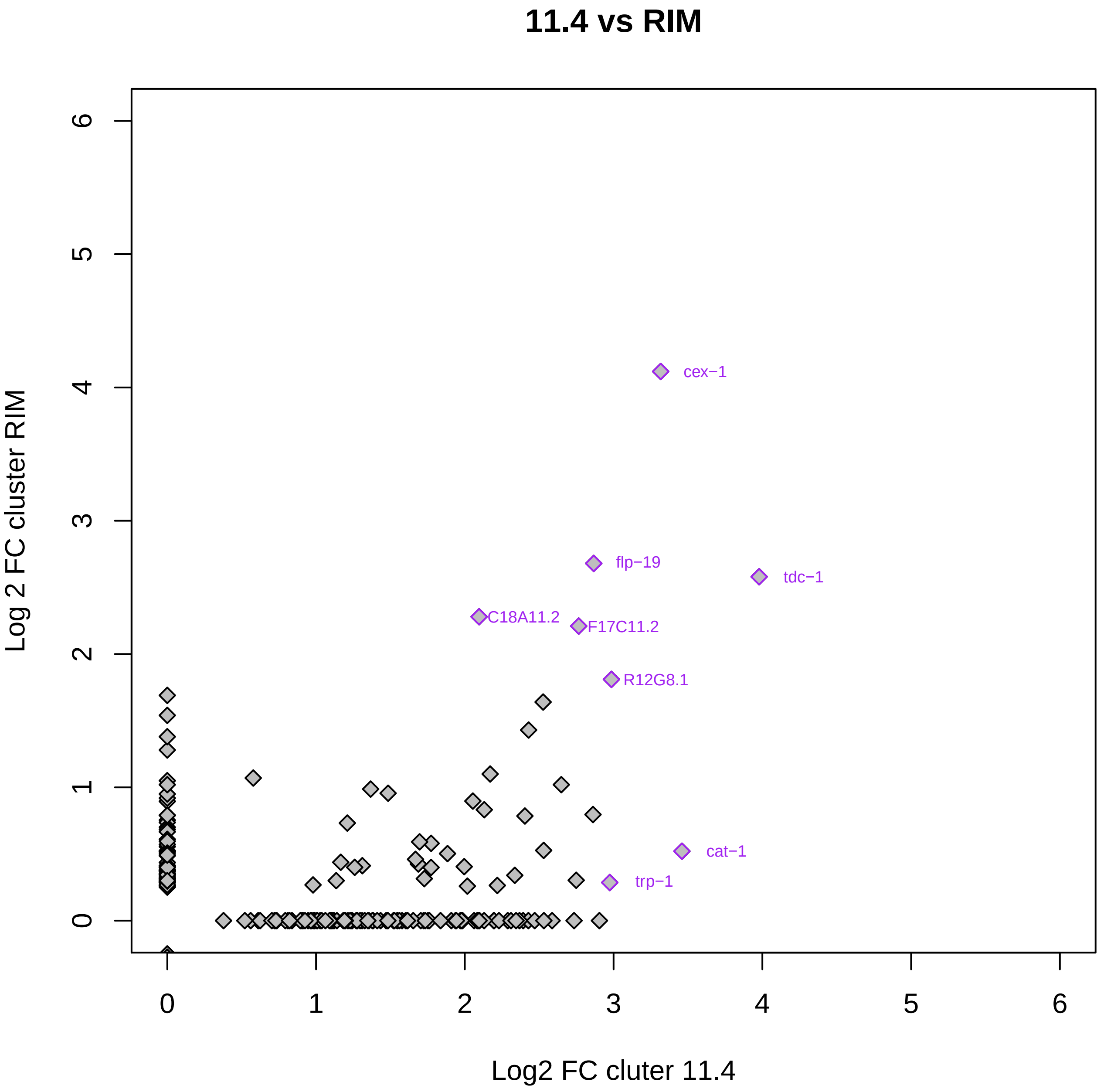
11.4 vs RIM
Log 2 FC cluster RIM
Log2 FC cluter 11.4
0
1
2
3
4
5
6
cex−1
flp−19
tdc−1
C18A11.2
F17C11.2
R12G8.1
cat−1
trp−1

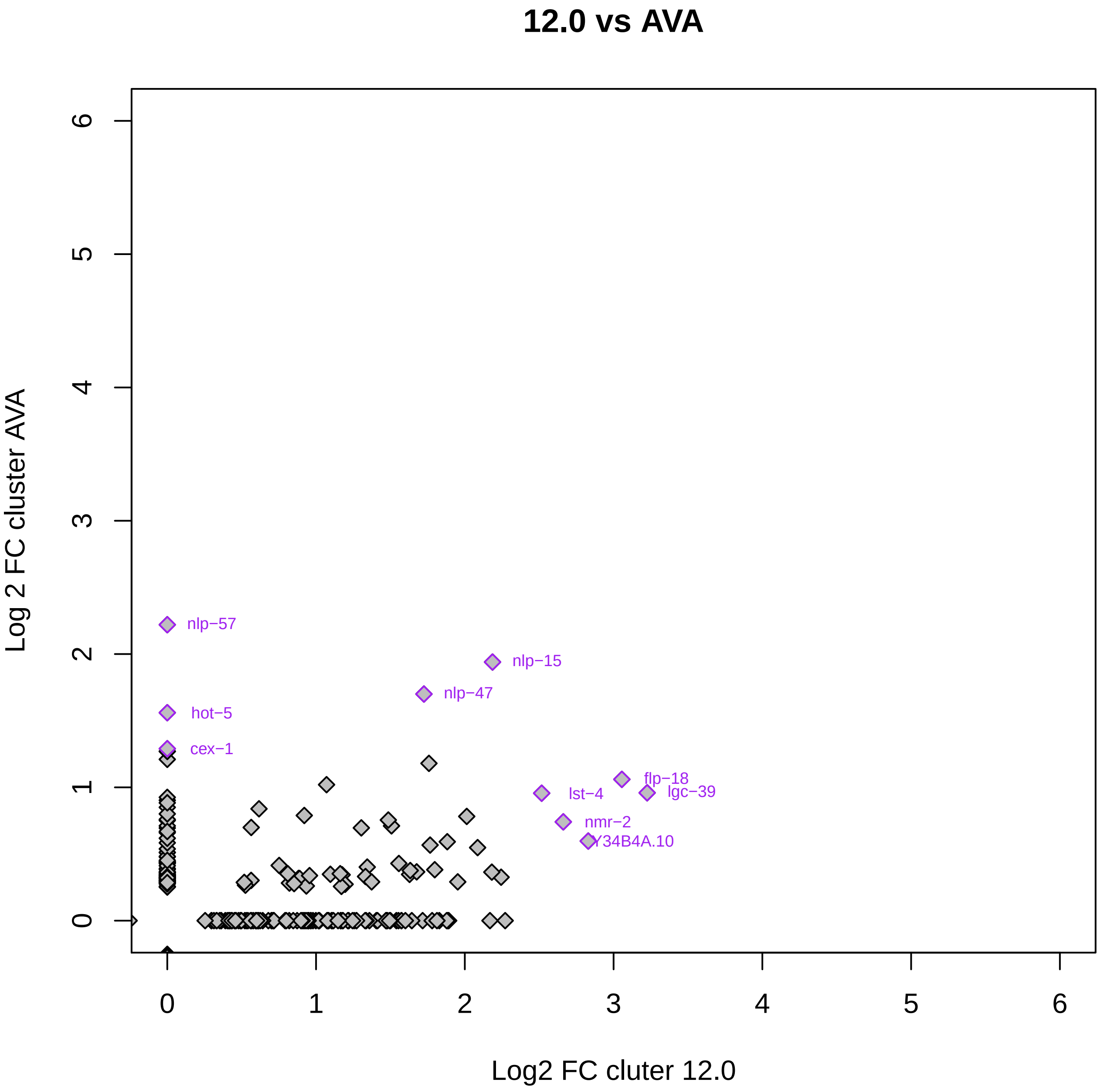

12.0 vs AVA
nlp−57
nlp−15
nlp−47
hot−5
cex−1
flp−18
lgc−39
lst−4
nmr−2
Y34B4A.10
Log 2 FC cluster AVA
Log2 FC cluter 12.0
0
1
2
3
4
5
6

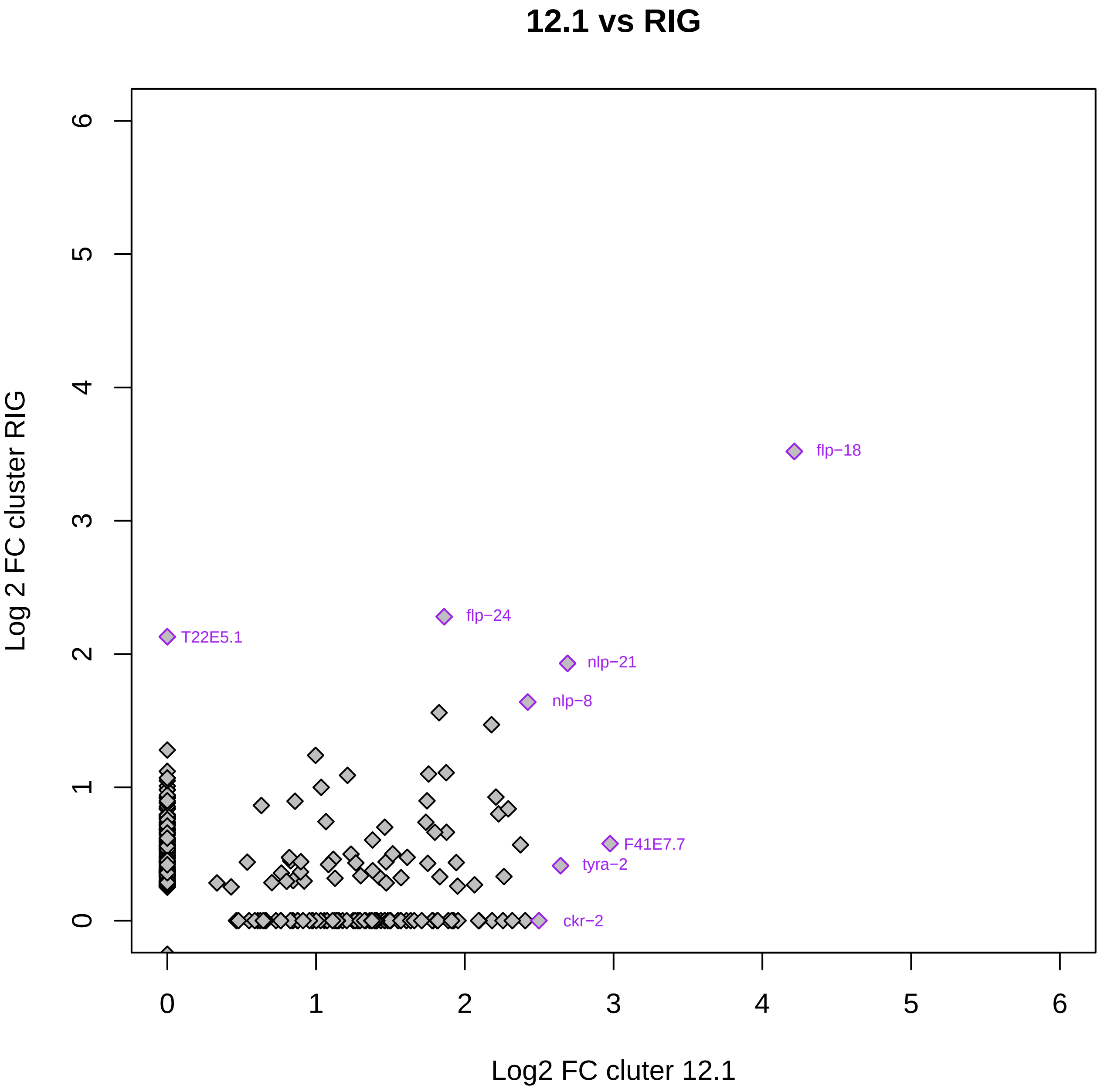
12.1 vs RIG
Log 2 FC cluster RIG
Log2 FC cluter 12.1
0
1
2
3
4
5
6
flp−18
flp−24
T22E5.1
nlp−21
nlp−8
F41E7.7
tyra−2
ckr−2

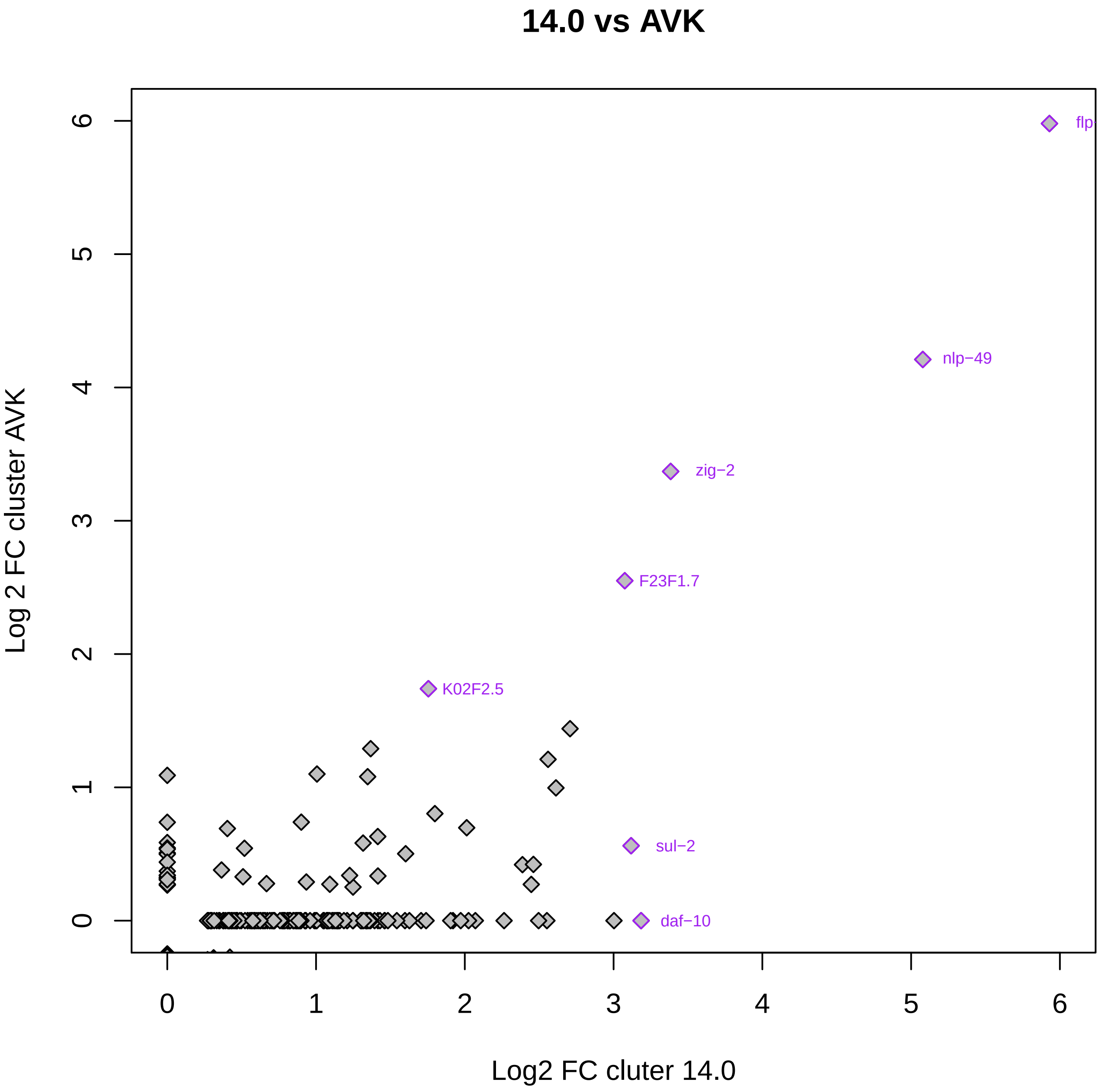

14.0 vs AVK
Log 2 FC cluster AVK
Log2 FC cluter 14.0
0
1
2
3
4
5
6
0
1
2
3
4
5
6
flp
nlp−49
zig−2
F23F1.7
K02F2.5
sul−2
daf−10

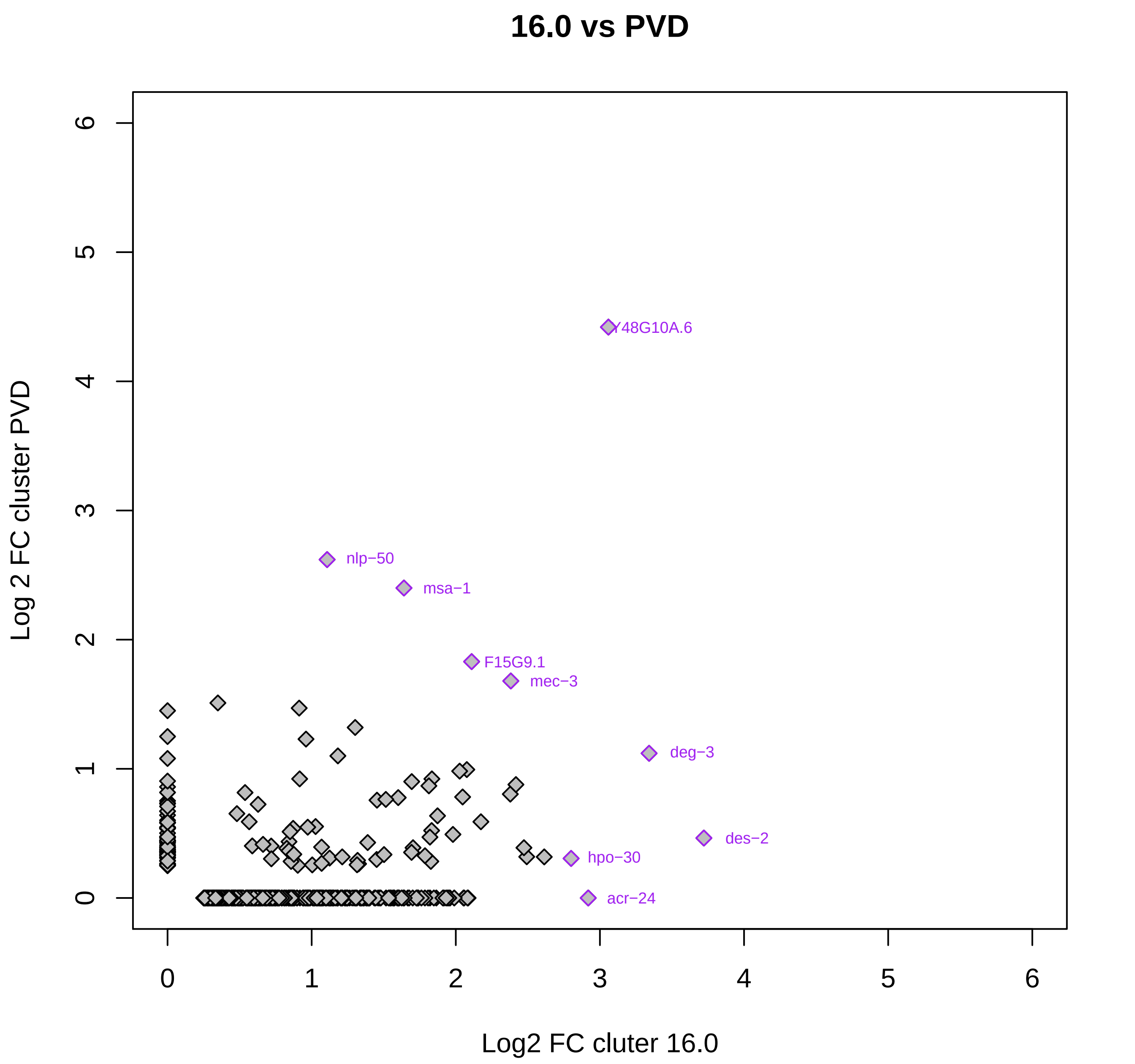
16.0 vs PVD
Y48G10A.6
nlp−50
msa−1
F15G9.1
mec−3
deg−3
des−2
hpo−30
acr−24
Log 2 FC cluster PVD
Log2 FC cluter 16.0
0
1
2
3
4
5
6

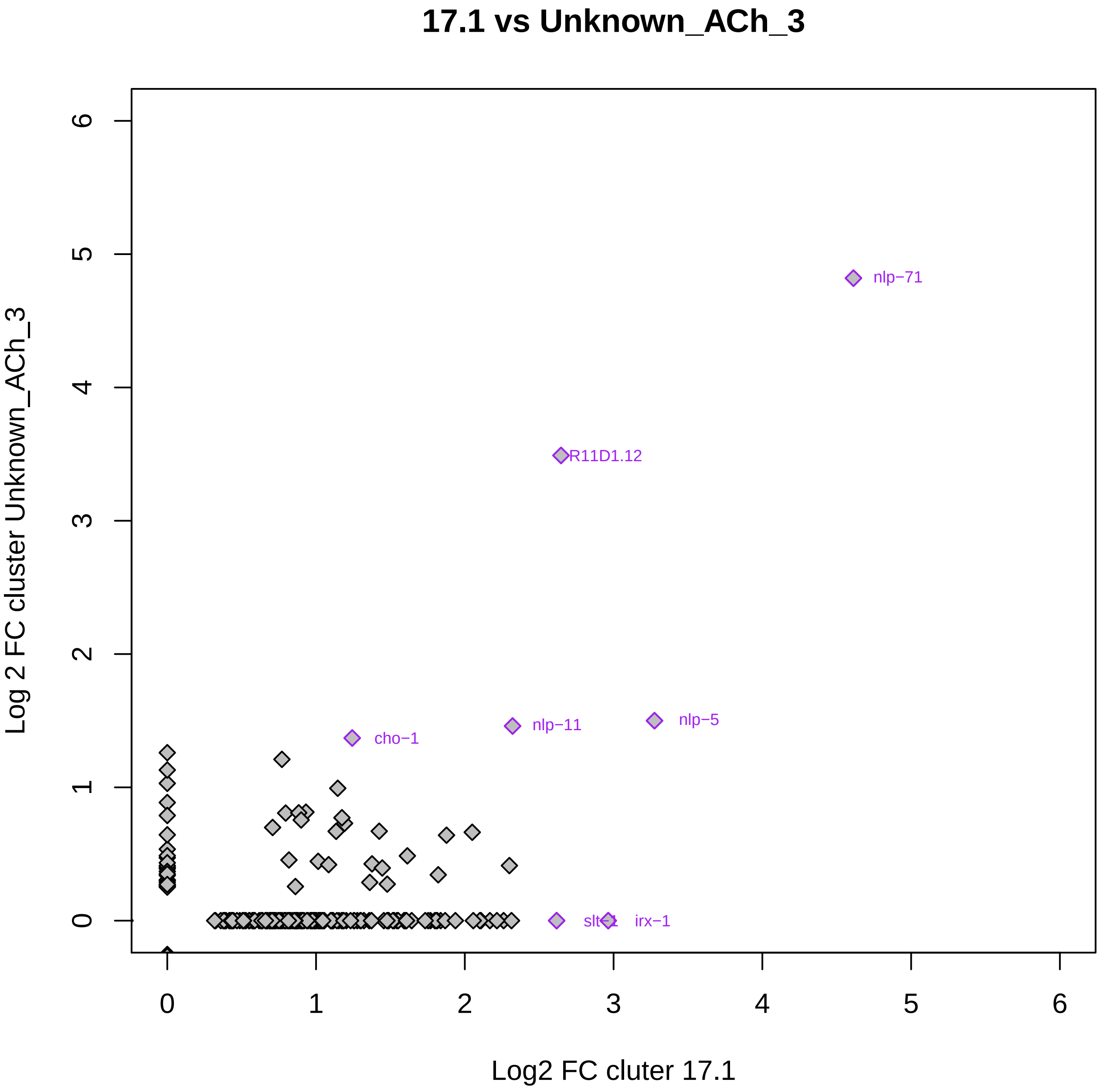

17.1 vs Unknown_ACh_3
Log 2 FC cluster Unknown_ACh_3
Log2 FC cluter 17.1
0
1
2
3
4
5
6
nlp−71
R11D1.12
cho−1
nlp−11
nlp−5
irx−1

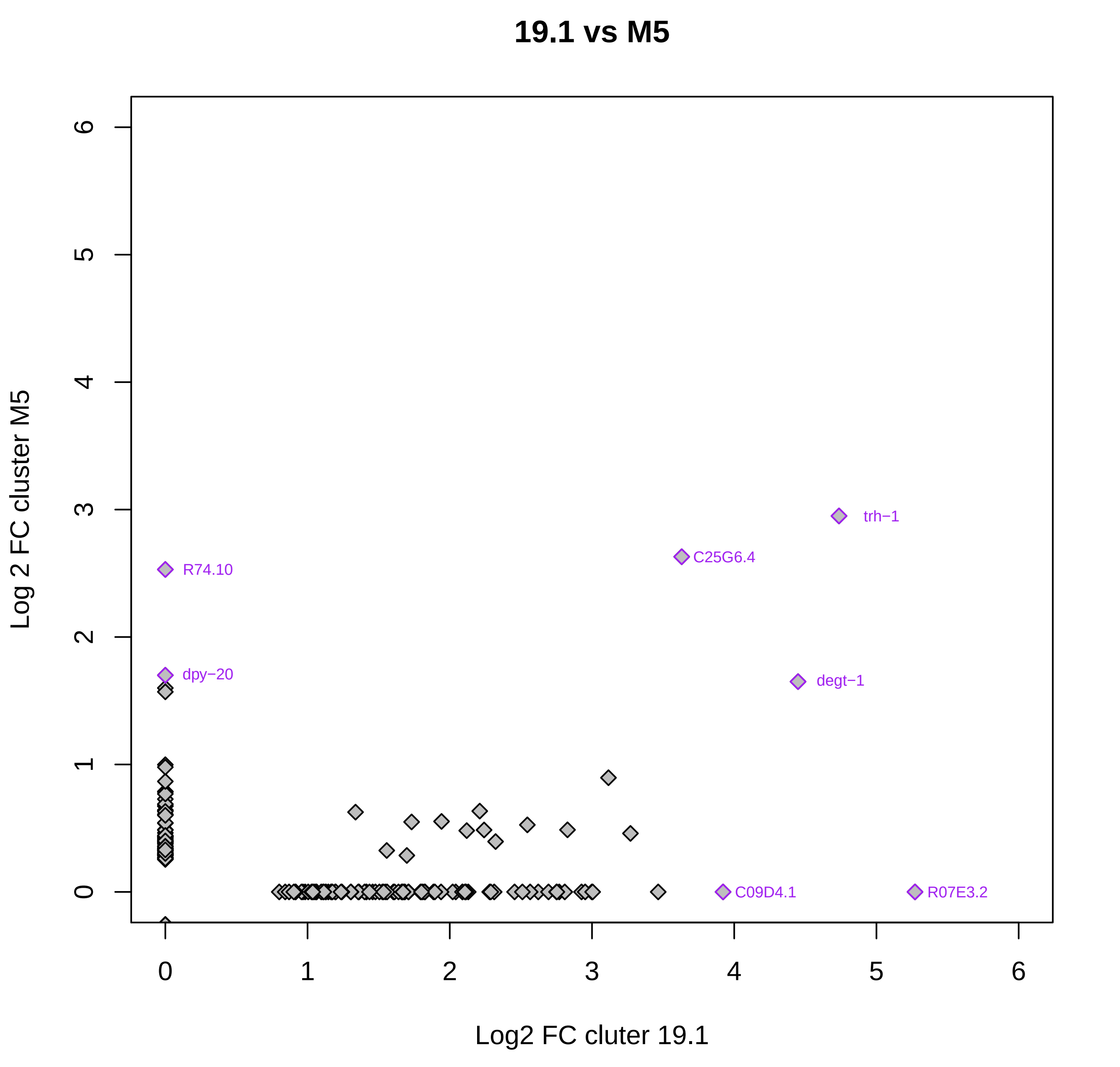
19.1 vs M5
Log 2 FC cluster M5
Log2 FC cluter 19.1
0
1
2
3
4
5
6
R74.10
dpy−20
C25G6.4
trh−1
degt−1
C09D4.1
R07E3.2

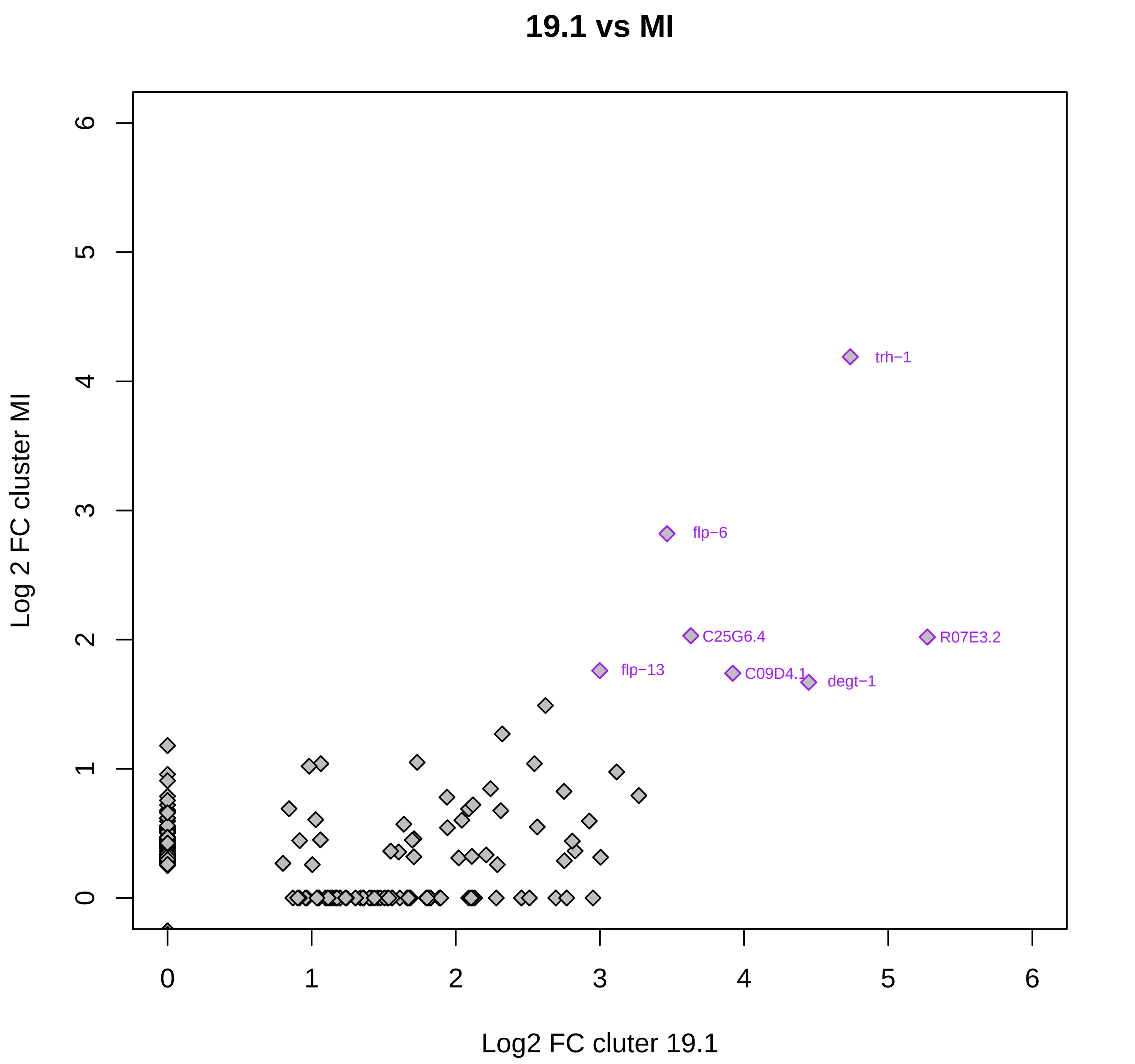

19.1 vs MI
Log 2 FC cluster MI
Log2 FC cluter 19.1
0
1
2
3
4
5
6
0
1
2
3
4
5
6
trh−1
flp−6
C25G6.4
R07E3.2
flp−13
C09D4.1
degt−1

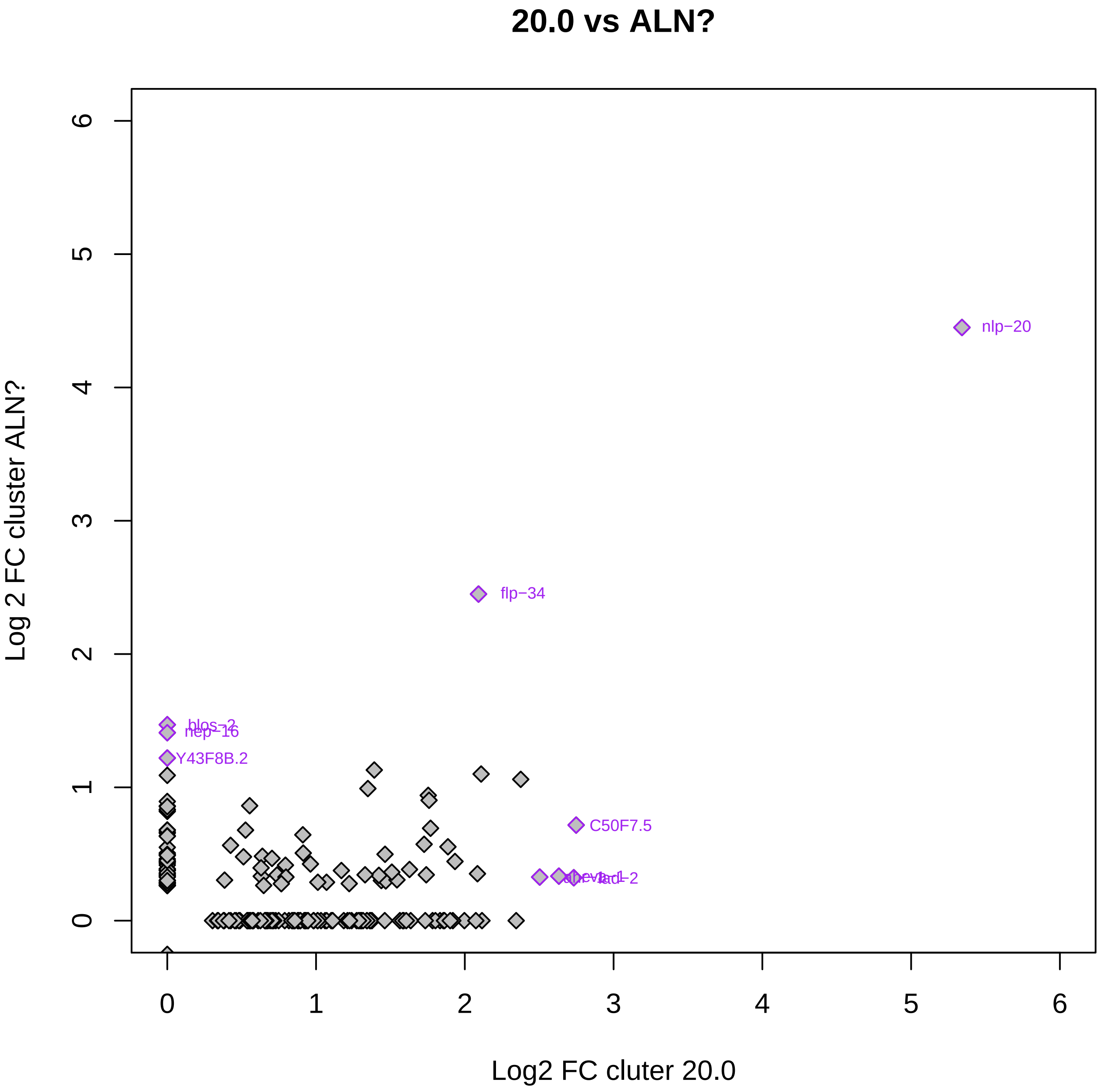

20.0 vs ALN?
Log 2 FC cluster ALN?
Log2 FC cluter 20.0
0
1
2
3
4
5
6
nlp−20
flp−34
blos−2
nep−16
Y43F8B.2
C50F7.5
lad−2

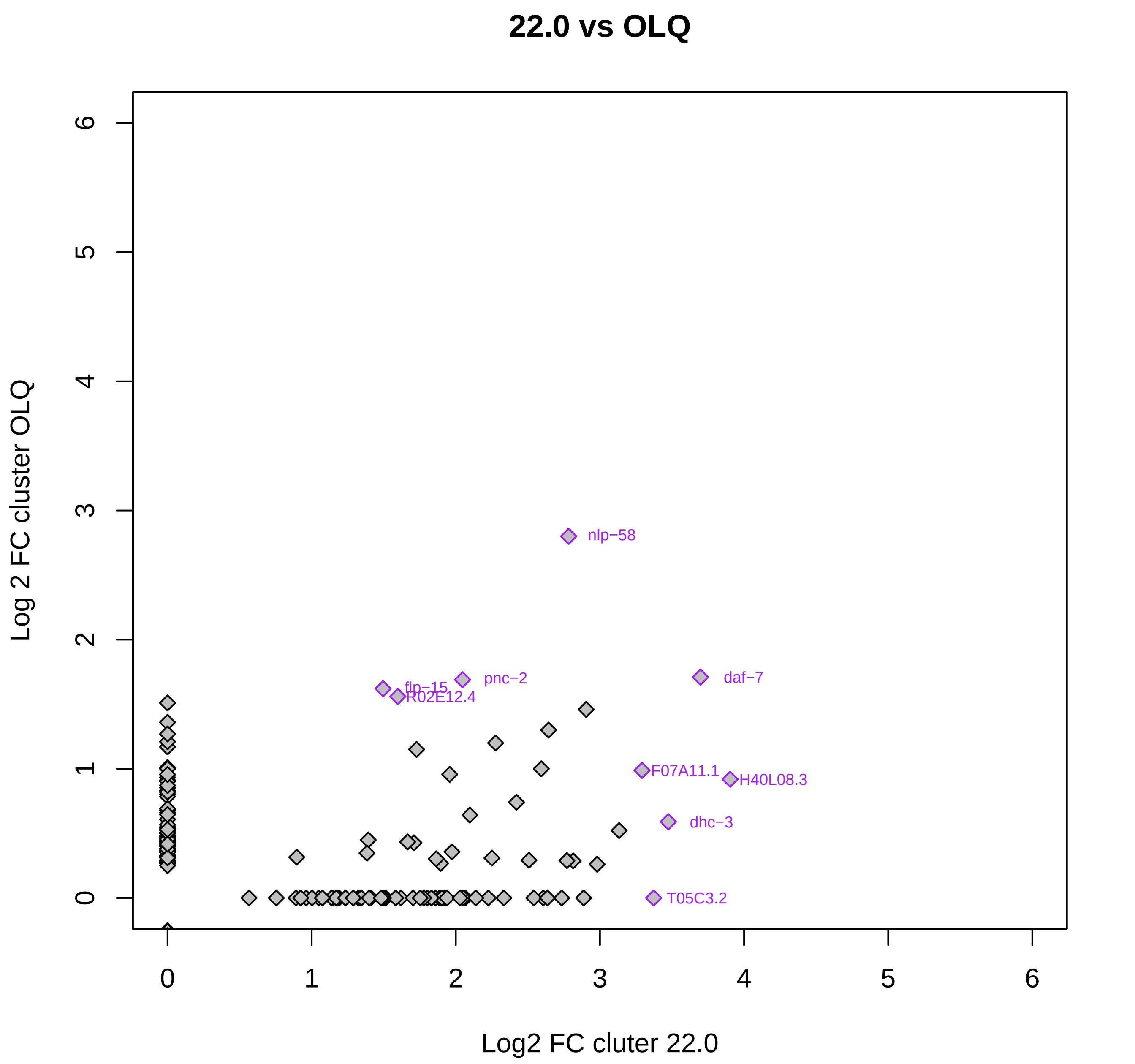

22.0 vs OLQ
Log 2 FC cluster OLQ
Log2 FC cluter 22.0
0
1
2
3
4
5
6
nlp−58
pnc−2
daf−7
flp−15
R02E12.4
F07A11.1
H40L08.3
dhc−3
T05C3.2

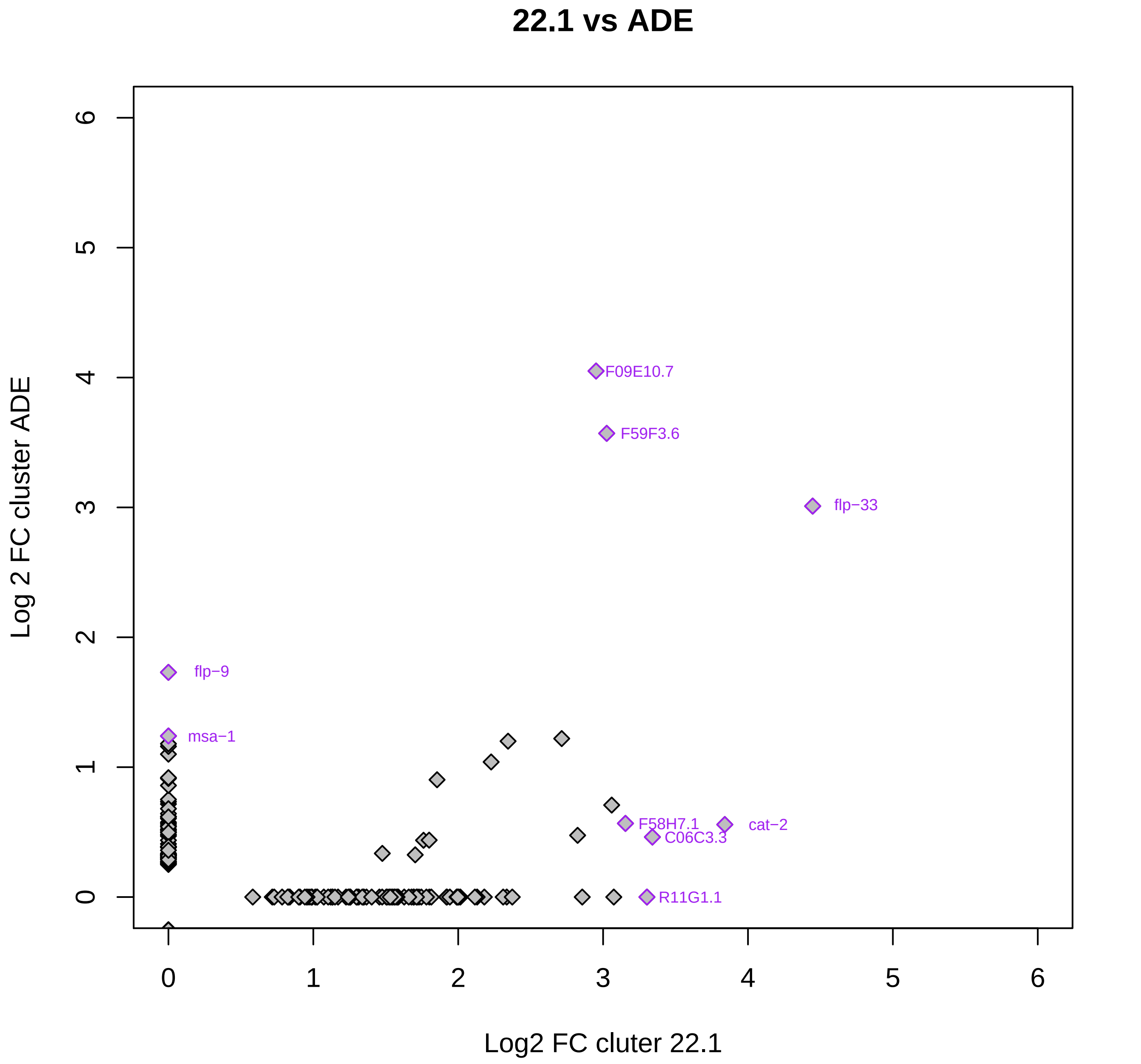

22.1 vs ADE
Log 2 FC cluster ADE
Log2 FC cluter 22.1
0
1
2
3
4
5
6
F09E10.7
F59F3.6
flp−33
flp−9
msa−1
F58H7.1
cat−2
C06C3.3
R11G1.1

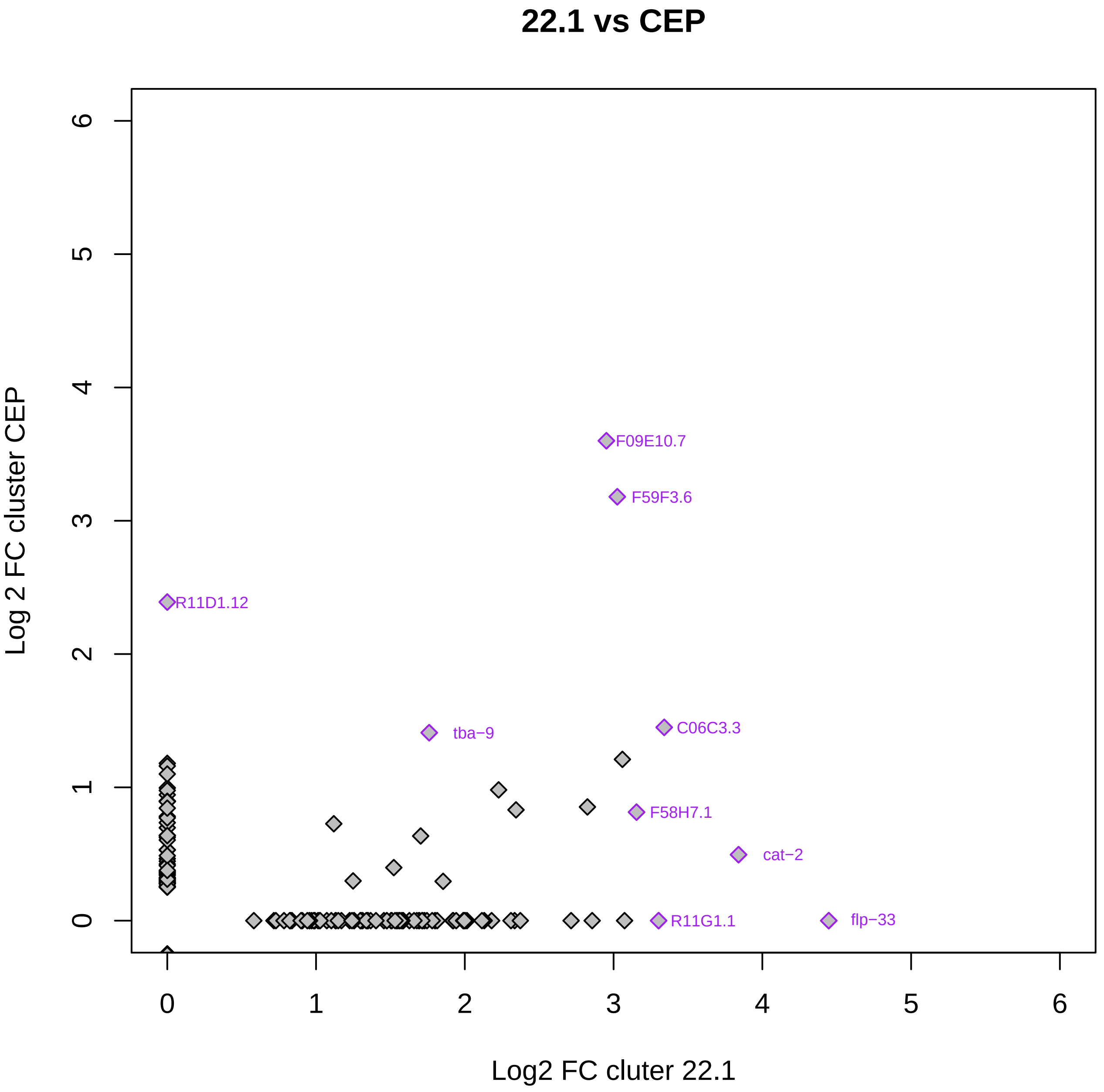
22.1 vs CEP
Log 2 FC cluster CEP
Log2 FC cluter 22.1
0
1
2
3
4
5
6
F09E10.7
F59F3.6
R11D1.12
tba−9
C06C3.3
F58H7.1
cat−2
R11G1.1
flp−33

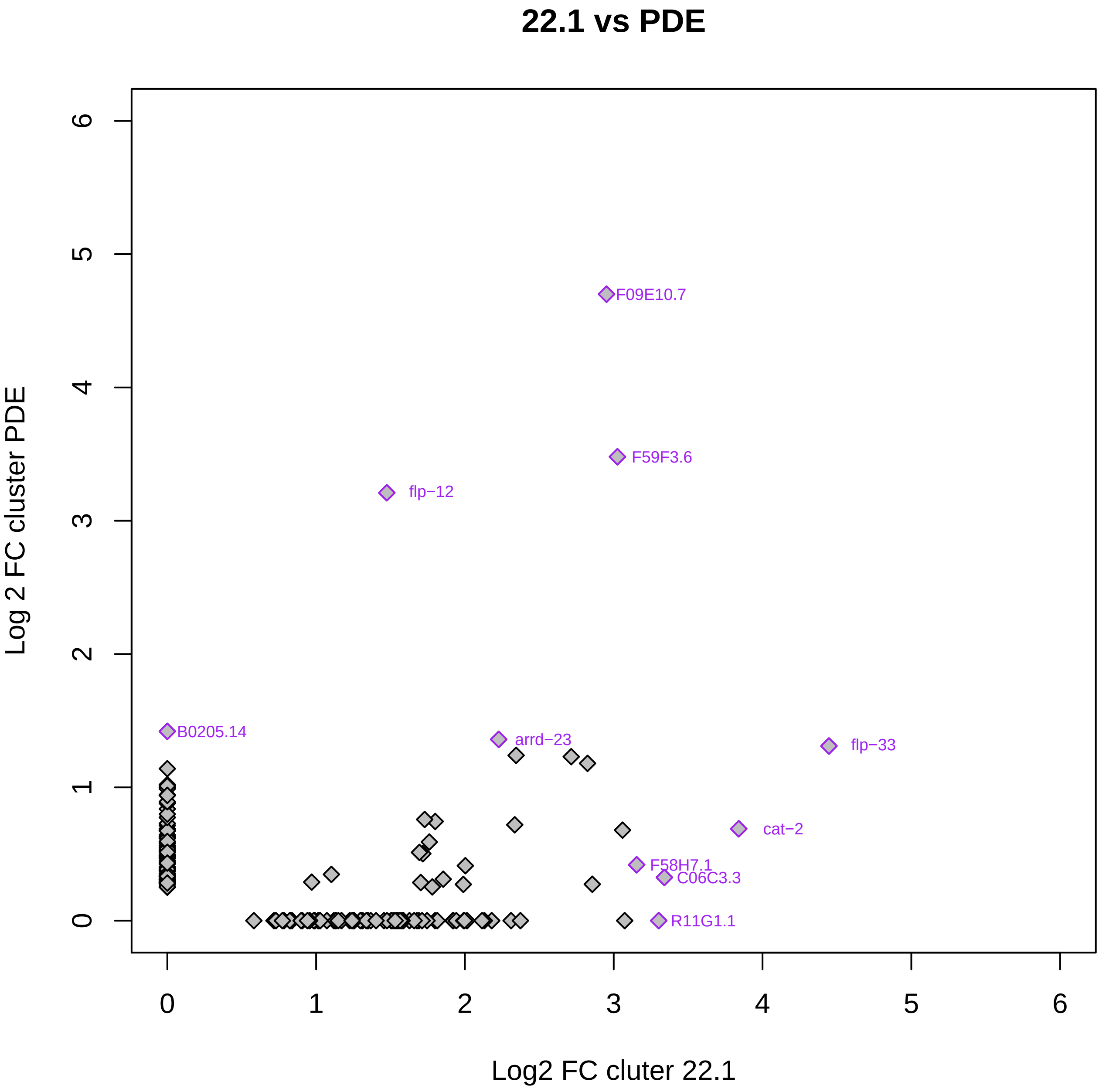
22.1 vs PDE
Log 2 FC cluster PDE
Log2 FC cluter 22.1
0
1
2
3
4
5
6
0
1
2
3
4
5
6
F09E10.7
F59F3.6
flp−12
B0205.14
arrd−23
flp−33
cat−2
F58H7.1
C06C3.3
R11G1.1

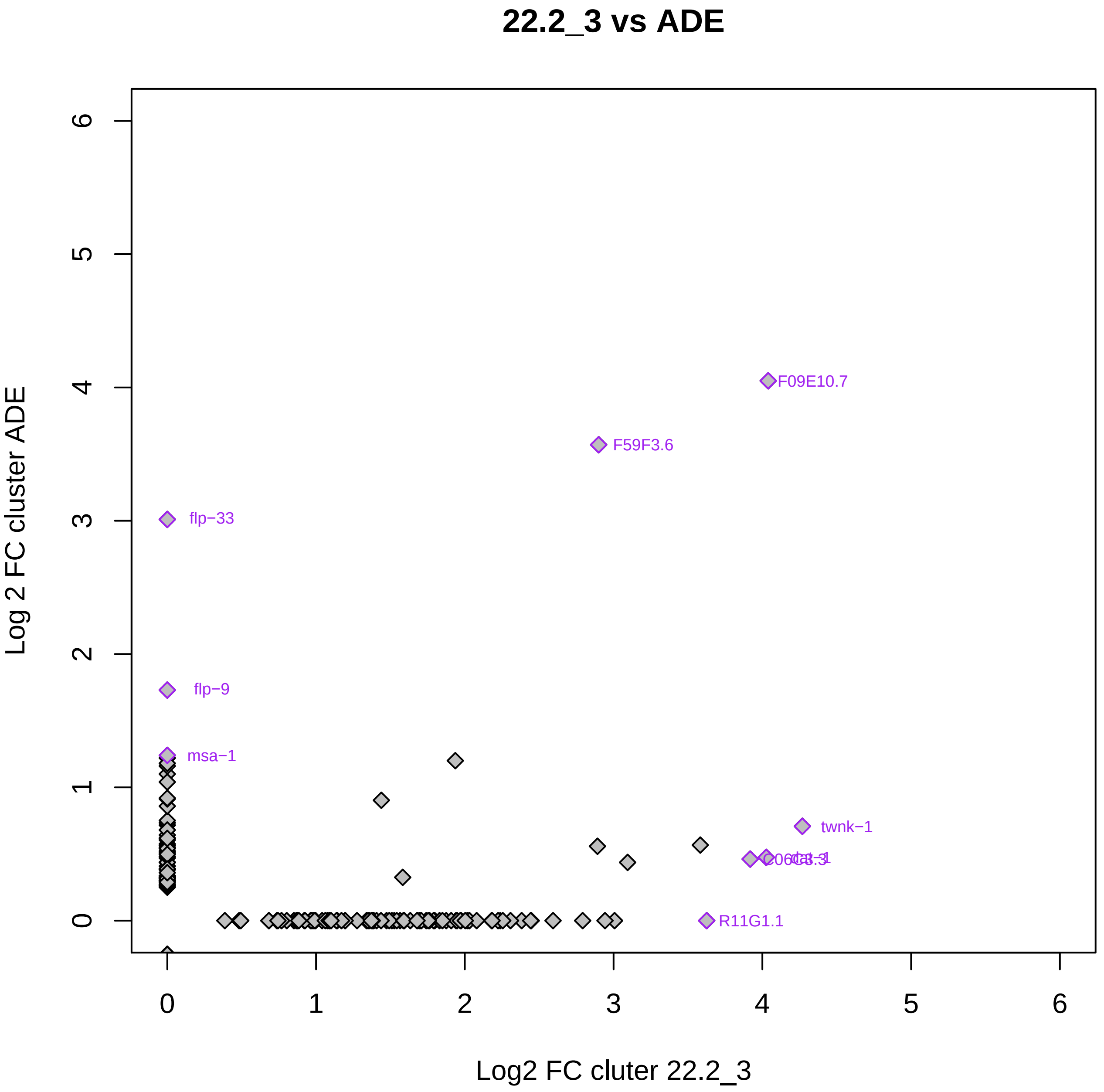
22.2_3 vs ADE
Log 2 FC cluster ADE
Log2 FC cluter 22.2_3
0
1
2
3
4
5
6
F09E10.7
F59F3.6
flp−33
flp−9
msa−1
twnk−1
R11G1.1

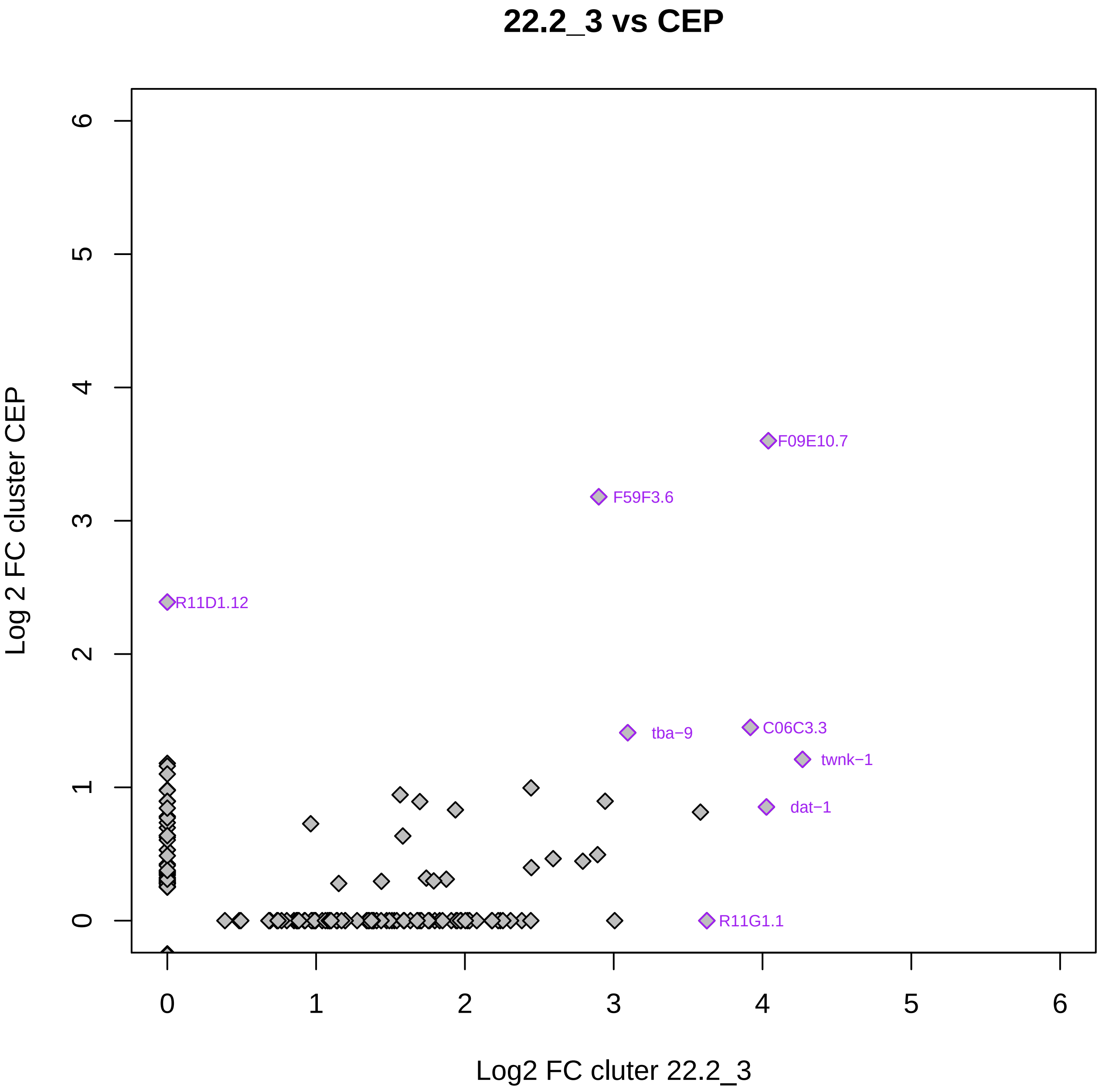

22.2_3 vs CEP
Log 2 FC cluster CEP
Log2 FC cluter 22.2_3
0
1
2
3
4
5
6
F09E10.7
F59F3.6
R11D1.12
tba−9
C06C3.3
twnk−1
dat−1
R11G1.1

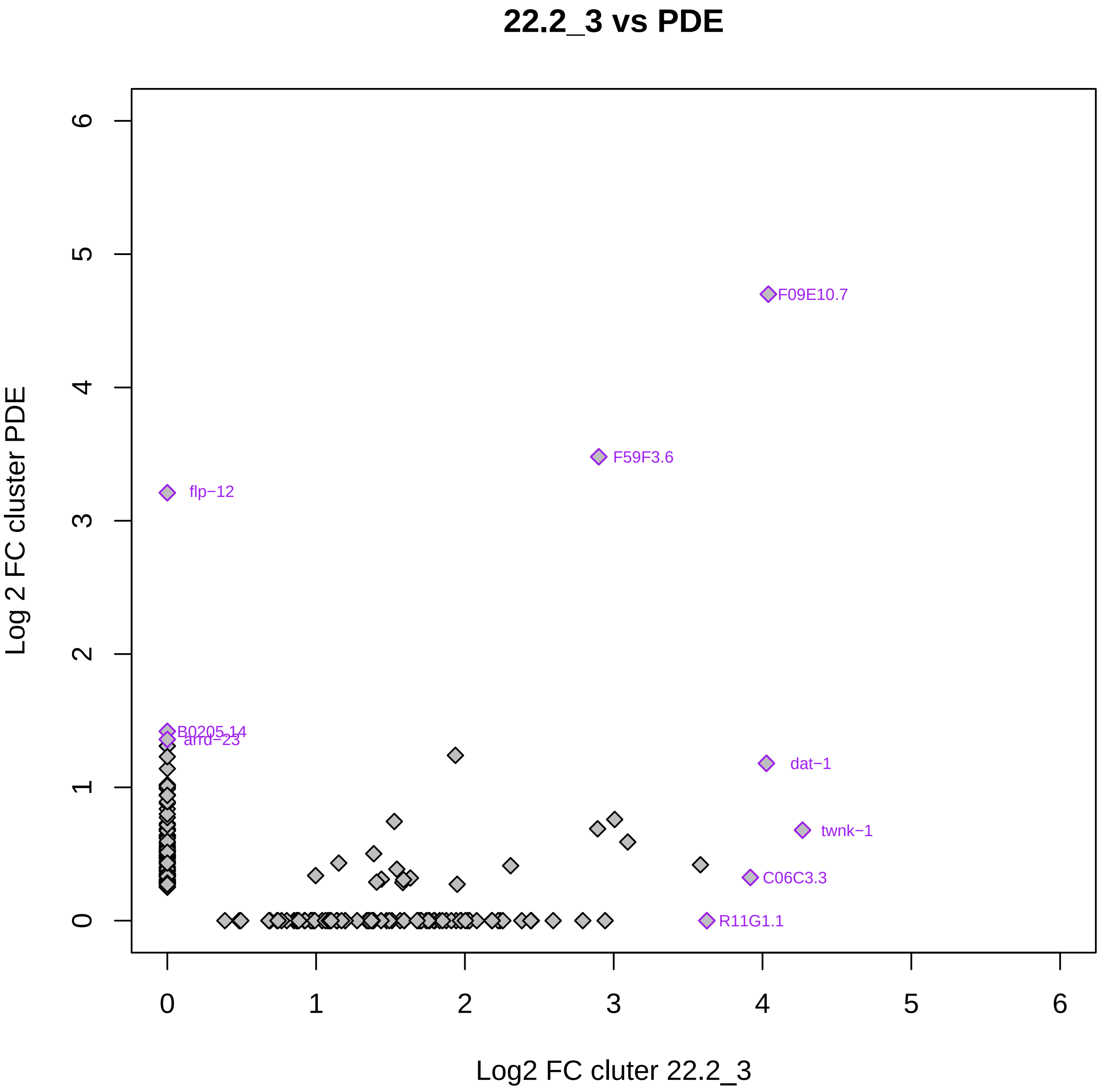
22.2_3 vs PDE
Log 2 FC cluster PDE
Log2 FC cluter 22.2_3
0
1
2
3
4
5
6
F09E10.7
F59F3.6
flp−12
B0205.14
arrd−23
dat−1
twnk−1
C06C3.3
R11G1.1

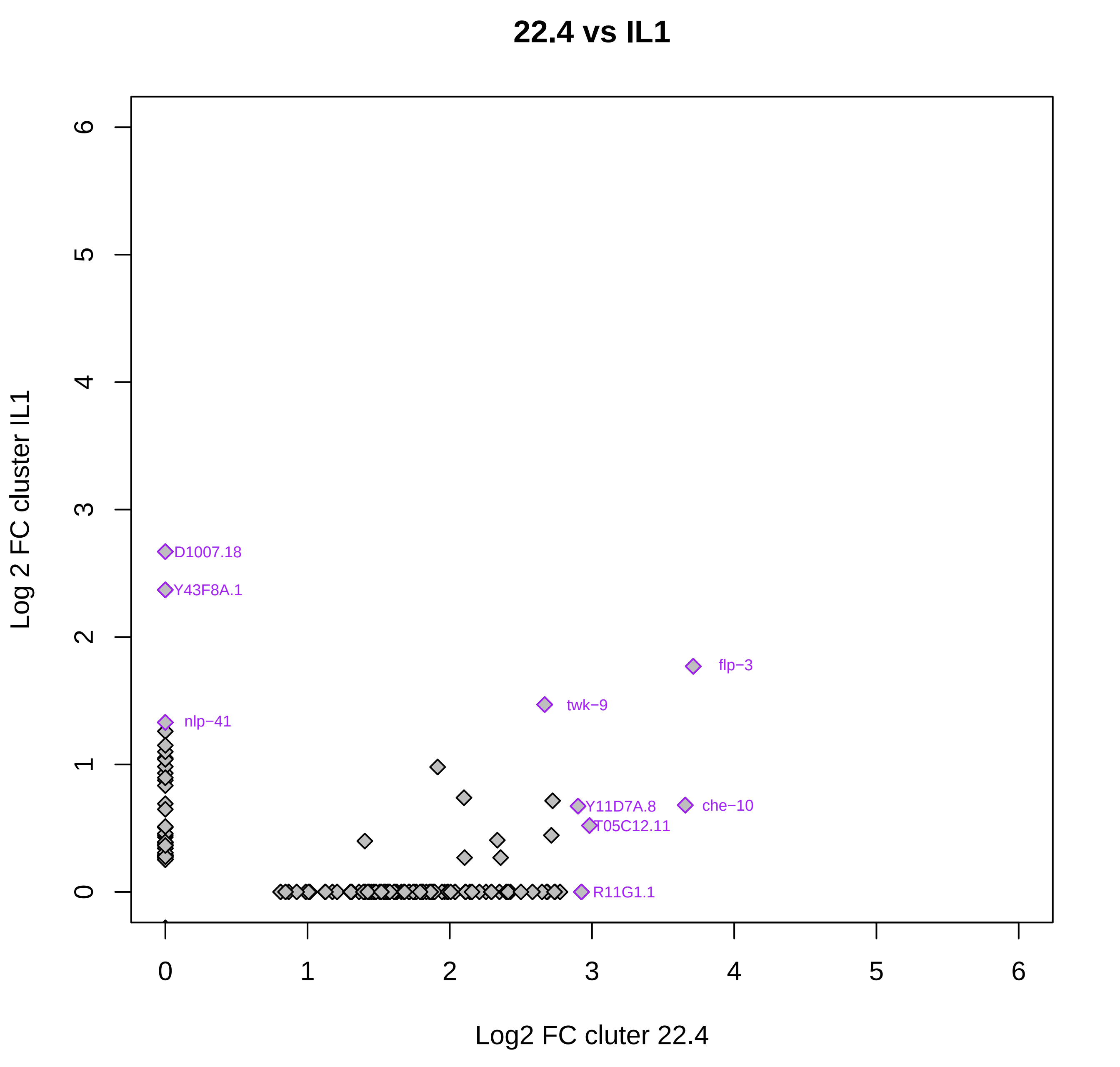

22.4 vs IL1
Log 2 FC cluster IL1
Log2 FC cluter 22.4
0
1
2
3
4
5
6
D1007.18
Y43F8A.1
flp−3
twk−9
nlp−41
Y11D7A.8
che−10
T05C12.11
R11G1.1

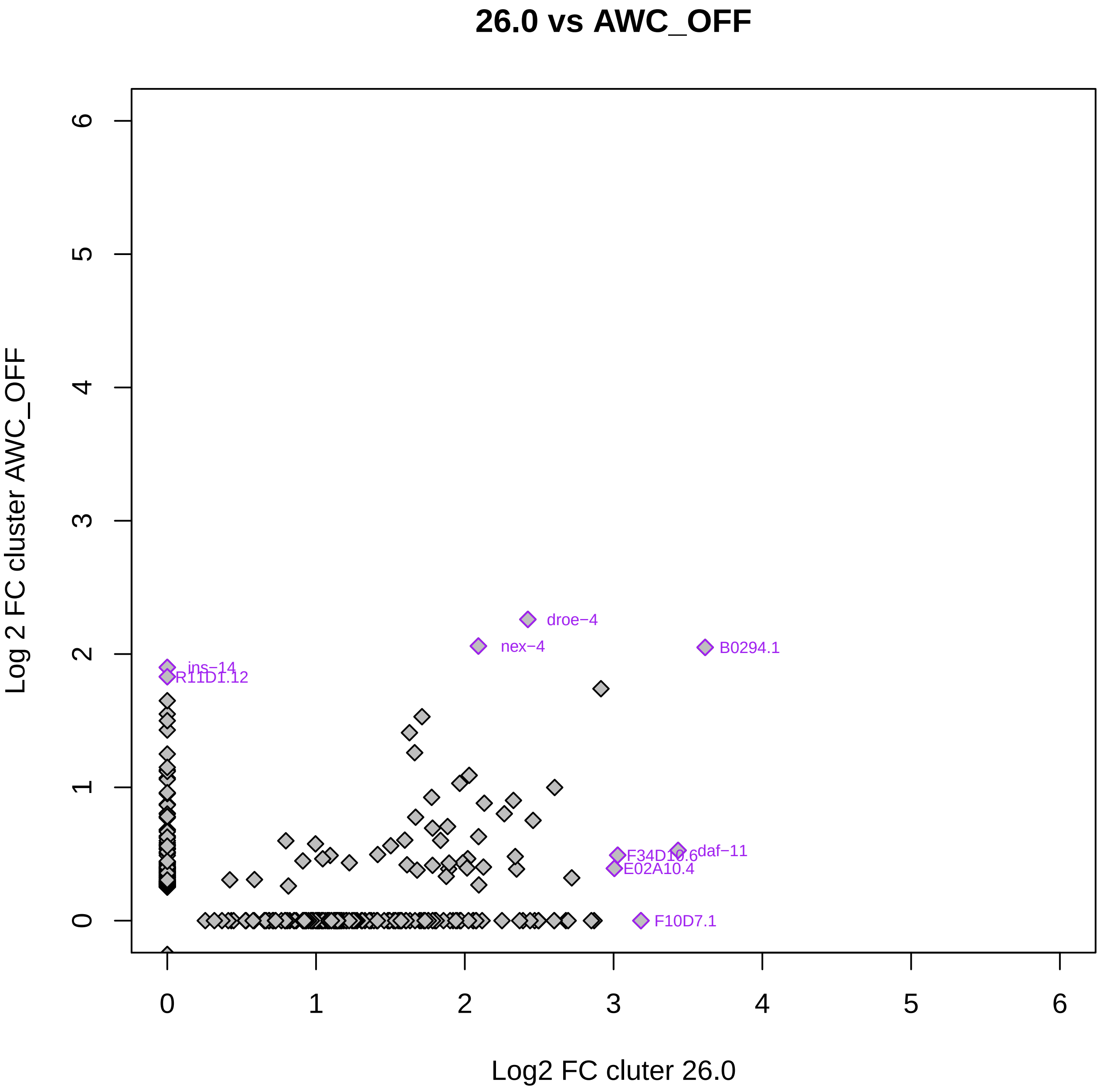

26.0 vs AWC_OFF
Log 2 FC cluster AWC_OFF
Log2 FC cluter 26.0
0
1
2
3
4
5
6
droe−4
nex−4
B0294.1
ins−14
R11D1.12
F34D10.6
daf−11
E02A10.4
F10D7.1

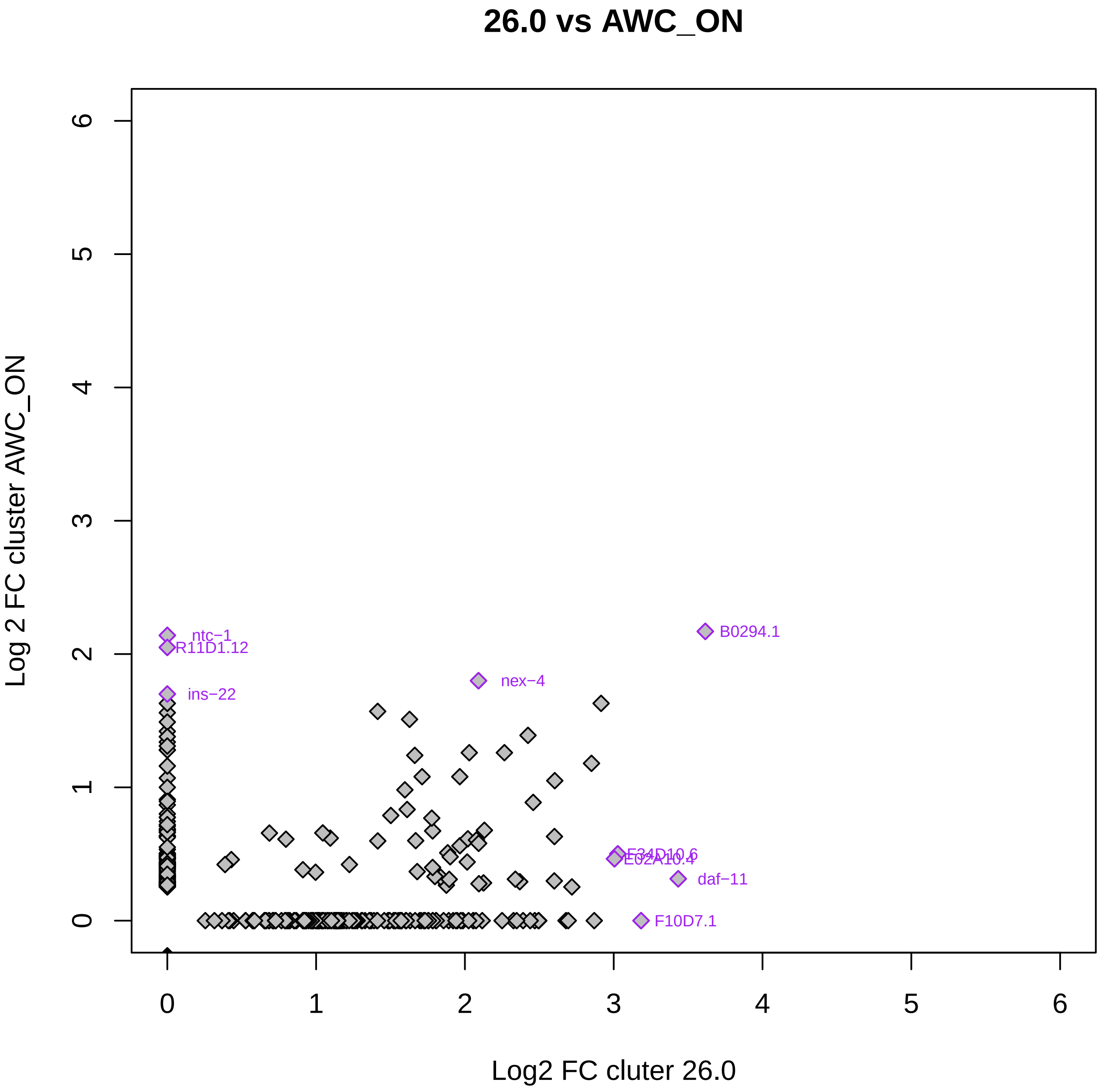

26.0 vs AWC_ON
Log 2 FC cluster AWC_ON
Log2 FC cluter 26.0
0
1
2
3
4
5
6
ntc−1
R11D1.12
ins−22
B0294.1
nex−4
F34D10.6
E02A10.4
daf−11
F10D7.1

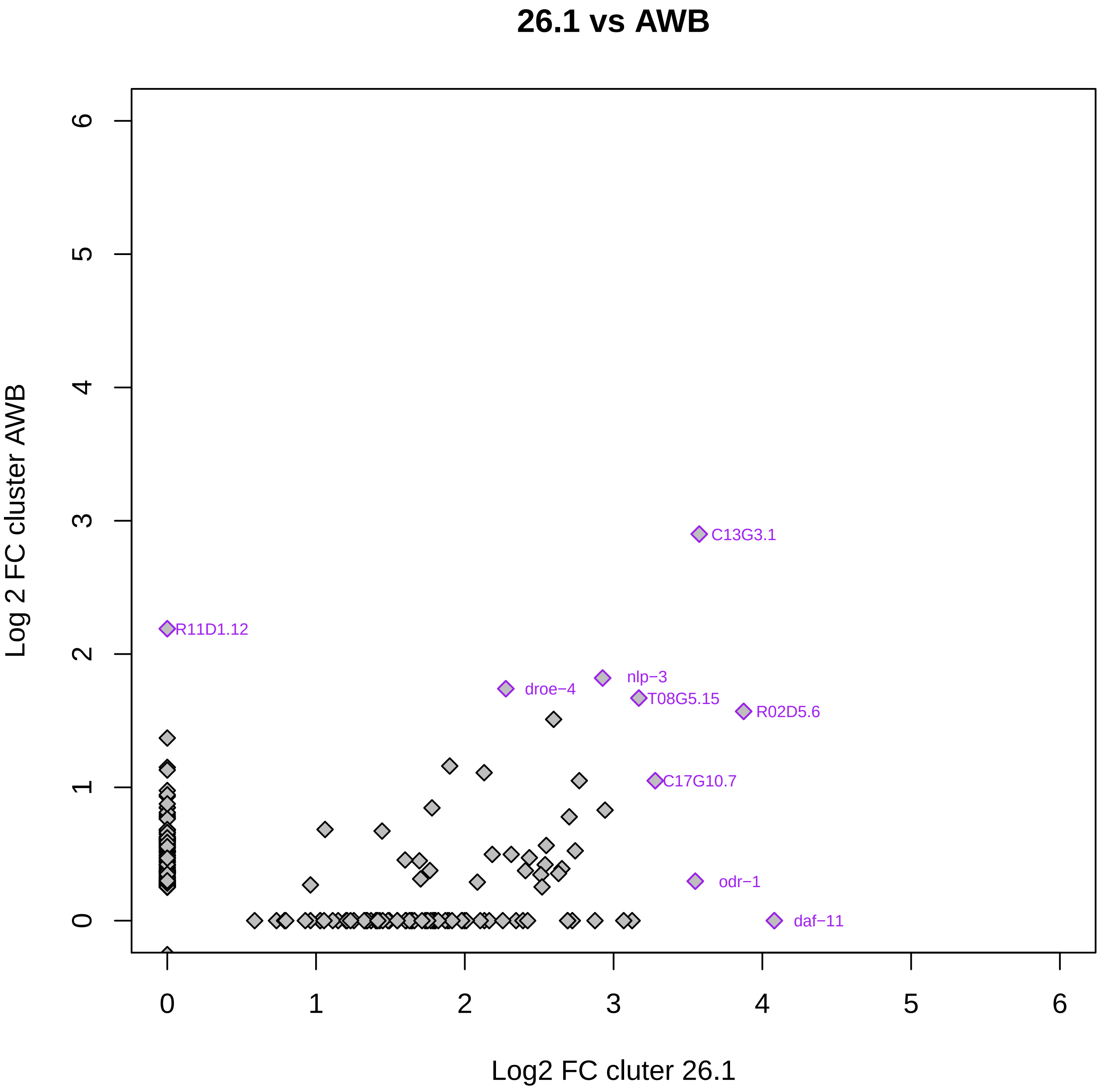
26.1 vs AWB
Log 2 FC cluster AWB
Log2 FC cluter 26.1
0
1
2
3
4
5
6
C13G3.1
R11D1.12
droe−4
nlp−3
T08G5.15
R02D5.6
C17G10.7
odr−1
daf−11

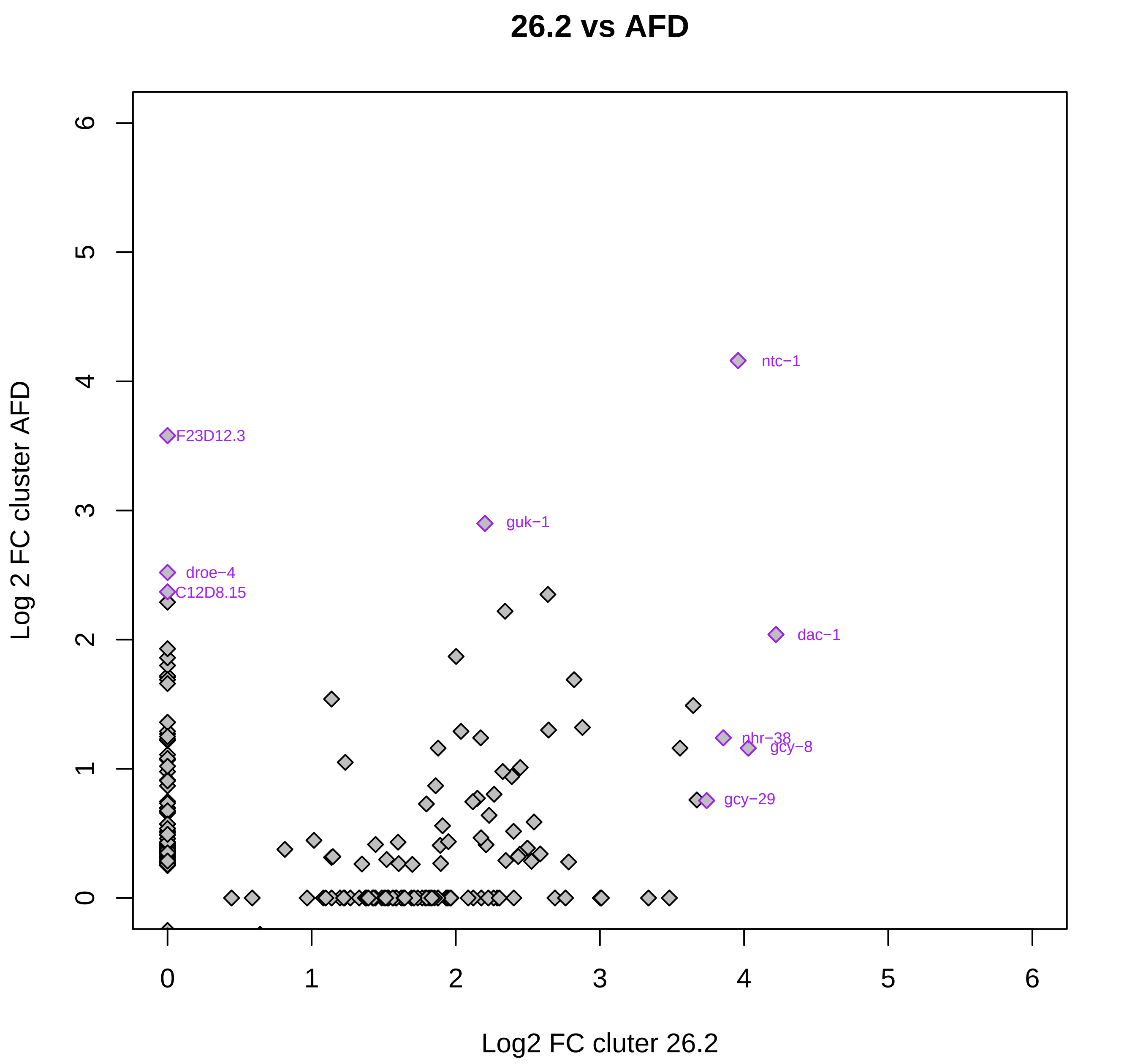
26.2 vs AFD
Log 2 FC cluster AFD
Log2 FC cluter 26.2
0
1
2
3
4
5
6
ntc−1
F23D12.3
guk−1
droe−4
C12D8.15
dac−1
nhr−38
gcy−8
gcy−29

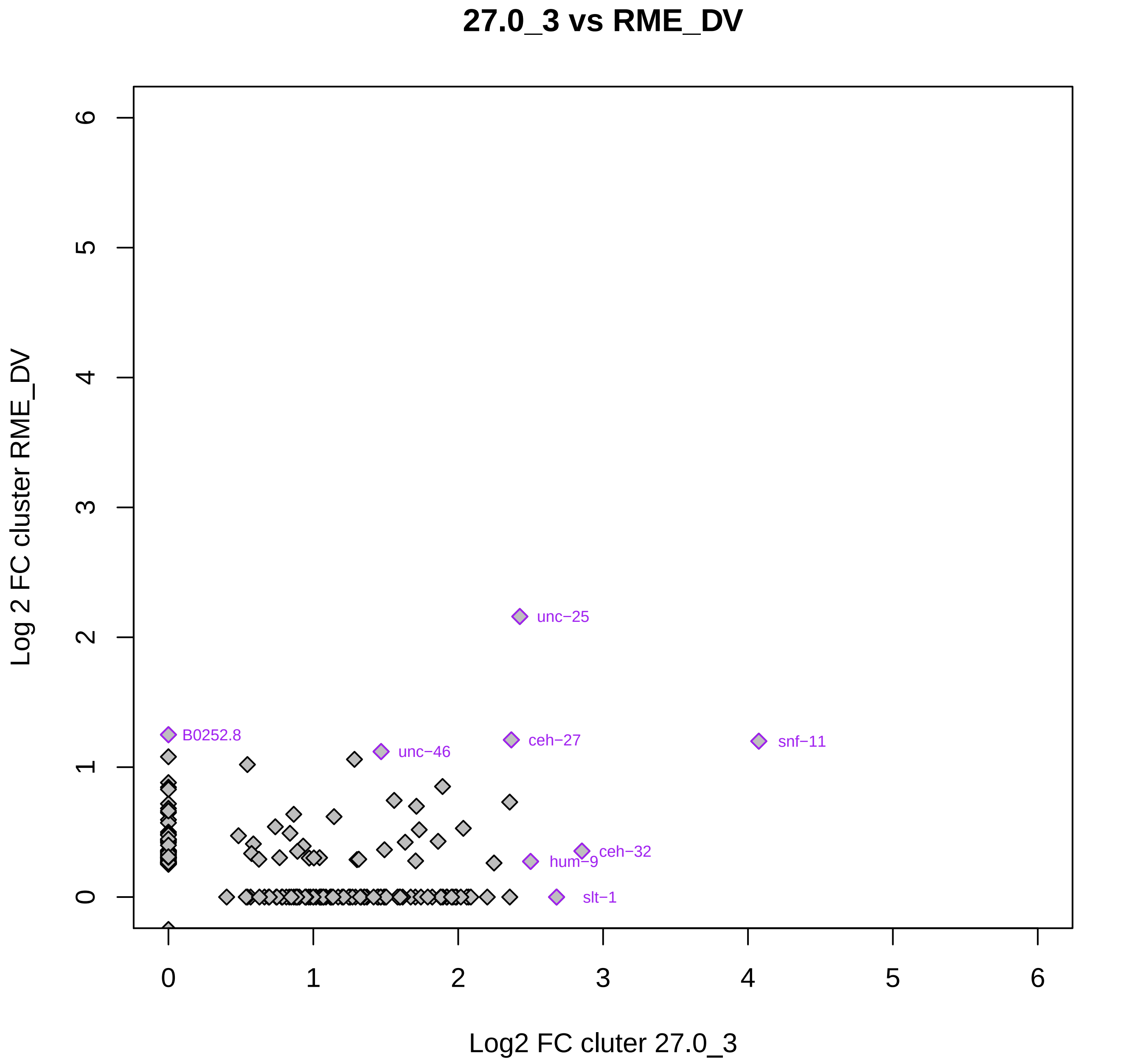

27.0_3 vs RME_DV
Log 2 FC cluster RME_DV
Log2 FC cluter 27.0_3
0
1
2
3
4
5
6
unc−25
B0252.8
unc−46
ceh−27
snf−11
hum−9
ceh−32
slt−1

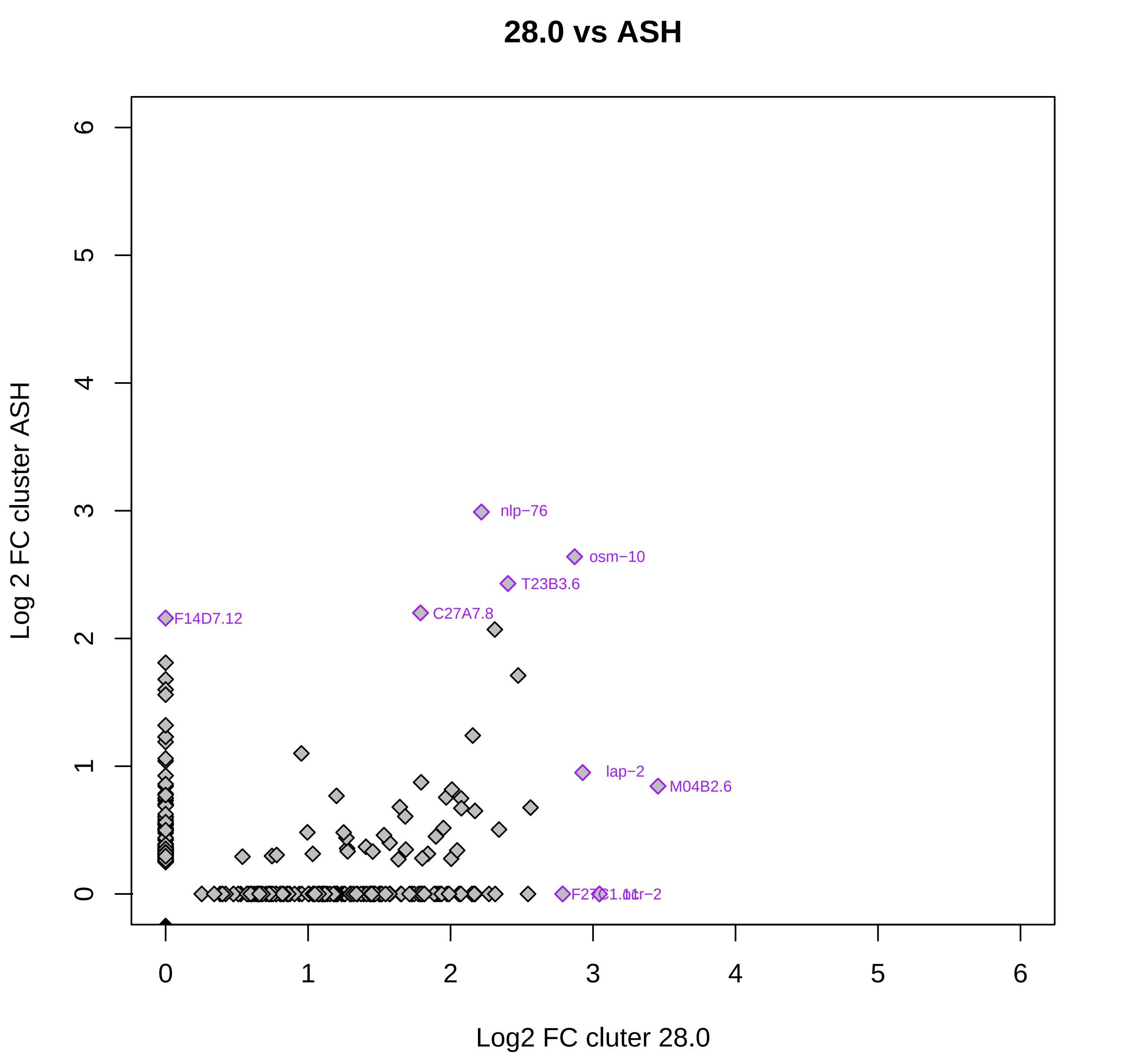
28.0 vs ASH
Log 2 FC cluster ASH
Log2 FC cluter 28.0
0
1
2
3
4
5
6
nlp−76
osm−10
T23B3.6
C27A7.8
F14D7.12
lap−2
M04B2.6
F27C1.11
ttr−2

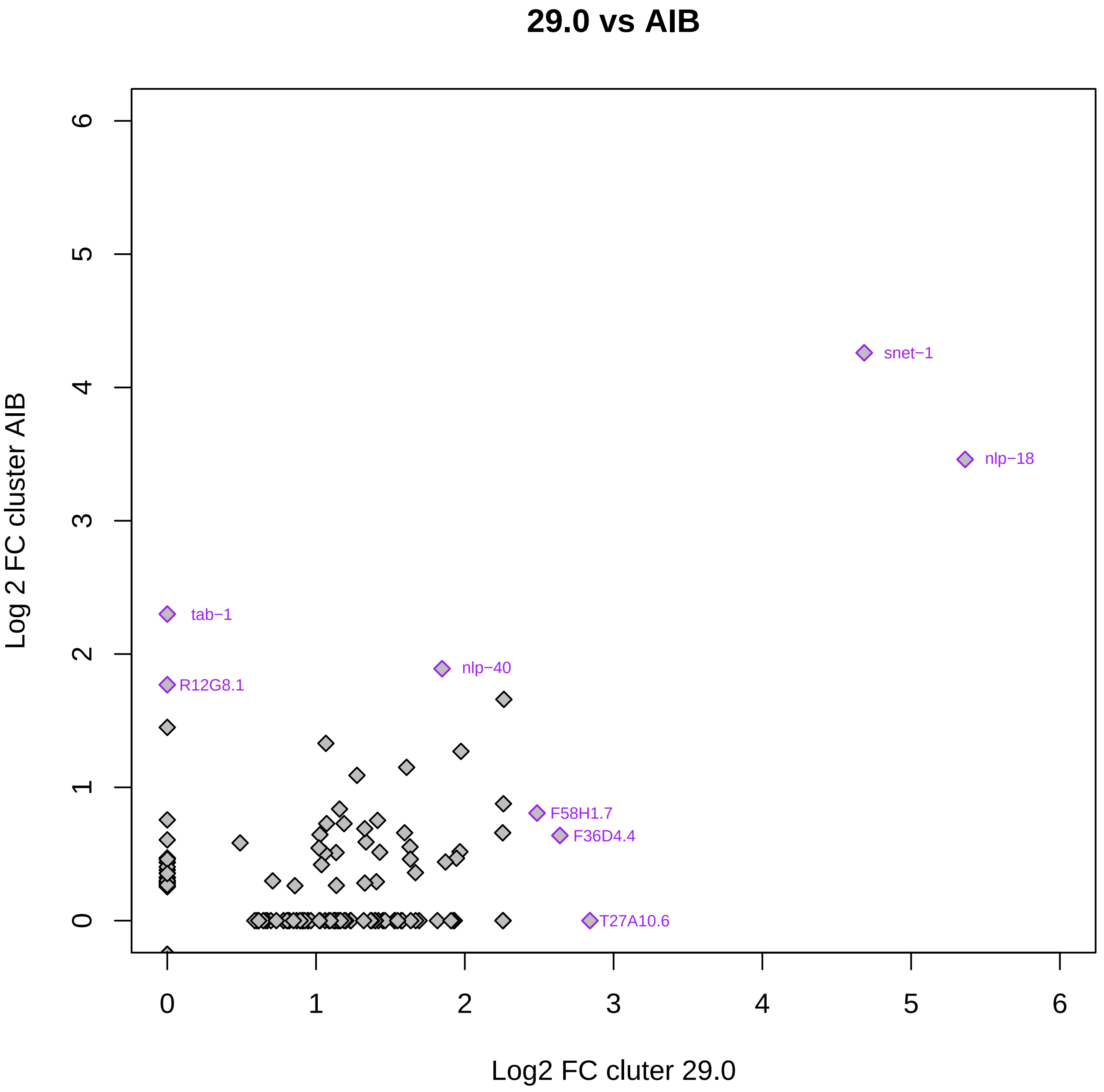
29.0 vs AIB
Log 2 FC cluster AIB
Log2 FC cluter 29.0
0
1
2
3
4
5
6
snet−1
nlp−18
tab−1
R12G8.1
nlp−40
F58H1.7
F36D4.4
T27A10.6

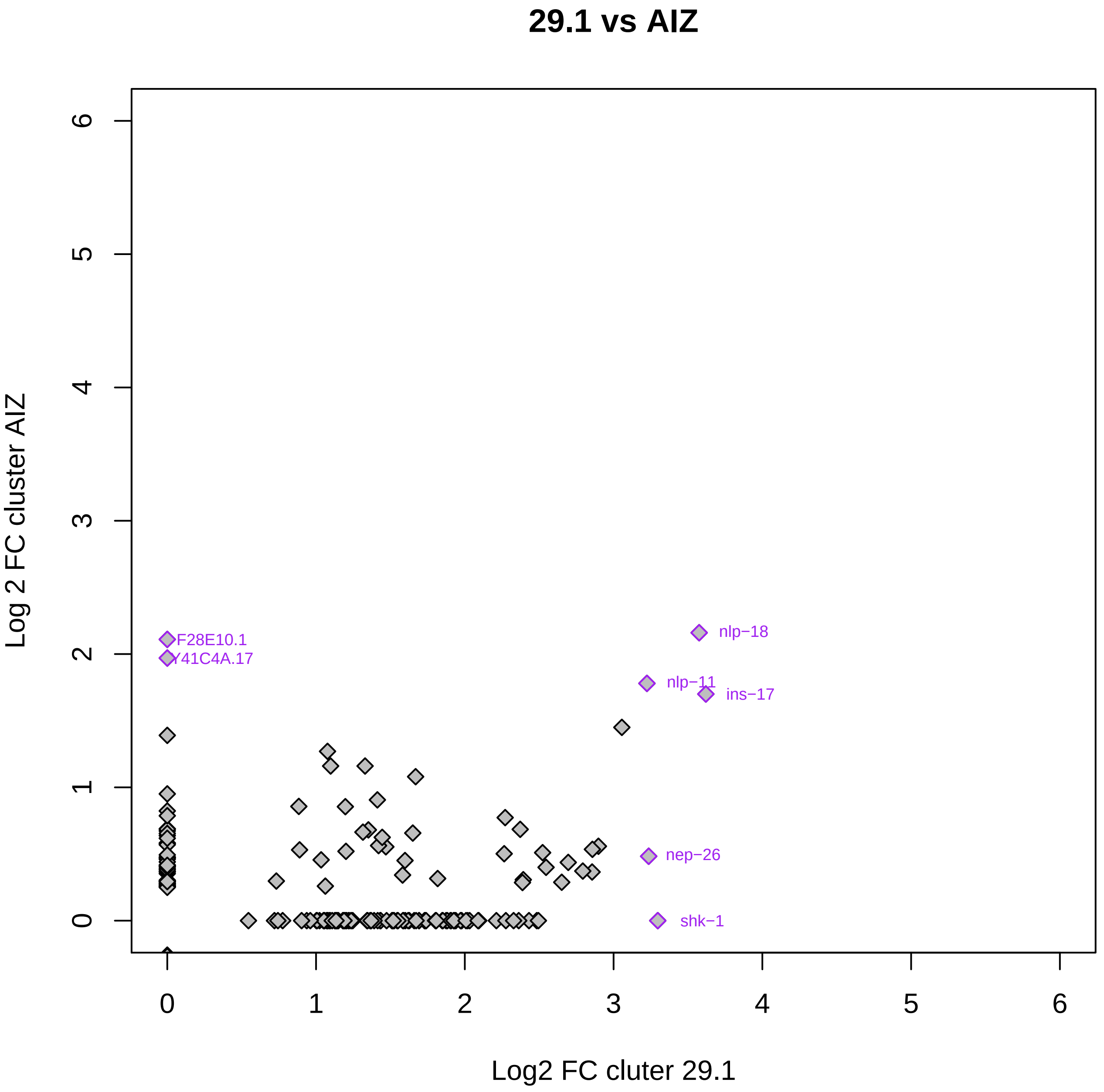
29.1 vs AIZ
Log 2 FC cluster AIZ
Log2 FC cluter 29.1
0
1
2
3
4
5
6
F28E10.1
Y41C4A.17
nlp−18
nlp−11
ins−17
nep−26
shk−1

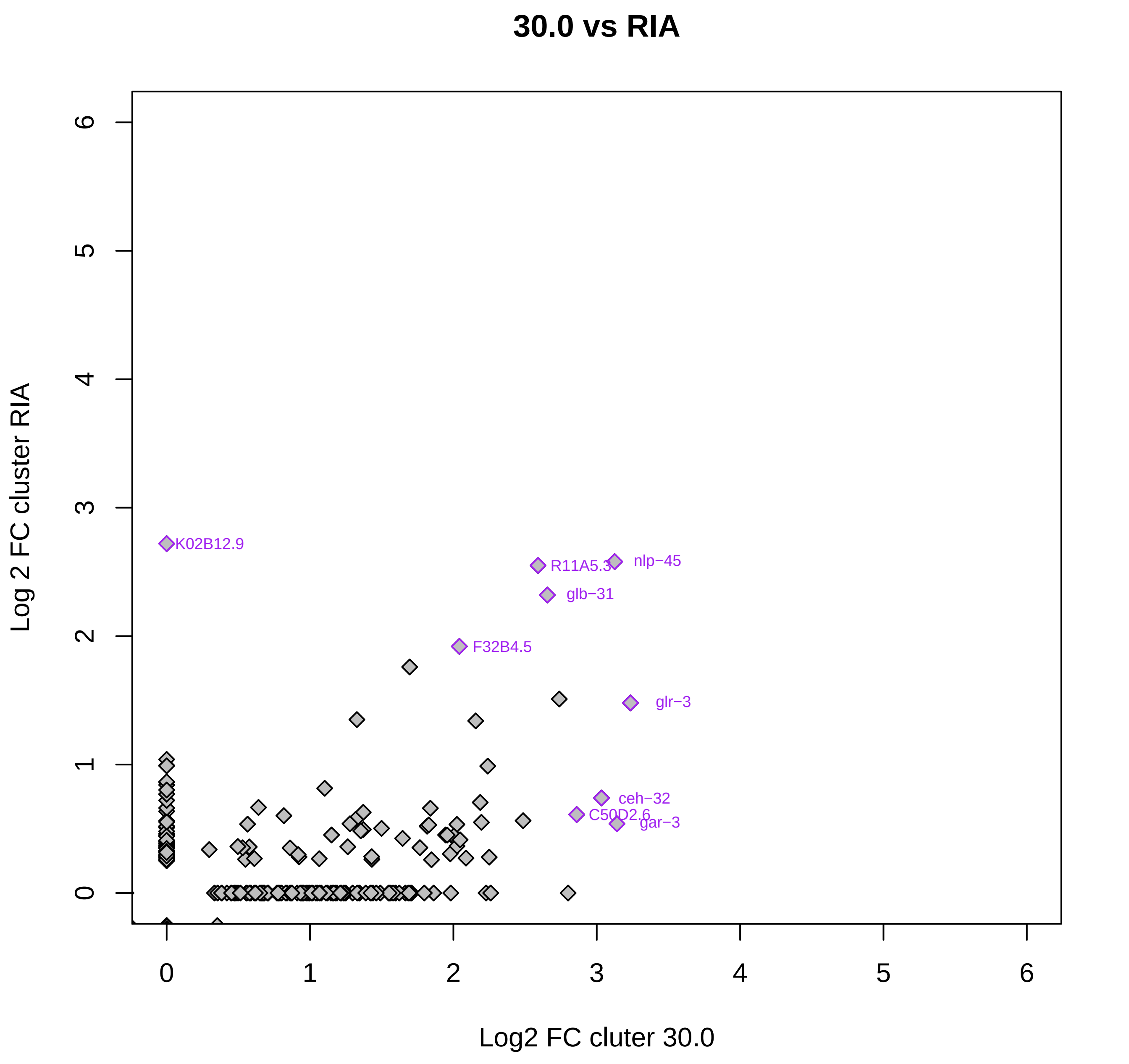
30.0 vs RIA
Log 2 FC cluster RIA
Log2 FC cluter 30.0
0
1
2
3
4
5
6
K02B12.9
R11A5.3
nlp−45
glb−31
F32B4.5
glr−3
ceh−32
C50D2.6
gar−3

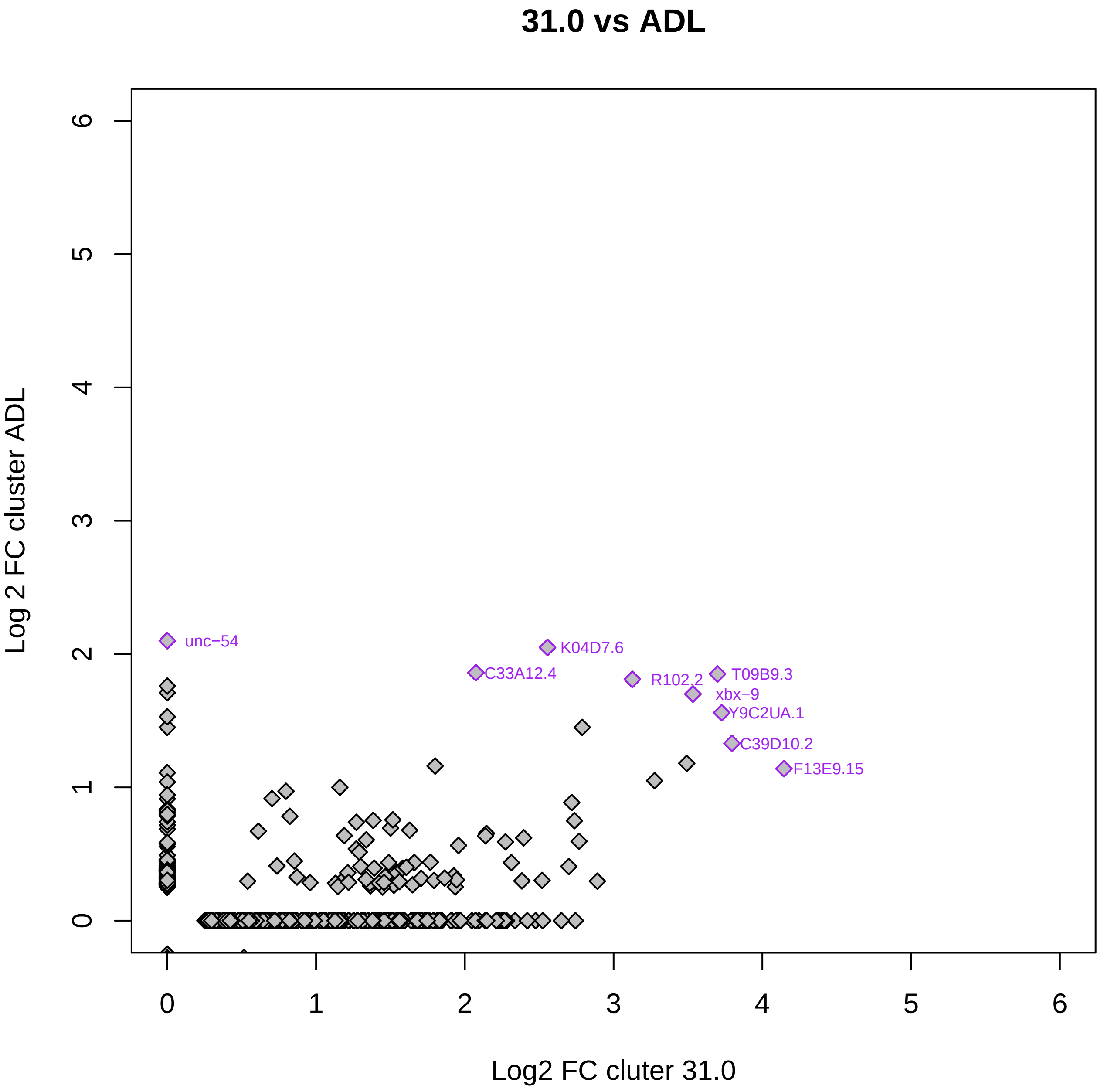

31.0 vs ADL
Log 2 FC cluster ADL
Log2 FC cluter 31.0
0
1
2
3
4
5
6
unc−54
K04D7.6
C33A12.4
R102.2
T09B9.3
xbx−9
Y9C2UA.1
C39D10.2
F13E9.15

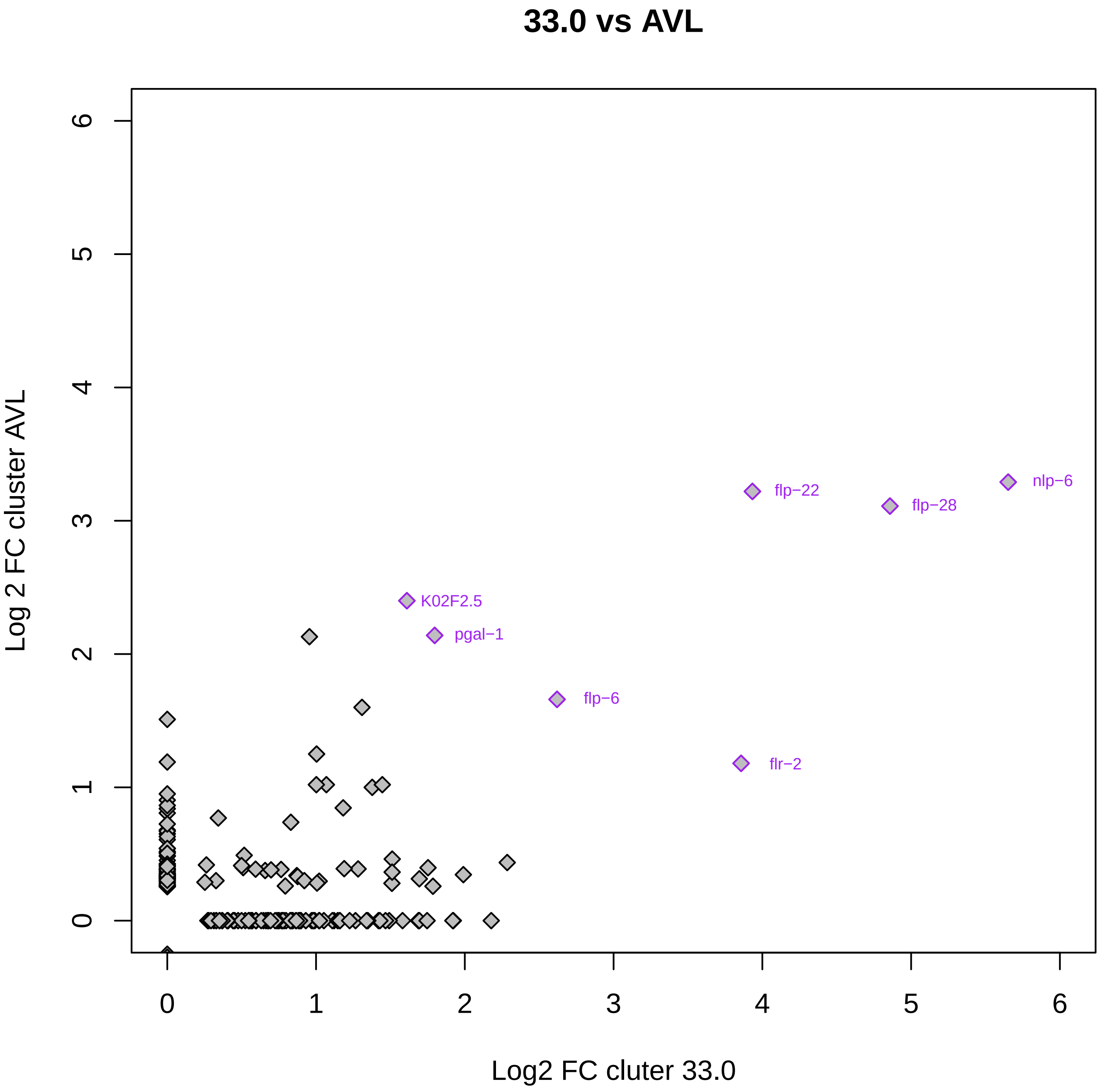

33.0 vs AVL
Log 2 FC cluster AVL
Log2 FC cluter 33.0
0
1
2
3
4
5
6
nlp−6
flp−22
flp−28
K02F2.5
pgal−1
flp−6
flr−2

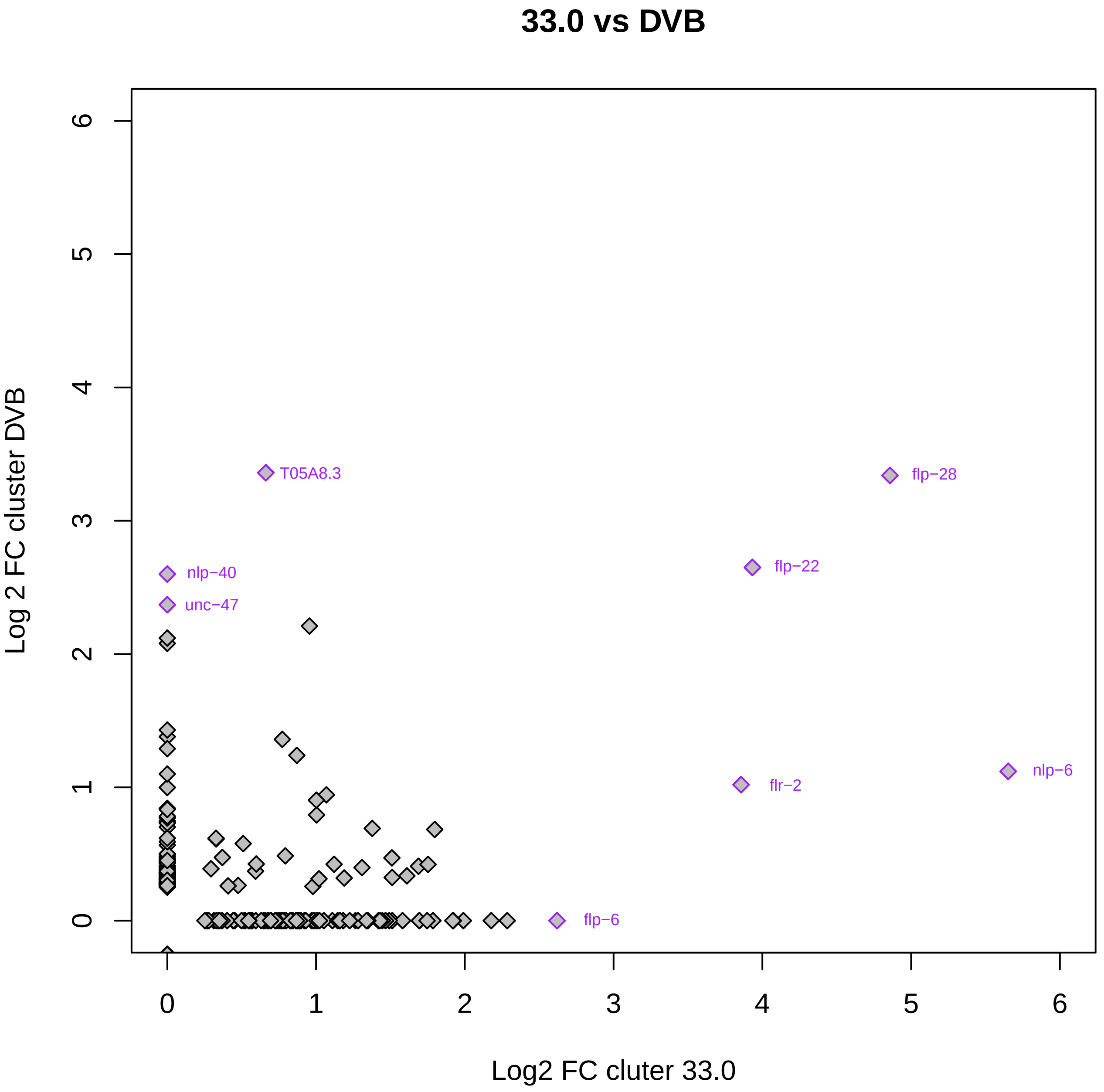

33.0 vs DVB
Log 2 FC cluster DVB
Log2 FC cluter 33.0
0
1
2
3
4
5
6
T05A8.3
flp−28
flp−22
nlp−40
unc−47
flr−2
nlp−6
flp−6

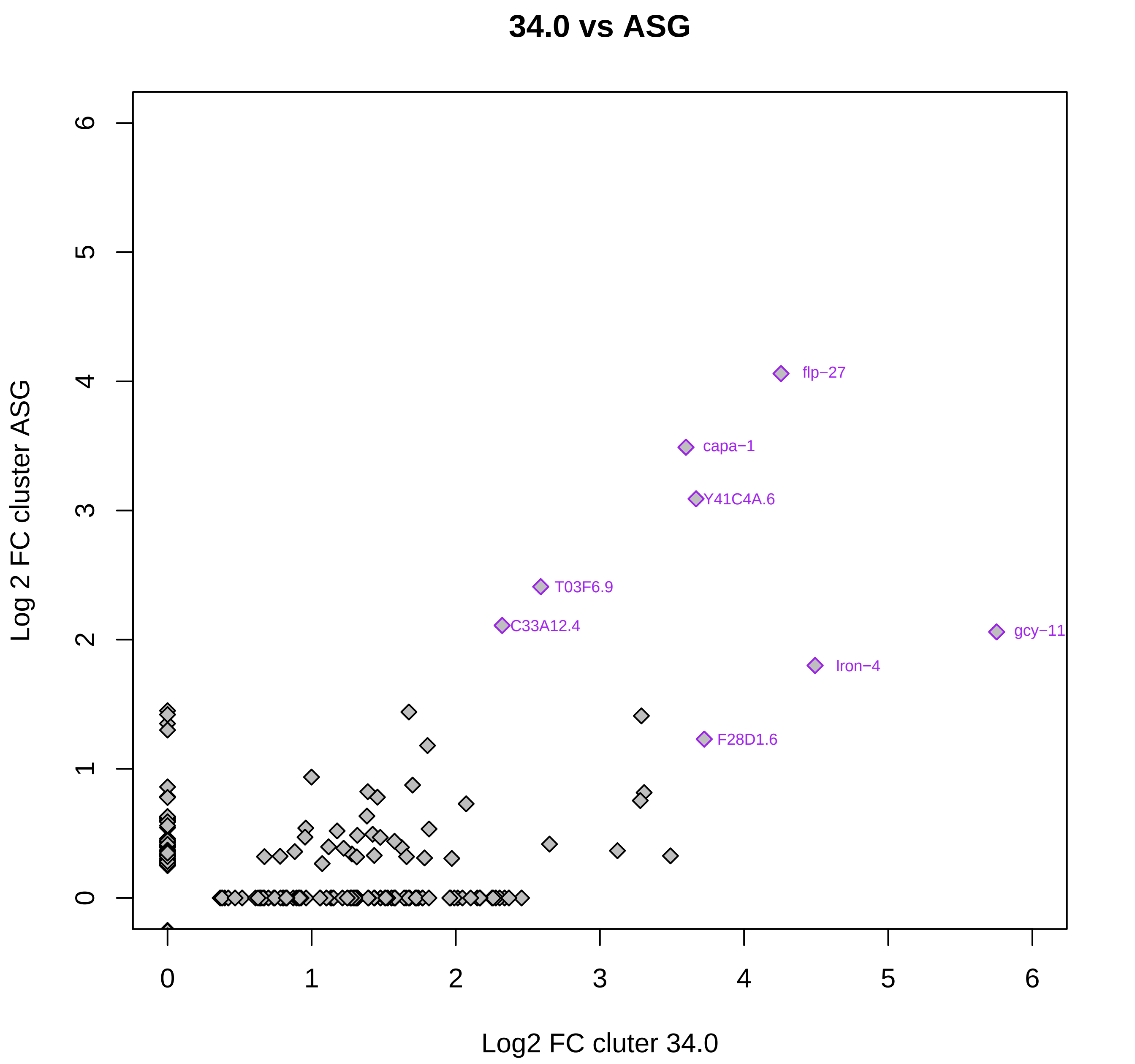
34.0 vs ASG
Log 2 FC cluster ASG
Log2 FC cluter 34.0
0
1
2
3
4
5
6
flp−27
capa−1
Y41C4A.6
T03F6.9
C33A12.4
gcy−11
lron−4
F28D1.6

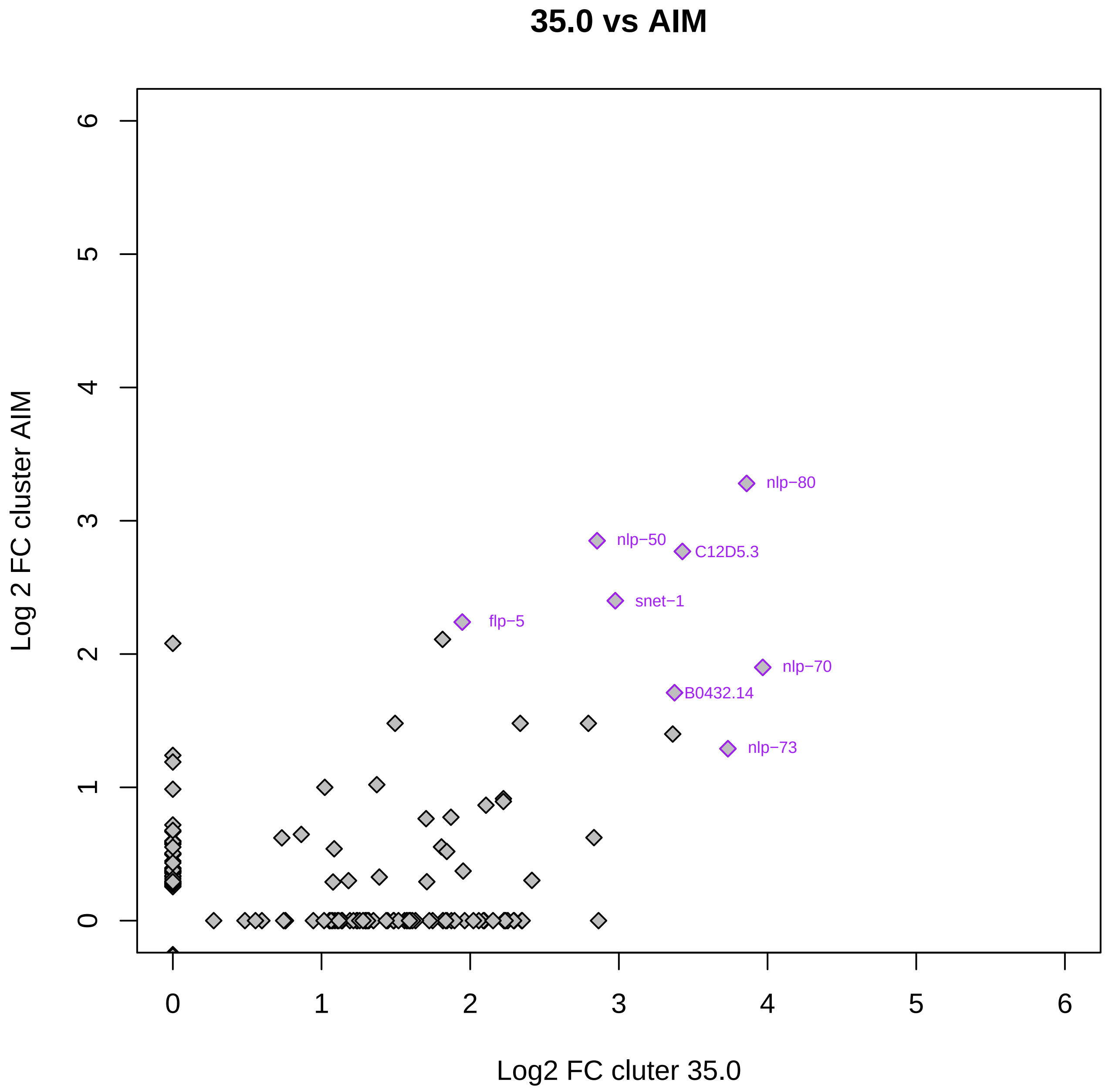

35.0 vs AIM
Log 2 FC cluster AIM
Log2 FC cluter 35.0
0
1
2
3
4
5
6
nlp−80
nlp−50
C12D5.3
snet−1
flp−5
nlp−70
B0432.14
nlp−73

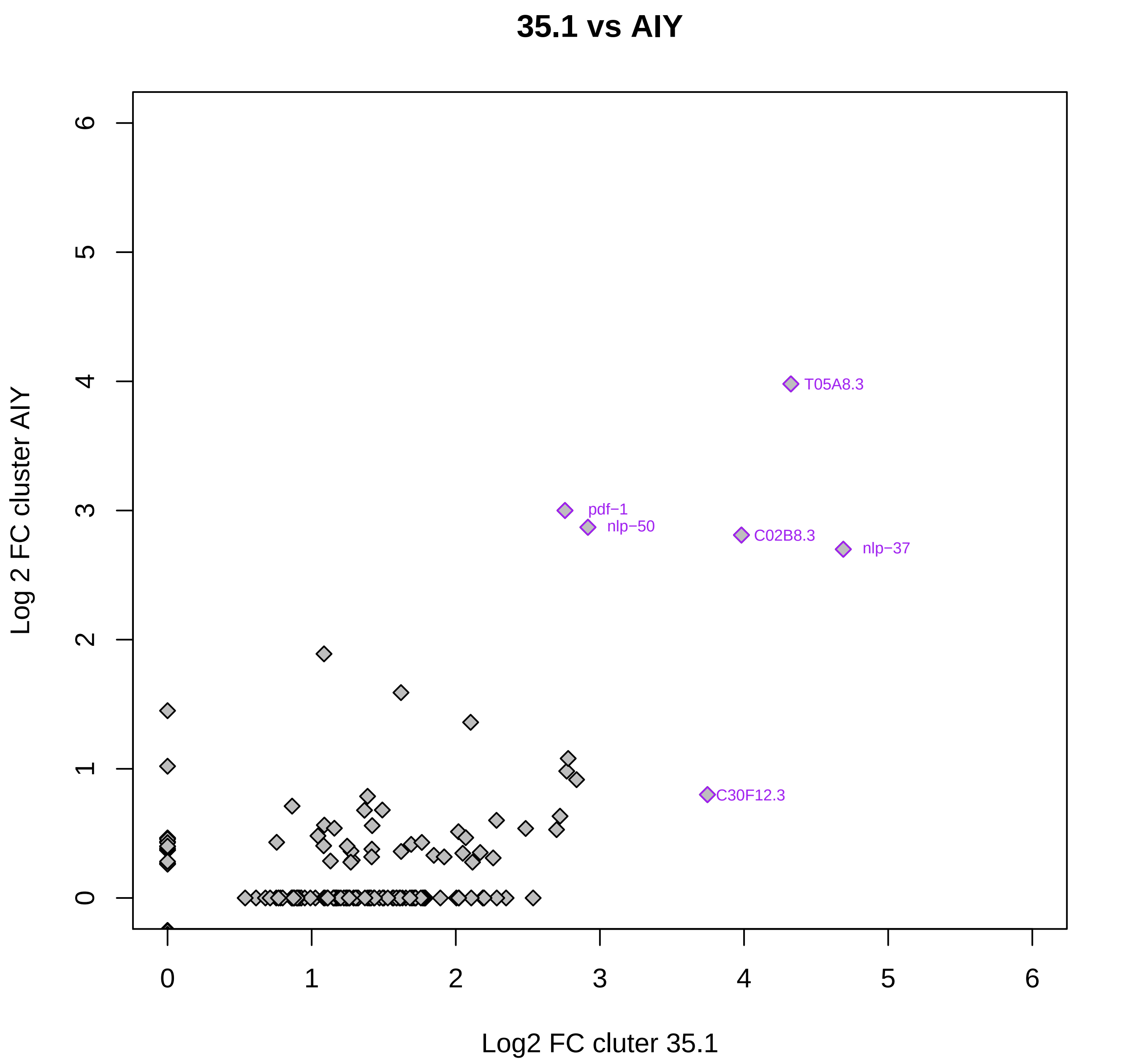
35.1 vs AIY
T05A8.3
pdf−1
nlp−50
C02B8.3
nlp−37
C30F12.3
Log 2 FC cluster AIY
Log2 FC cluter 35.1
0
1
2
3
4
5
6

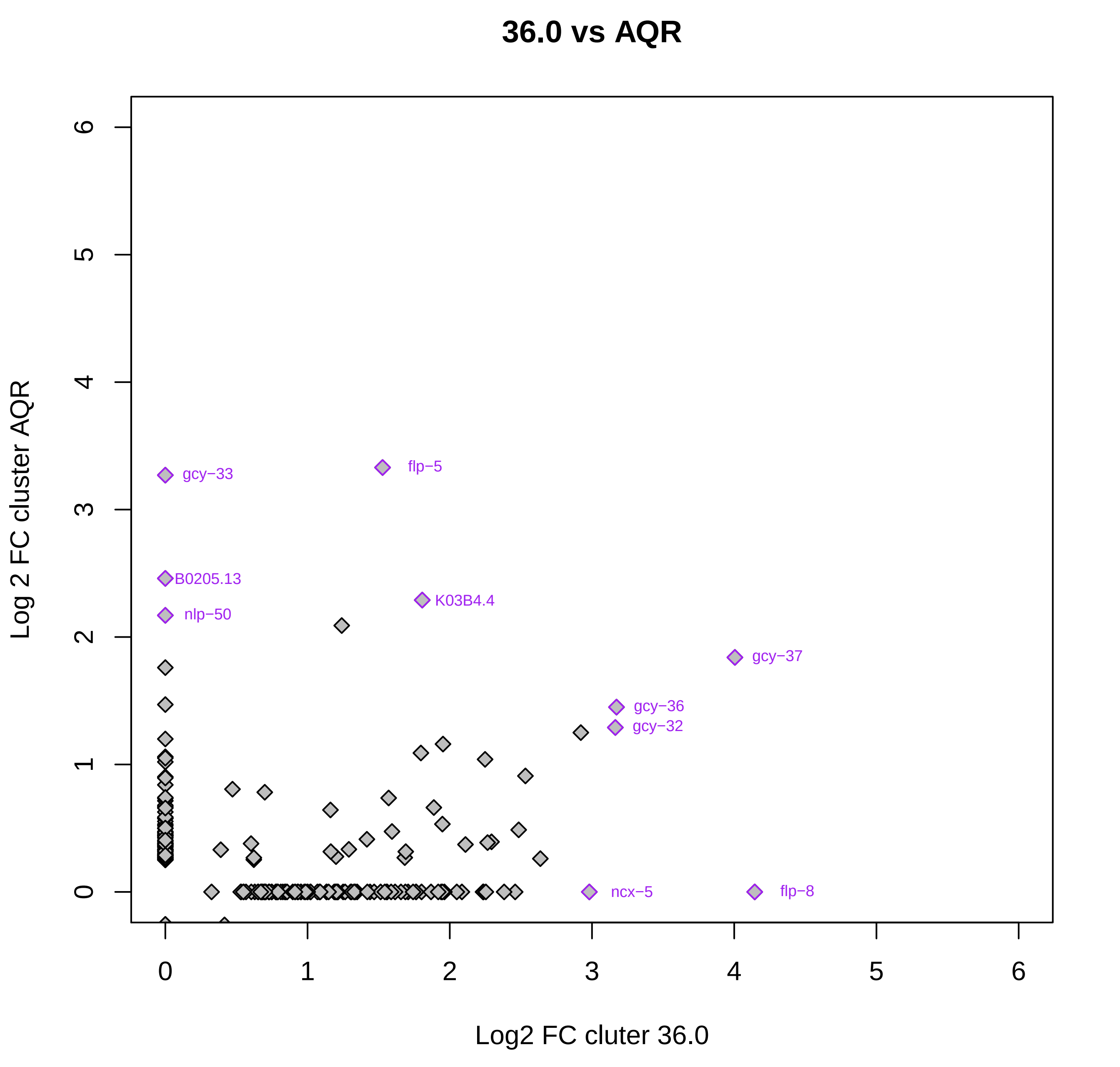
36.0 vs AQR
Log 2 FC cluster AQR
Log2 FC cluter 36.0
0
1
2
3
4
5
6
gcy−33
flp−5
B0205.13
K03B4.4
nlp−50
gcy−37
gcy−36
gcy−32
ncx−5
flp−8

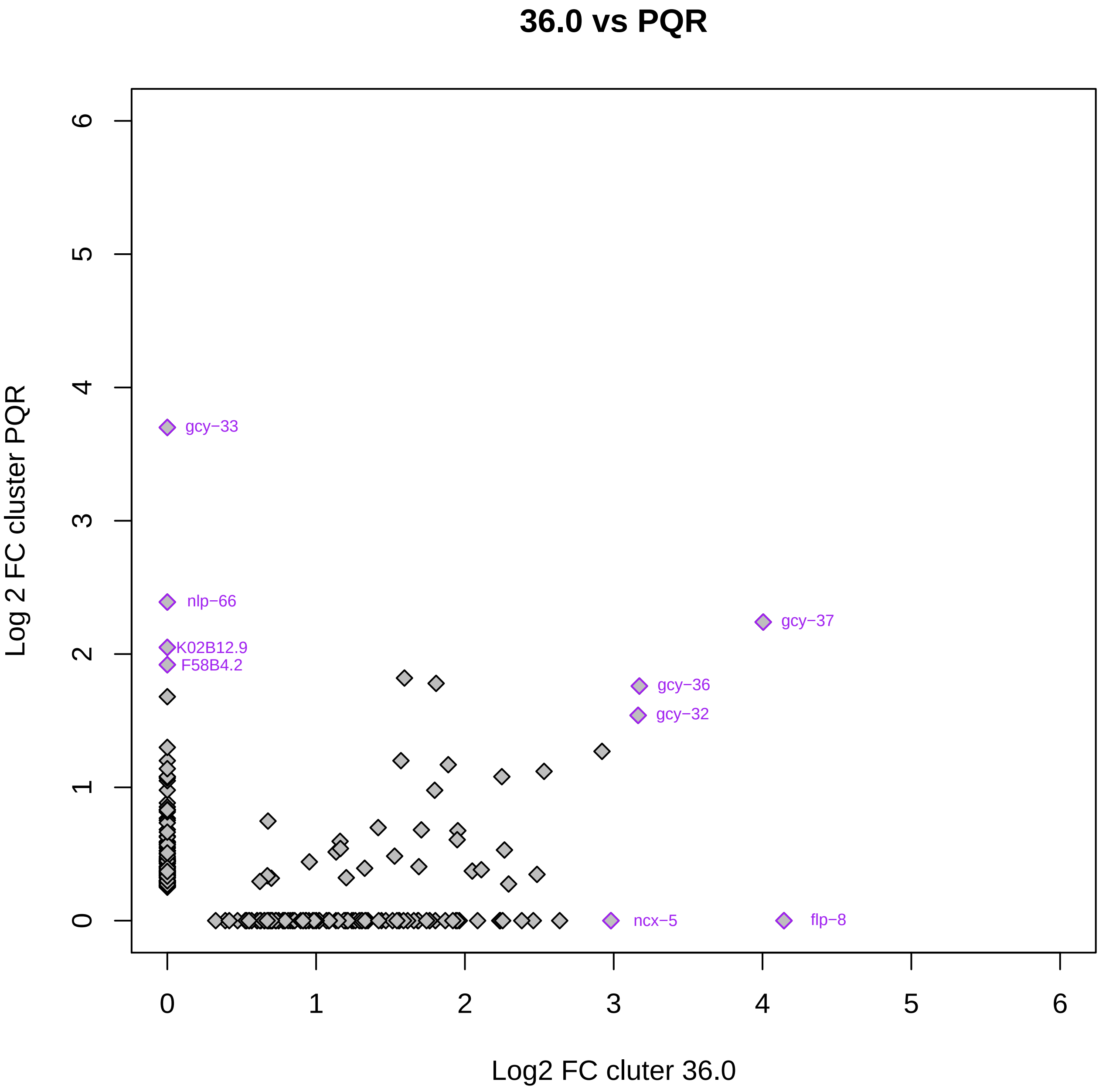
36.0 vs PQR
Log 2 FC cluster PQR
Log2 FC cluter 36.0
gcy−33
nlp−66
K02B12.9
F58B4.2
gcy−37
gcy−36
gcy−32
ncx−5
flp−8
0
1
2
3
4
5
6

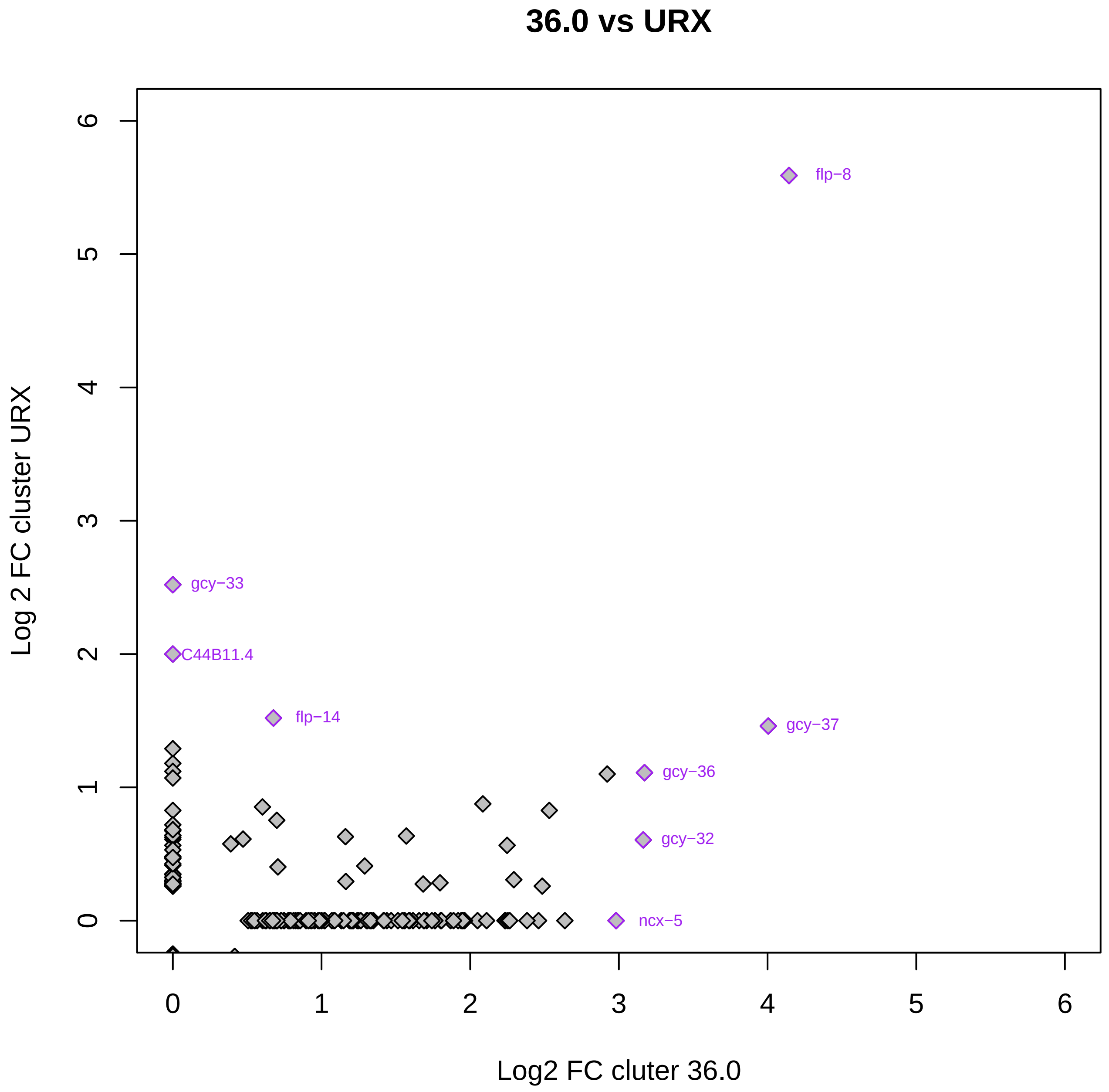
36.0 vs URX
Log 2 FC cluster URX
Log2 FC cluter 36.0
0
1
2
3
4
5
6
flp−8
gcy−33
C44B11.4
flp−14
gcy−37
gcy−36
gcy−32
ncx−5

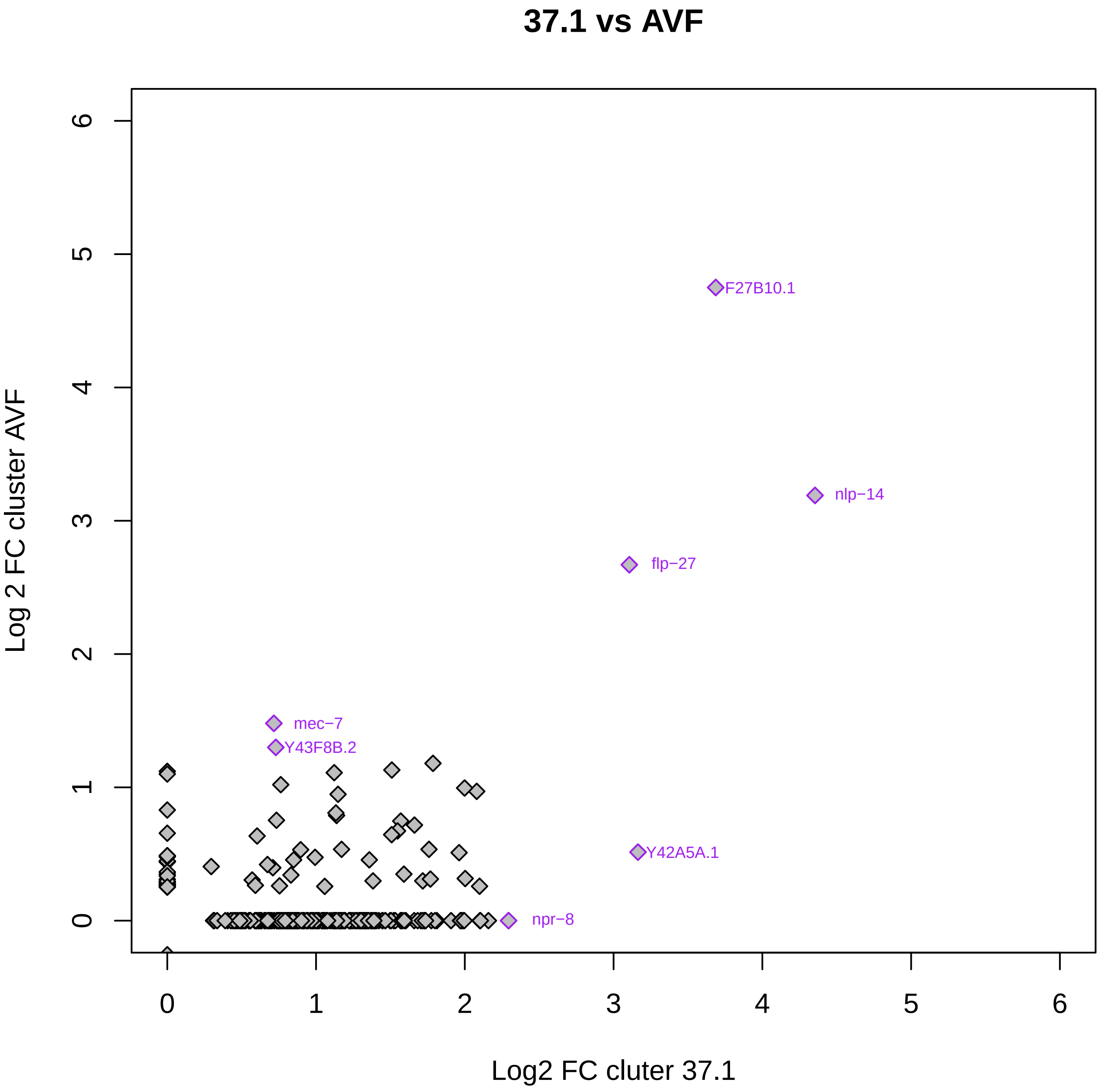

37.1 vs AVF
Log 2 FC cluster AVF
Log2 FC cluter 37.1
0
1
2
3
4
5
6
F27B10.1
nlp−14
flp−27
mec−7
Y43F8B.2
Y42A5A.1
npr−8

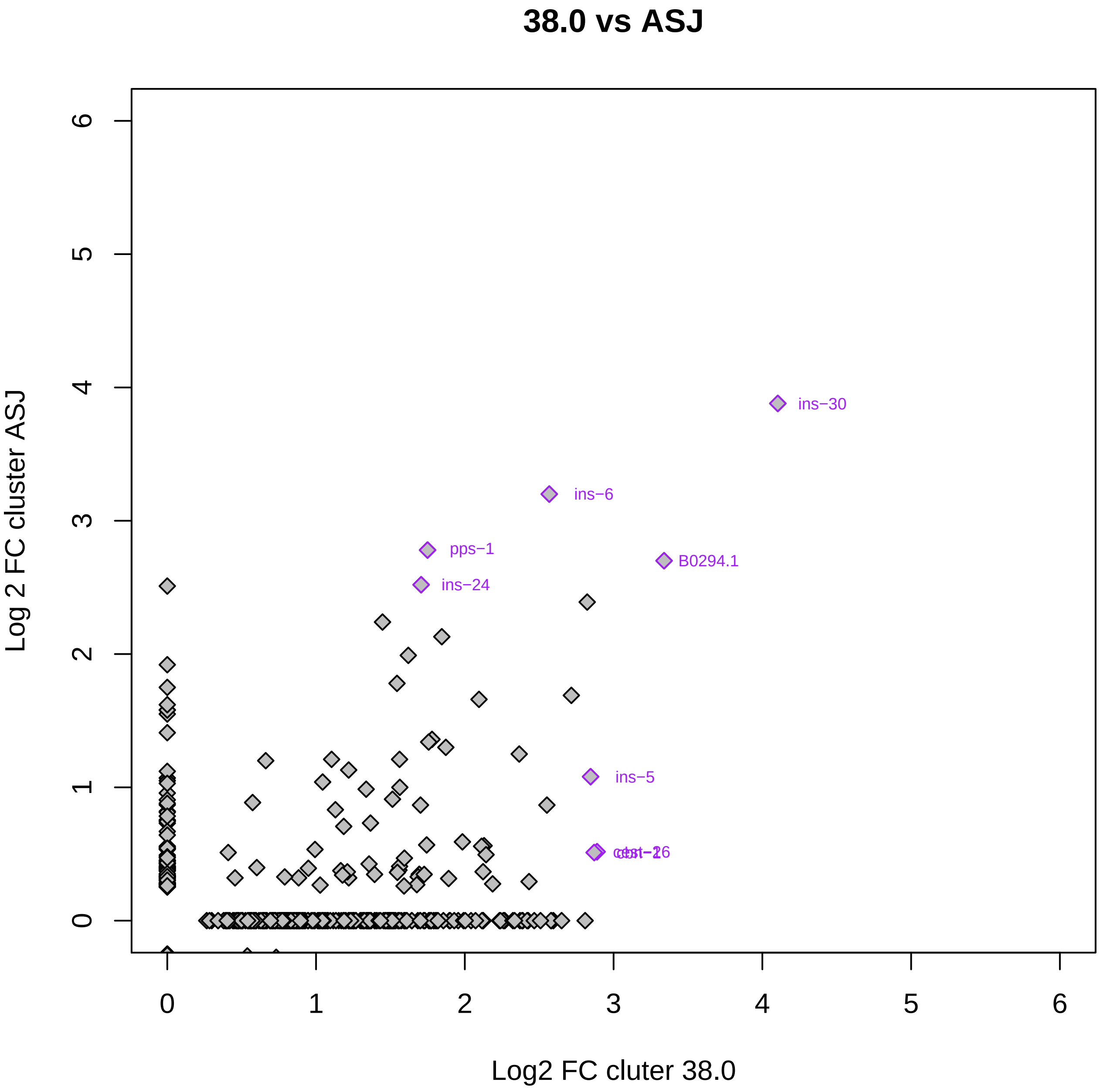

38.0 vs ASJ
ins−30
ins−6
pps−1
B0294.1
ins−24
ins−5
Log 2 FC cluster ASJ
Log2 FC cluter 38.0

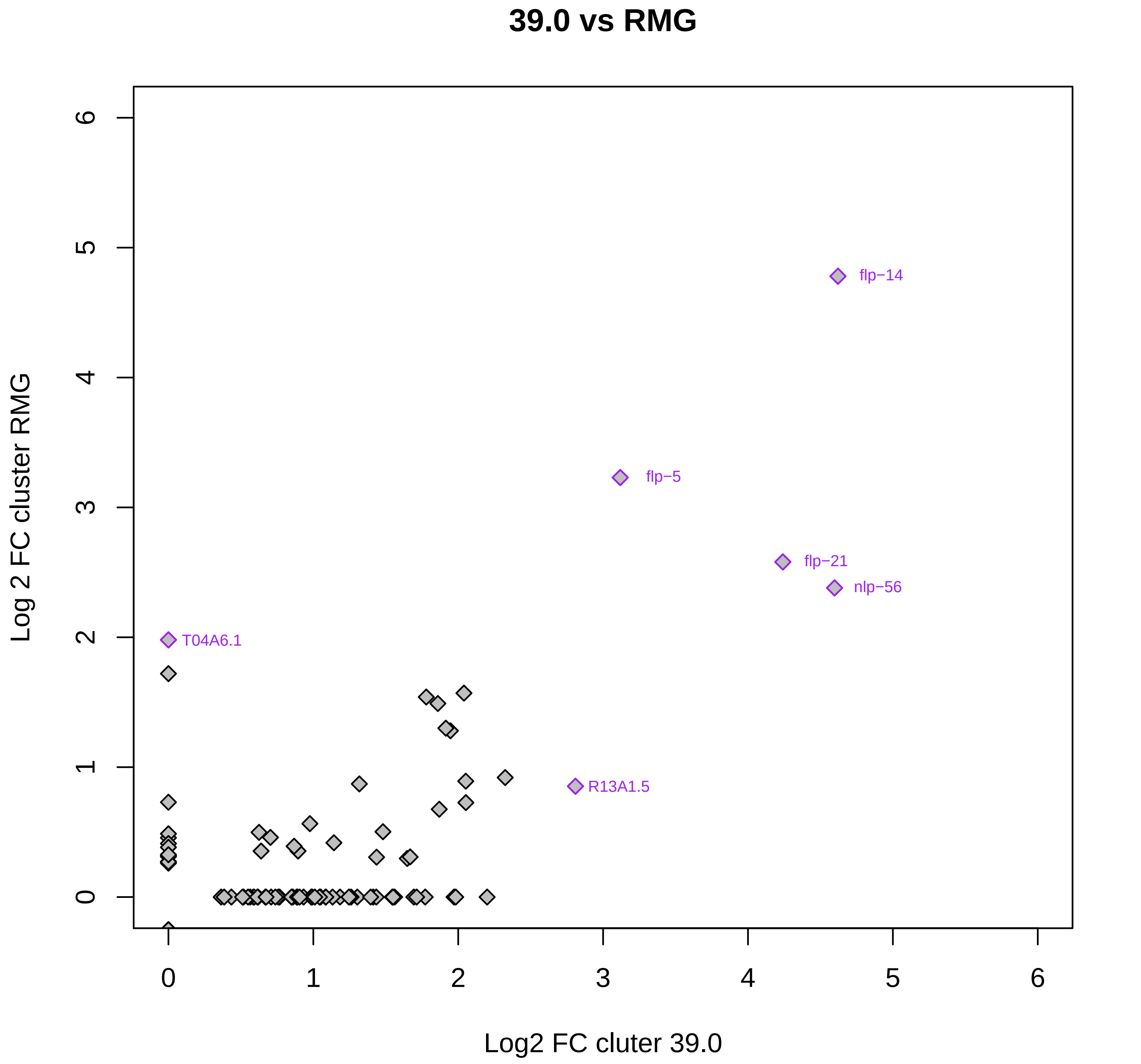
39.0 vs RMG
Log 2 FC cluster RMG
Log2 FC cluter 39.0
flp−14
flp−5
flp−21
nlp−56
T04A6.1
R13A1.5
0
1
2
3
4
5
6

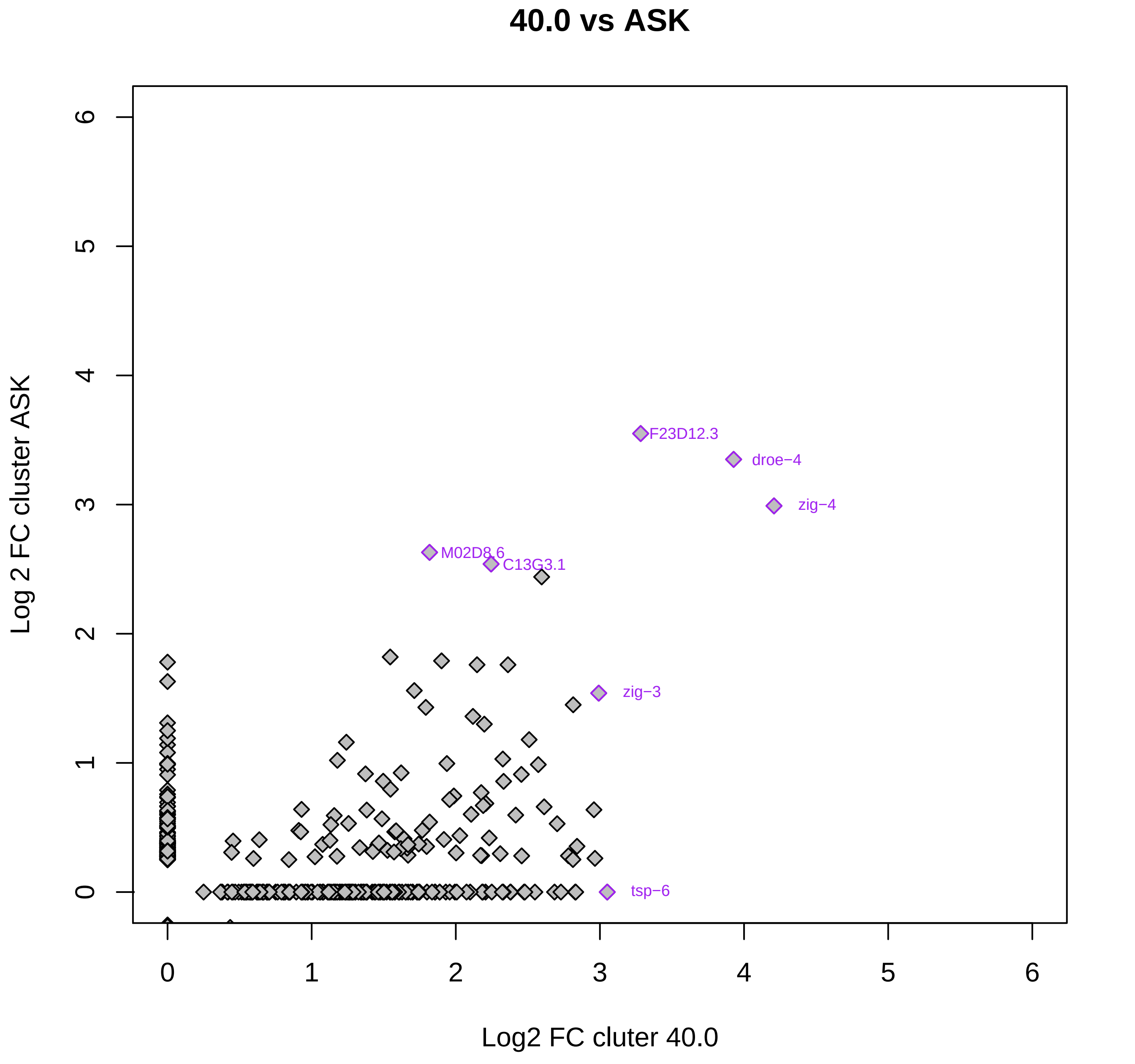

40.0 vs ASK
Log 2 FC cluster ASK
Log2 FC cluter 40.0
F23D12.3
droe−4
zig−4
M02D8.6
C13G3.1
zig−3
tsp−6
0
1
2
3
4
5
6

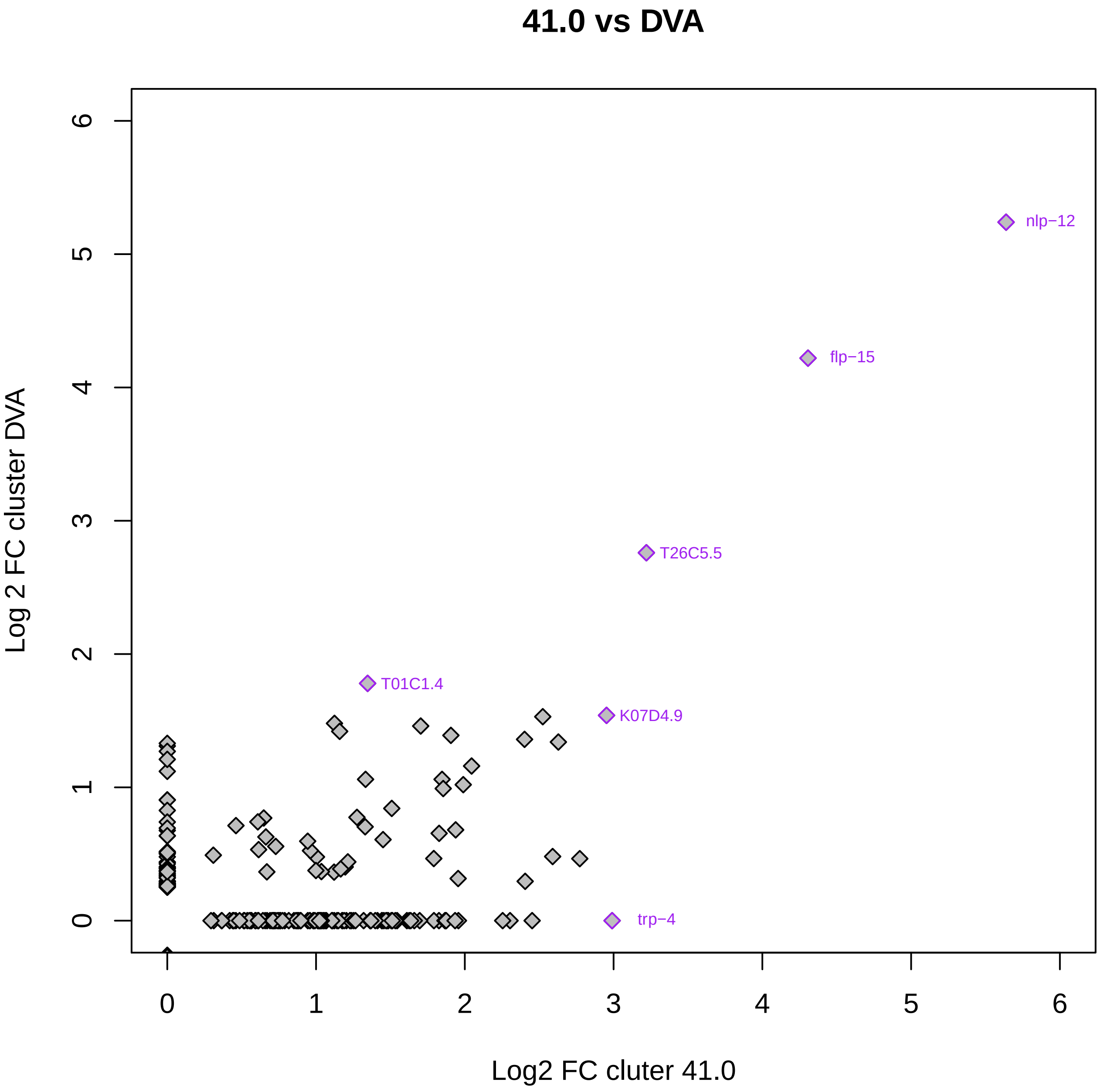
41.0 vs DVA
Log 2 FC cluster DVA
Log2 FC cluter 41.0
nlp−12
flp−15
T26C5.5
T01C1.4
K07D4.9
trp−4
0
1
2
3
4
5
6

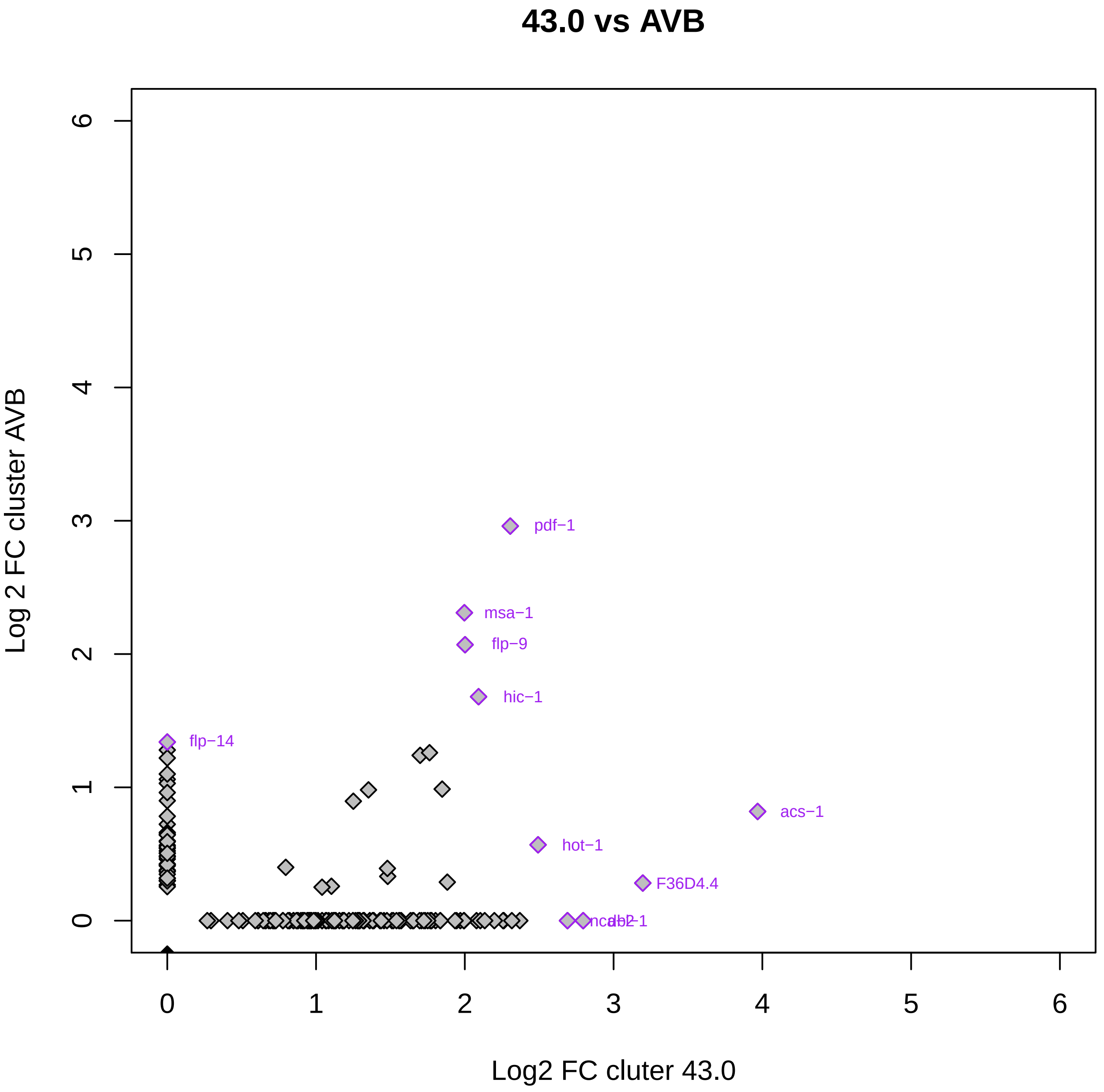

43.0 vs AVB
Log 2 FC cluster AVB
Log2 FC cluter 43.0
0
1
2
3
4
5
6
pdf−1
msa−1
flp−9
hic−1
flp−14
acs−1
hot−1
F36D4.4

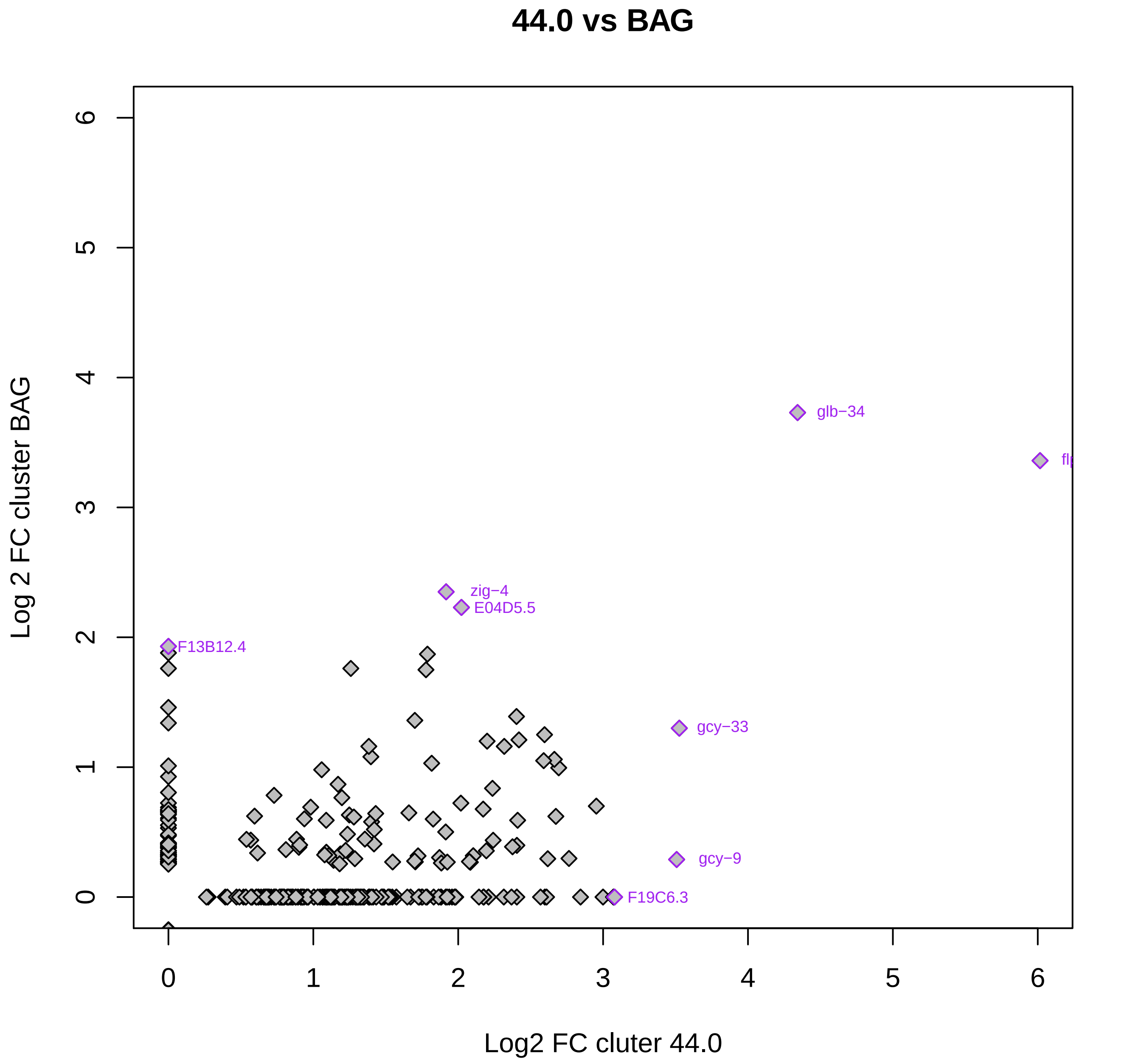

44.0 vs BAG
Log 2 FC cluster BAG
Log2 FC cluter 44.0
0
1
2
3
4
5
6
glb−34
zig−4
E04D5.5
F13B12.4
gcy−33
gcy−9
F19C6.3

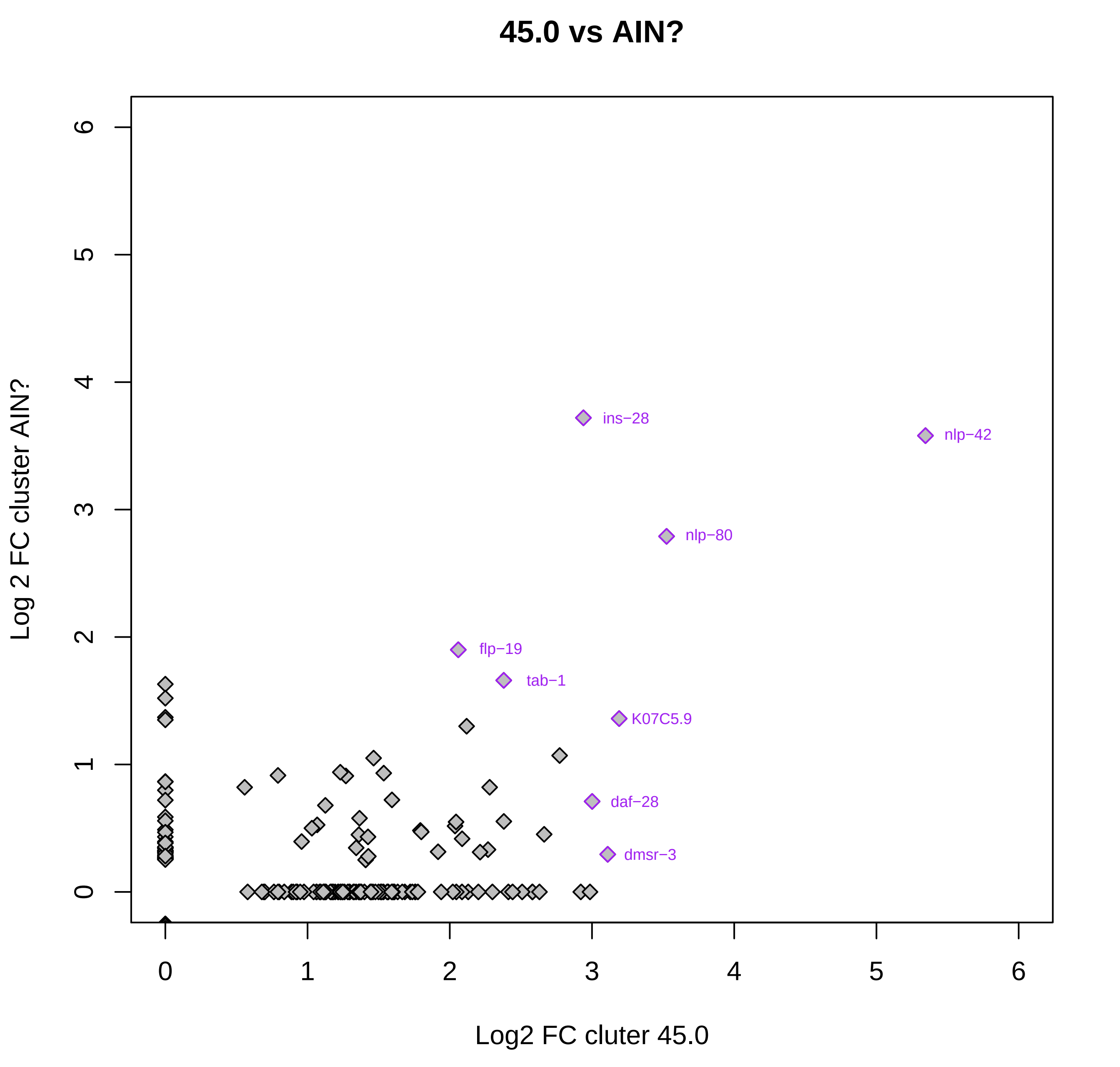

45.0 vs AIN?
Log 2 FC cluster AIN?
Log2 FC cluter 45.0
0
1
2
3
4
5
6
ins−28
nlp−42
nlp−80
flp−19
tab−1
K07C5.9
daf−28
dmsr−3

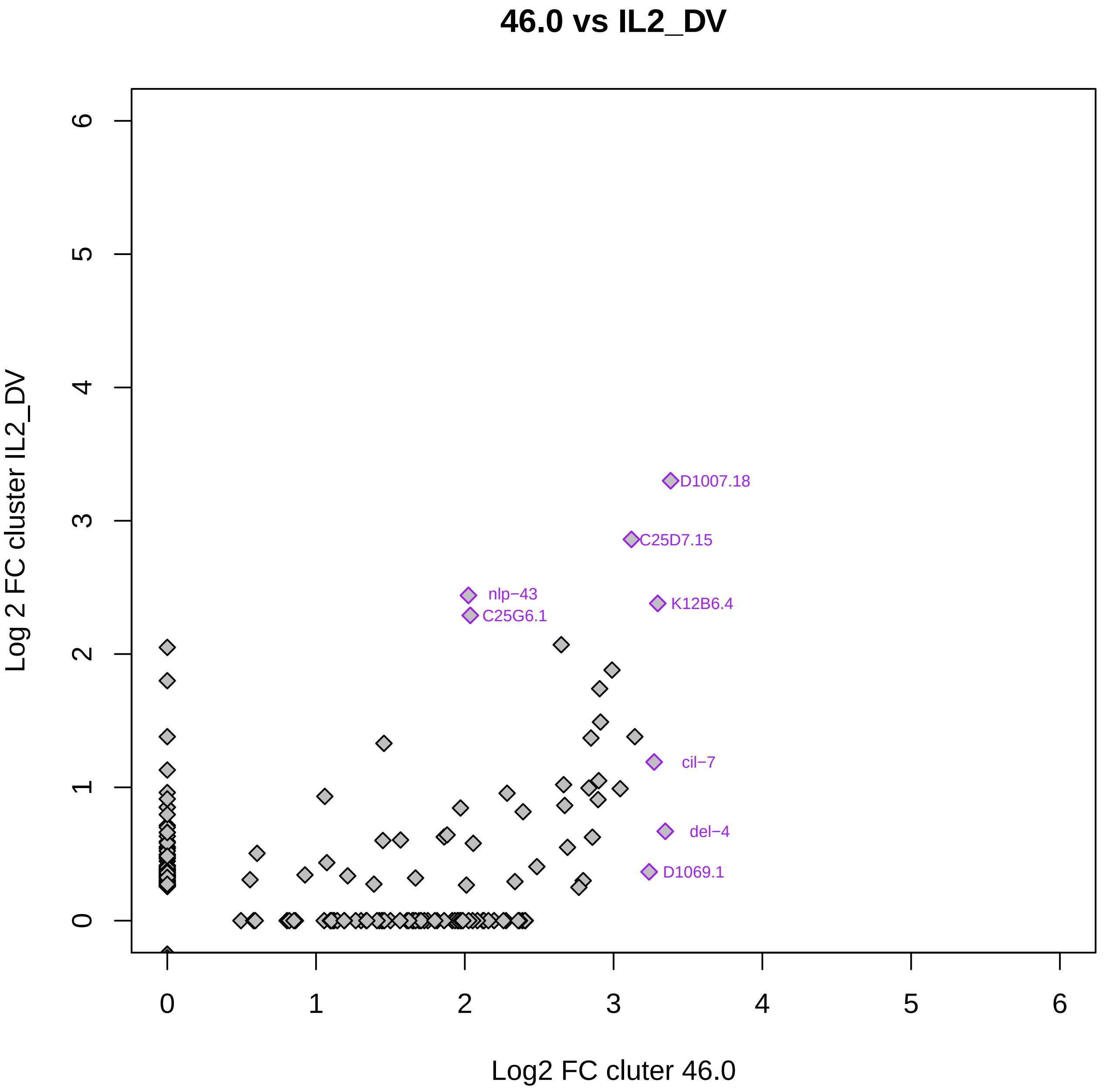
46.0 vs IL2_DV
Log 2 FC cluster IL2_DV
Log2 FC cluter 46.0
0
1
2
3
4
5
6
D1007.18
C25D7.15
nlp−43
C25G6.1
K12B6.4
cil−7
del−4
D1069.1

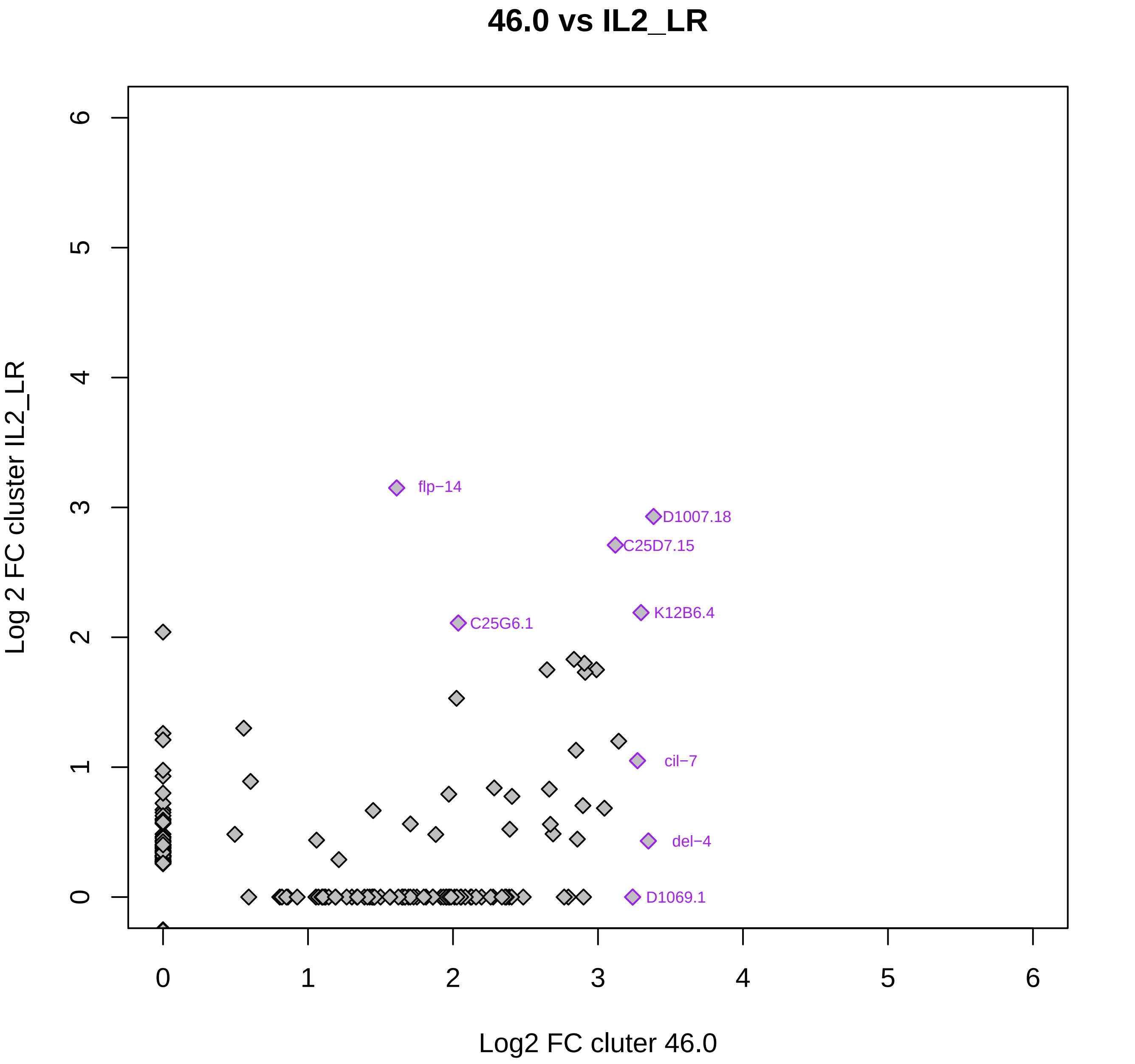

46.0 vs IL2_LR
Log 2 FC cluster IL2_LR
Log2 FC cluter 46.0
0
1
2
3
4
5
6
0
1
2
3
4
5
6
flp−14
D1007.18
C25D7.15
K12B6.4
C25G6.1
cil−7
del−4
D1069.1

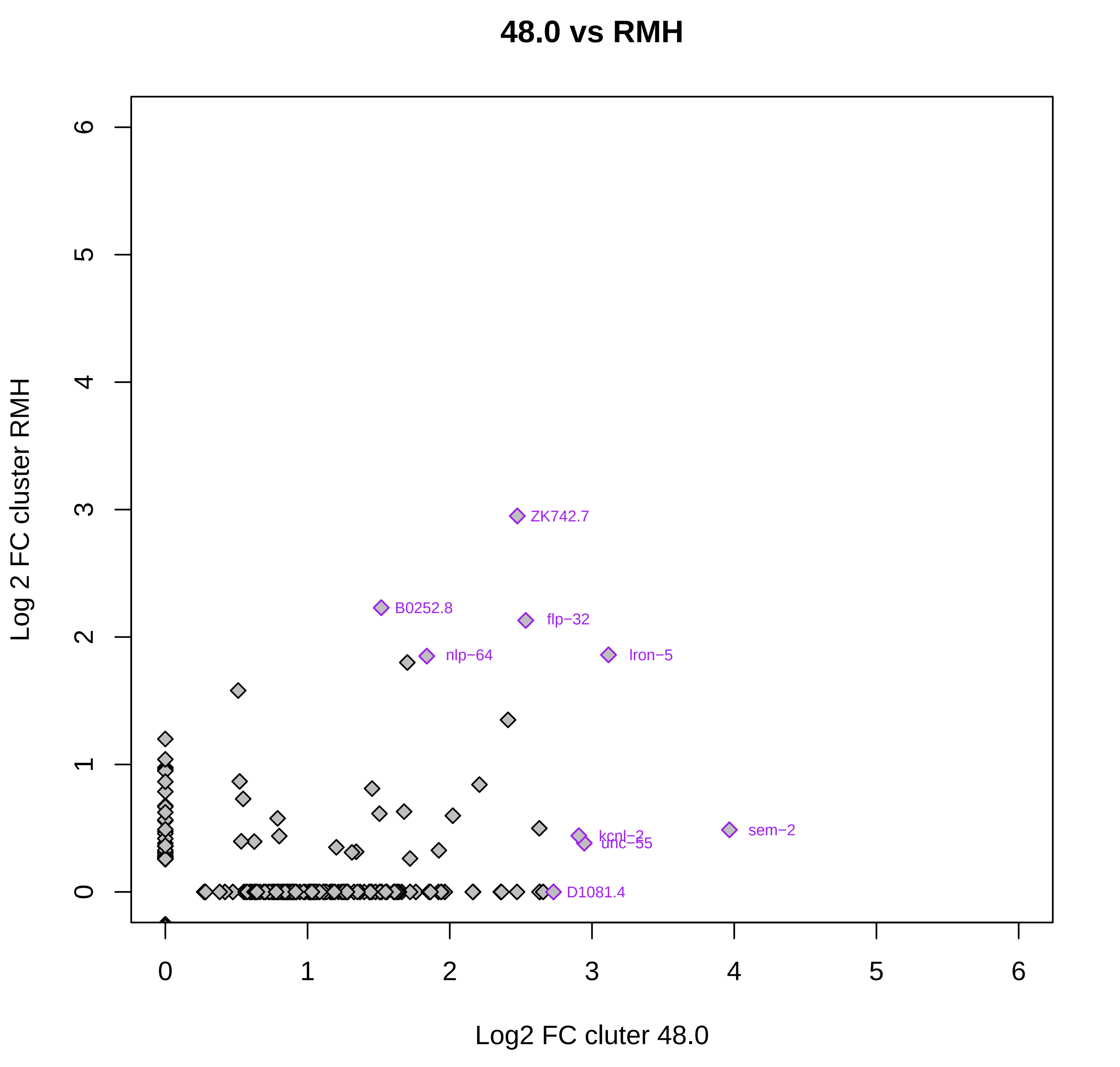

48.0 vs RMH
Log 2 FC cluster RMH
Log2 FC cluter 48.0
0
1
2
3
4
5
6
ZK742.7
B0252.8
flp−32
nlp−64
lron−5
sem−2
kcnl−2
unc−55
D1081.4

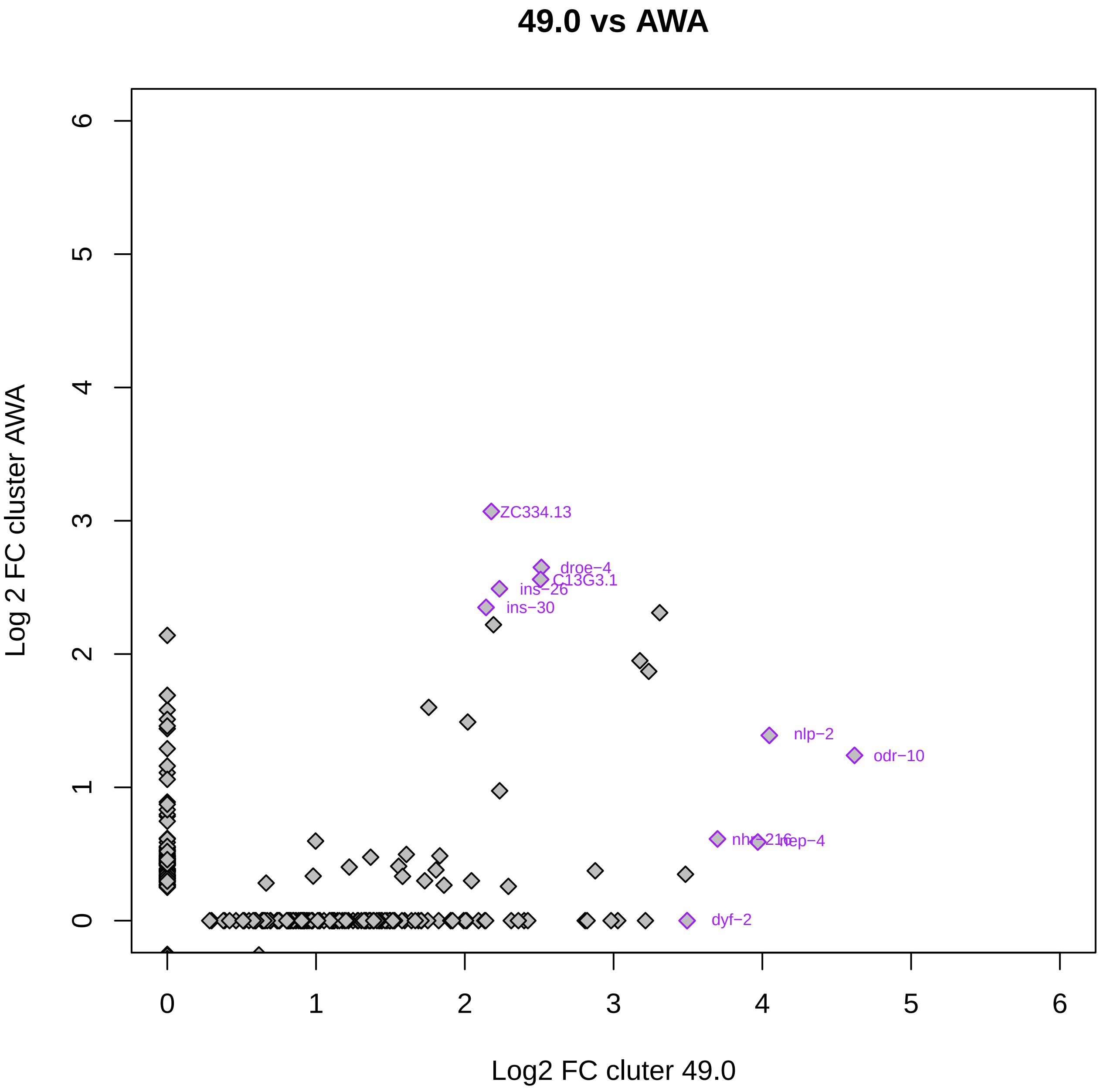

49.0 vs AWA
Log 2 FC cluster AWA
Log2 FC cluter 49.0
0
1
2
3
4
5
6
ZC334.13
droe−4
C13G3.1
ins−26
ins−30
nlp−2
odr−10
nhr−216
nep−4
dyf−2

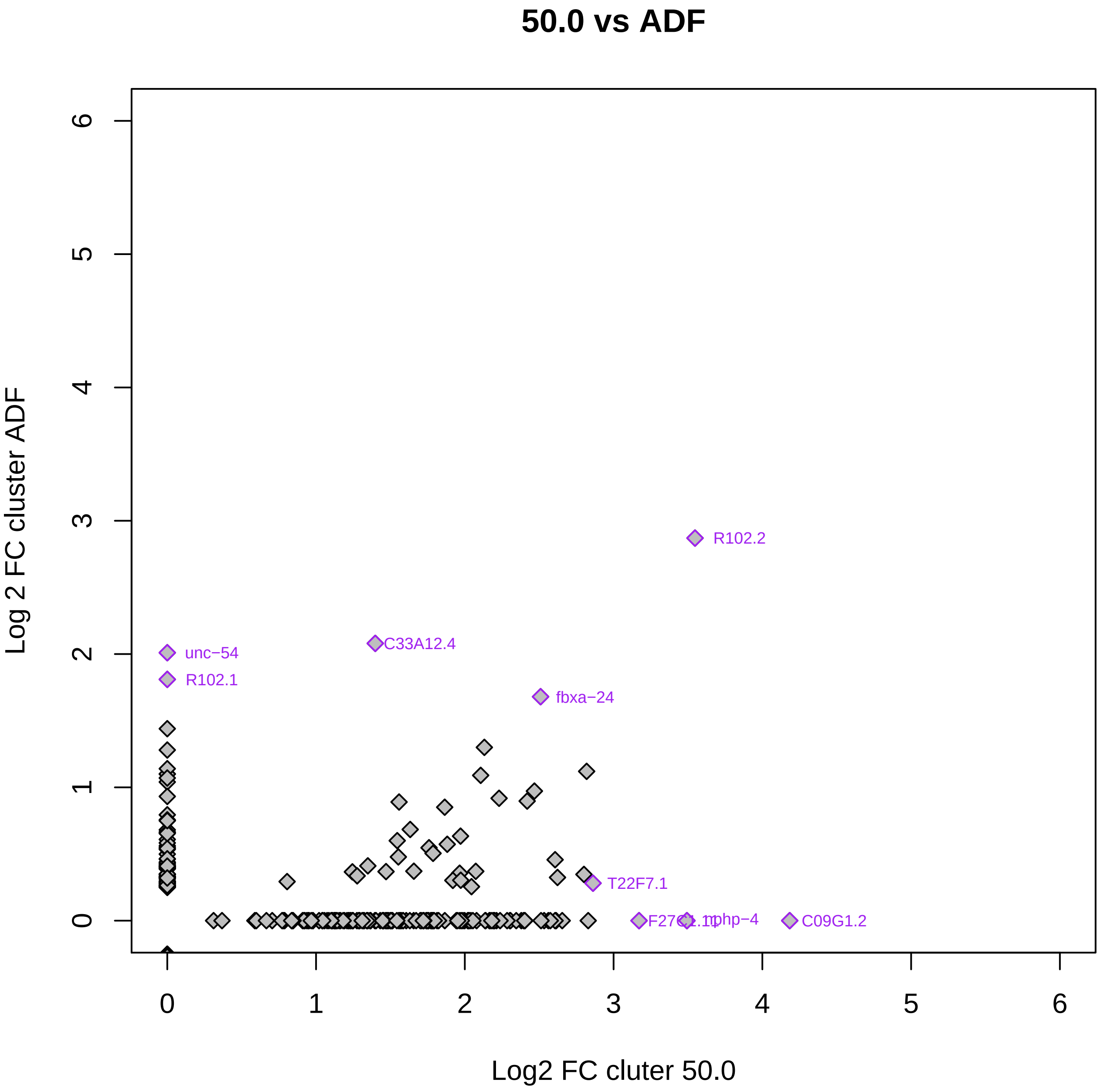

50.0 vs ADF
Log 2 FC cluster ADF
Log2 FC cluter 50.0
0
1
2
3
4
5
6
R102.2
C33A12.4
unc−54
R102.1
fbxa−24
T22F7.1
nphp−4
C09G1.2

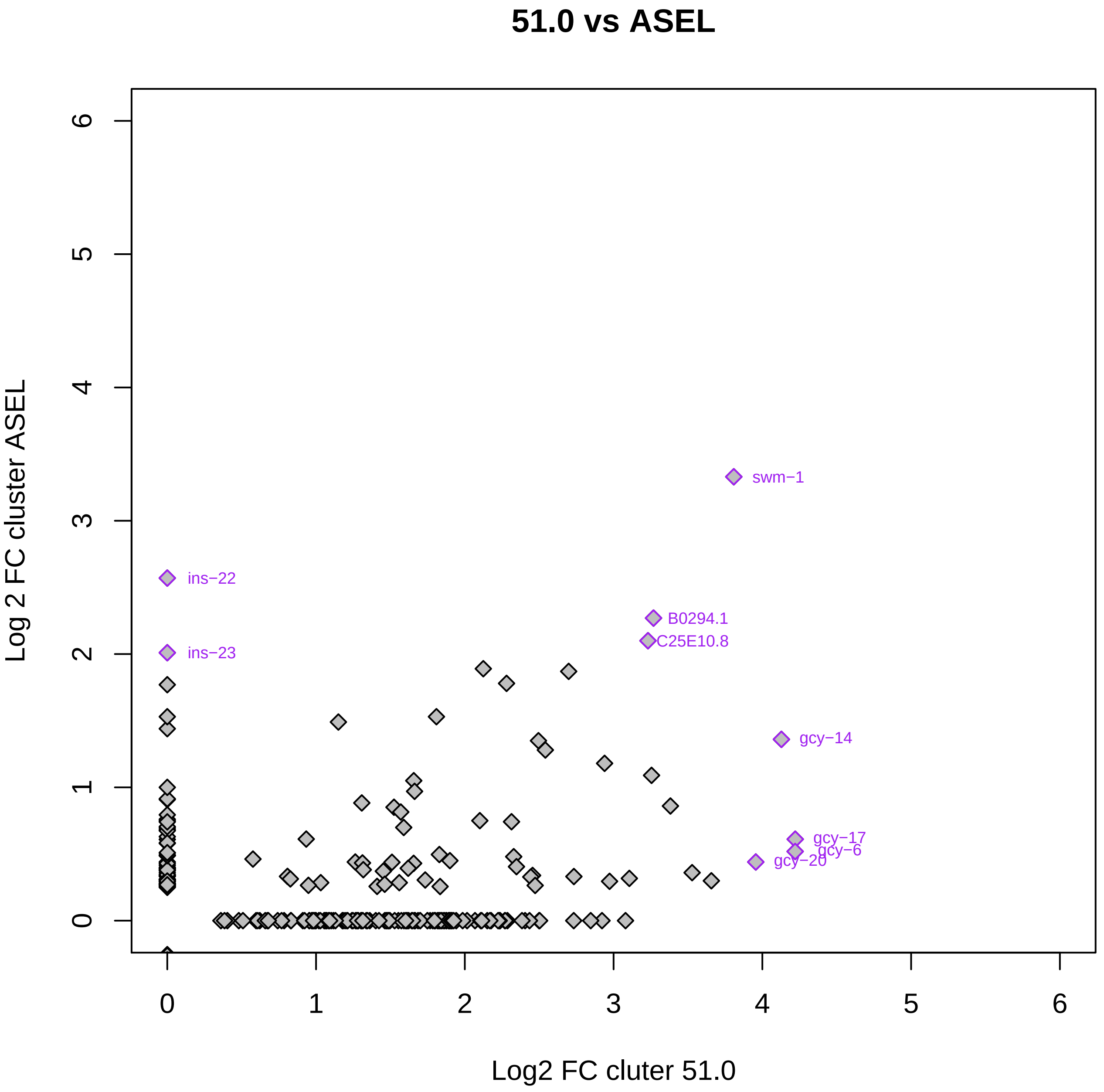

51.0 vs ASEL
Log 2 FC cluster ASEL
Log2 FC cluter 51.0
0
1
2
3
4
5
6
swm−1
ins−22
ins−23
B0294.1
C25E10.8
gcy−14
gcy−17
gcy−6
gcy−20

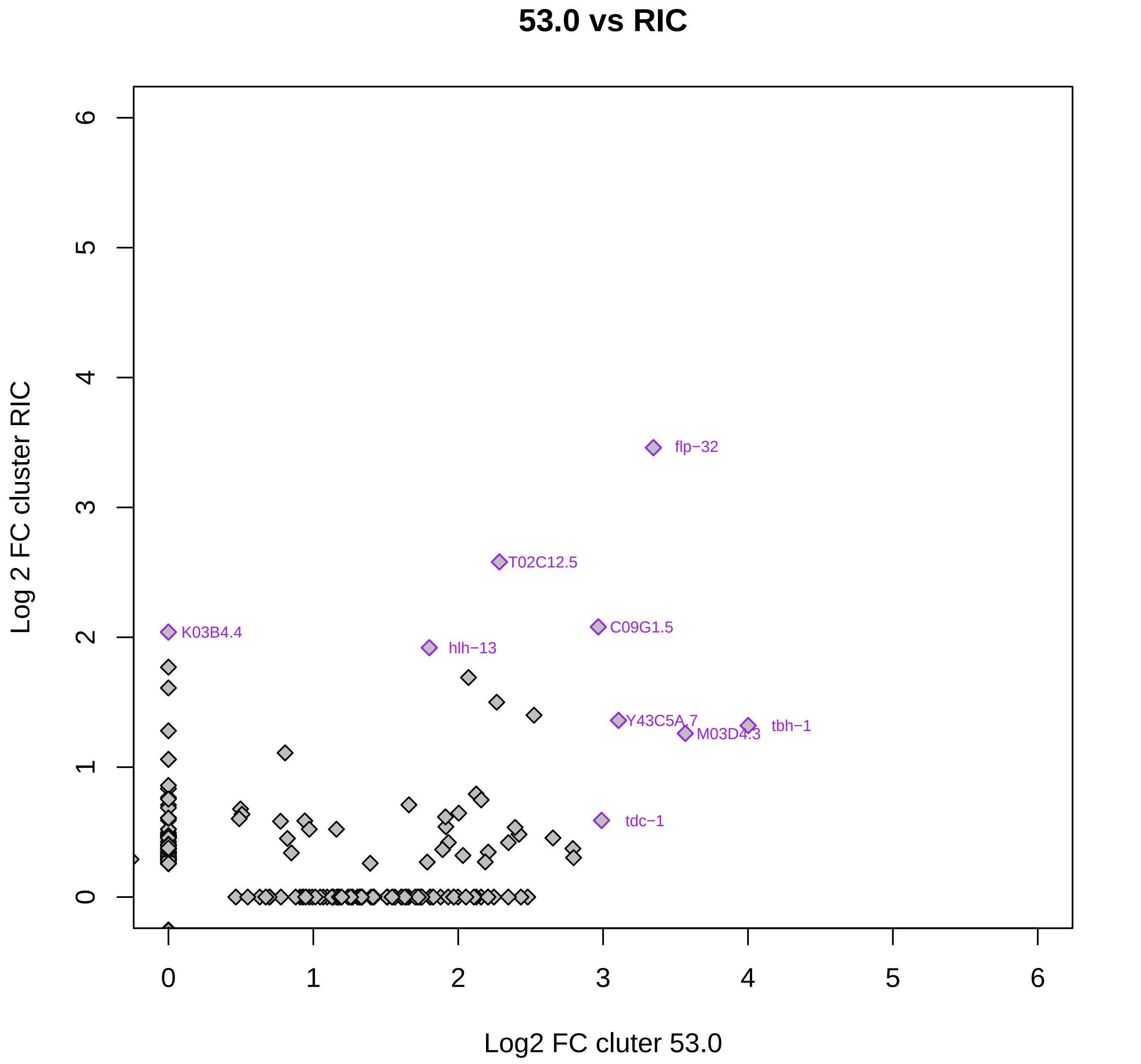

53.0 vs RIC
Log 2 FC cluster RIC
Log2 FC cluter 53.0
0
1
2
3
4
5
6
flp−32
T02C12.5
C09G1.5
K03B4.4
hlh−13
Y43C5A.7
M03D4.3
tbh−1
tdc−1

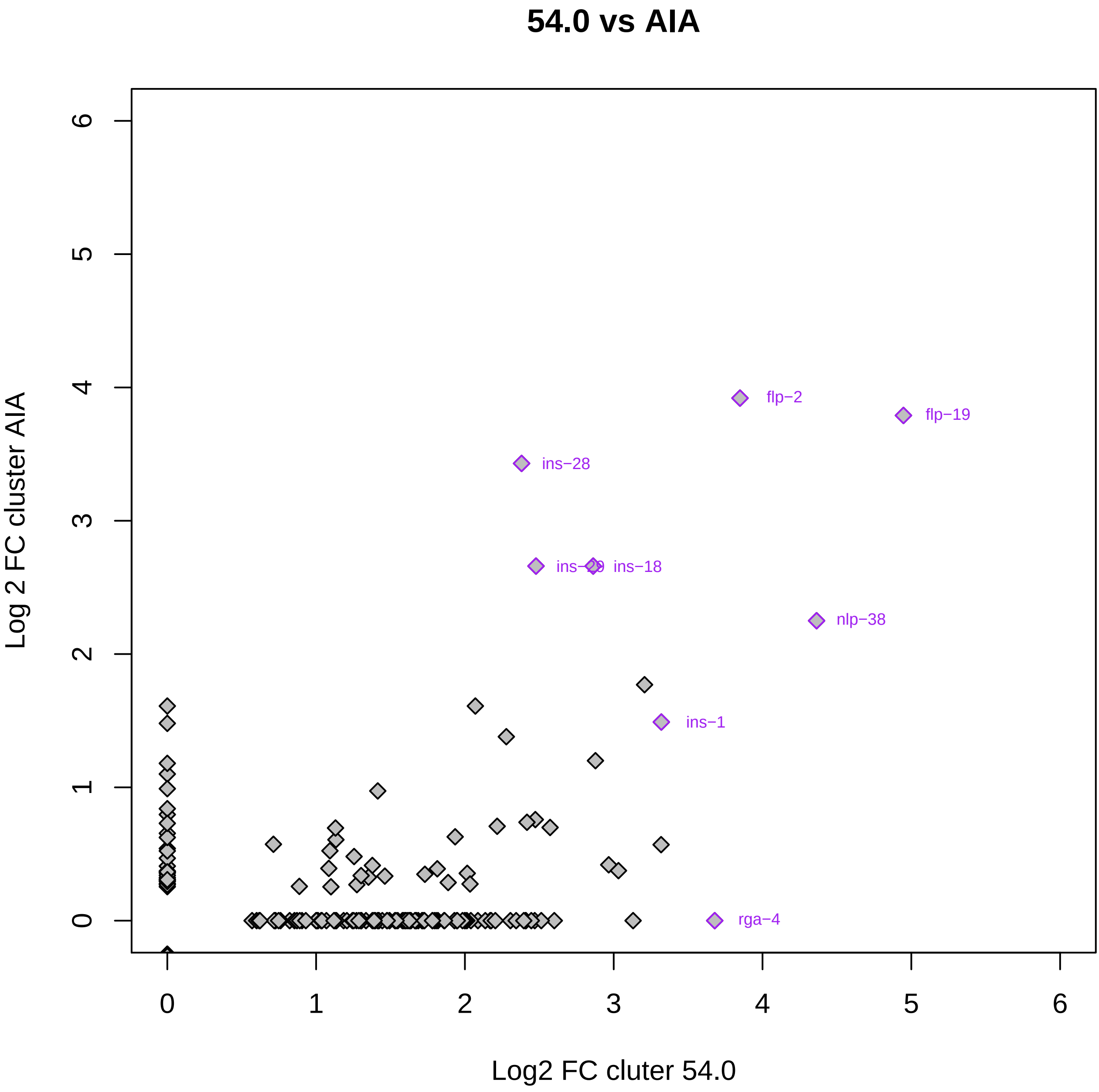

54.0 vs AIA
Log 2 FC cluster AIA
Log2 FC cluter 54.0
0
1
2
3
4
5
6
flp−2
flp−19
ins−28
ins−29
ins−18
nlp−38
ins−1
rga−4

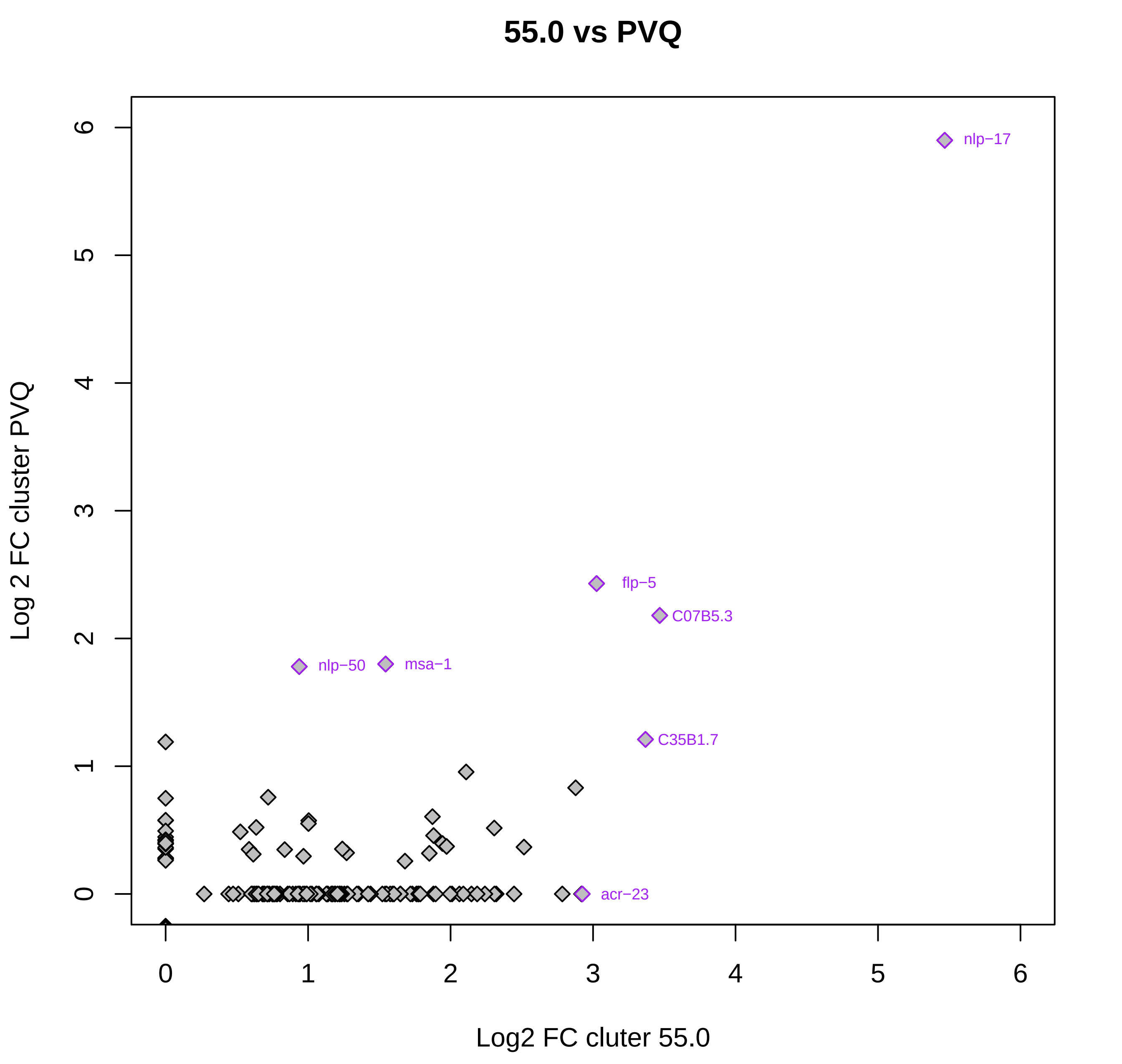
55.0 vs PVQ
Log 2 FC cluster PVQ
Log2 FC cluter 55.0
0
1
2
3
4
5
6
0
1
2
3
4
5
6
nlp−17
flp−5
C07B5.3
nlp−50
msa−1
C35B1.7
acr−23

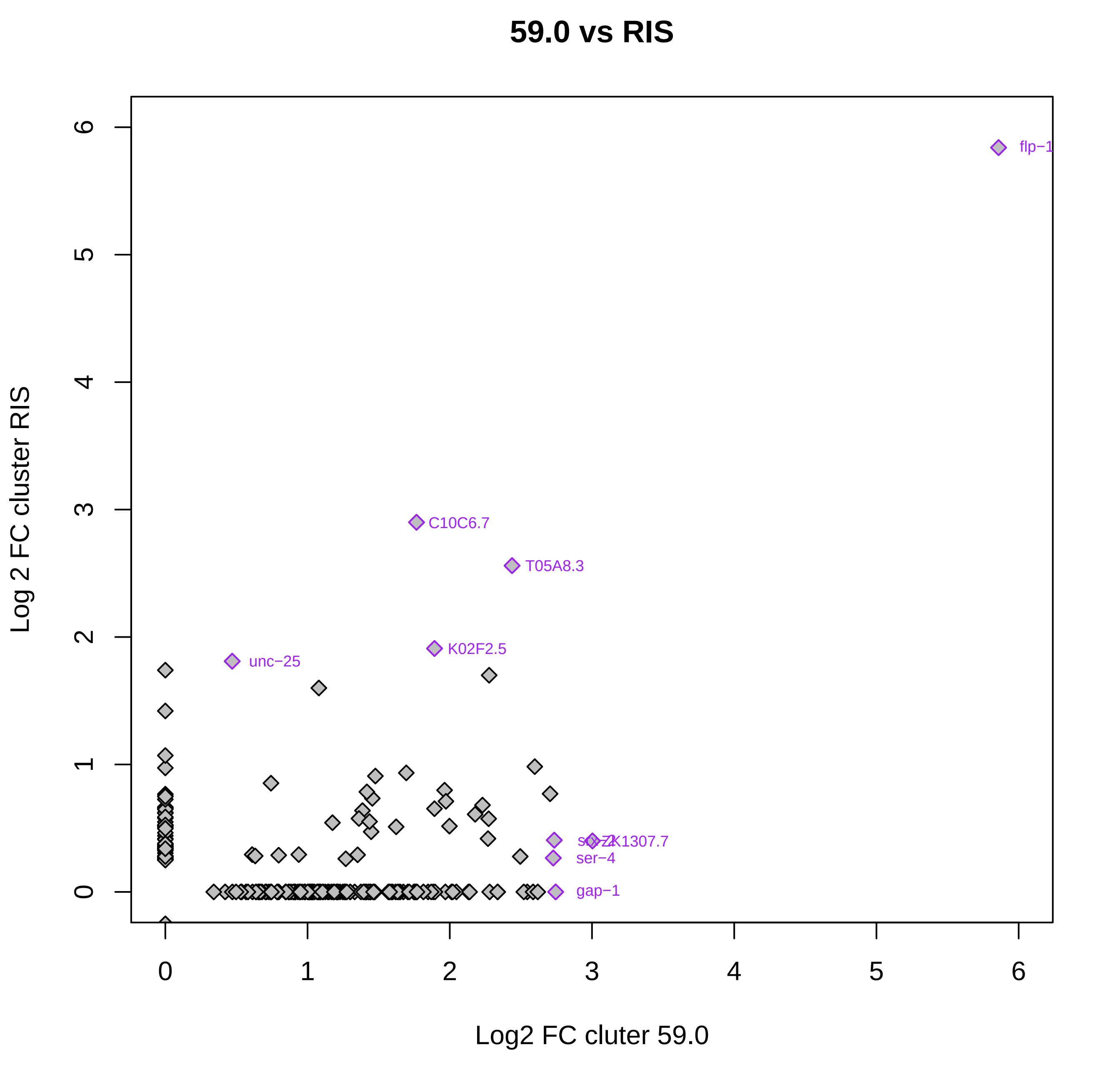

59.0 vs RIS
Log 2 FC cluster RIS
Log2 FC cluter 59.0
0
1
2
3
4
5
6
flp−1
C10C6.7
T05A8.3
K02F2.5
unc−25
ZK1307.7
ser−4
gap−1

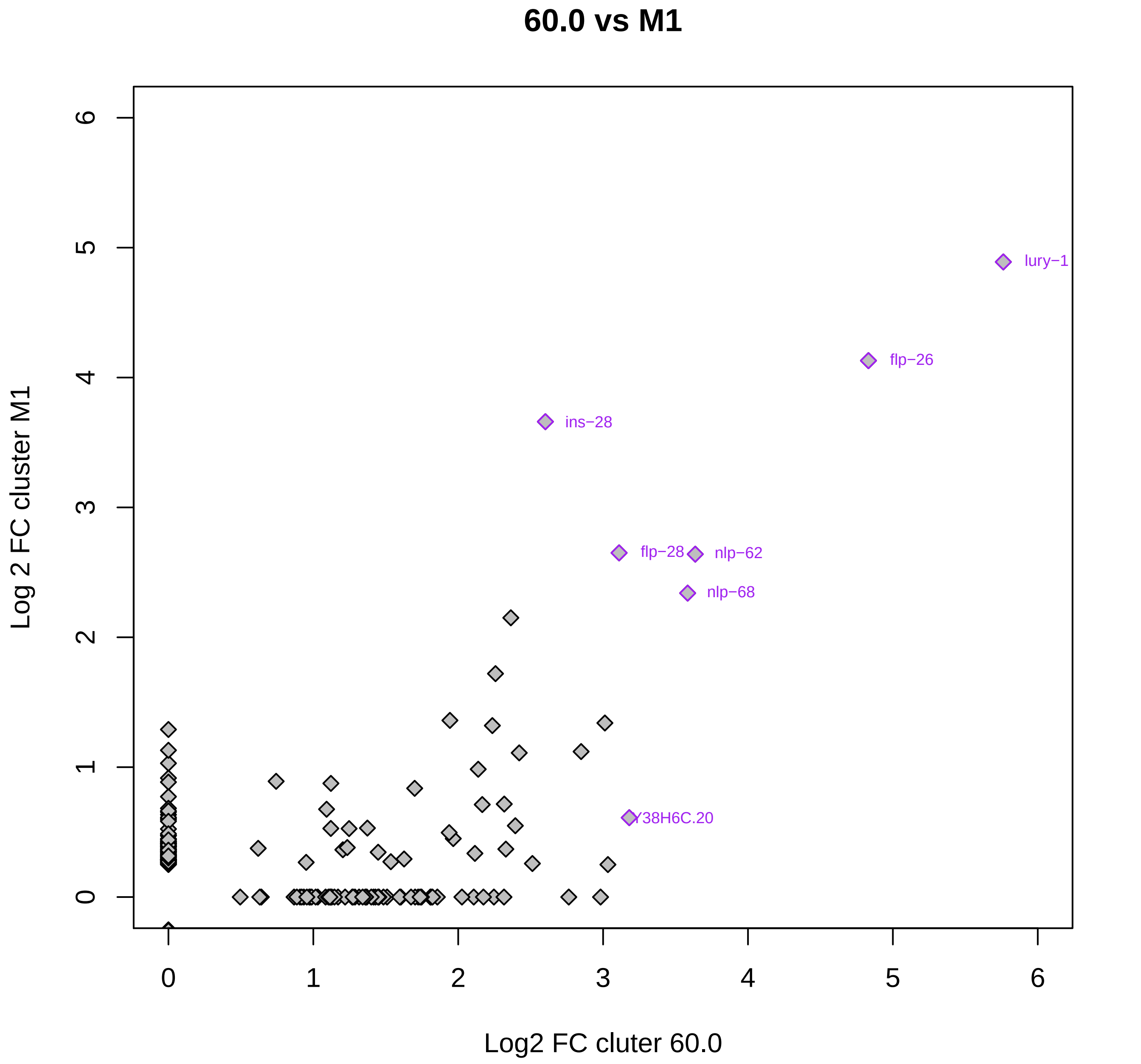

60.0 vs M1
lury−1
flp−26
ins−28
flp−28
nlp−62
nlp−68
Y38H6C.20
0
1
2
3
4
5
6
Log 2 FC cluster M1
Log2 FC cluter 60.0

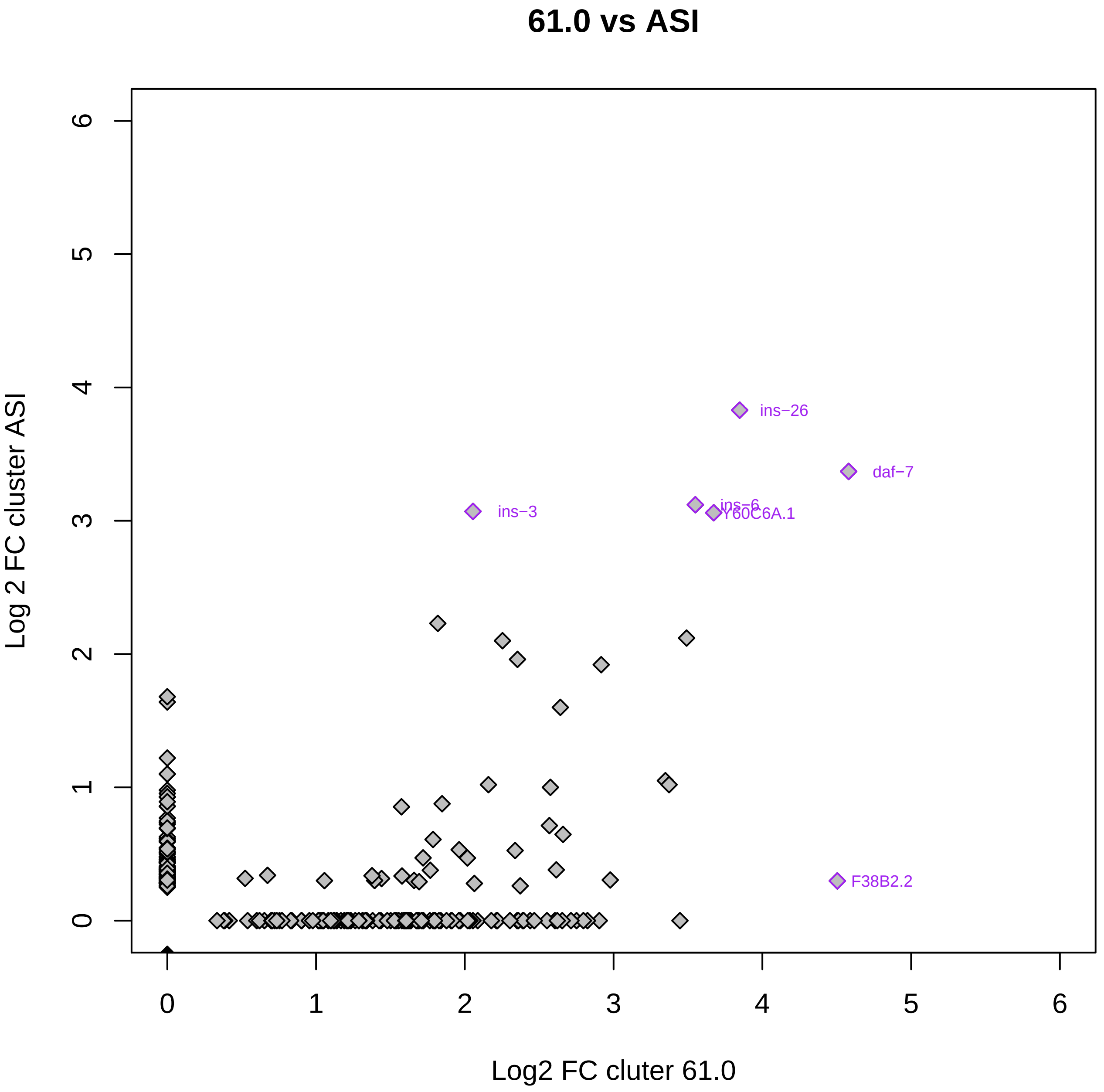
61.0 vs ASI
Log 2 FC cluster ASI
Log2 FC cluter 61.0
0
1
2
3
4
5
6
ins−26
daf−7
ins−3
ins−6
Y60C6A.1
F38B2.2

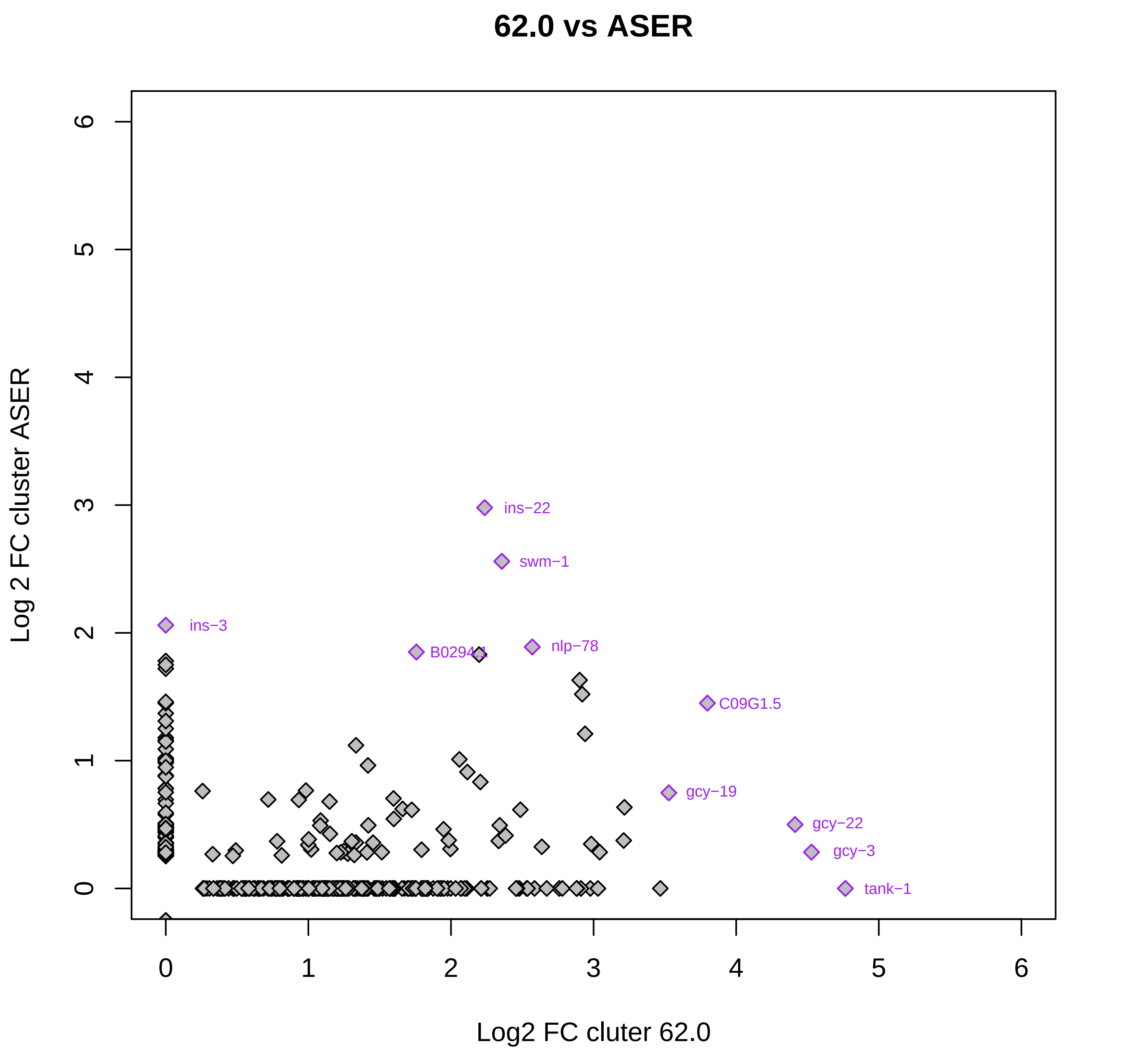
62.0 vs ASER
Log 2 FC cluster ASER
Log2 FC cluter 62.0
0
1
2
3
4
5
6
ins−22
swm−1
ins−3
B0294.1
nlp−78
C09G1.5
gcy−19
gcy−22
gcy−3
tank−1

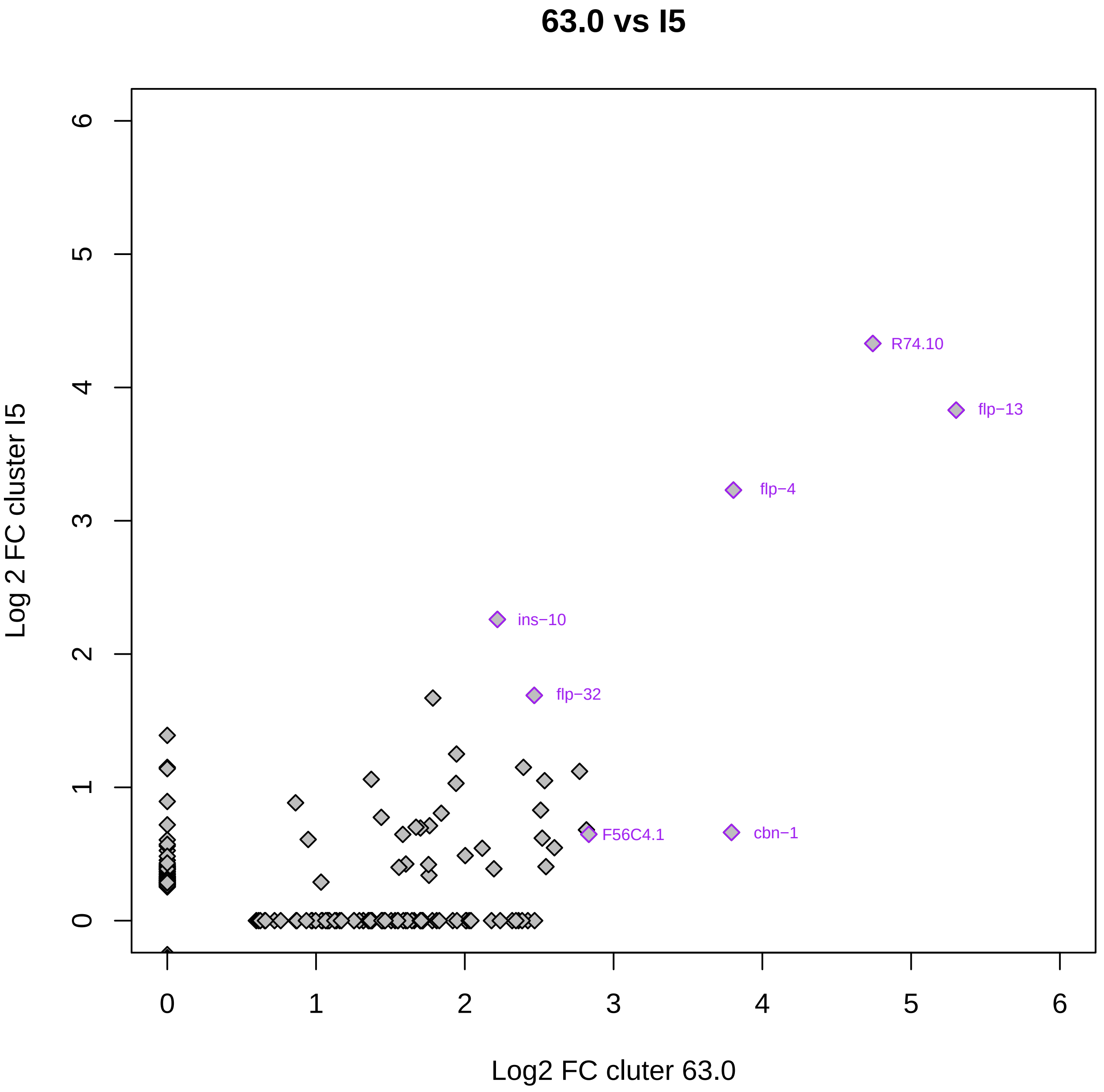

63.0 vs I5
Log 2 FC cluster I5
Log2 FC cluter 63.0
0
1
2
3
4
5
6
R74.10
flp−13
flp−4
ins−10
flp−32
F56C4.1
cbn−1
